# Old Age Drives Fusion and Expansion of Vitellogenin Vesicles in the Intestine of *Caenorhabditis elegans*

**DOI:** 10.1101/2022.07.21.501051

**Authors:** Chao Zhai, Nan Zhang, Xi-Xia Li, Xi Chen, Fei Sun, Meng-Qiu Dong

## Abstract

Some of the most conspicuous aging phenotypes of *C. elegans* are related to post-reproductive production of vitellogenins (Vtg), which form yolk protein (YP) complexes after processing and lipid loading. Vtg/YP levels show huge increases with age, and inhibition of this extends lifespan, but how subcellular and organism-wide distribution of these proteins changes with age has not been systematically explored. Here this has been done to understand how vitellogenesis promotes aging. The age-associated changes of intestinal vitellogenin vesicles (VVs), pseudocoelomic yolk patches (PYPs), and gonadal yolk organelles (YOs) have been characterized by immuno-electron microscopy. We find that from reproductive adult day 2 (AD 2) to post-reproductive AD 6 and AD 9, intestinal VVs expand from 0.2 μm to 3-4 μm in diameter or by > 3,000 times in volume, PYPs increase by > 3 times in YP concentration and volume, while YOs in oocytes shrink slightly from 0.5 μm to 0.4 μm in diameter or by 60% in volume. In AD 6 and AD 9 worms, mis-localized YOs found in the hypodermis, uterine cells, and the somatic gonadal sheath can reach a size of 10 μm across in the former two tissues. This remarkable size increase of VVs and that of mis-localized YOs in post-reproductive worms are accompanied by extensive fusion between these Vtg/YP-containing vesicular structures in somatic cells. In contrast, no fusion is seen between YOs in oocytes. We propose that in addition to the continued production of Vtg, excessive fusion between VVs and mislocalized YOs in the soma worsen the aging pathologies seen in *C. elegans.*

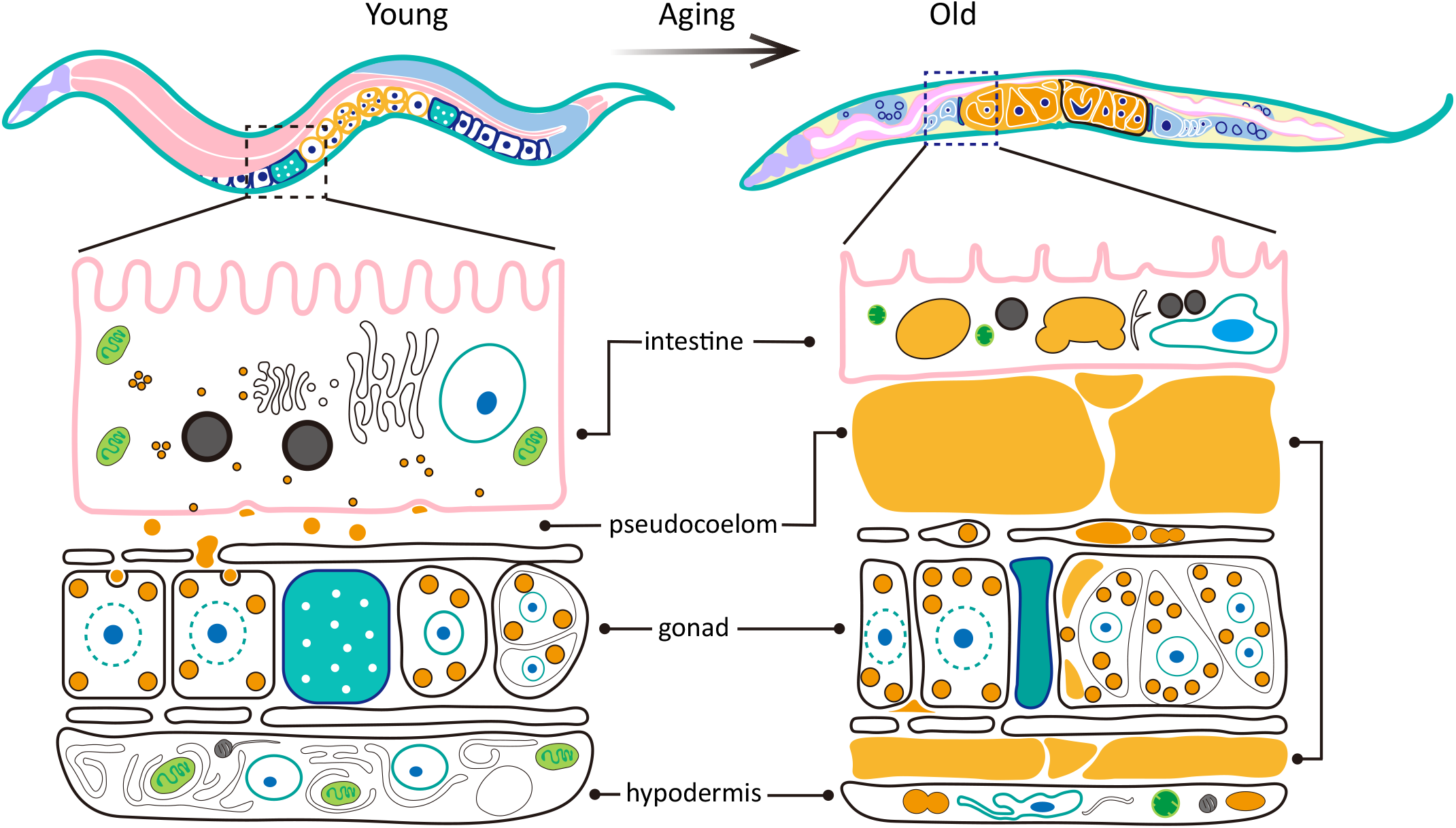
Graphical Summary: Age-dependent changes of vitellogenins-/yolk proteins-containing subcellular structures in *C. elegans*. On the left is a young wild-type *C. elegans* of adult day 1 or 2, and on the right is an older worm of adult day 6 or 9. The orange-colored structures are: in the intestine, vitellogenin vesicles (VVs), of a median diameter of 200 nm on the left and 3-4 micrometers on the right, with a frequent fusion between VVs in the older worm; in the pseudocoelom, pseudocoelomic yolk patches, greatly expanded in the older worm; in oocytes and eggs, yolk organelles (YOs), of a median diameter of 500 nm on the left and 400 nm on the right; in the hypodermis, sheath or uterine cells of the older worm, mislocalized YOs.

## INTRODUCTION

All metazoan animals synthesize in large quantities a tiny number of highly conserved, specialized proteins as provisions of nutrients to their progeny. These include vitellogenins (Vtg), the precursors of yolk proteins (YP) that are deposited in the eggs of nearly all oviparous animals (Sun & Zhang, 2015). In viviparous mammals, which have no Vtg genes (Zhou et al., 2021), this functional role is taken up in a sense by casein, the major protein component of milk. Interestingly, unlike casein proteins, Vtgs are not merely nutrients allocated to the young. For example, Vtg affects the division of labor in honey bees, that is, hive bees vs foragers (Amdam et al., 2003). In addition, Vtgs can scavenge free radicals, carry metal ions, and exert immunological activities in insects, fish, and corals. (Du et al., 2017; Leipart et al.; Sun & Zhang, 2015). There are reports of Vtg/YP participating in intergenerational signal transduction. In *C. elegans*, maternal age and early-life starvation experience of the mother affect maternal provision of YPs to the progeny, which in turn affects growth, fecundity, and several other physiological traits of the progeny (Perez et al., 2017; Jordan et al., 2019). Another way by which YPs could influence intergenerational inheritance in *C. elegans* is to act as carriers of double-stranded RNAs and deposit these messenger molecules from the mother to the progeny (Marré et al., 2016).

There is also a connection between Vtg and aging in perennial social insects. In ants and honey bees, the production of Vtg is negatively correlated with that of juvenile hormone (Amdam et al., 2004; Amsalem et al., 2014), which prevents precocious metamorphosis during development and promotes aging of adults (Jindra et al., 2013). Reducing the honey bee Vtg protein levels by RNA interference (RNAi) elevates juvenile hormone and shortens lifespan (Nelson et al., 2007).

Studies of the nematode *C. elegans* have found repeatedly that expression of the Vtg genes affects adult lifespan in a negative way. There are six Vtg genes in *C. elegans*, from *vit-1* to *vit-6.* RNAi of one or more Vtg genes has been shown to extend the lifespan of wild-type (WT) *C. elegans* by ~20% or less (Ezcurra et al., 2018; Murphy et al., 2003; Seah et al., 2016). A loss-of-function mutation of *ceh-60*, a transcription factor that activates the expression of all six Vtg genes, extends lifespan by 35% (Dowen, 2019). Conversely, overexpression of *vit-2::gfp* suppresses the longevity phenotype of *daf-2, glp-1*, and *eat-2* mutants, although it has no effect on WT lifespan (Seah et al., 2016).

Mechanistic explanation of this negative relationship between Vtg expression and *C. elegans* lifespan is provided by a series of in-depth investigations in recent years (Ezcurra et al., 2018; Kern et al., 2020; Murphy et al., 2003; Sornda et al., 2019; Wang et al., 2018). It is shown that *C. elegans* does not shut down Vtg production in the intestine after the worm lays the last eggs, which happens typically by AD 5 under the standard culture condition at 20°C. In fact, the yolk protein levels continue to increase up till AD 14, accompanied by atrophy of the intestine, growth of the so-called uterine tumors, and a notable increase of pseudocoelomic yolk patches (previously called pseudocoelomic lipoprotein pools, renamed because these “pools” are too small in young adults) (Ezcurra et al., 2018; Kern et al., 2020; Sornda et al., 2019; Wang et al., 2018). Knocking down the Vtg transcripts is shown to ameliorate all aging pathologies described above, and to extend lifespan (Ezcurra et al., 2018; Sornda et al., 2019; Wang et al., 2018). Therefore, post-reproductive vitellogenin production promotes senescent pathologies and accelerates aging (Ezcurra et al., 2018).

Interestingly, it was recently found that this seemingly self-harming act of Vtg production by post-reproductive hermaphrodites is actually beneficial to the reproductive fitness of *C. elegans*, for the yolk vented by old worms can be consumed by larvae and thus promote larval growth (Kern et al., 2021).

Although the senescent pathologies related to Vtg/YP have been investigated in detail in the *C. elegans* system by means of genetics or molecular biology, they have not been examined systematically using immuno-electron microscopy (immuno-EM). In the previous EM studies of *C. elegans* yolk proteins, the lipid membrane structures were not preserved in the best way, and the somatic tissues were missed as the focus was placed on the gonad and the pseudocoelom (Britton & Murray, 2004; Hall et al., 1999; Herndon et al., 2002; Paupard et al., 2001).

Here, using high-pressure freezing to preserve membrane structures and immuno-gold labeling, we inspected age-dependent changes of vitellogenin vesicles (VVs), pseudocoelomic yolk patches (PYPs), and yolk organelles (YOs) in multiple tissues. We find that in post-reproductive hermaphrodites of AD 6 and AD 9, intestinal VVs, which are 0.2 μm in diameter on AD 2, fuse with one another at high frequencies and form VVs that are 3-4 μm in median diameter. Occasionally, intestinal VVs of AD 6 and AD 9 worms can exceed 10 μm in diameter and fill up the cytoplasmic space of intestinal cells. For PYPs, we identified two subtypes based on the density of anti-YP170B gold particles. Only the high-density ones accumulate in post-reproductive animals. YOs in oocytes become slightly smaller, from ~0.5 μm in diameter on AD 2 to ~0.4 μm on AD 6 and AD 9. Unexpectedly, YOs, which should be limited to oocytes, are found mislocalized in the hypodermis and in the gonad sheath in post-reproductive worms. Both YOs and the membrane-less yolk are seen in high abundance in the tumor-like masses or oocyte clusters in the uterus, confirming the notion that YP complexes fuel the growth of the uterine tumors. Graphical summary and table 1 summarize the age-dependent changes of Vtg/YP-containing structures as found in this and the earlier EM studies.

**Table 1.** Age-dependent changes in the distribution of yolk proteins in *C. elegans* hermaphrodites.

|  |  |  |  |
| --- | --- | --- | --- |
| Tissues | AD 1/AD 2 | AD 6/AD 9 | AD 18 |
| intestine | VITs are found localized in the rough ER, the Golgi apparatus, and VVs. The median diameter of VVs: 200 nm (Fig. 2A-B, E). Exocytosis likely mediates VITs secretion from the intestine to the pseudocoelom (Zhai et al., 2022). | The median diameter of VVs increased to 3-4 μm, occasionally >10 μm. VVs frequently fuse with one another and likely expand in this way (Fig. 3). | Almost no detectable VVs in the intestine (file S3E). |
| pseudocoelom | Weak and diffuse VIT-2::GFP and VIT-1/3/6::mCherry signals can sometimes be seen in the pseudocoelom (Zhai et al., 2022). By DIC and fluorescence imaging, most PYPs look like droplets, <5 μm in diameter (Fig. S2). | PYPs increase dramatically (Fig. S2). PYPs can fuse with each other (video 1). | PYPs are dispersed throughout the pseudocoelomic space (Fig. S3E). |
| sheath cell | No YPs found inside sheath cells (Hall et al., 1999). | Occasionally, YOs are found inside sheath cells (Fig. 6E). | Not seen in sheath cells |
| oviduct | YPs are barely detectable | YPs accumulate in the oviduct (Fig. D-F, Fig. S3E, G). |  |
| oocyte | Mature oocytes have more YOs than less mature ones, ~500 nm in diameter. | The diameter of YOs decreases slightly to ~400 nm (Fig. 5A-D). | YOs are still present (Fig. S3E, G). |
| eggs or uterine tumors in the uterus | YOs in eggs look the same as YOs in oocytes, with a diameter of ~500 nm (Zhai et al., 2022). | There are YOs and amorphous yolk in uterine tumors (Fig. 5E-G, and Fig. S3E, I), and the diameter of YOs inside uterine tumors is ~400 nm (Fig. S3J). |  |
| uterine cells | Not seen | Yolk substances are frequently seen inside the uterine cells of WT worms at AD 6/9/18 (Fig 6E, Fig. 3SH). |  |
| distal gonad | Not seen | Although in worms expressing *vit-2::gfp* and *mCherry*-tagged *vit-1/6*, weak fluorescent signals were detected in the distal gonad on AD 6-9 (Fig. S3A-D), immuno-EM did not detect YPs in the distal gonad of WT worms of the same age (Fig. S3E). |  |
| hypodermis | Not seen | YOs found inside the hypodermal cells of WT worms on AD 6, 9, and 18 by Immuno-EM |  |
| muscle | Yolk proteins were not detected by immuno-EM in the body wall muscles of WT worms of any age. This is in disagreement with a published study using a GFP reporter (Turek et al., 2021). |  |  |

The immuno-EM documentation of Vtg/YP-related senescent pathologies in this study confirms and extends earlier studies (Ezcurra et al., 2018; Wang et al., 2018) at the ultrastructural level. Our data indicate that Vtg/YP-related senescent pathologies affect more tissues than previously thought and that vesicular fusion is a prominent and previously unknown aspect of those pathological phenotypes. The increase of VV-occupied regions in the intestine and the accumulation of PYPs suggest that gut-to-yolk biomass conversion occurs both inside and outside of the intestine.

## RESULTS

### The size of intestinal VVs increases dramatically with age

We examined the age-associated morphological changes of vitellogenin/yolk protein (Vtg/YP) containing structures in wild-type *C. elegans* by immuno-EM. Specifically, we used an anti-VIT-1/2 antibody and indirectly conjugated colloidal gold particles to label vitellogenin vesicles (VVs), PYPs, and yolk organelles (YOs). We compared these three types of Vtg/YP structures seen in reproductive young adults (adult day 2 or AD 2) with their counterparts in post-reproductive adults (AD 6 and AD 9) and quantified comprehensively the ultrastructural changes.

For context, we illustrate the known developmental relationships between these three Vtg/YP structures in figure 1. Briefly, vitellogenins are synthesized in the adult intestine, packed into VVs, and then exocytosed out of the intestine to become PYPs (Zhai et al., 2022). Through openings in the gonad sheath, oocytes take up pseudocoelomic yolk and store it in YOs (Hall et al., 1999). The characteristics of VVs, PYPs, and YOs in AD 2 hermaphrodites are presented in figure 1, as a reference for comparison later with the same structures in older worms.

**Figure 1.**
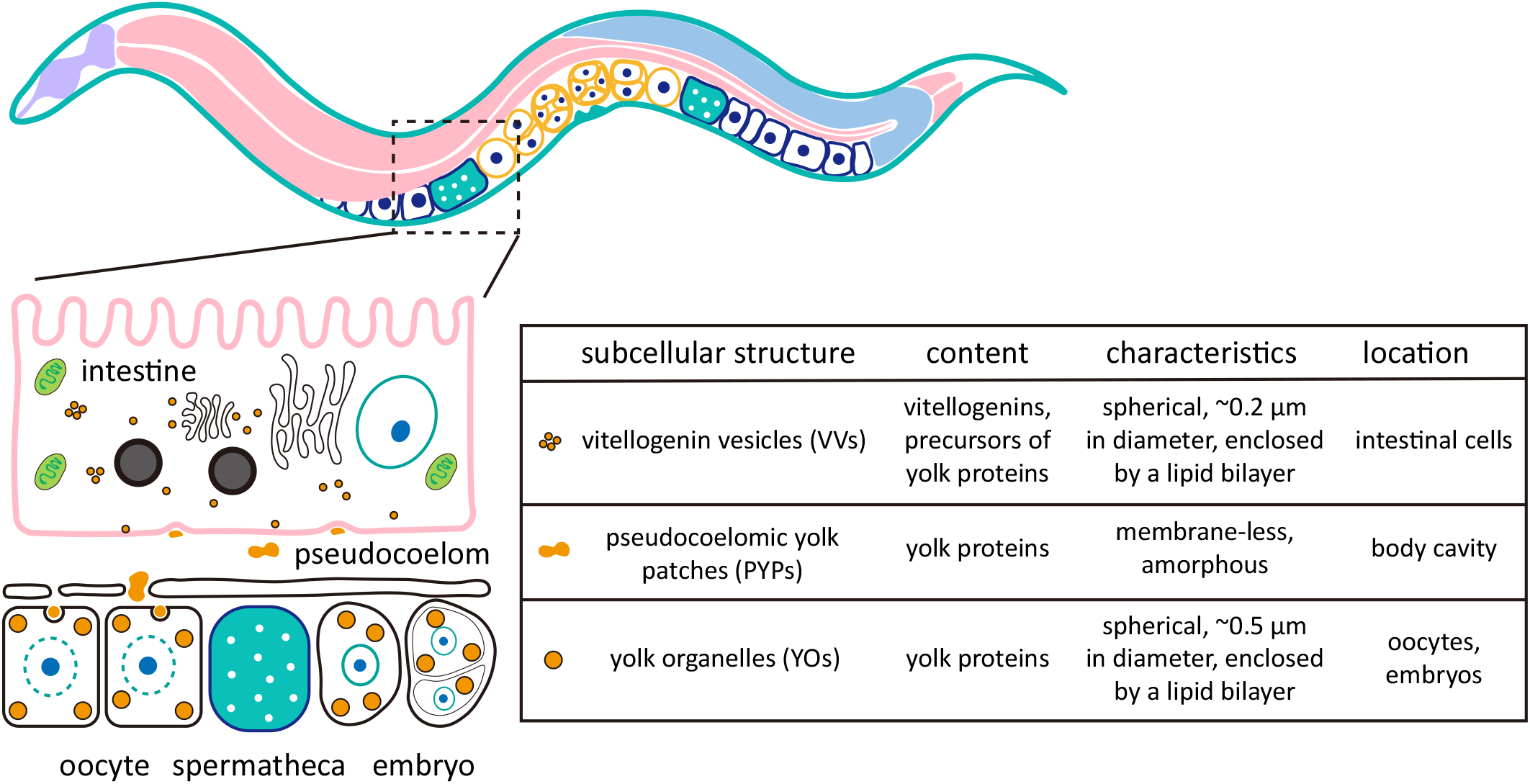
Vitellogenin- or yolk protein-containing structures in young adult *C. elegans.* The cartoon illustrates that vitellogenins are synthesized in the intestine, and secreted through exocytosis to the pseudocoelom where vitellogenins are processed into mature yolk proteins. Oocytes take up yolk proteins and store them in yolk organelles. The characteristics of VVs, PYPs, and YOs are summarized on the right.

Figure 2 displays the micrographs of VVs on AD 2, AD 6, and AD 9. On AD 2, VVs are 0.2 μm in median diameter (Fig. 2A-B). On AD 6 and AD 9, the median diameter of VVs expanded to 3 μm and 4 μm, respectively (Fig. 2C-E). These data indicate that the volume of VVs has enlarged by 3,000-8,000 times going from AD 2 to AD 6 and AD 9. Using lipid droplets in the same micrographs as a visual reference, AD 2 VVs look minuscule, whereas AD 6 and AD 9 VVs appear gigantic (Fig. 2A-D). In the extreme case, AD 6 and AD 9 VVs can reach above 10 μm across (Fig. 2E) and fill up almost the entire cellular space of an intestinal cell (Fig. 2F-G).

**Figure 2.**
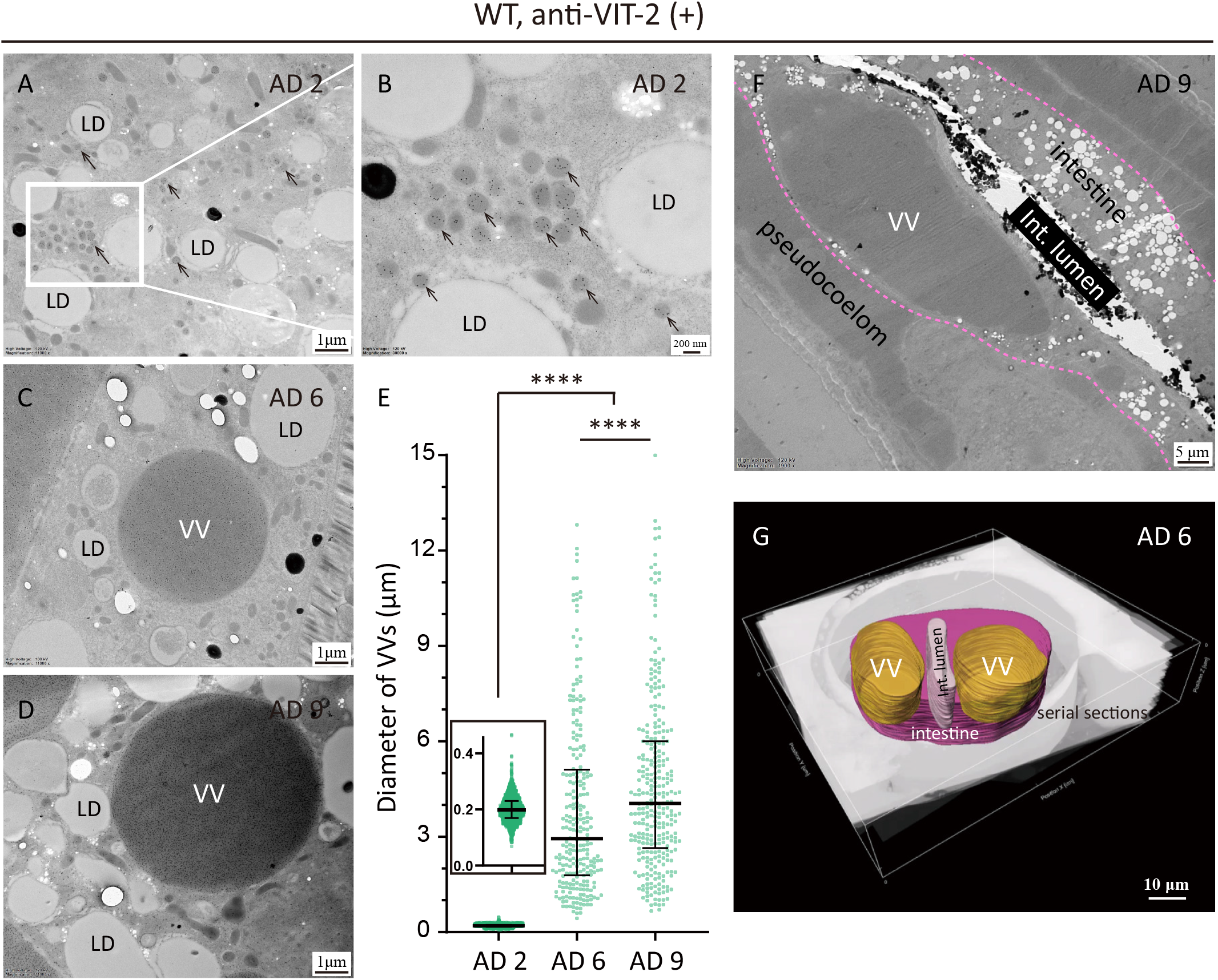
The size of intestinal VVs increases dramatically in older worms. (A-D, F) Immuno-EM images of WT worms at AD 2 (A, B), AD 6 (C), and AD 9 (D, F), labeled with the VIT-2 antibody. Black arrows point to VVs. Int.: intestine; LD: lipid droplet. The basal membrane of the intestine is traced by pink dotted lines (F). (E) Quantification of the diameters of VVs in AD 2, AD 6, and AD 9 *C. elegans* from 217, 127, and 129 micrographs, respectively. Median and the interquartile range are indicated. \*\*\*\**p* < 0.0001; one-way ANOVA with Tukey’s multiple comparisons test was performed to compare with each two data sets. (G) A 3-dimensional reconstruction of two VVs and a part of the intestine, based on 200 serial sections imaged by scanning electron microscopy (SEM). The intestine, the intestinal lumen, and VVs are colored in pink, grey, and yellow, respectively.

We also examined the size of VVs of the three *gfp*- and *mCherry-tagged* vits strains *(vit-1::mCherry vit-2::gfp, vit-2::gfp vit-3::mCherry, vit-2::gfp; vit-6::mCherry)* via fluorescent microscopy. The size of VVs increased dramatically with age (from AD 2 to AD 9) (Fig. S1A-O). For example, in *vit-1::mCherry vit-2::gfp* KI worms, the median diameter of VVs increased from 1.4-micron meters at AD 2 to 12.9-micron meters at AD 9 (Fig. S1P), and the latter ones occupied almost the space of the intestinal cell.

### Fusion of intestinal VVs

Fusion events between two or multiple VVs are readily detectable in post-reproductive worms (Fig. 3A-D). Quantification of the occurrence frequency of VVs caught in the middle of a fusion event indicates that on AD 6 and AD 9, 49.4% and 29.7% of the VVs captured in micrographs are respectively in the act of coalescing with one another (Fig. 3E). In comparison, this number is only 6.2% for the VVs captured in micrographs on AD 2 (Fig. 3E). These findings suggest that intestinal VVs grow by fusion in older worms.

**Figure 3.**
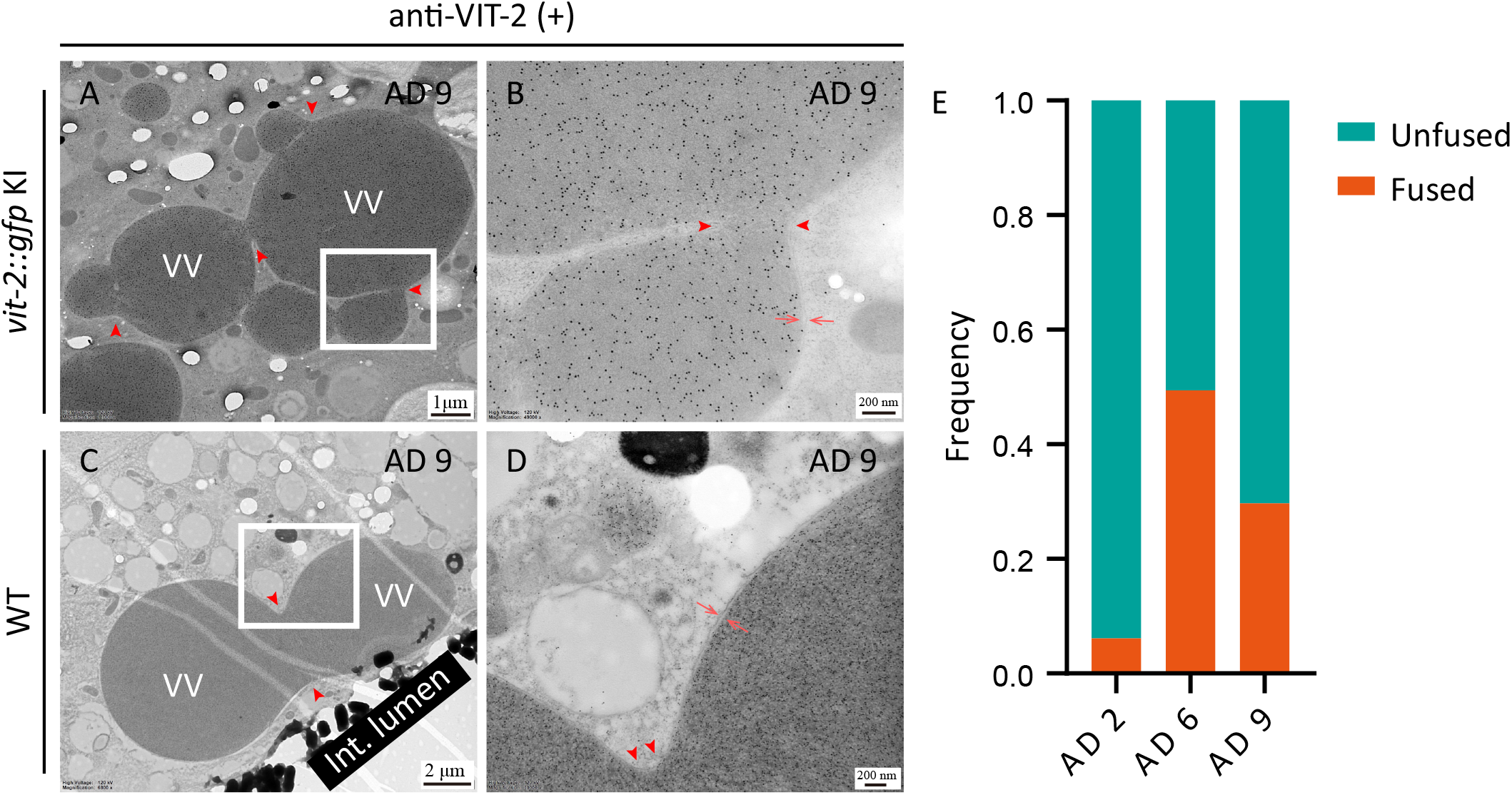
Fusion of intestinal VVs. (A-D) Anti-VIT-2 immuno-labeling of a *vit-2::gfp* KI worm (A, B) and a WT (N2) worm (C, D), both on AD 9. Red arrowheads point to the fusion sites between VVs, and paired red arrows point to the membrane. (E) Frequency of fused VVs and that of unfused VVs as observed in the immuno-EM images of WT worms. A total of 79, 92, and 84 micrographs were analyzed for AD 2, AD 6, and AD 9, respectively.

### Yolk accumulated in the expanded pseudocoelom in older worms

Turning from the intestine to the pseudocoelom, we found that AD 2 PYPs were categorically distinct from the AD 6 and AD 9 counterparts. Although gold particle labeling is found throughout the pseudocoelom regardless of the age of the adult worm, the density of immuno-gold particles attached onto AD 2 PYPs is markedly lower compared to AD 6 and AD 9 PYPs (median value: 19, 130, and 90 gold particles per μm^2^ for AD 2, AD 6, and AD 9, respectively) (Fig. 4A-D). This suggests that the concentration of yolk proteins of AD 6 and AD 9 PYPs is more than 4 times as much as that of AD 2 PYPs. This confirms the previous observations of pseudocoelomic yolk accumulation in old worms by fluorescence microscopy (Ezcurra et al., 2018; Garigan et al., 2002; Herndon et al., 2002). Supporting this conclusion, analysis of the pseudocoelom by conventional EM showed that they have relatively low electron density (expressed as grey value in micrograph) on AD 2 and high electron density on AD 6 and AD 9 (median value 13.1, 31.3, and 31.6, respectively) (Fig. 4E-H). Drawing a cutoff of 50 gold particles/μm^2^ for immuno-EM (Fig. 4D) or a grey value of 20 for conventional EM (Fig. 4H), we classified PYPs into two categories: the low-density ones are predominant on AD 2 and the high-density ones are predominant on AD 6 and AD 9.

**Figure 4.**
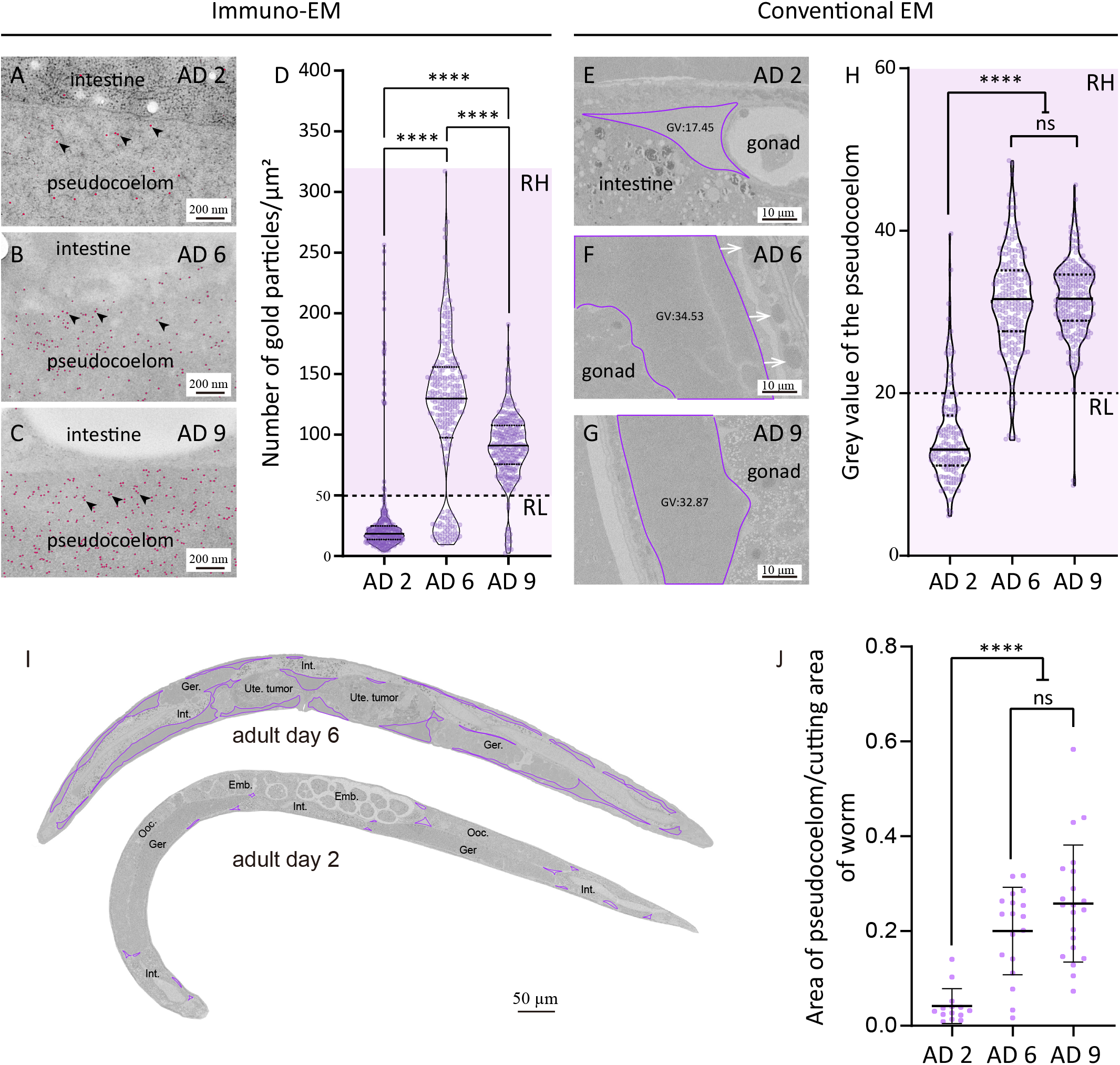
Yolk accumulates in the expanded pseudocoelom in older worms. (A-C) Anti-VIT-2 immuno-EM images of pseudocoelomic regions bordering the intestine of AD 2, AD 6, and AD 9 WT worms. Red dots mark the 10-nm gold particles, some of which are indicated by black arrowheads for clarity. The quantification results are shown in (D). (D, H) Violin plots showing the median and interquartile of each data set. Every point represents one value measured of one selected rectangle region in the pseudocoelom. Four regions were selected from each image. RH: region of high electron density; RL: region of low electron density. (E-G) Conventional SEM images of WT worms at the indicated age, focusing on the pseudocoelomic space traced by purple lines. GV: grey value. The quantification results are shown in (H). (I) Longitudinal sections of WT worms on AD 2 and AD 6. Purple lines trace the pseudocoelomic space. Int., Ger., Ooc., Ute., Emb., and Pha. are abbreviations for intestine, germline, oocyte, uterus, embryo, and pharynx. (J) Fraction of the pseudocoelomic space in longitudinal sections. In (D), (H), and (J), the median and the interquartile range are shown for each data set. For AD 2, AD 6, and AD 9, respectively, 75, 62, and 66 micrographs were quantified in (D), 40, 53, and 62 micrographs in (H), 11, 12, and 11 worms were stitched and analyzed in (J). \*\*\*\**p* < 0.0001; \*\**p* < 0.01; ns: not significant; one-way ANOVA with Tukey’s multiple comparisons test.

To characterize the dynamic process of pseudocoelomic yolk accumulation with age, we examined PYPs in worms expressing VIT-2::GFP and mCherry-tagged VIT-1/3/6. From AD 1 to AD 4, most PYPs look like droplets, and they can fuse to become bigger ones (Fig. S2A-I, and video 1). Video 1 shows that fusion is fast and dynamic. In post-reproductive worms, PYPs are milk-like and dispersed throughout the pseudocoelom (Fig. S2J-0). Video 2 shows that milk-like PYPs slosh back and forth as the worm moves.

As more PYPs accumulate in the pseudocoelom of older animals, the pseudocoelomic space expands. Using longitudinal EM sections, we quantified the pseudocoelomic area relative to the area occupied by the worm and found that from AD 2 to AD 6 and AD 9, the relative pseudocoelomic area increased from 4% to 20% and 26%, respectively (Fig. 4I-J).

To summarize, in post-reproductive *C. elegans*, while the intestine continues to produce vitellogenins and secret YP complexes to the pseudocoelom, large amounts of high-density PYPs accumulate in and expand the pseudocoelom.

### YOs in oocytes hardly changed with age

In contrast to the dramatic changes of VVs in the intestine and of yolk in the pseudocoelom, YOs found in oocytes remain unchanged by and large. The diameter of YOs decreases only very slightly, from an average of 0.5 μm on AD 2 to 0.4 μm on both AD 6 and AD 9 (Fig. 5A-D).

**Figure 5.**
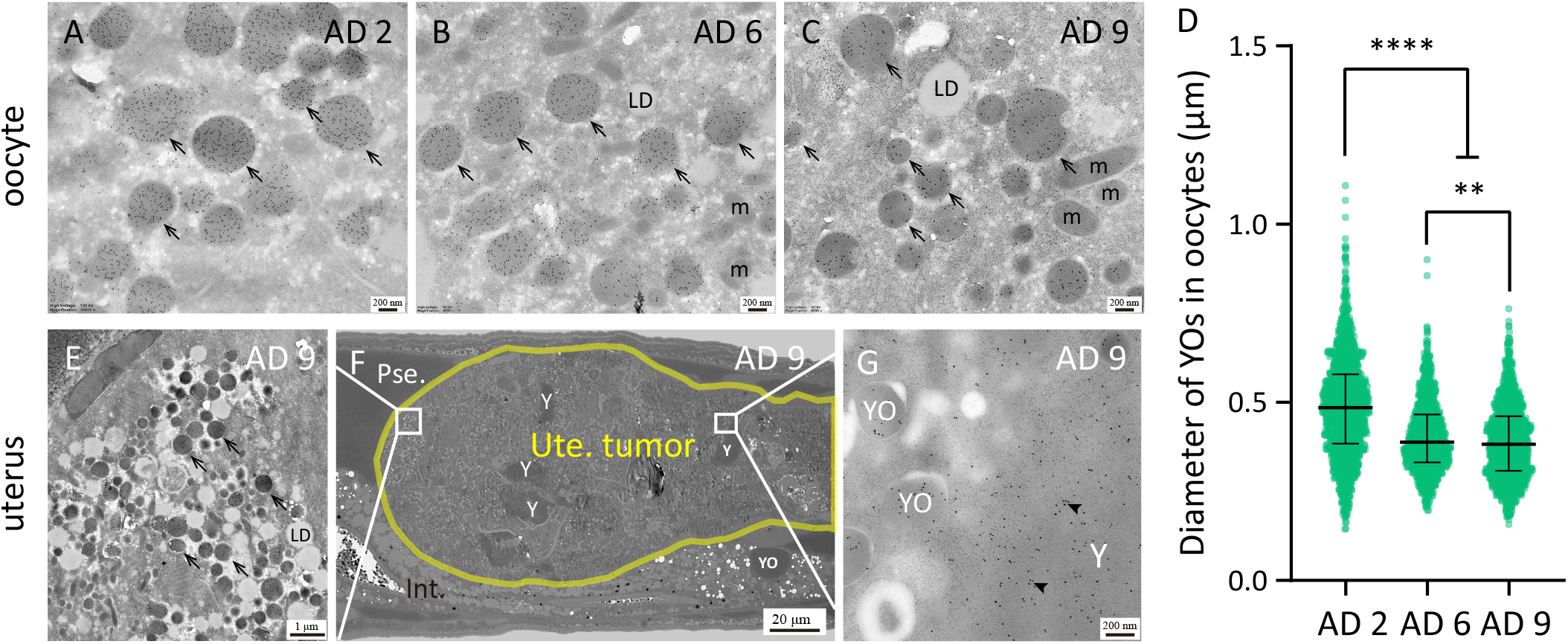
Gonadal YOs become slightly smaller in older worms. (A-C) Anti-VIT-2 immuno-EM images of WT worms on AD 2, AD 6, and AD 9. Black arrows indicate YOs in oocytes. (D) The diagram shows the median and interquartile of each data set. Every point represents one YO. A total of 30, 58, and 37 micrographs of WT worms on AD 2, AD 6, and AD 9 were analyzed, respectively. All YOs were measured for every micrograph analyzed. \*\*\*\**p* < 0.0001; \*\**p* < 0.01; one-way ANOVA with Tukey’s multiple comparisons test. (E-G) Anti-VIT-2 immuno-EM images of YOs and membrane-less yolk in a uterine tumor found in a WT worm on AD 9. The yellow line delineates the uterine tumor. Pse., Int., Ute., Y, m, and LD are abbreviations for pseudocoelom, intestine, uterus, yolk substance, mitochondrion, and lipid droplet, respectively.

Post-reproductive worms frequently develop uterine tumors, which originate from oocytes (Wang et al., 2018). We detected in uterine tumors immuno-gold labeling in two types of structures: those that looked exactly like YOs and those that were amorphous and not enclosed by a membrane, resembling PYPs (Fig. 5E-G). Although speculative, it seems plausible that these amorphous patches may originate from YOs after membrane rupture.

### Mislocalized yolk in the hypodermis and somatic gonad

Apart from pseudocoelom, we also observed that yolk substances also accumulated in the oviduct, which suggested that the yolk flood was overwhelming or the yolk endocytic capacity of old oocytes was compromised in old worms (Fig. 6D). In AD 6 and AD 9 but not AD 2 hermaphrodites, we observed YO-like structures in the hypodermal cells (Fig. 6A-C), the gonad sheath cells and the uterine cells (Fig. 6E-F). We verified that these mislocalized Vtg/YP structures are enclosed by a lipid bilayer membrane (Fig. 6C). These ectopic YOs in the hypodermis and uterine cells resemble the YOs found in oocytes, but can be much larger, sometimes reaching several micrometers in diameter (Fig. 6A-B, Fig. 7A-B, D-E).

**Figure 6.**
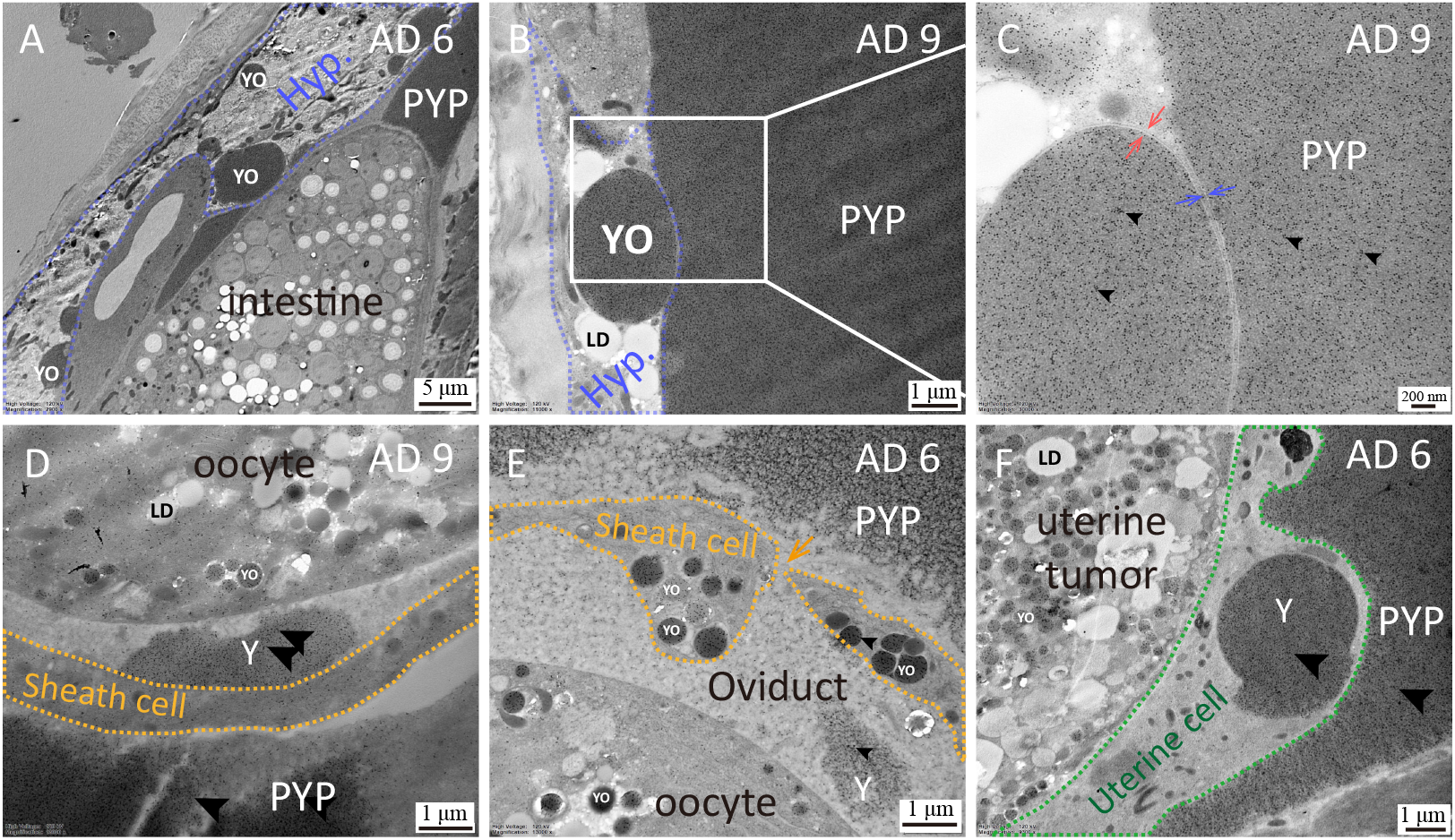
Mislocalized yolk in old worms. (A-F) Immuno-EM micrographs showing mislocalized YOs or membrane-less yolk in AD 6 (A, E-F) or AD 9 (B-D) WT *C. elegans.* (D) shows a patch of membrane-less yolk found in the cavity of the oviduct. YOs in the hypodermis (A-C) or the uterine cells (F) of old worms can be of a size of several micrometers or more. YOs also appeared inside the gonad sheath (E). In (F), yolk appeared to be leaking out of a YO (labeled “Y”) with a presumably ruptured membrane. Black arrowheads point to the 10-nm gold particles. Red and blue paired arrows indicate the lipid bilayer membrane of a hypodermal YO and a hypodermal cell, respectively. The blue and orange dotted lines mark the boundary of the hypodermal cells and the gonad sheath cells, respectively. The green dotted line outlines the uterine cell. The orange arrow points to the sheath pore. Hyp., PYPs and Y are abbreviations for hypodermis, pseudocoelomic yolk patches, and yolk substance.

### Absence of YO fusion in oocytes

We observed fusion of YOs not only in the intestine, but also in hypodermal cells and uterine cells (Fig. 7A-B, D-E). As worms age, the frequency of YO appearing in hypodermal cells increases, as does the frequency of fusion of hypodermal YOs, from 0% on AD 2 to 20% on AD 6 and then to 27% on AD 9 (Fig. 7C). The frequencies of YO appeared in uterine cells and sheath cells are low, and there are not enough images for quantification.

**Figure 7.**
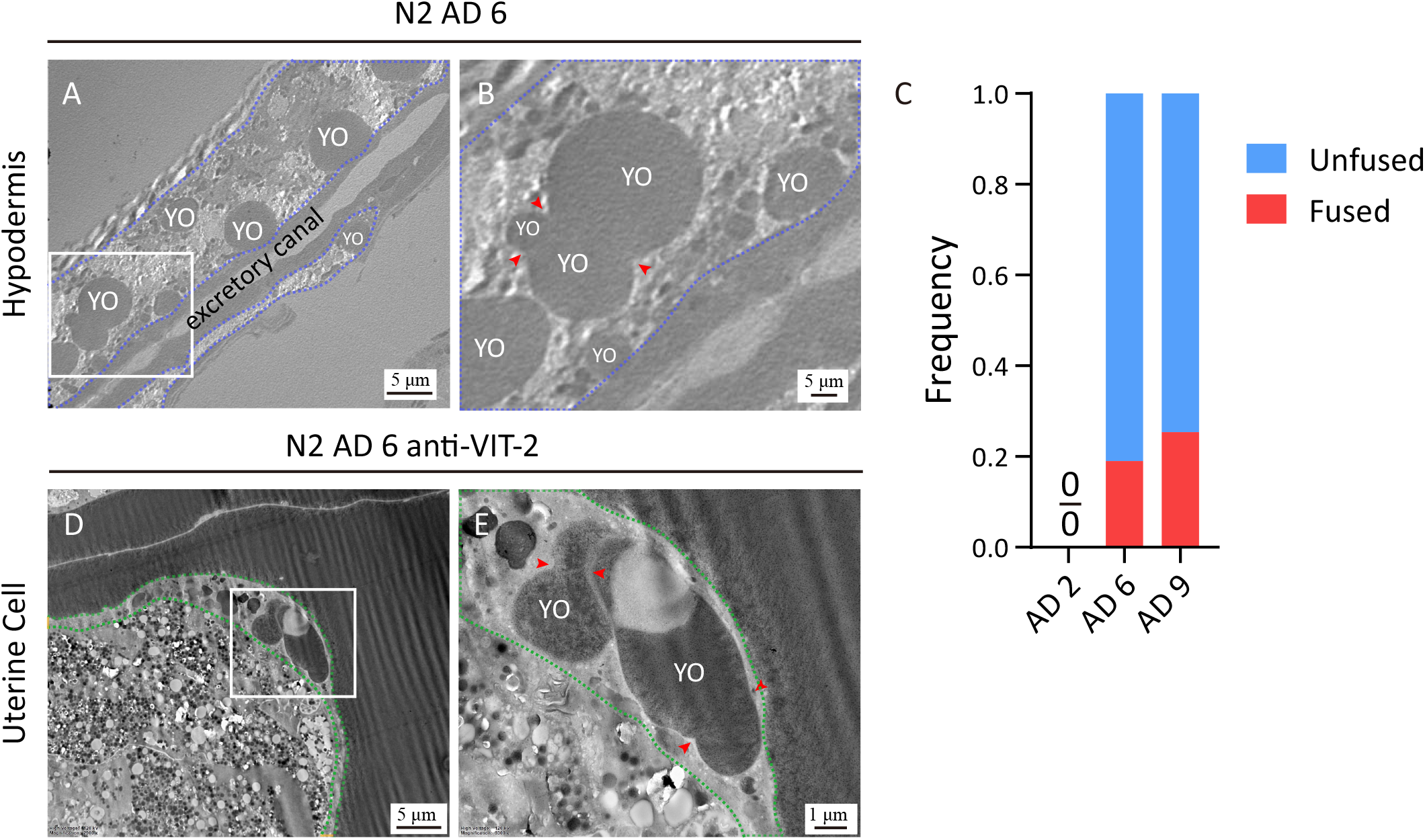
Fusion of YOs in the hypodermis and the gonad sheath. (A-B) Fusion of YOs in the hypodermal cells of a WT worm on AD 6. The blue trace marks the hypodermis. Arrowheads indicate the fusion sites. (C) Occurrence frequency of fused and unfused YOs seen in the hypodermis of AD 2, AD 6, and AD 9 WT worms, quantified from 15, 50, and 36 transmission electron microscopy (TEM) images, respectively. (D-E) Immuno-EM images show fused YOs inside the uterine cell, which is marked by green dotted lines.

YOs are abundant in the oocytes of post-reproductive adult worms, but fusion between oocyte YOs was not observed. Among all the cell types examined, it seems that oocytes have a mechanism to prevent YO fusion, while somatic cells do not.

### Intestinal atrophy and deterioration in older worms

Intestinal atrophy during aging was previously measured by the relative intestinal width, *i.e.*, subtracting the width of the intestinal lumen from the width of the intestine and then normalizing it against the width of the worm body (Ezcurra et al., 2018; Kern et al., 2020). Here, using stereological analysis, we quantified age-associated changes of the intestinal volume in both absolute and relative terms (Fig. 8B).

**Figure 8.**
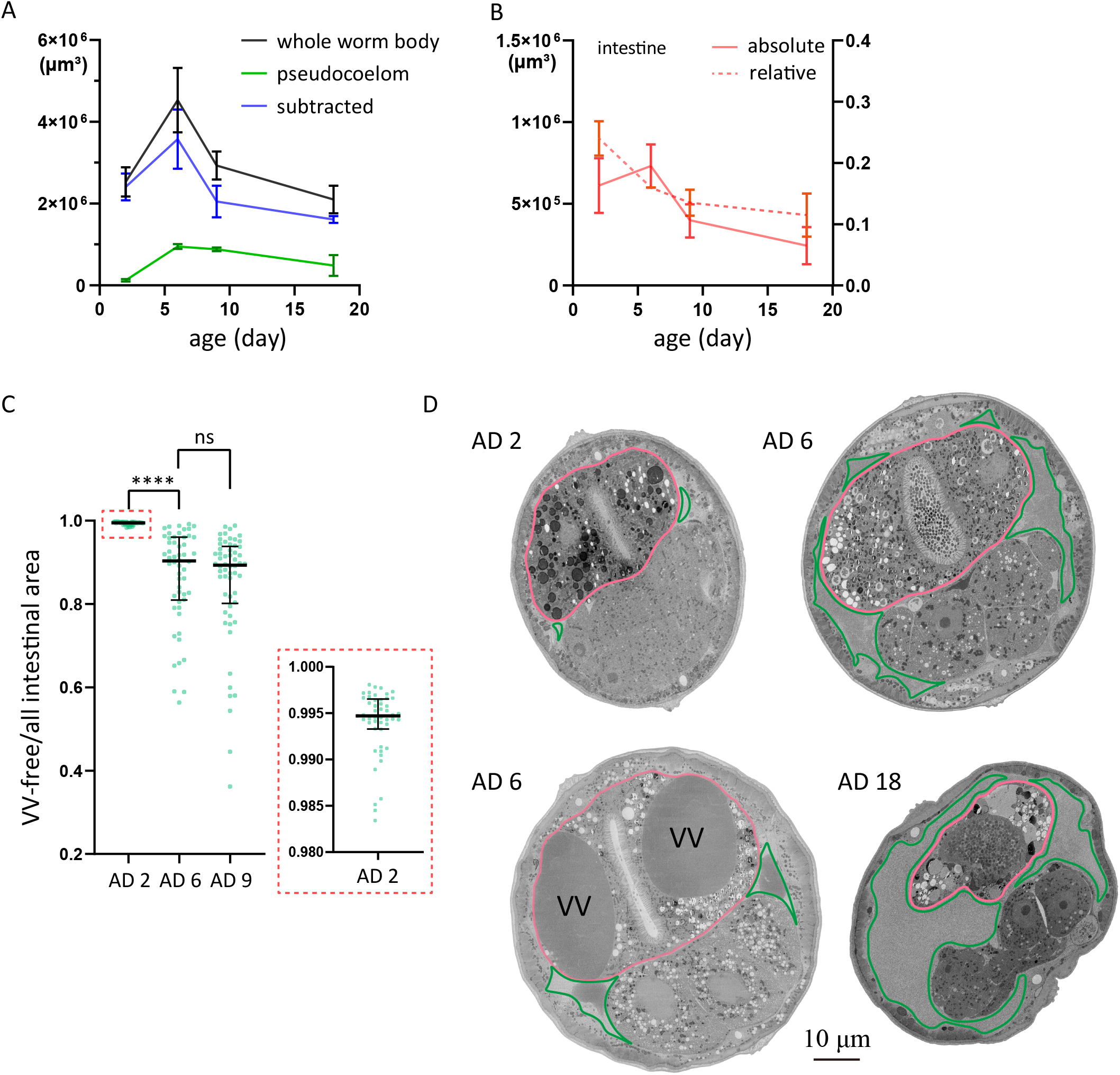
Quantification of age-associated intestinal atrophy in WT *C. elegans*. (A-B) Stereological analysis of the volume of the worm body, the pseudocoelom, and the intestine on adult days 2, 6, 9 and 18, based on serial EM sections from head to tail. Two worms were analyzed for each time point. (C) Fraction of VV-free intestinal area. A total of 49, 54, and 58 micrographs were analyzed for adult days 2, 6, and 9. \*\*\*\**p* < 0.0001; ns: not significant; one-way ANOVA with Tukey’s multiple comparisons test. (D) Micrographs contrasting the morphological of the intestine on AD 2, AD 6, and AD 18. The upper right micrograph is typical for AD 6, and the lower left one is an extreme example for AD 6. The other two micrographs are representative for AD 2 and AD 18, respectively. All images are cross-sections of the second intestinal ring, with pink and blue traces marking the intestine and the pseudocoelom, respectively.

Stereology is a methodology for quantifying three-dimensional characteristics by examining evenly spaced, two-dimensional sections that sample through an entire three-dimensional object (Ferguson et al., 2017). We take advantage of the Cavalieri principle to quantify the absolute volume of the tissue of interest. By analyzing the micrographs of 16-17 cross-sections that were evenly spaced from the head to the tail of a worm, we measured the absolute volume of the body, the pseudocoelom, and the intestine (excluding the luminal space). Two worms each were examined on AD 2, AD 6, AD 9, and AD 18 (Fig. 8A-B).

As shown, the absolute volumes of the worm body (with or without the pseudocoelomic space subtracted), the pseudocoelom, and the intestine all peaked on AD 6 (Fig. 8A-B). The relative volume of the intestine kept declining, from 21% on AD 2 to 11% on AD 18 (Fig. 8B). Hence, intestinal atrophy is evident after AD 6, but arguable from AD 2 to AD 6 because the absolute volume of the intestine increases (from 6.1E5 μm^3^ to 7.3E5 μm^3^) whereas the relative volume decreases.

Knowing that intestinal VVs grow dramatically from AD 2 to AD 6, with a 15-fold increase in diameter or > 3000-fold increase in volume (Fig. 2), we wondered whether this underlies the increase of the absolute volume of the intestine. From a random selection of immuno-EM sections, we quantified the total intestinal area and the summed area of VVs in each section and calculated the relative VV-free area. On AD 2, AD 6, and AD 9, the mean value of the percentage of VV-free intestinal area is 99.4%, 86.6%, and 84.8%, respectively (Fig. 8D). If the intestinal volume is corrected with the percentage of VV-free intestinal area, then the enlargement of the intestine on AD 6 becomes marginal (6.1E5 and 6.3E5 μm^3^ for AD2 and AD 6, respectively). This suggests that the apparent enlargement of the intestine on AD 6 can be accounted for by the expansion of VVs. In other words, the external enlargement reflects internal deterioration.

Some of the EM sections recorded impressive examples of intestinal deterioration. In figure 8D, a representative EM section of an AD 2 worm shows the normal structures, and the top right, of an AD 6 worm, displays the increased intestinal area in a cross-section compared with the one in AD 2. At the bottom left, a cross-section of AD 6 indicated the VV enlargement counted for the age-related increment in the absolute intestinal volume, while the VV-free volume didn’t get increase. The AD 6 micrograph features two large VVs and an expanded pseudocoelom (Fig. 8D). The two VVs almost fill up the entire cross-section of the intestine. At the bottom right, the intestine shrinks dramatically in worms at AD 18. What is shown in the EM images of figure 8D is consistent with the quantified results (Fig. 8A-C).

## DISCUSSION

The nematode *C. elegans* employs a precise mechanism to turn on the vit genes, so that their expression starts exactly at the beginning of adulthood and is usually limited to the intestine of a worm with a female gonad (Kimble & Sharrock, 1983; Klass et al., 1979). This makes sense because the purpose of *vit* genes is to generate nutrient supply for the progeny, but this is costly for the mother. Analogously, expression of vitellogenins of mated *C. elegans* males promotes post-mating death of those animals (Shi et al., 2017). Another intriguing phenomenon is that *C. elegans* hermaphrodites do not turn off these genes after the task of reproduction is completed. Post-reproductive mothers continue to make yolk at the cost of intestinal atrophy and shortened lifespan, which has been characterized in detail (Ezcurra et al., 2018; Sornda et al., 2019), albeit not at the EM level. Here, using immuno-EM, we observed a previously unreported aspect of this intestine-to-yolk biomass conversion: the intestinal atrophy starts internally before the intestine shrinks visibly. It has been shown that the relative intestinal width decreases by about one-third from AD 1 to AD 7 (by ~22% from AD 1-AD 4, and by ~25% from AD 1-AD 8) (Sornda et al., 2019), and by half or more after AD 11 (Ezcurra et al., 2018). The intestinal atrophy occurs internally in a concealed manner in addition to the visible shrinkage. In other words, the intestinal atrophy is worse than how it looks on the outside, as VVs grow huge from ~0.2 μm to 3-4 μm across, and occupy more and more space inside the intestine (Fig. 8C).

In contrast, YOs in oocytes are able to maintain a nearly constant size, with a diameter of 0.5 μm on AD 2 and 0.4 μm on AD 6 and AD 9. This seems to be a unique property of oocytes, because mislocalized YOs in AD 6 and AD 9 hypodermal cells or uterine cells can be several micrometers across, almost as big as the VVs in old intestinal cells.

We find that VVs can grow bigger by fusion with one another (Fig. 3). Fusion between mislocalized YOs in somatic tissues was also seen (Fig. 7). In contrast, no fusion events were detected for YOs in oocytes, nor for YOs in uterine tumors, which originate from oocytes. We thus conclude that oocytes have a mechanism to prevent fusion between YOs, which is worth investigating in the future.

RME-2 is the only yolk protein receptor so far identified in *C. elegans* (Grant & Hirsh, 1999). Only oocytes express *rme-2* and only late-stage oocytes have an abundance of RME-2 on the cell surface (Grant & Hirsh, 1999). YOs form through RME-2-mediated endocytosis of yolk from the pseudocoelom. Normally, YOs are present only in oocytes and after fertilization, in embryos. It is unclear how the somatic cells of old worms acquire YOs. It could result from mis-expression of RME-2 in the hypodermis, gonad sheath, and uterine cells of old worms, or through an RME-2 independent mechanism.

Compared with the dramatic aging pathologies associated with Vtg/YP, there is only a modest lifespan extension of 20% or so by knocking down the expression of all six vitellogenin genes (Ezcurra et al., 2018; Murphy et al., 2003; Seah et al., 2016; Sornda et al., 2019). This could suggest that post-reproductive production of yolk proteins may be less detrimental to the mother than one might expect from the associated, severe-looking aging pathologies. Alternatively, we speculate that post-reproductive production of yolk proteins, although costly to the mother, might also benefit the mother in some way. It has been shown that Vtgs scavenge oxidants, and enhance immunity in honeybees (Park et al., 2018). Although knocking down the vitellogenin genes did not make worms more resistant to oxidants (Sornda et al., 2019), in those experiments, RNAi started at L4 and the treated worms were assayed on adult day 1 (Sornda et al., 2019). Mutations of multiple vit genes have been shown to cause Vtg accumulation and ER stress in the intestine and also sensitivity to pathogenic *P. aeruginosa* (Singh & Aballay, 2017), but the immunity defects are likely a secondary phenotype of ER stress. In any case, it remains to be tested whether post-reproductive Vtg production affords protection to the mother from oxidants or pathogens.

## MATERIALS AND METHODS

### Worm Culture and Strains

*C. elegans* was fed with *E. coli* OP50 on nematode growth medium (NGM) plates and cultured at 20°C. To produce synchronized cohorts of worms, 25 gravid hermaphrodites were put on a plate and allowed to lay eggs for 4 hours before being taken away. Worms were regarded as one-day-old within 24 hours after reaching sexual maturity. In this study, two worm strains were used, including wild type (N2) and BCN9071 *vit-2(crg9070[vit-2::gfp]) X;* both were from the Caenorhabditis Genetics Center.

### Antibodies

The rat polyclonal anti-VIT-2 antibody (diluted 1:100 for immuno-EM labeling) was kindly provided by Dr. Xiao-Chen Wang (Institute of Biophysics, Chinese Academy of Sciences, Beijing, China) (Liu et al., 2012). The epitope of the antibody is a recombinant protein VIT-2 (83-620 amino acid)::6xHIS. The rabbit-derived second antibody (anti-rat) conjugated with 10-nm colloidal gold (Sigma, lot #SLBZ8963) is available as a commercial product.

### Immuno-EM Methods

The immuno-EM workflow including sample preparation, sectioning, immuno-labeling, and transmission electron microscopy (TEM) imaging are described clearly (Zhai et al., 2022).

### Conventional EM Sample Preparation

Methods of conventional EM sample preparation were developed by Li et al. (Li et al., 2017). Based on the differences in worm samples, we adjusted the methods slightly. High-pressure freezing and freeze-substitution were used for sample fixation and dehydration as described before (Zhai et al., 2022). The one difference was that the component of the substitution solution contained 1% OsO_4_, 0.1% UAc, and 98.9% acetone. After that, samples were put into 1 ml UAc saturated acetone solution and stained for 3.5 hrs at room temperature on a shaker.

After dehydration, samples were infiltrated in SPI-PON 812 resin mixture. The pure resin mixture was made by mixing 19.5 g SPI-PON 812 (SPI-CHEM, lot #1230517), 10 g DDSA (SPI-CHEM, lot #1230425), 12 g NMA (SPI-CHEM, lot #1230514), and 1.5% (v/v) BDMA (SPI-CHEM, lot #1220601). Pure resin mixture and acetone were mixed as 1:3, 1:1, and 3:1 for sample filtration for 2 hrs, overnight, and 48 hrs, respectively. The worm cakes were taken out from carriers carefully using a pair of needles on a 1 ml syringe under a stereo microscope. Then, nematodes were separated, and individually transferred into a cell of the embedding plate that was already filled with the pure resin mixture. Polymerization was performed in a 60°C oven for 48 hrs.

### Serial Sectioning and Scanning Electron Microscopy Imaging

Serial sectioning and scanning electron microscopy (SEM) imaging were conducted according to the methods developed by Li et al. (Li et al., 2017). Before SEM imaging, the tapes carrying sections were adhered to SEM Cylinder Specimen Mounts (Electron Microscopy China, Cat. #DP16232) by carbon conductive double-faced adhesive tape (NISSHIN EM Co. Ltd, Japan). The specimen mounts carrying samples were transferred under the SEM (FEI Helios NanoLab 600i) equipped with a CBS detector. Images were acquired by the software xT microscope control (FEI, version 5.2.2.2898) and iFast (FEI) with parameter settings of 2 kV accelerating voltage, 0.69 nA current, and 5 μs dwell time.

### Image Analysis

All quantitative data came from the manual measurement of cellular structures by ImageJ software. The pixel size of the TEM images was calibrated using a standard sample (diffraction grating replica with latex spheres, TED PELLA, INC, prod. #673) at different magnifications, and the pixel size has been reported before (Zhai et al., 2022). The quantitative data were analyzed by GraphPad Prism 8.4.3.

### SEM Image Reconstruction

The reconstructed intestine and VVs in figure 2G were based on 200 serial 70 nm-thick sections. The methods for image alignment were described before (Li et al., 2017), and the aligned continuous images were processed with Imaris (version 9.0.1) for 3D reconstruction.

### Stereological Analysis

The methods of collecting whole worm serial SEM sections and SEM images were described by Li et al. (Li et al., 2017). For adults of different ages, each worm can be cut into over 10,000 serial sections in 50-to 70-nm thick (the exact thickness of every sample and images analyzed are in table S1 and file S1). Worms at AD 6 and AD 9 could even be cut into about 20,000 serial sections. For stereological analysis, 16 or 17 SEM images (attached in Supplementary File 1) were selected using the systematic random sampling method. The equation was based on the Cavalieri principle: Volume = T x Area_point_ x ∑points; where T means the interval thickness of every adjacent two sections sampled. T values of every worm are in table S1. Stereological Analyzer (version 4.3.3) software was used to show an evenly distributed point grid covered on an EM image of a cross-cut section. The Area_point_ in the equation means the absolute area of each point represented. Here, Area_point_ is set as 21.34 μm^2^ for worm body volume, and 5.34 μm^2^ for intestinal and pseudocoelomic volume. ∑points means the total number of point hits on the cellular structures of interest. Relative volume equals the absolute volume over the volume of the whole worm body.

## ACKNOWLEDGEMENT

We thank Dr. Xiao-Chen Wang of the Institute of Biophysics, Chinese Academy of Science, for providing the anti-VIT-2 antibody; Drs. Cheng-Gang Zhou and Bin Liang, both of Yunnan University, for providing the *ceh-60* mutant, the *vit-2* mutant, and the *vit-2::gfp* knock-in worm strains; and the Caenorhabditis Genetics Center, which is supported by the NIH Office of Infrastructure Programs (P40 OD010440), for providing the wild-type N2 strain. We thank Drs. Wan-Zhong He of Institute of Chemistry, Chinese Academy of Sciences (ICCAS) and Zhao-Di Jiang of National Institute of Biological Sciences (NIBS), Beijing for technical guidance in EM sample preparation. We thank Dr. Jian-Guo Zhang, Dr. Gang Ji, and Can Peng of the Institute of Biophysics, Chinese Academy of Science, for guidance in EM imaging. We are grateful to Dr. John Hugh Snyder for critical reading and editing of this manuscript. This work was funded by National Natural Science Foundation of China (NSFC-ISF 31925026 to F.S., 31501160 to X.-X.L., and 32061143020 to M.-Q.D.), and Ministry of Science and Technology of China (institutional grants to NIBS, Beijing, a fund of the National High-Level Talents Special Support Program to M.-Q.D.), Beijing Municipal Science and Technology Commission (institutional grants to NIBS, Beijing and a fund for cultivation and development of innovation base to M.-Q.D.).

## STATEMENT

The authors confirm that the data supporting the findings of this study are available within the supplementary material and corresponding authors, upon reasonable request. The work described is original research that has not been published previously, and is not under consideration for publication elsewhere, in whole or in part. No conflicts of interest exist in the submission of this manuscript.

## SUPPORTING INFORMATION

| Filename | Description |
| --- | --- |
| Figure_S1 | The size of VV increases dramatically with age seen by fluorescent microscopy |
| Figure_S2 | Yolk substances accumulate in the pseudocoelom with age seen by fluorescent microscopy |
| Figure_S3 | The distribution of yolk proteins in old worms |
| Table_S1 | Raw and processed data of the stereological analysis of *C. elegans* at the indicated ages |
| File_S1 | SEM images for stereological analysis of *C. elegans* at the indicated ages |
| Video_1 | Lipid droplet-like pseudocoelomic yolk patches move dynamically and fuse with each other in *C. elegans*. |
| Video_2 | The milk-like pseudocoelomic yolk patches flow in the pseudocoelom as the worm moves. |

---

**Figure S1.**
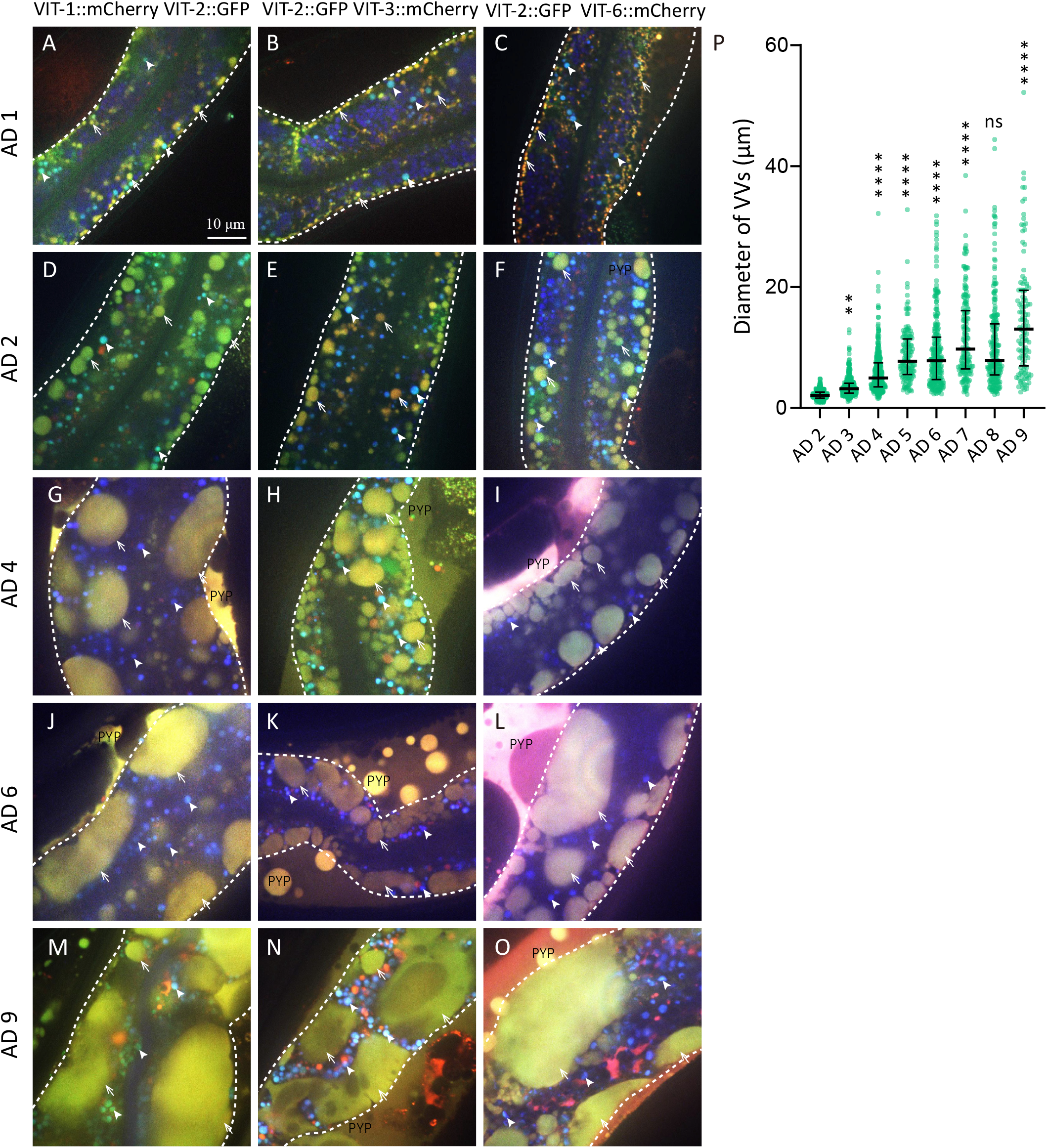
The size of VV increases dramatically with age seen by fluorescent microscopy. White arrows point to VVs, and arrowheads indicate gut granules. White dashed lines outline the intestine. PYP means pseudocoelomic yolk patch. (P) Quantification of the diameters of VVs based on fluorescent and DIC merged micrographs of *vit-1::mCherry vit-2::gfp* KI worms of different ages. 11, 11, 22, 15, 27, 24, 24, 22 micrographs were quantified for AD 2, 3, 4, 5, 6, 7, 8, 9, respectively. Median and the interquartile range are indicated. \*\*\*\**p* < 0.0001; \*\**p* < 0.01; ns: not significant; all compared to the day before; one-way ANOVA with Tukey’s multiple comparisons test was performed to compare each two data sets.

**Figure S2.**
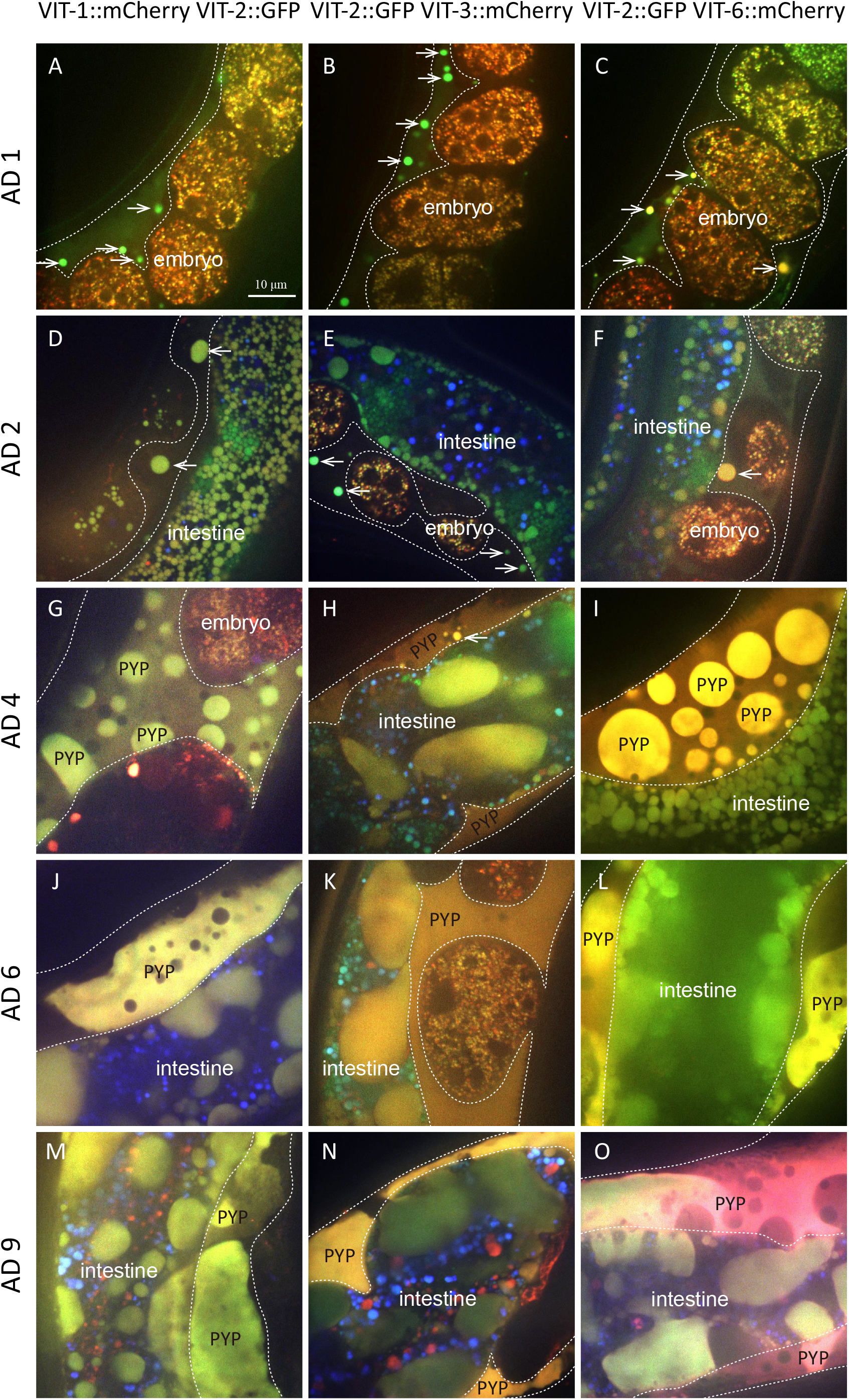
Yolk substances accumulate in the pseudocoelom with age seen by fluorescent microscopy. (A-O) White dots outline the pseudocoelomic region that can be recognized in epifluorescent images. White arrows and PYPs indicate pseudocoelomic yolk patches.

**Figure S3.**
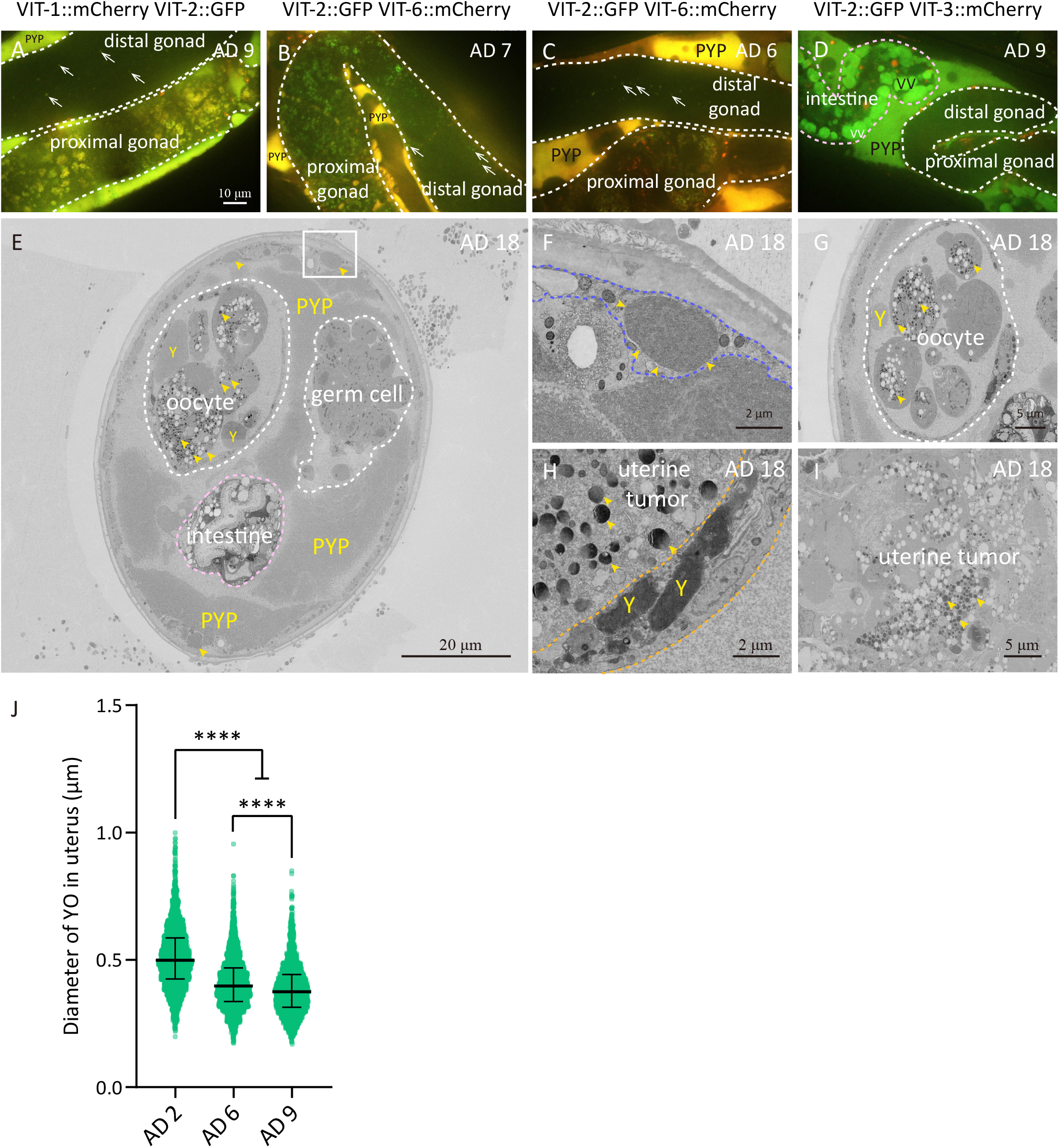
The distribution of yolk proteins in old worms. (A-D) White arrows indicate the fluorescent signals of yolk proteins. (E-I) SEM images show the cross-sections of old WT worms. White, pink, lavender-blue, and orange dots outline the gonad, intestine, hypodermis, and uterine cells, respectively. Yellow arrowheads point to YOs in the hypodermis (E, F), the oocyte (E, G), and uterine tumors (H, I). Y represents amorphous yolk. F is the larger view of the white rectangle region in E. (J) Quantification of the diameters of YOs in the uterus (that is embryos at AD 2 or uterine tumors at AD 6 and AD 9) is based on 50, 88, 73 immuno-EM micrographs. Median and the interquartile range are indicated. \*\*\*\**p* < 0.0001; one-way ANOVA with Tukey’s multiple comparisons test was performed to compare with each two data sets.

**Video 1. Lipid droplet-like pseudocoelomic yolk patches move dynamically and fuse with each other in *C. elegans.***

A *vit-2::gfp vit-3::mCherry* KI worm at AD 5 was imaged via fluorescent microscopy, and a view of the mid-body is shown in this video.

**Video 2. The milk-like pseudocoelomic yolk patches flow in the pseudocoelom as the worm moves.**

A *vit-2::gfp; vit-6::mCherry* KI worm at AD 6 was examined via fluorescent microscopy, and this video shows its posterior body.

