## Supplementary material for "Old Age Drives Fusion and Expansion of Vitellogenin Vesicles in the Intestine of *Caenorhabditis elegans*": file S1

**Supplementary File 1. Stereological Analysis of *C. elegans* at the indicated ages**

**Notes:**

- ✧ This file contains all the scanning electron microscopy (SEM) images of *C. elegans* used for stereological analysis. The voltage setting, magnification, scale bar, and other information related to EM imaging are indicated below each image.
- ✧ A total of eight worms were analyzed (two each for AD 2, 6, 9, and 18) by serial sectioning from head to tail. Each worm was given an ID following an "Age-number" format. For example, "day18-18(1)" means worm No. 1 of resin block No. 18 for adult day 18.
- ✧ For each worm, 16-17 evenly spaced sections were imaged by SEM and further analyzed stereologically. The images of the same worm are grouped together, and each image is named by its section number.
- ✧ Each SEM image was analyzed twice, once for the measurement of body volume and the second time for the measurement of tissue volumes.
- ✧ In each image, the "+" marks are uniformly distributed, and the checkmark on the "+" mark represents the attribution of cell structures. See "Materials and Methods" for details.
- ✧ All the marked images are hyperlinked to the table of CONTENT for a quick lookup.
- ✧ See Supplementary Table 1 for the raw statistics and parameters of stereological analysis of each image.

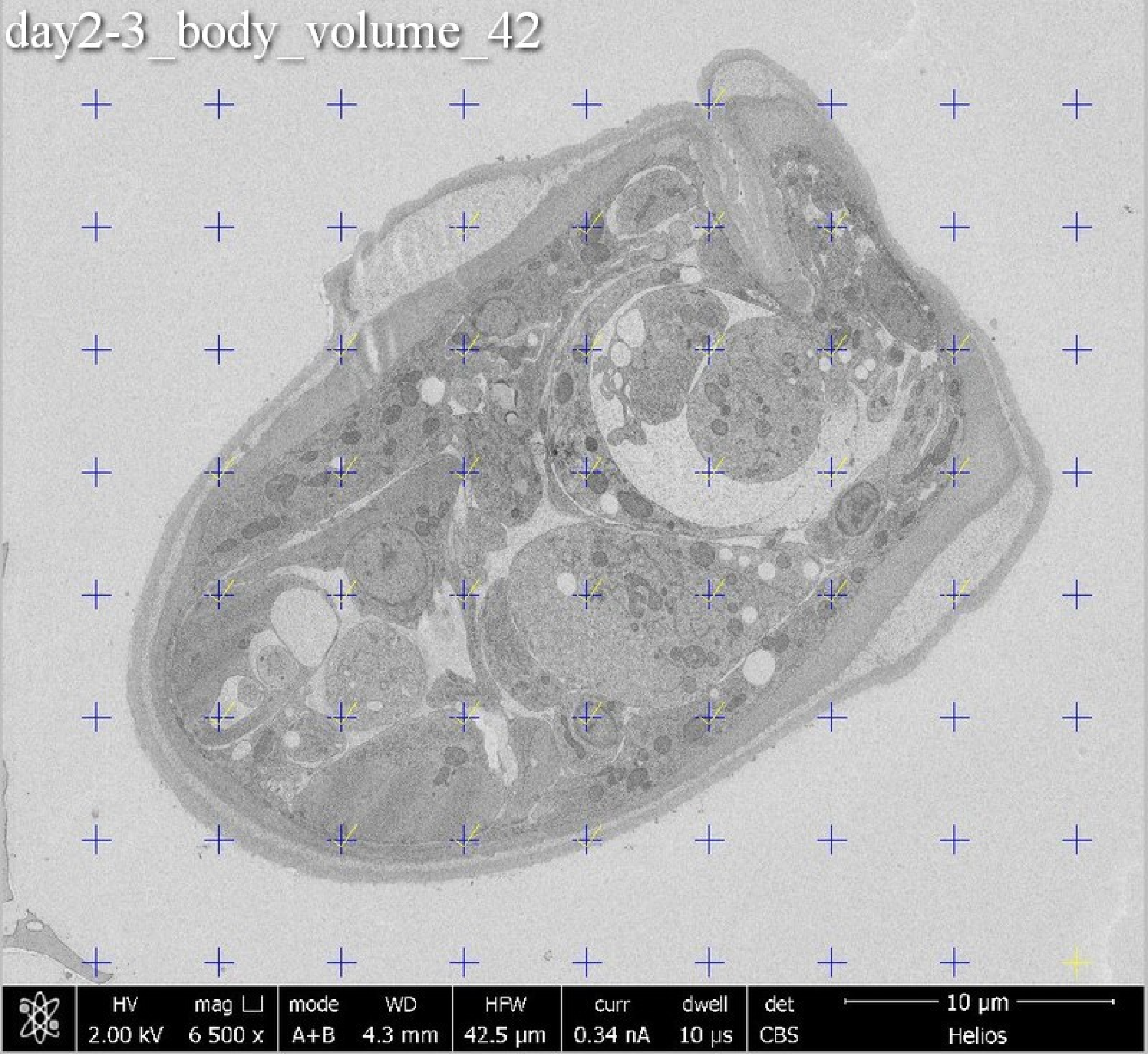
day2-3_body_volume_42
HV 2.00 kV
mag ⊔ 6 500 x
mode A+B
WD 4.3 mm
HFW 42.5 µm
curr 0.34 nA
dwell 10 µs
det CBS
10 µm
Helios

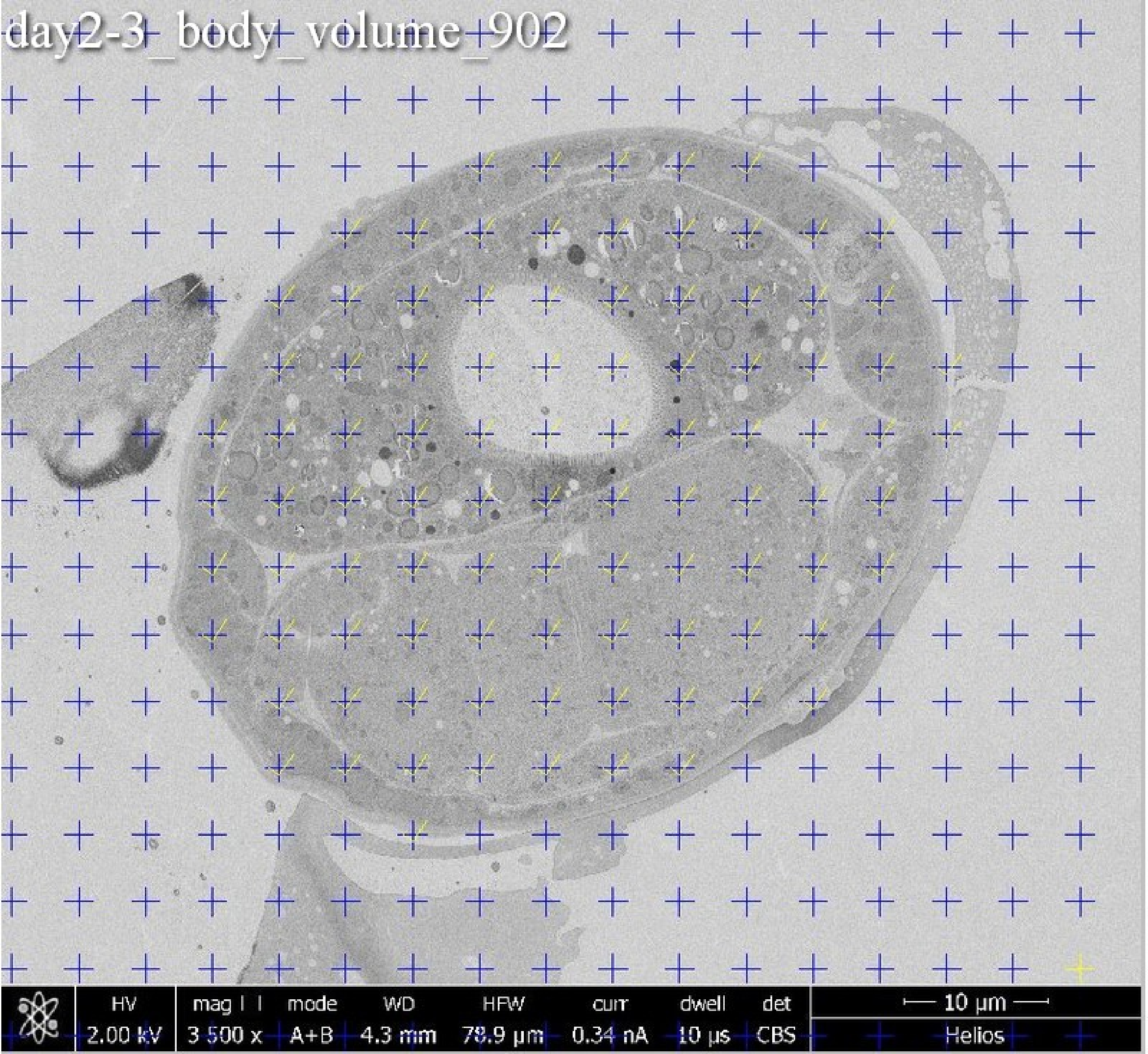
day2-3_body_volume_902
HV
2.00 kV
mag
3 500 x
mode
A+B
WD
4.3 mm
HFW
78.9 µm
curr
0.34 nA
dwell
10 µs
det
CBS
10 µm
Helios

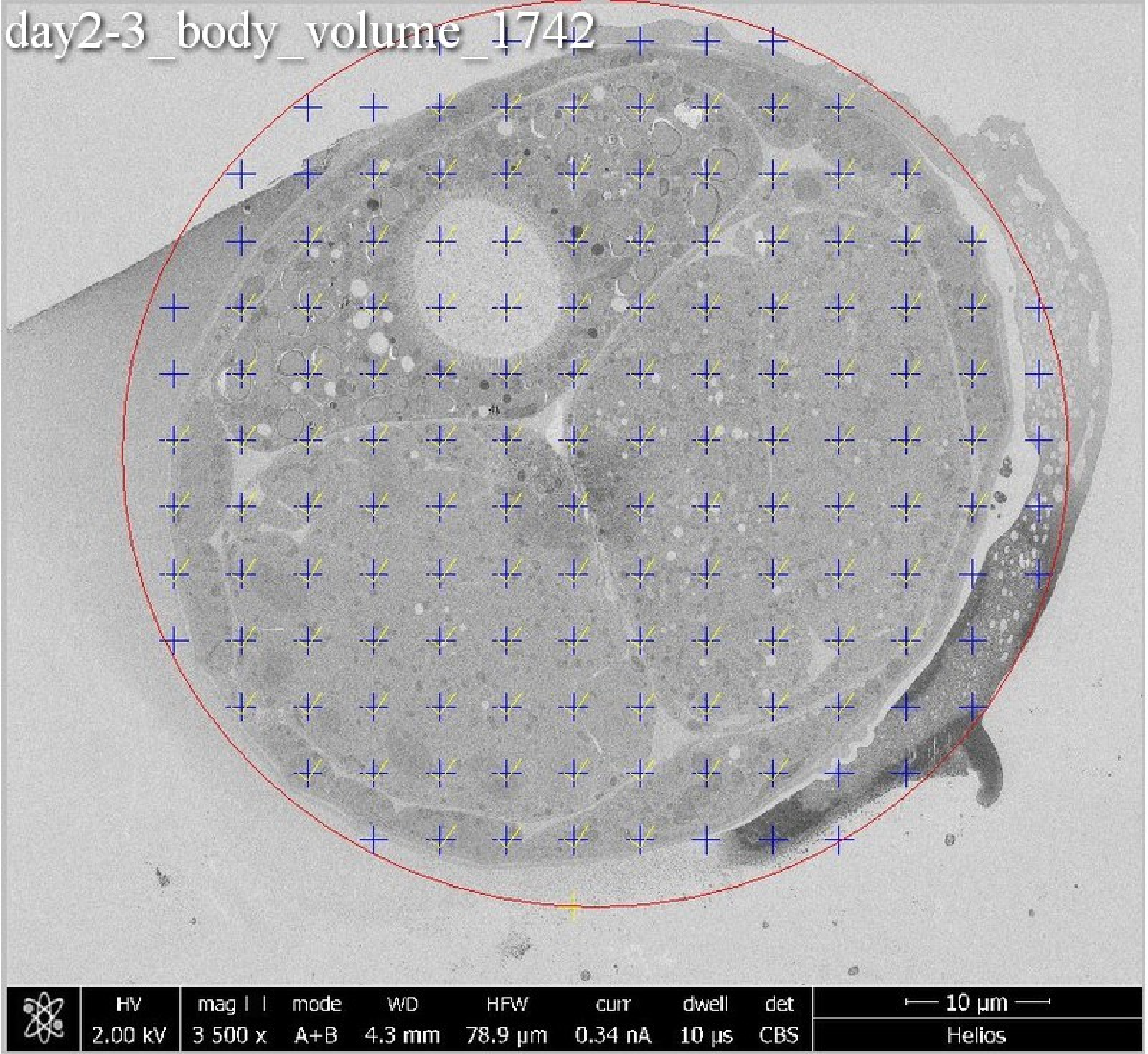
day2-3_body_volume_1742
HV
2.00 kV
mag
3 500 x
mode
A+B
WD
4.3 mm
HFW
78.9 μm
curr
0.34 nA
dwell
10 μs
det
CBS
10 μm
Helios

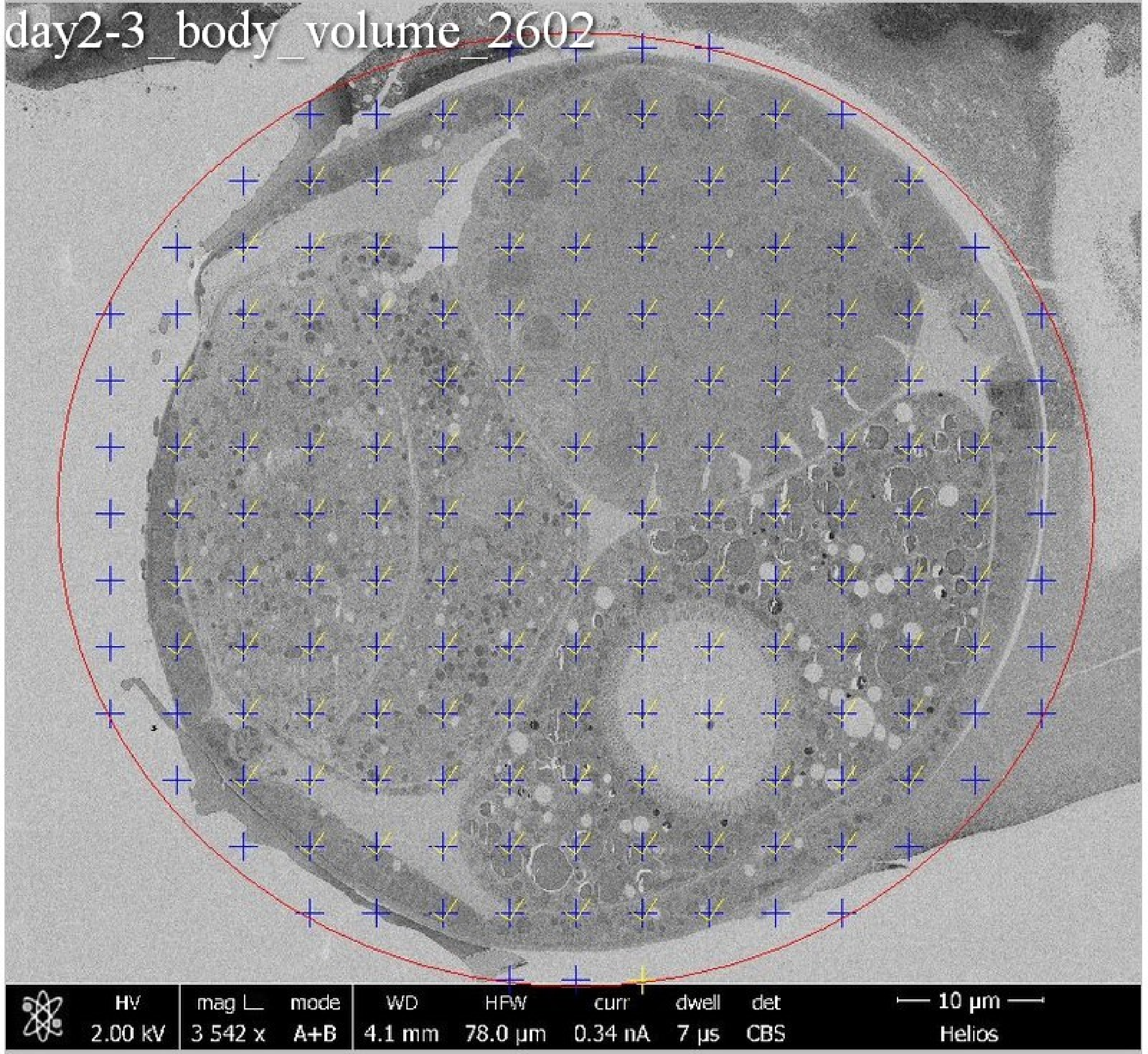

day2-3_body_volume_2602
HV 2.00 kV
mag 3 542 x
mode A+B
WD 4.1 mm
HFW 78.0 μm
curr 0.34 nA
dwell 7 μs
det CBS
10 μm
Helios

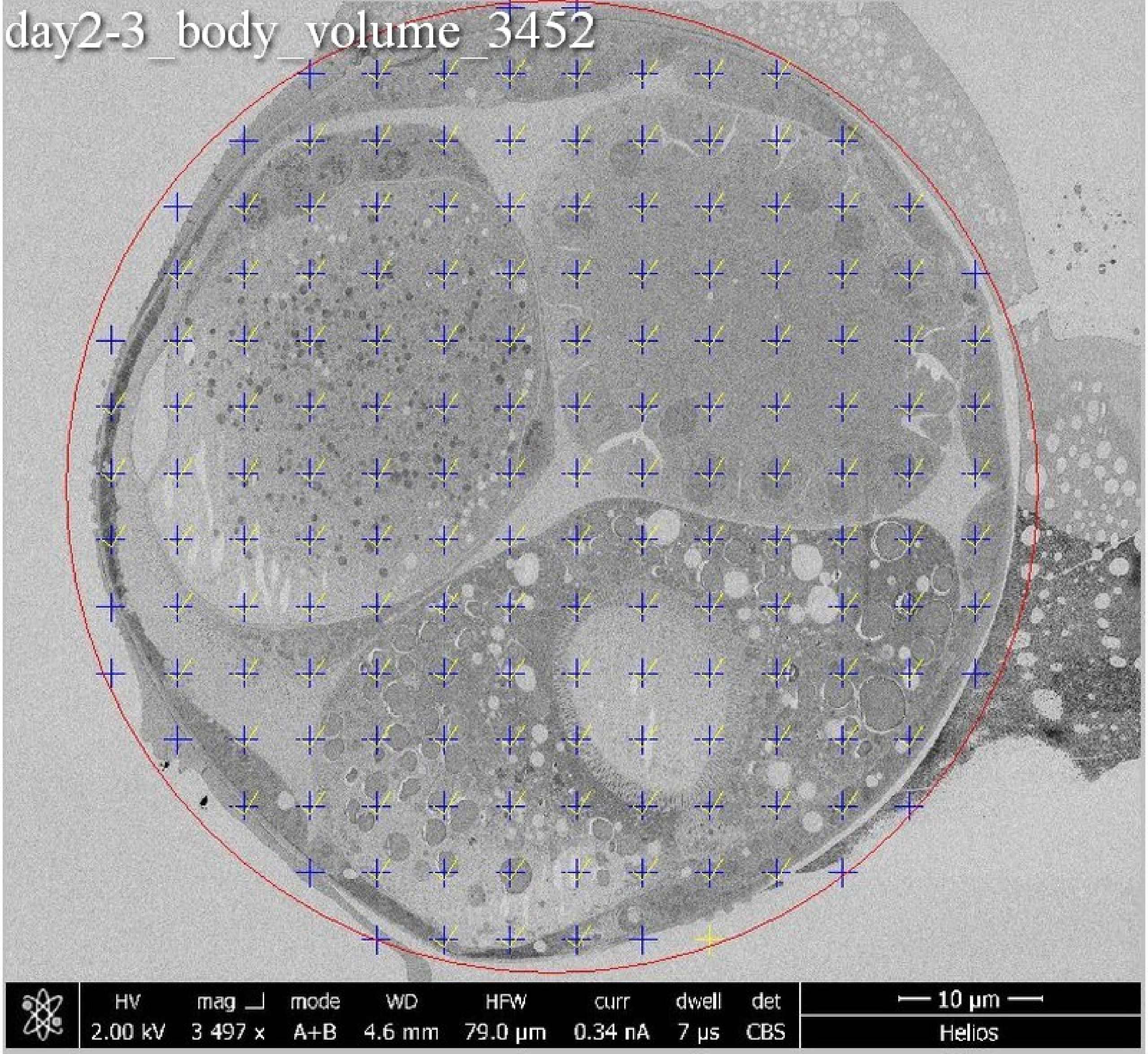
day2-3_body_volume_3452
HV
2.00 kV
mag
3 497 x
mode
A+B
WD
4.6 mm
HFW
79.0 μm
curr
0.34 nA
dwell
7 μs
det
CBS
10 μm
Helios

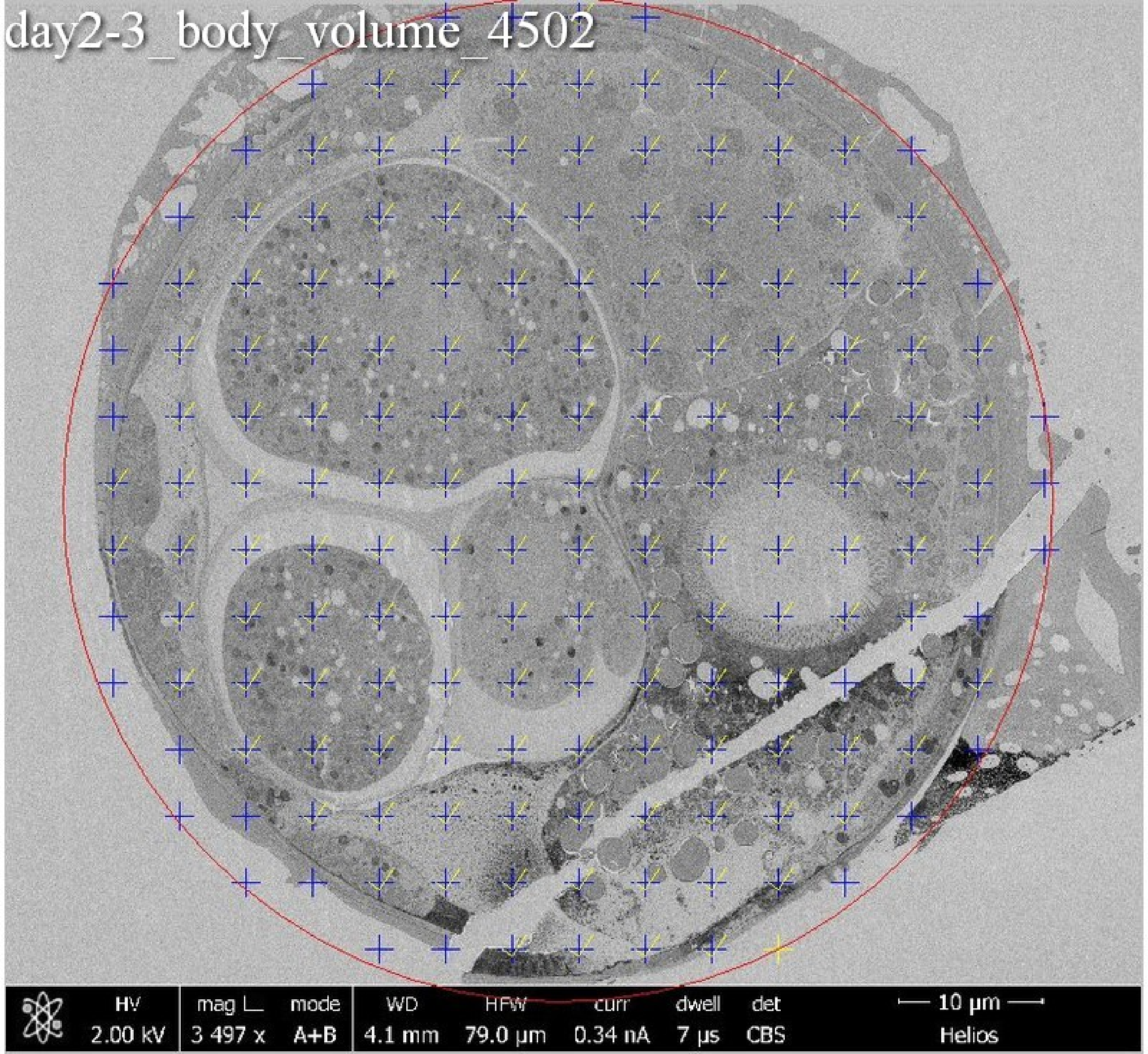
day2-3_body_volume_4502
HV
2.00 kV
mag ⊔
3 497 x
mode
A+B
WD
4.1 mm
HFW
79.0 µm
curr
0.34 nA
dwell
7 µs
det
CBS
10 µm
Helios

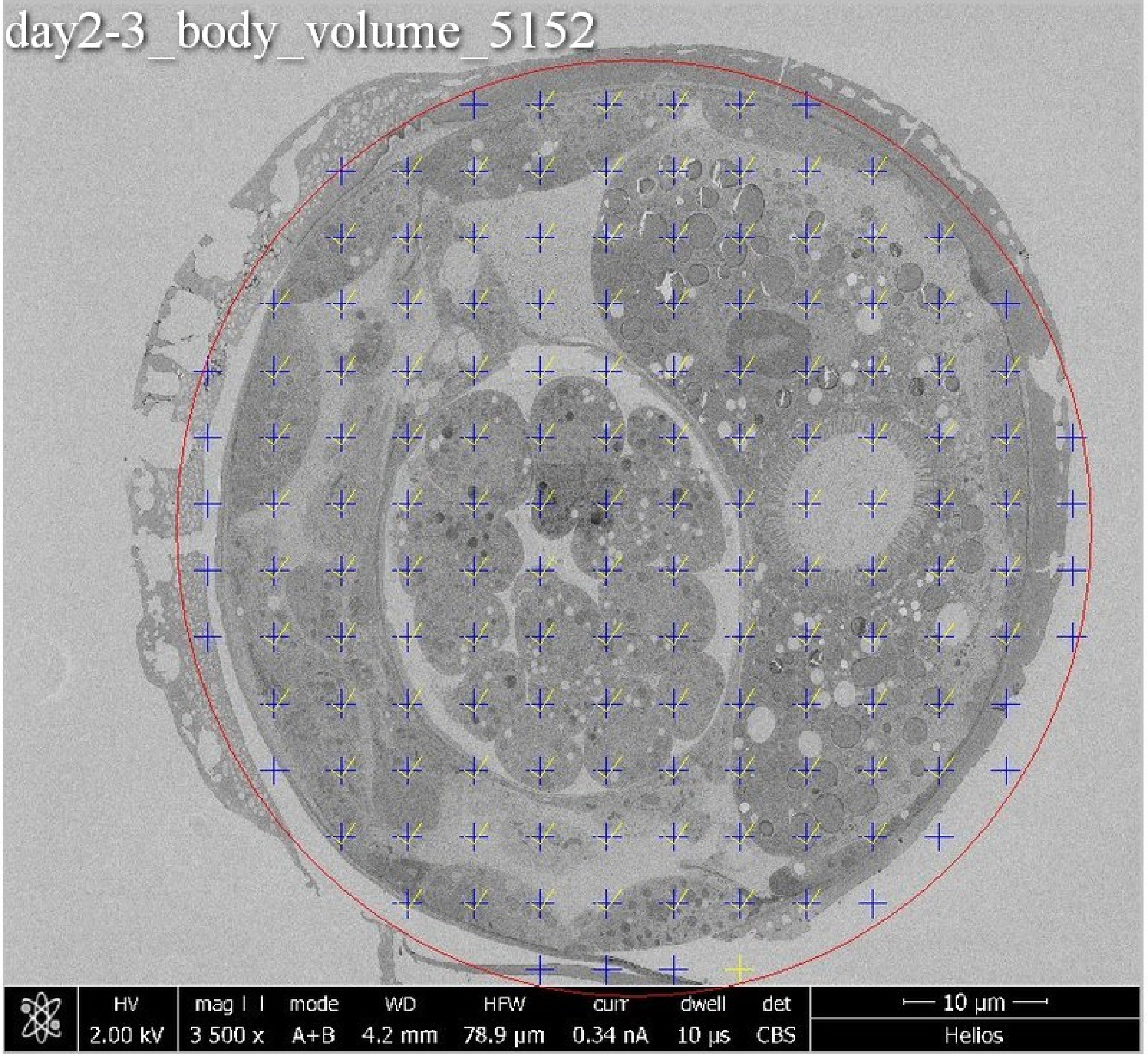

day2-3_body_volume_5152
HV 2.00 kV
mag 3 500 x
mode A+B
WD 4.2 mm
HFW 78.9 µm
curr 0.34 nA
dwell 10 µs
det CBS
10 µm
Helios

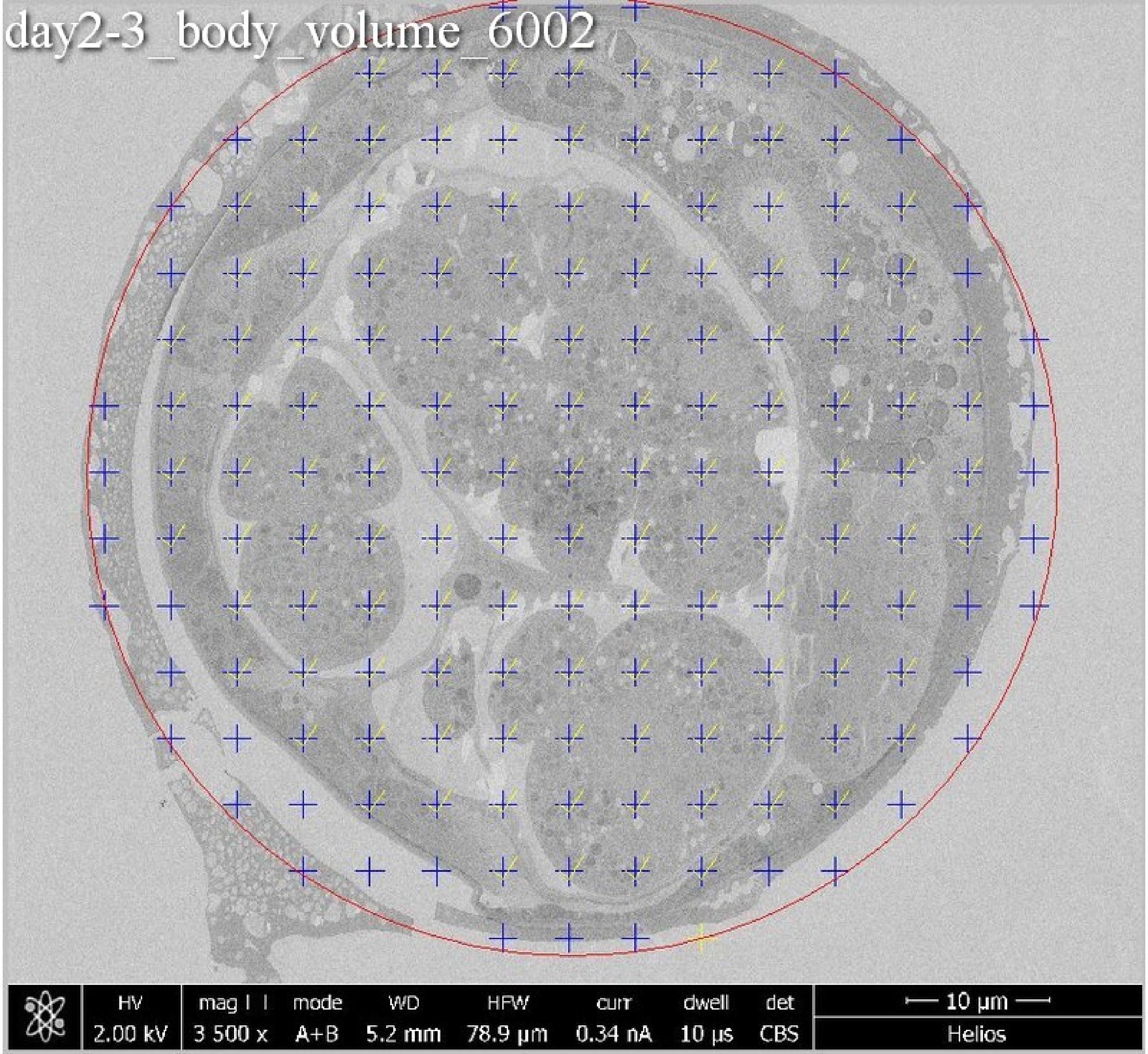
day2-3_body_volume_6002
HV 2.00 kV
mag ⊞ 3 500 x
mode A+B
WD 5.2 mm
HFW 78.9 µm
curr 0.34 nA
dwell 10 µs
det CBS
10 µm
Helios

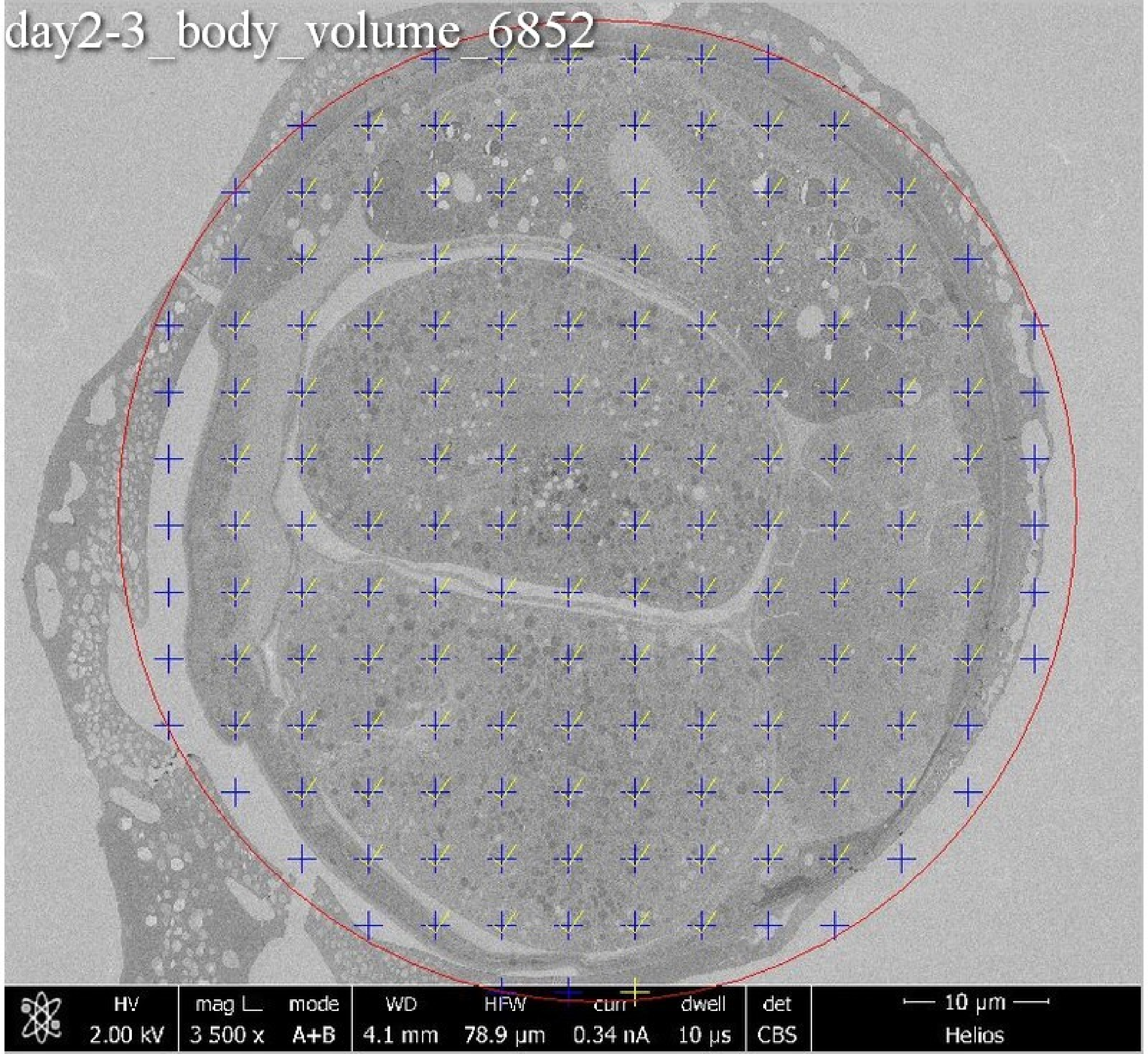
day2-3_body_volume_6852
HV
2.00 kV
mag ⊔
3 500 x
mode
A+B
WD
4.1 mm
HFW
78.9 µm
curr
0.34 nA
dwell
10 µs
det
CBS
10 µm
Helios

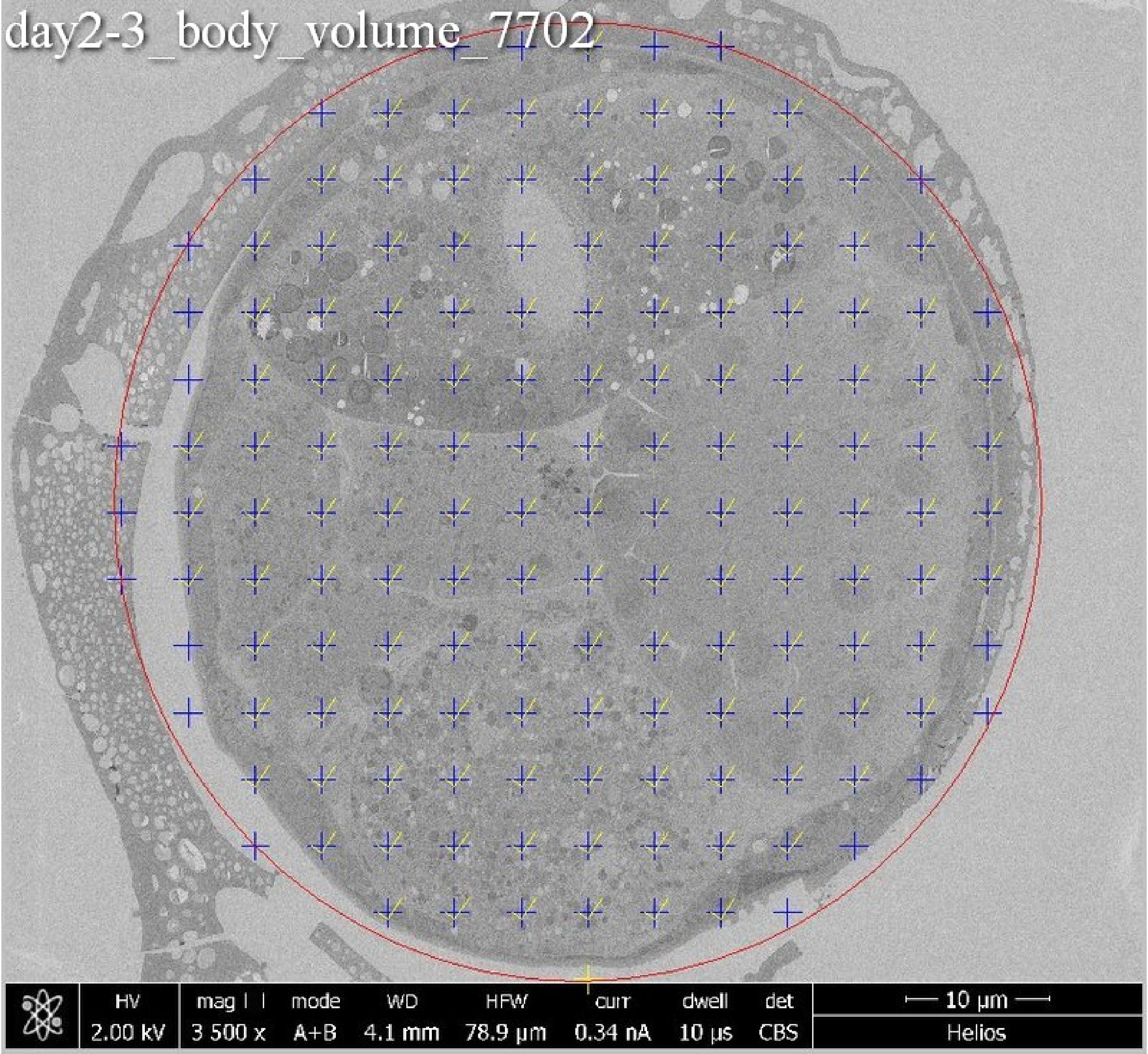

day2-3_body_volume_7702
HV
2.00 kV
mag
3 500 x
mode
A+B
WD
4.1 mm
HFW
78.9 µm
curr
0.34 nA
dwell
10 µs
det
CBS
10 µm
Helios

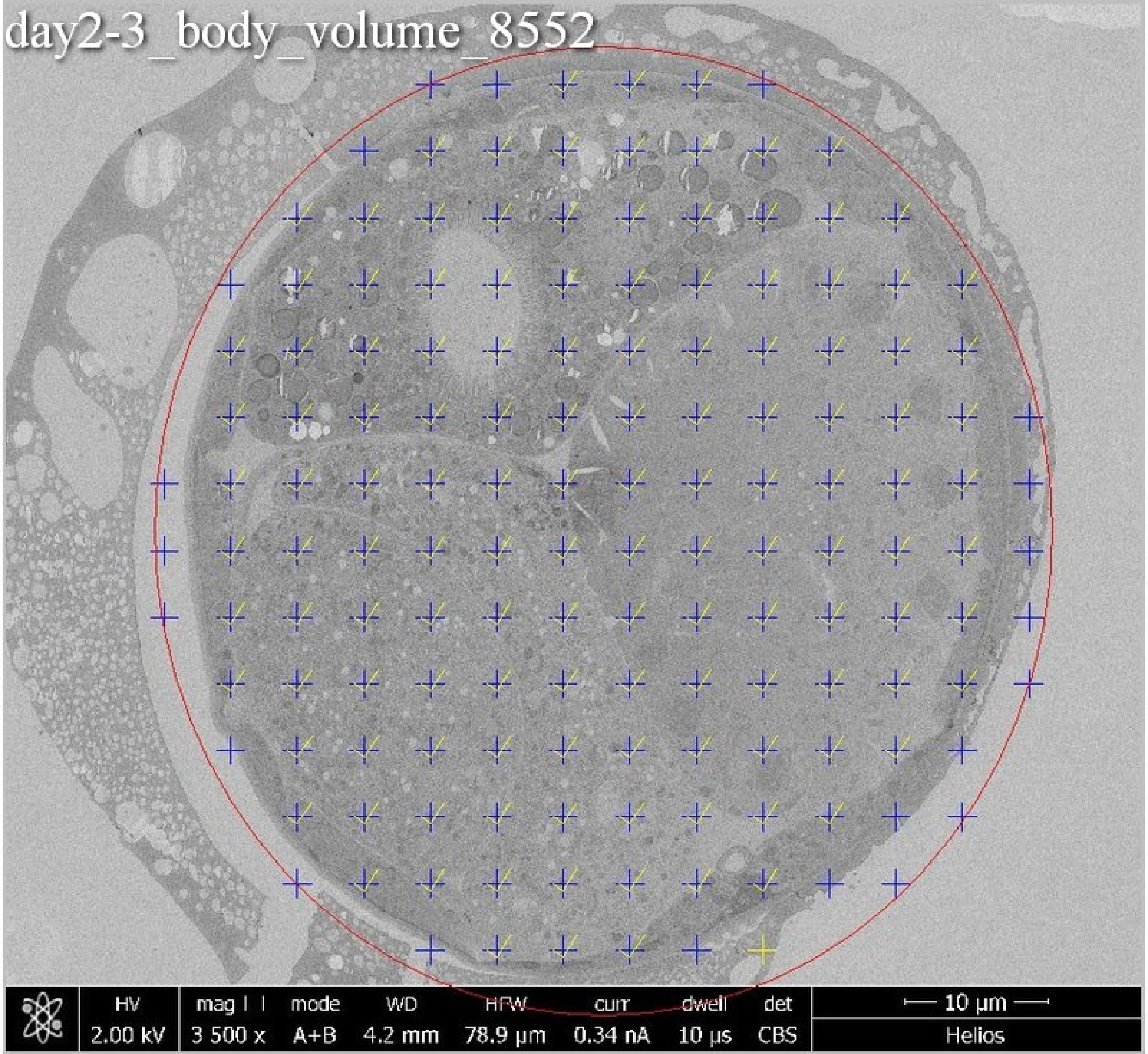
day2-3_body_volume_8552
HV 2.00 kV
mag 3 500 x
mode A+B
WD 4.2 mm
HFW 78.9 µm
curr 0.34 nA
dwell 10 µs
det CBS
10 µm
Helios

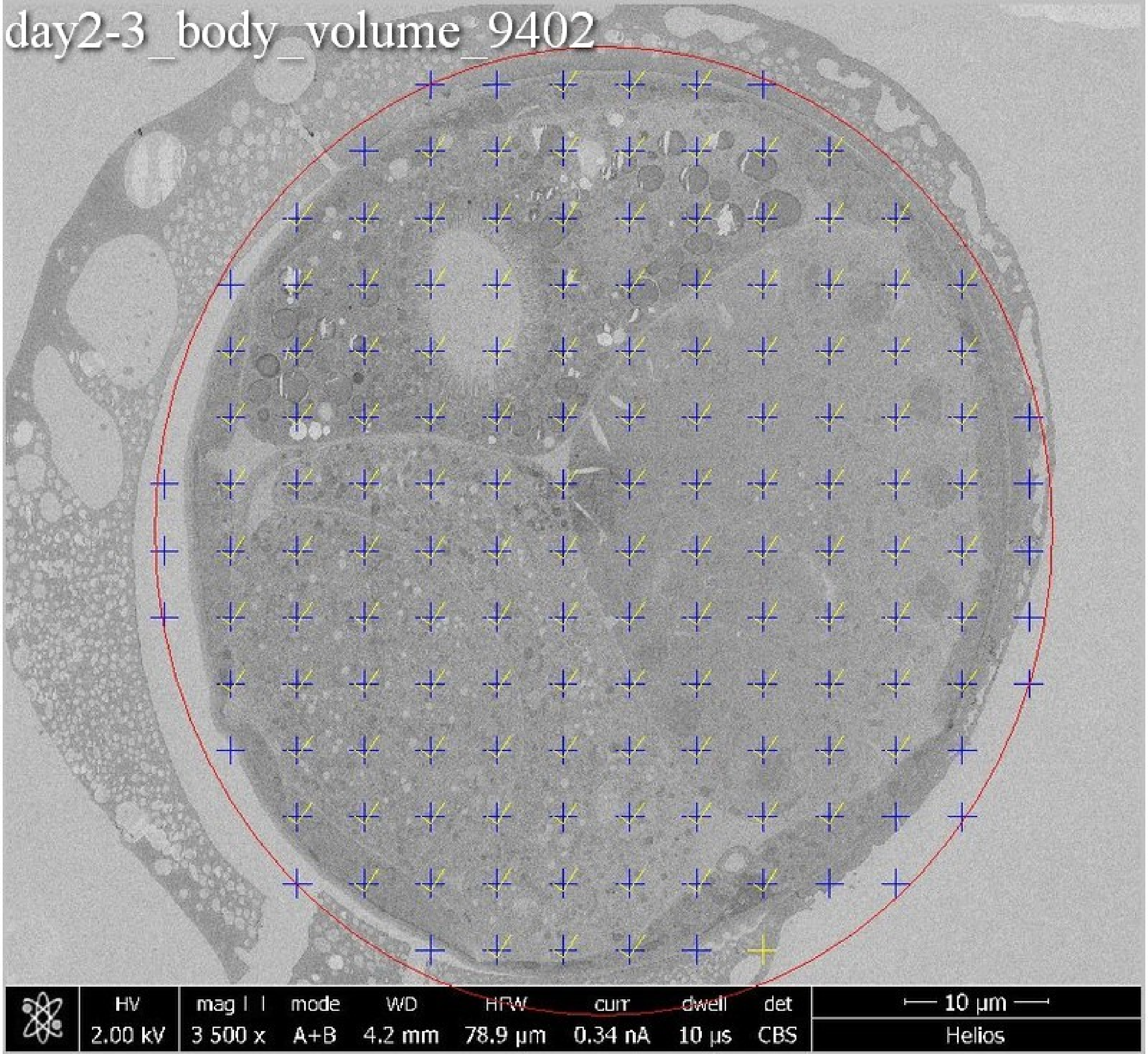
day2-3_body_volume_9402
HV
2.00 kV
mag
3 500 x
mode
A+B
WD
4.2 mm
HFW
78.9 µm
curr
0.34 nA
dwell
10 µs
det
CBS
10 µm
Helios

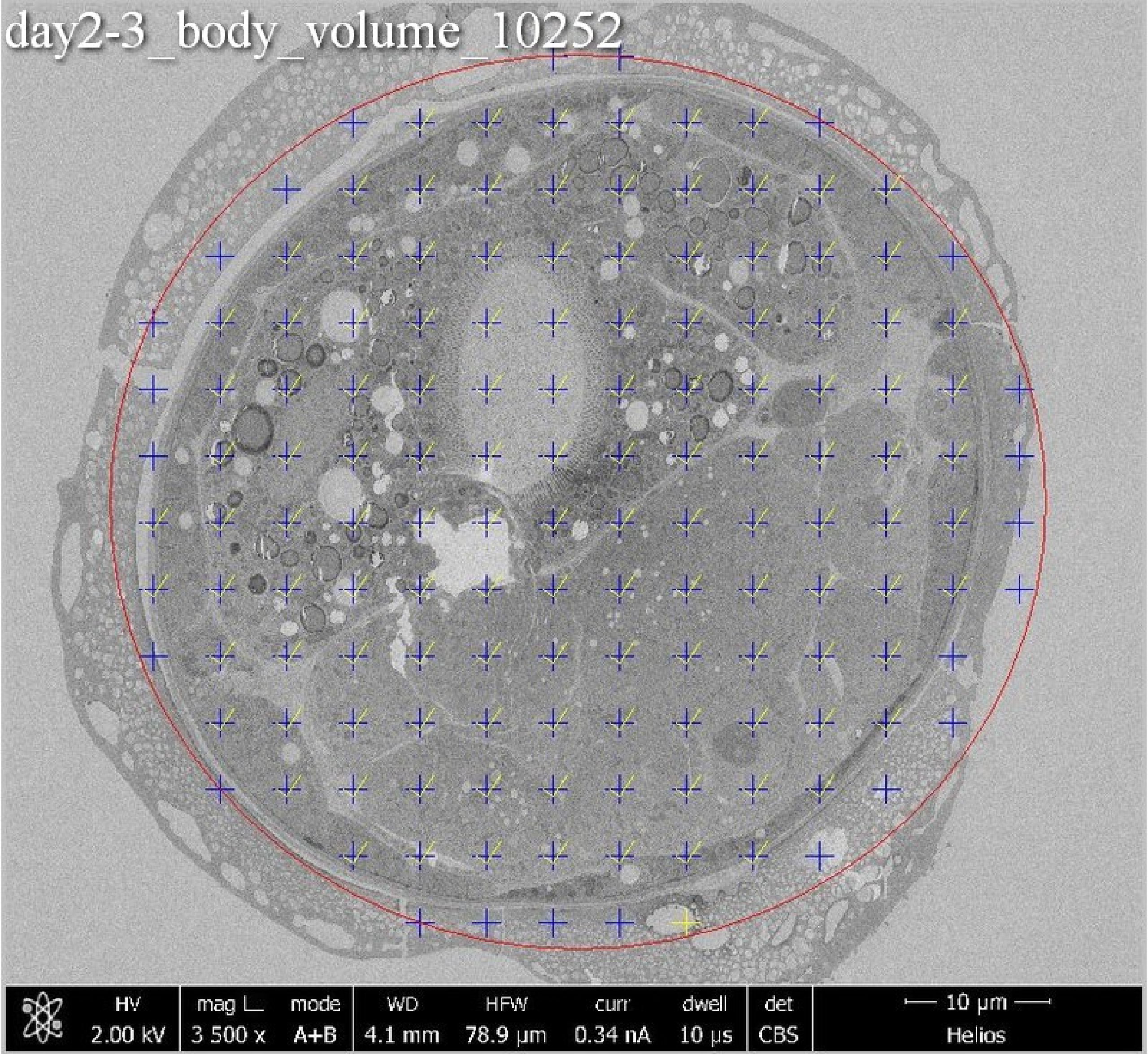
day2-3_body_volume_10252
HV
2.00 kV
mag ⌙
3 500 x
mode
A+B
WD
4.1 mm
HFW
78.9 µm
curr
0.34 nA
dwell
10 µs
det
CBS
10 µm
Helios

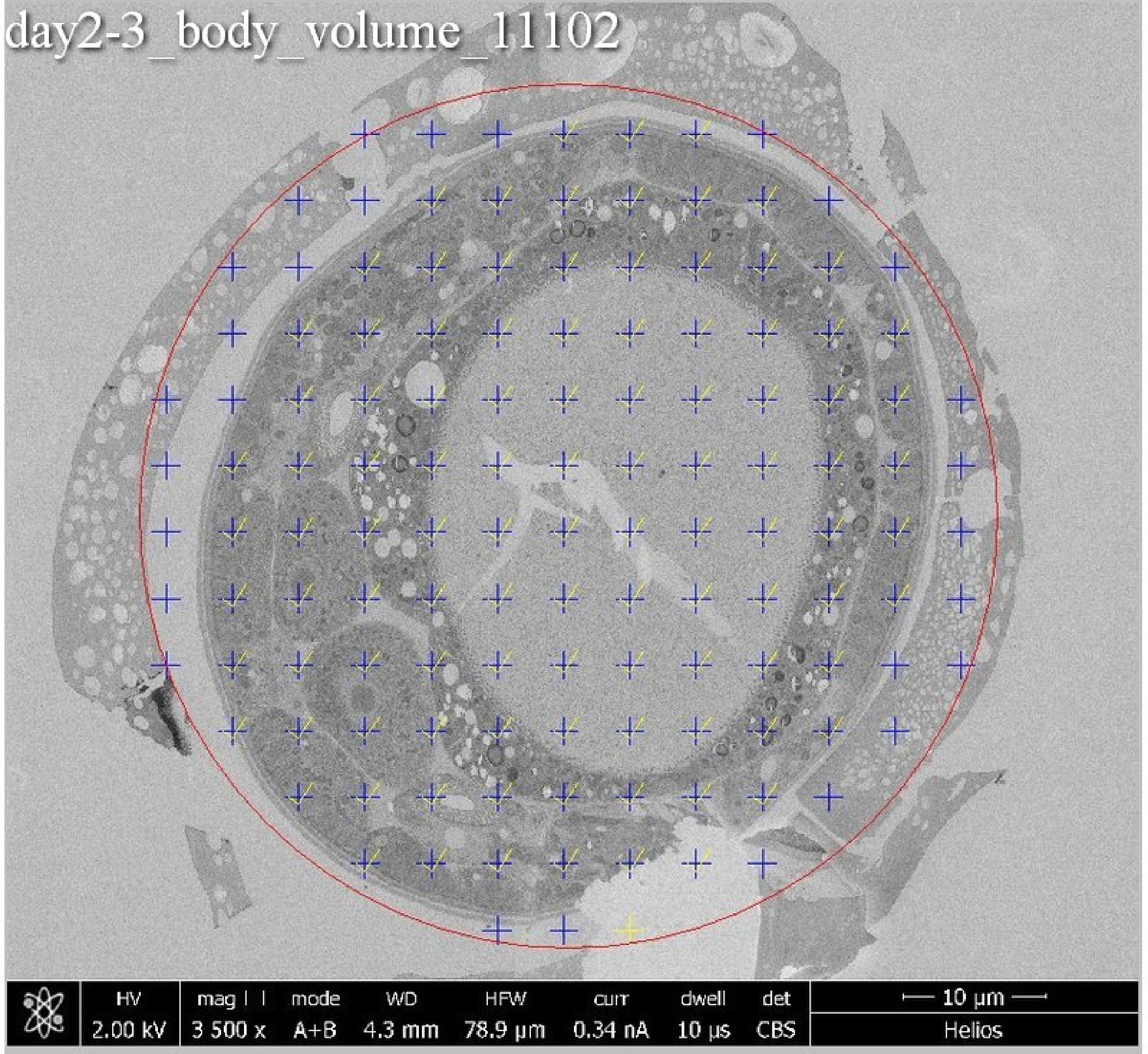
day2-3_body_volume_11102
HV
2.00 kV
mag
3 500 x
mode
A+B
WD
4.3 mm
HFW
78.9 µm
curr
0.34 nA
dwell
10 µs
det
CBS
10 µm
Helios

day2-3_body_volume_11952

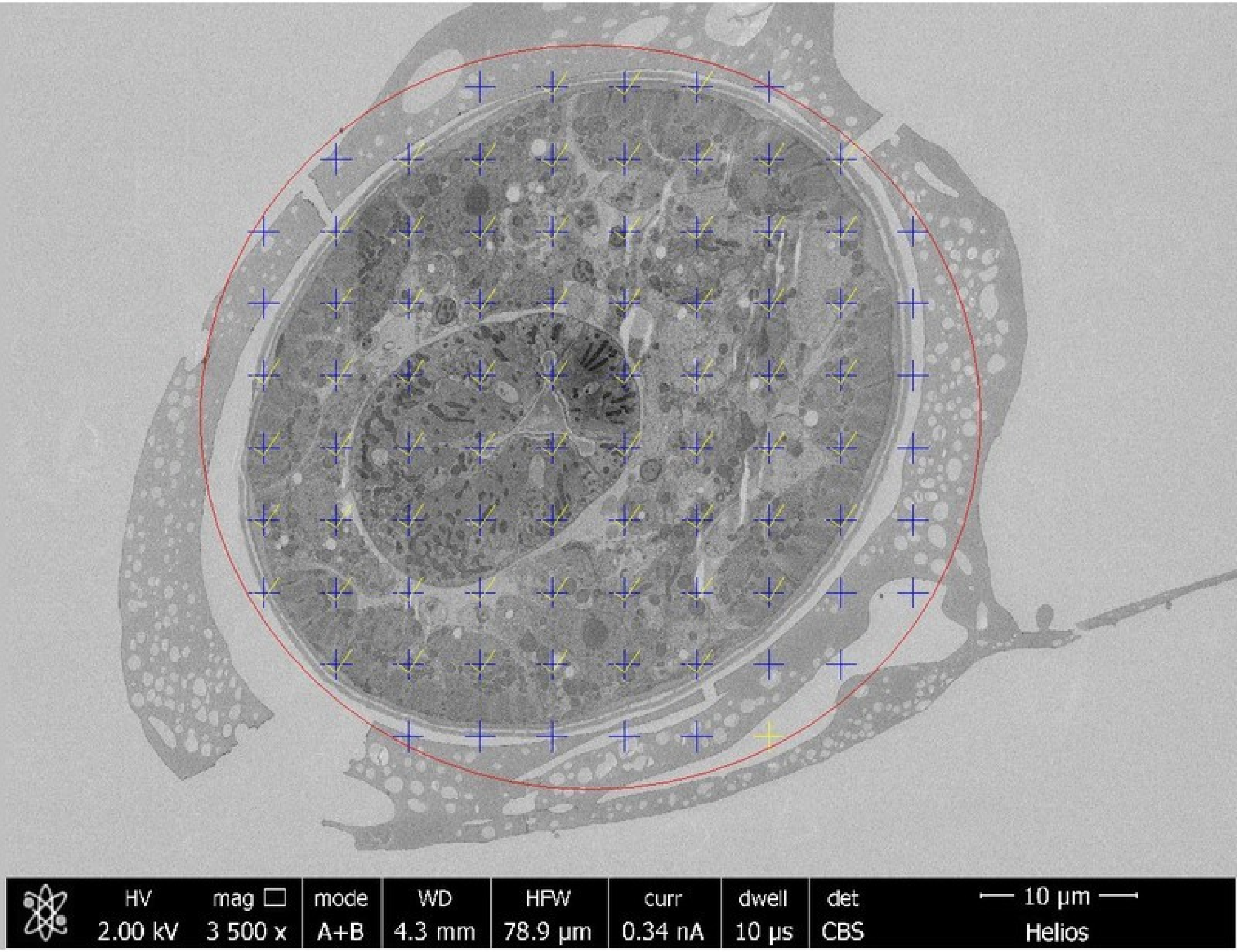

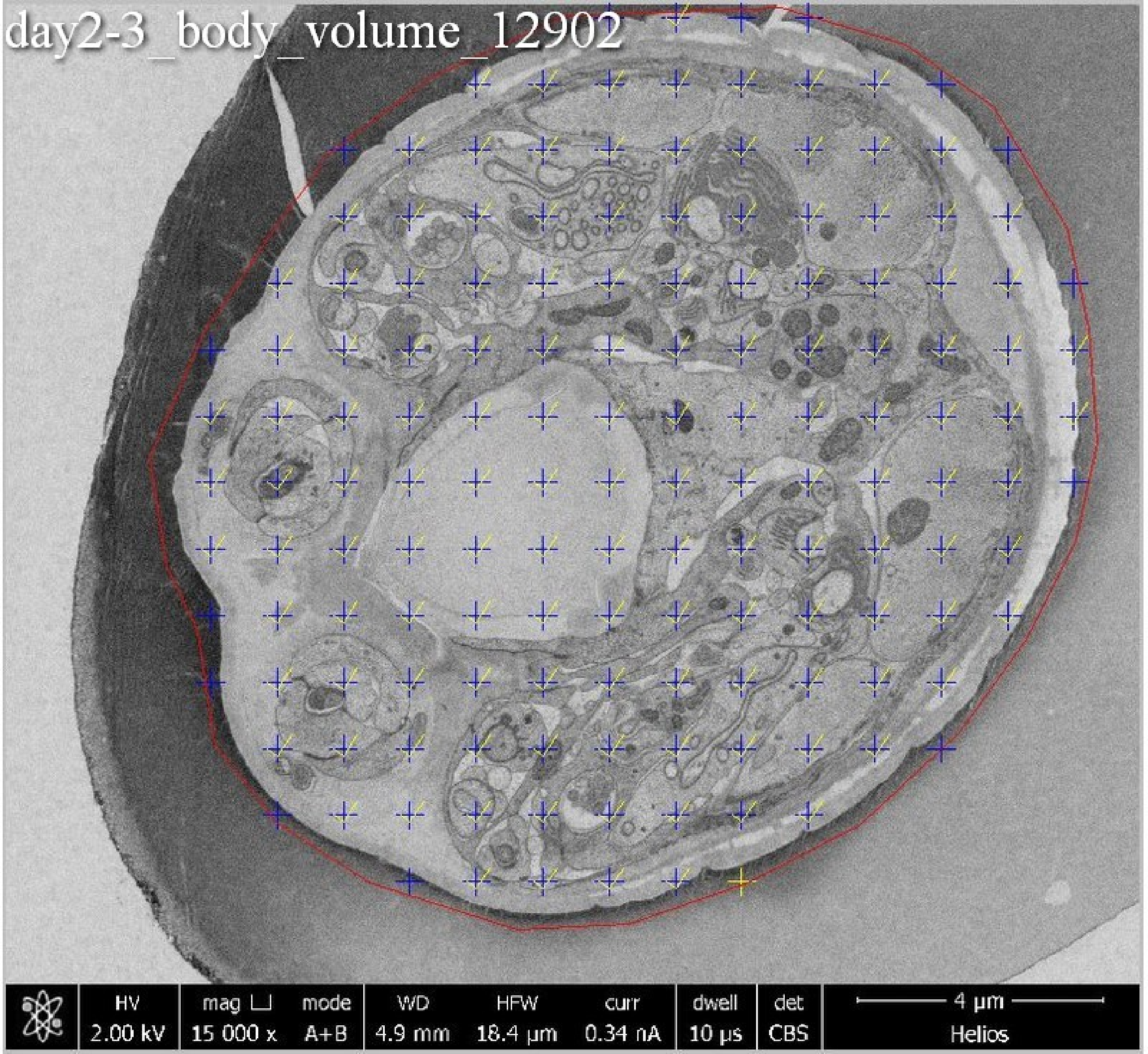
day2-3_body_volume_12902
HV
2.00 kV
mag
15 000 x
mode
A+B
WD
4.9 mm
HFW
18.4 µm
curr
0.34 nA
dwell
10 µs
det
CBS
4 µm
Helios

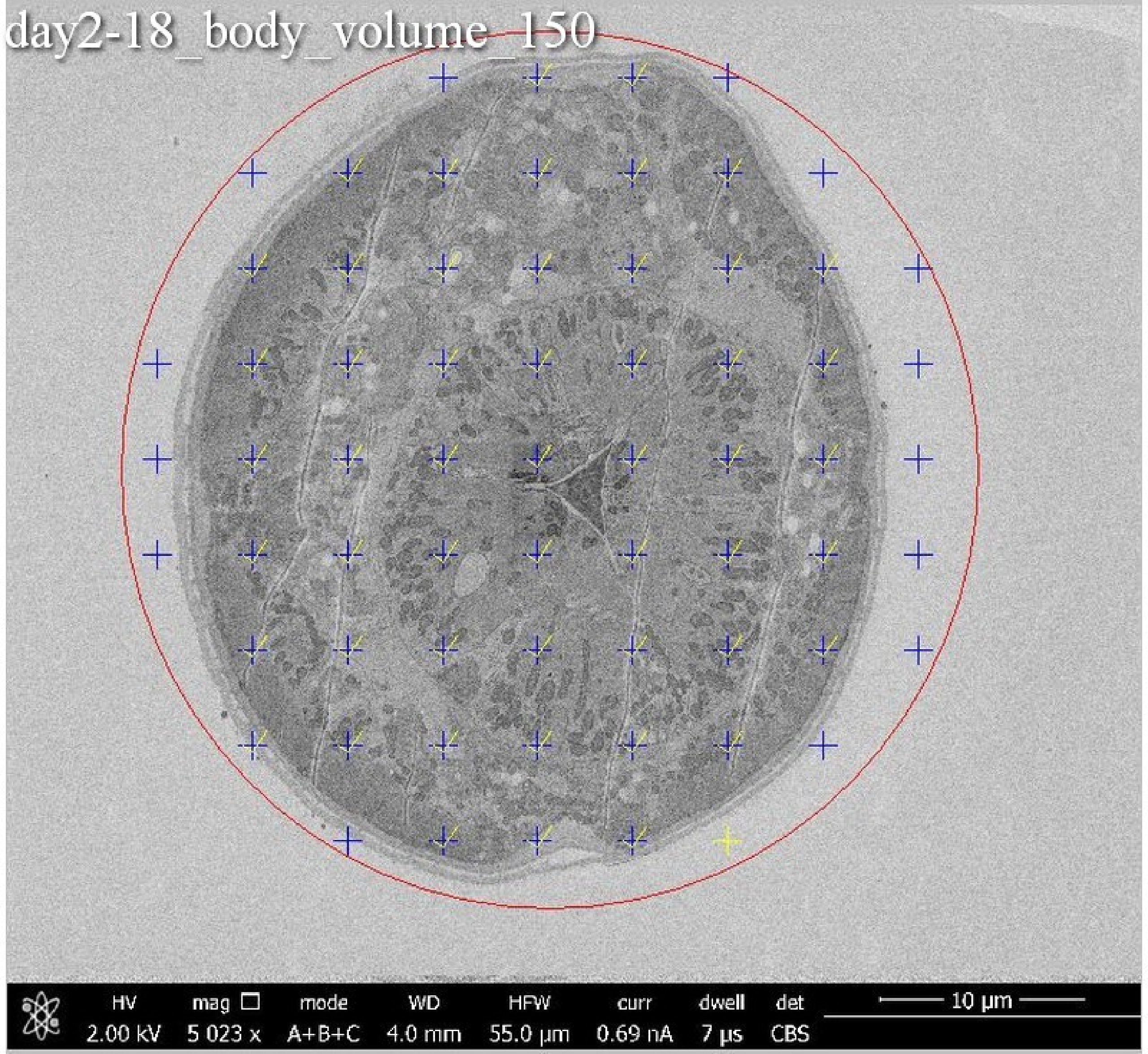
day2-18_body_volume_150
HV
2.00 kV
mag
5 023 x
mode
A+B+C
WD
4.0 mm
HFW
55.0 µm
curr
0.69 nA
dwell
7 µs
det
CBS
10 µm

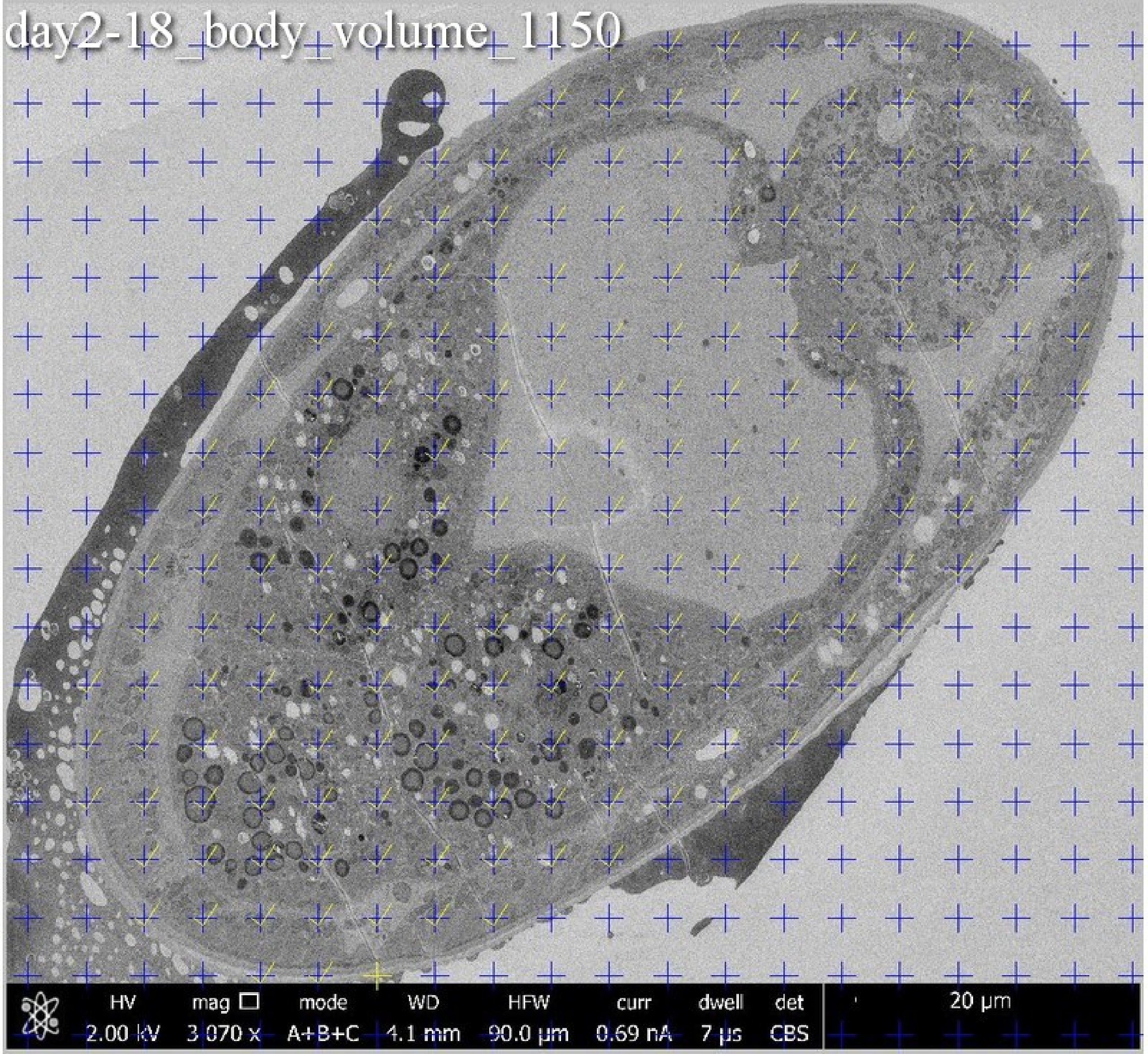
day2-18_body_volume_1150
HV 2.00 kV
mag □ 3 070 x
mode A+B+C
WD 4.1 mm
HFW 90.0 µm
curr 0.69 nA
dwell 7 µs
det CBS
20 µm

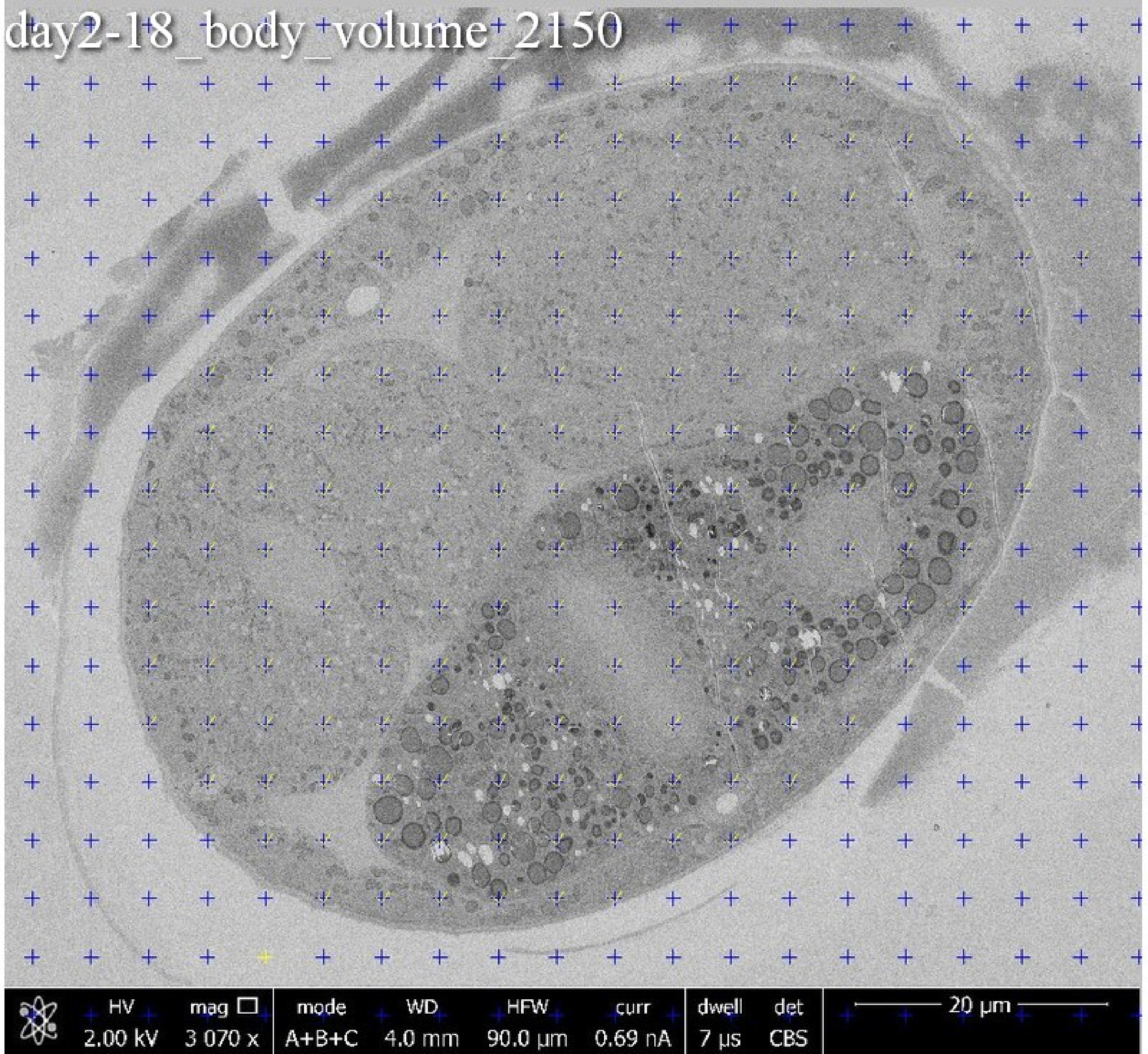
day2-18_body_volume_2150
HV 2.00 kV
mag 3 070 x
mode A+B+C
WD 4.0 mm
HFW 90.0 µm
curr 0.69 nA
dwell 7 µs
det CBS
20 µm

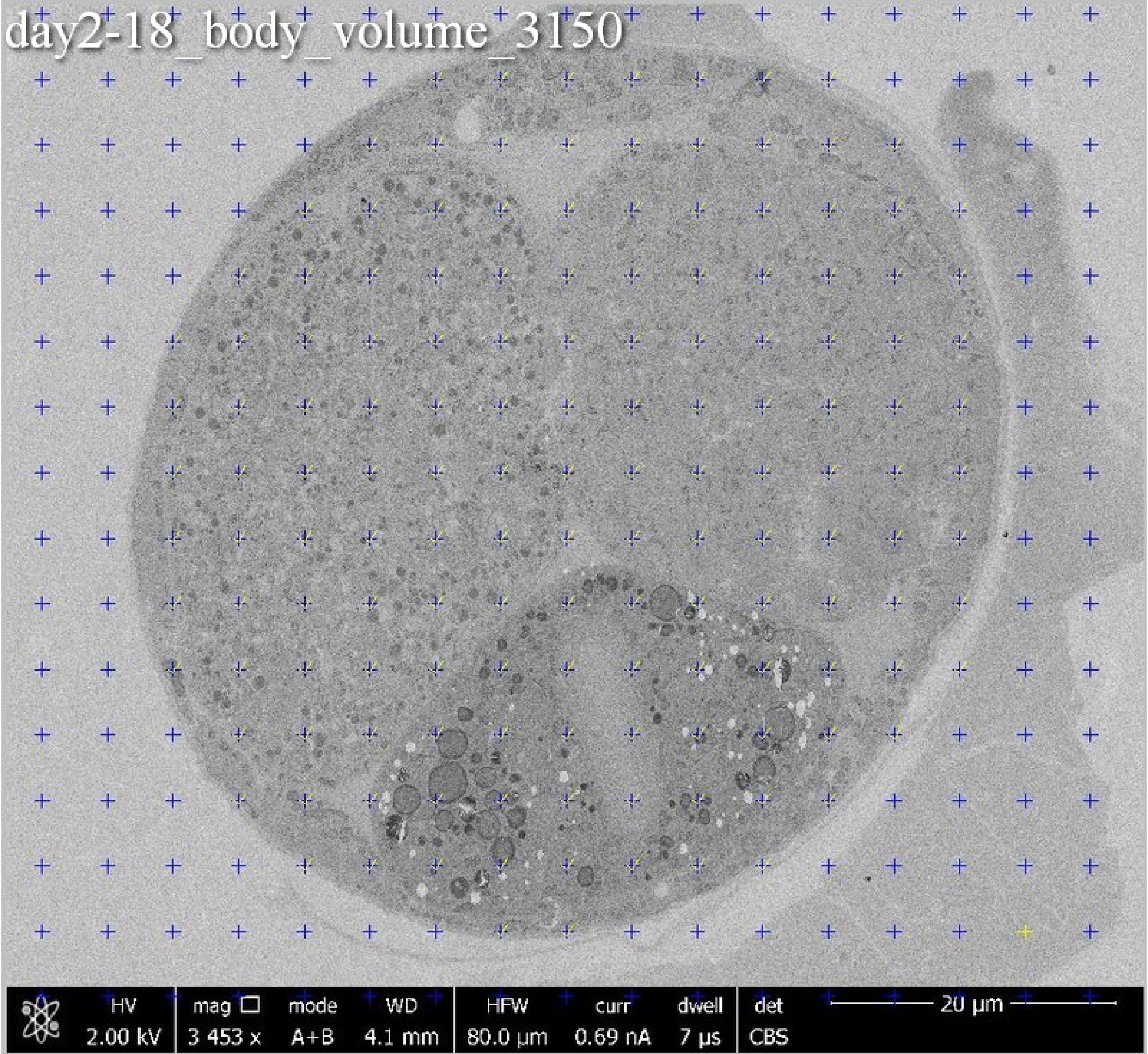
day2-18_body_volume_3150
HV 2.00 kV
mag 3 453 x
mode A+B
WD 4.1 mm
HFW 80.0 µm
curr 0.69 nA
dwell 7 µs
det CBS
20 µm

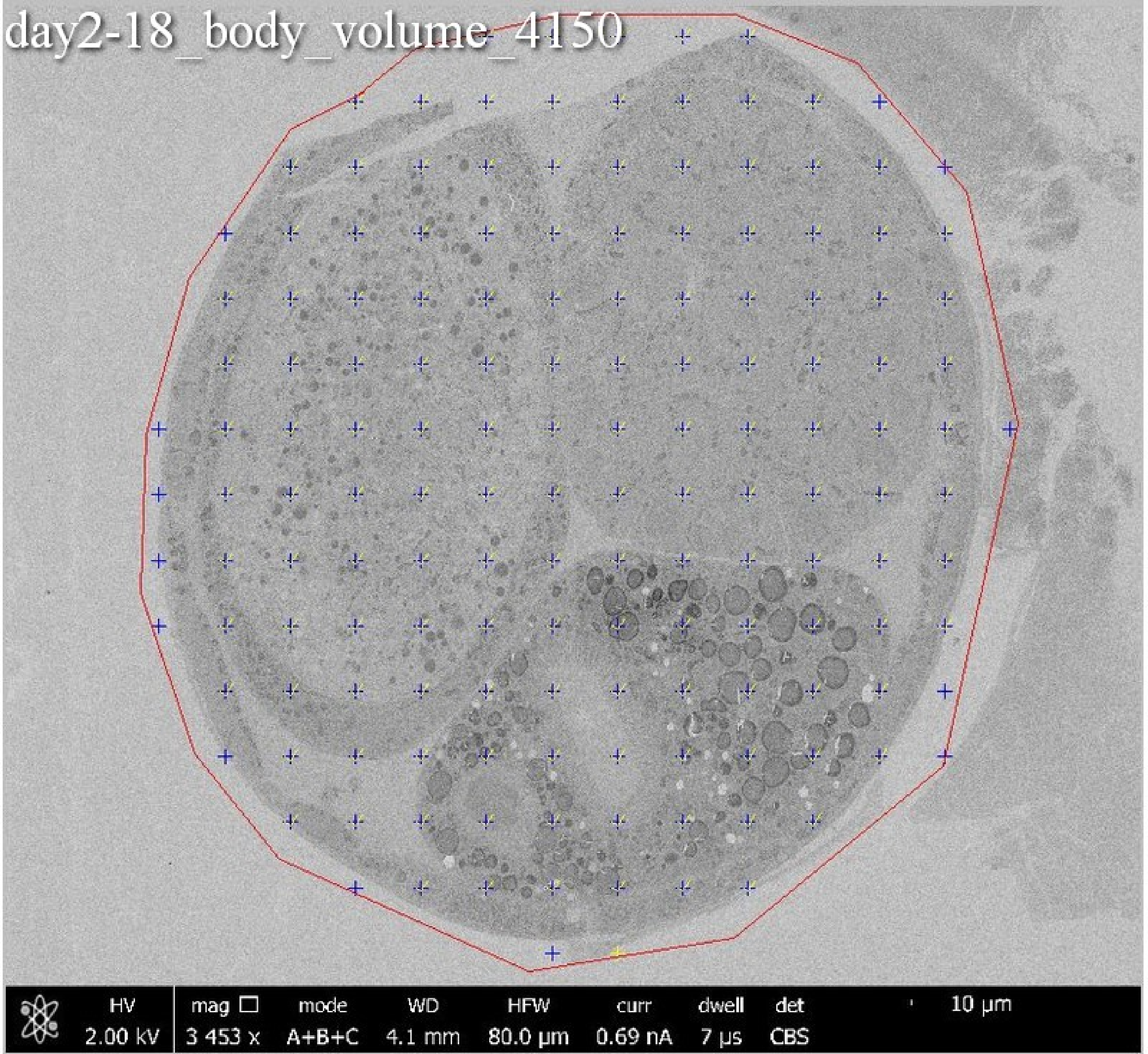
day2-18_body_volume_4150
HV 2.00 kV
mag 3 453 x
mode A+B+C
WD 4.1 mm
HFW 80.0 µm
curr 0.69 nA
dwell 7 µs
det CBS
10 µm

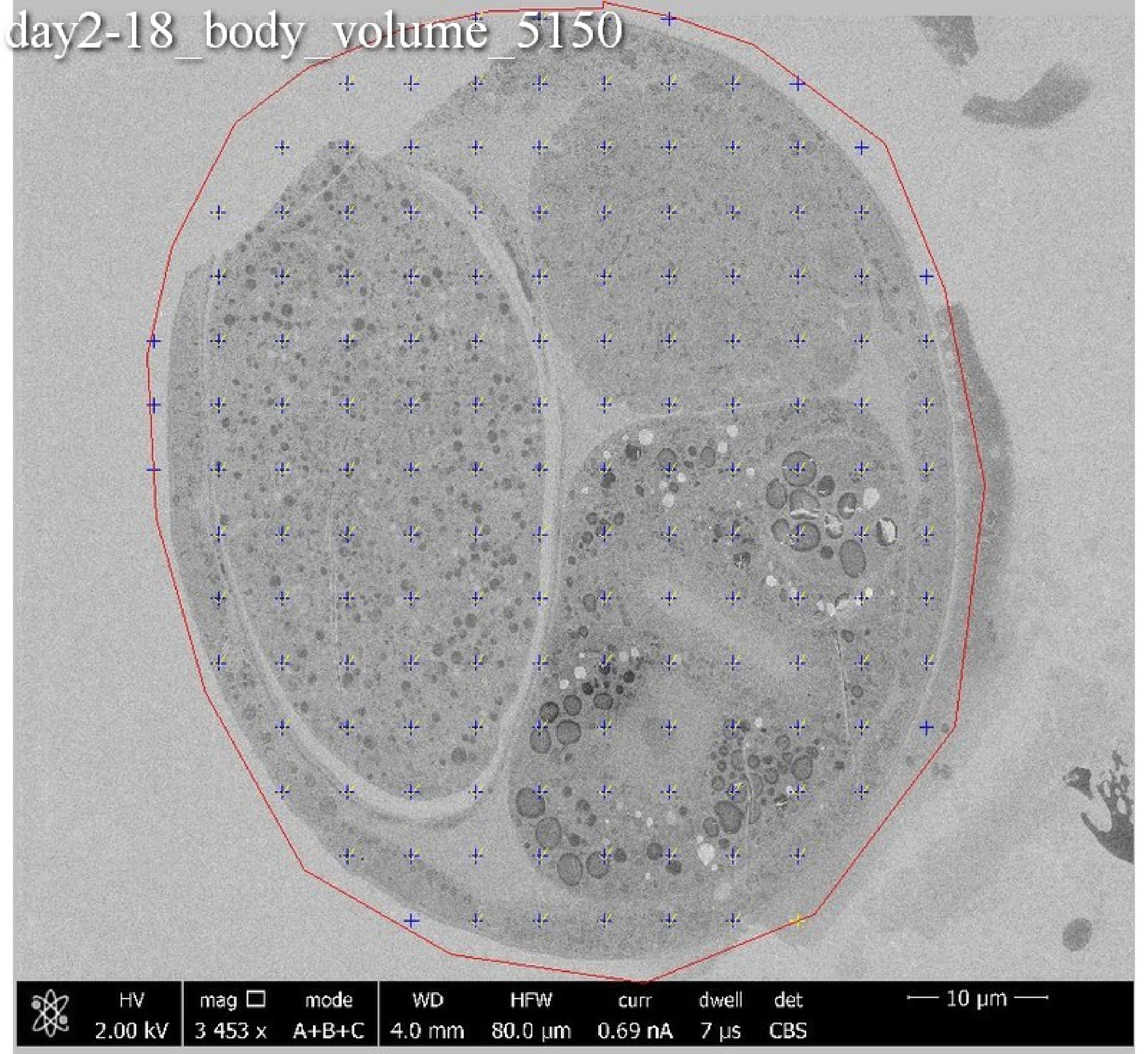
day2-18_body_volume_5150
HV
2.00 kV
mag ☐
3 453 x
mode
A+B+C
WD
4.0 mm
HFW
80.0 µm
curr
0.69 nA
dwell
7 µs
det
CBS
10 µm

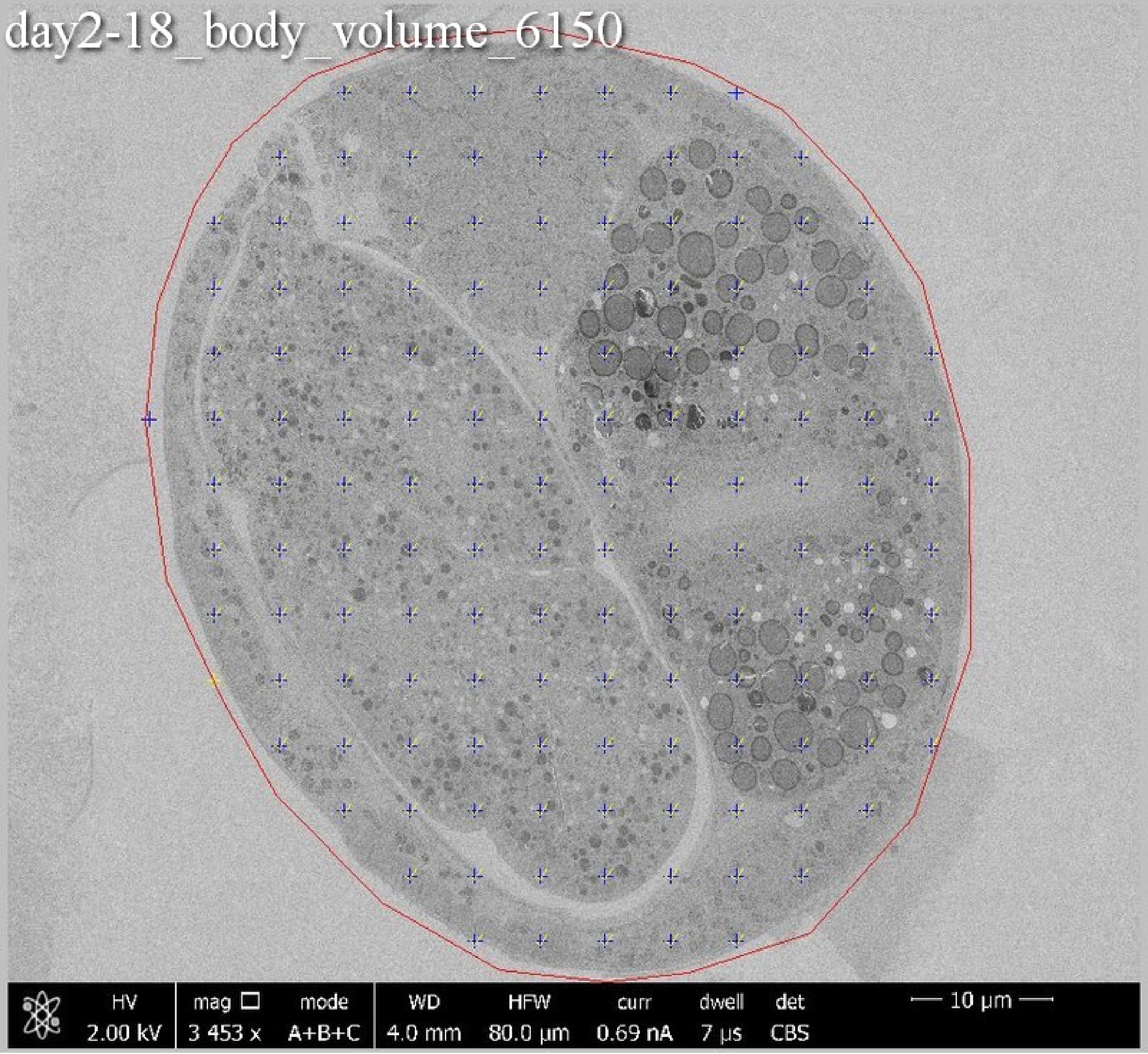
day2-18_body_volume_6150
HV 2.00 kV
mag □ 3 453 x
mode A+B+C
WD 4.0 mm
HFW 80.0 µm
curr 0.69 nA
dwell 7 µs
det CBS
10 µm

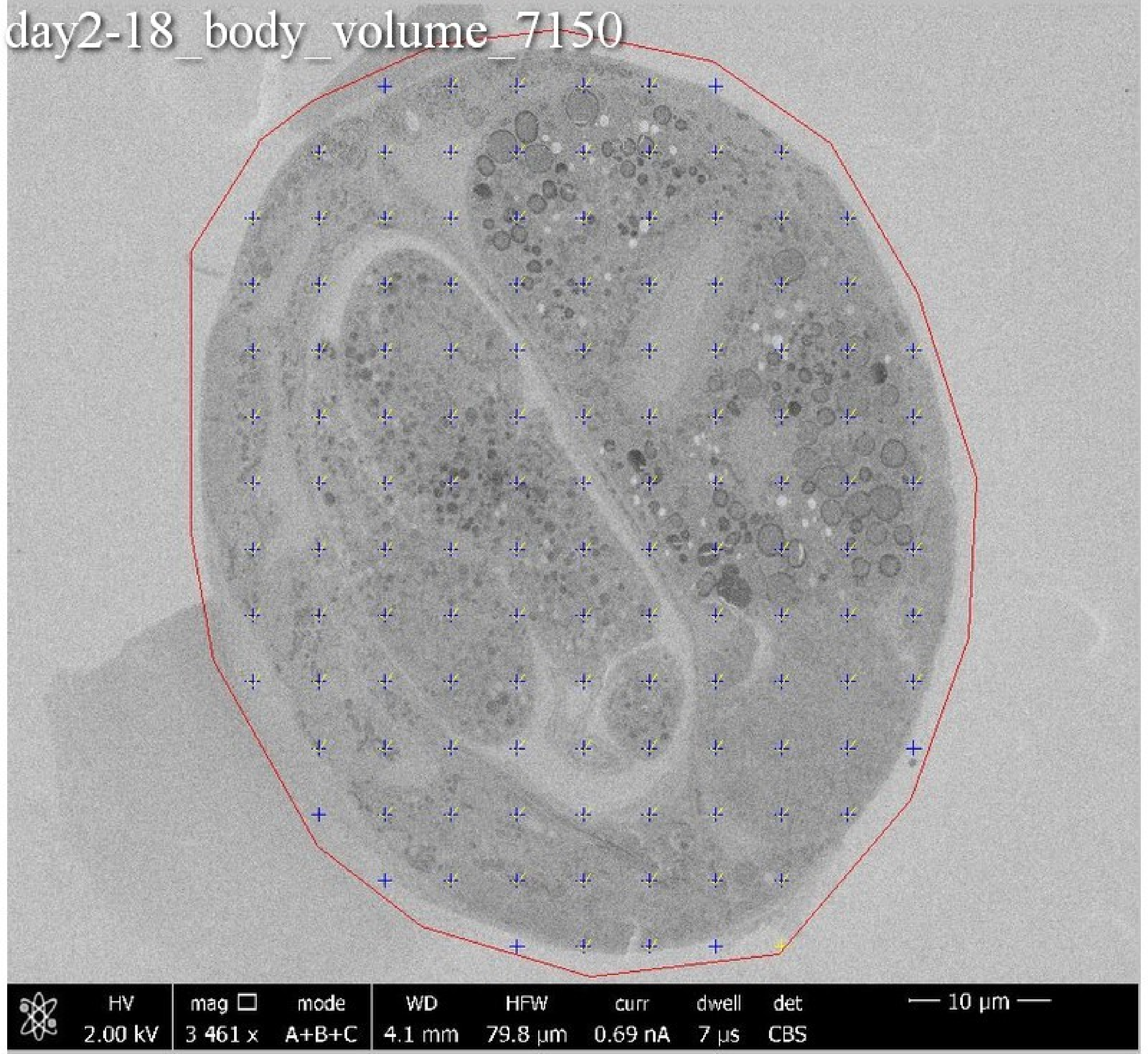
day2-18_body_volume_7150
HV
2.00 kV
mag
3 461 x
mode
A+B+C
WD
4.1 mm
HFW
79.8 μm
curr
0.69 nA
dwell
7 μs
det
CBS
10 μm

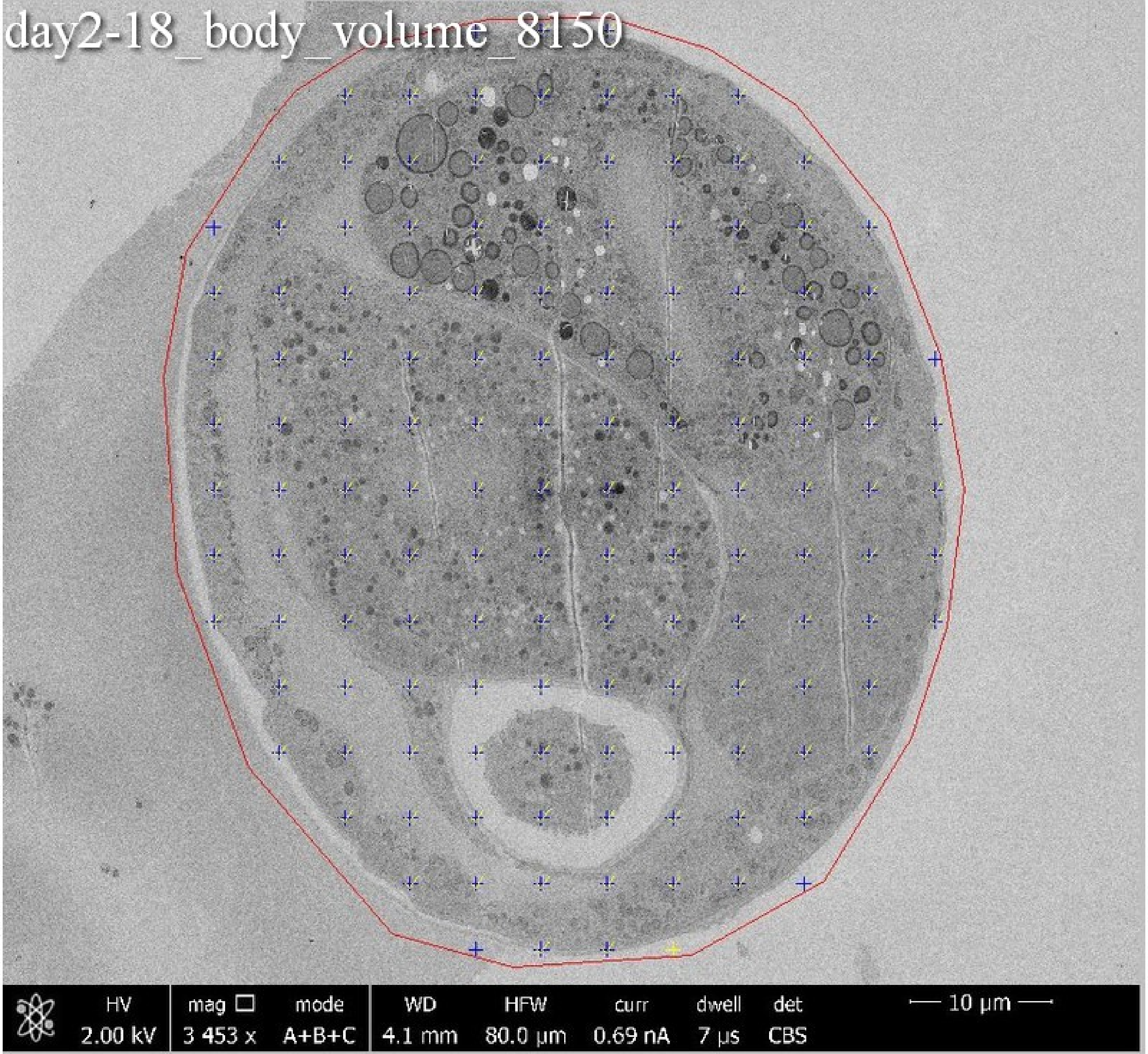
day2-18_body_volume_8150
HV
2.00 kV
mag ☐
3 453 x
mode
A+B+C
WD
4.1 mm
HFW
80.0 µm
curr
0.69 nA
dwell
7 µs
det
CBS
10 µm

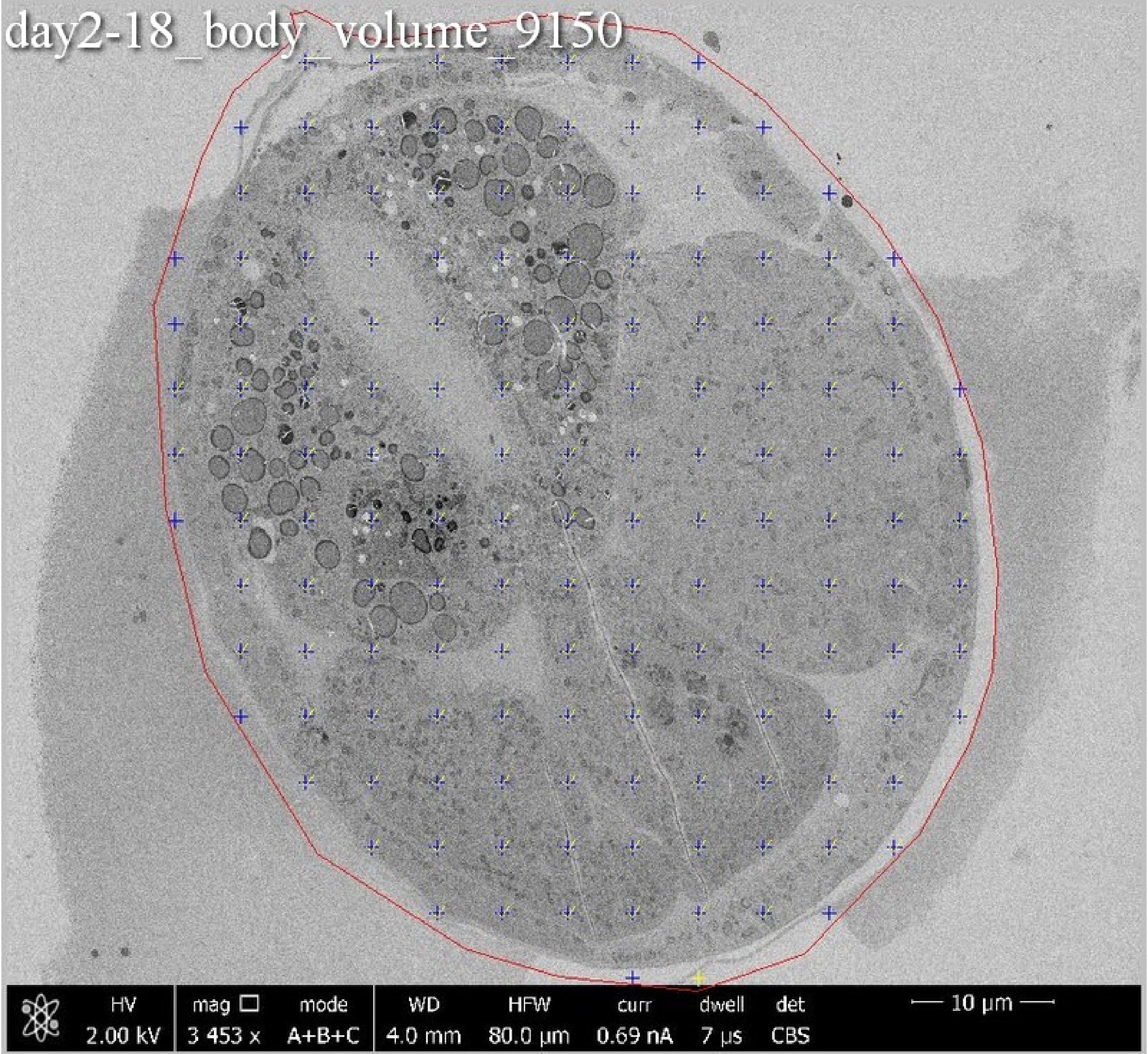
day2-18_body_volume_9150
HV 2.00 kV
mag 3 453 x
mode A+B+C
WD 4.0 mm
HFW 80.0 μm
curr 0.69 nA
dwell 7 μs
det CBS
10 μm

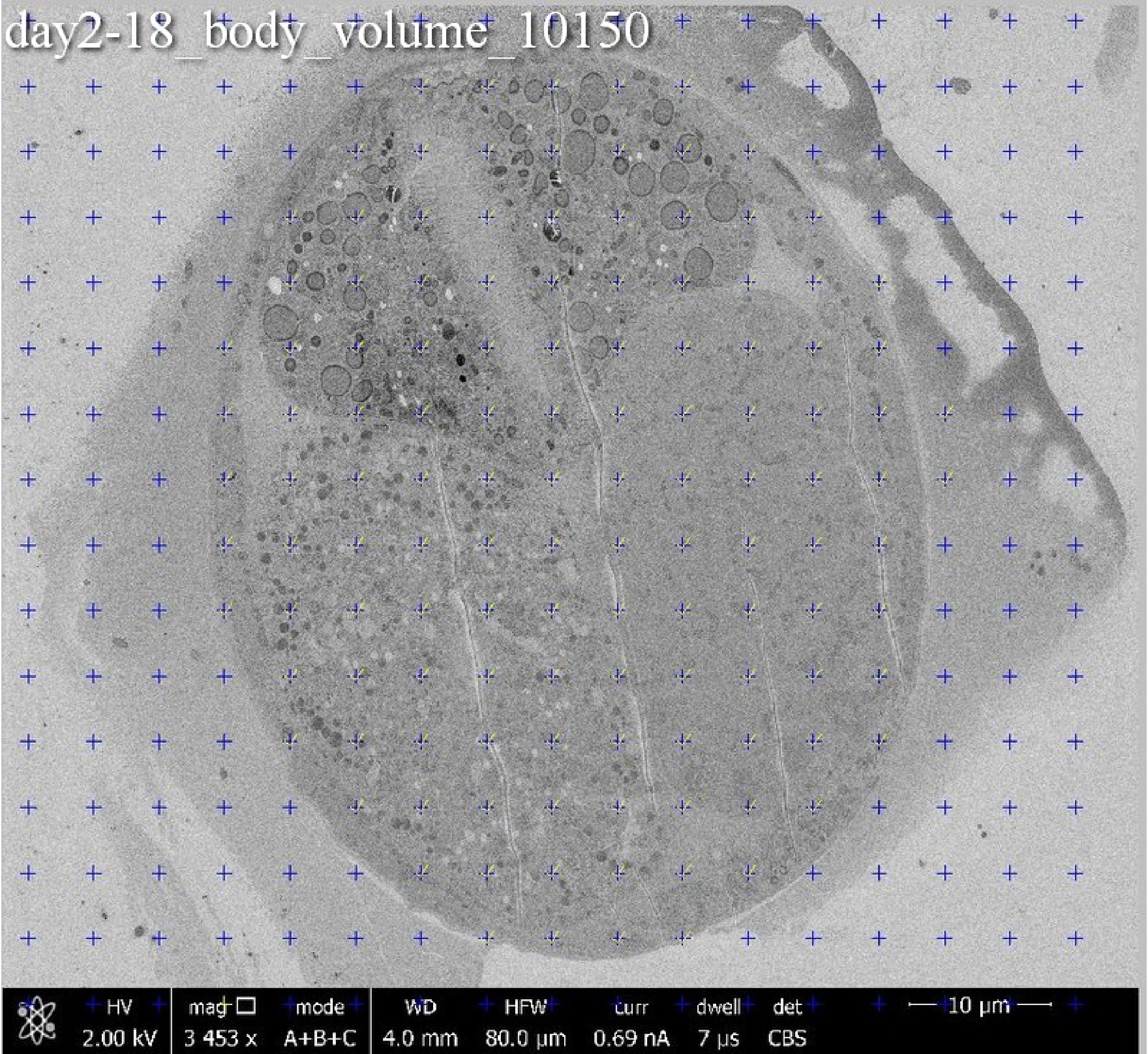
day2-18_body_volume_10150
HV
2.00 kV
mag
3 453 x
mode
A+B+C
WD
4.0 mm
HFW
80.0 µm
curr
0.69 nA
dwell
7 µs
det
CBS
10 µm

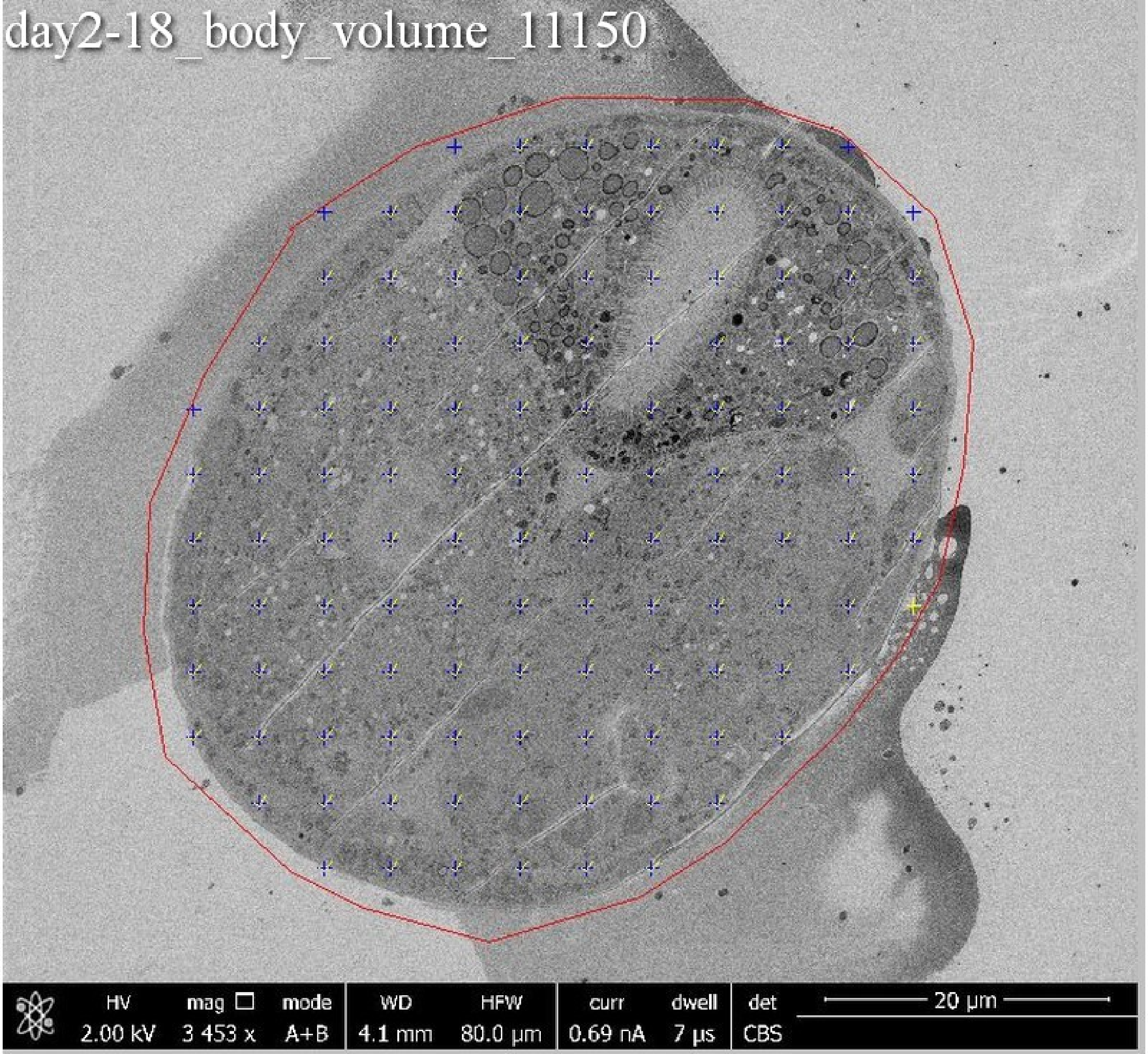

day2-18_body_volume_11150
HV
2.00 kV
mag
3 453 x
mode
A+B
WD
4.1 mm
HFW
80.0 µm
curr
0.69 nA
dwell
7 µs
det
CBS
20 µm

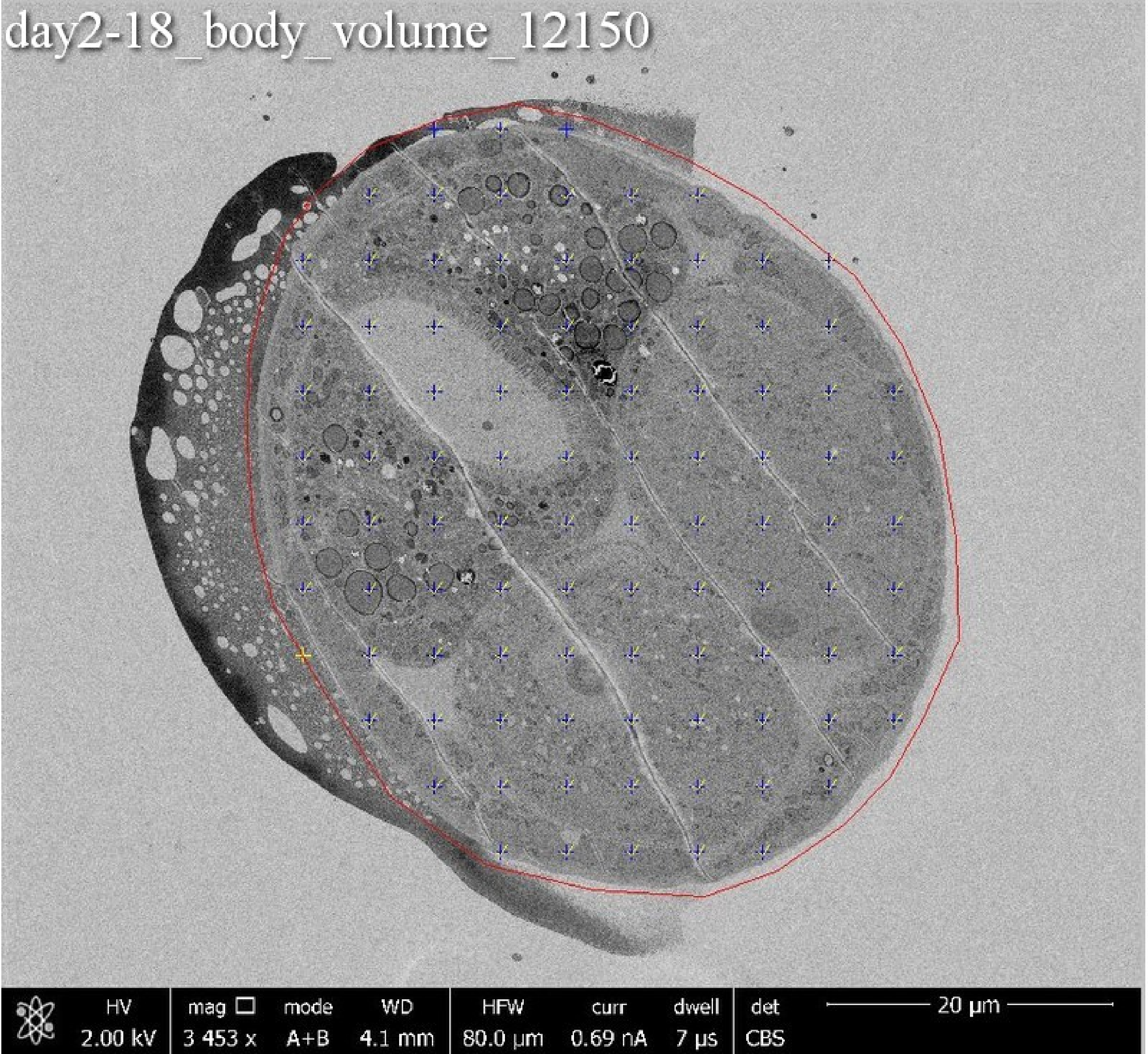
day2-18_body_volume_12150
HV
2.00 kV
mag
3 453 x
mode
A+B
WD
4.1 mm
HFW
80.0 µm
curr
0.69 nA
dwell
7 µs
det
CBS
20 µm

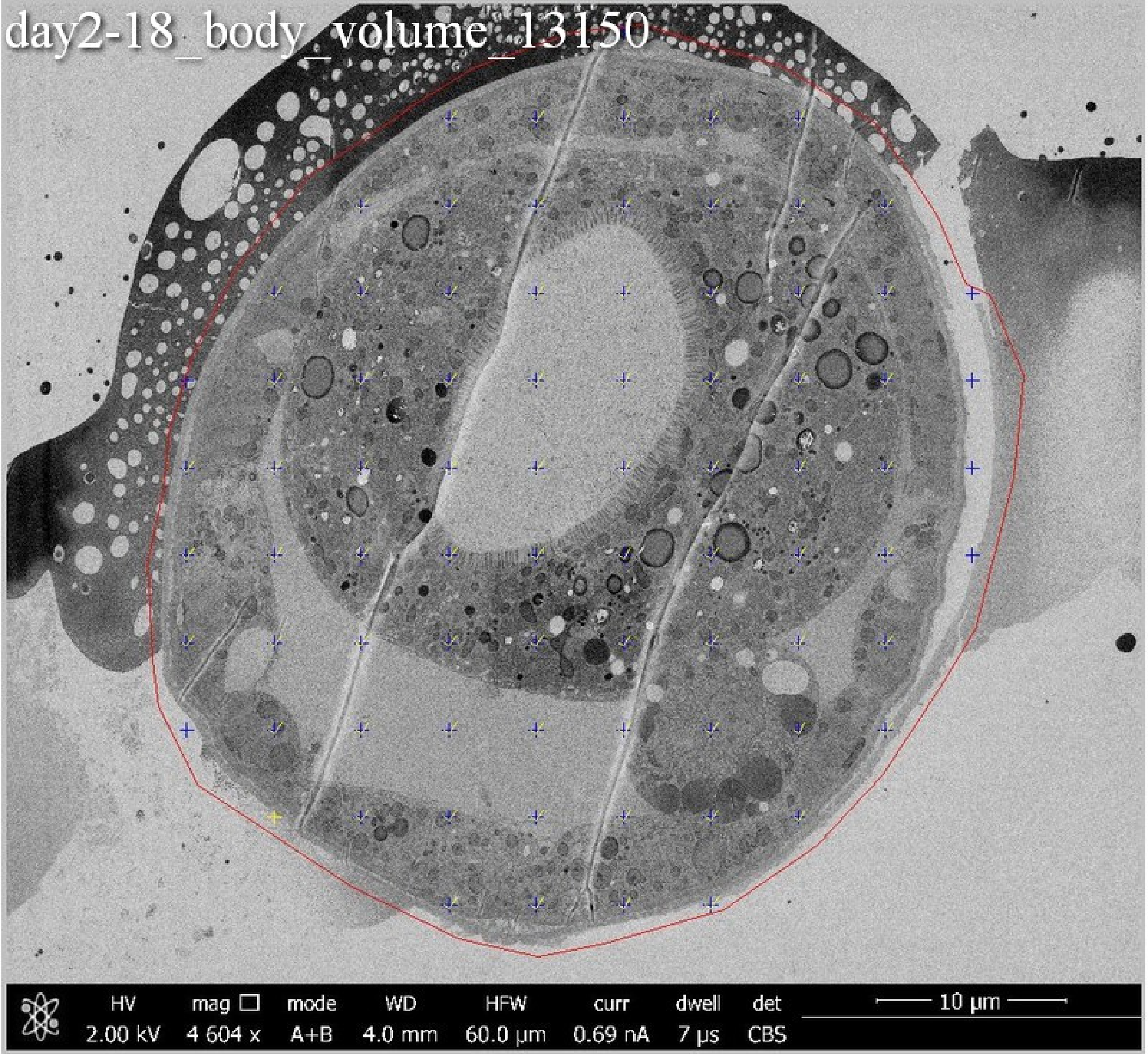
day2-18_body_volume_13150
HV 2.00 kV
mag 4 604 x
mode A+B
WD 4.0 mm
HFW 60.0 µm
curr 0.69 nA
dwell 7 µs
det CBS
10 µm

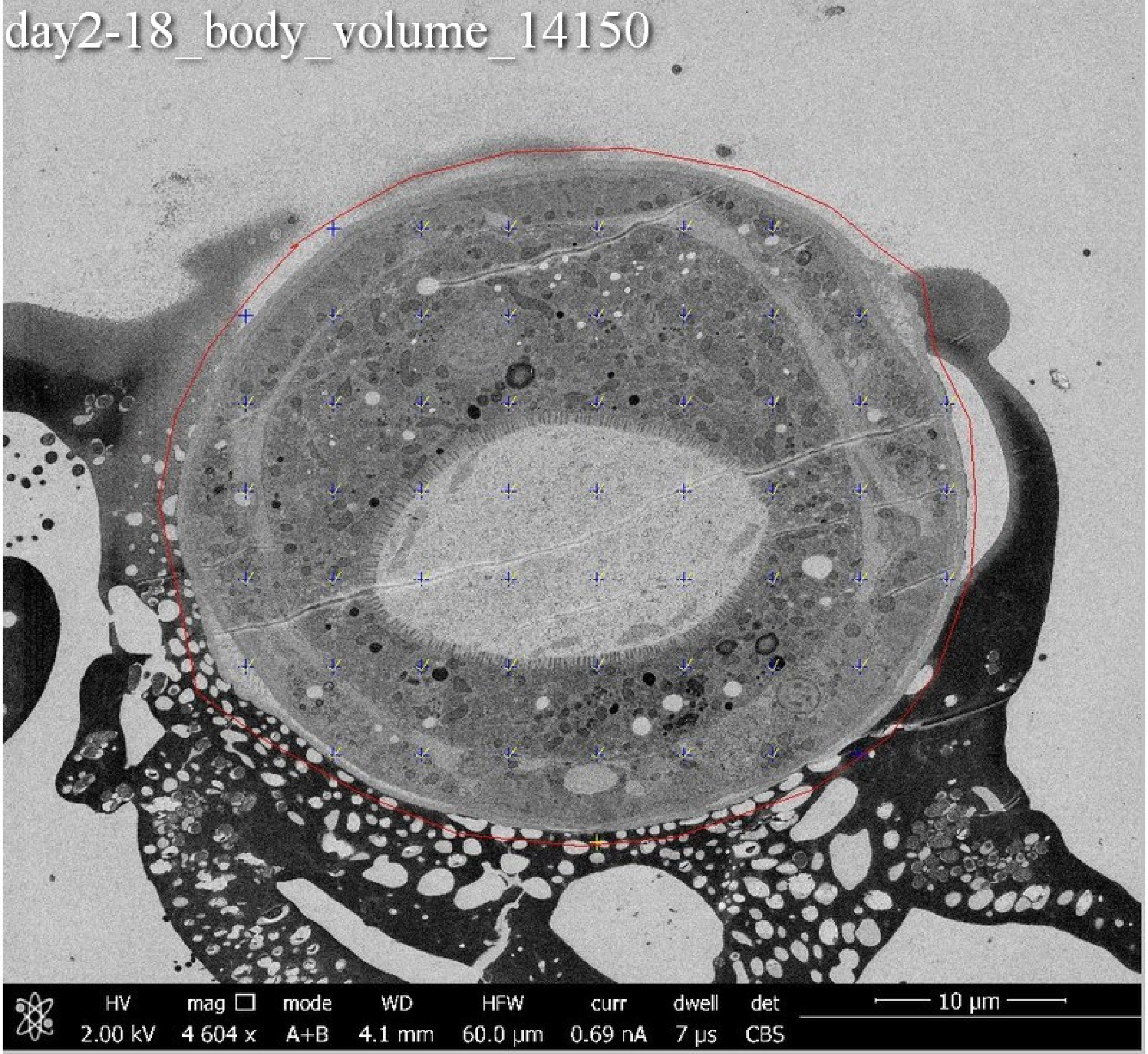

day2-18_body_volume_14150
HV
2.00 kV
mag
4 604 x
mode
A+B
WD
4.1 mm
HFW
60.0 µm
curr
0.69 nA
dwell
7 µs
det
CBS
10 µm

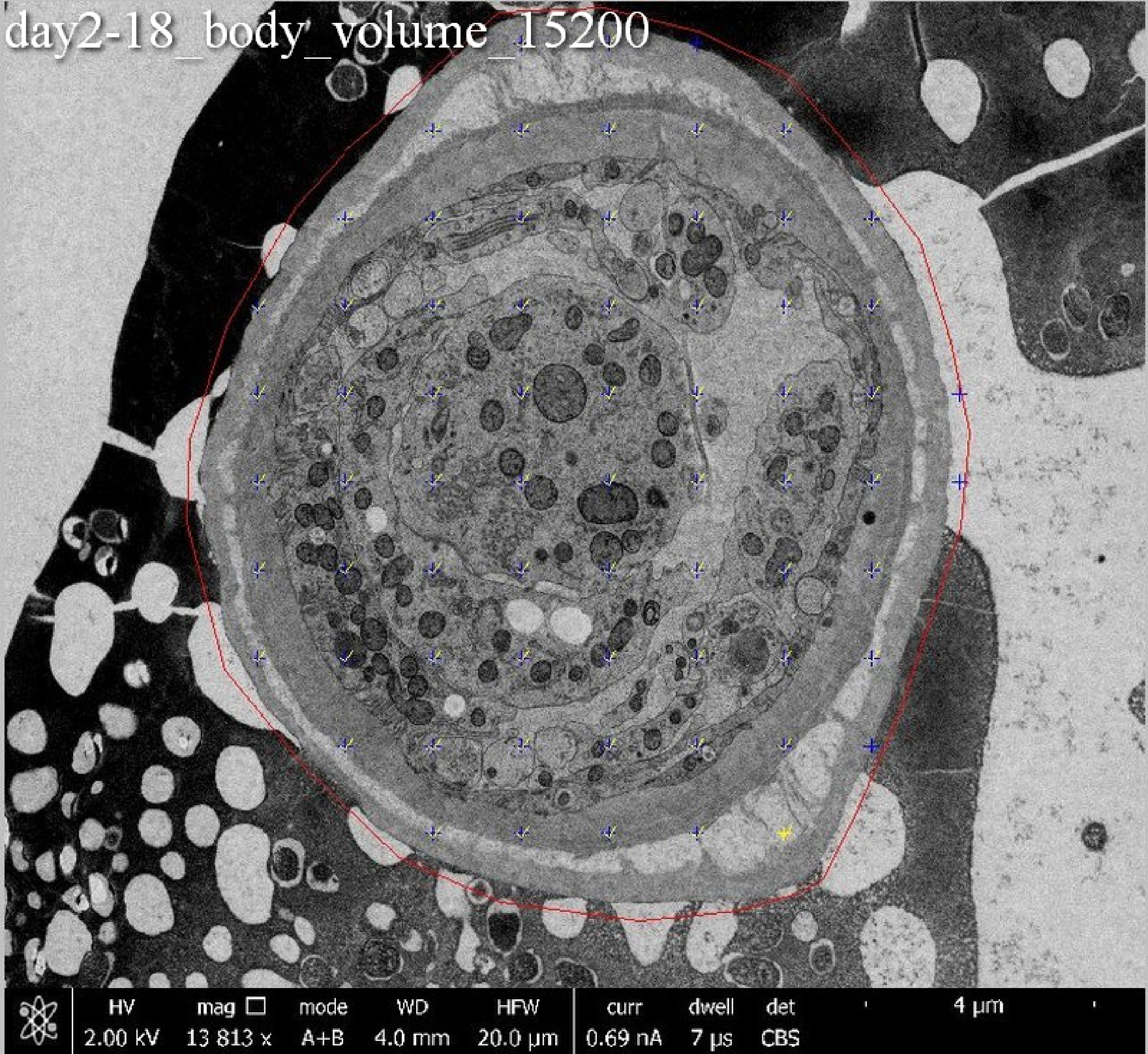
day2-18_body_volume_15200
HV
2.00 kV
mag
13 813 x
mode
A+B
WD
4.0 mm
HFW
20.0 μm
curr
0.69 nA
dwell
7 μs
det
CBS
4 μm

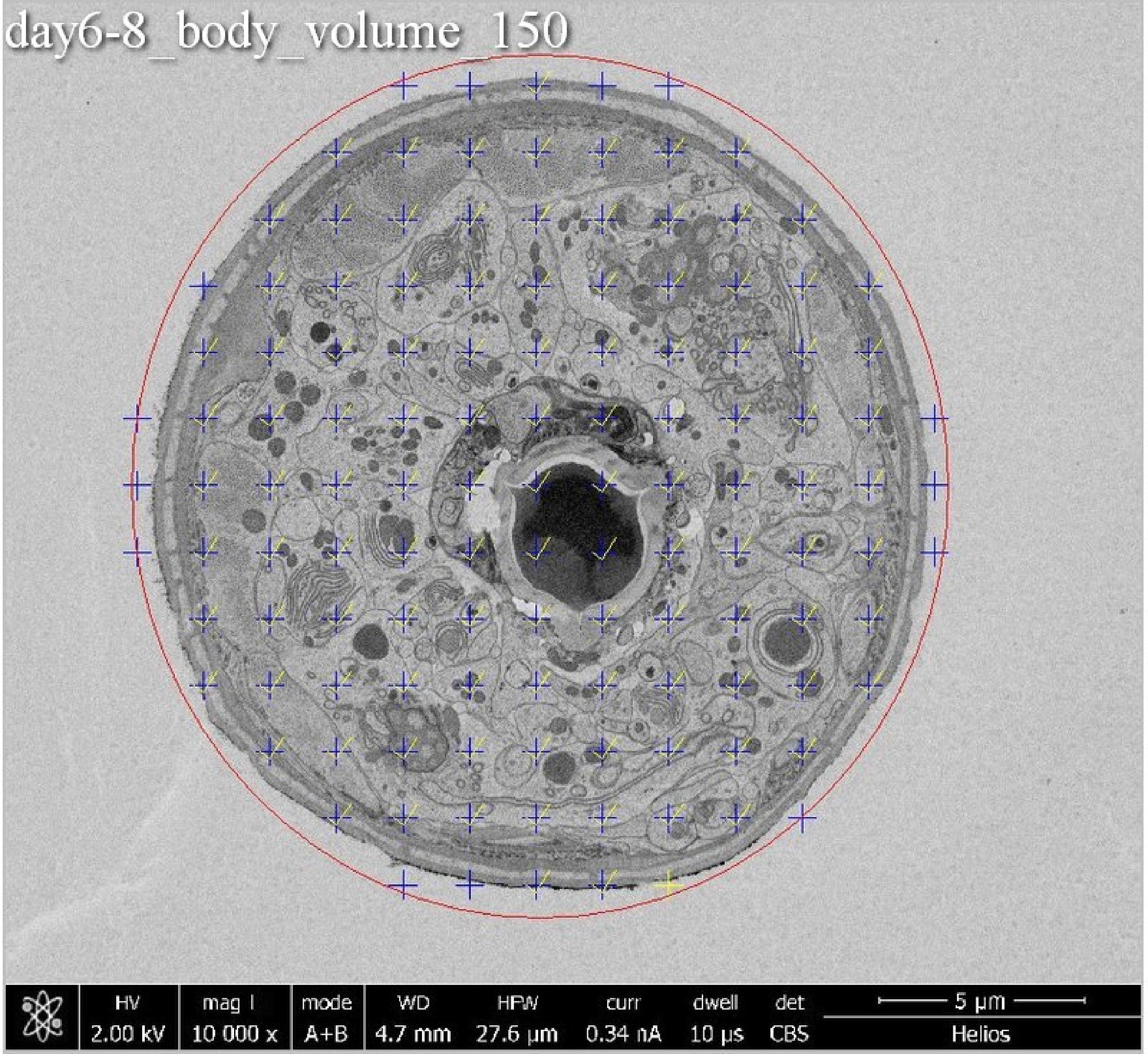
day6-8_body_volume_150
HV
2.00 kV
mag I
10 000 x
mode
A+B
WD
4.7 mm
HFW
27.6 µm
curr
0.34 nA
dwell
10 µs
det
CBS
5 µm
Helios

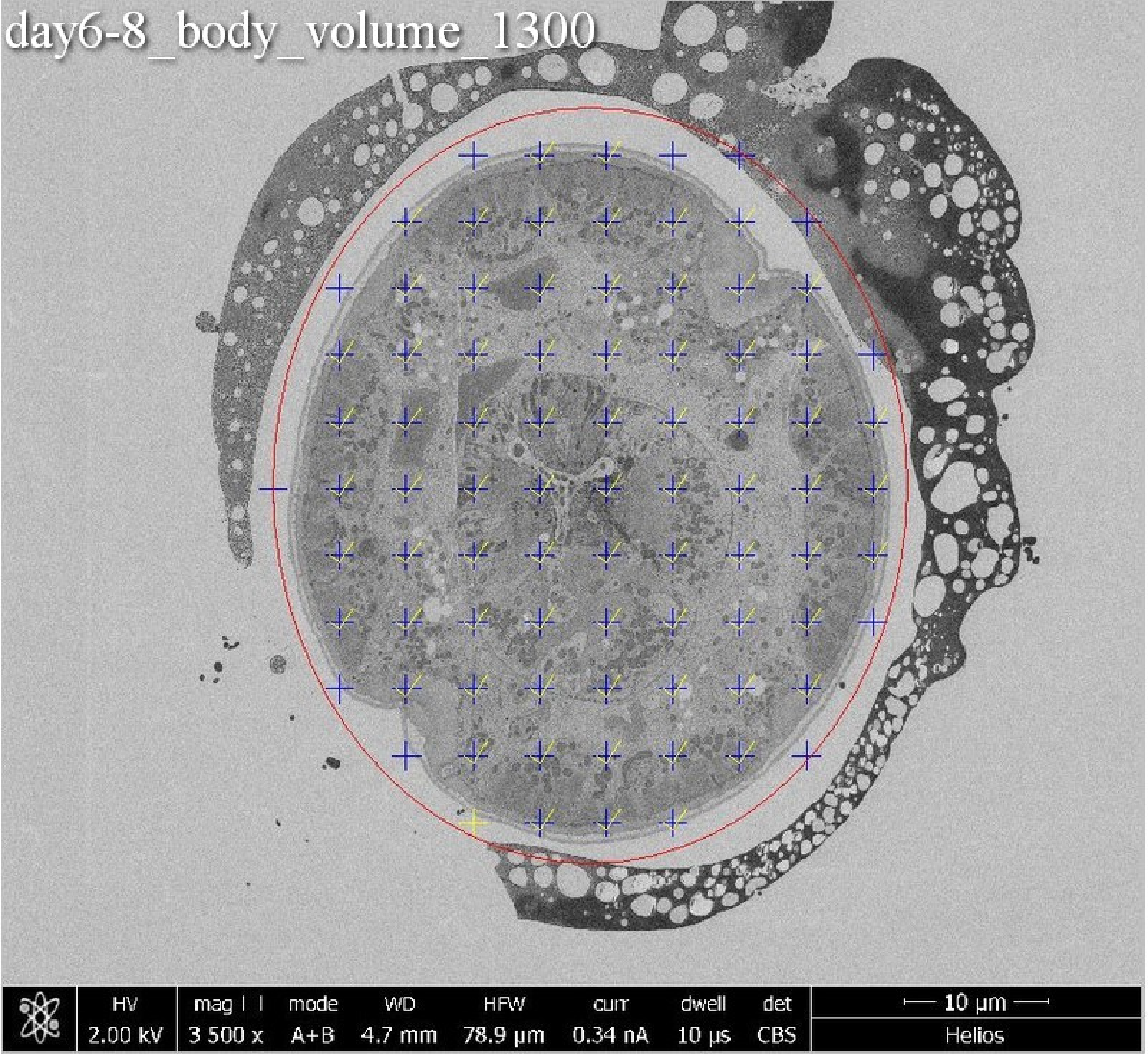
day6-8_body_volume_1300
HV
2.00 kV
mag
3 500 x
mode
A+B
WD
4.7 mm
HFW
78.9 µm
curr
0.34 nA
dwell
10 µs
det
CBS
10 µm
Helios

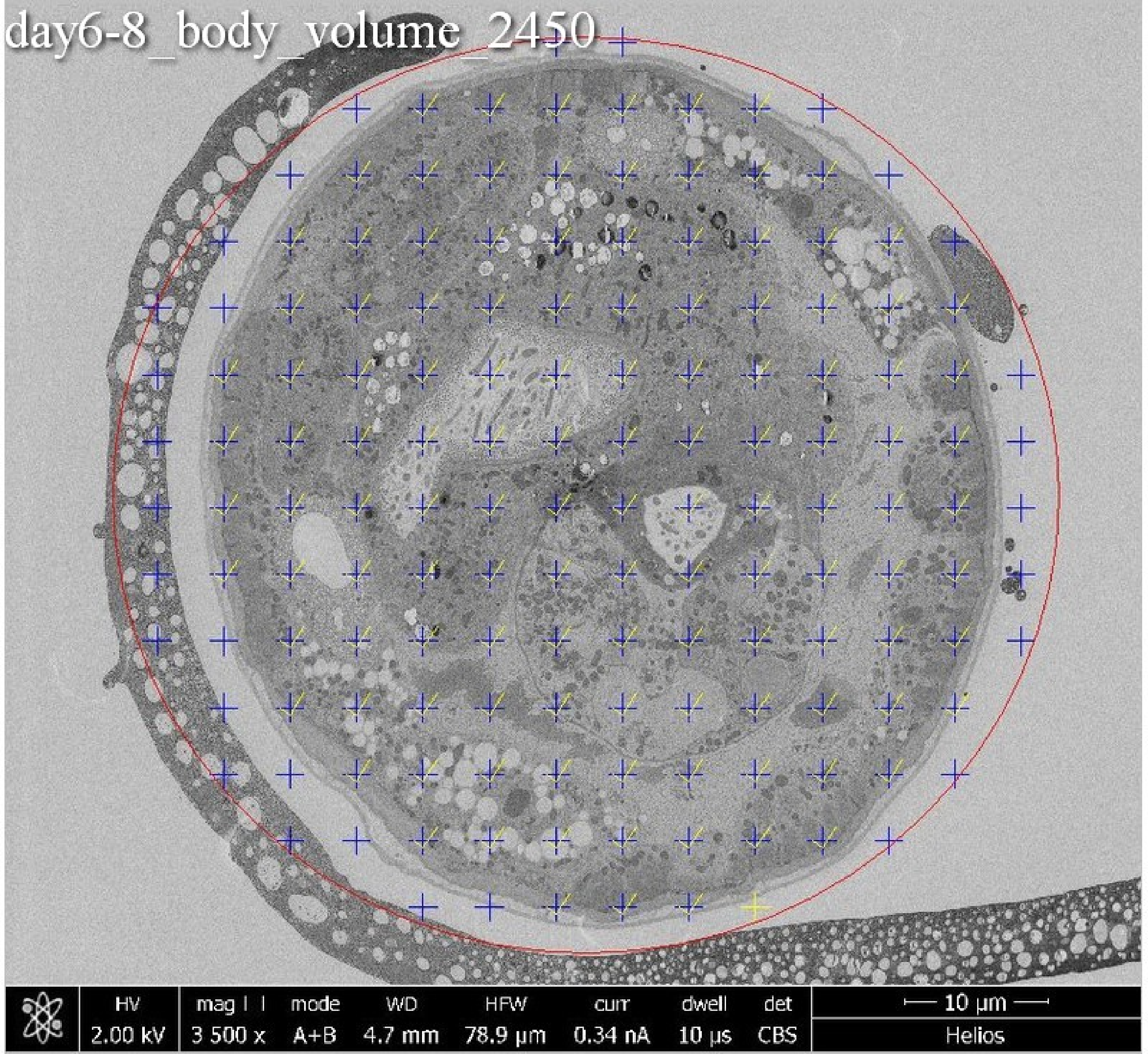
day6-8_body_volume_2450
HV 2.00 kV
mag 3 500 x
mode A+B
WD 4.7 mm
HFW 78.9 µm
curr 0.34 nA
dwell 10 µs
det CBS
10 µm
Helios

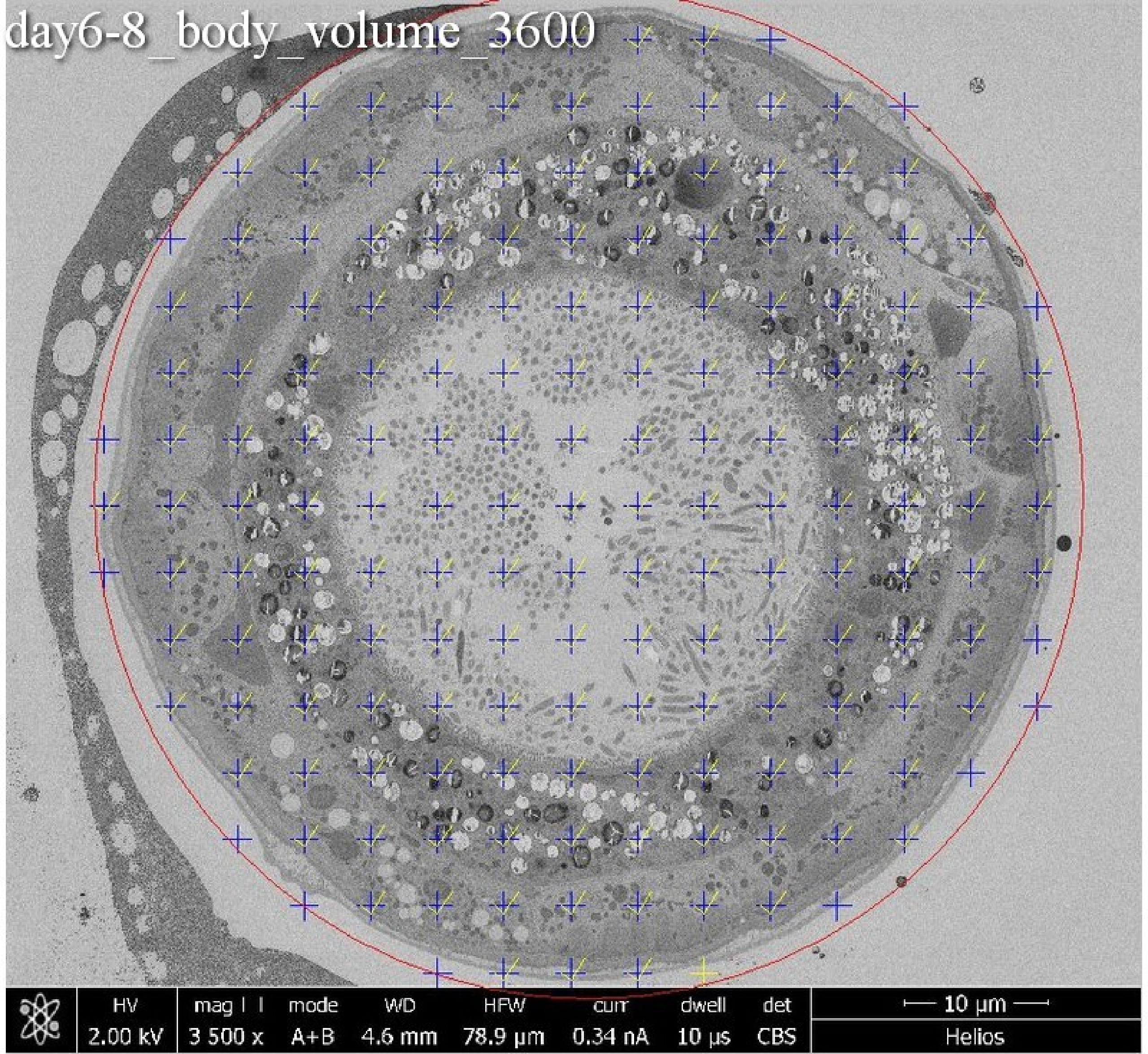
day6-8_body_volume_3600
HV
2.00 kV
mag
3 500 x
mode
A+B
WD
4.6 mm
HFW
78.9 µm
curr
0.34 nA
dwell
10 µs
det
CBS
10 µm
Helios

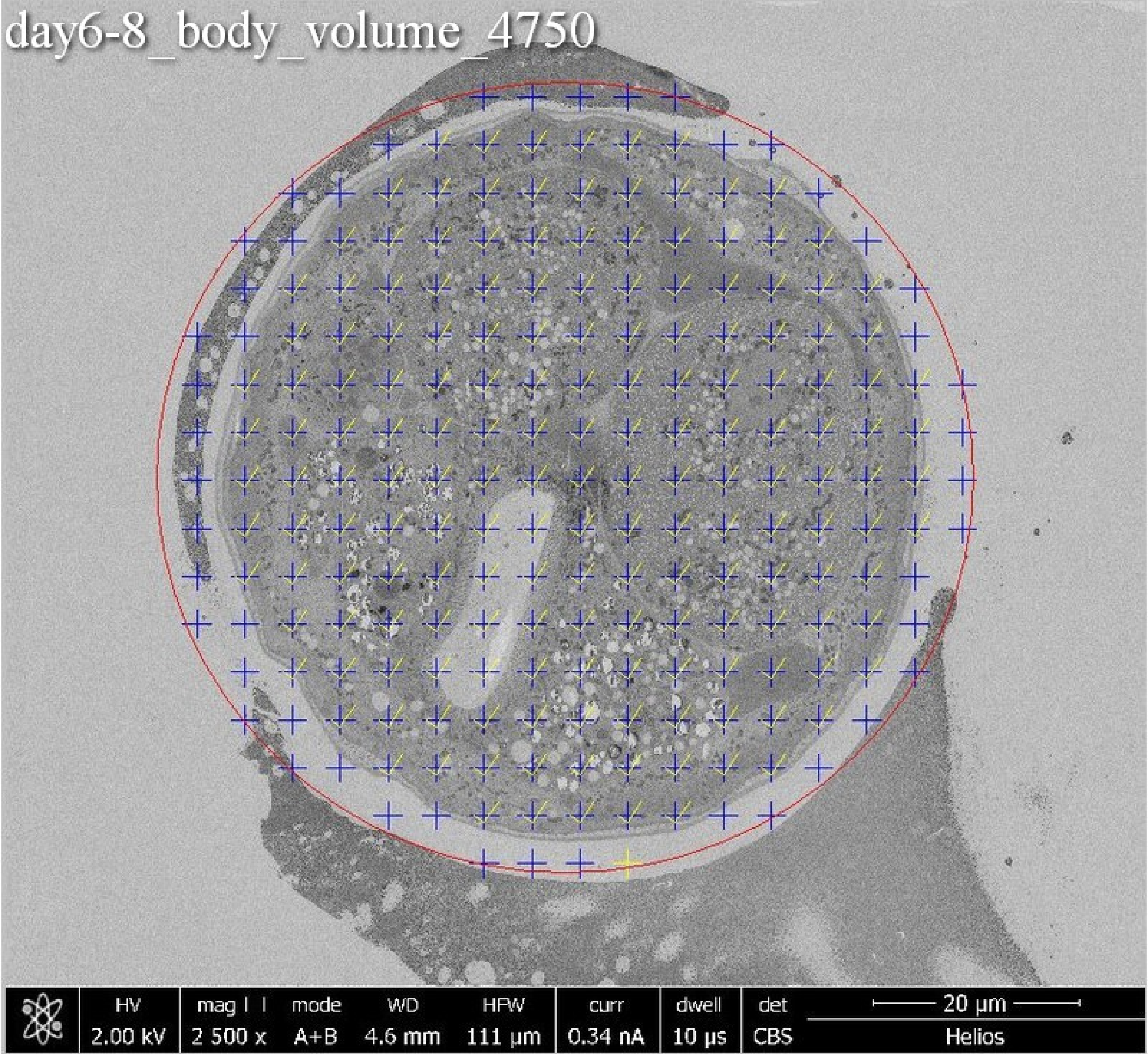
day6-8_body_volume_4750
HV
2.00 kV
mag
2 500 x
mode
A+B
WD
4.6 mm
HFW
111 μm
curr
0.34 nA
dwell
10 μs
det
CBS
20 μm
Helios

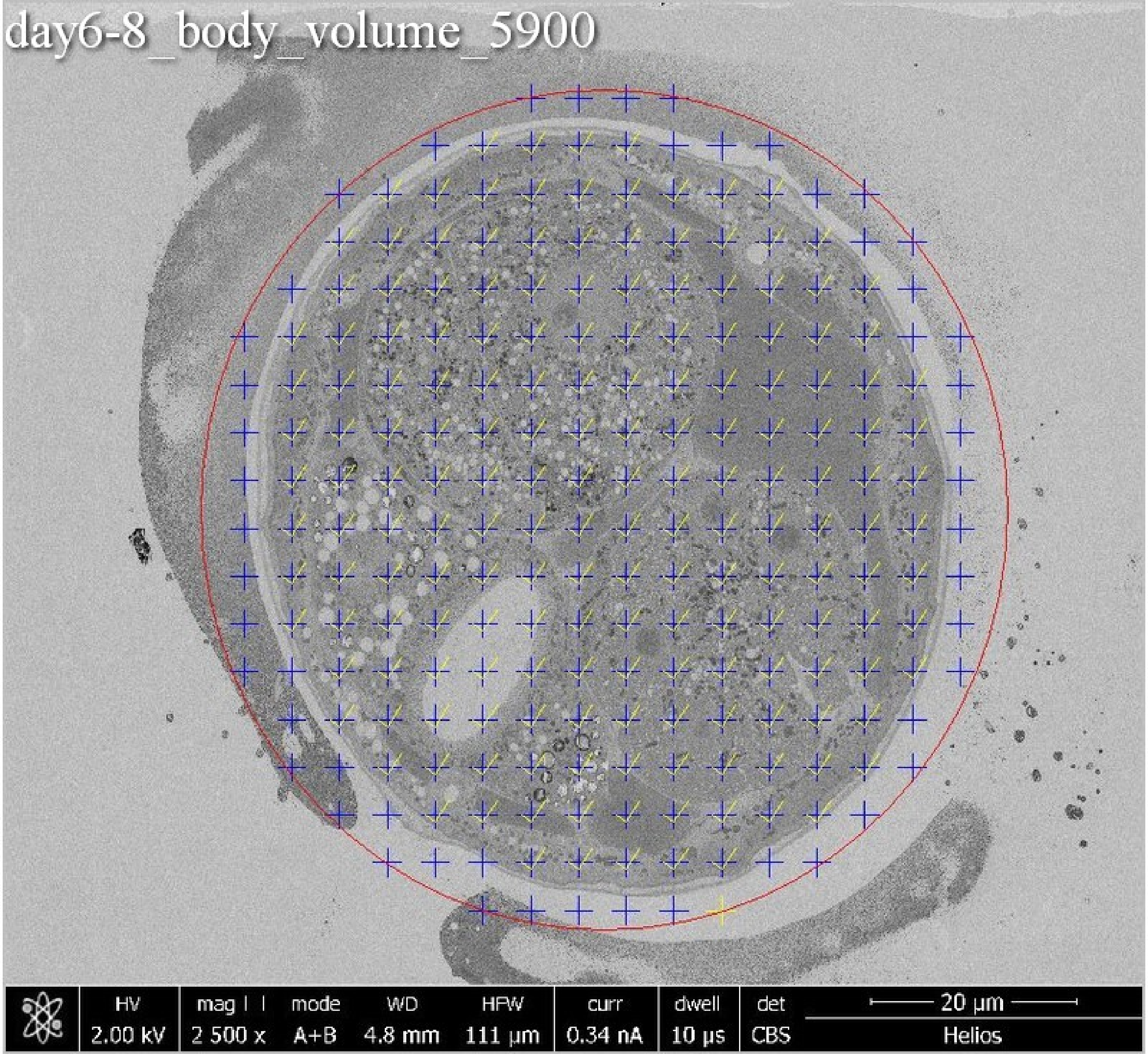
day6-8_body_volume_5900
HV
2.00 kV
mag
2 500 x
mode
A+B
WD
4.8 mm
HFW
111 µm
curr
0.34 nA
dwell
10 µs
det
CBS
20 µm
Helios

day6-8_body_volume_7050

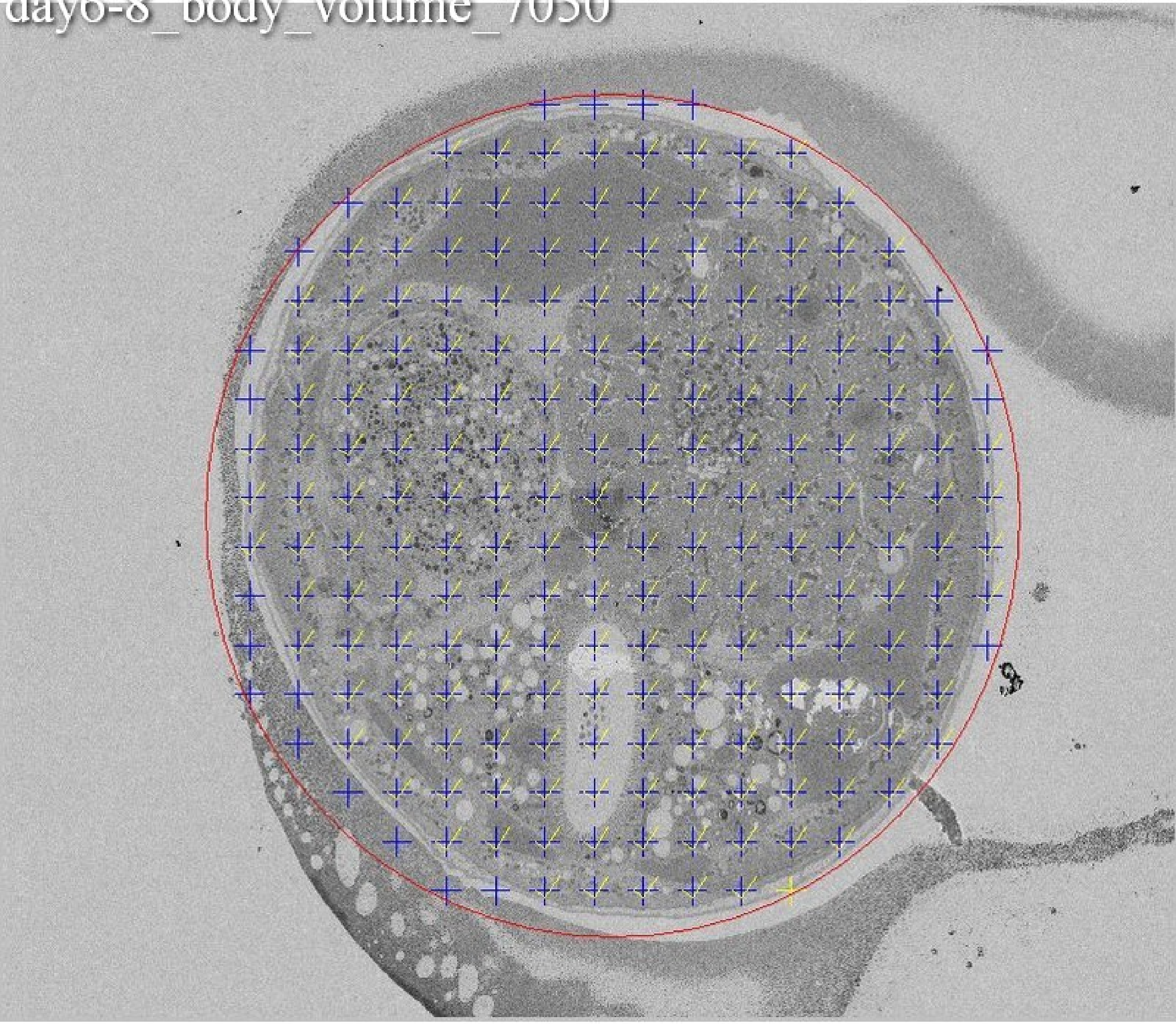

day6-8_body_volume_8200

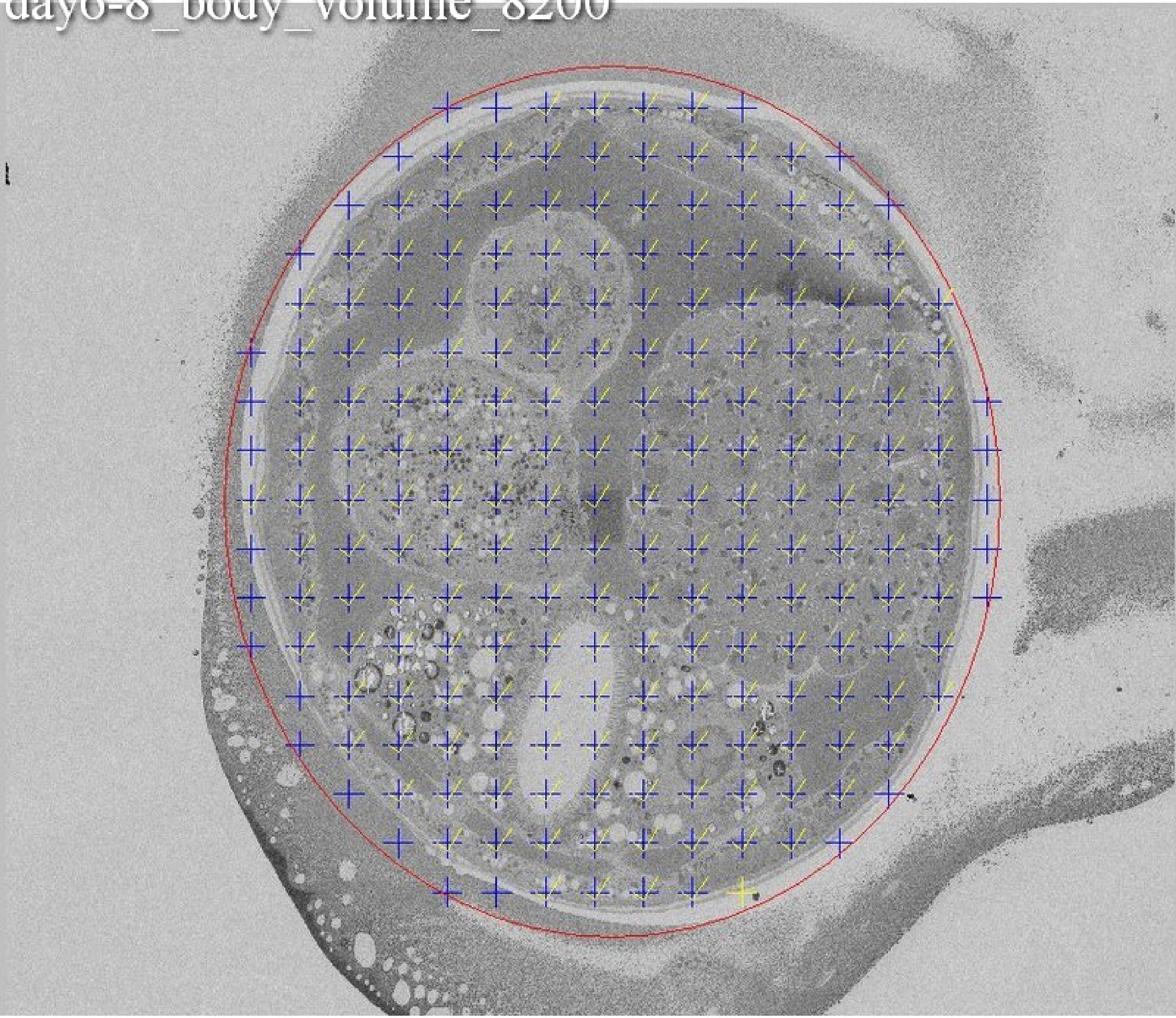

day6-8_body_volume_9350

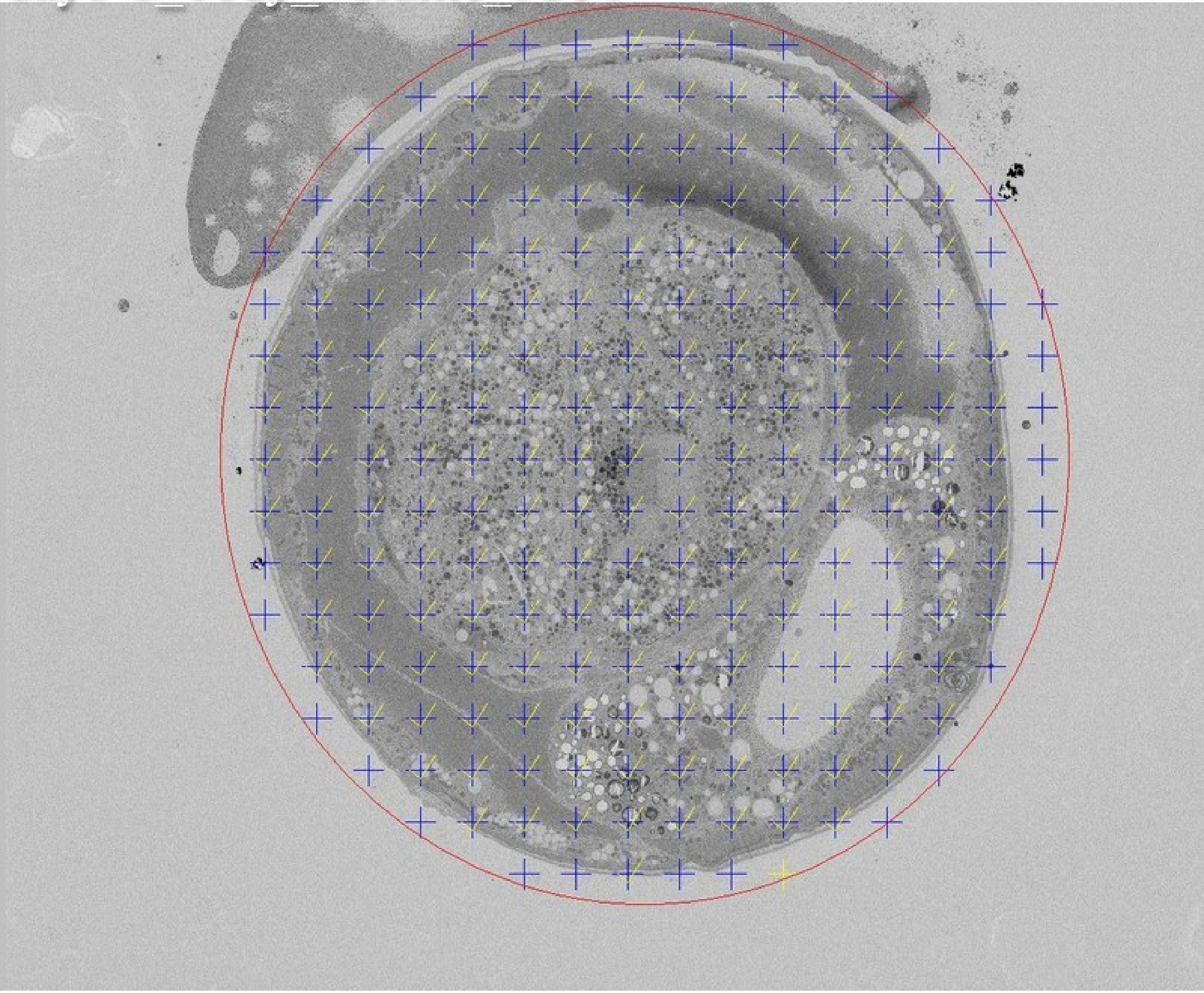

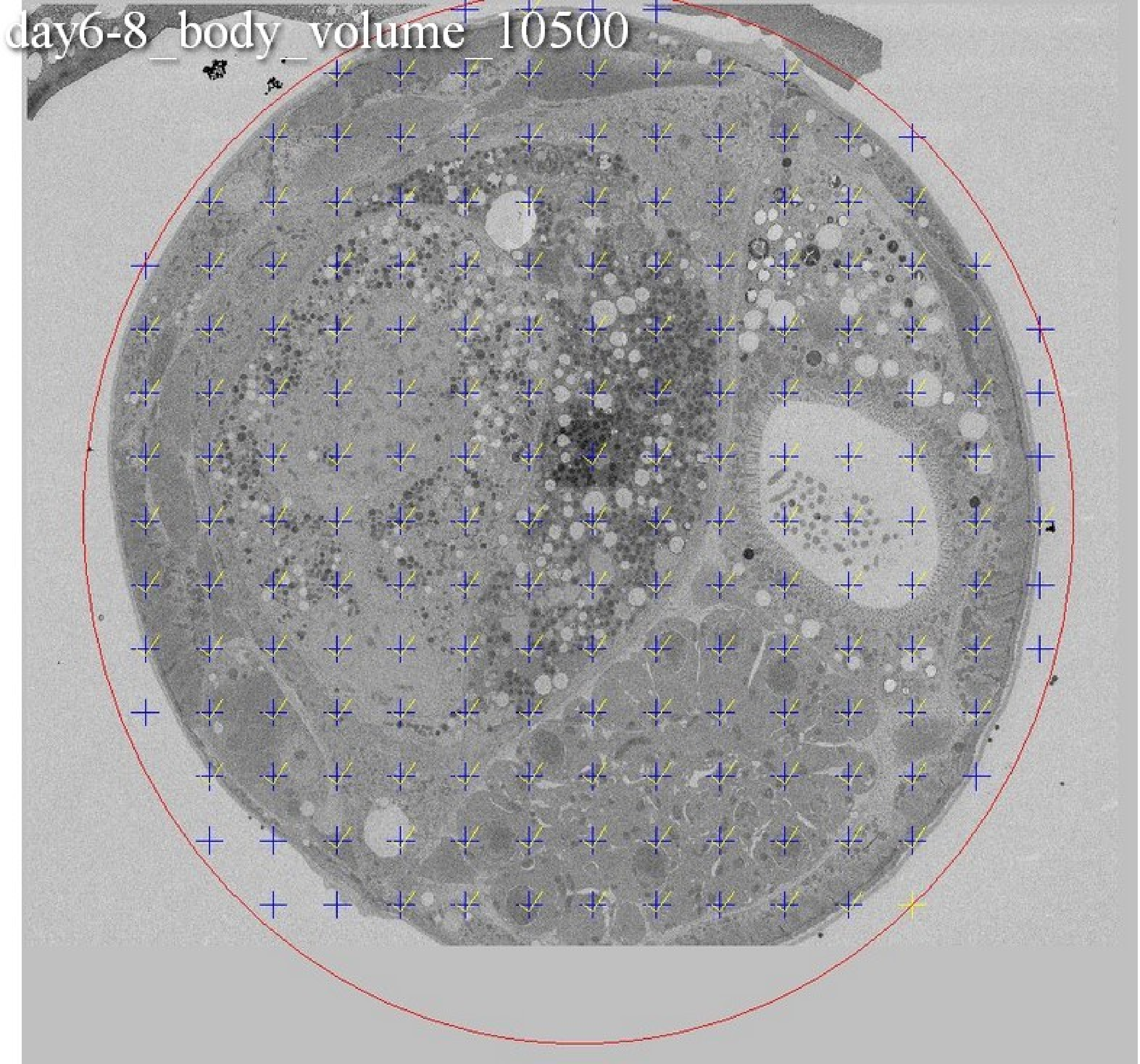
day6-8_body_volume_10500

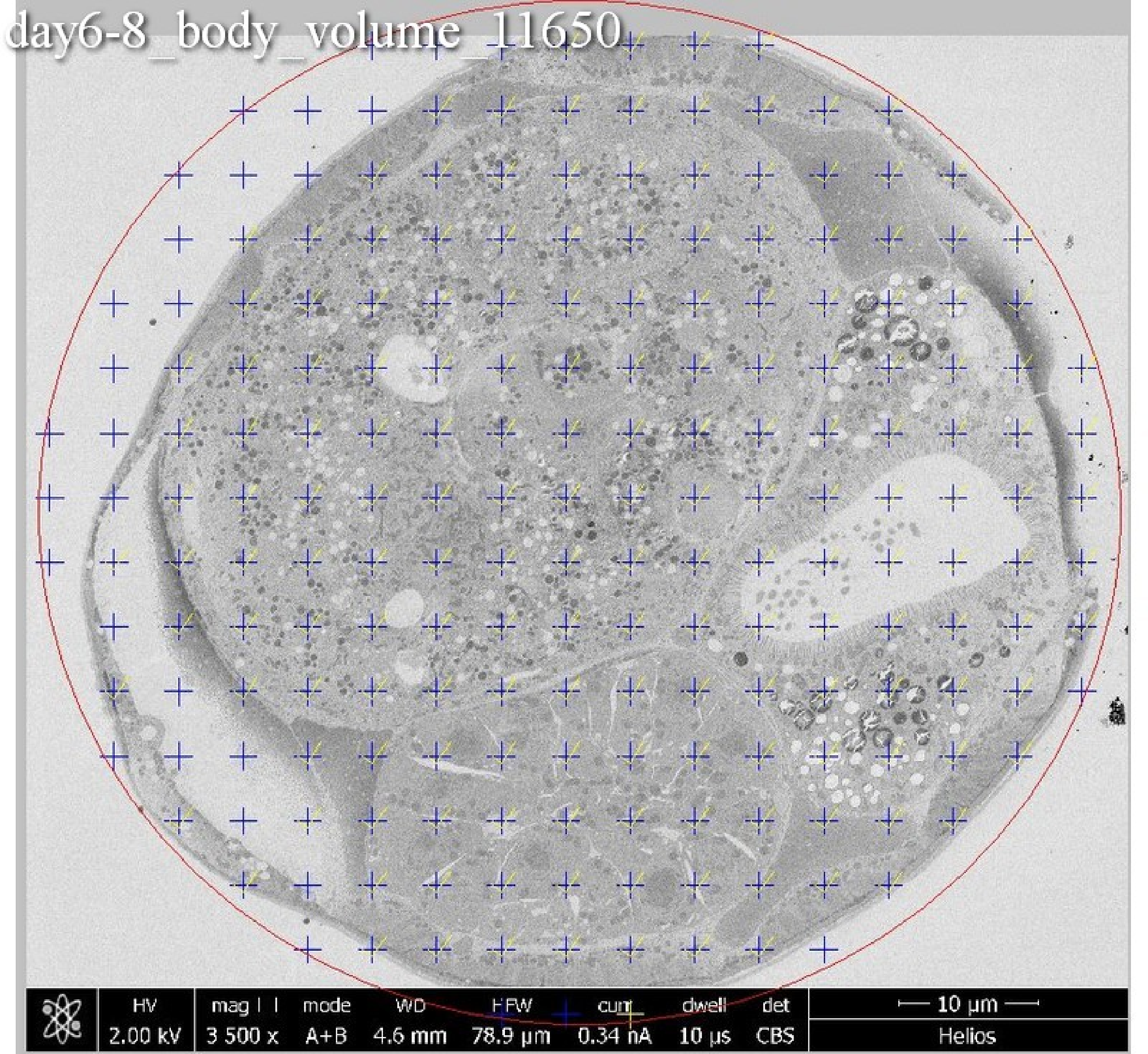
day6-8_body_volume_11650
HV
2.00 kV
mag
3 500 x
mode
A+B
WD
4.6 mm
HFW
78.9 µm
curr
0.34 nA
dwell
10 µs
det
CBS
10 µm
Helios

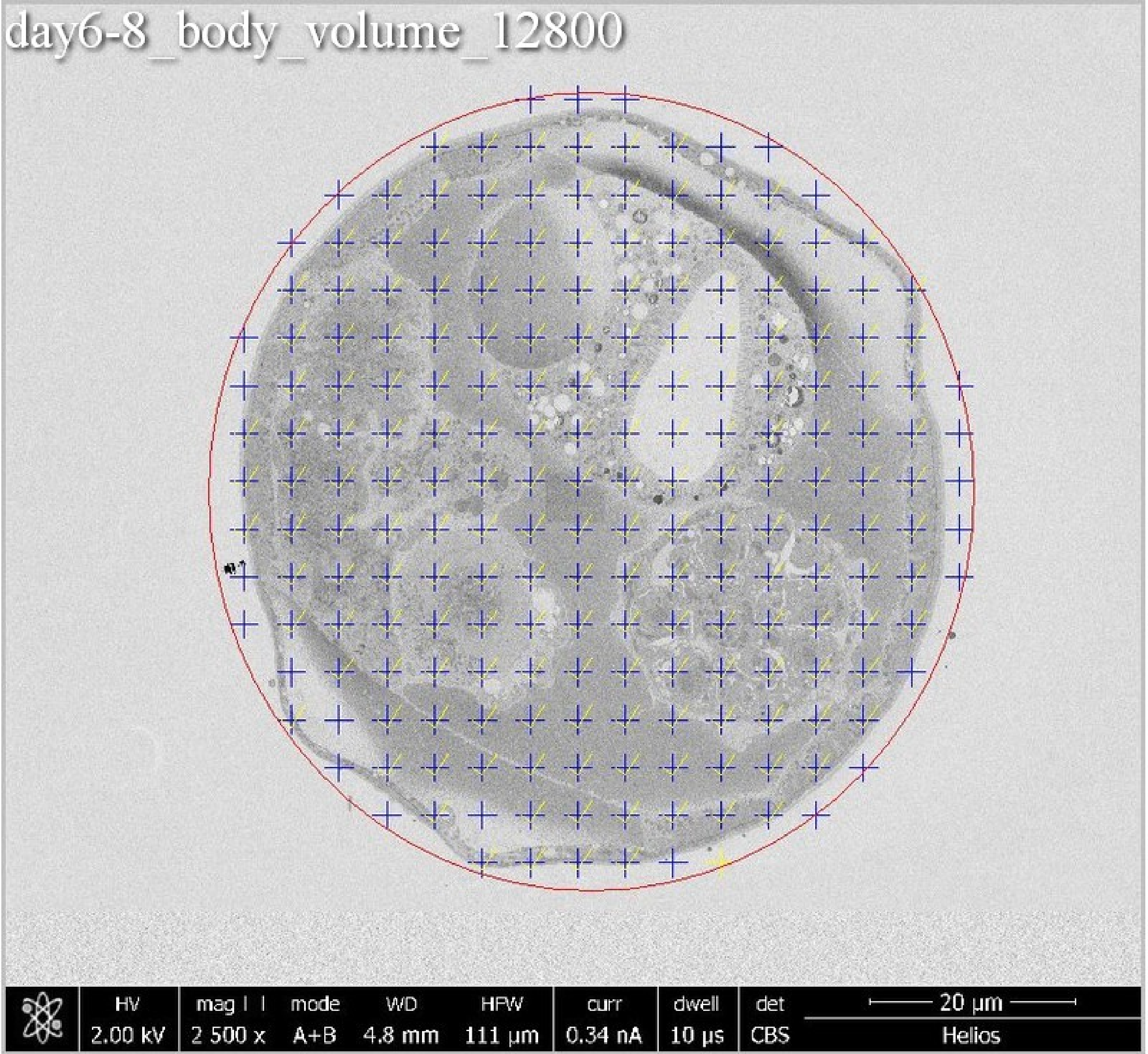
day6-8_body_volume_12800
HV
2.00 kV
mag
2 500 x
mode
A+B
WD
4.8 mm
HFW
111 µm
curr
0.34 nA
dwell
10 µs
det
CBS
20 µm
Helios

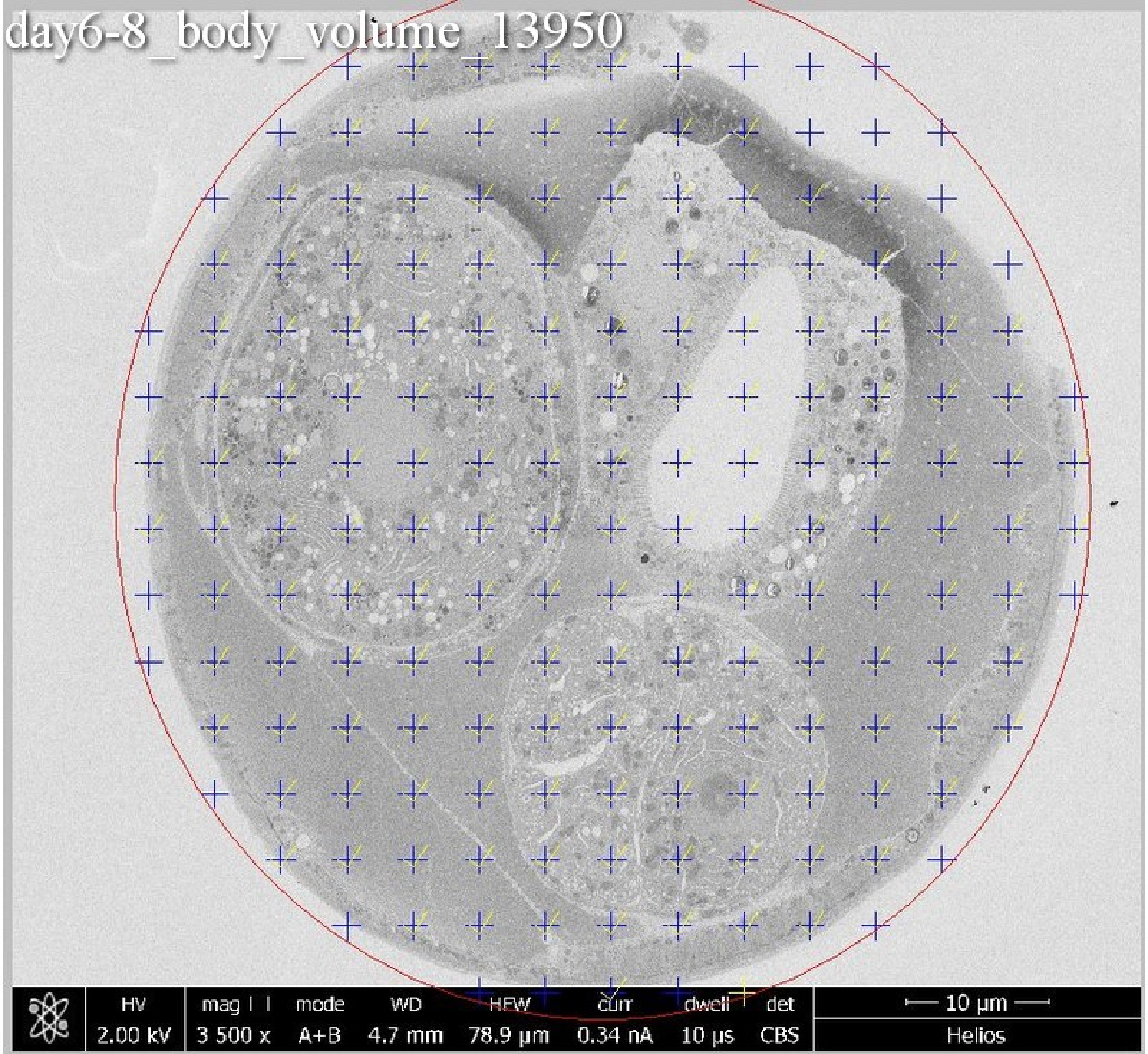
day6-8_body_volume_13950
HV 2.00 kV
mag 3 500 x
mode A+B
WD 4.7 mm
HFW 78.9 µm
curr 0.34 nA
dwell 10 µs
det CBS
10 µm
Helios

day6-8_body_volume_15100

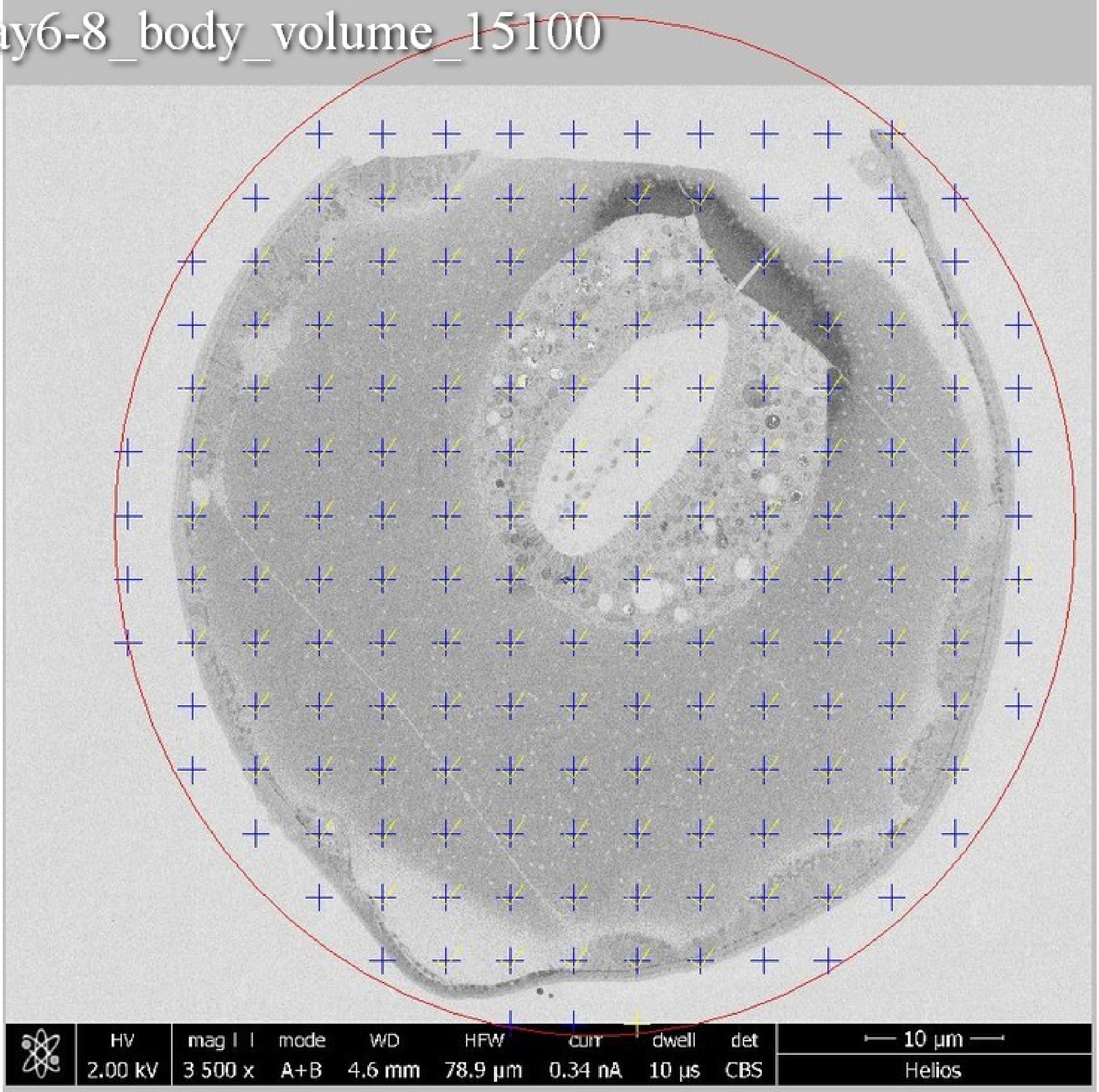

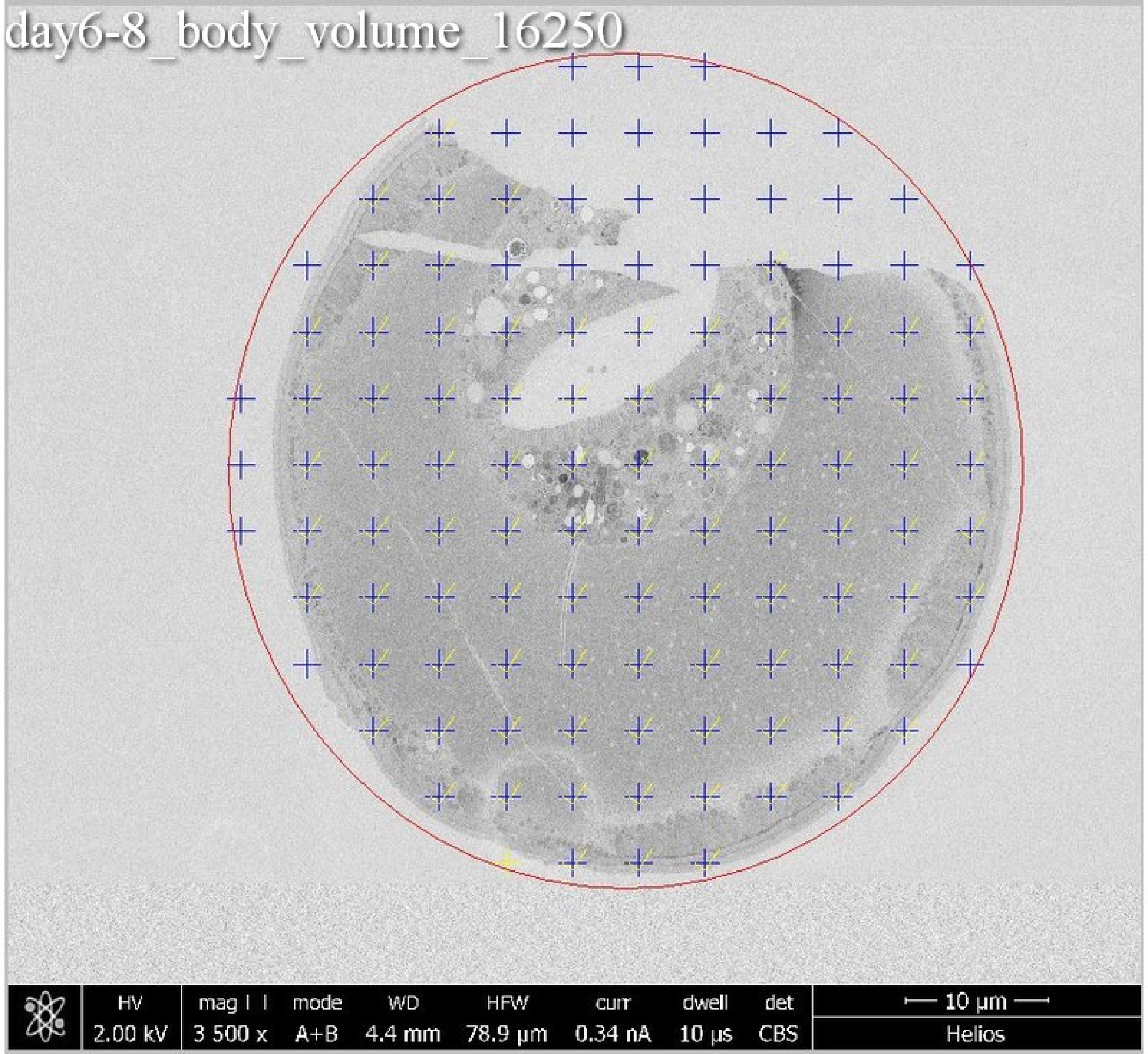
day6-8_body_volume_16250
HV 2.00 kV
mag 3 500 x
mode A+B
WD 4.4 mm
HFW 78.9 µm
curr 0.34 nA
dwell 10 µs
det CBS
10 µm
Helios

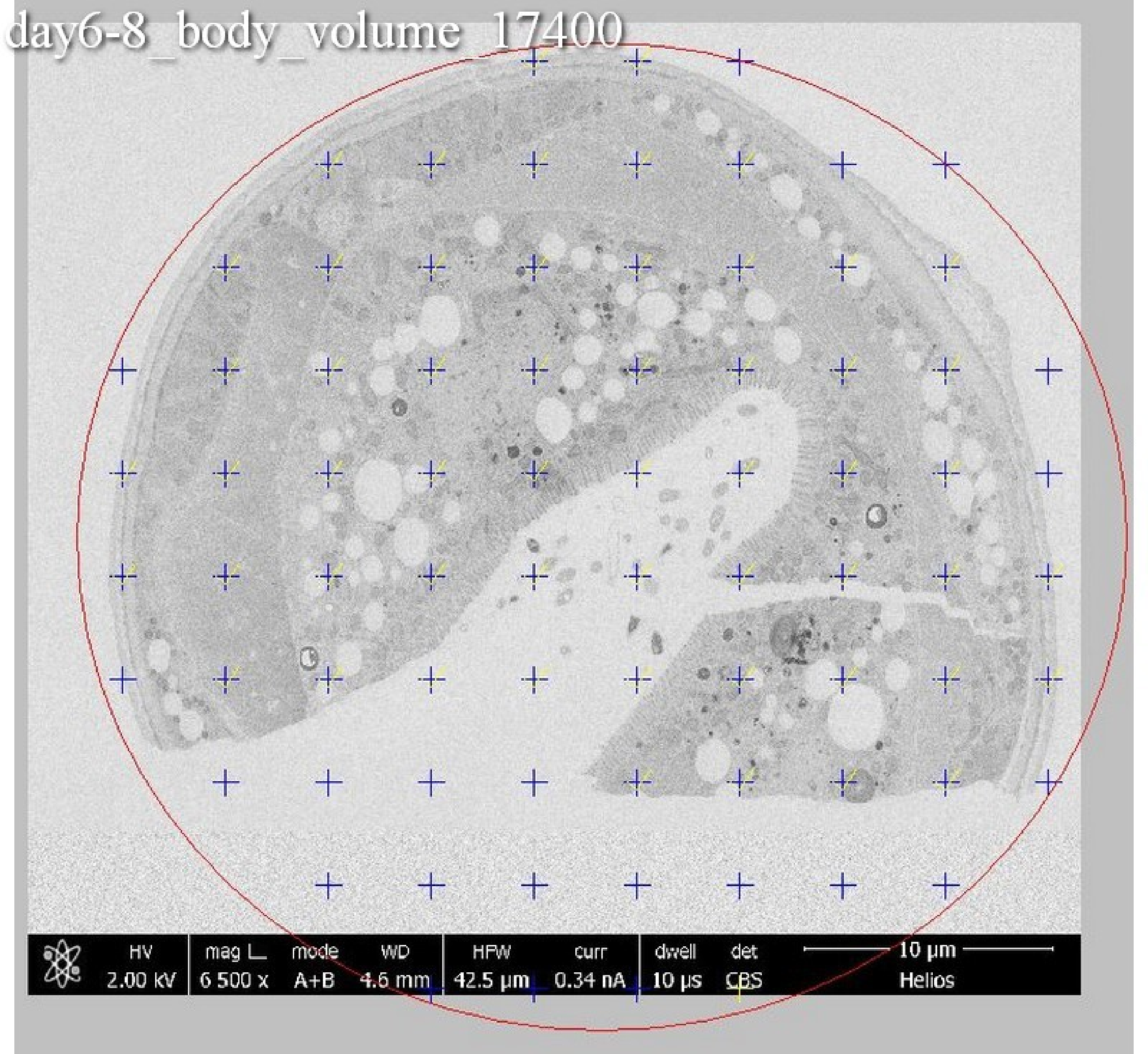
day6-8_body_volume_17400
HV
2.00 kV
mag ⊔
6 500 x
mode
A+B
WD
4.6 mm
HFW
42.5 µm
curr
0.34 nA
dwell
10 µs
det
CBS
10 µm
Helios

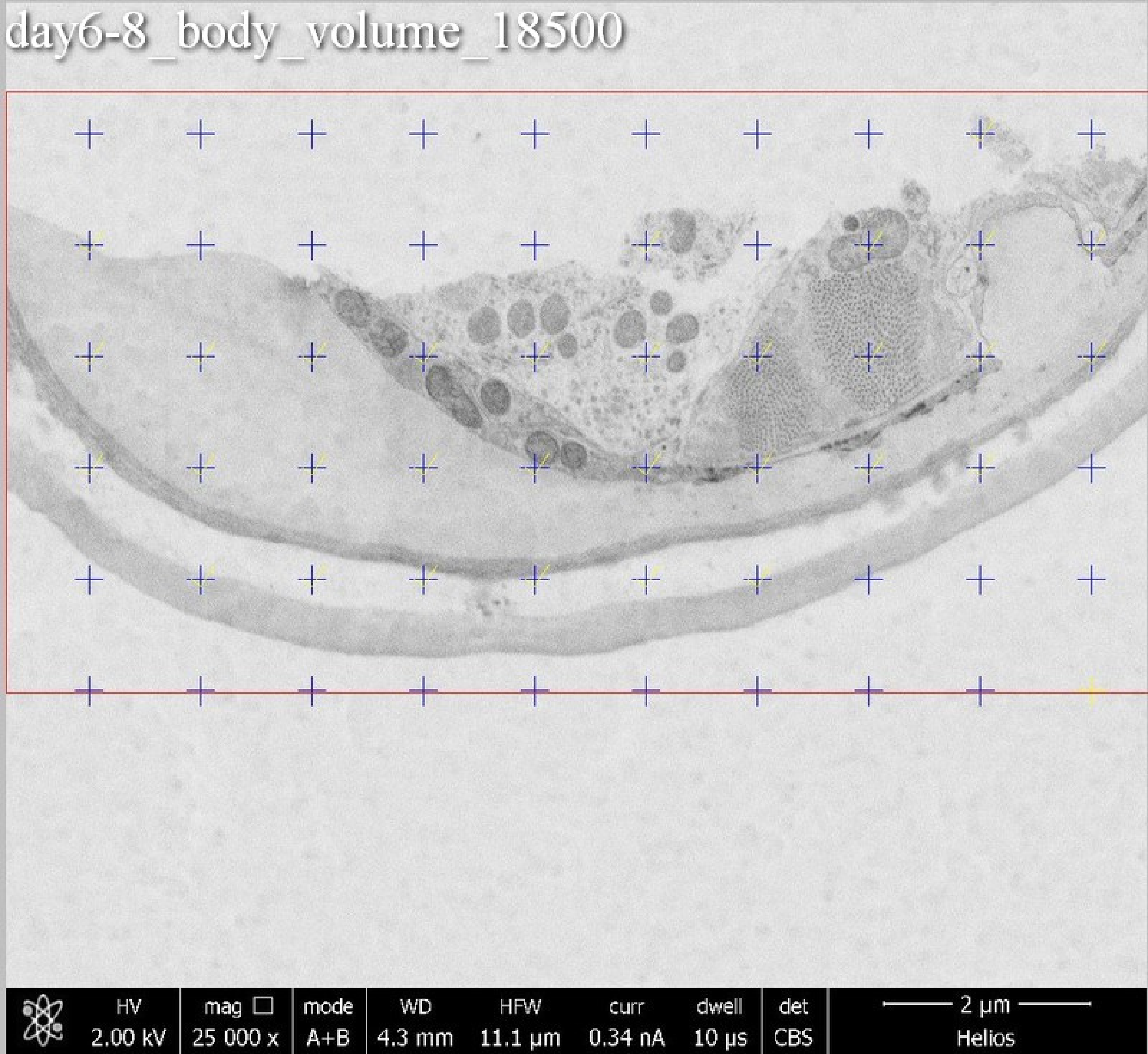
day6-8_body_volume_18500
HV 2.00 kV
mag 25 000 x
mode A+B
WD 4.3 mm
HFW 11.1 µm
curr 0.34 nA
dwell 10 µs
det CBS
2 µm
Helios

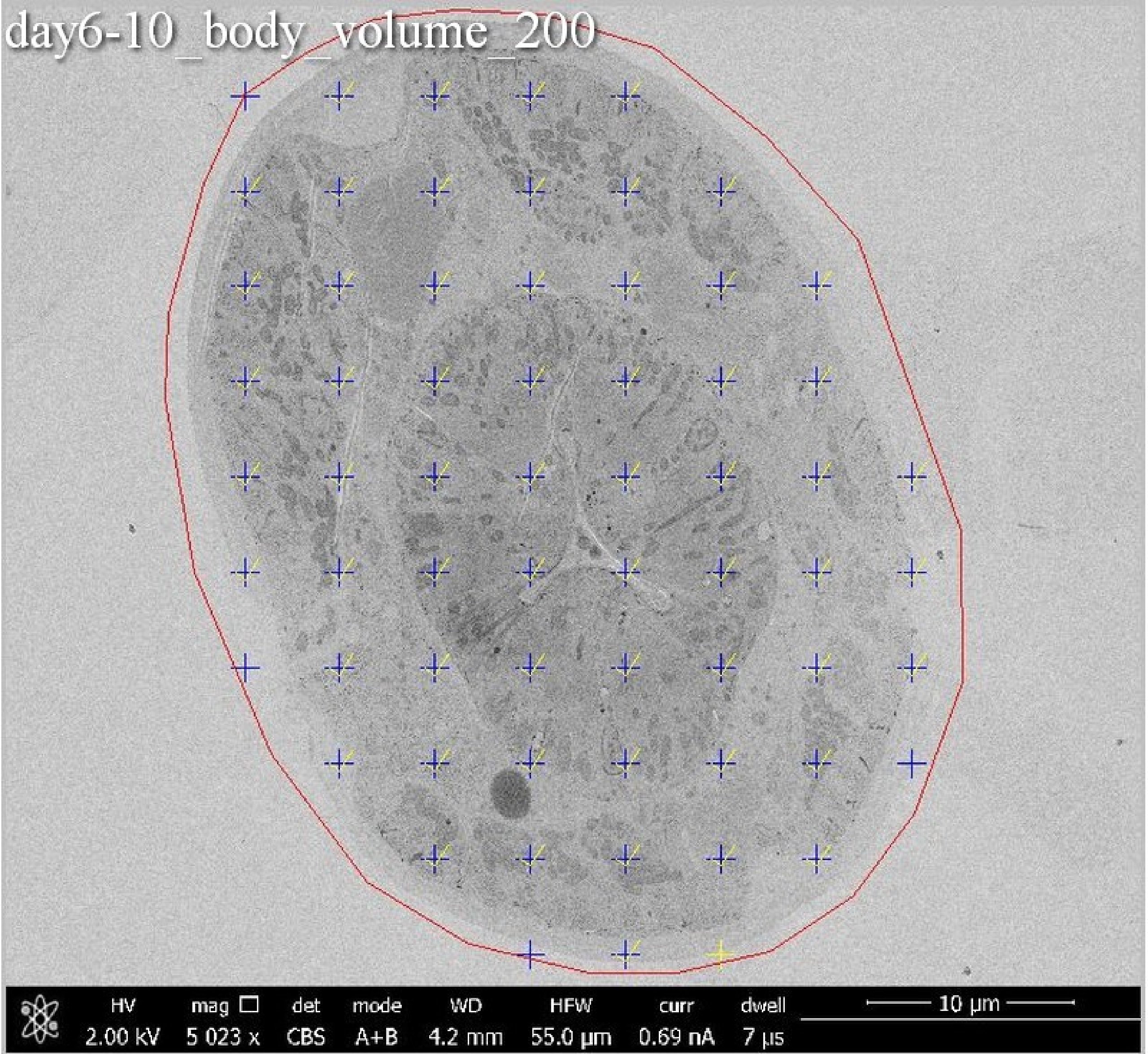
day6-10_body_volume_200
HV
2.00 kV
mag
5 023 x
det
CBS
mode
A+B
WD
4.2 mm
HFW
55.0 µm
curr
0.69 nA
dwell
7 µs
10 µm

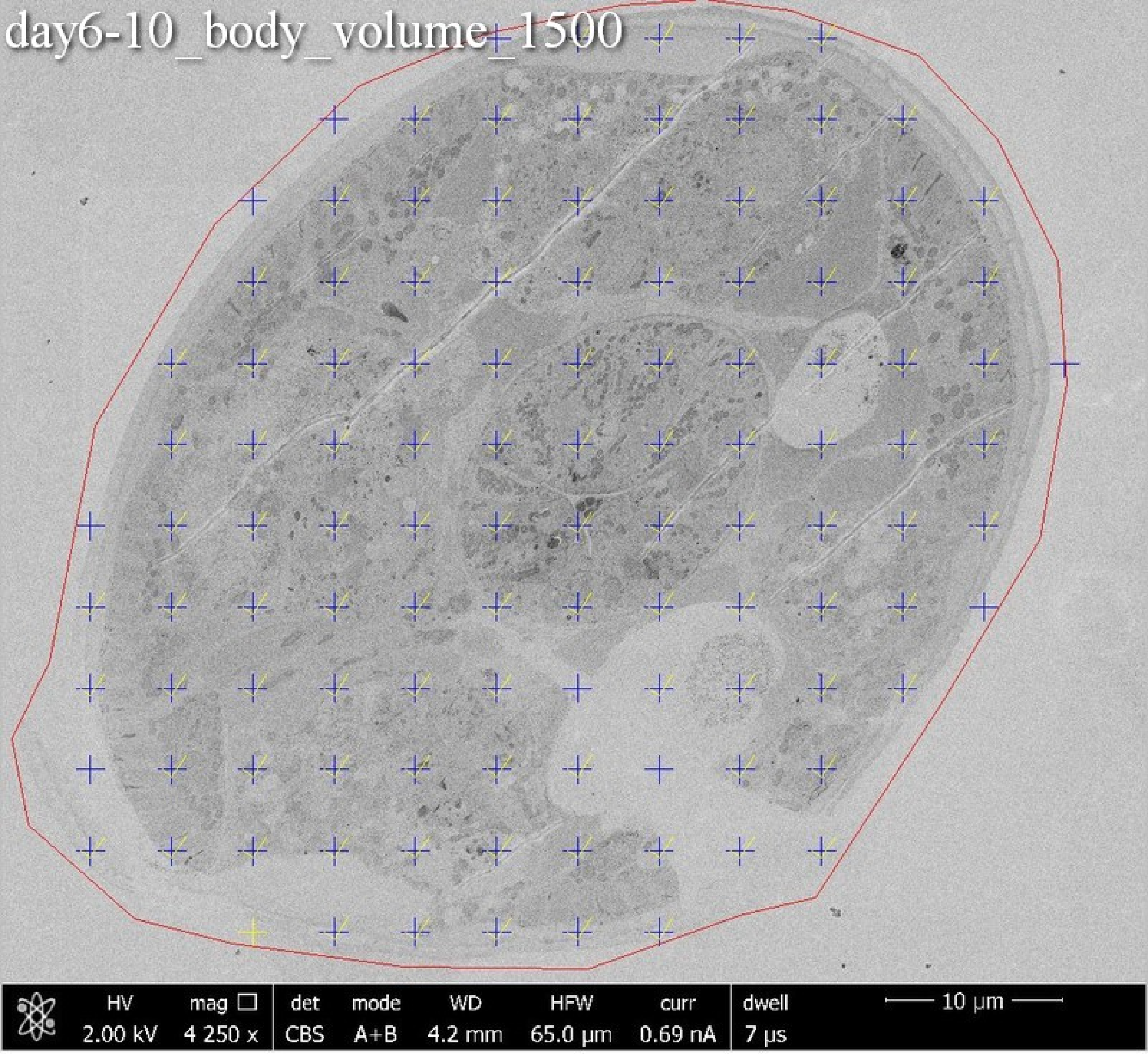
day6-10_body_volume_1500
HV 2.00 kV
mag ☐ 4 250 x
det CBS
mode A+B
WD 4.2 mm
HFW 65.0 µm
curr 0.69 nA
dwell 7 µs
10 µm

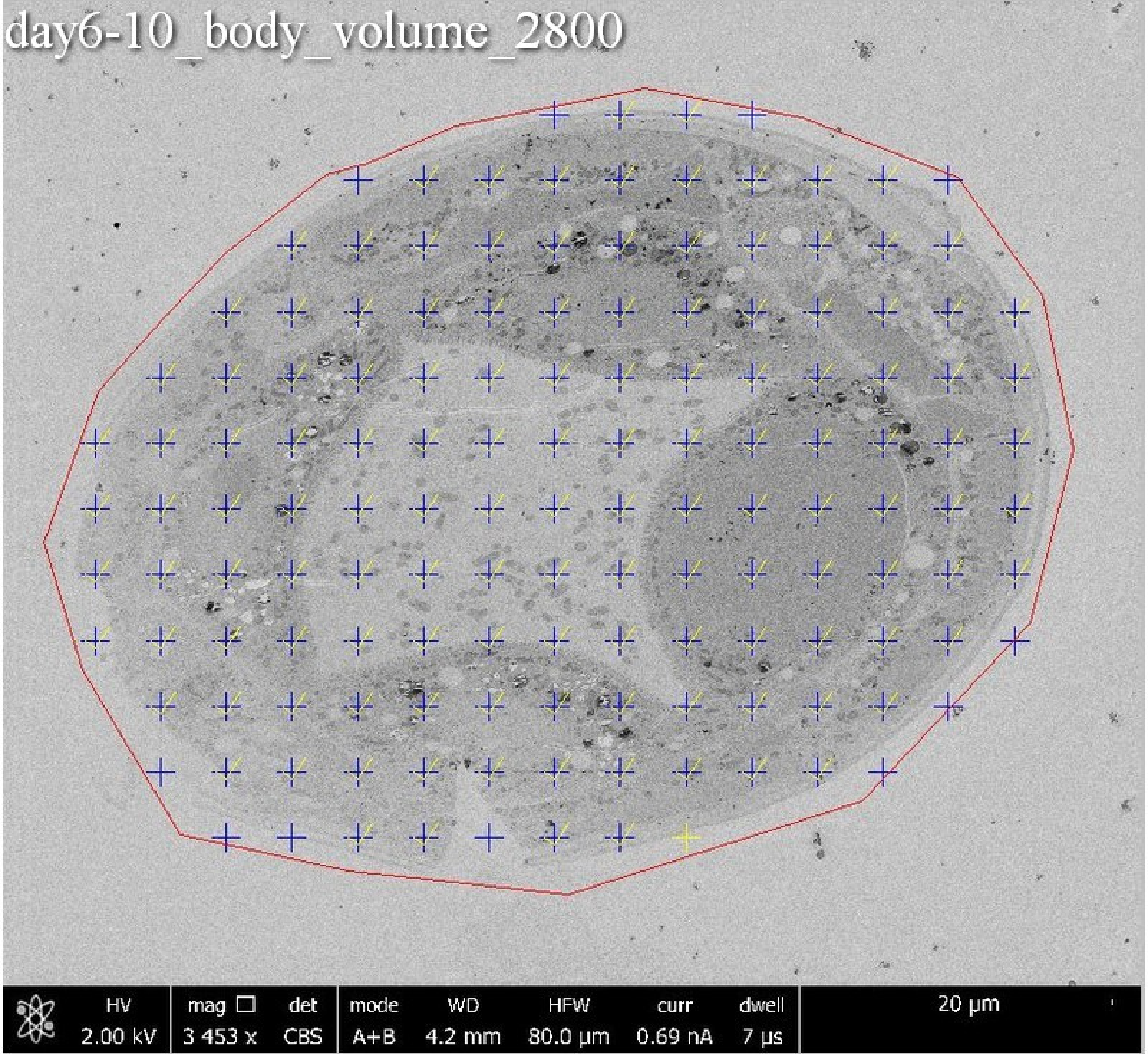
day6-10_body_volume_2800
HV
2.00 kV
mag
3 453 x
det
CBS
mode
A+B
WD
4.2 mm
HFW
80.0 μm
curr
0.69 nA
dwell
7 μs
20 μm

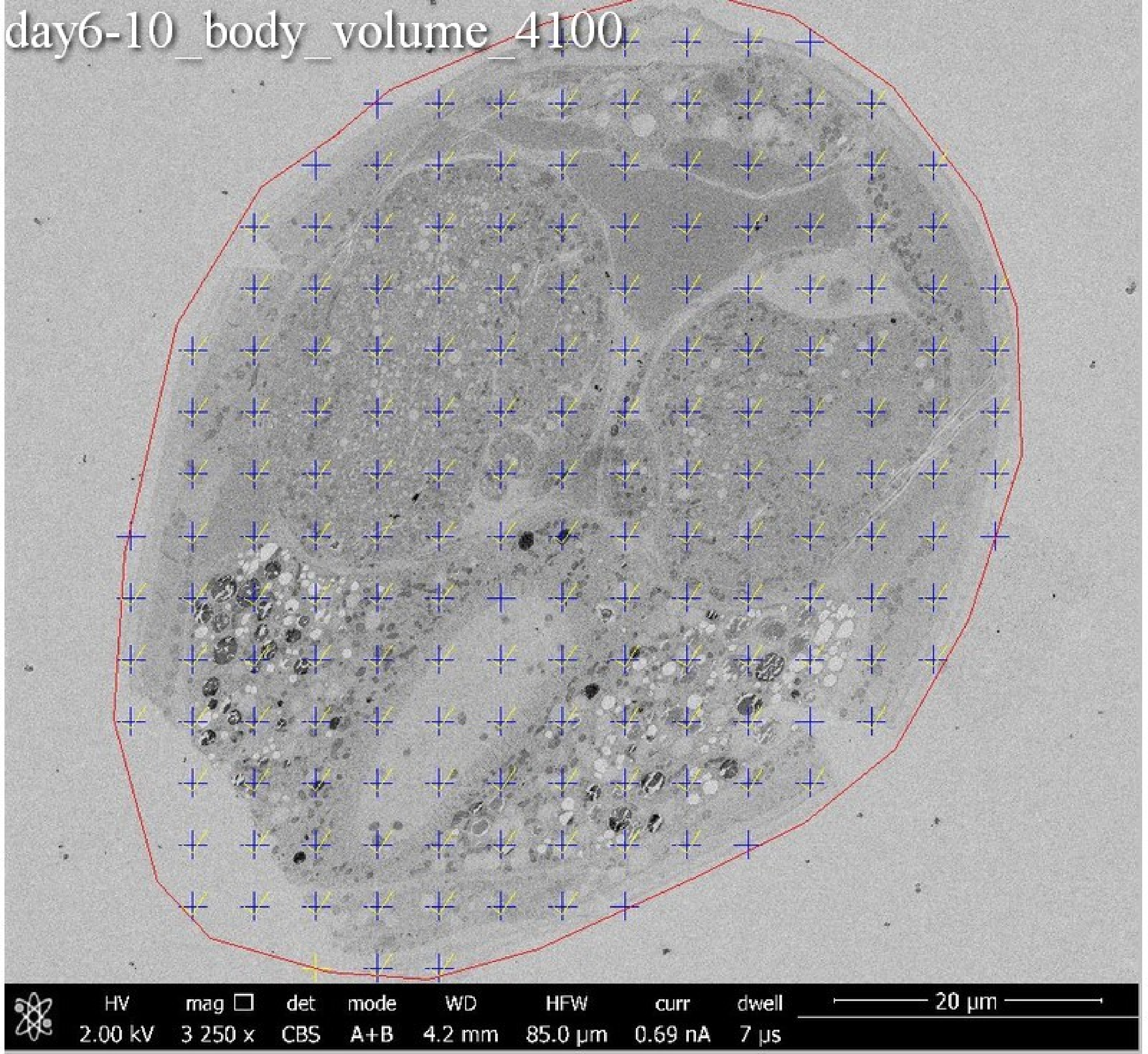

day6-10_body_volume_4100
HV 2.00 kV
mag □ 3 250 x
det CBS
mode A+B
WD 4.2 mm
HFW 85.0 µm
curr 0.69 nA
dwell 7 µs
20 µm

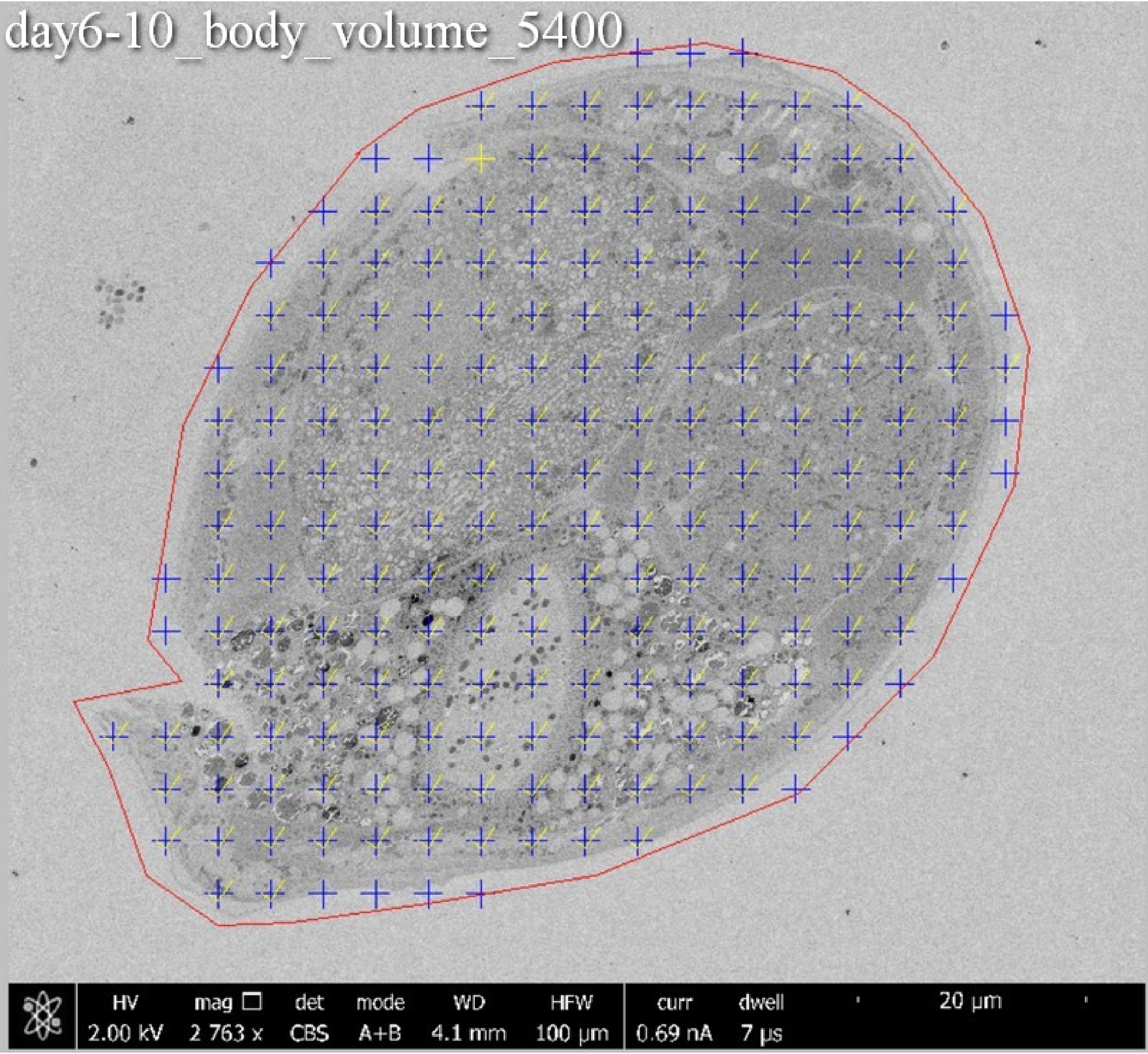

day6-10_body_volume_5400
HV 2.00 kV
mag □ 2 763 x
det CBS
mode A+B
WD 4.1 mm
HFW 100 μm
curr 0.69 nA
dwell 7 μs
20 μm

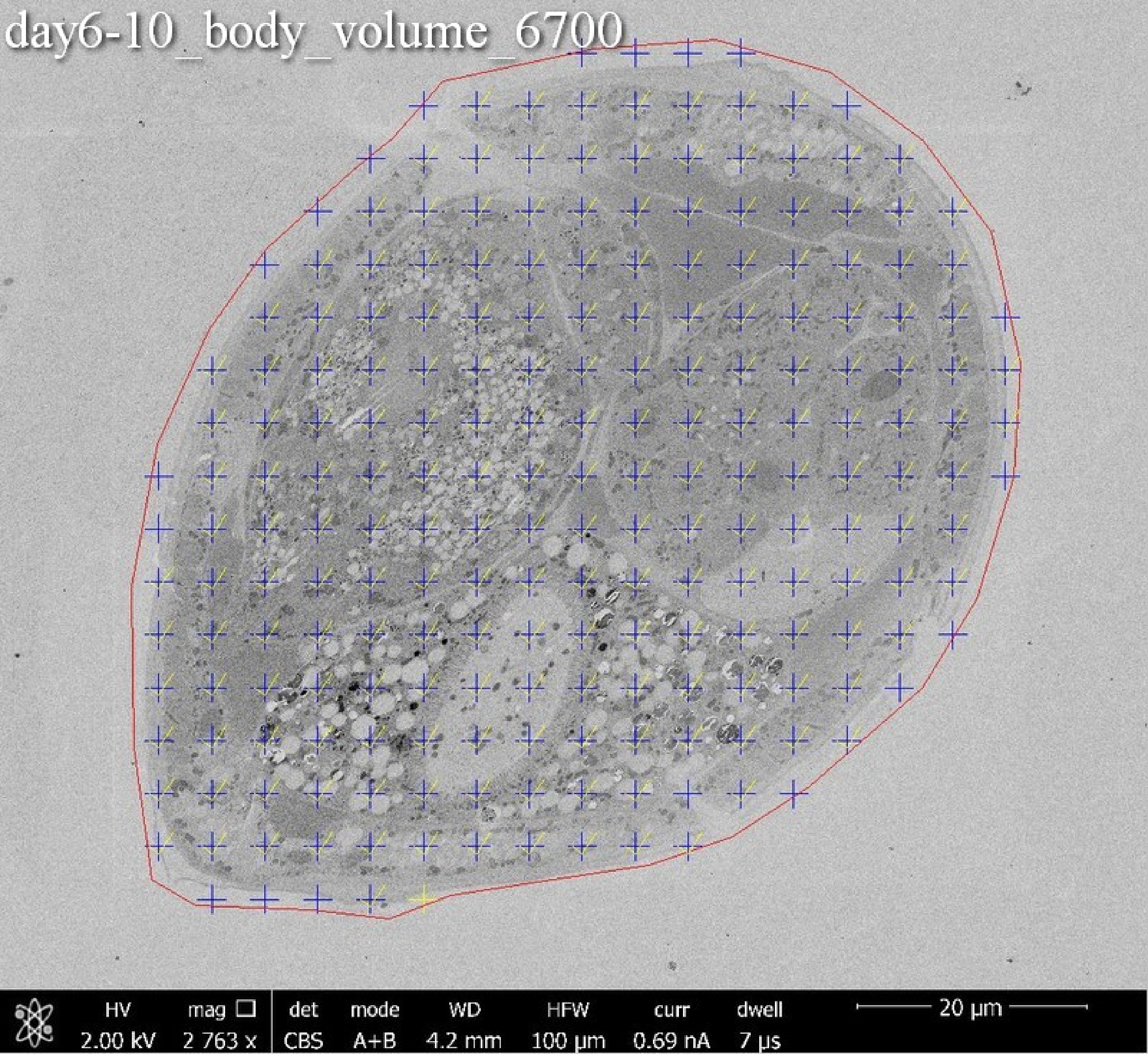
day6-10_body_volume_6700
HV
2.00 kV
mag
2 763 x
det
CBS
mode
A+B
WD
4.2 mm
HFW
100 µm
curr
0.69 nA
dwell
7 µs
20 µm

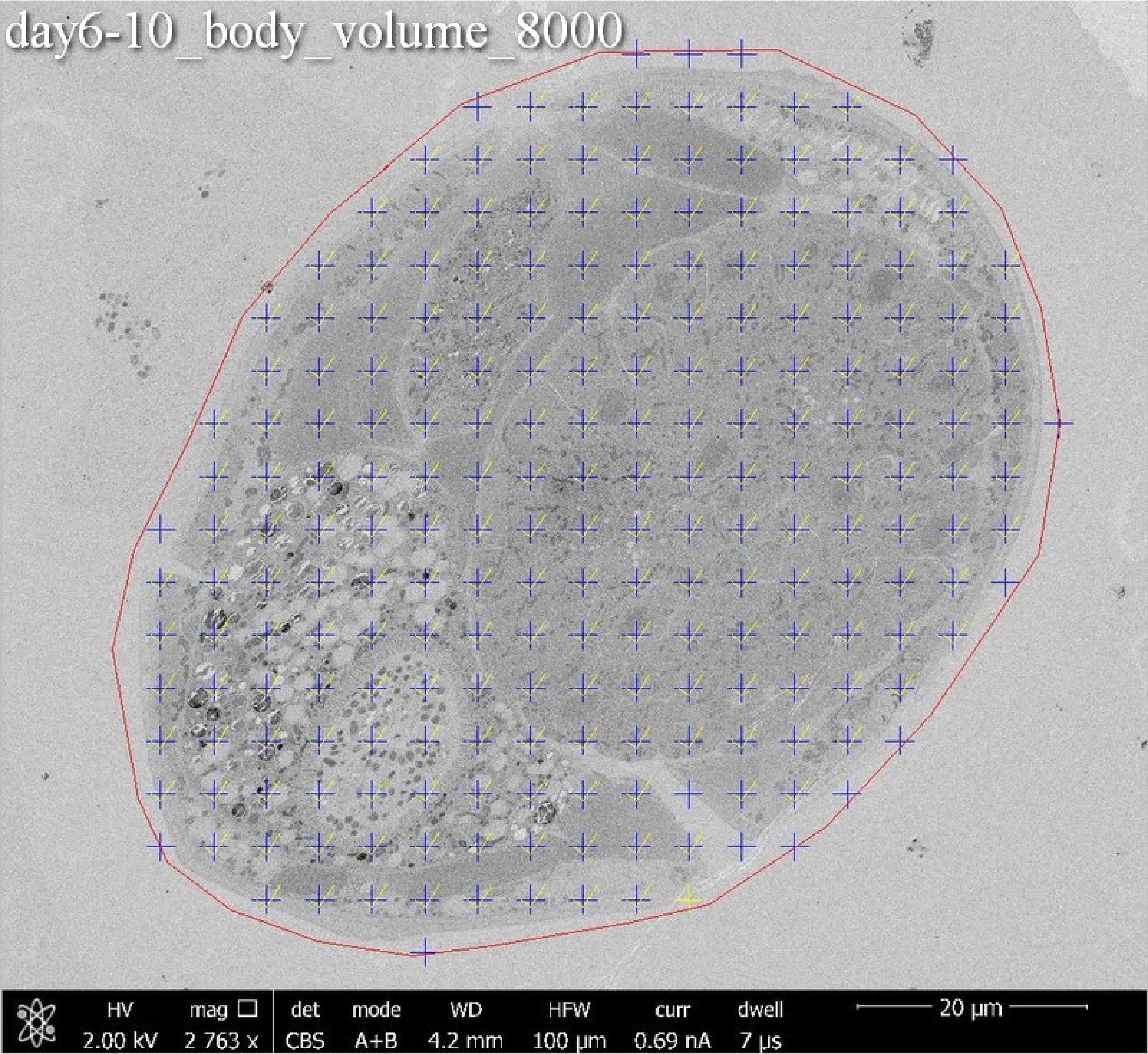
day6-10_body_volume_8000
HV 2.00 kV
mag 2 763 x
det CBS
mode A+B
WD 4.2 mm
HFW 100 µm
curr 0.69 nA
dwell 7 µs
20 µm

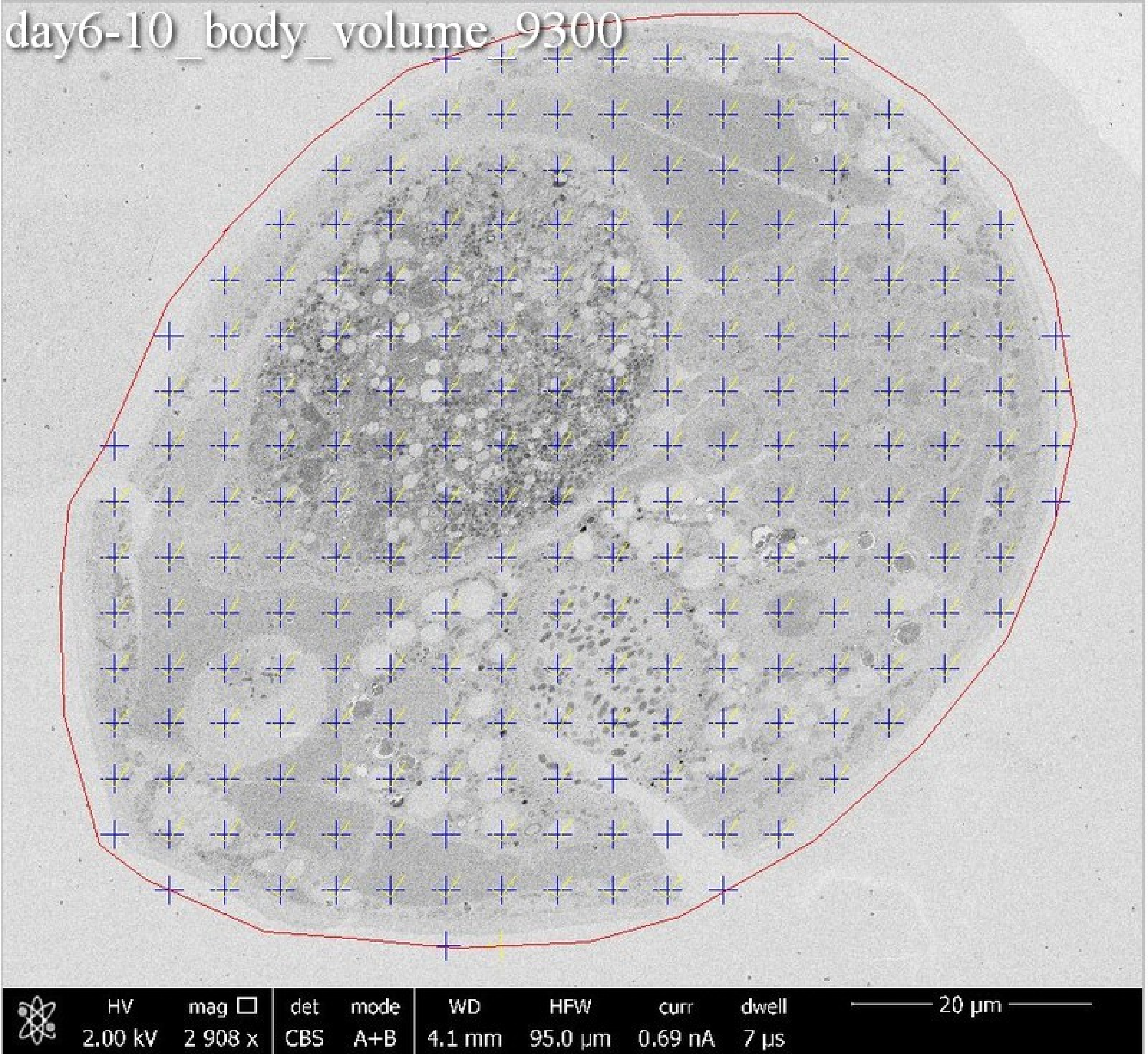
day6-10_body_volume_9300
HV 2.00 kV
mag 2 908 x
det CBS
mode A+B
WD 4.1 mm
HFW 95.0 μm
curr 0.69 nA
dwell 7 μs
20 μm

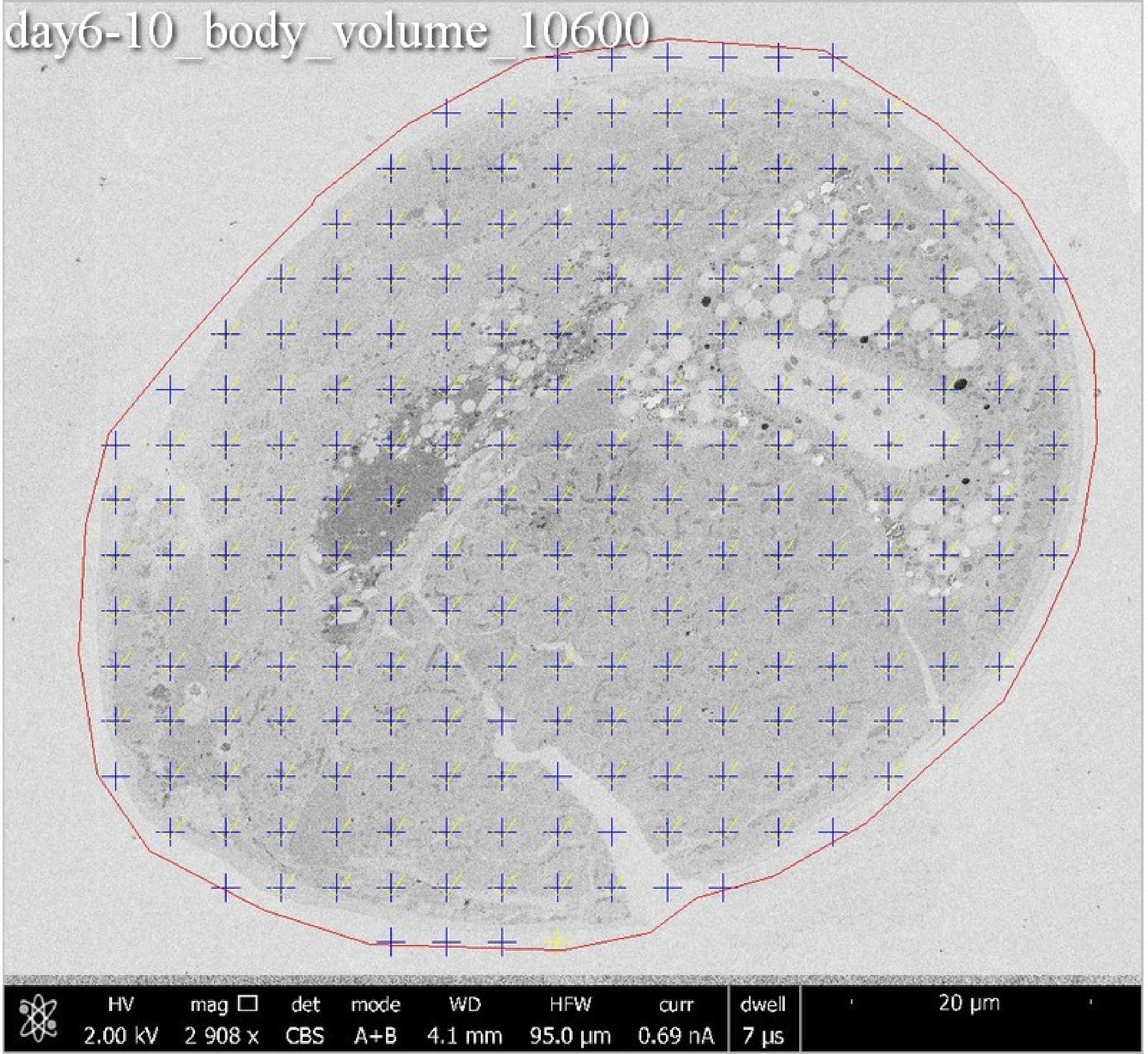

day6-10_body_volume_10600
HV
2.00 kV
mag
2 908 x
det
CBS
mode
A+B
WD
4.1 mm
HFW
95.0 μm
curr
0.69 nA
dwell
7 μs
20 μm

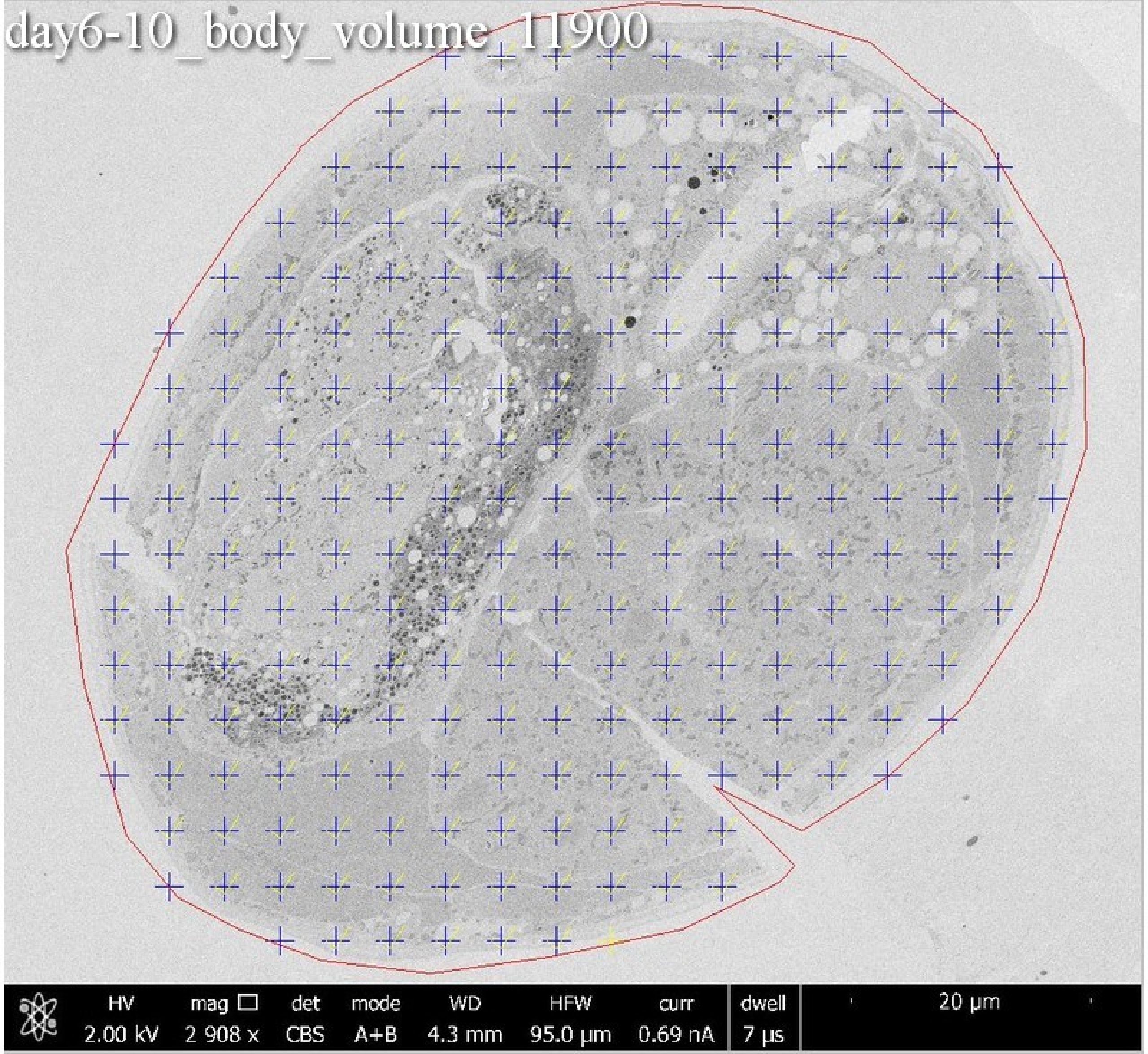
day6-10_body_volume_11900
HV 2.00 kV
mag ☐ 2 908 x
det CBS
mode A+B
WD 4.3 mm
HFW 95.0 µm
curr 0.69 nA
dwell 7 µs
20 µm

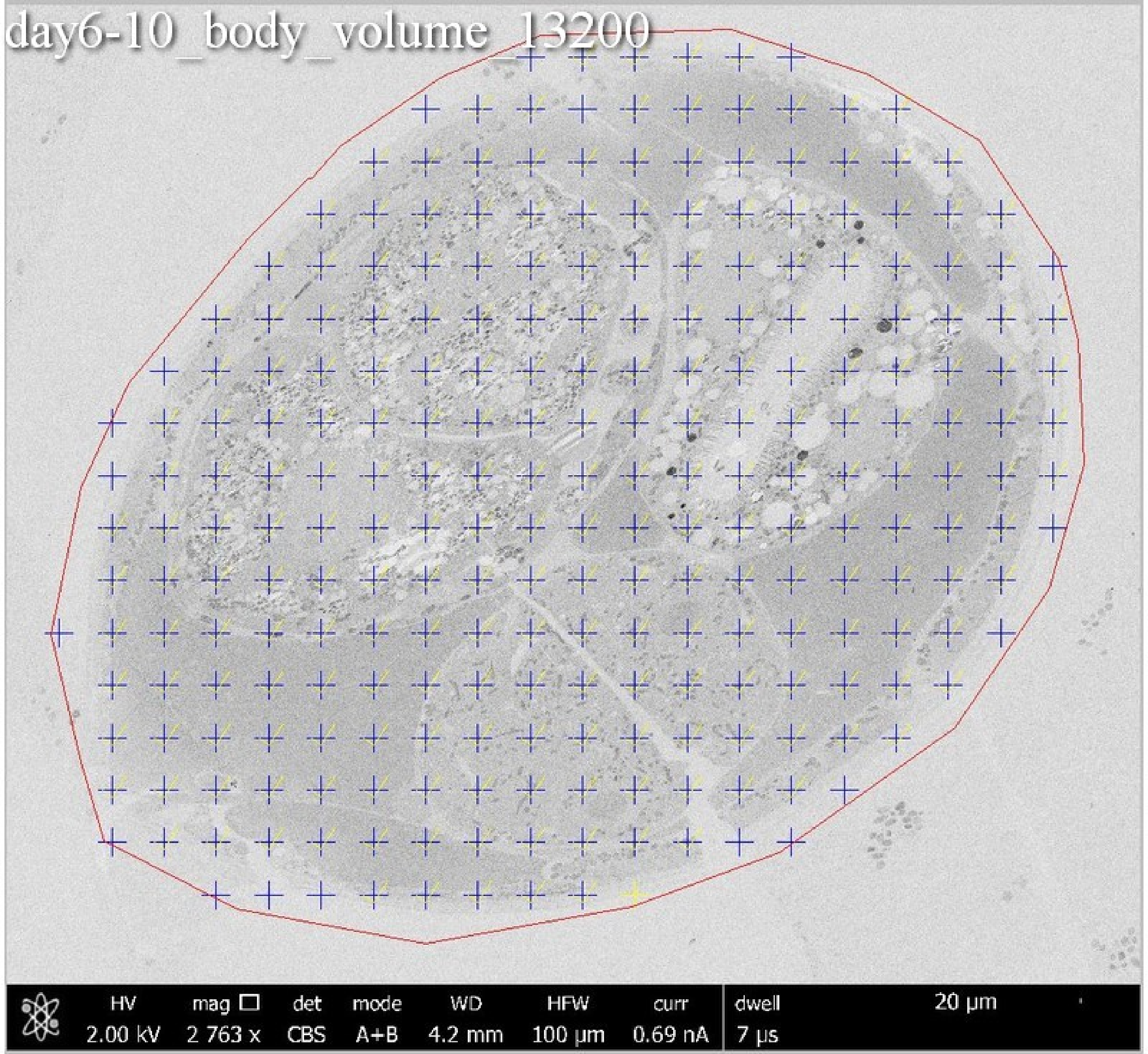
day6-10_body_volume_13200
HV 2.00 kV
mag □ 2 763 x
det CBS
mode A+B
WD 4.2 mm
HFW 100 µm
curr 0.69 nA
dwell 7 µs
20 µm

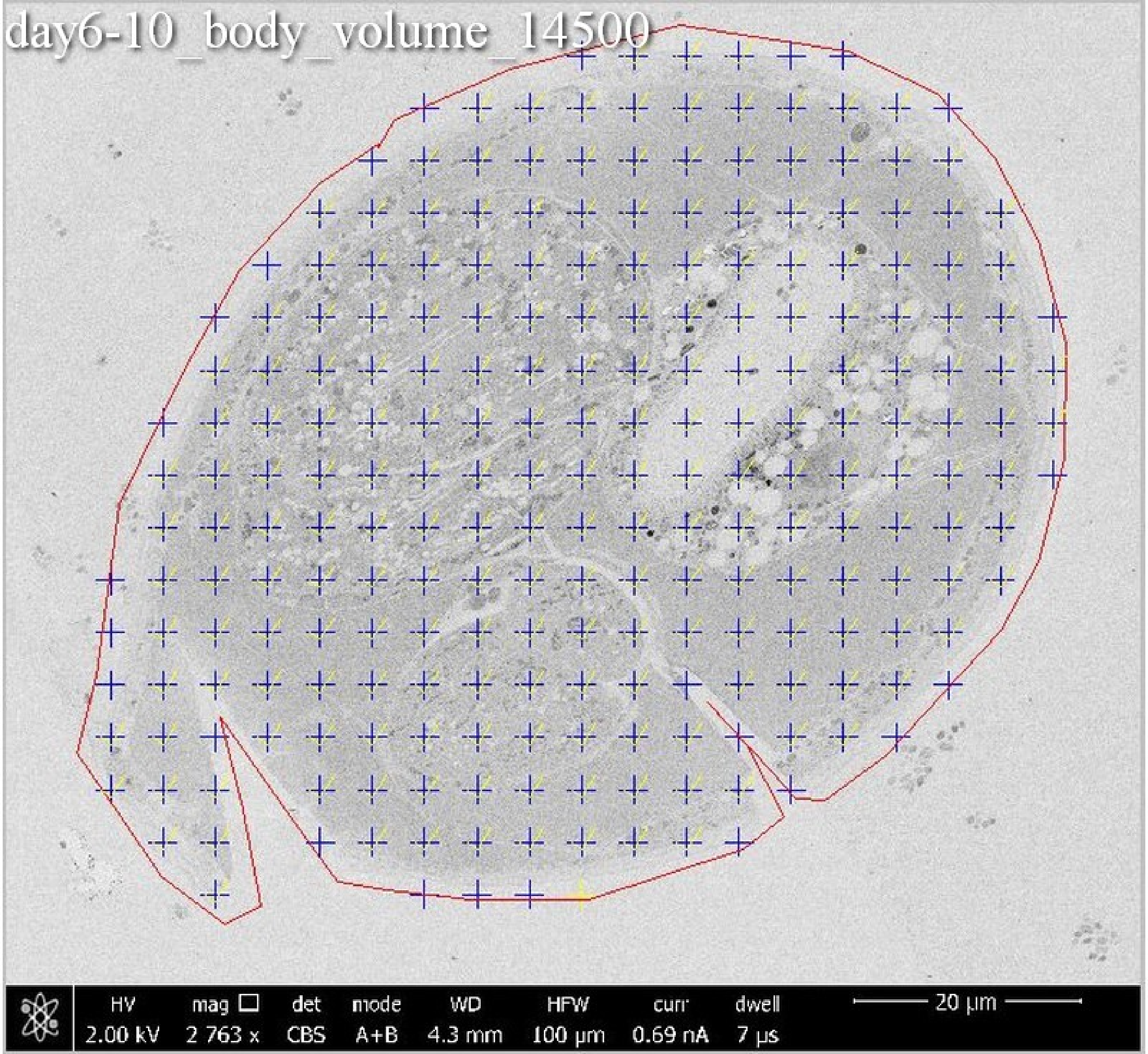
day6-10_body_volume_14500
HV 2.00 kV
mag 2 763 x
det CBS
mode A+B
WD 4.3 mm
HFW 100 μm
curr 0.69 nA
dwell 7 μs
20 μm

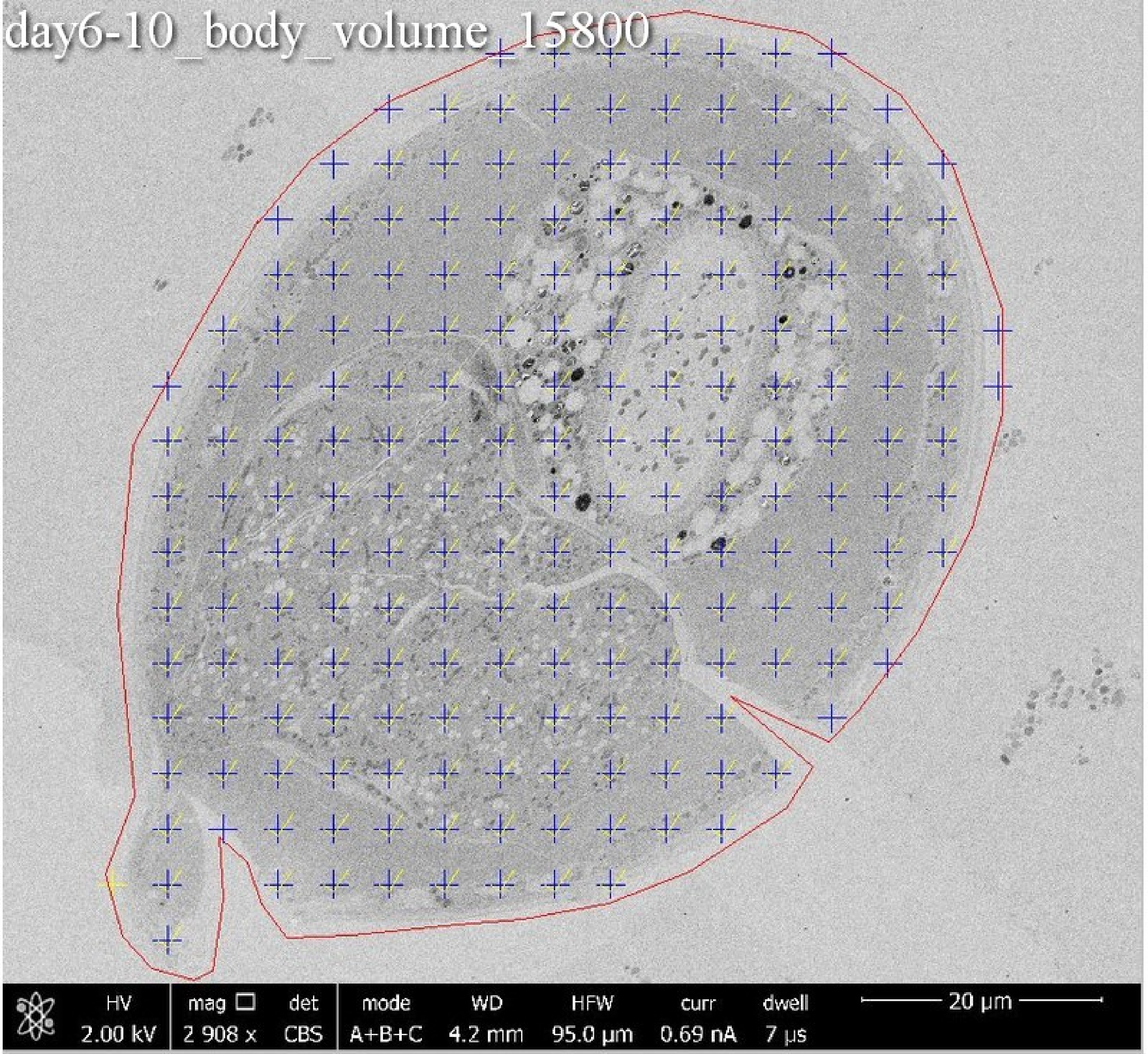
day6-10_body_volume_15800
HV 2.00 kV
mag 2 908 x
det CBS
mode A+B+C
WD 4.2 mm
HFW 95.0 µm
curr 0.69 nA
dwell 7 µs
20 µm

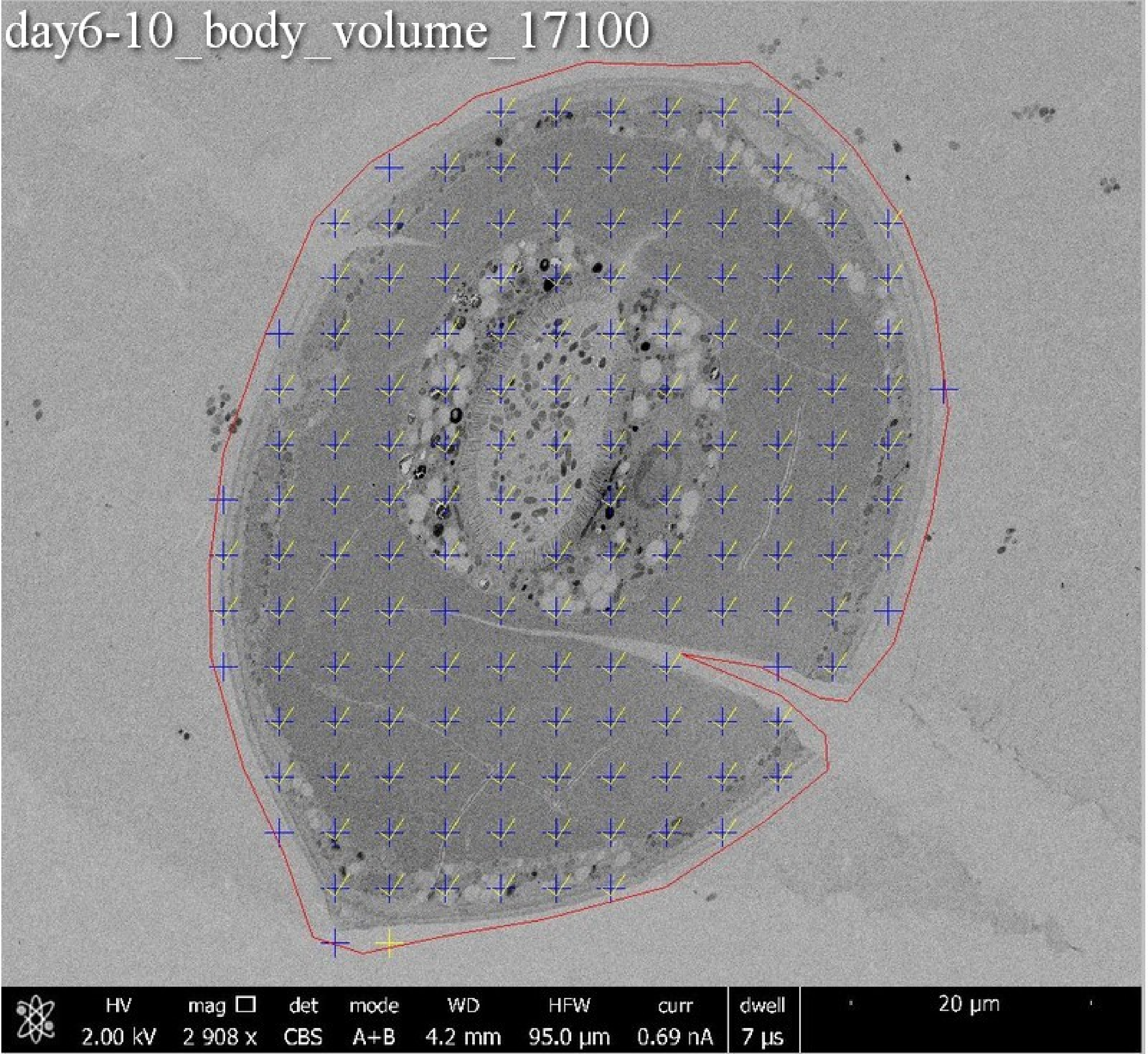

day6-10_body_volume_17100
HV 2.00 kV
mag □ 2 908 x
det CBS
mode A+B
WD 4.2 mm
HFW 95.0 µm
curr 0.69 nA
dwell 7 µs
20 µm

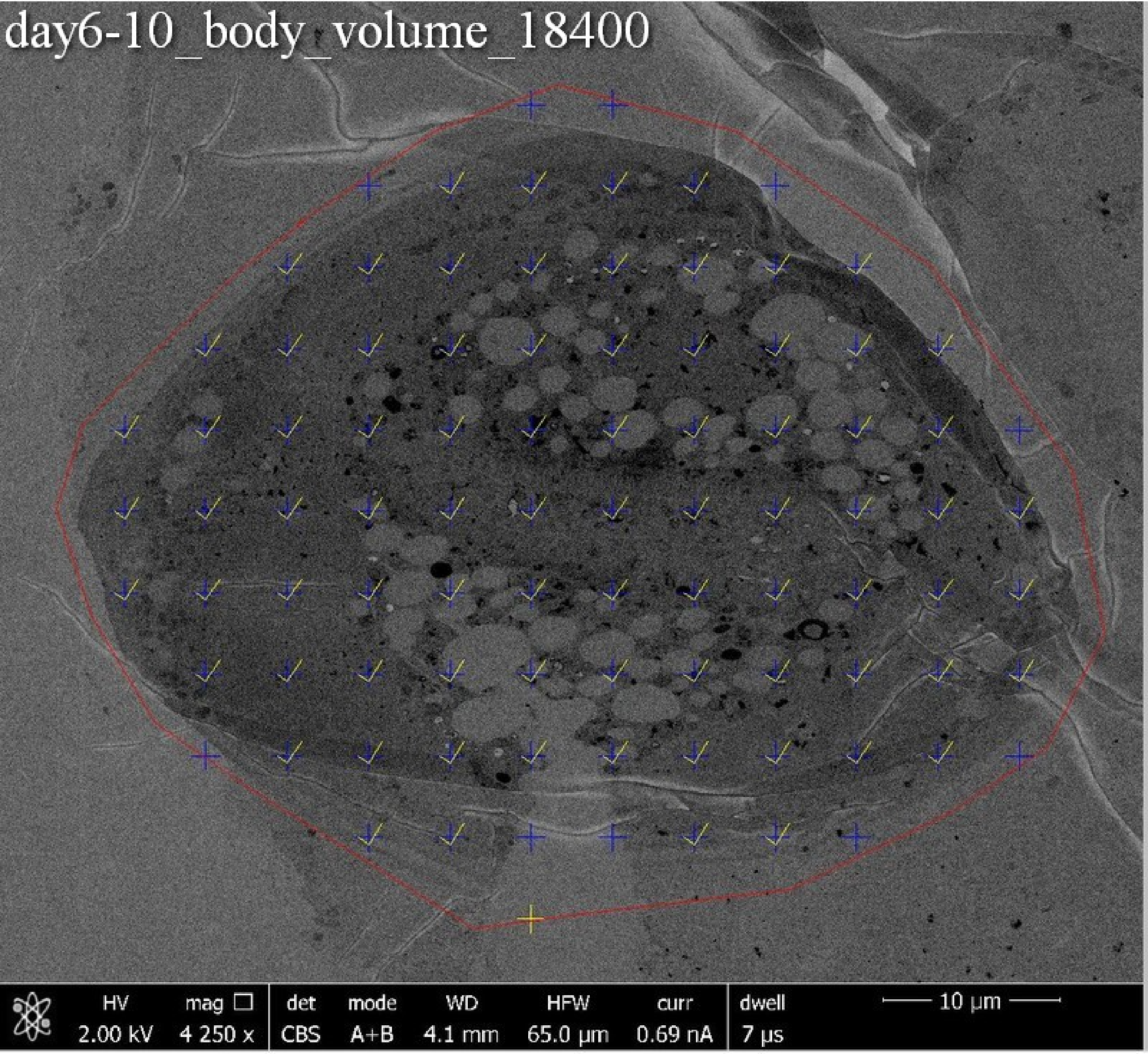
day6-10_body_volume_18400
HV 2.00 kV
mag 4 250 x
det CBS
mode A+B
WD 4.1 mm
HFW 65.0 µm
curr 0.69 nA
dwell 7 µs
10 µm

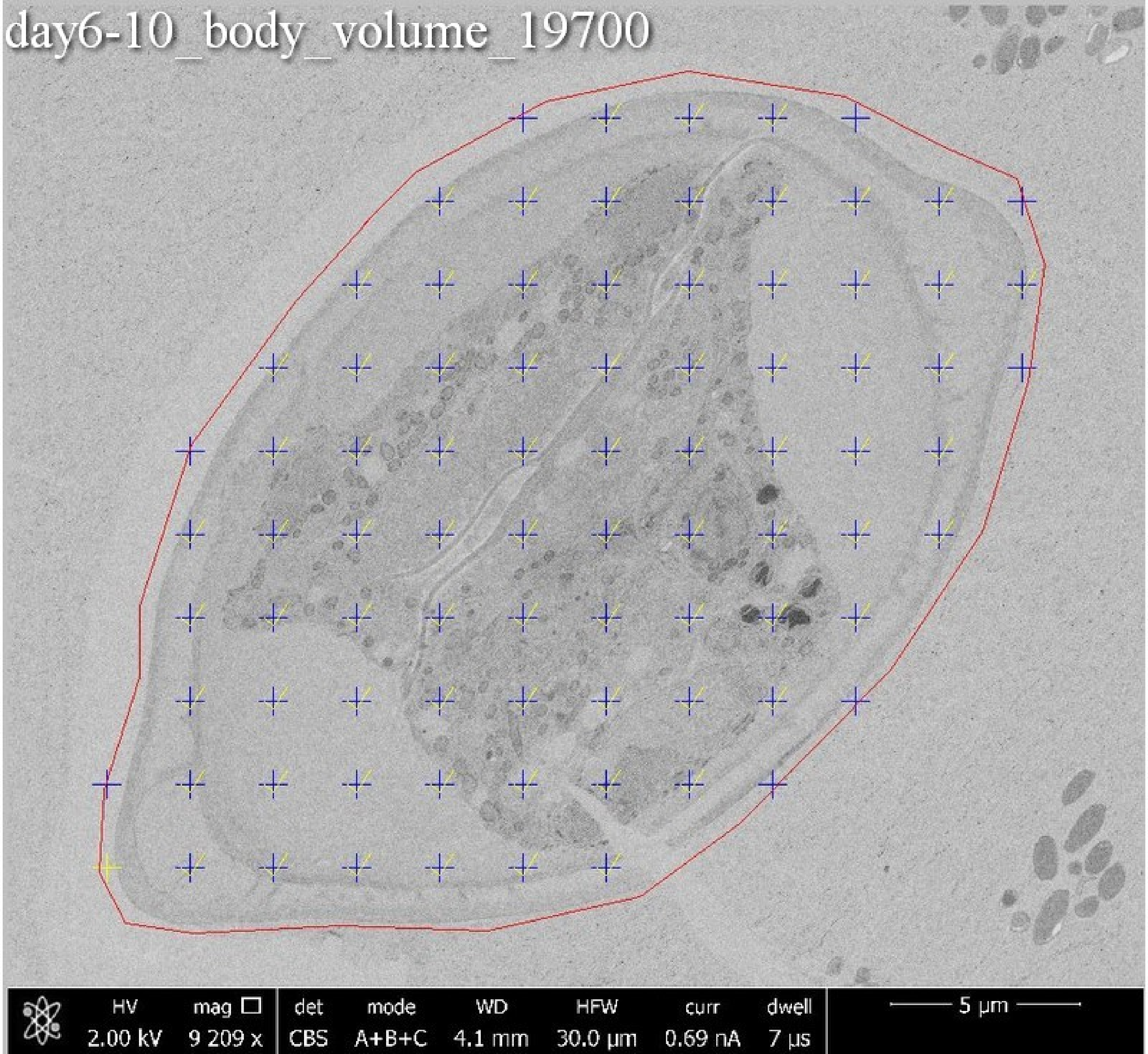
day6-10_body_volume_19700
HV
2.00 kV
mag
9 209 x
det
CBS
mode
A+B+C
WD
4.1 mm
HFW
30.0 µm
curr
0.69 nA
dwell
7 µs
5 µm

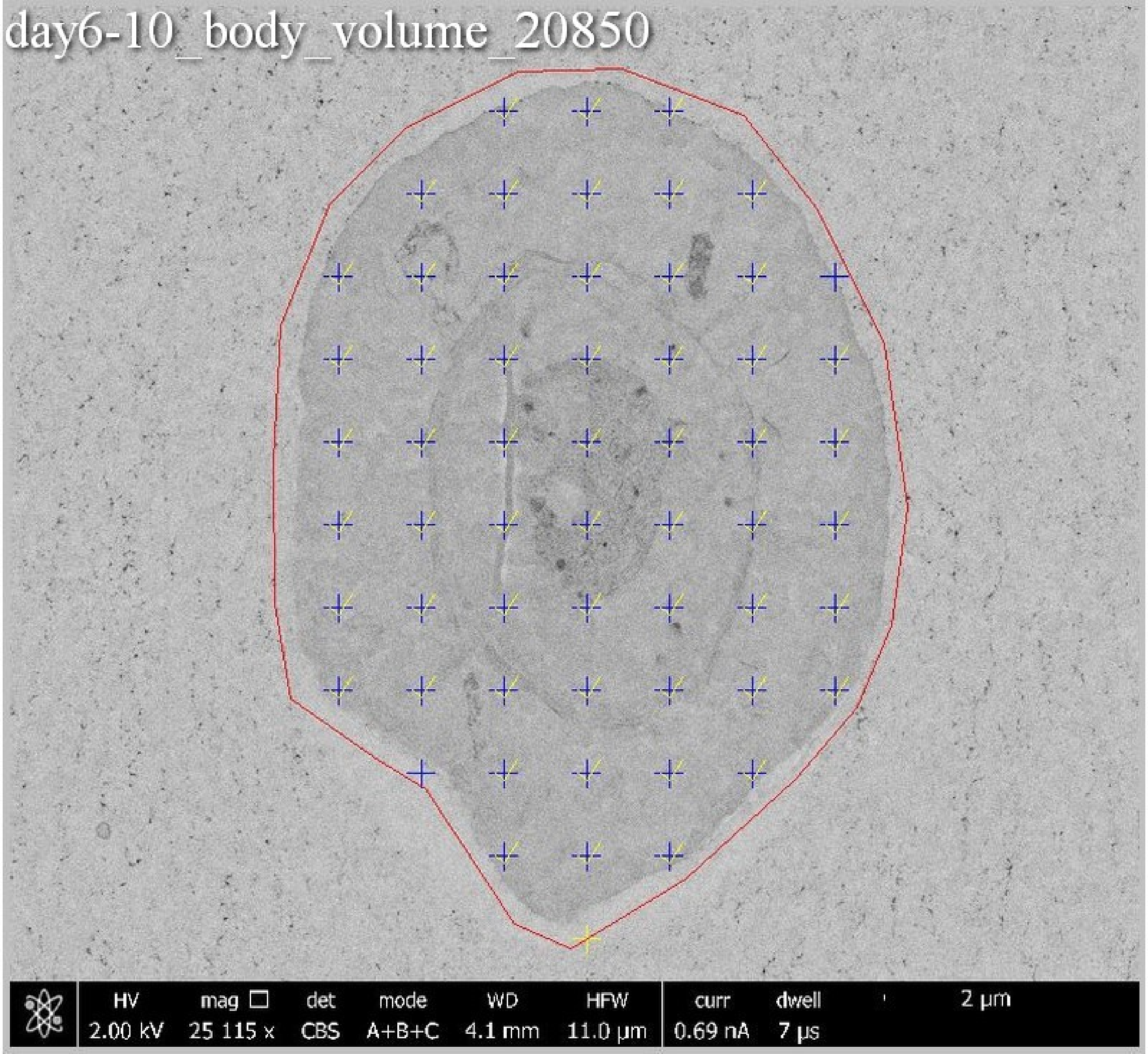
day6-10_body_volume_20850
HV 2.00 kV
mag □ 25 115 x
det CBS
mode A+B+C
WD 4.1 mm
HFW 11.0 µm
curr 0.69 nA
dwell 7 µs
2 µm

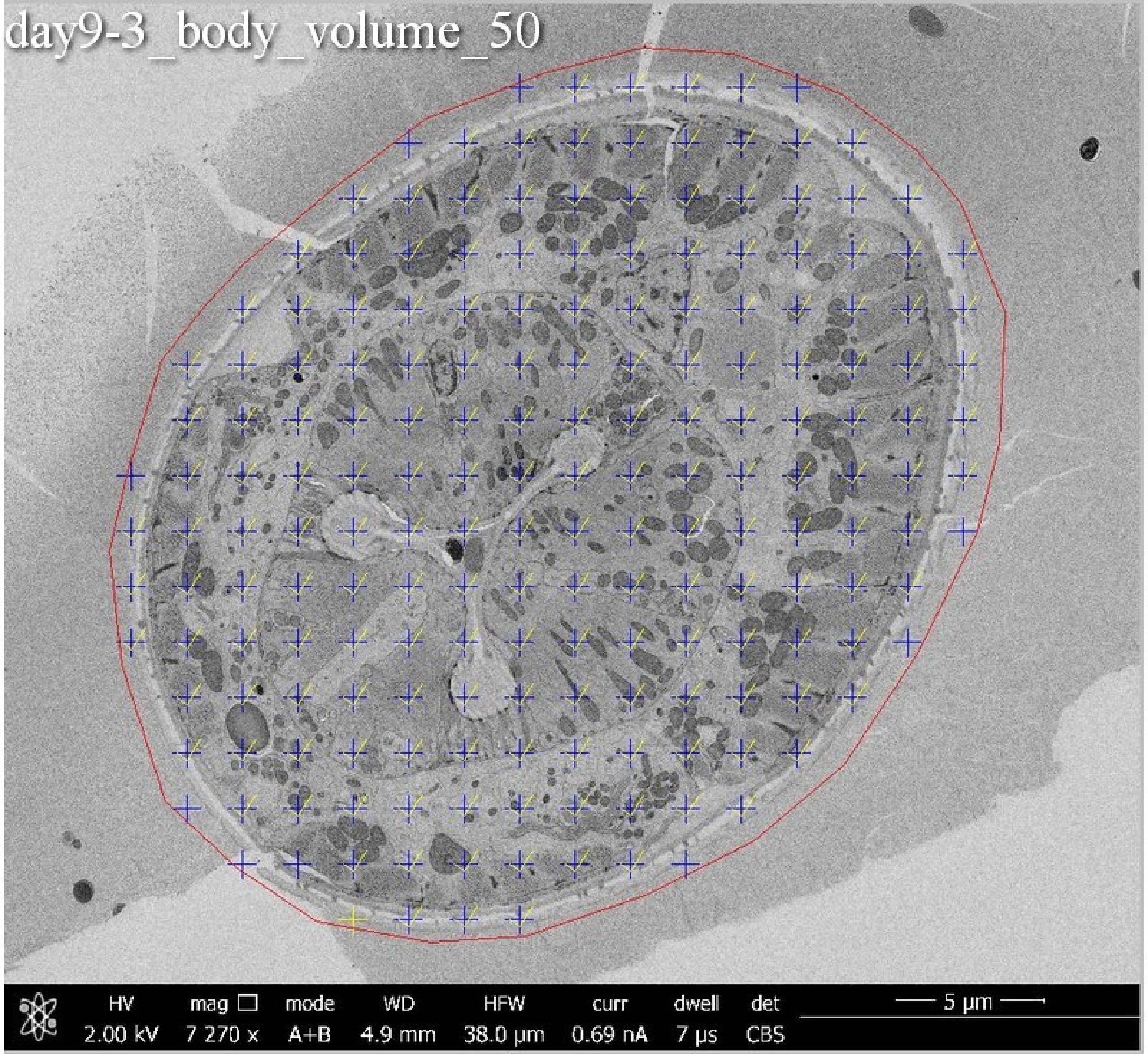
day9-3_body_volume_50
HV
2.00 kV
mag ☐
7 270 x
mode
A+B
WD
4.9 mm
HFW
38.0 μm
curr
0.69 nA
dwell
7 μs
det
CBS
5 μm

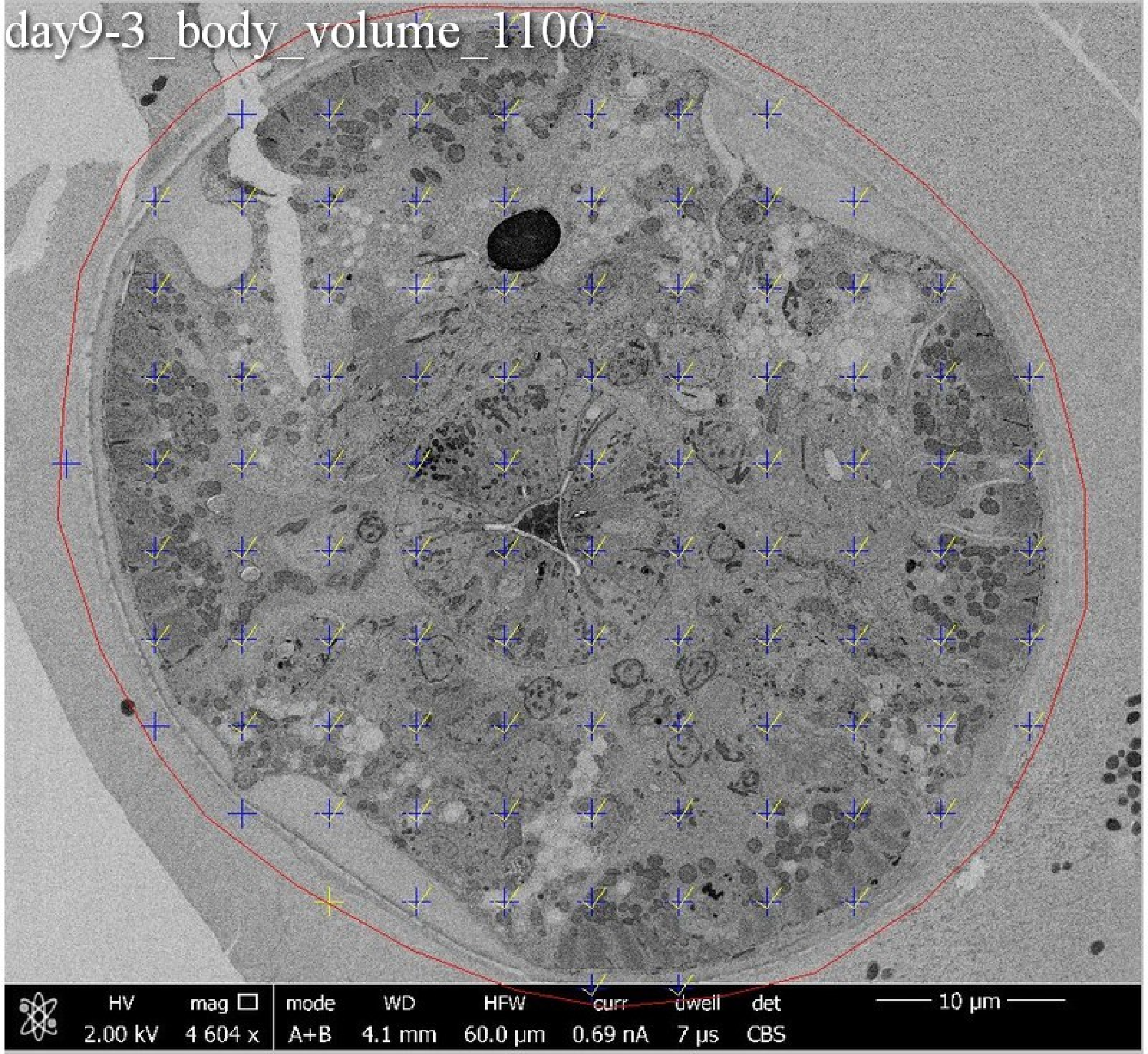

day9-3_body_volume_1100
HV 2.00 kV
mag □ 4 604 x
mode A+B
WD 4.1 mm
HFW 60.0 µm
curr 0.69 nA
dwell 7 µs
det CBS
10 µm

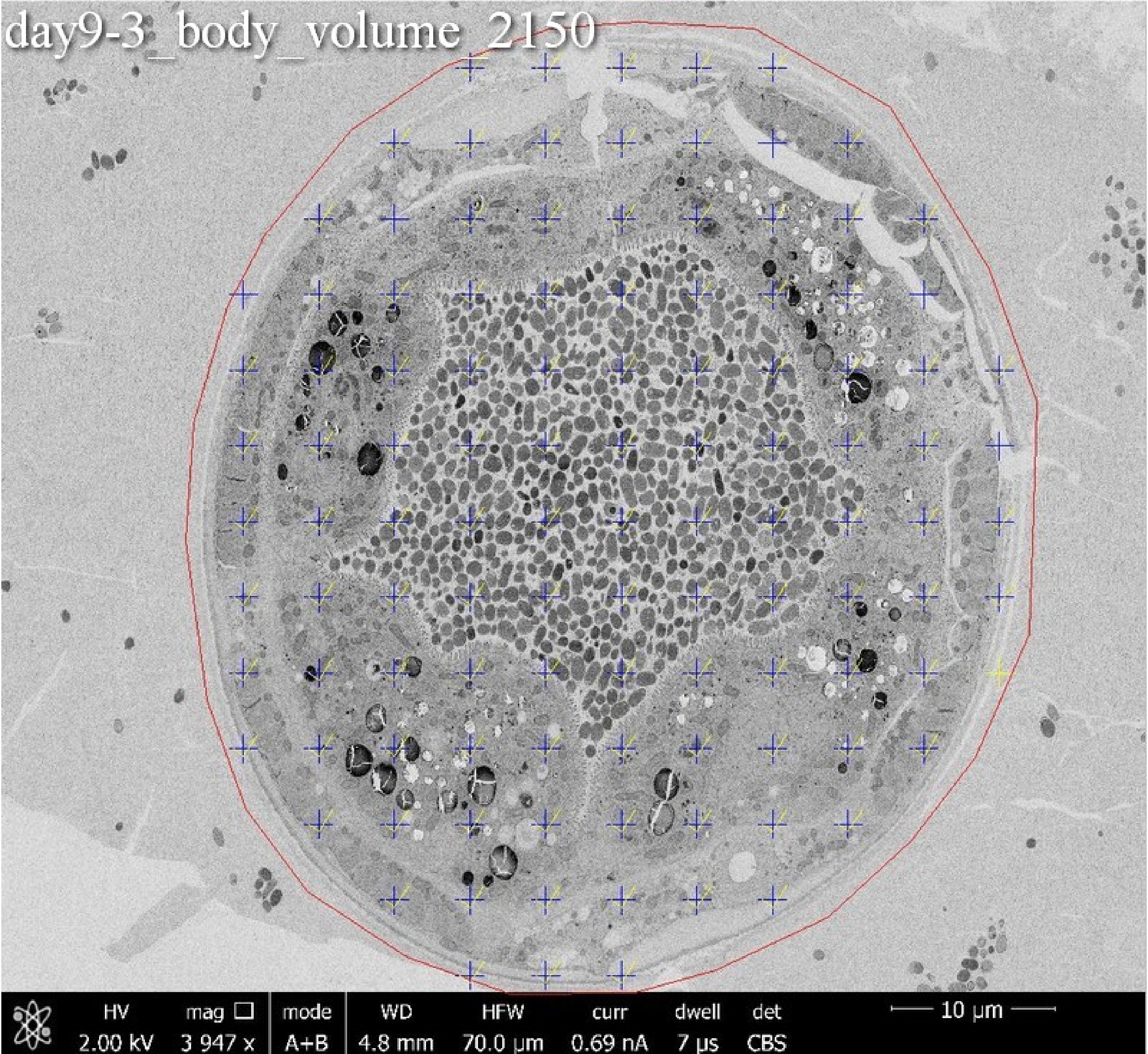
day9-3_body_volume_2150
HV
2.00 kV
mag
3 947 x
mode
A+B
WD
4.8 mm
HFW
70.0 μm
curr
0.69 nA
dwell
7 μs
det
CBS
10 μm

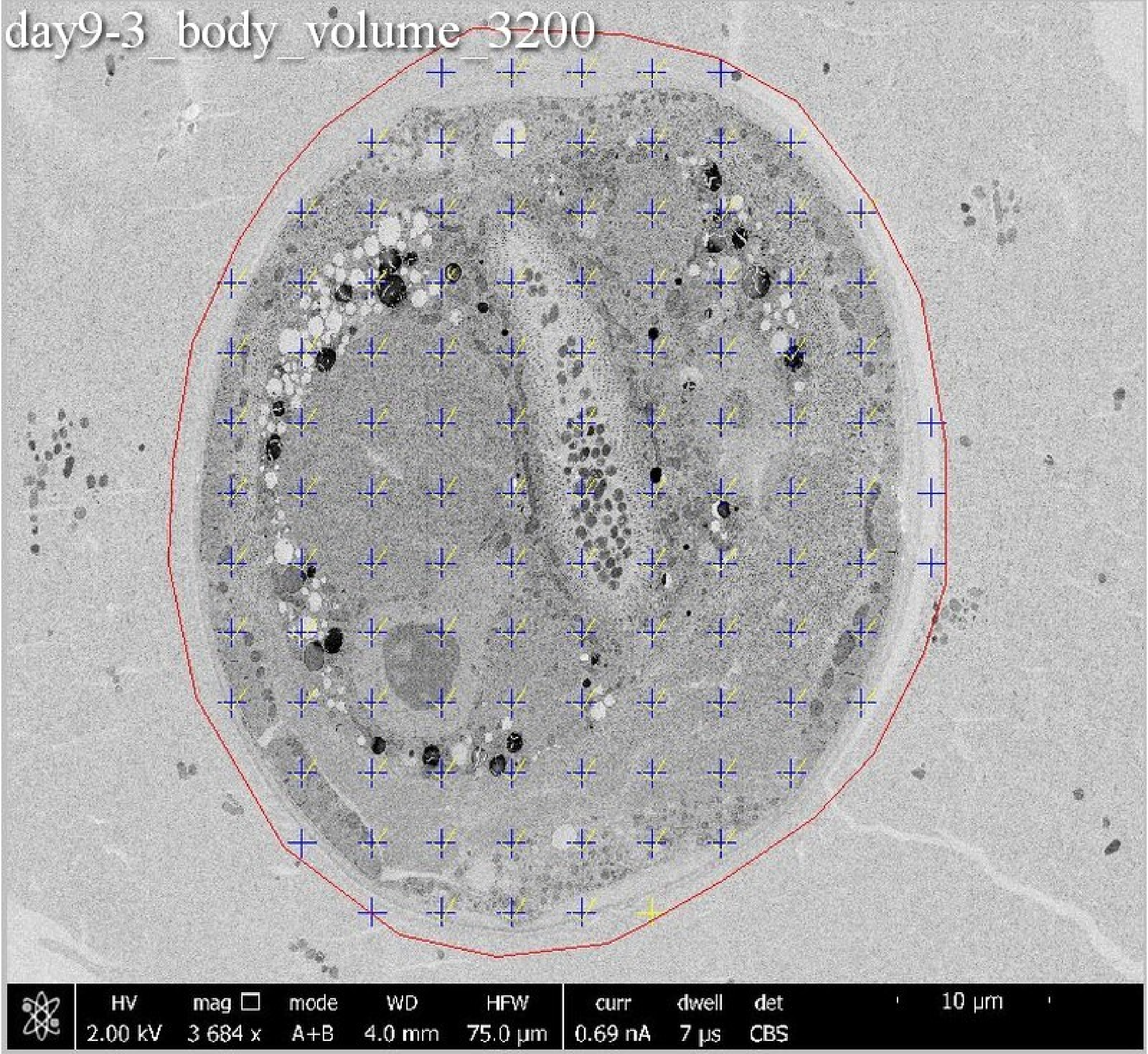
day9-3_body_volume_3200
HV
2.00 kV
mag
3 684 x
mode
A+B
WD
4.0 mm
HFW
75.0 μm
curr
0.69 nA
dwell
7 μs
det
CBS
10 μm

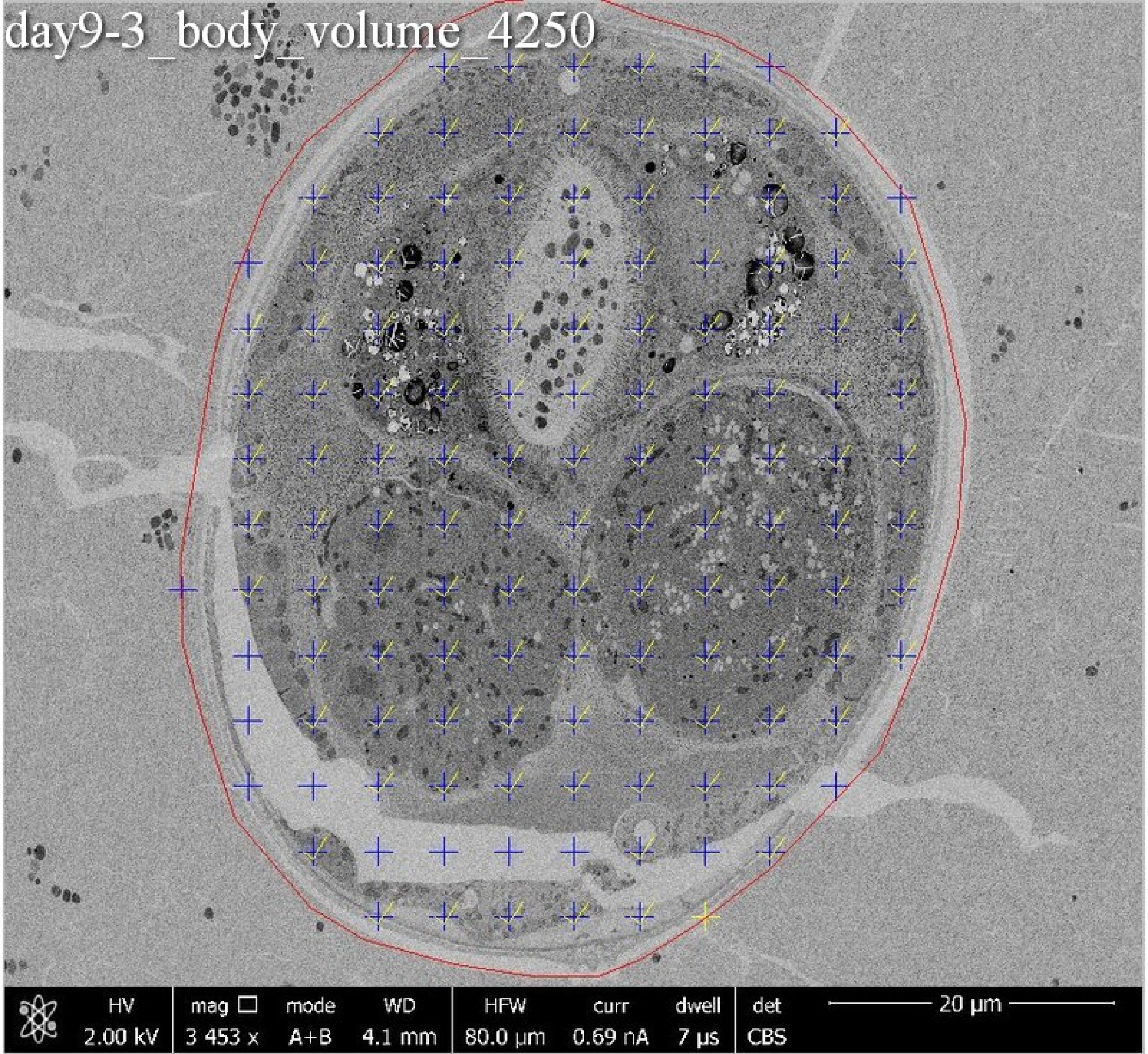
day9-3_body_volume_4250
HV 2.00 kV
mag □ 3 453 x
mode A+B
WD 4.1 mm
HFW 80.0 µm
curr 0.69 nA
dwell 7 µs
det CBS
20 µm

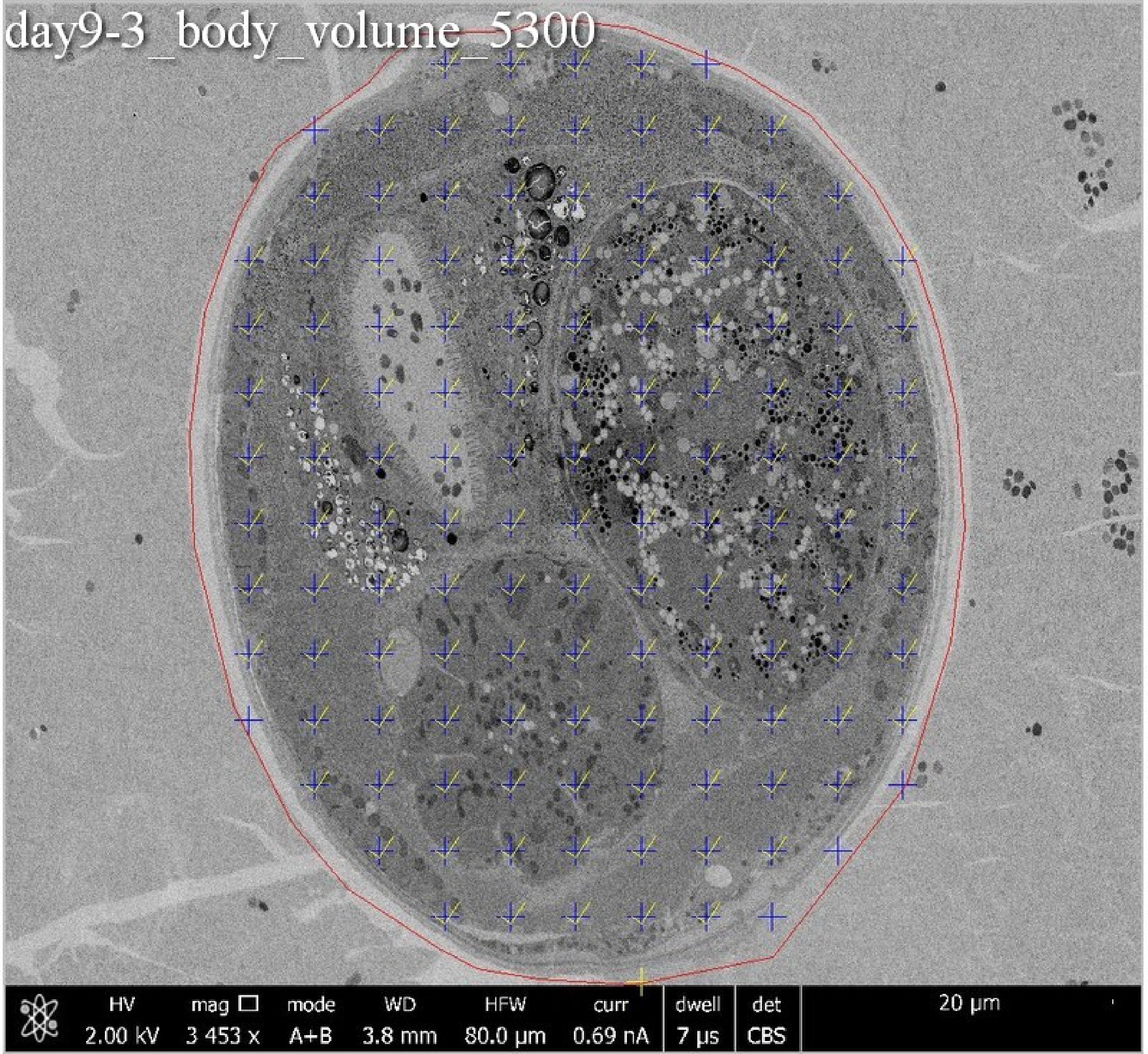
day9-3_body_volume_5300
HV
2.00 kV
mag
3 453 x
mode
A+B
WD
3.8 mm
HFW
80.0 µm
curr
0.69 nA
dwell
7 µs
det
CBS
20 µm

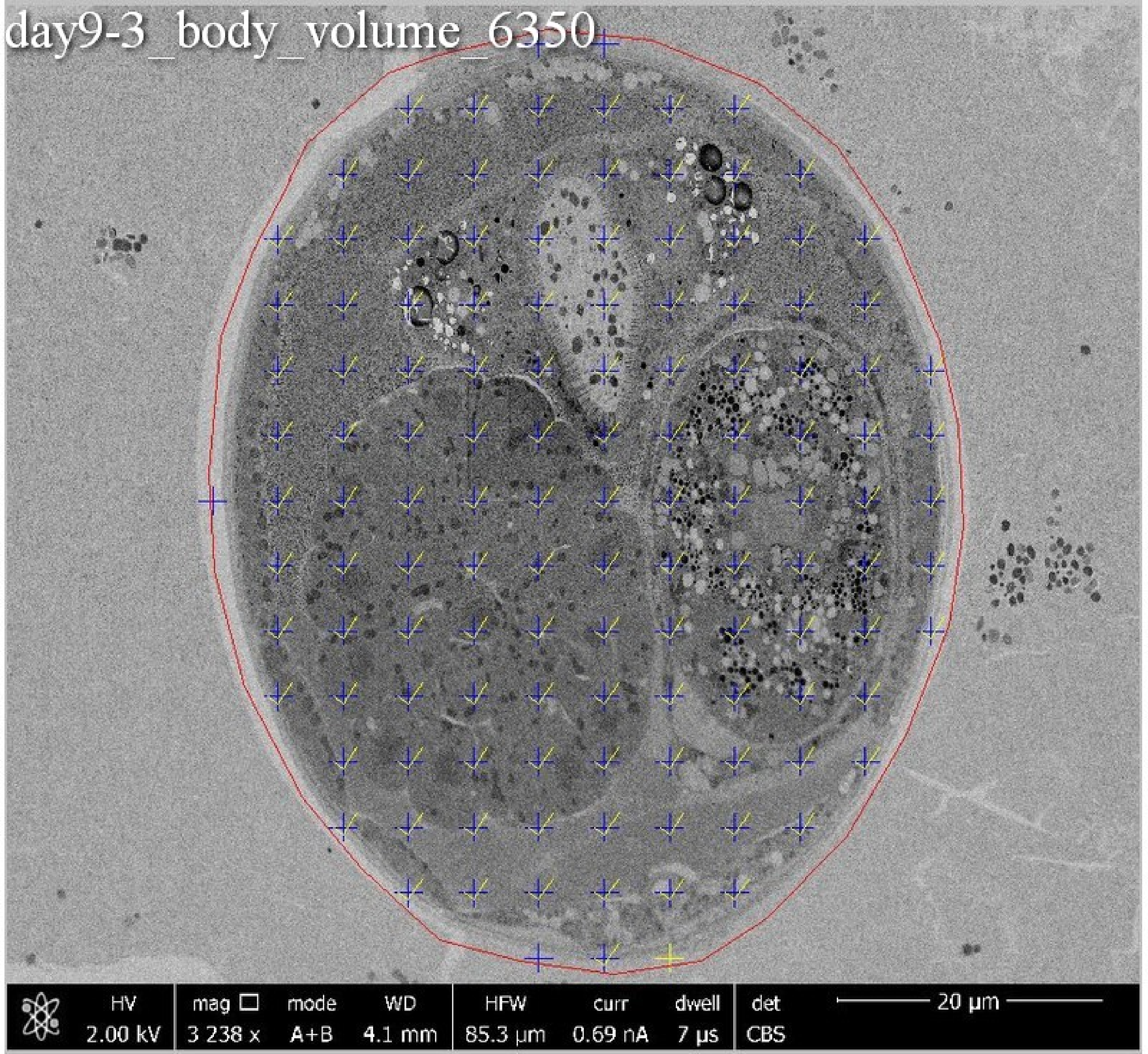
day9-3_body_volume_6350
HV
2.00 kV
mag □
3 238 x
mode
A+B
WD
4.1 mm
HFW
85.3 µm
curr
0.69 nA
dwell
7 µs
det
CBS
20 µm

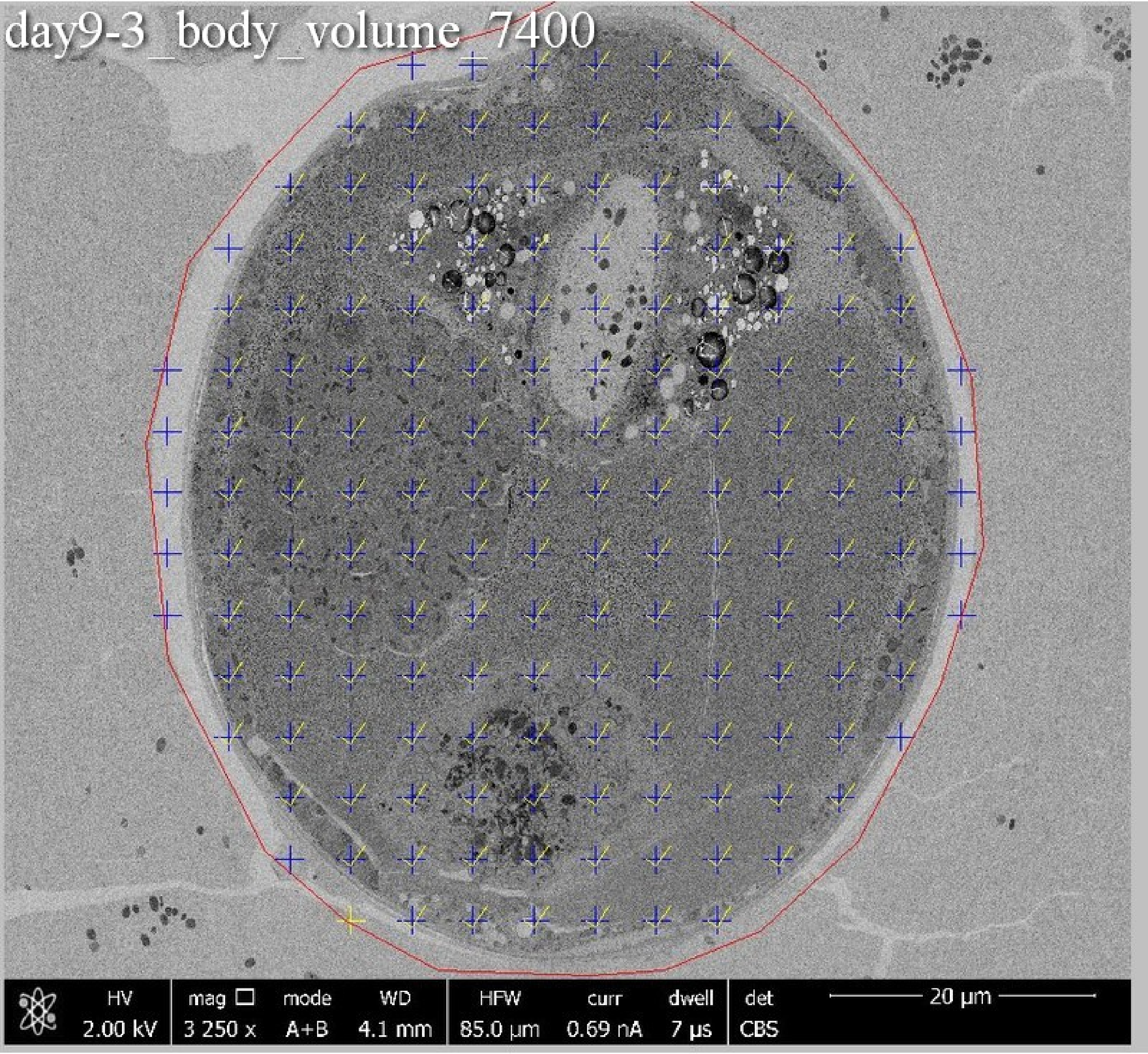
day9-3_body_volume_7400
HV 2.00 kV
mag 3 250 x
mode A+B
WD 4.1 mm
HFW 85.0 µm
curr 0.69 nA
dwell 7 µs
det CBS
20 µm

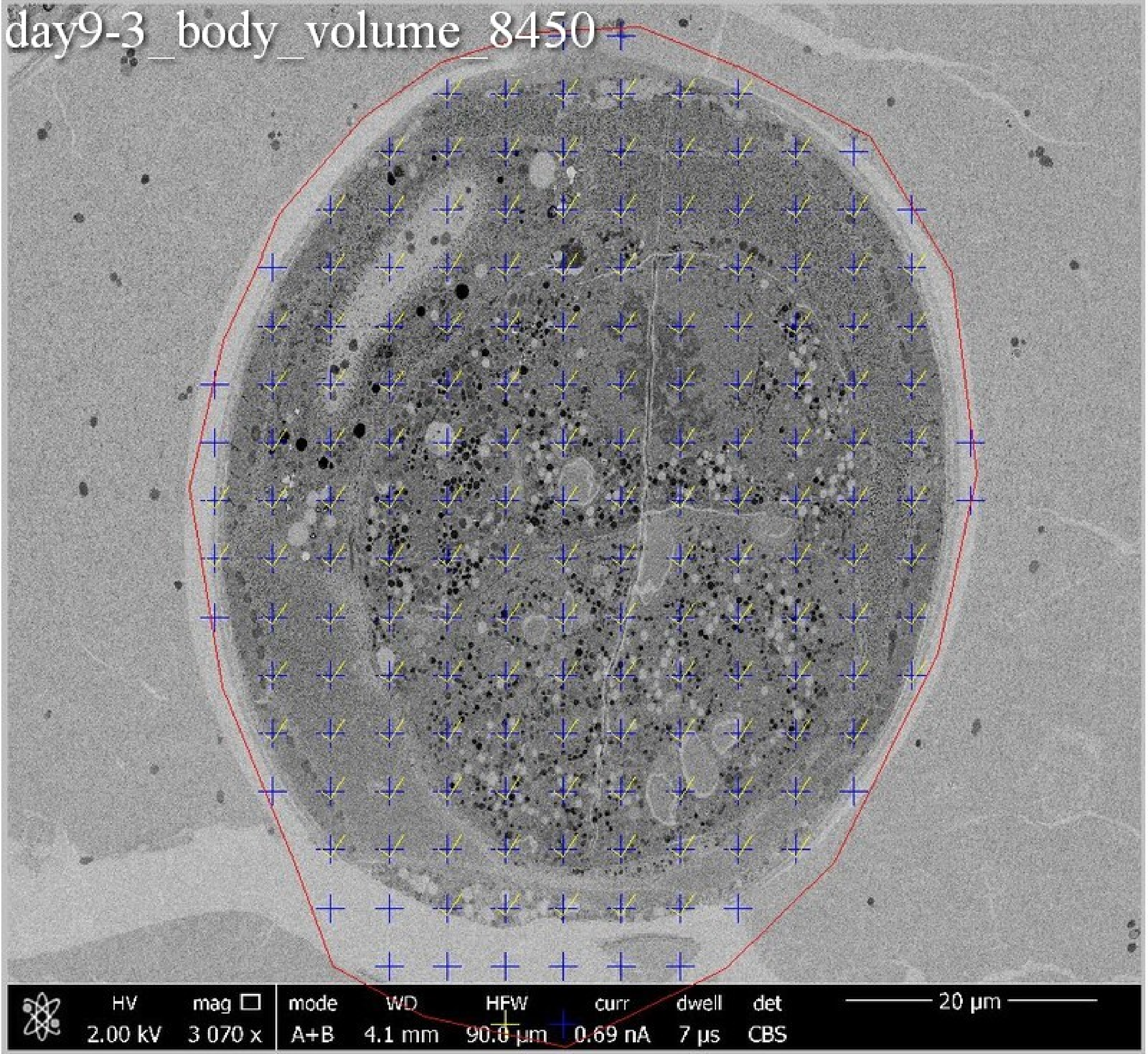

day9-3_body_volume_8450
HV
2.00 kV
mag
3 070 x
mode
A+B
WD
4.1 mm
HFW
90.0 µm
curr
0.69 nA
dwell
7 µs
det
CBS
20 µm

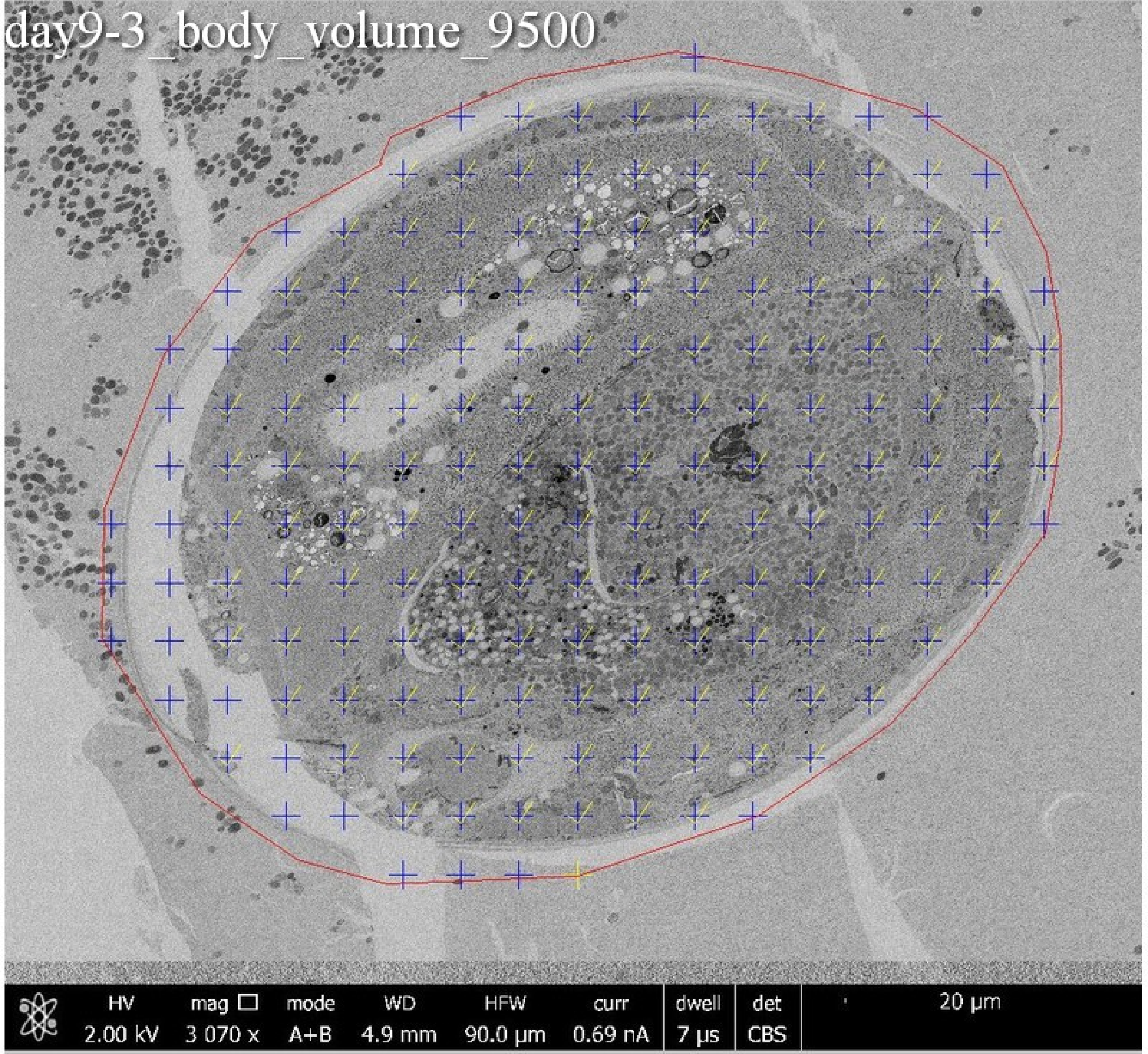

day9-3_body_volume_9500
HV
2.00 kV
mag ☐
3 070 x
mode
A+B
WD
4.9 mm
HFW
90.0 μm
curr
0.69 nA
dwell
7 μs
det
CBS
20 μm

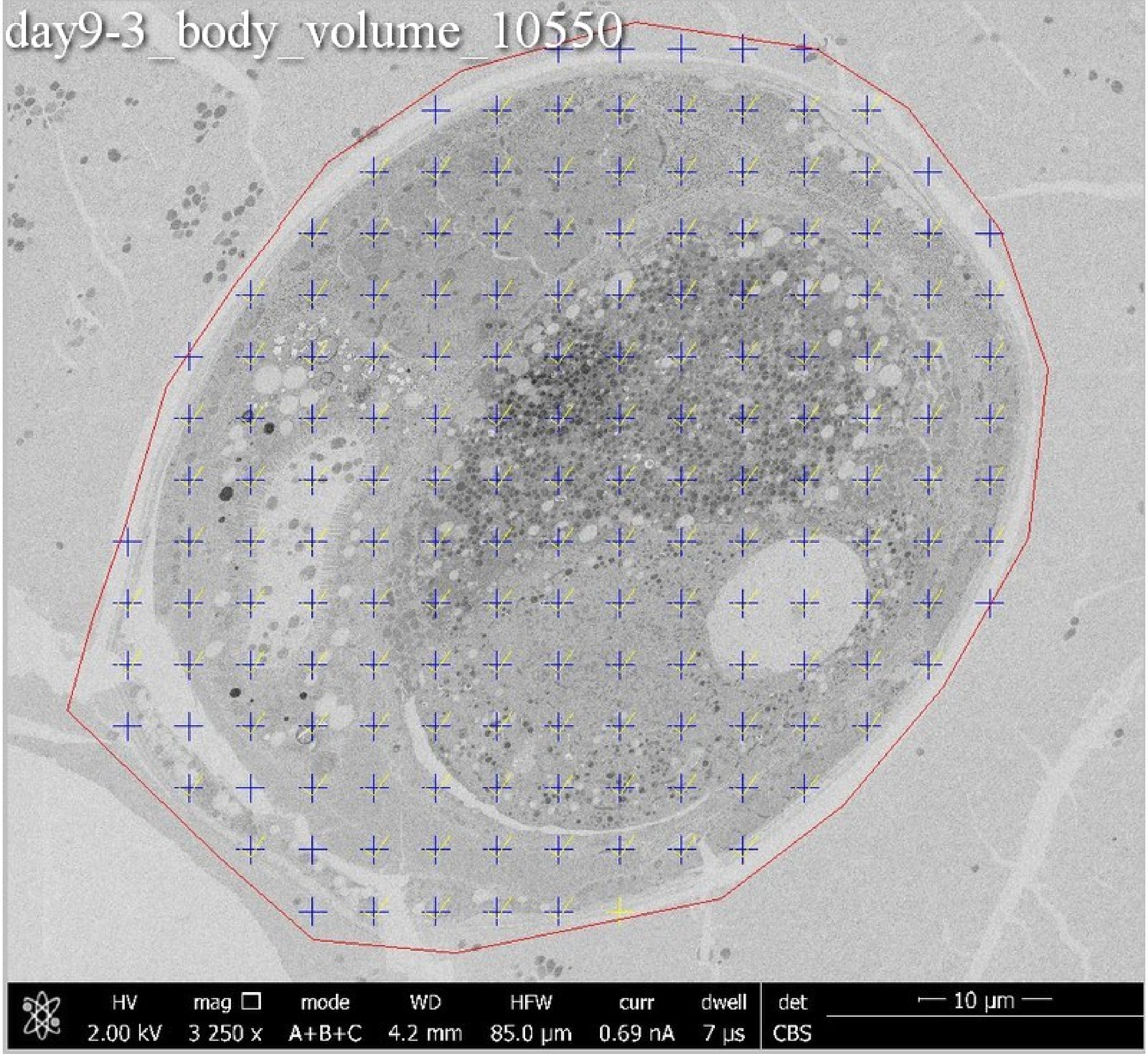
day9-3_body_volume_10550
HV 2.00 kV
mag 3 250 x
mode A+B+C
WD 4.2 mm
HFW 85.0 µm
curr 0.69 nA
dwell 7 µs
det CBS
10 µm

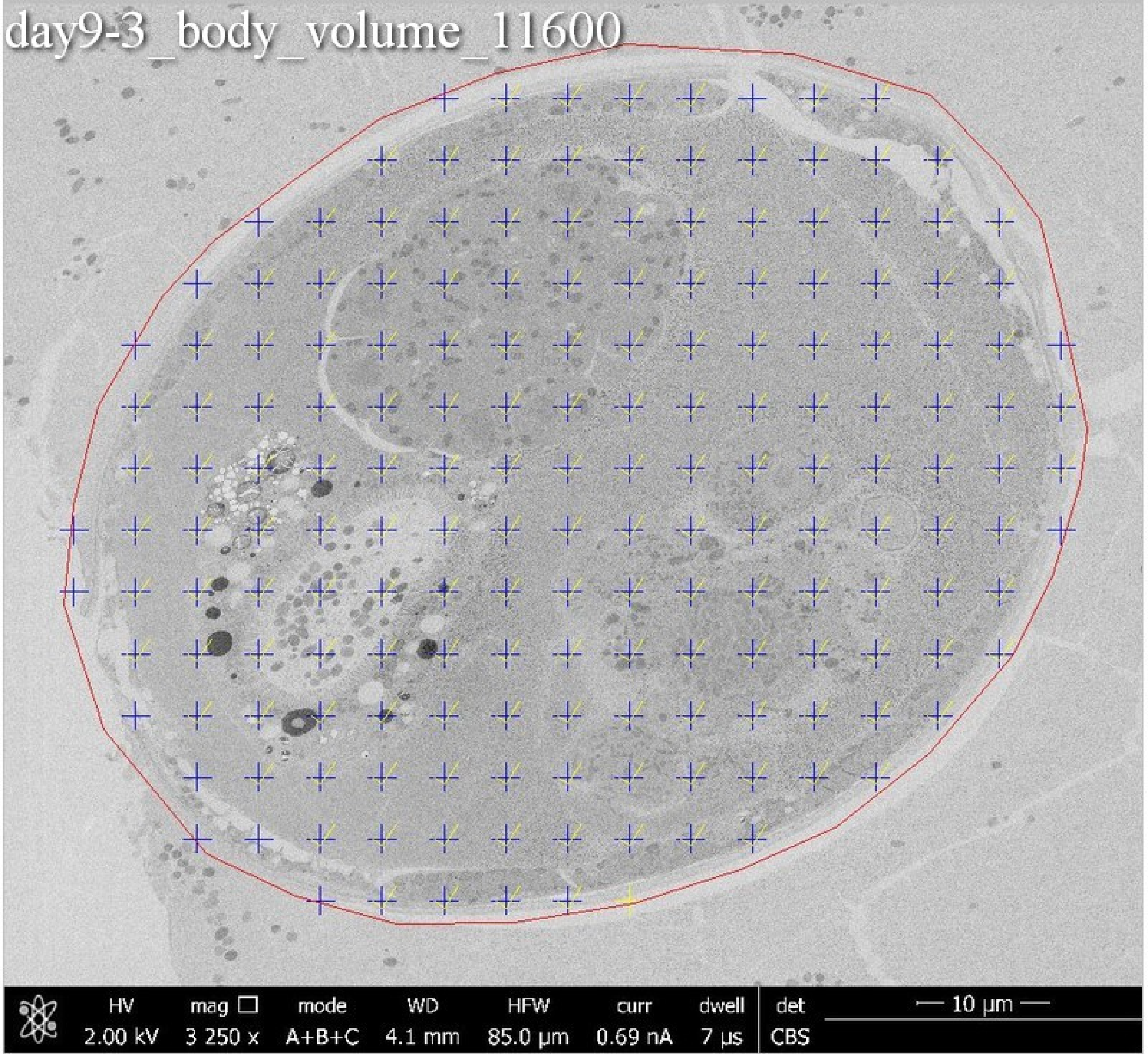
day9-3_body_volume_11600
HV 2.00 kV
mag 3 250 x
mode A+B+C
WD 4.1 mm
HFW 85.0 µm
curr 0.69 nA
dwell 7 µs
det CBS
10 µm

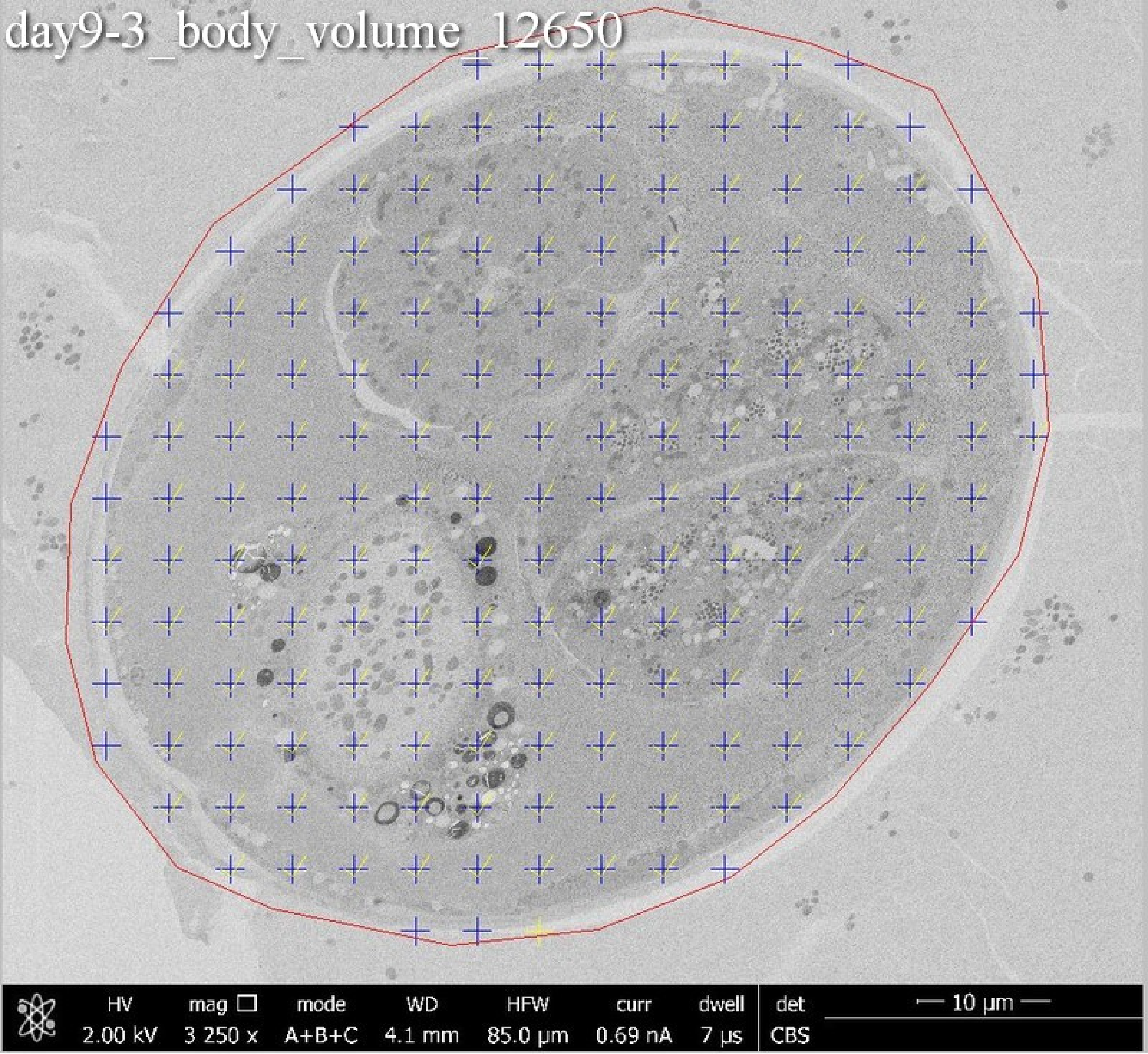
day9-3_body_volume_12650
HV
2.00 kV
mag
3 250 x
mode
A+B+C
WD
4.1 mm
HFW
85.0 µm
curr
0.69 nA
dwell
7 µs
det
CBS
10 µm

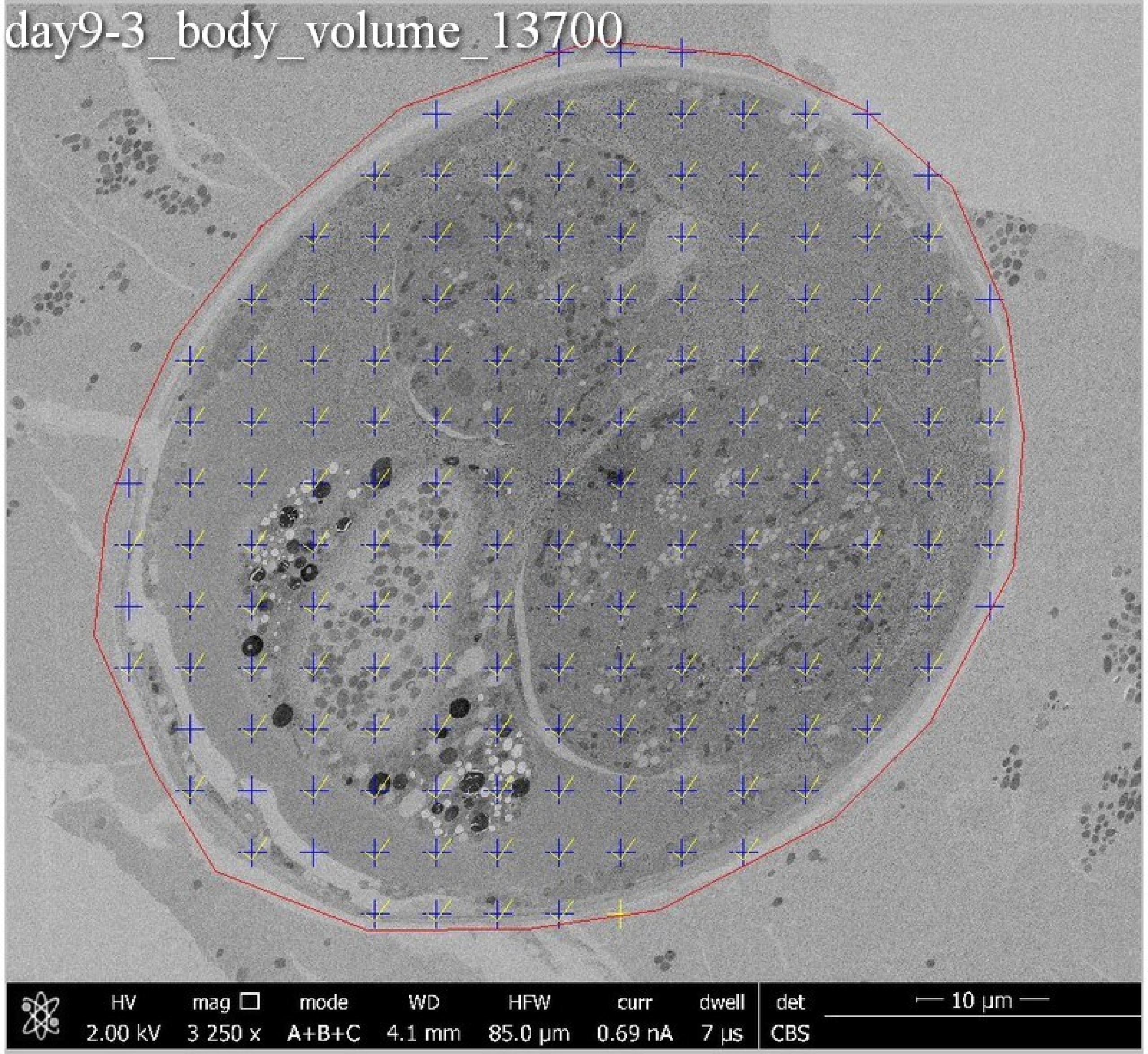
day9-3_body_volume_13700
HV
2.00 kV
mag
3 250 x
mode
A+B+C
WD
4.1 mm
HFW
85.0 µm
curr
0.69 nA
dwell
7 µs
det
CBS
10 µm

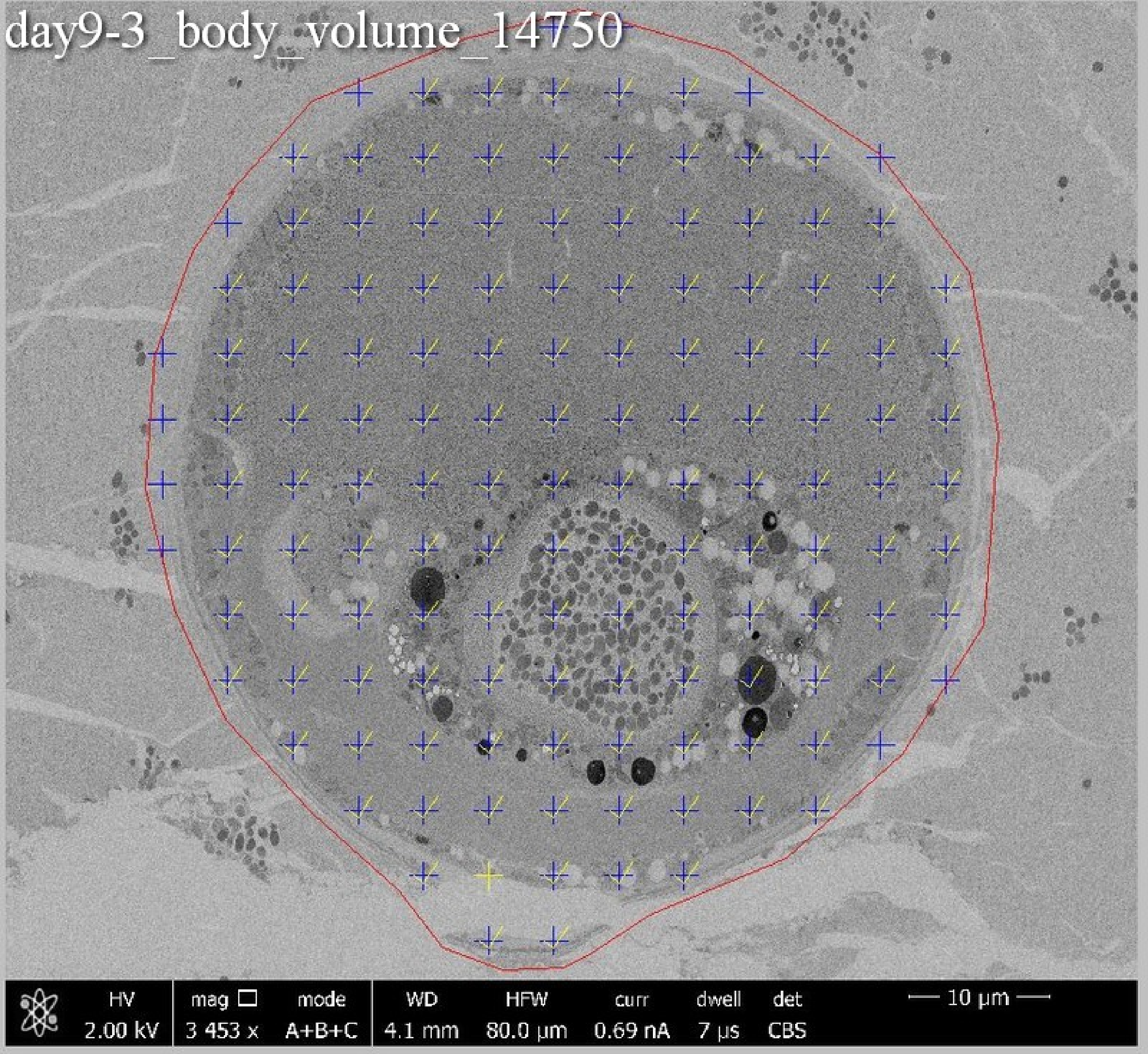
day9-3_body_volume_14750
HV 2.00 kV
mag 3 453 x
mode A+B+C
WD 4.1 mm
HFW 80.0 µm
curr 0.69 nA
dwell 7 µs
det CBS
10 µm

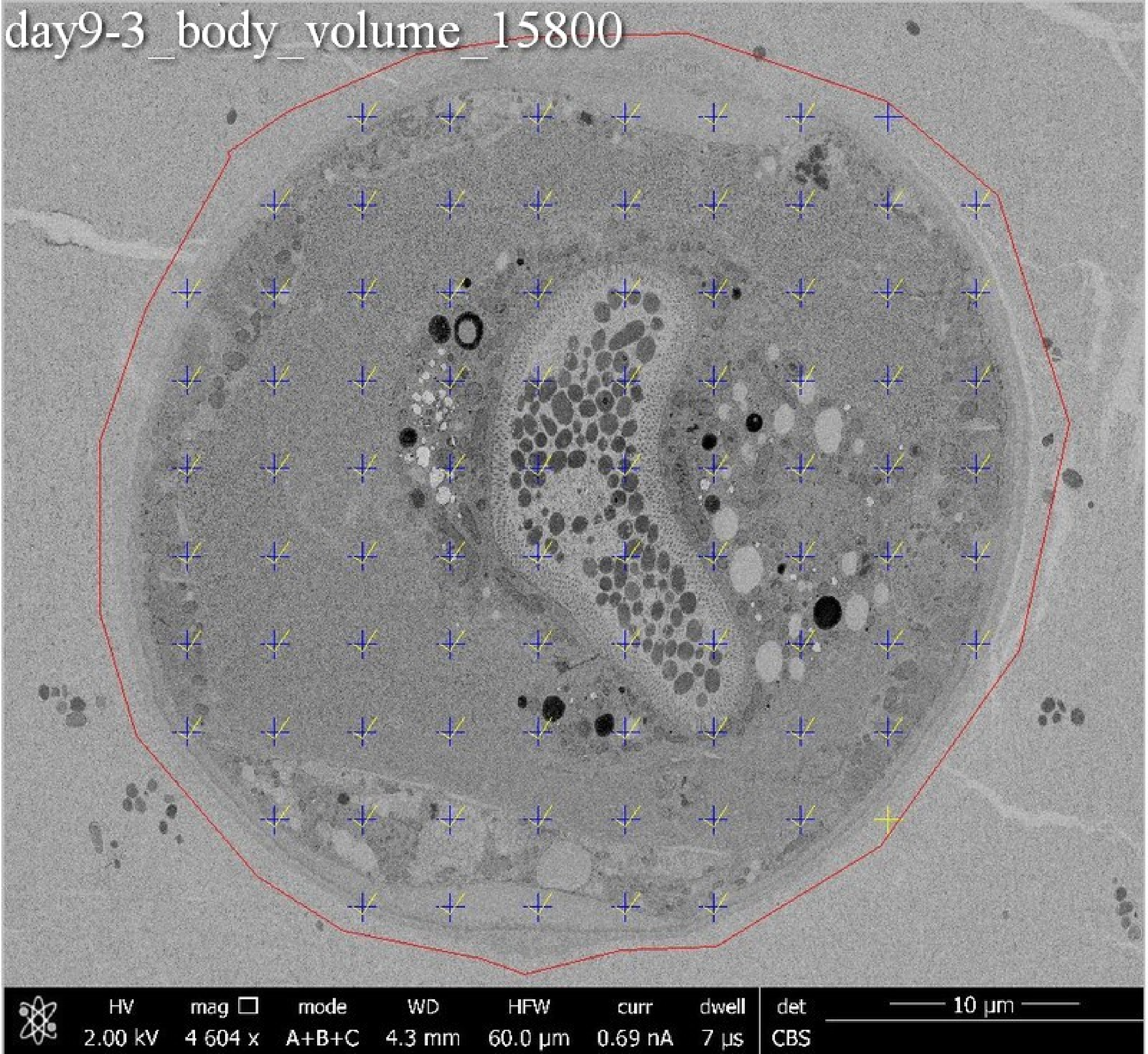
day9-3_body_volume_15800
HV 2.00 kV
mag 4 604 x
mode A+B+C
WD 4.3 mm
HFW 60.0 μm
curr 0.69 nA
dwell 7 μs
det CBS
10 μm

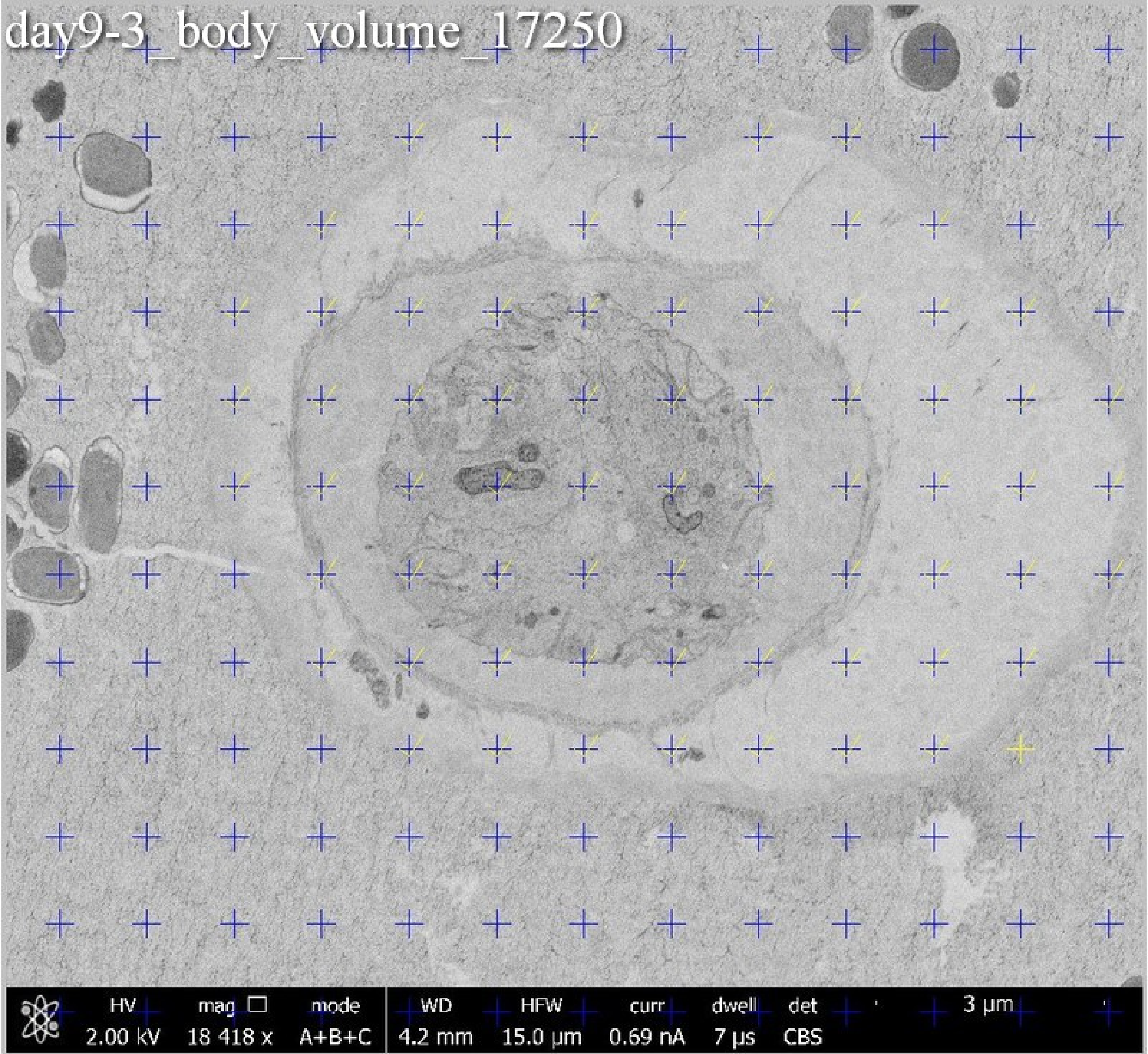

day9-3_body_volume_17250
HV 2.00 kV
mag 18 418 x
mode A+B+C
WD 4.2 mm
HFW 15.0 μm
curr 0.69 nA
dwell 7 μs
det CBS
3 μm

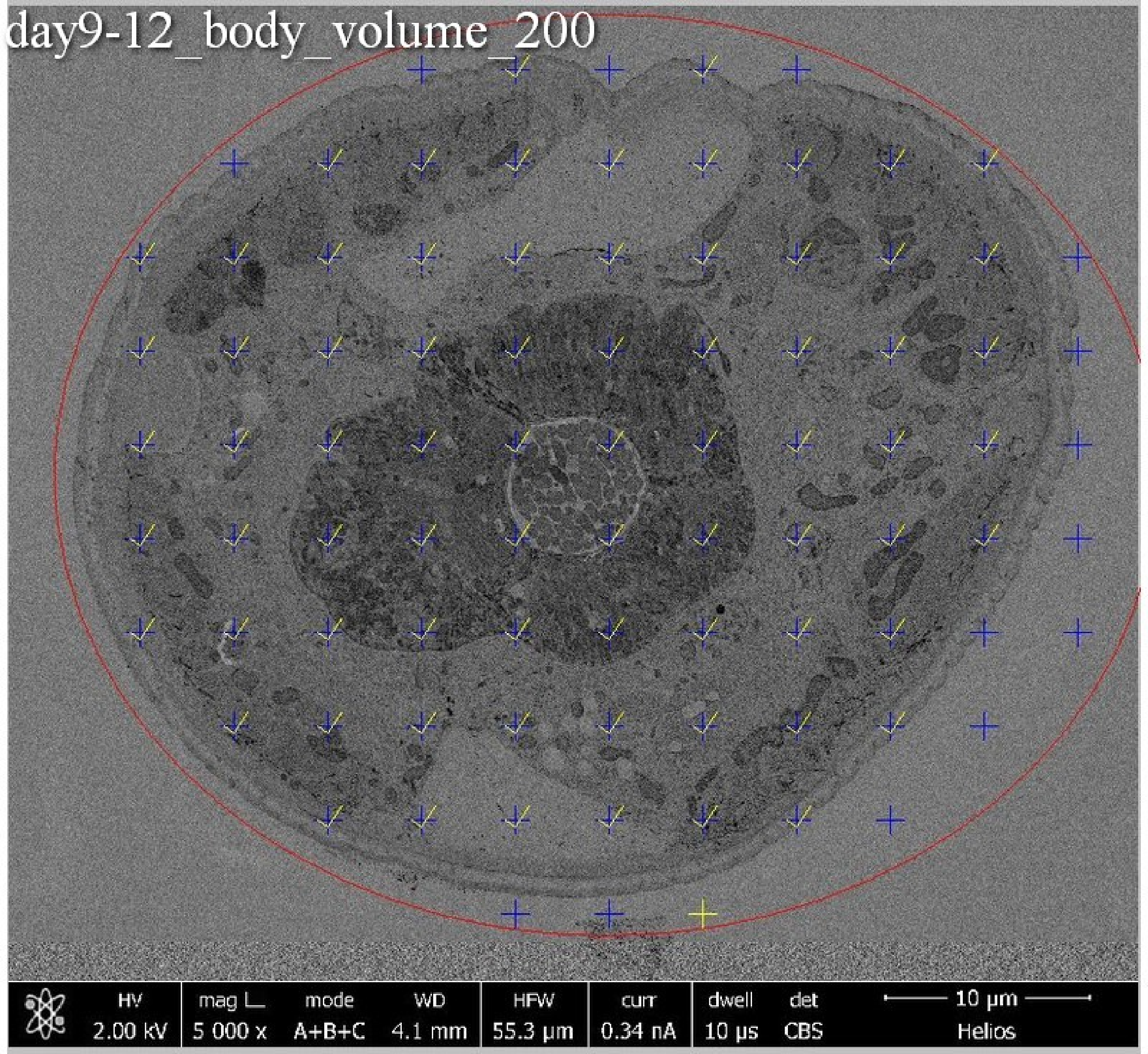
day9-12_body_volume_200
HV
2.00 kV
mag
5 000 x
mode
A+B+C
WD
4.1 mm
HFW
55.3 µm
curr
0.34 nA
dwell
10 µs
det
CBS
10 µm
Helios

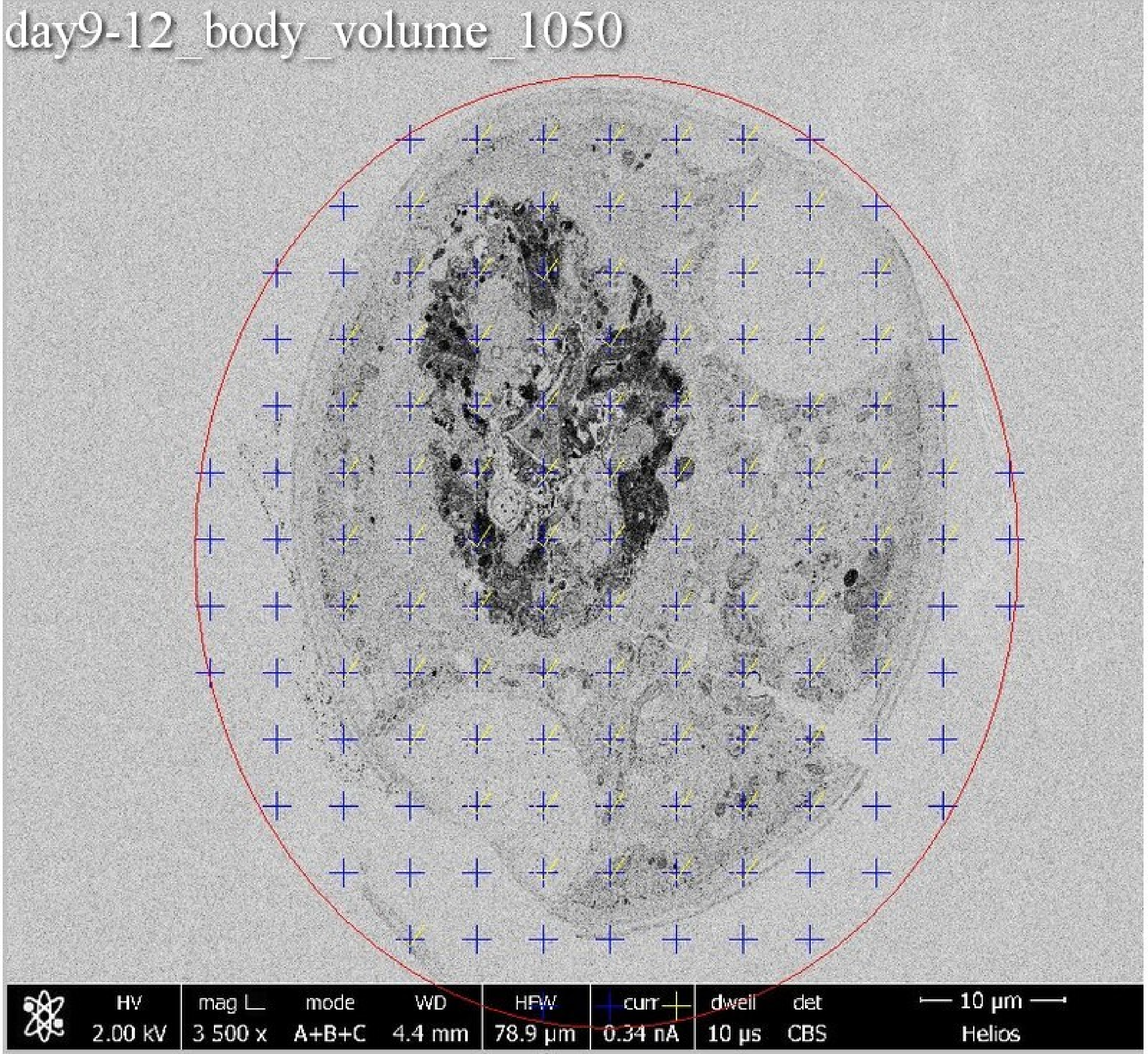
day9-12_body_volume_1050
HV 2.00 kV
mag 3 500 x
mode A+B+C
WD 4.4 mm
HFW 78.9 µm
curr 0.34 nA
dwell 10 µs
det CBS
10 µm
Helios

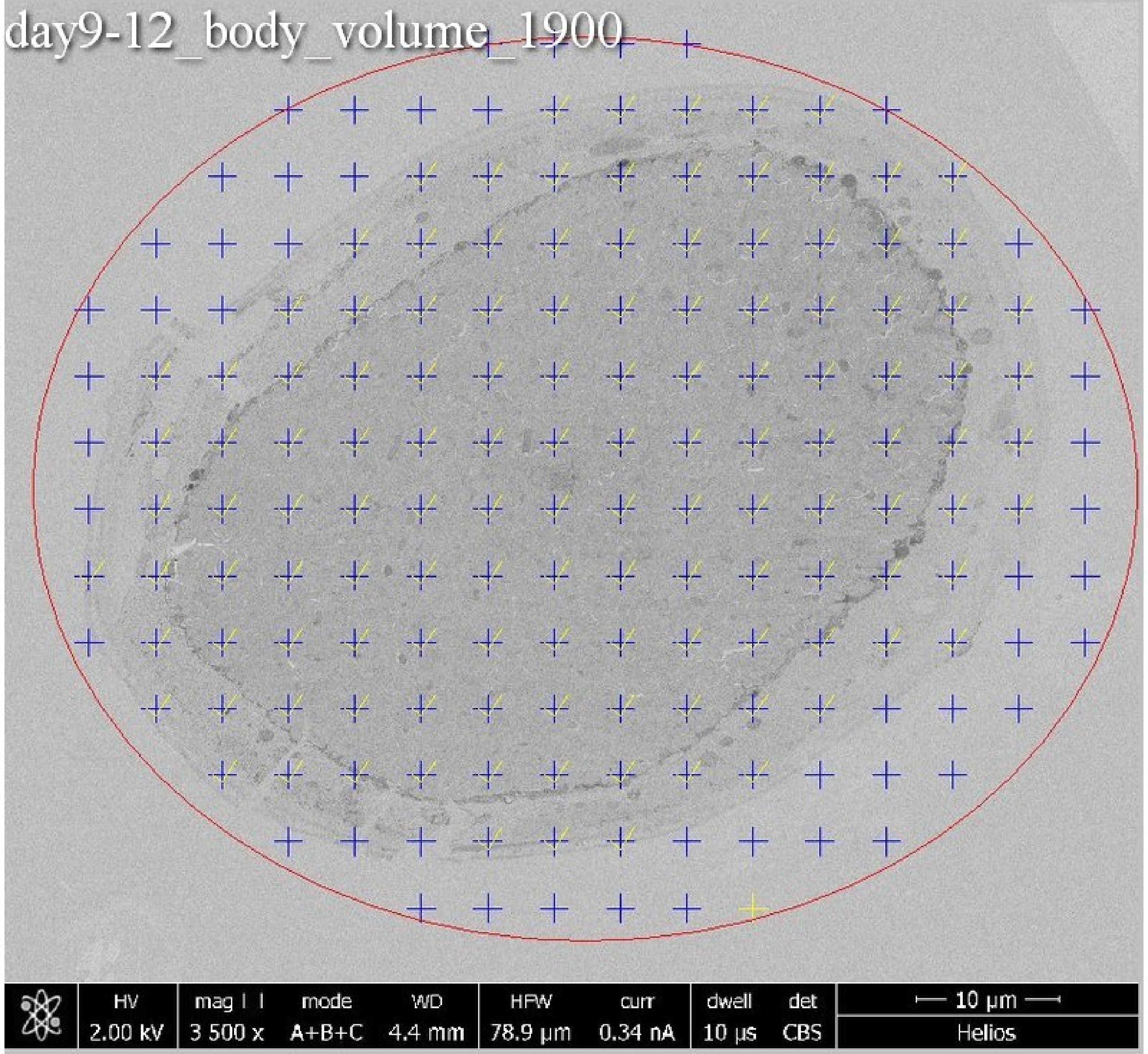
day9-12_body_volume_1900
HV
2.00 kV
mag
3 500 x
mode
A+B+C
WD
4.4 mm
HFW
78.9 µm
curr
0.34 nA
dwell
10 µs
det
CBS
10 µm
Helios

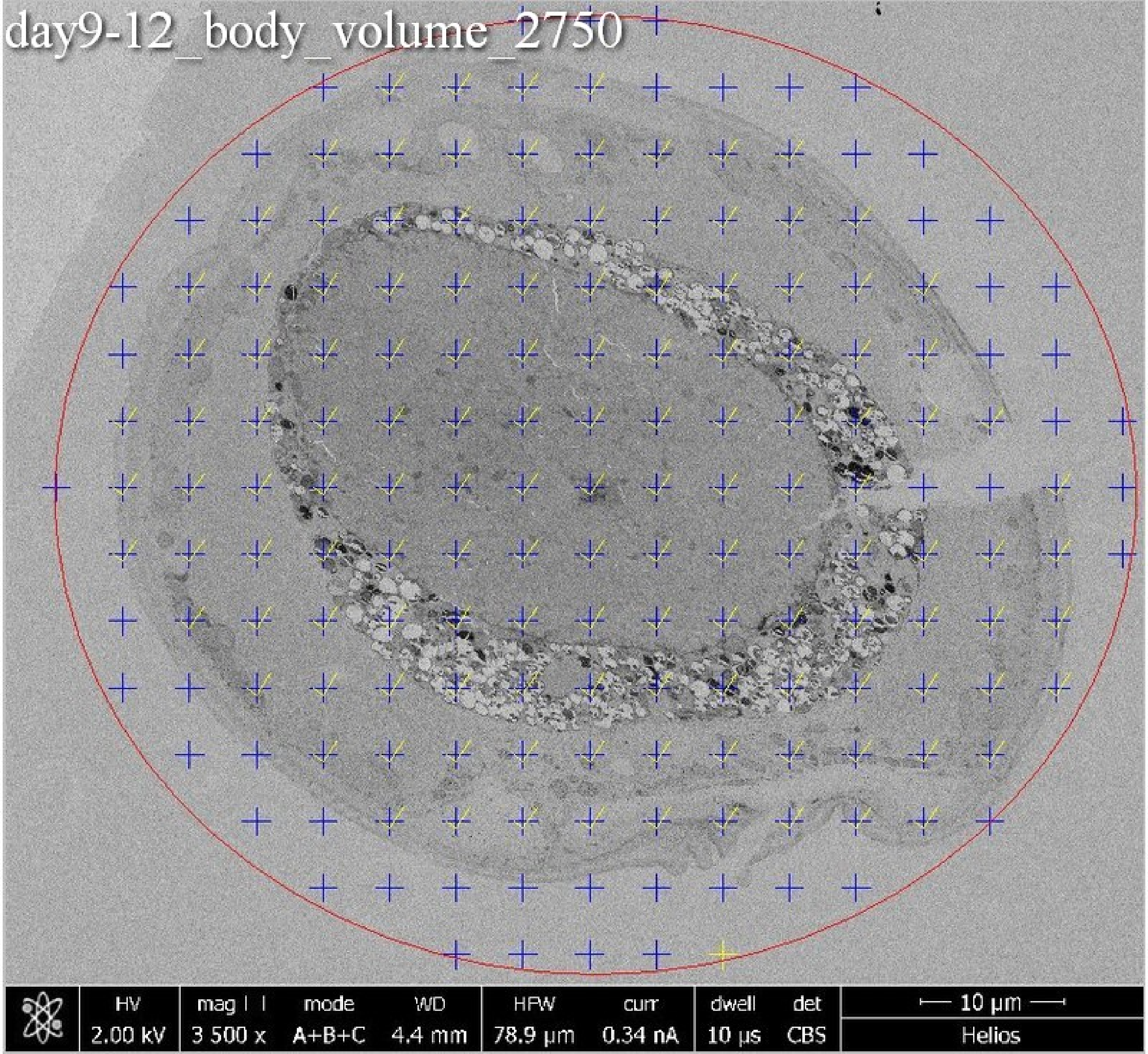
day9-12_body_volume_2750
HV
2.00 kV
mag ⊞
3 500 x
mode
A+B+C
WD
4.4 mm
HFW
78.9 µm
curr
0.34 nA
dwell
10 µs
det
CBS
10 µm
Helios

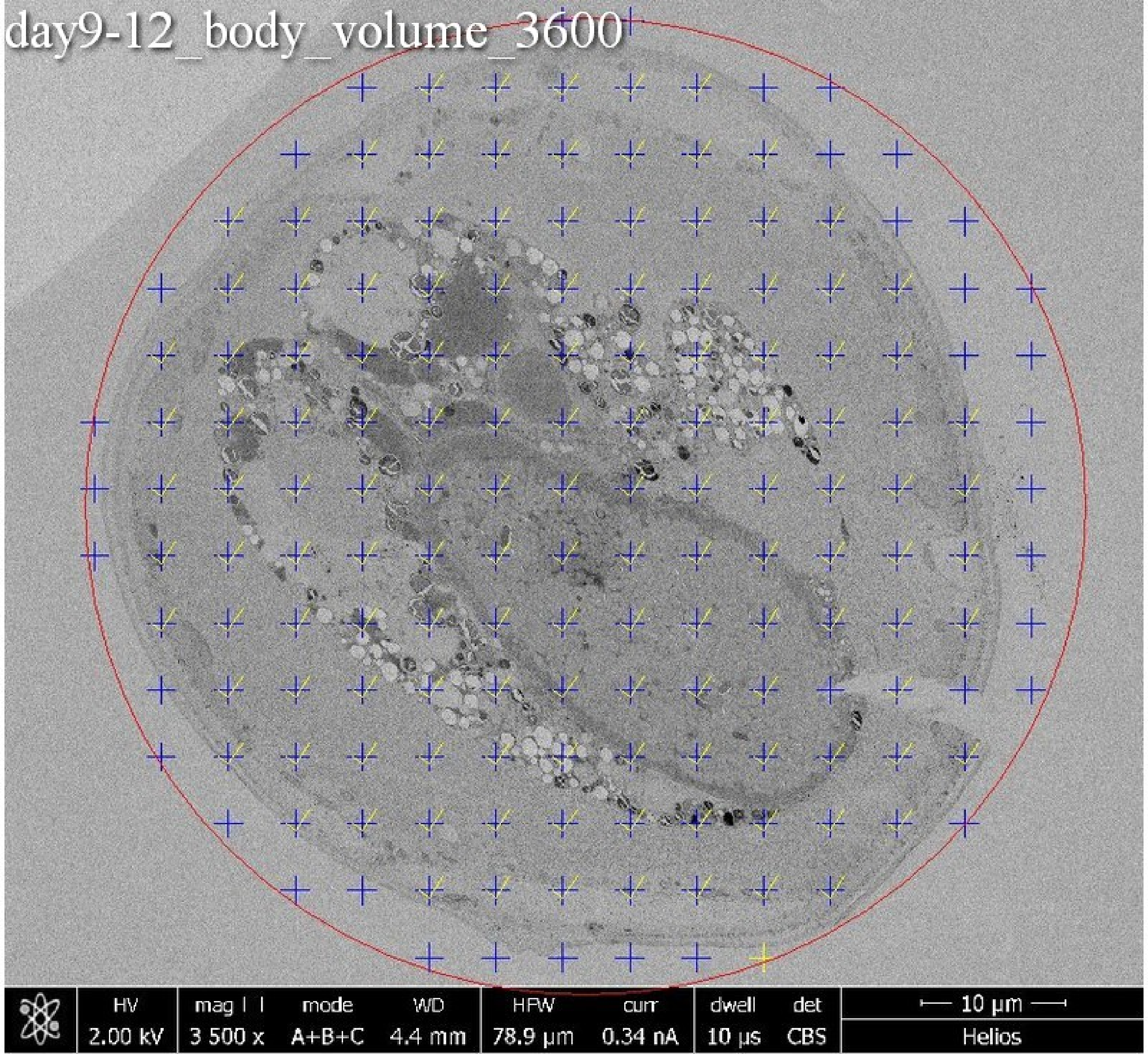

day9-12_body_volume_3600
HV
2.00 kV
mag
3 500 x
mode
A+B+C
WD
4.4 mm
HFW
78.9 µm
curr
0.34 nA
dwell
10 µs
det
CBS
10 µm
Helios

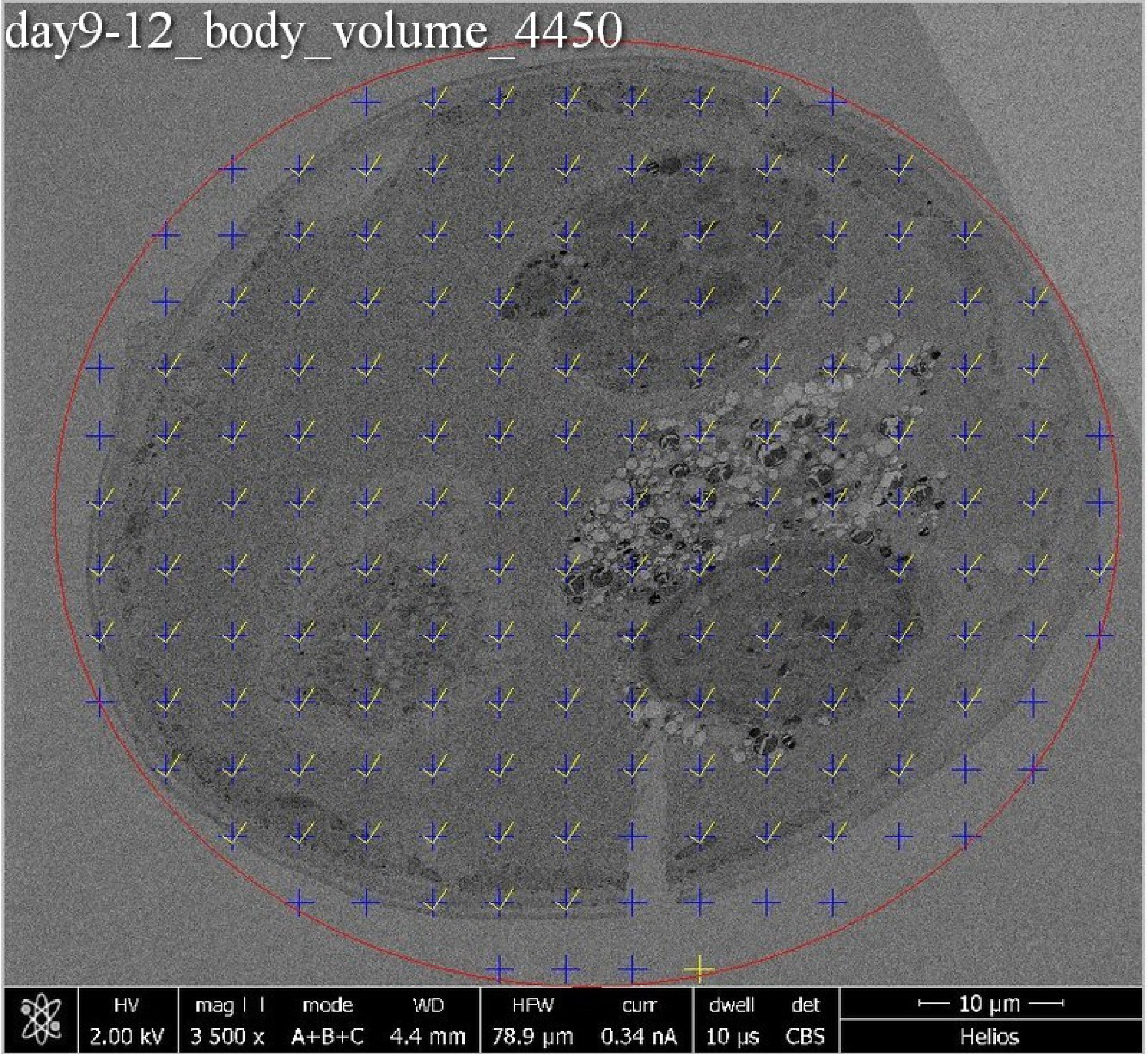
day9-12_body_volume_4450
HV 2.00 kV
mag 3 500 x
mode A+B+C
WD 4.4 mm
HFW 78.9 µm
curr 0.34 nA
dwell 10 µs
det CBS
10 µm
Helios

day9-12_body_volume_5300

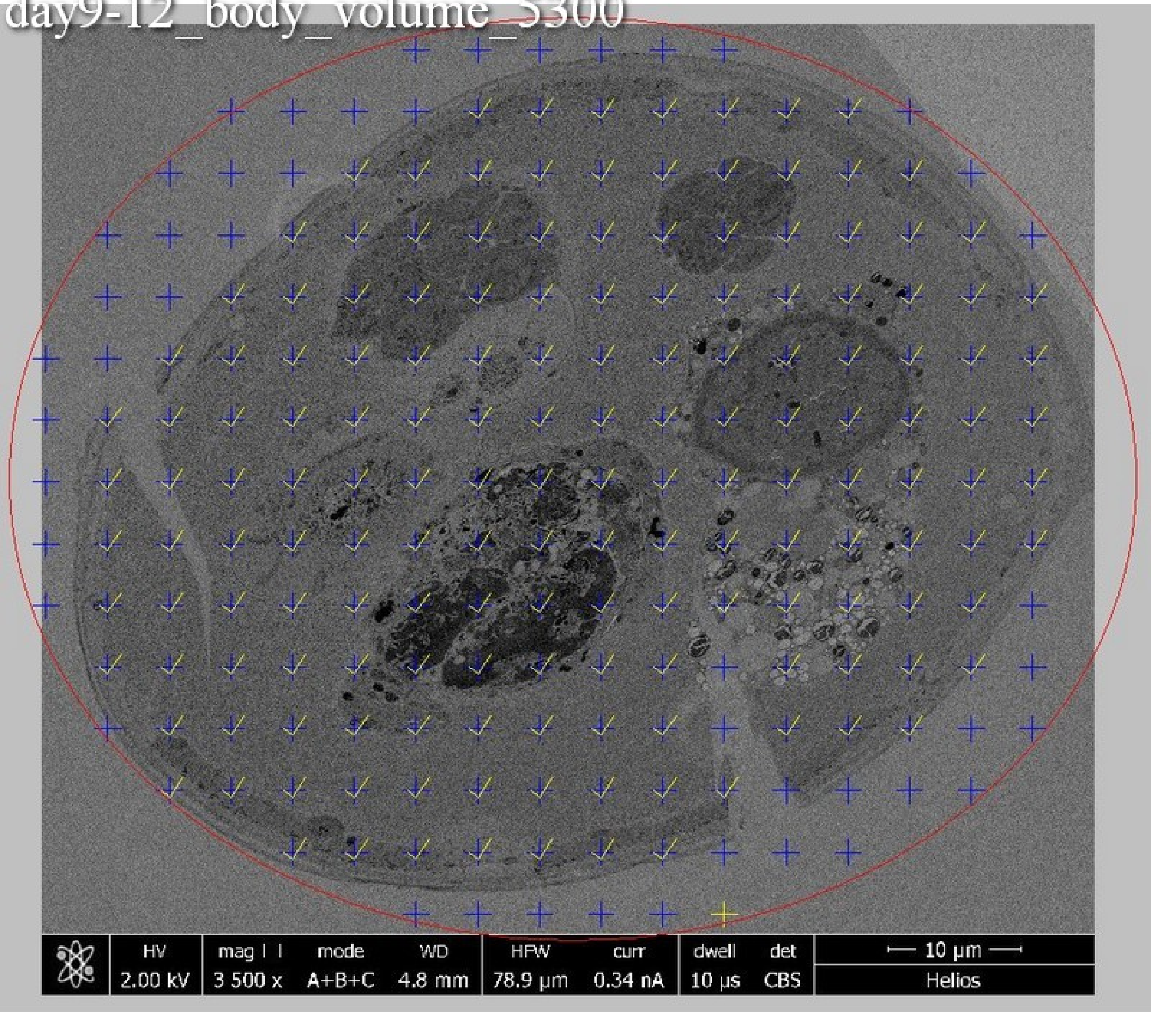

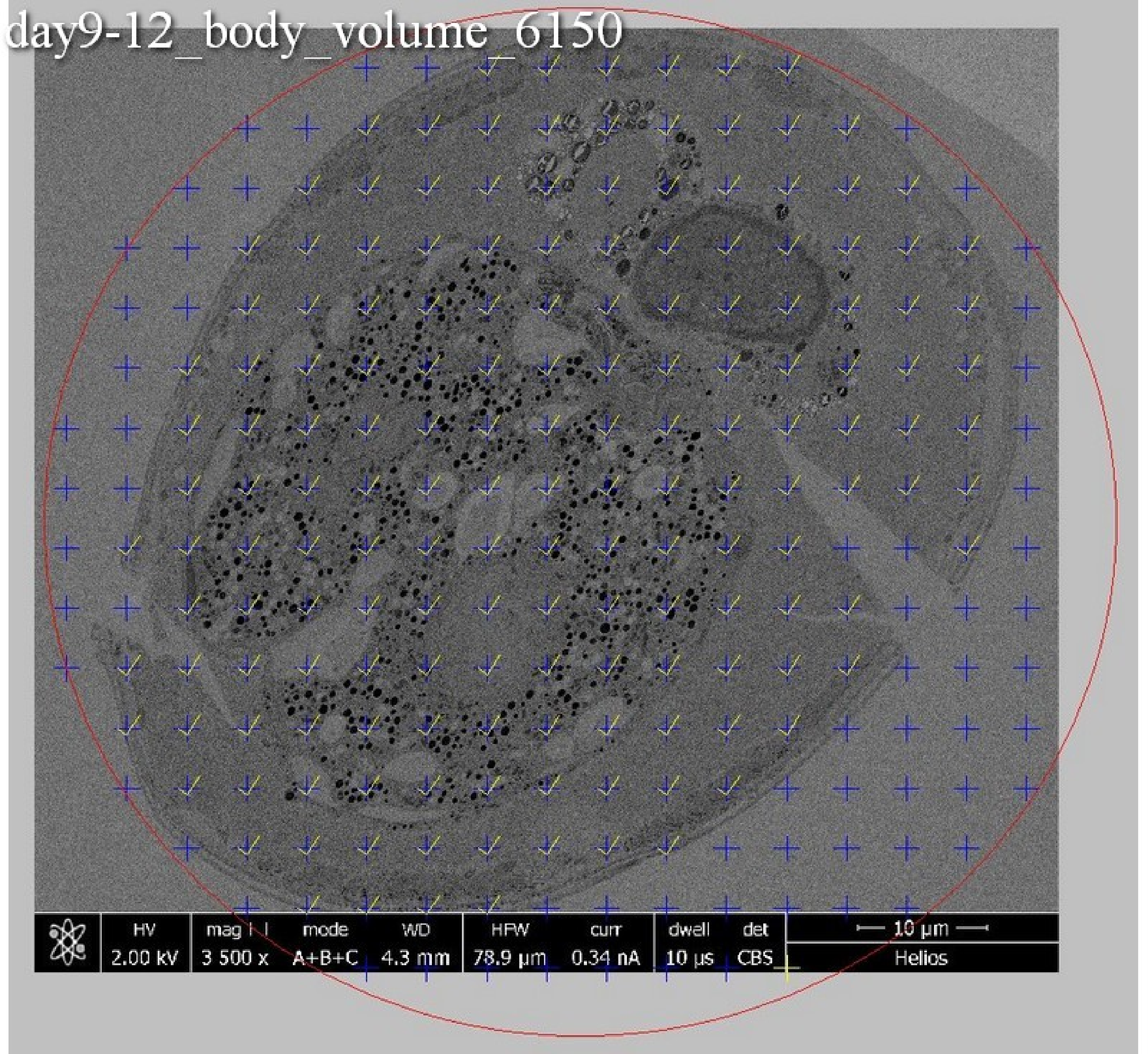
day9-12_body_volume_6150
HV
2.00 kV
mag
3 500 x
mode
A+B+C
WD
4.3 mm
HFW
78.9 µm
curr
0.34 nA
dwell
10 µs
det
CBS
10 µm
Helios

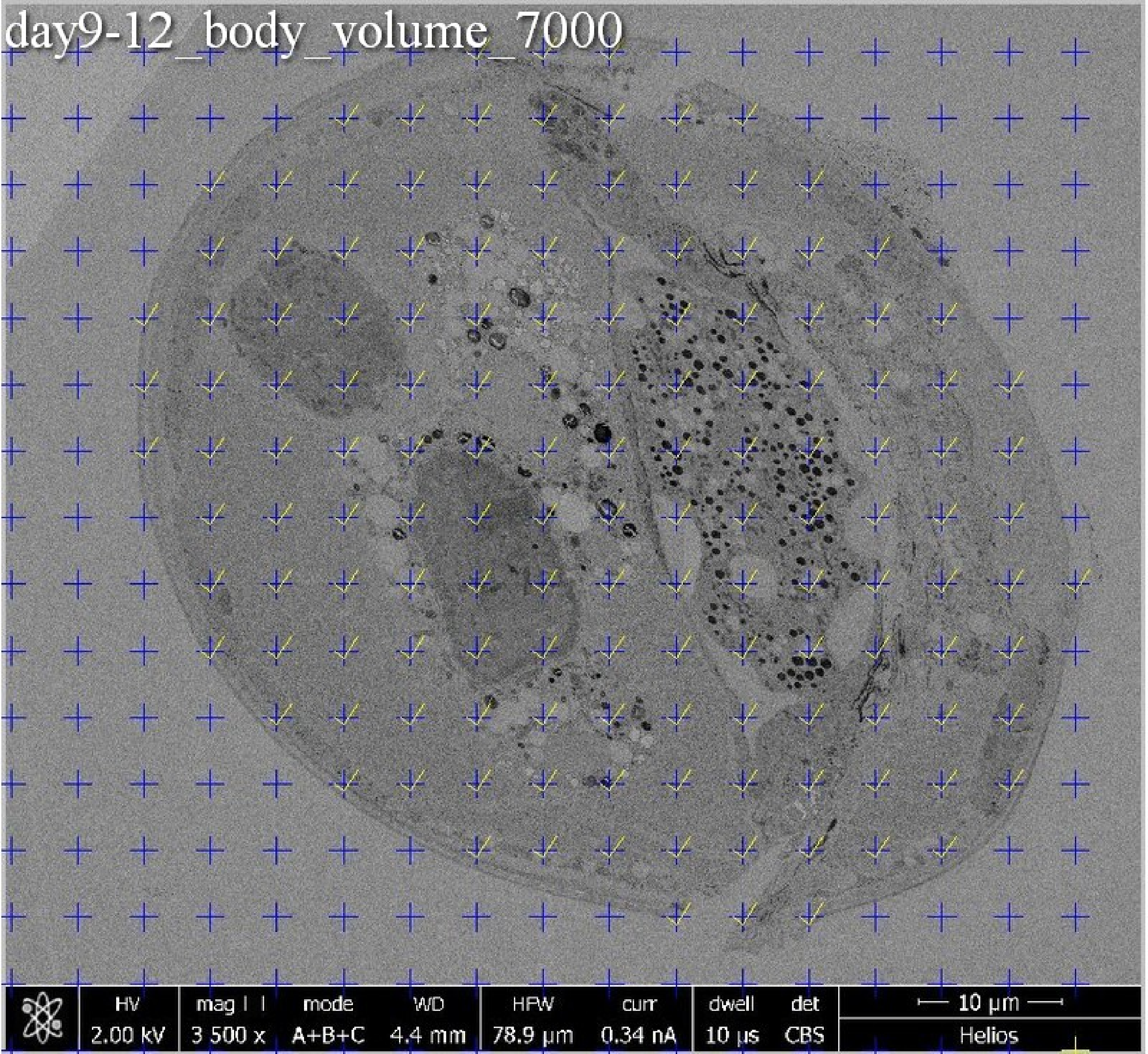
day9-12_body_volume_7000
HV
2.00 kV
mag
3 500 x
mode
A+B+C
WD
4.4 mm
HFW
78.9 µm
curr
0.34 nA
dwell
10 µs
det
CBS
10 µm
Helios

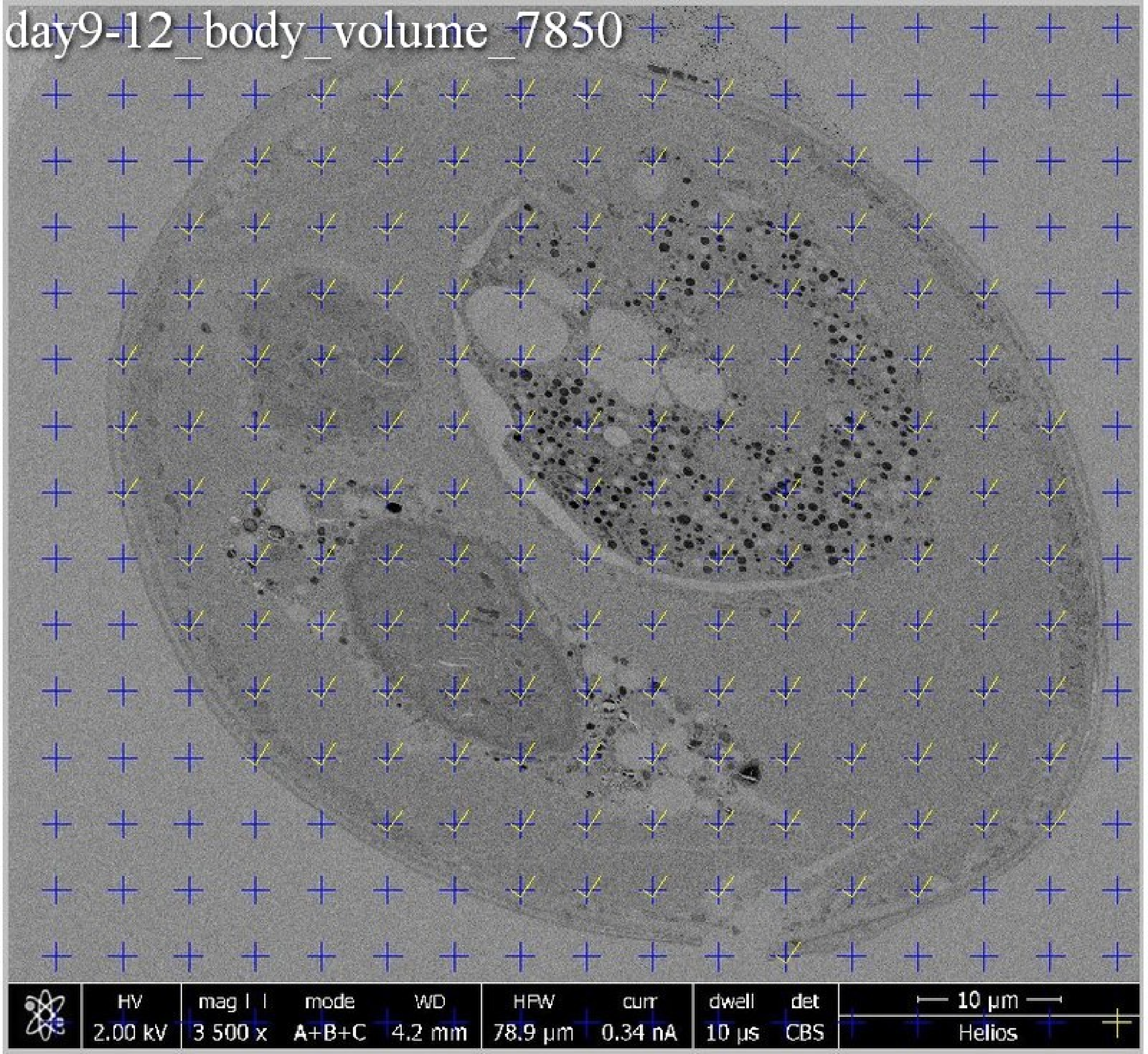

day9-12_body_volume_7850
HV
2.00 kV
mag
3 500 x
mode
A+B+C
WD
4.2 mm
HFW
78.9 µm
curr
0.34 nA
dwell
10 µs
det
CBS
10 µm
Helios

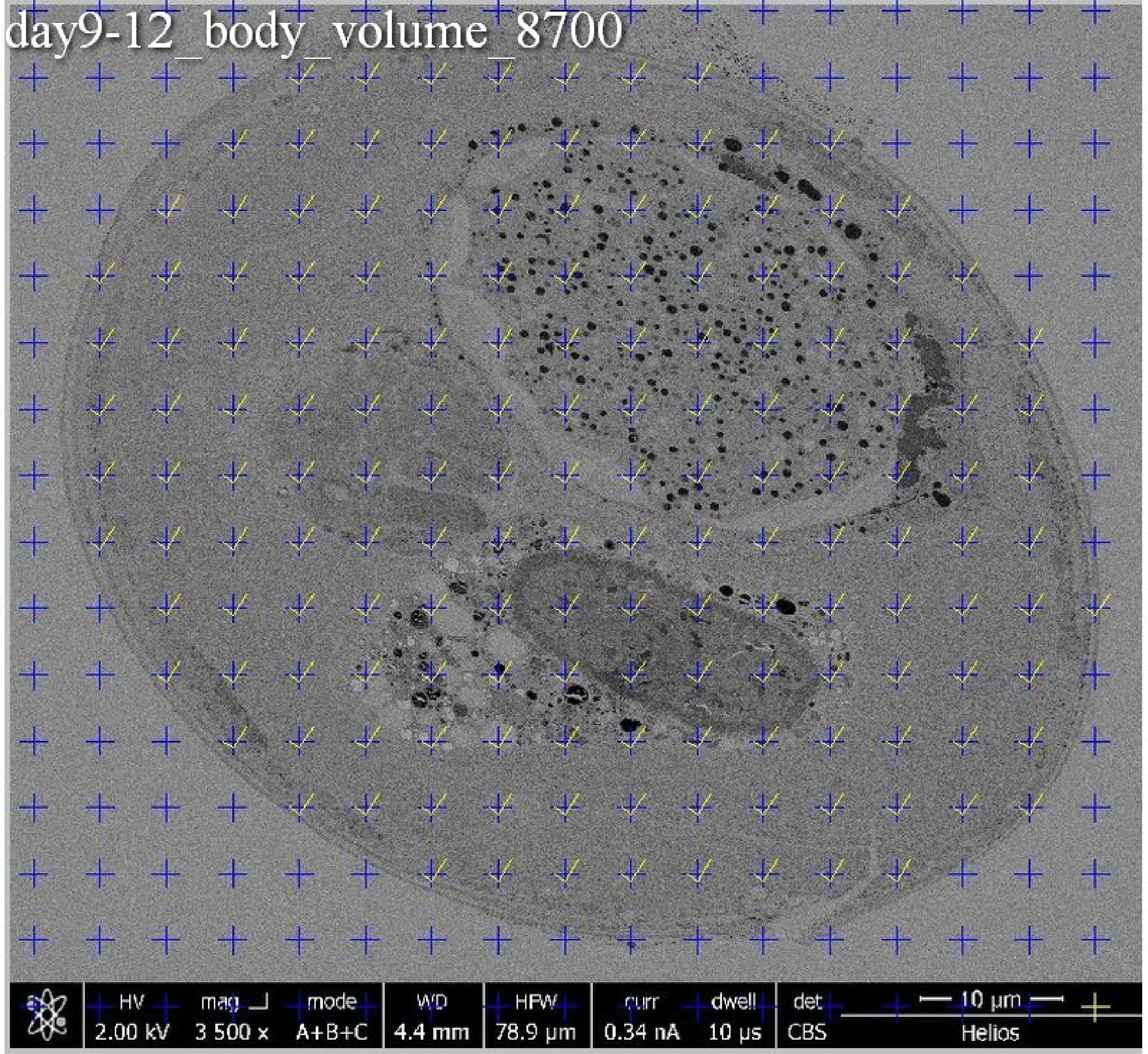
day9-12_body_volume_8700
HV 2.00 kV
mag 3 500 x
mode A+B+C
WD 4.4 mm
HFW 78.9 µm
curr 0.34 nA
dwell 10 µs
det CBS
10 µm
Helios

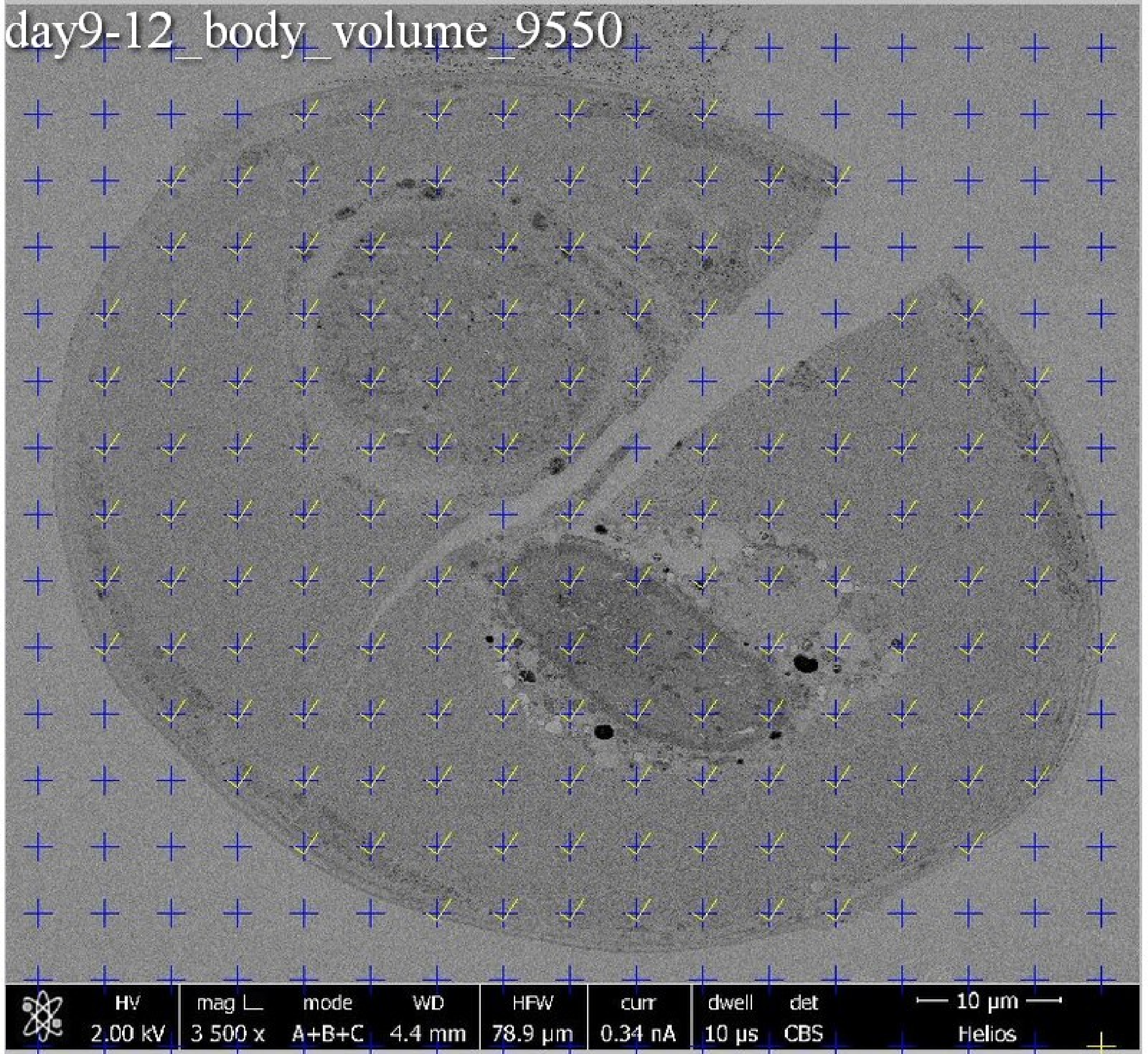
day9-12_body_volume_9550
HV
2.00 kV
mag
3 500 x
mode
A+B+C
WD
4.4 mm
HFW
78.9 µm
curr
0.34 nA
dwell
10 µs
det
CBS
10 µm
Helios

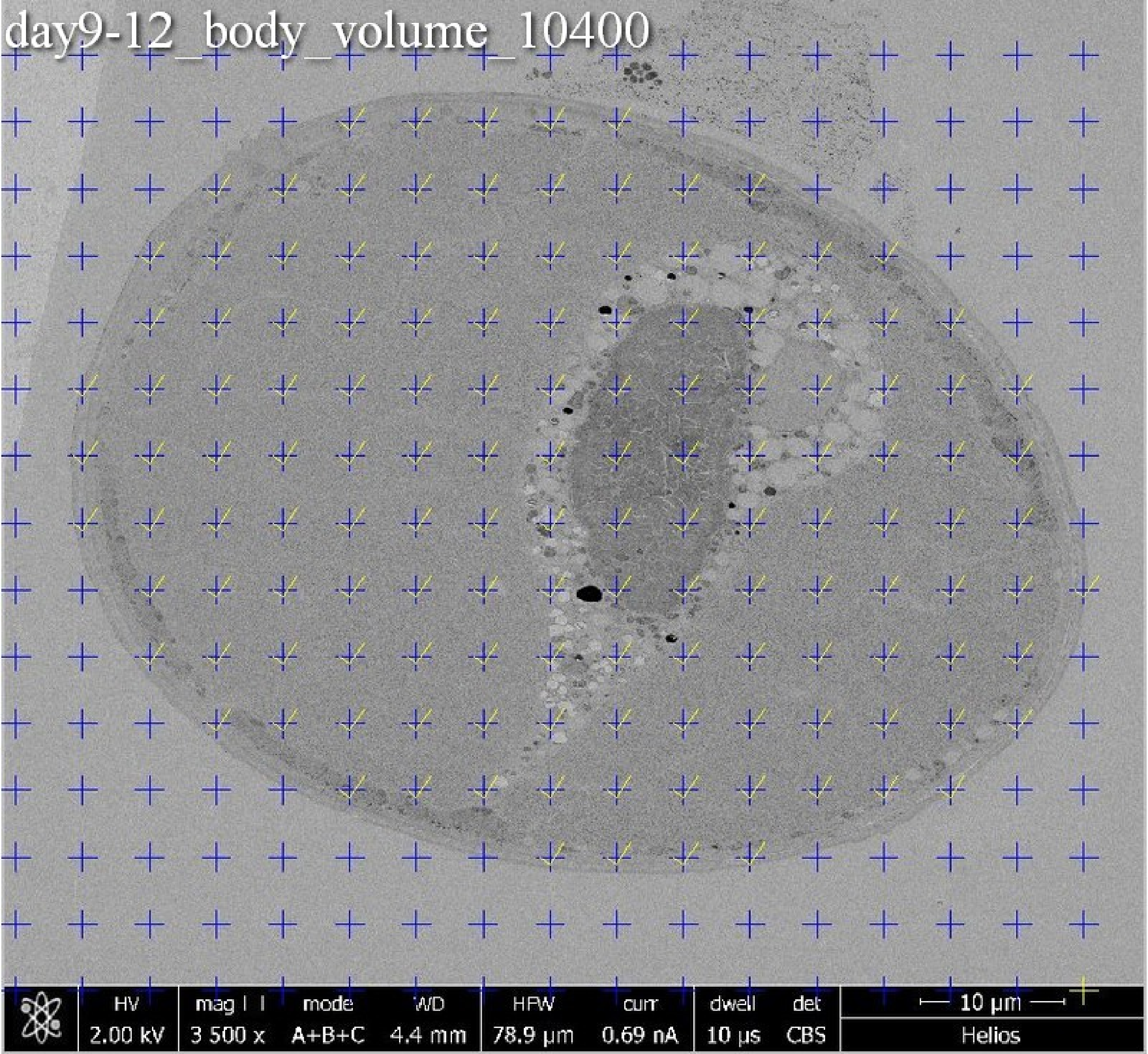
day9-12_body_volume_10400
HV 2.00 kV
mag 3 500 x
mode A+B+C
WD 4.4 mm
HFW 78.9 µm
curr 0.69 nA
dwell 10 µs
det CBS
10 µm
Helios

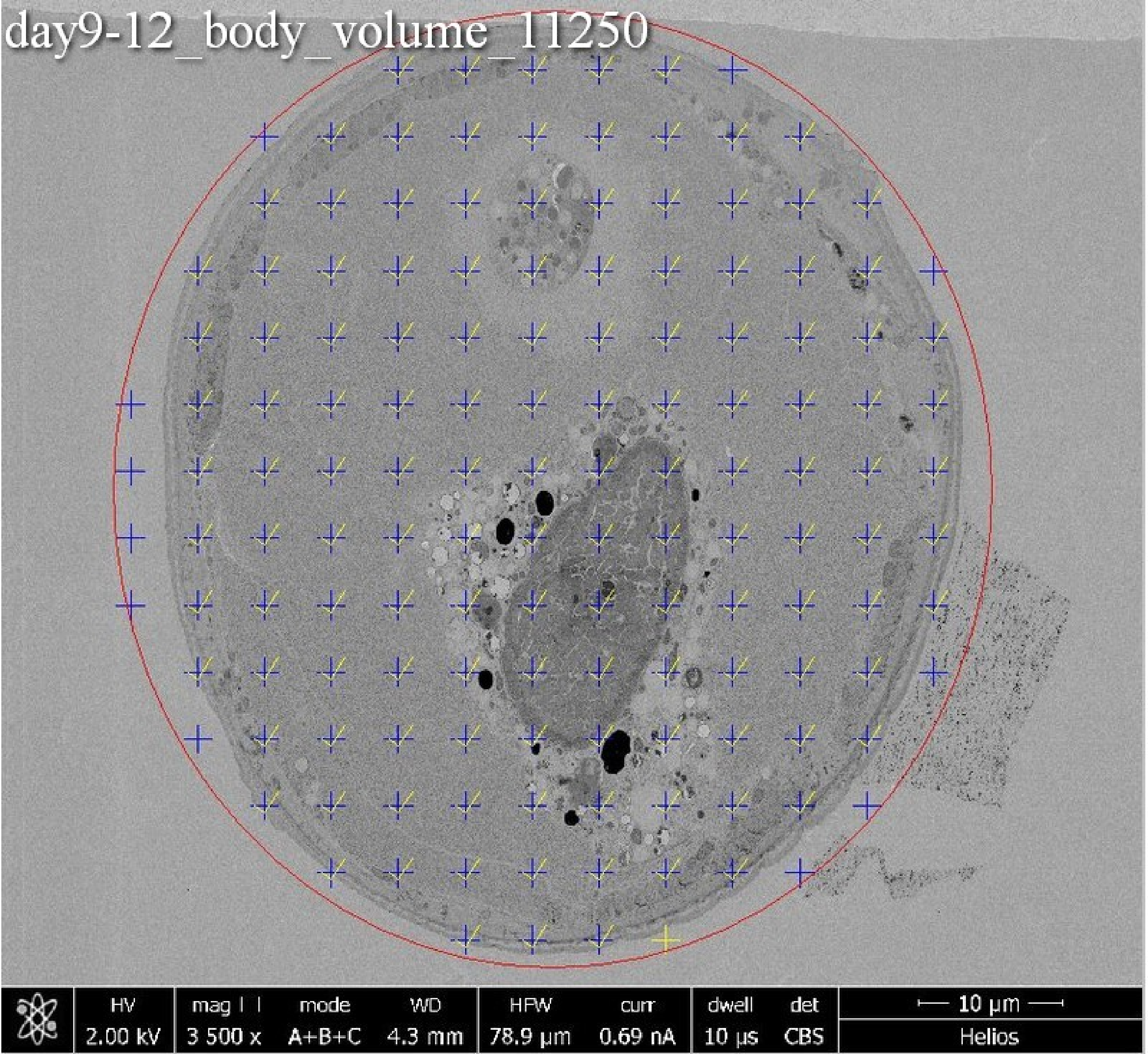
day9-12_body_volume_11250
HV
2.00 kV
mag
3 500 x
mode
A+B+C
WD
4.3 mm
HFW
78.9 µm
curr
0.69 nA
dwell
10 µs
det
CBS
10 µm
Helios

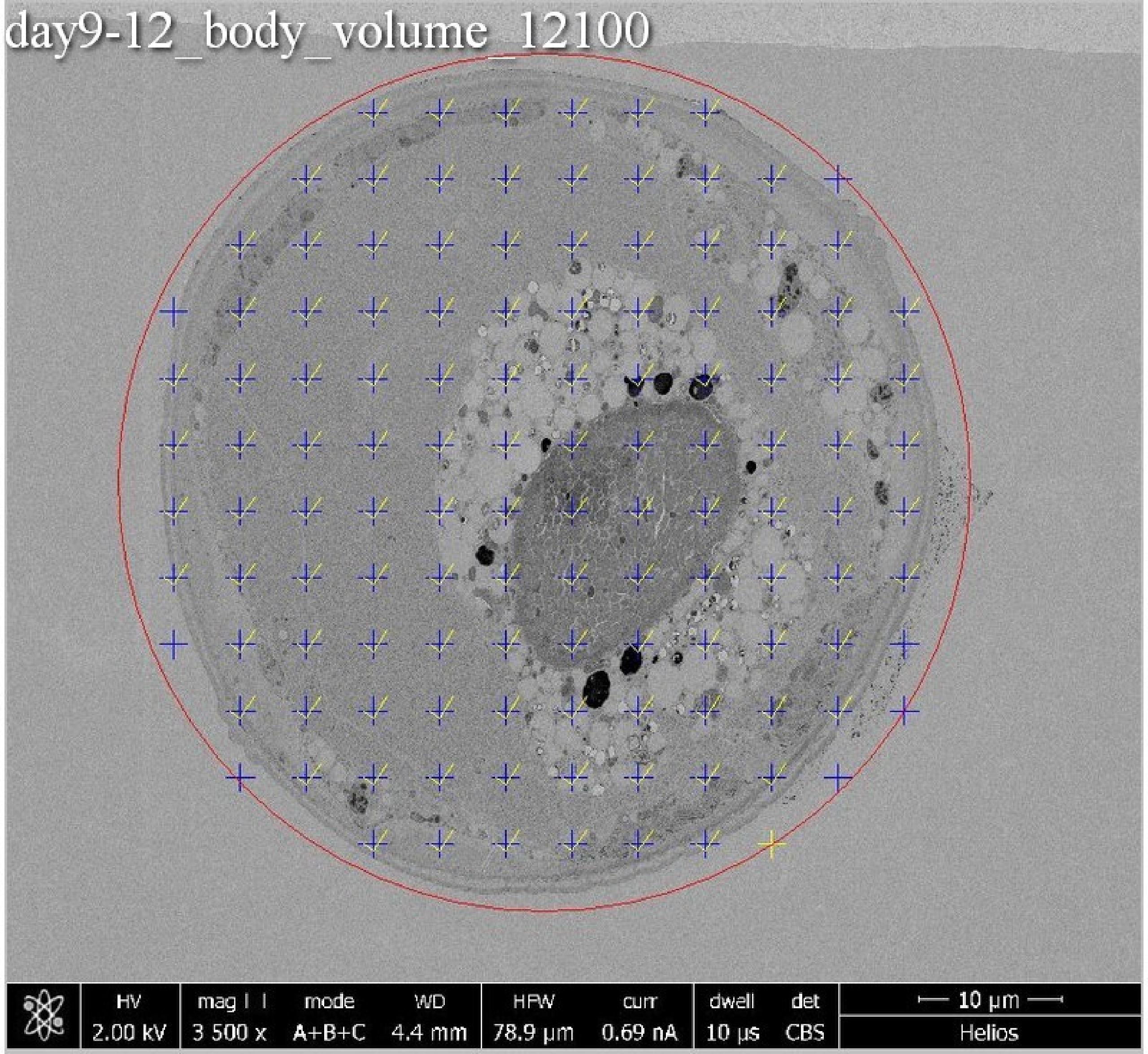
day9-12_body_volume_12100
HV 2.00 kV
mag ⊞ 3 500 x
mode A+B+C
WD 4.4 mm
HFW 78.9 µm
curr 0.69 nA
dwell 10 µs
det CBS
10 µm
Helios

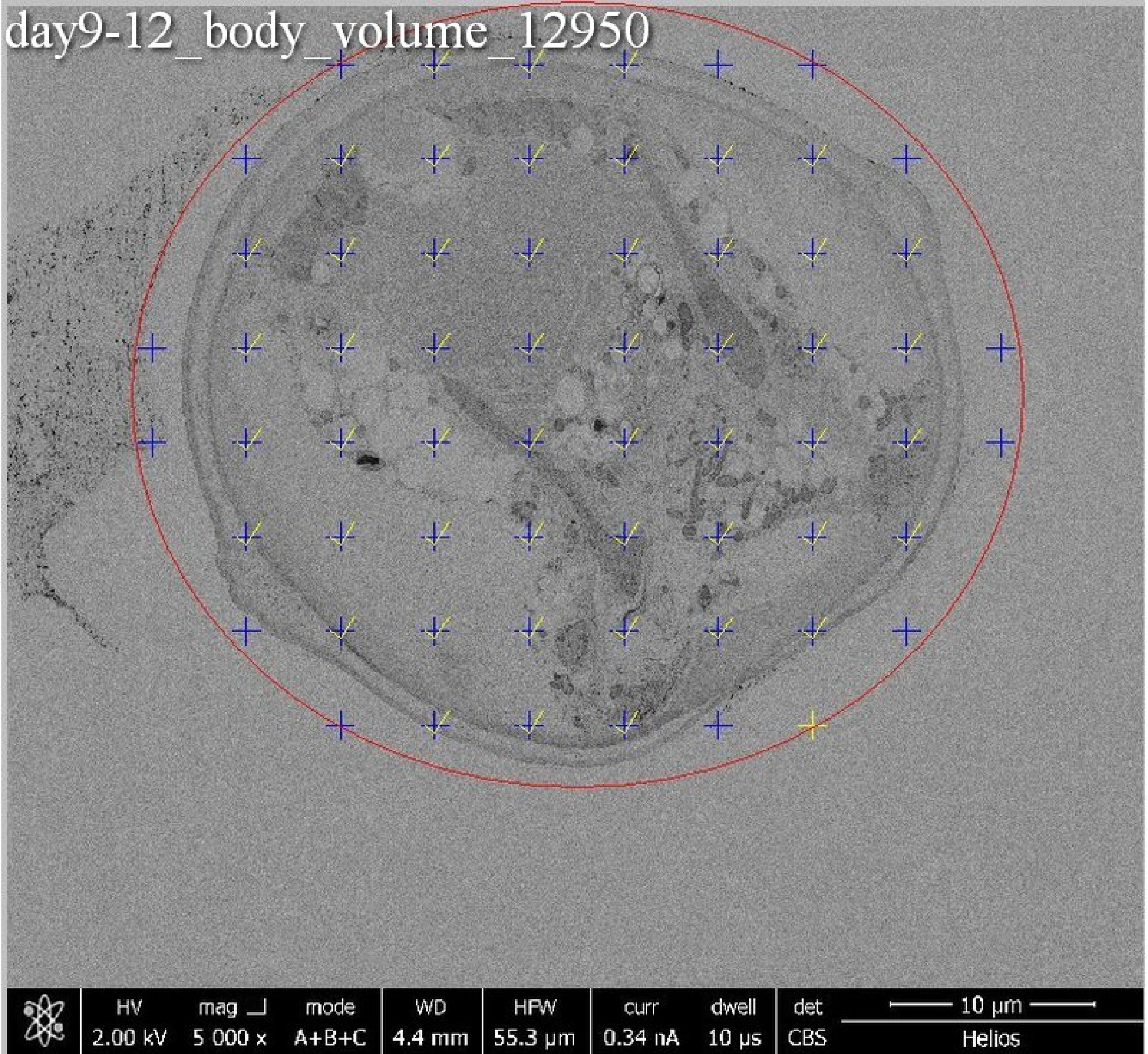
day9-12_body_volume_12950
HV 2.00 kV
mag 5 000 x
mode A+B+C
WD 4.4 mm
HFW 55.3 µm
curr 0.34 nA
dwell 10 µs
det CBS
10 µm
Helios

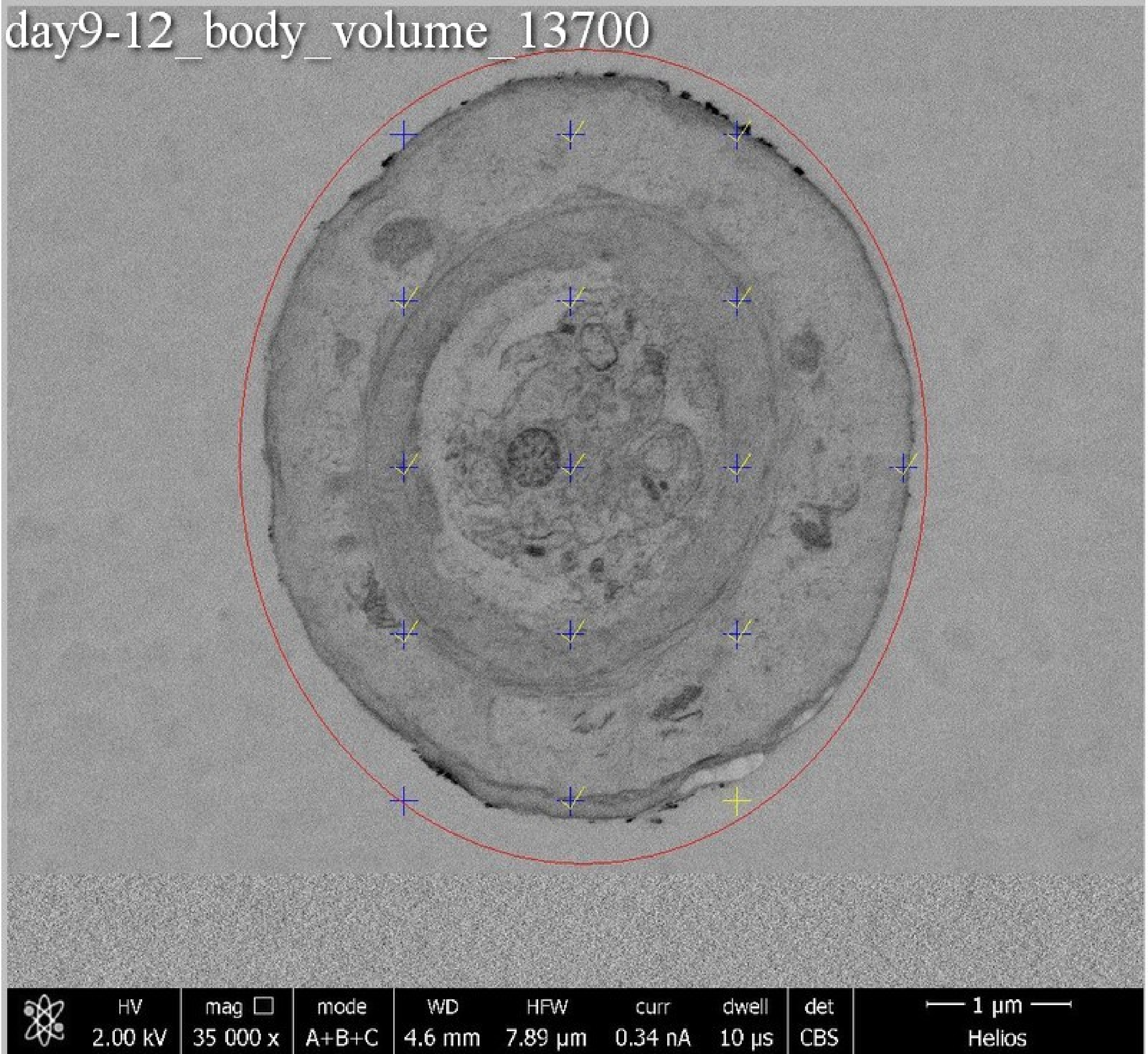
day9-12_body_volume_13700
HV 2.00 kV
mag 35 000 x
mode A+B+C
WD 4.6 mm
HFW 7.89 μm
curr 0.34 nA
dwell 10 μs
det CBS
1 μm
Helios

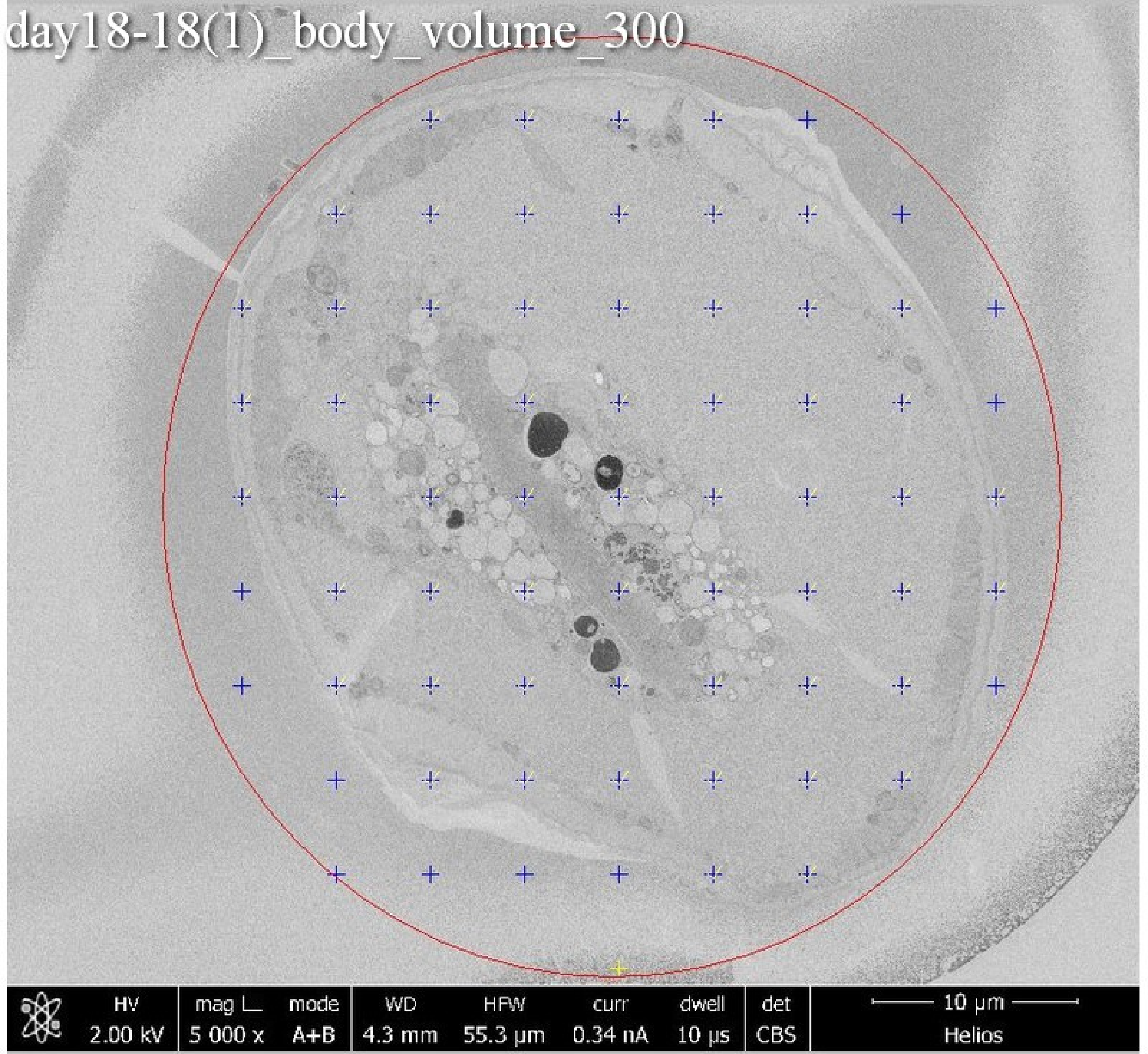

day18-18(1)_body_volume_300
HV
2.00 kV
mag
5 000 x
mode
A+B
WD
4.3 mm
HFW
55.3 µm
curr
0.34 nA
dwell
10 µs
det
CBS
10 µm
Helios

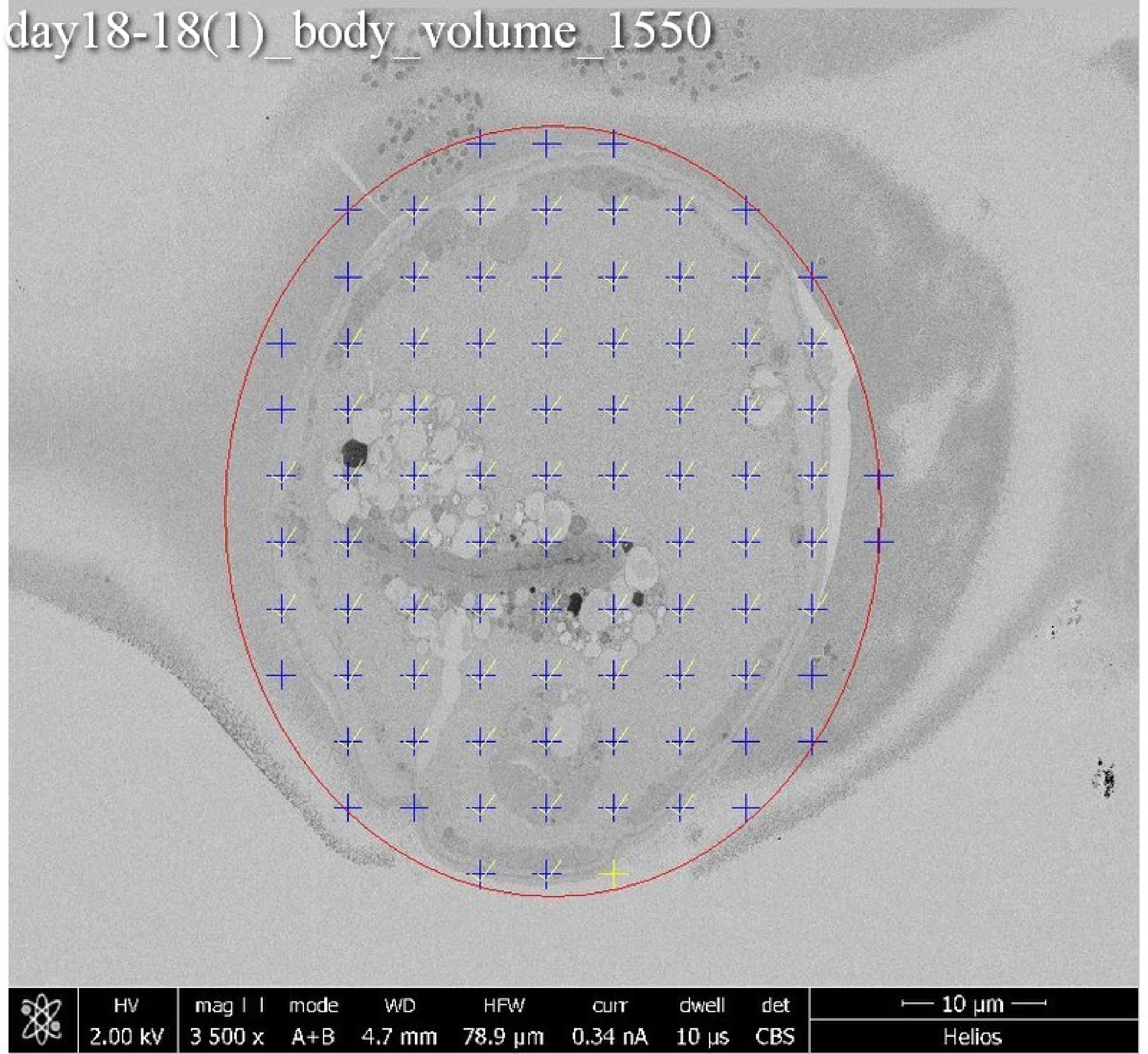
day18-18(1)_body_volume_1550
HV
2.00 kV
mag
3 500 x
mode
A+B
WD
4.7 mm
HFW
78.9 µm
curr
0.34 nA
dwell
10 µs
det
CBS
10 µm
Helios

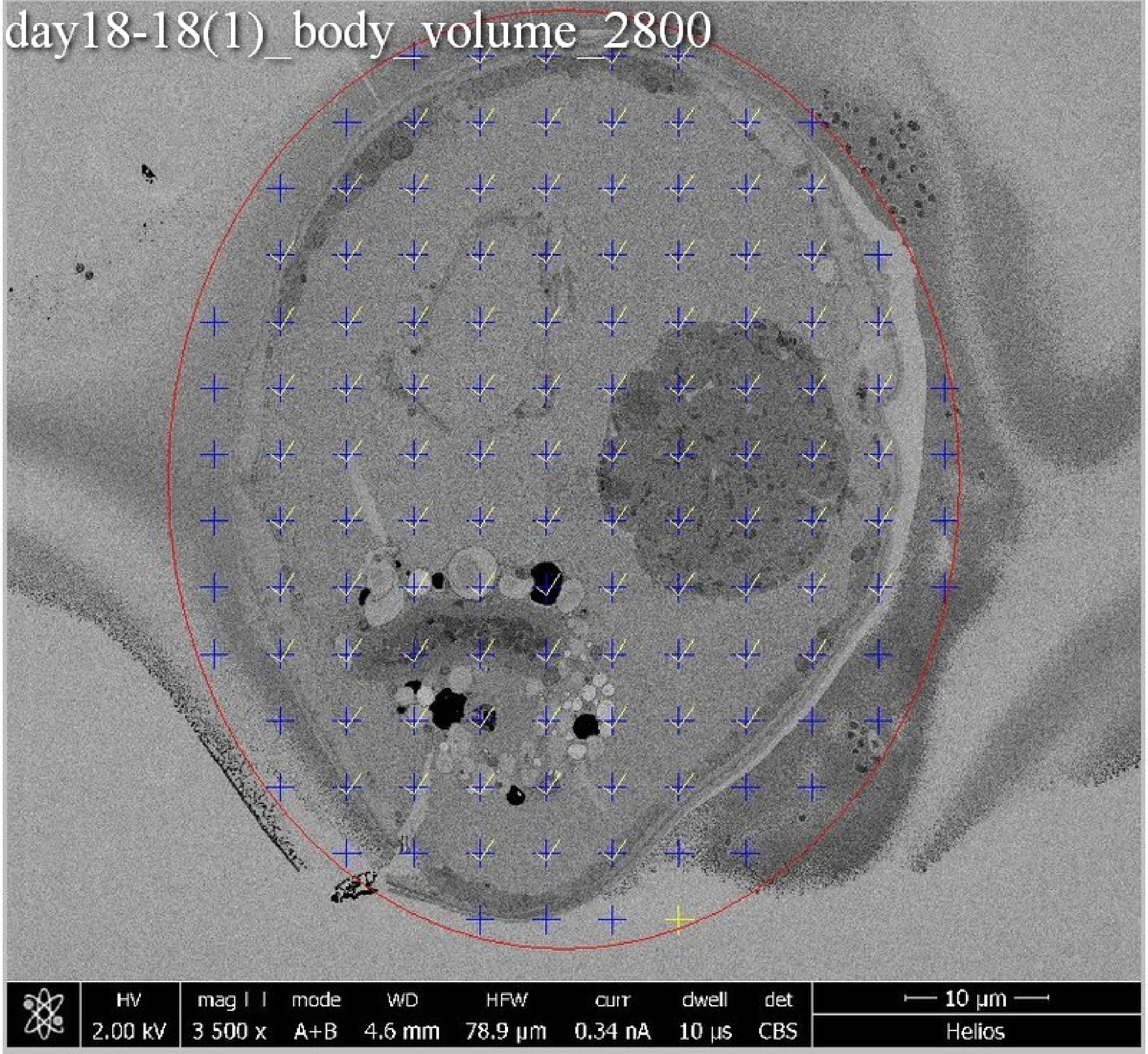
day18-18(1)_body_volume_2800
HV 2.00 kV
mag 3 500 x
mode A+B
WD 4.6 mm
HFW 78.9 µm
curr 0.34 nA
dwell 10 µs
det CBS
10 µm
Helios

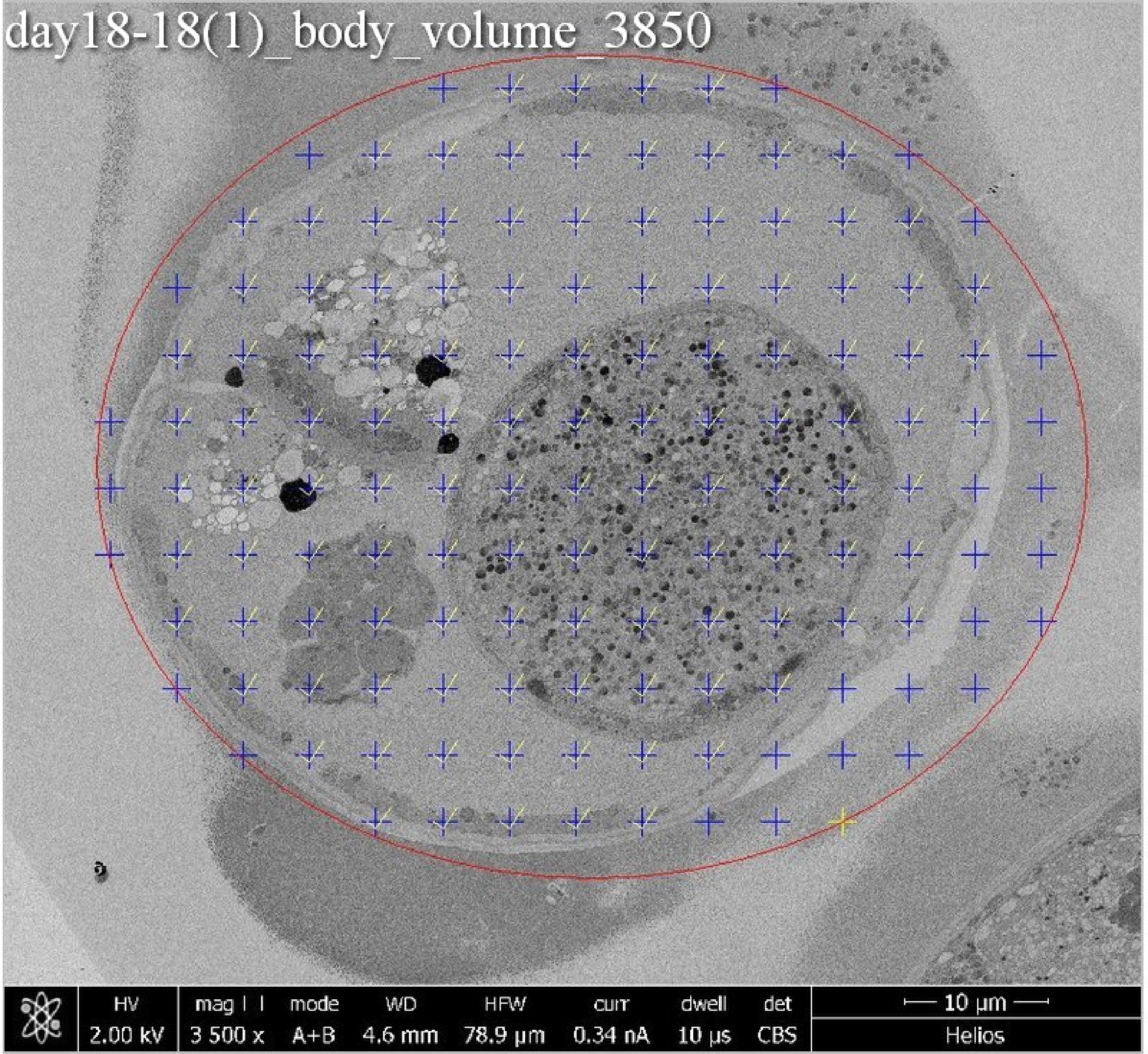
day18-18(1)_body_volume_3850
HV
2.00 kV
mag
3 500 x
mode
A+B
WD
4.6 mm
HFW
78.9 µm
curr
0.34 nA
dwell
10 µs
det
CBS
10 µm
Helios

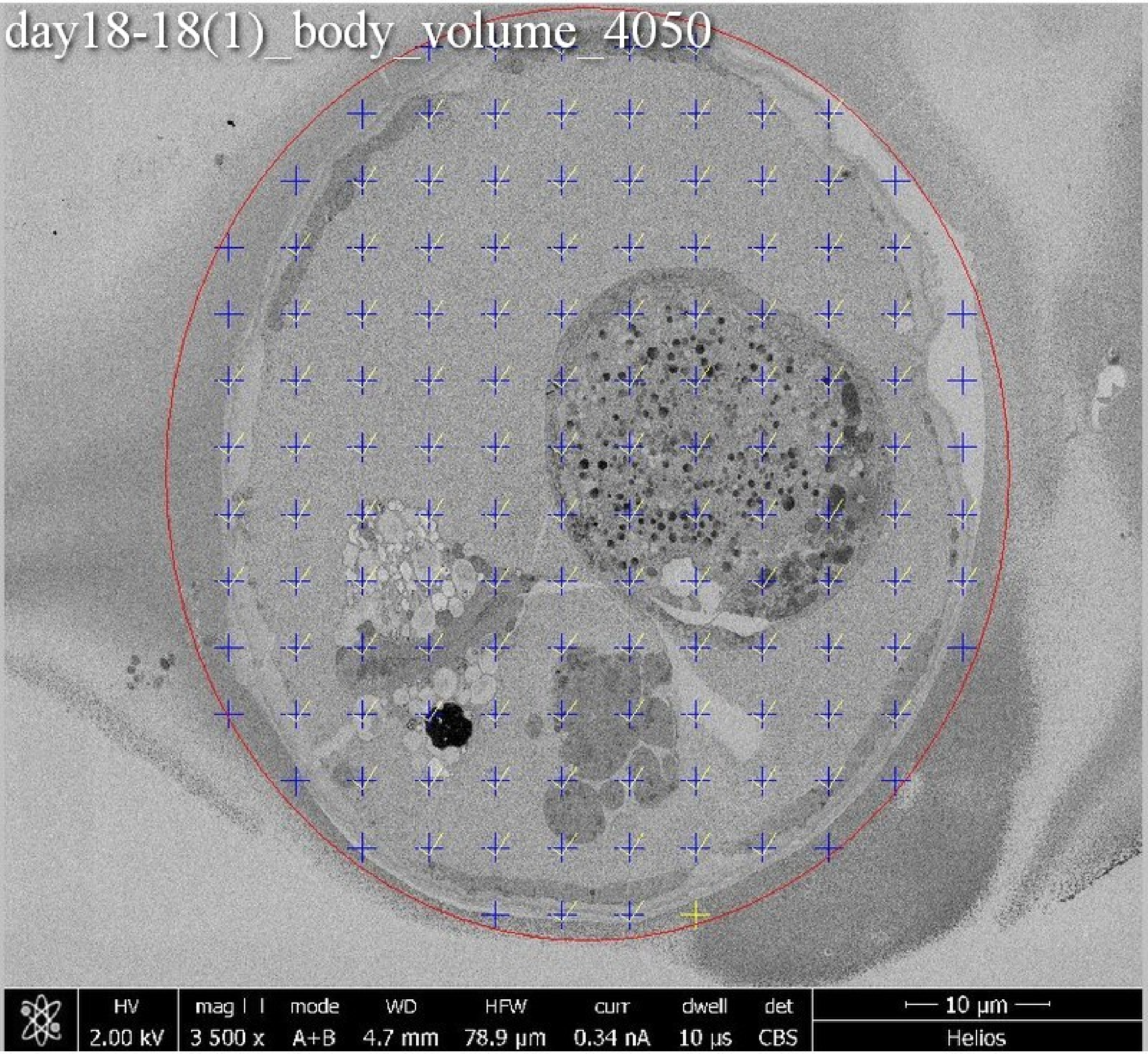
day18-18(1)_body_volume_4050
HV
2.00 kV
mag
3 500 x
mode
A+B
WD
4.7 mm
HFW
78.9 μm
curr
0.34 nA
dwell
10 μs
det
CBS
10 μm
Helios

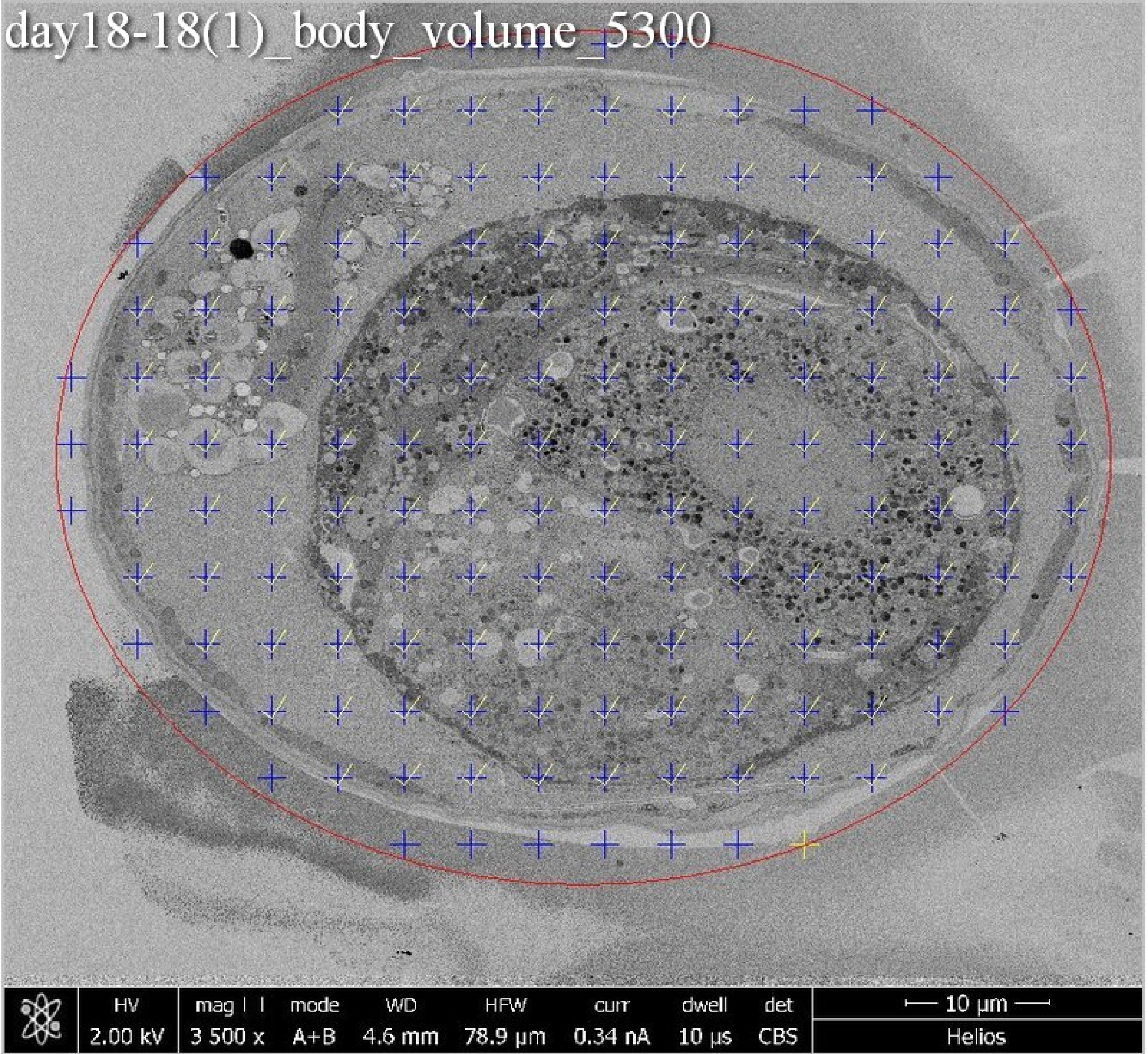
day18-18(1)_body_volume_5300
HV
2.00 kV
mag
3 500 x
mode
A+B
WD
4.6 mm
HFW
78.9 μm
curr
0.34 nA
dwell
10 μs
det
CBS
10 μm
Helios

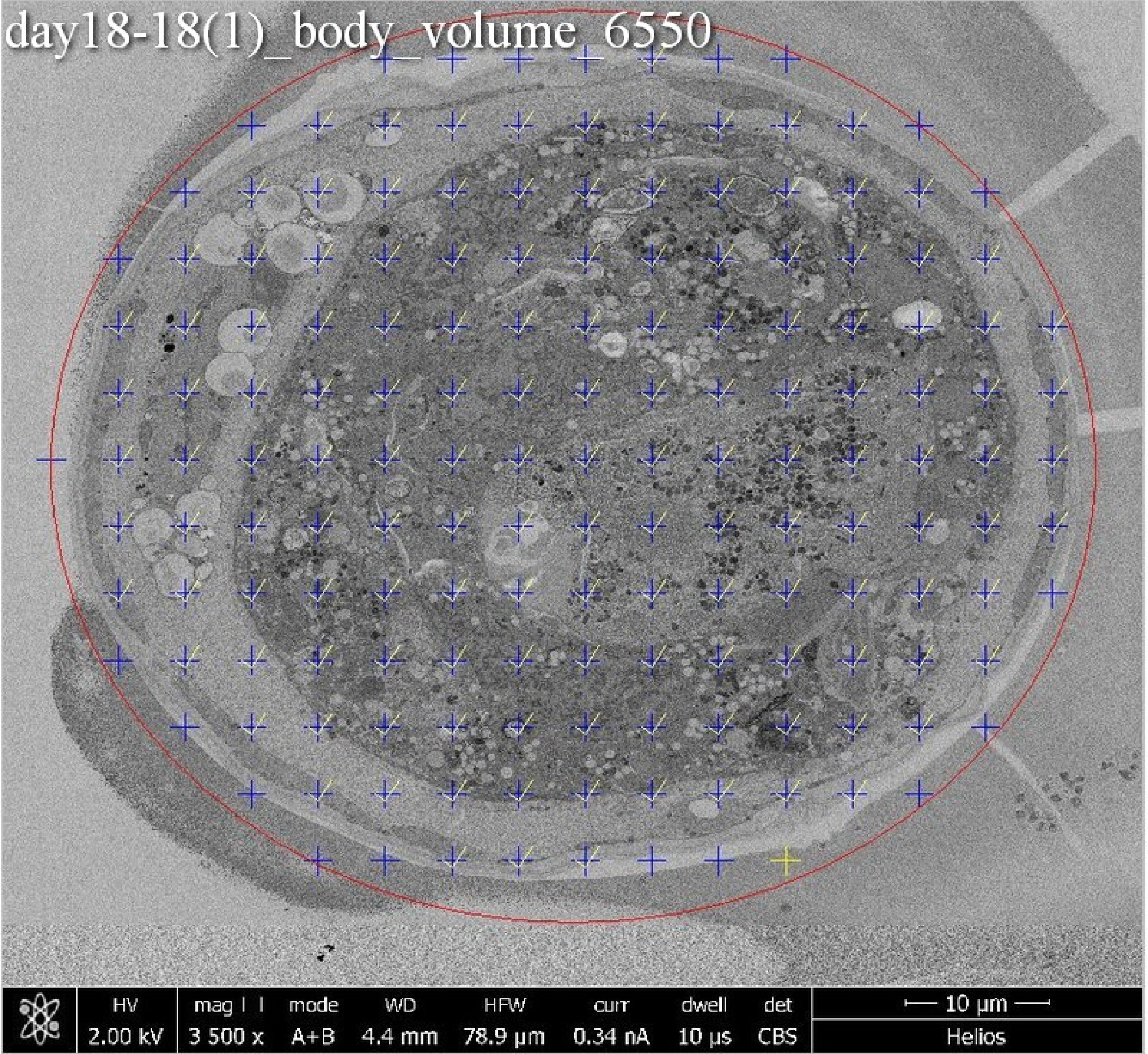
day18-18(1)_body_volume_6550
HV
2.00 kV
mag
3 500 x
mode
A+B
WD
4.4 mm
HFW
78.9 µm
curr
0.34 nA
dwell
10 µs
det
CBS
10 µm
Helios

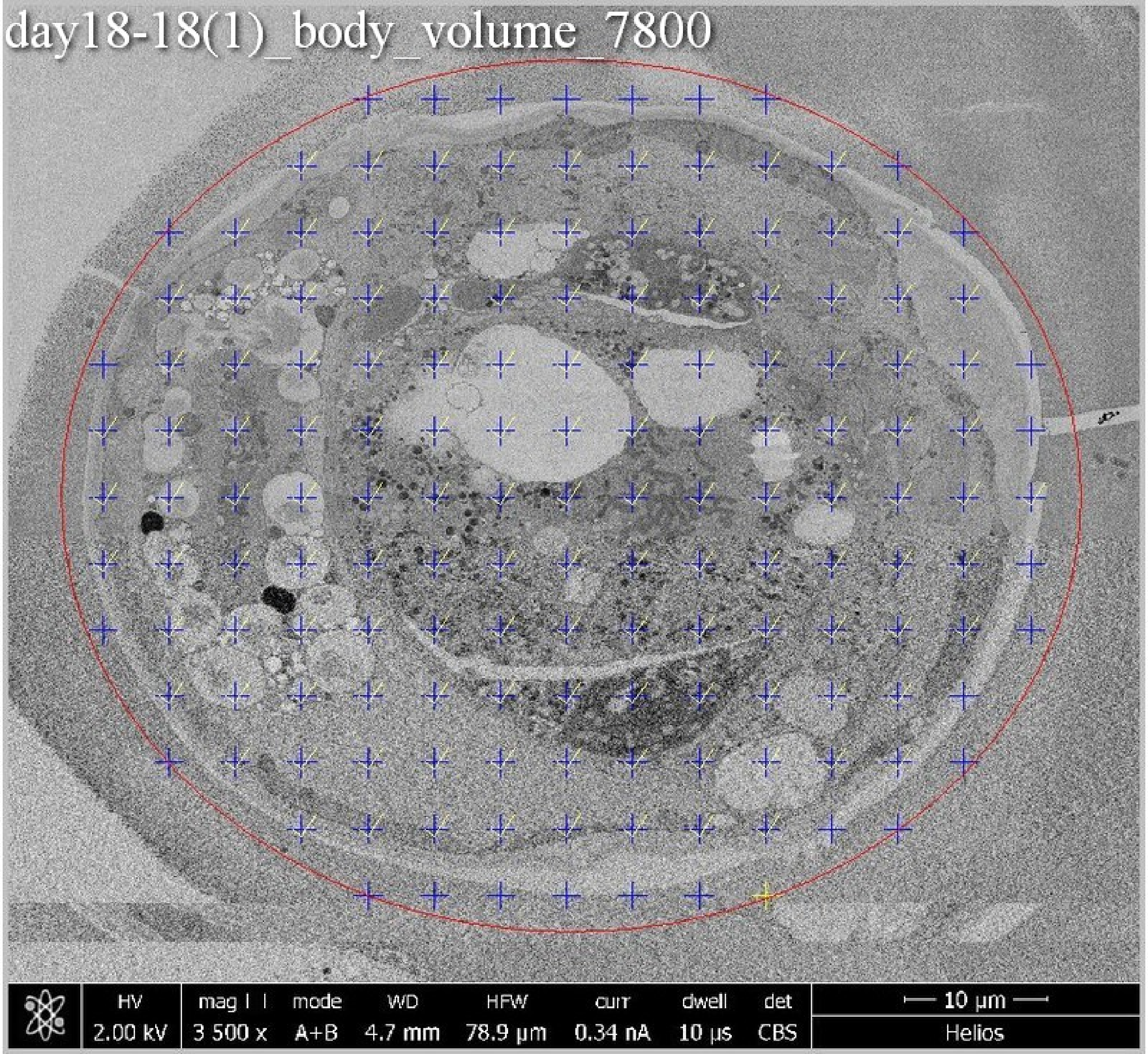

day18-18(1)_body_volume_7800
HV
2.00 kV
mag
3 500 x
mode
A+B
WD
4.7 mm
HFW
78.9 µm
curr
0.34 nA
dwell
10 µs
det
CBS
10 µm
Helios

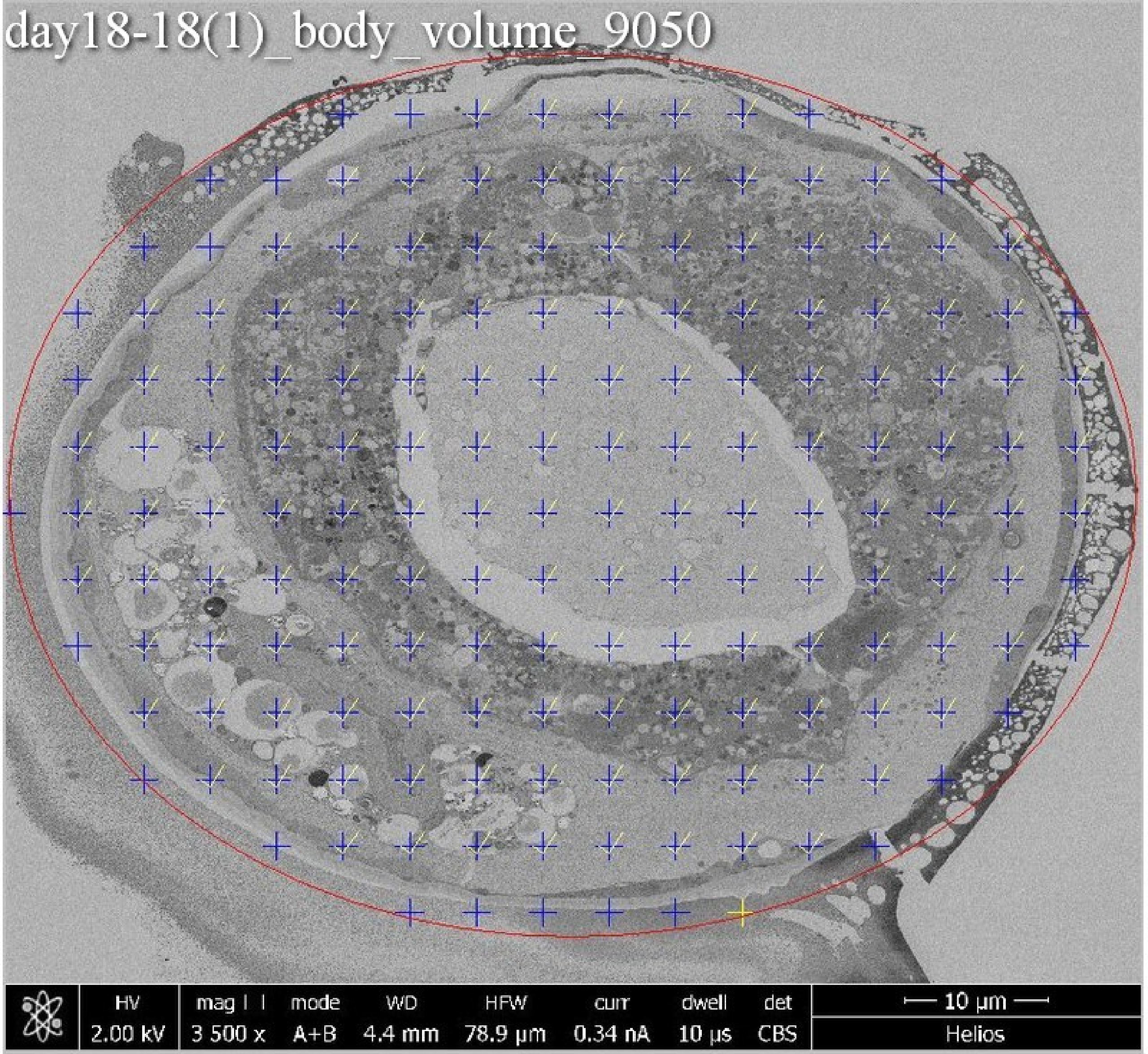
day18-18(1)_body_volume_9050
HV 2.00 kV
mag 3 500 x
mode A+B
WD 4.4 mm
HFW 78.9 μm
curr 0.34 nA
dwell 10 μs
det CBS
10 μm
Helios

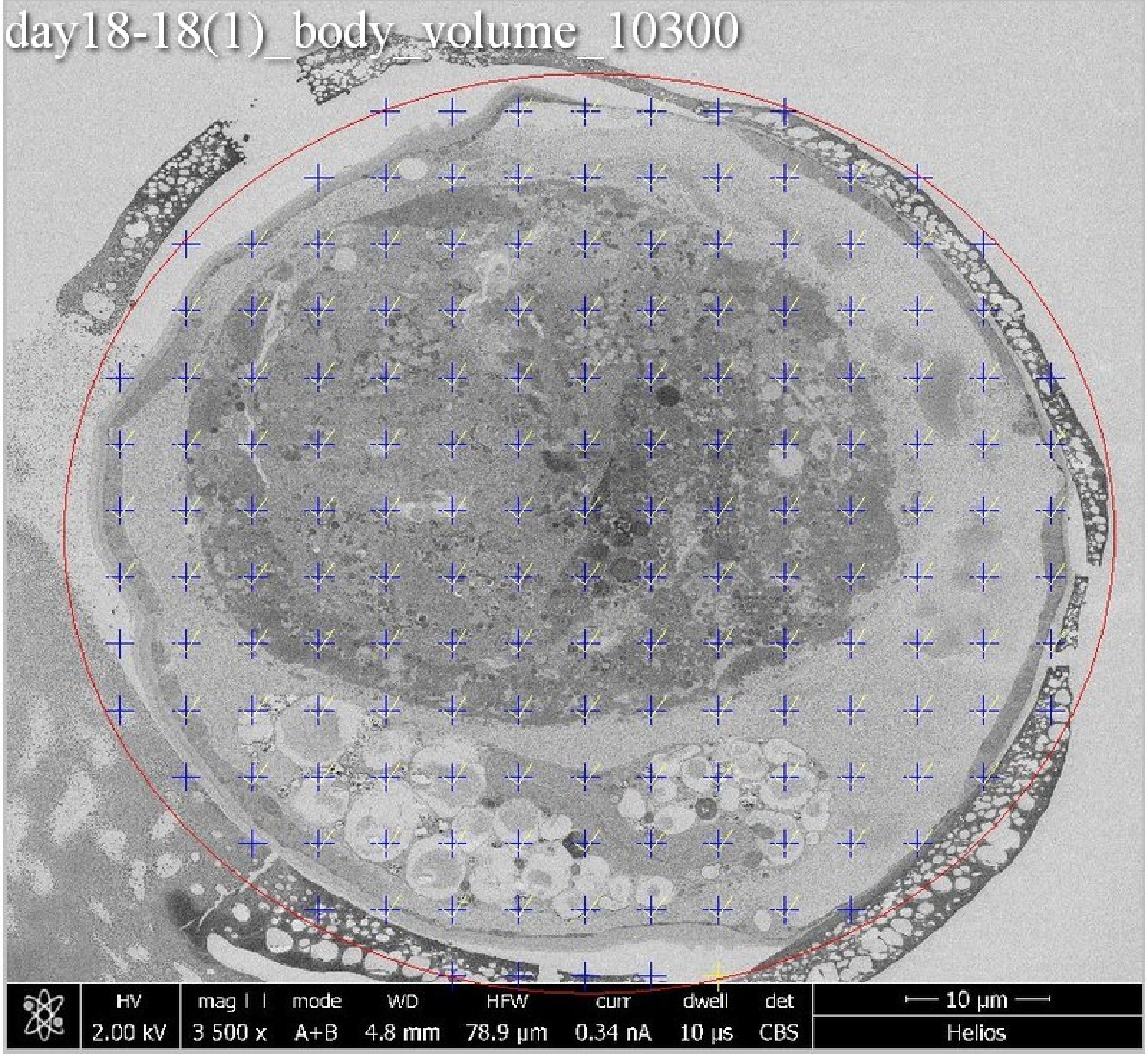
day18-18(1)_body_volume_10300
HV
2.00 kV
mag
3 500 x
mode
A+B
WD
4.8 mm
HFW
78.9 µm
curr
0.34 nA
dwell
10 µs
det
CBS
10 µm
Helios

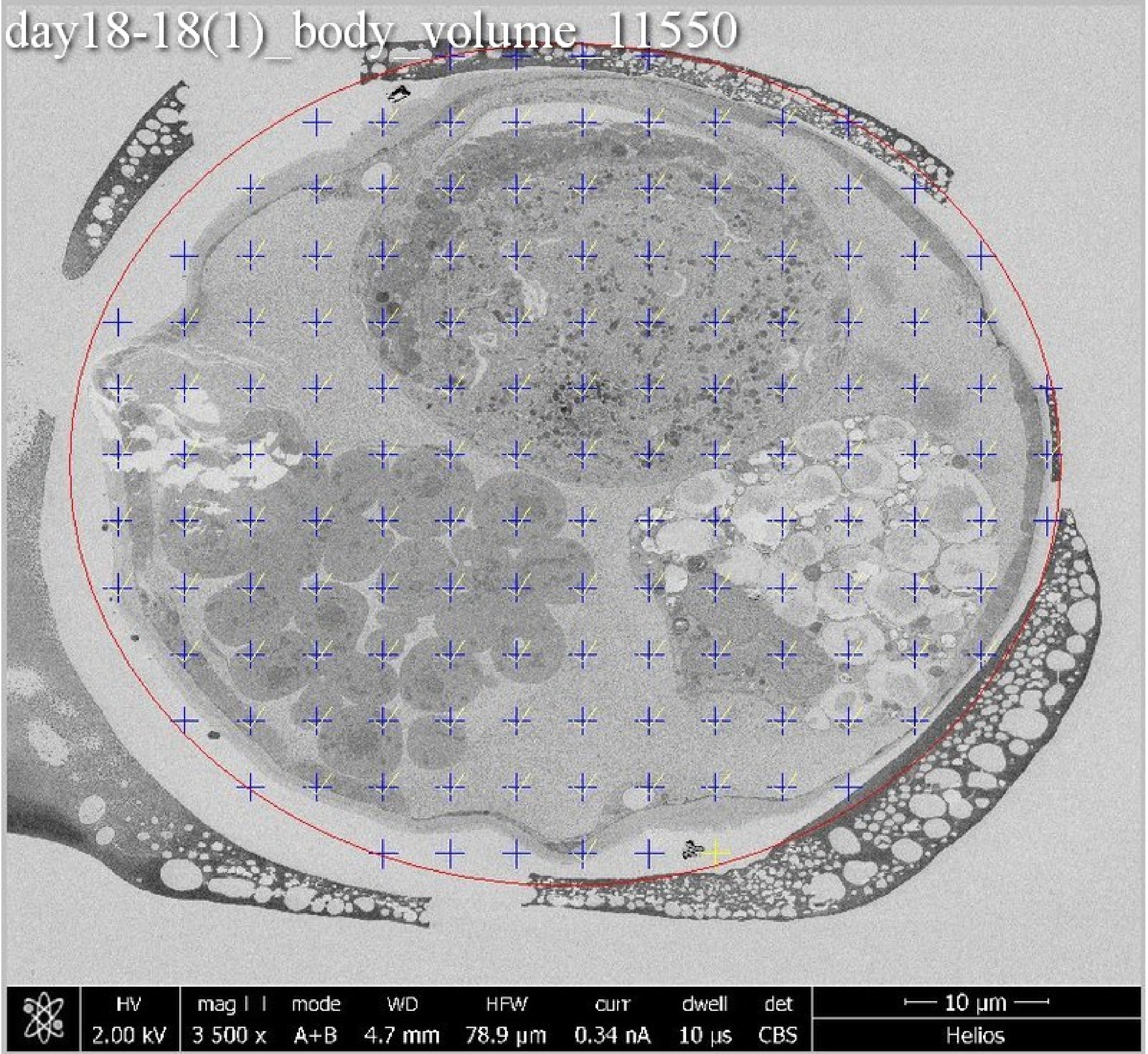
day18-18(1)_body_volume_11550
HV
2.00 kV
mag
3 500 x
mode
A+B
WD
4.7 mm
HFW
78.9 μm
curr
0.34 nA
dwell
10 μs
det
CBS
10 μm
Helios

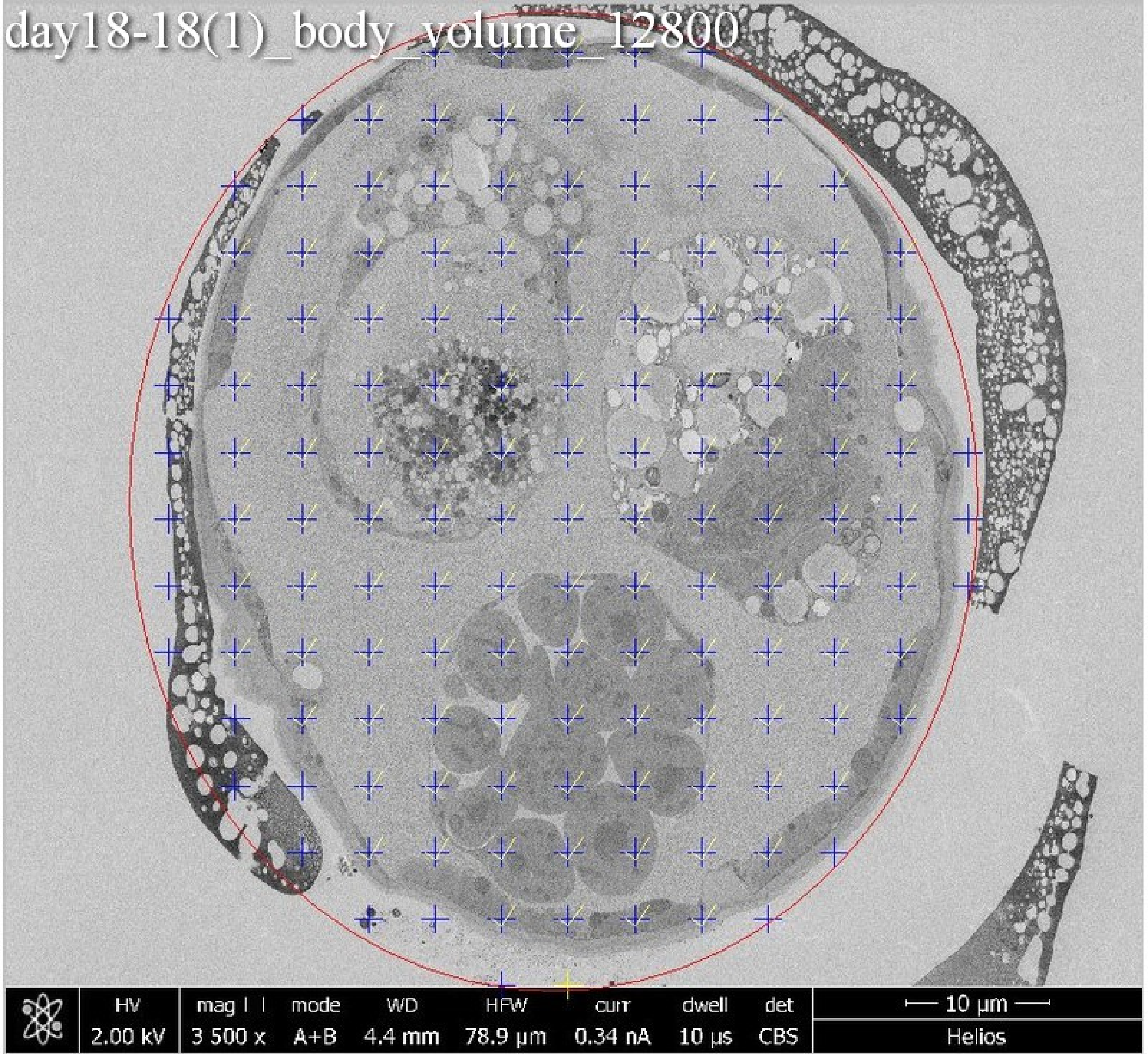

day18-18(1)_body_volume_12800
HV 2.00 kV
mag 3 500 x
mode A+B
WD 4.4 mm
HFW 78.9 μm
curr 0.34 nA
dwell 10 μs
det CBS
10 μm
Helios

day18-18(1)_body_volume_14900

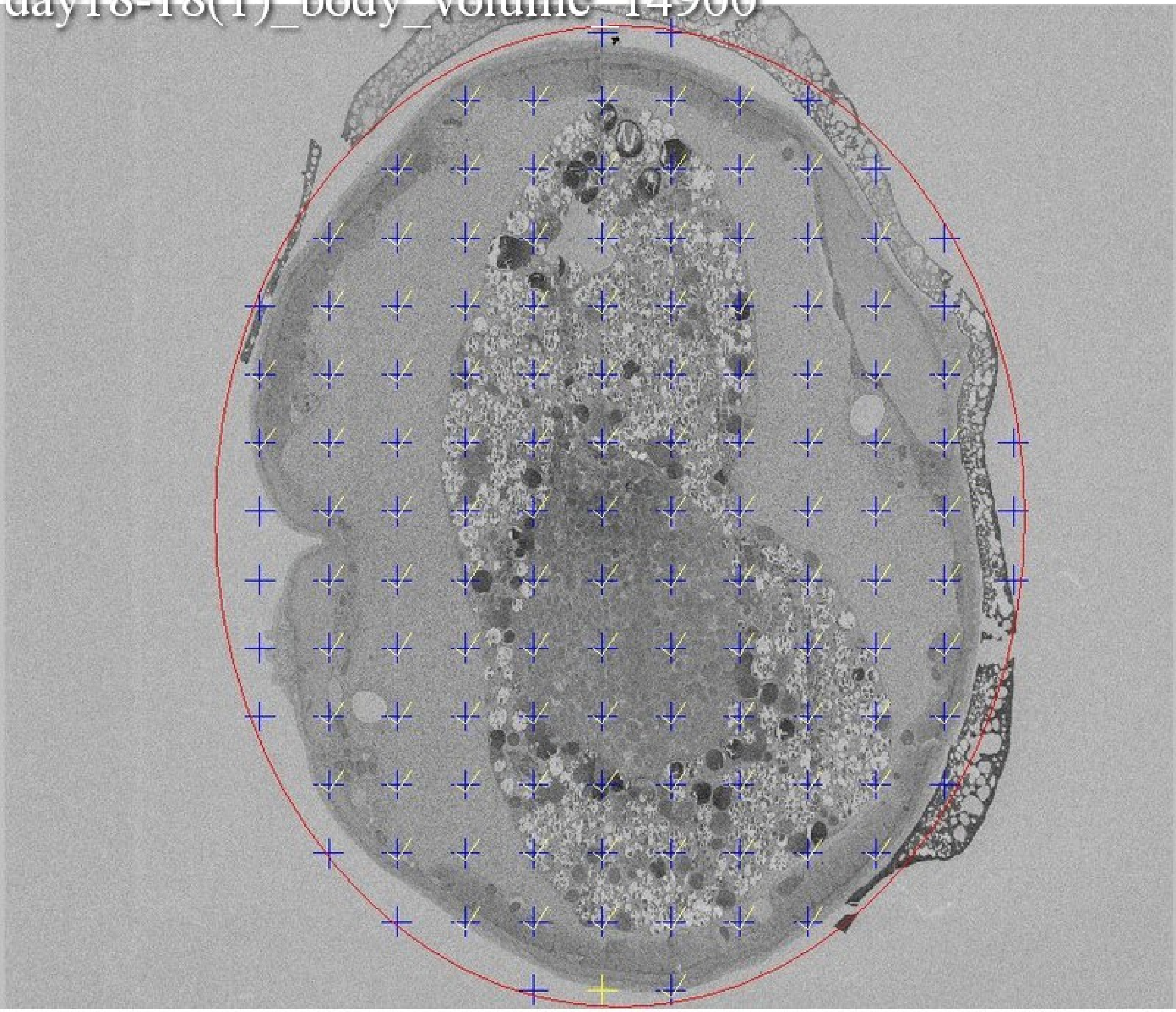

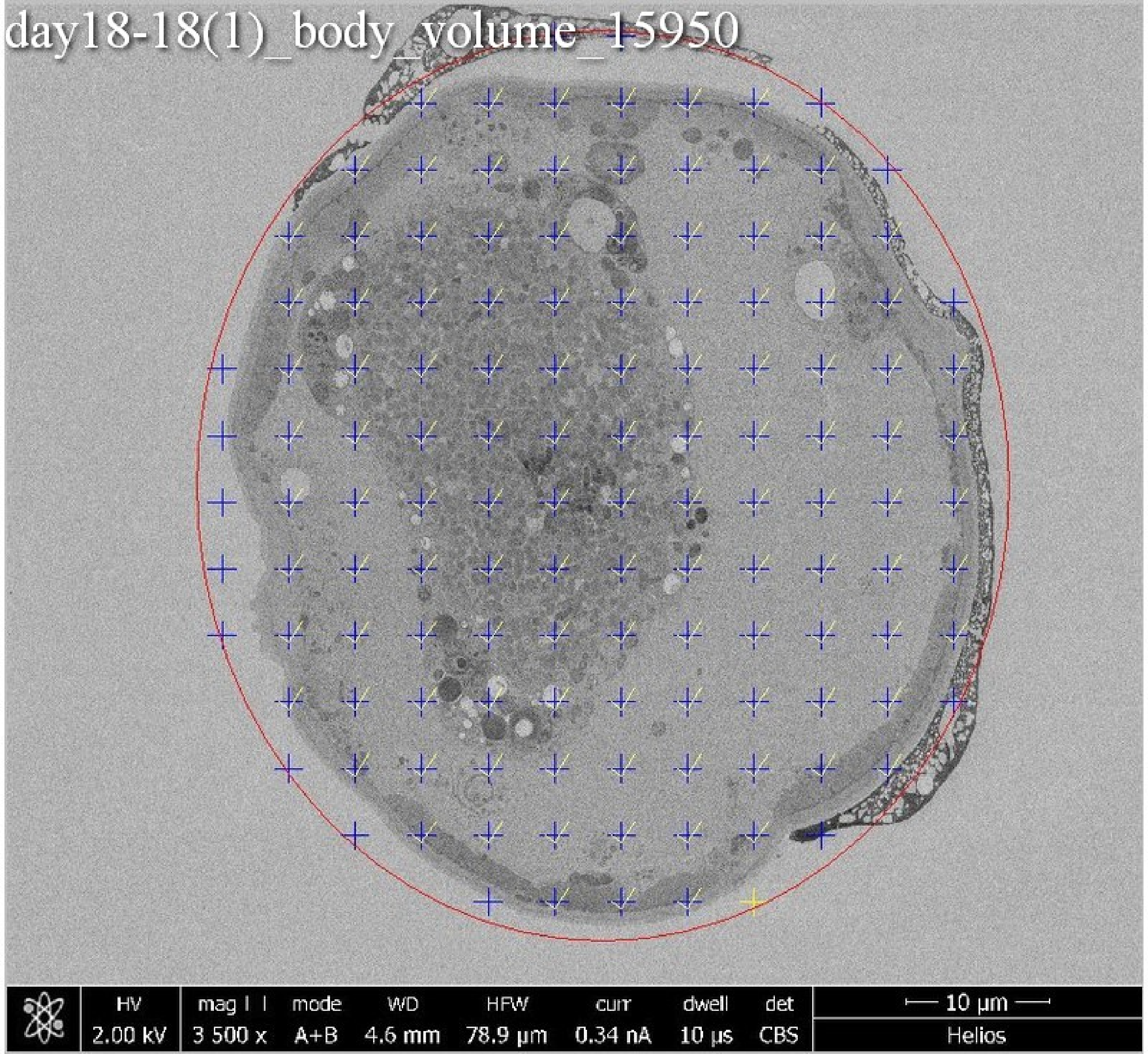
day18-18(1)_body_volume_15950
HV 2.00 kV
mag ⊡ 3 500 x
mode A+B
WD 4.6 mm
HFW 78.9 µm
curr 0.34 nA
dwell 10 µs
det CBS
10 µm
Helios

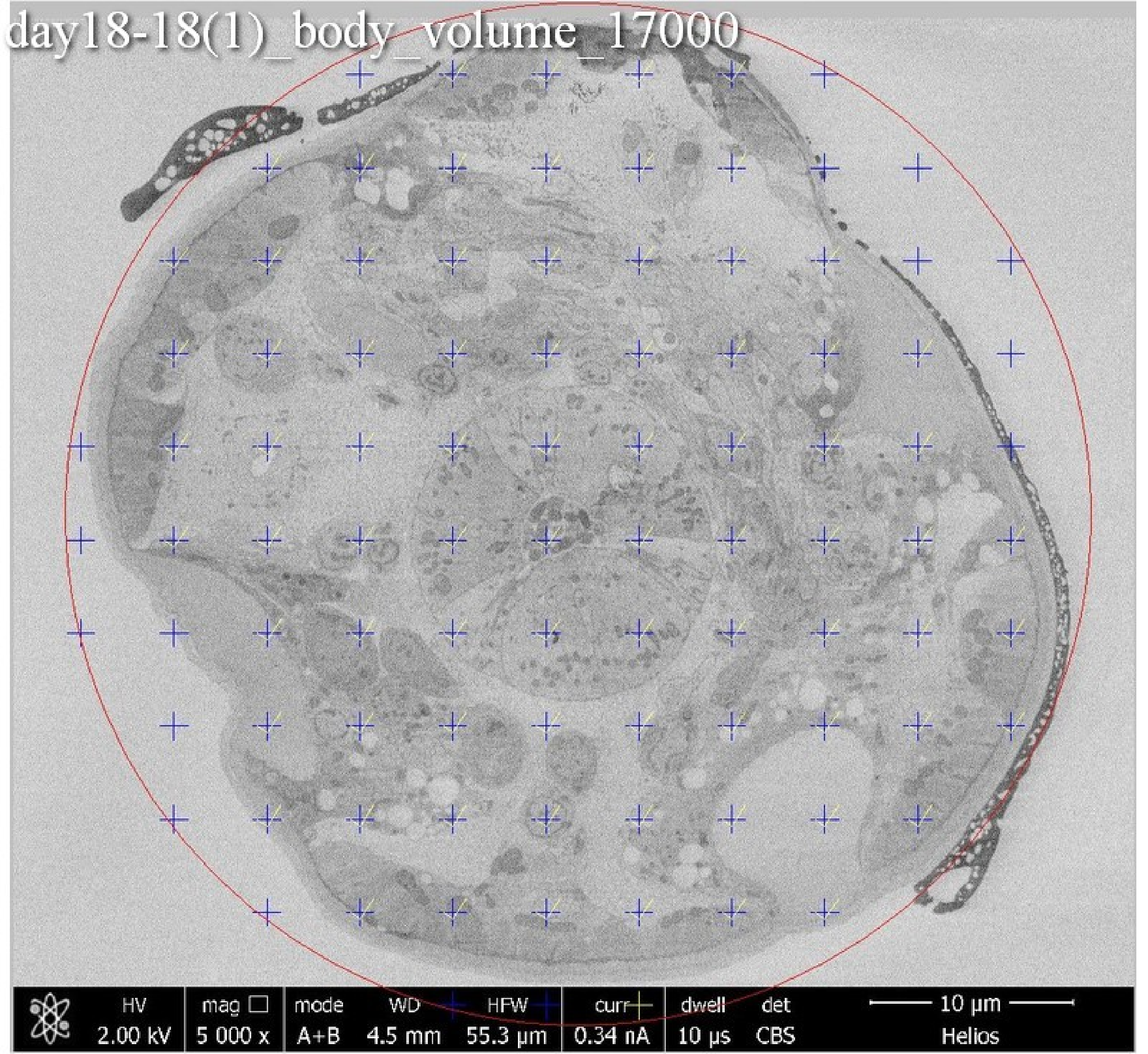
day18-18(1)_body_volume_17000
HV
2.00 kV
mag
5 000 x
mode
A+B
WD
4.5 mm
HFW
55.3 µm
curr
0.34 nA
dwell
10 µs
det
CBS
10 µm
Helios

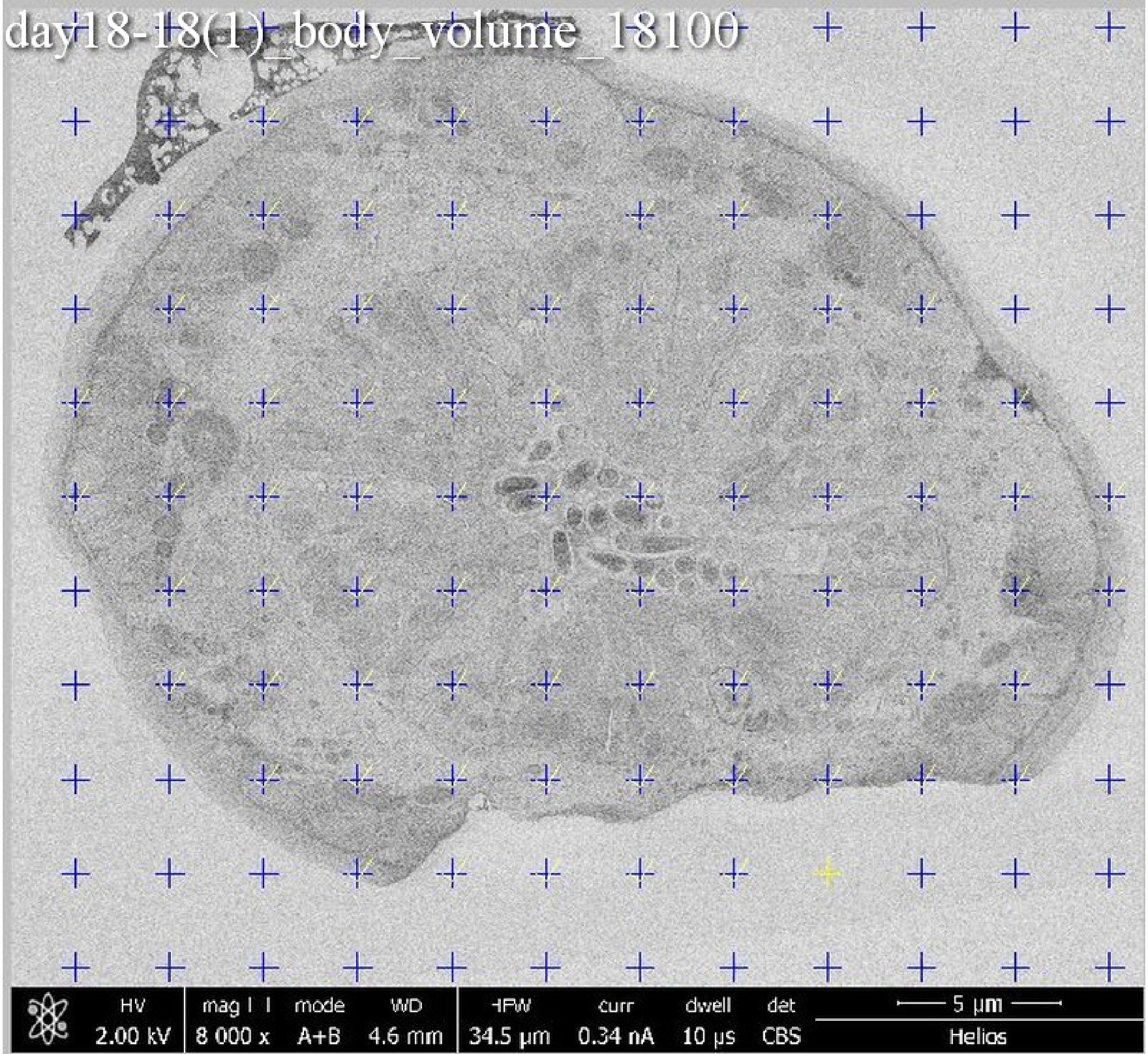
day18-18(1)_body_volume_18100
HV
2.00 kV
mag
8 000 x
mode
A+B
WD
4.6 mm
HFW
34.5 µm
curr
0.34 nA
dwell
10 µs
det
CBS
5 µm
Helios

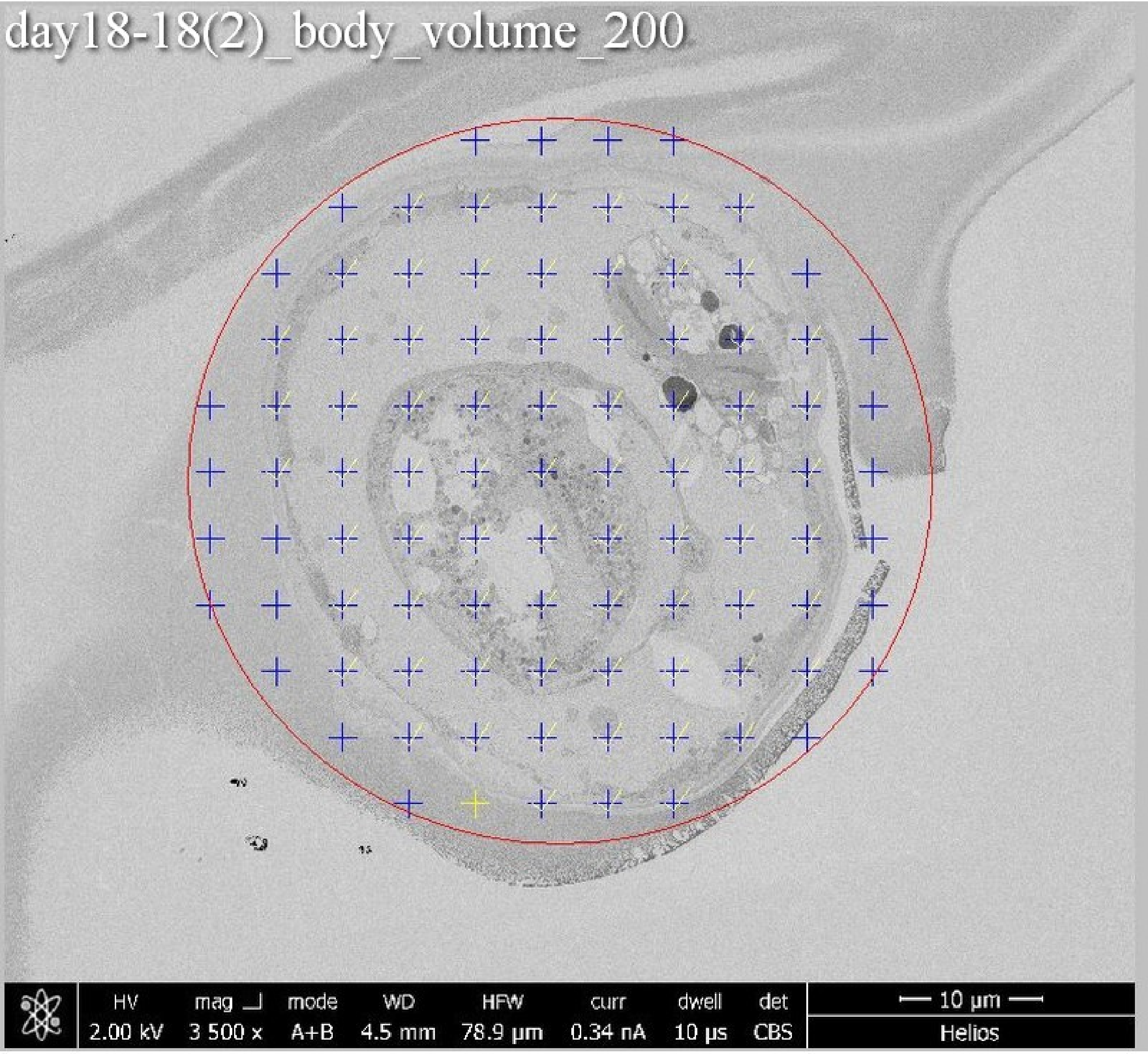

day18-18(2)_body_volume_200
HV 2.00 kV
mag 3 500 x
mode A+B
WD 4.5 mm
HFW 78.9 μm
curr 0.34 nA
dwell 10 μs
det CBS
10 μm
Helios

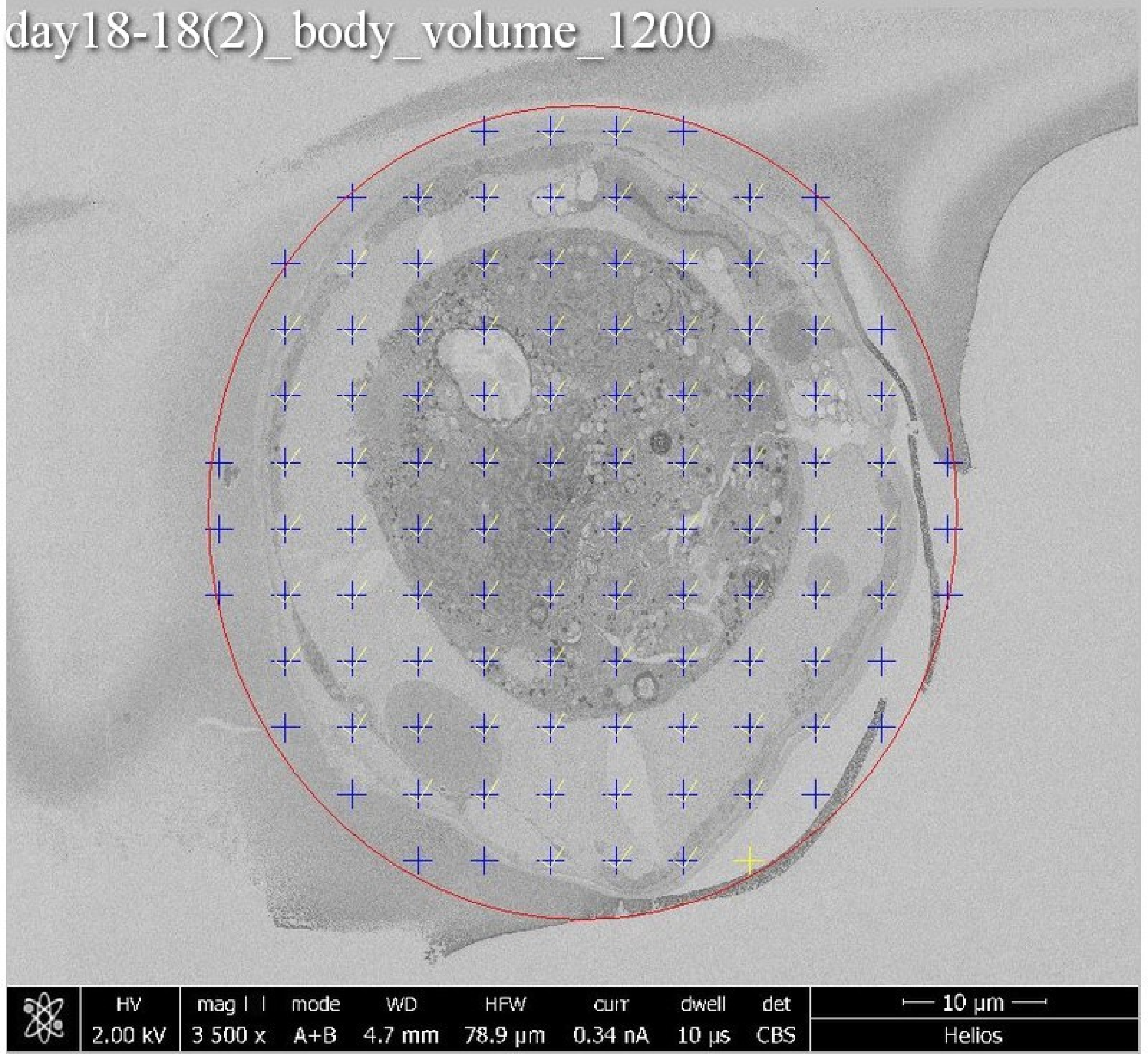
day18-18(2)_body_volume_1200
HV 2.00 kV
mag 3 500 x
mode A+B
WD 4.7 mm
HFW 78.9 µm
curr 0.34 nA
dwell 10 µs
det CBS
10 µm
Helios

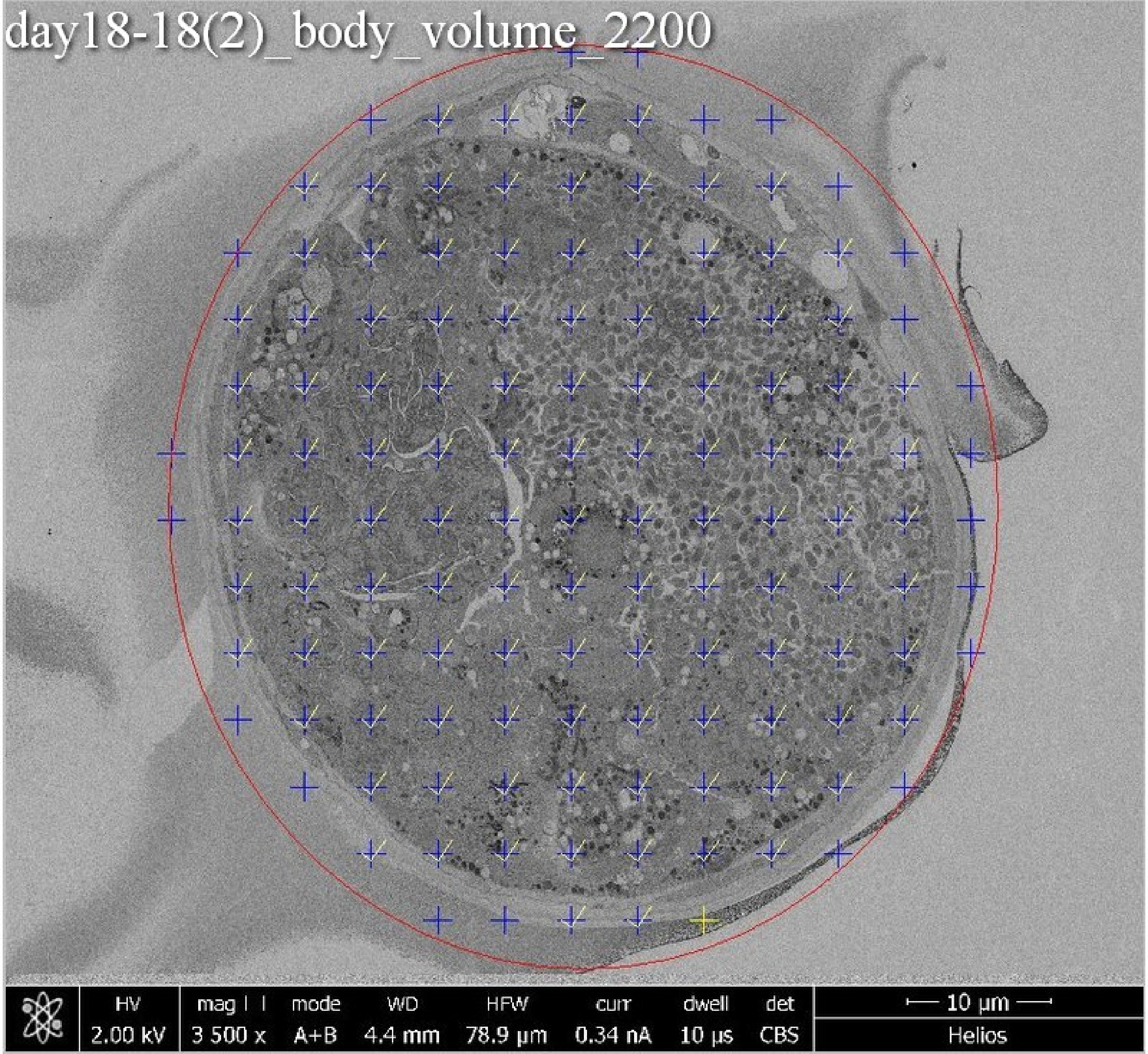

day18-18(2)_body_volume_2200
HV
2.00 kV
mag
3 500 x
mode
A+B
WD
4.4 mm
HFW
78.9 μm
curr
0.34 nA
dwell
10 μs
det
CBS
10 μm
Helios

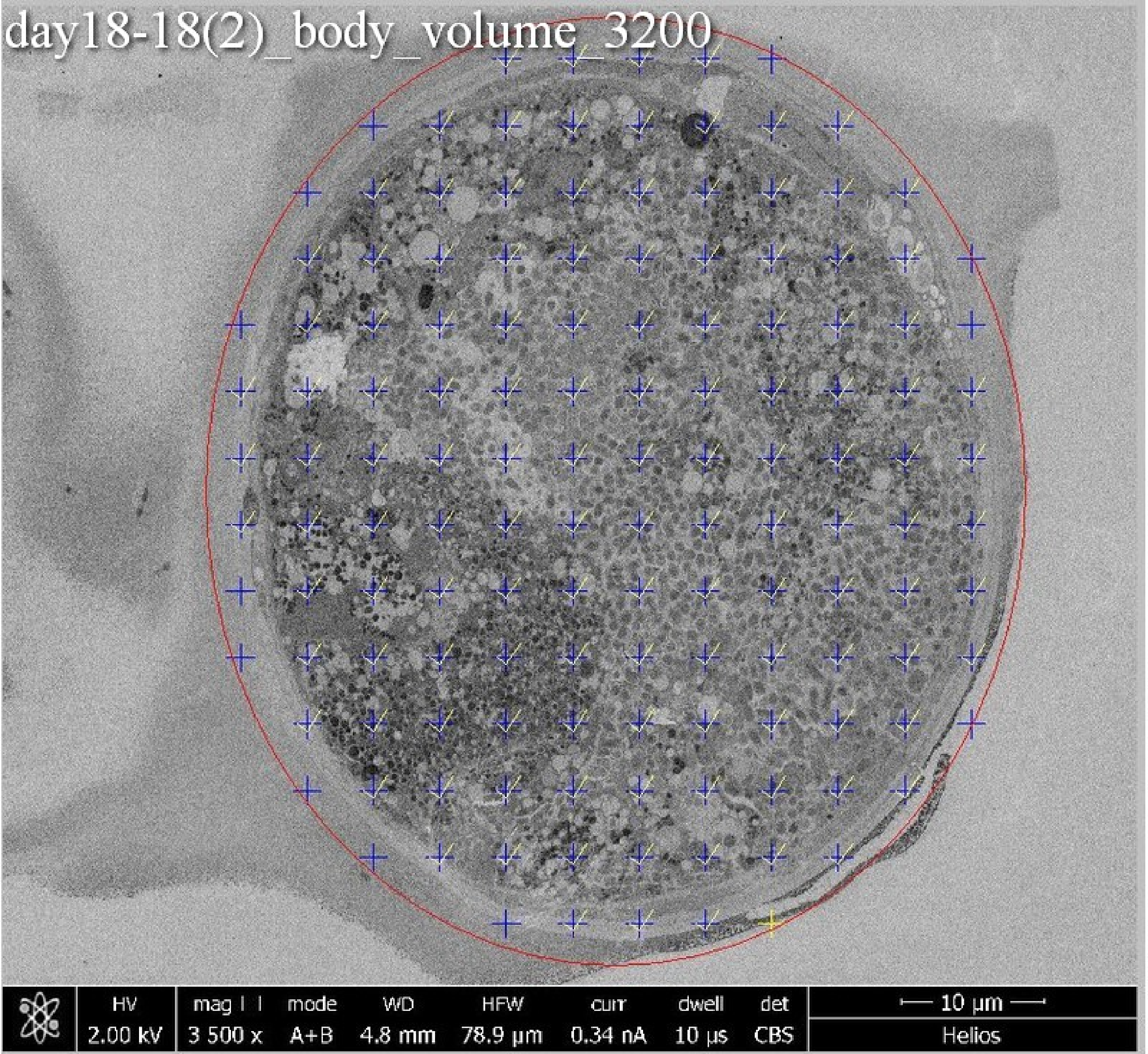
day18-18(2)_body_volume_3200
HV 2.00 kV
mag 3 500 x
mode A+B
WD 4.8 mm
HFW 78.9 µm
curr 0.34 nA
dwell 10 µs
det CBS
10 µm
Helios

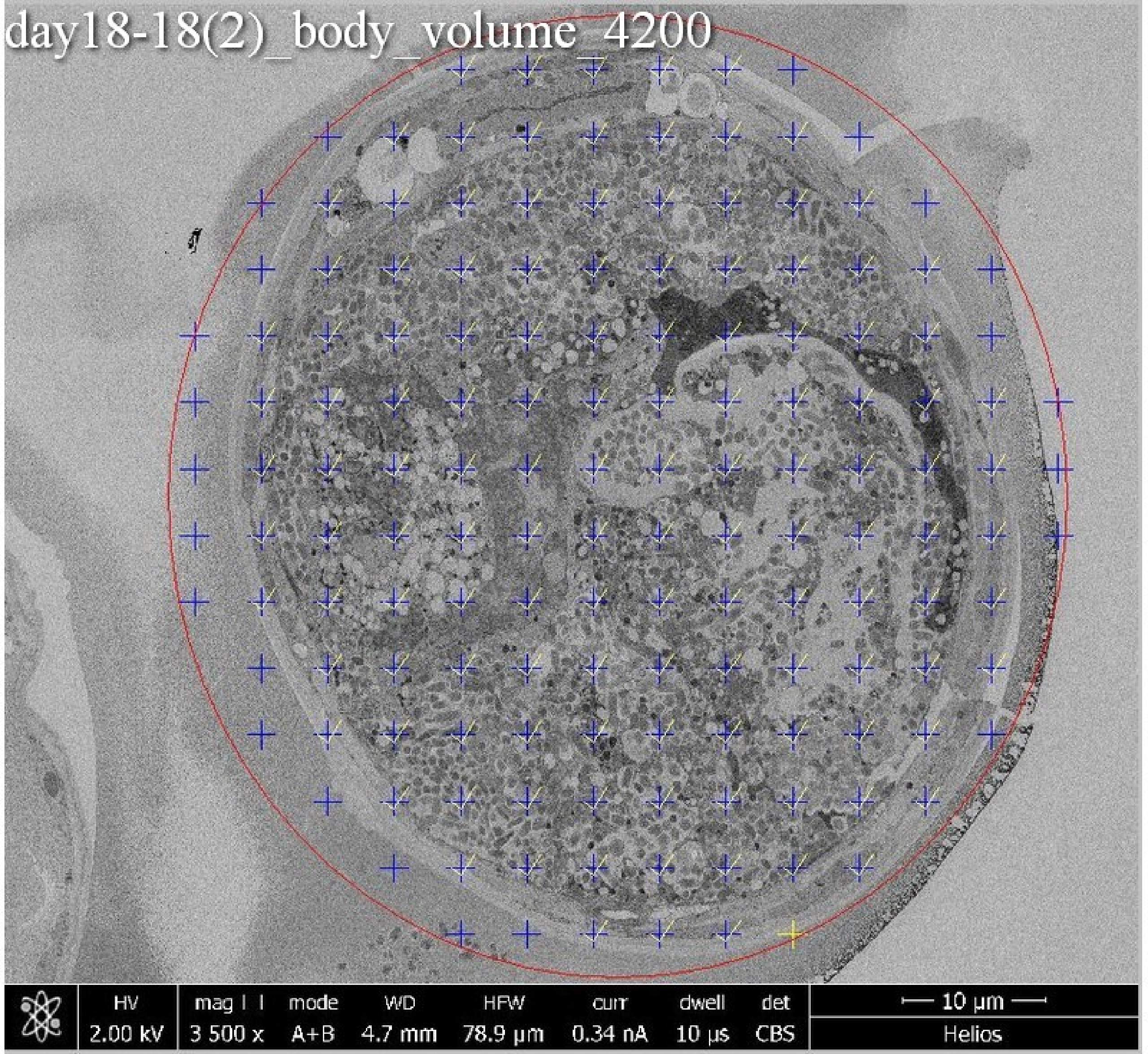

day18-18(2)_body_volume_4200
HV
2.00 kV
mag
3 500 x
mode
A+B
WD
4.7 mm
HFW
78.9 µm
curr
0.34 nA
dwell
10 µs
det
CBS
10 µm
Helios

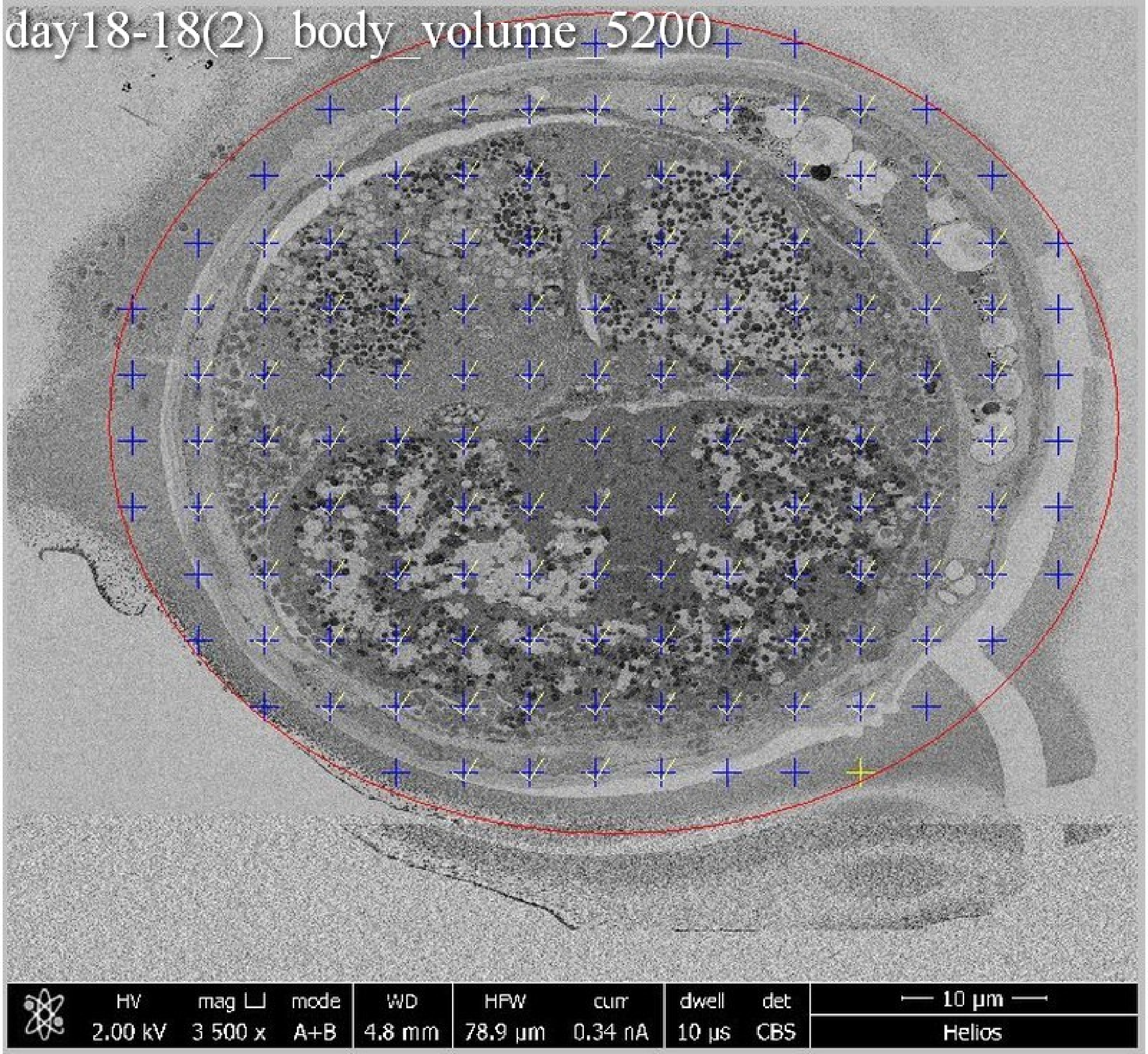
day18-18(2)_body_volume_5200
HV 2.00 kV
mag ⊔ 3 500 x
mode A+B
WD 4.8 mm
HFW 78.9 µm
curr 0.34 nA
dwell 10 µs
det CBS
10 µm
Helios

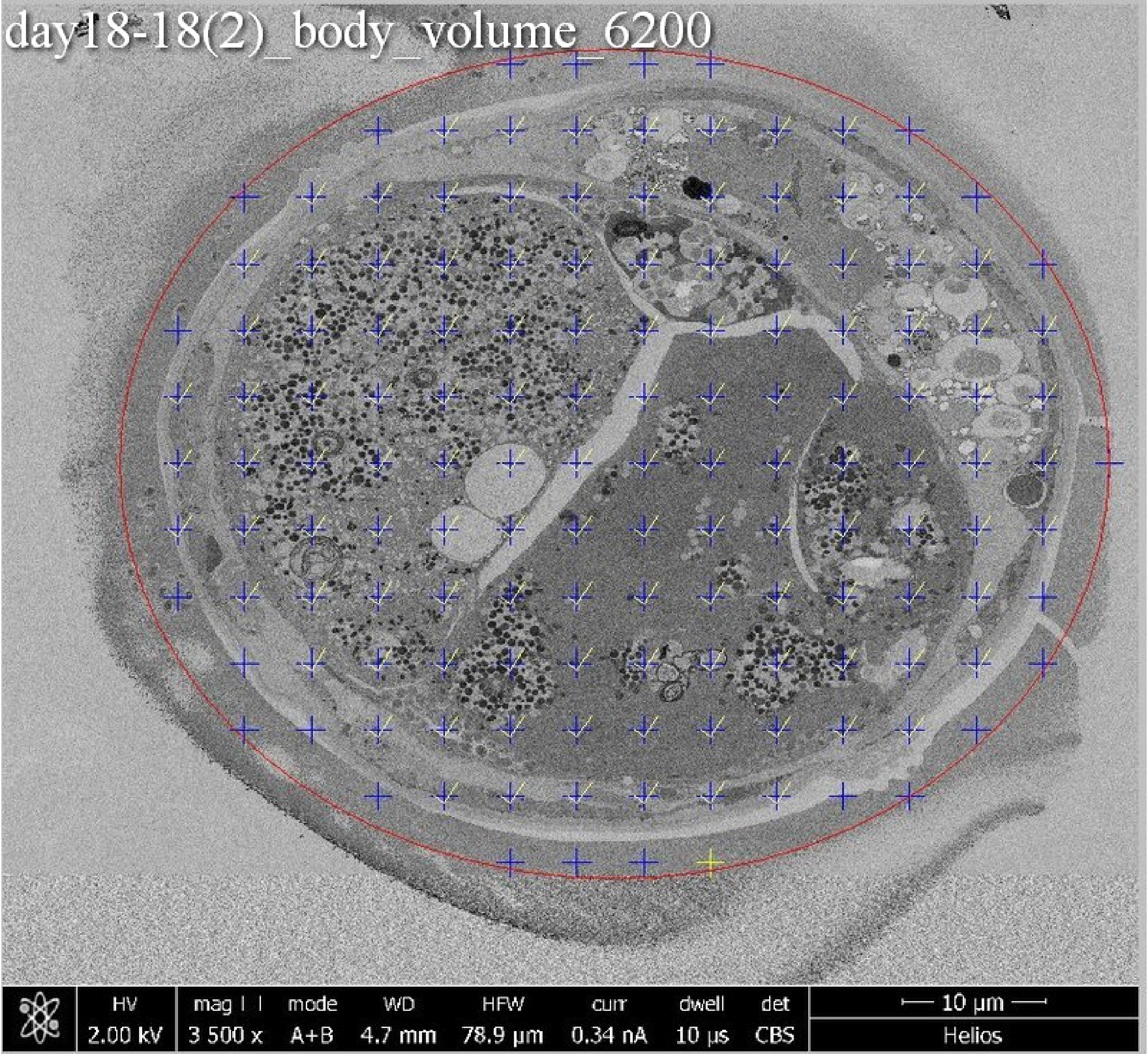
day18-18(2)_body_volume_6200
HV
2.00 kV
mag
3 500 x
mode
A+B
WD
4.7 mm
HFW
78.9 µm
curr
0.34 nA
dwell
10 µs
det
CBS
10 µm
Helios

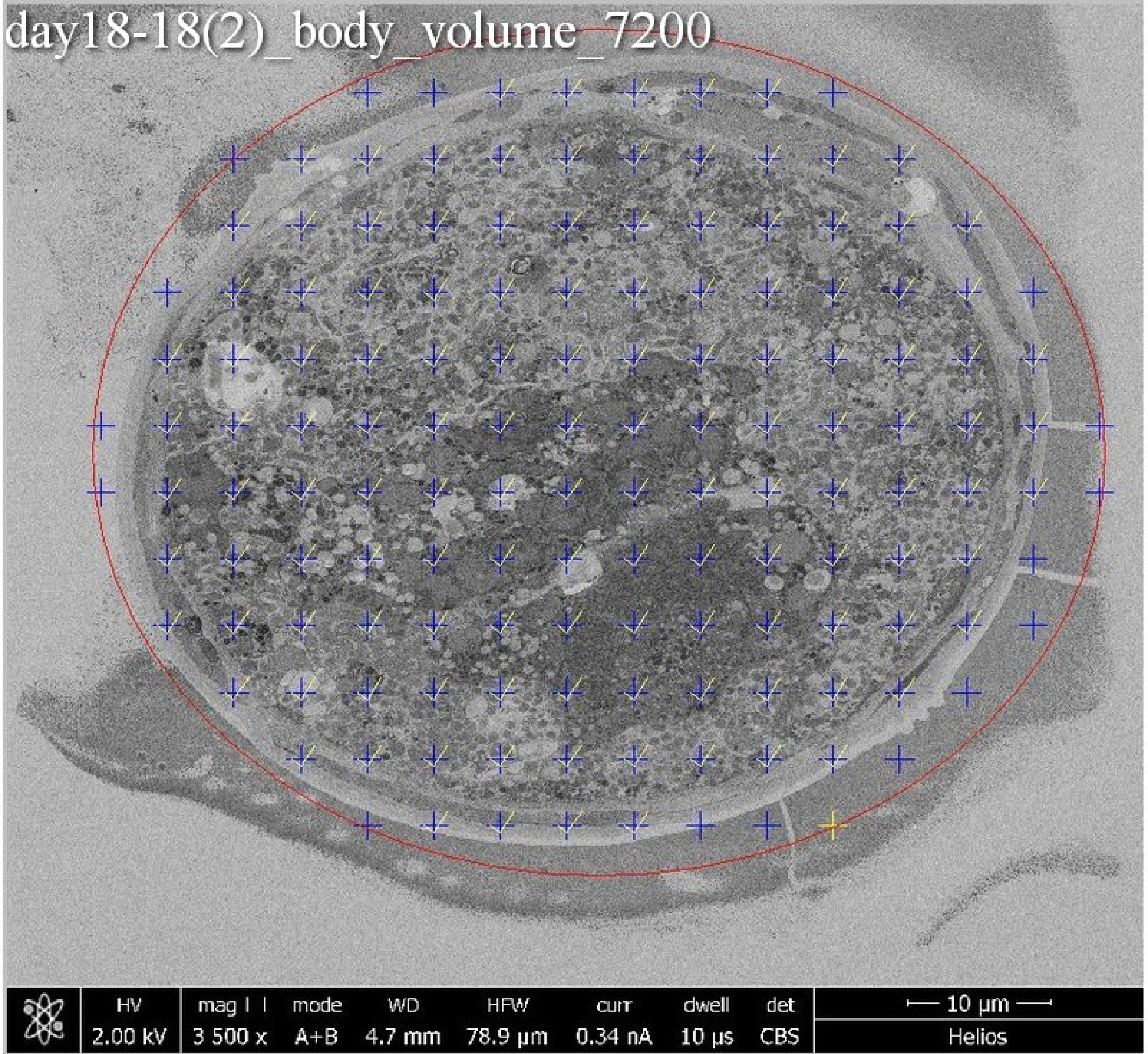
day18-18(2)_body_volume_7200
HV
2.00 kV
mag
3 500 x
mode
A+B
WD
4.7 mm
HFW
78.9 µm
curr
0.34 nA
dwell
10 µs
det
CBS
10 µm
Helios

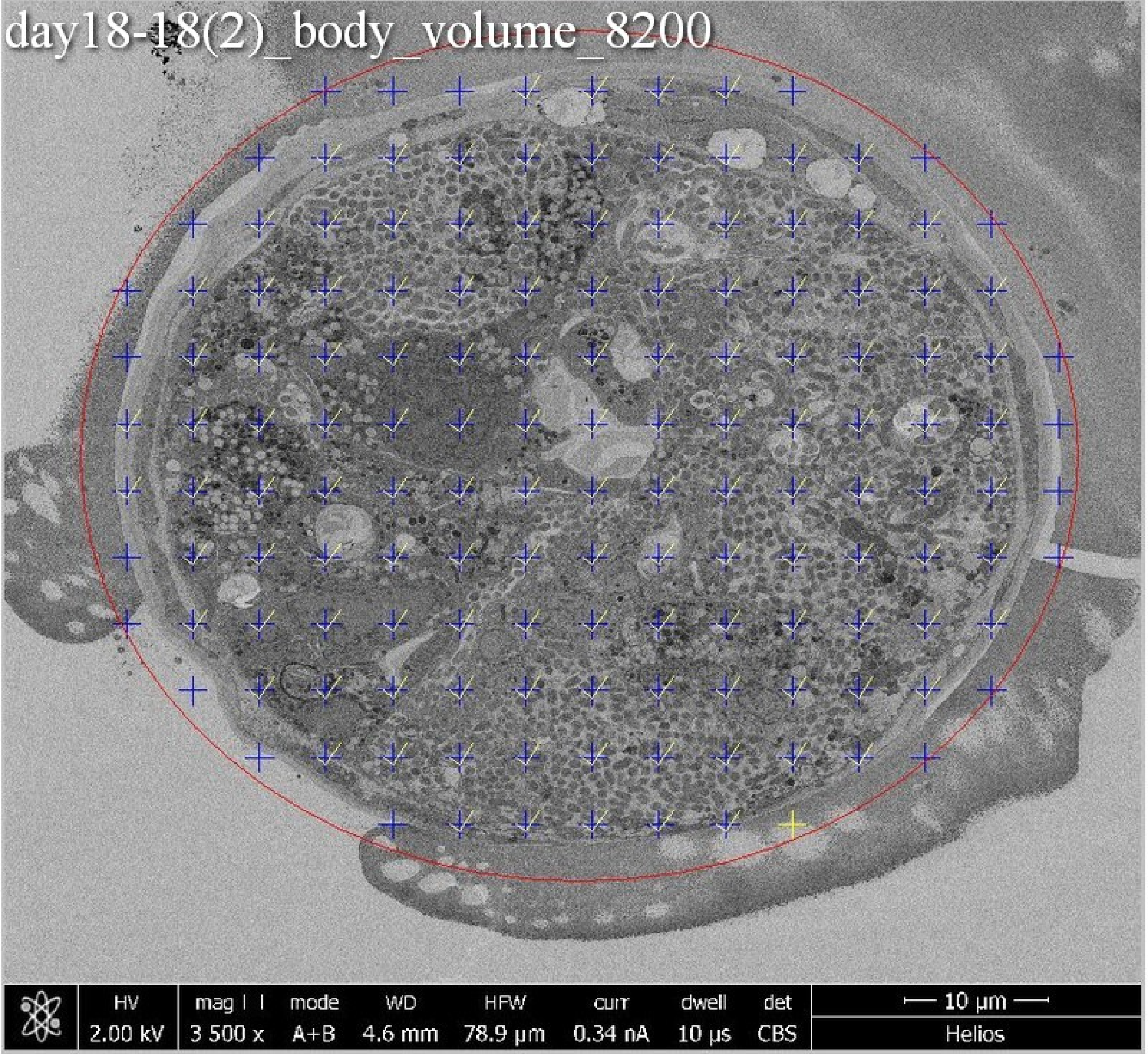

day18-18(2)_body_volume_8200
HV
2.00 kV
mag
3 500 x
mode
A+B
WD
4.6 mm
HFW
78.9 µm
curr
0.34 nA
dwell
10 µs
det
CBS
10 µm
Helios

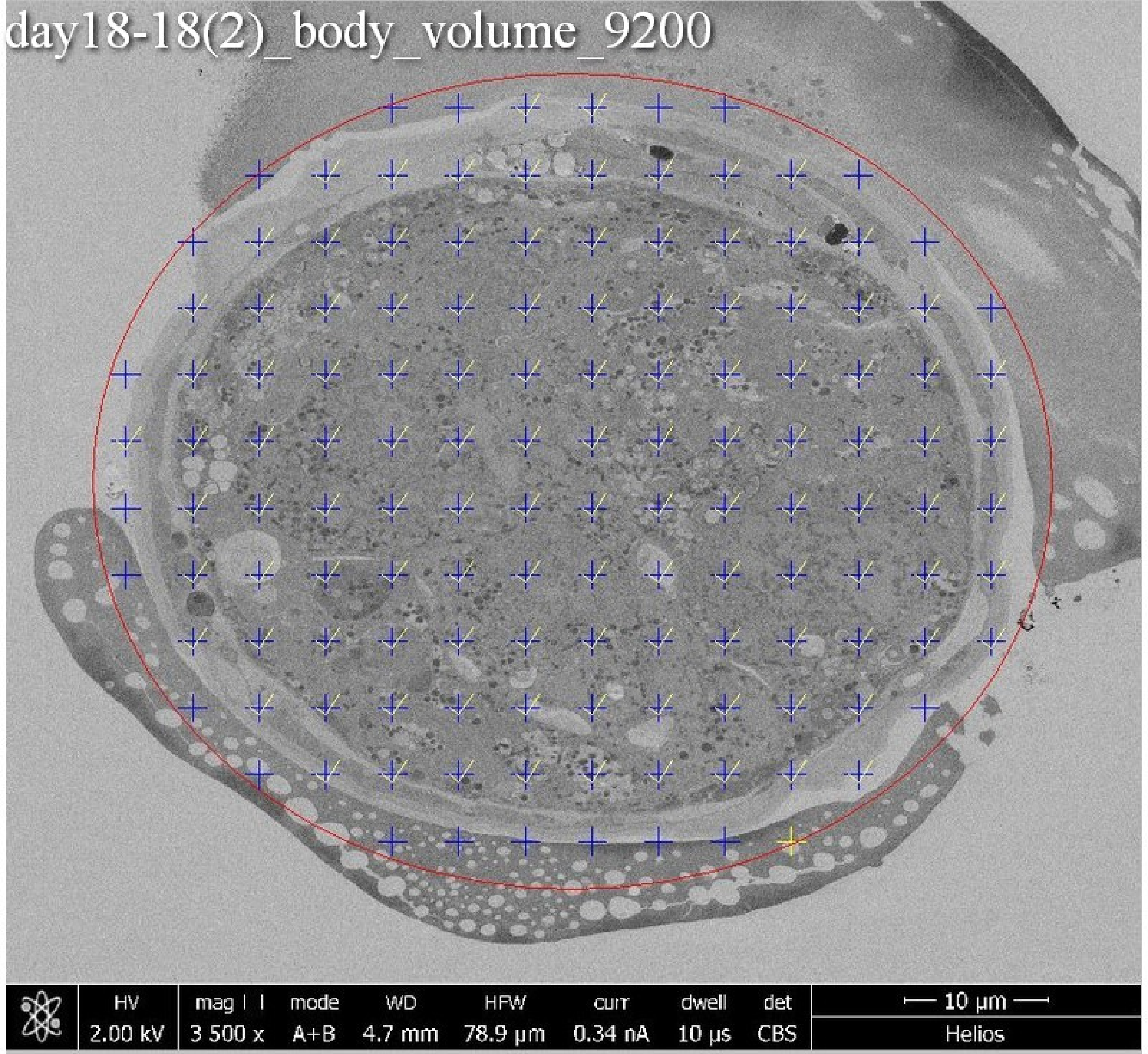
day18-18(2)_body_volume_9200
HV
2.00 kV
mag
3 500 x
mode
A+B
WD
4.7 mm
HFW
78.9 µm
curr
0.34 nA
dwell
10 µs
det
CBS
10 µm
Helios

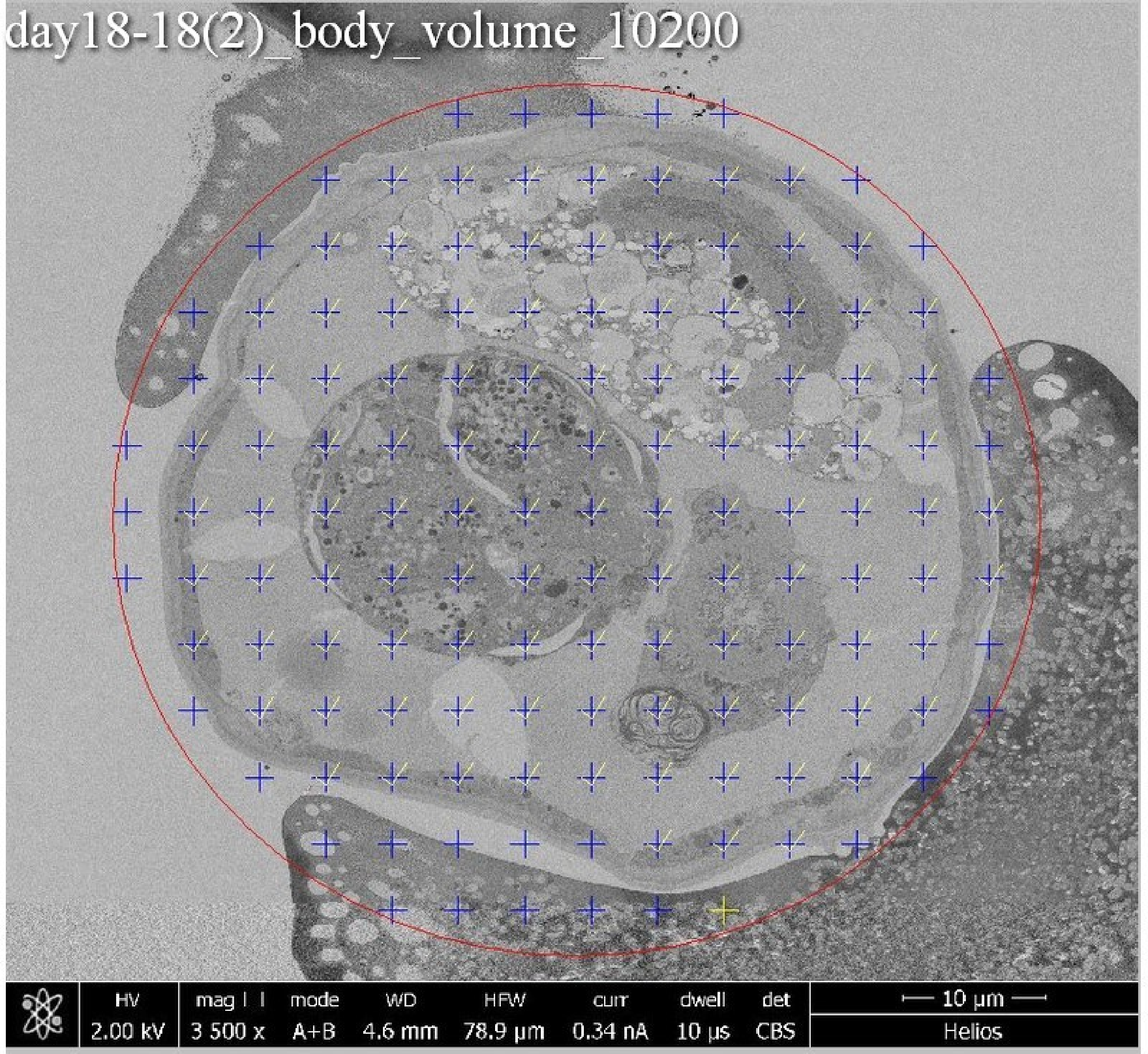
day18-18(2)_body_volume_10200
HV
2.00 kV
mag
3 500 x
mode
A+B
WD
4.6 mm
HFW
78.9 µm
curr
0.34 nA
dwell
10 µs
det
CBS
10 µm
Helios

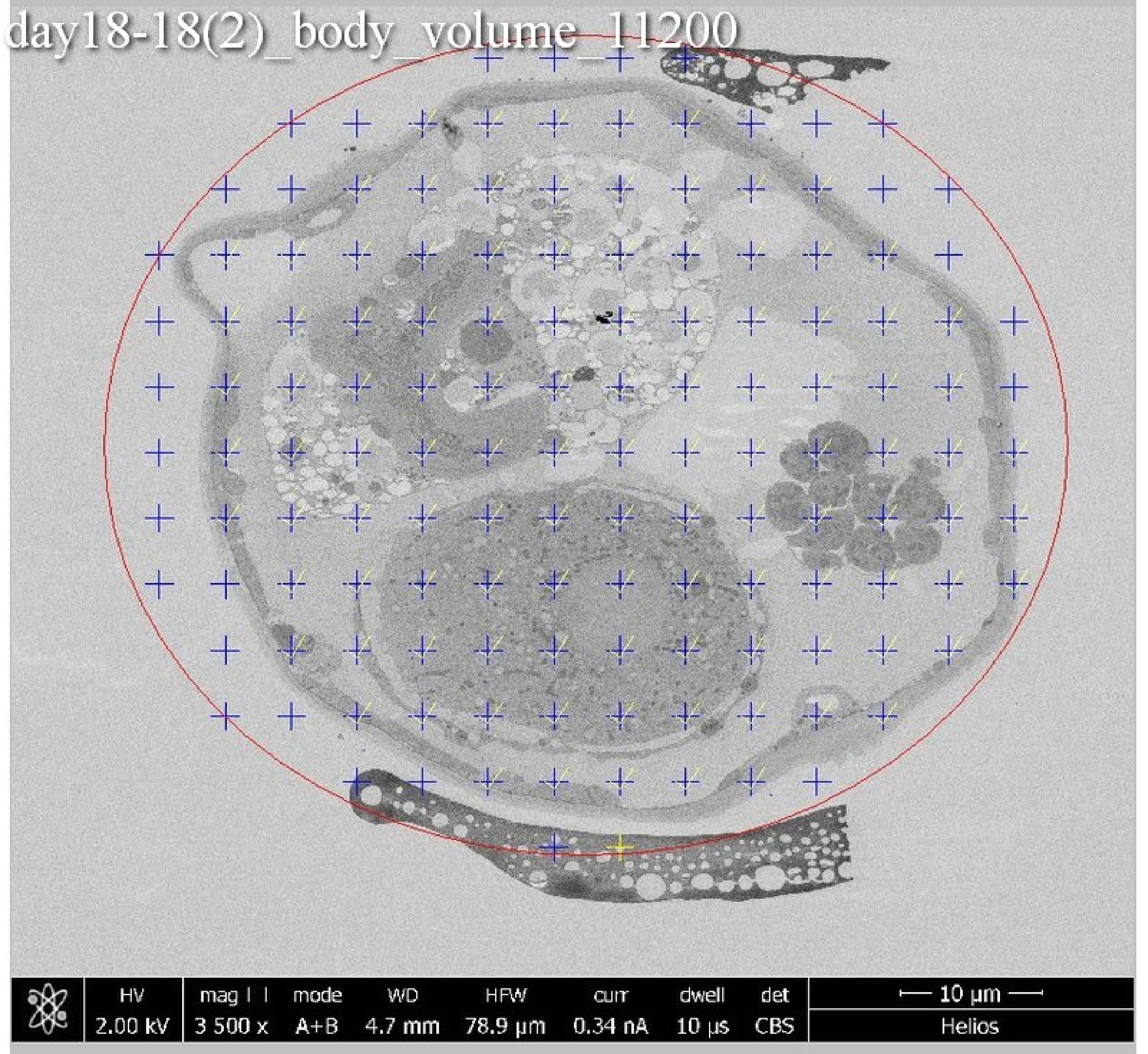
day18-18(2)_body_volume_11200
HV
2.00 kV
mag ⊡
3 500 x
mode
A+B
WD
4.7 mm
HFW
78.9 µm
curr
0.34 nA
dwell
10 µs
det
CBS
10 µm
Helios

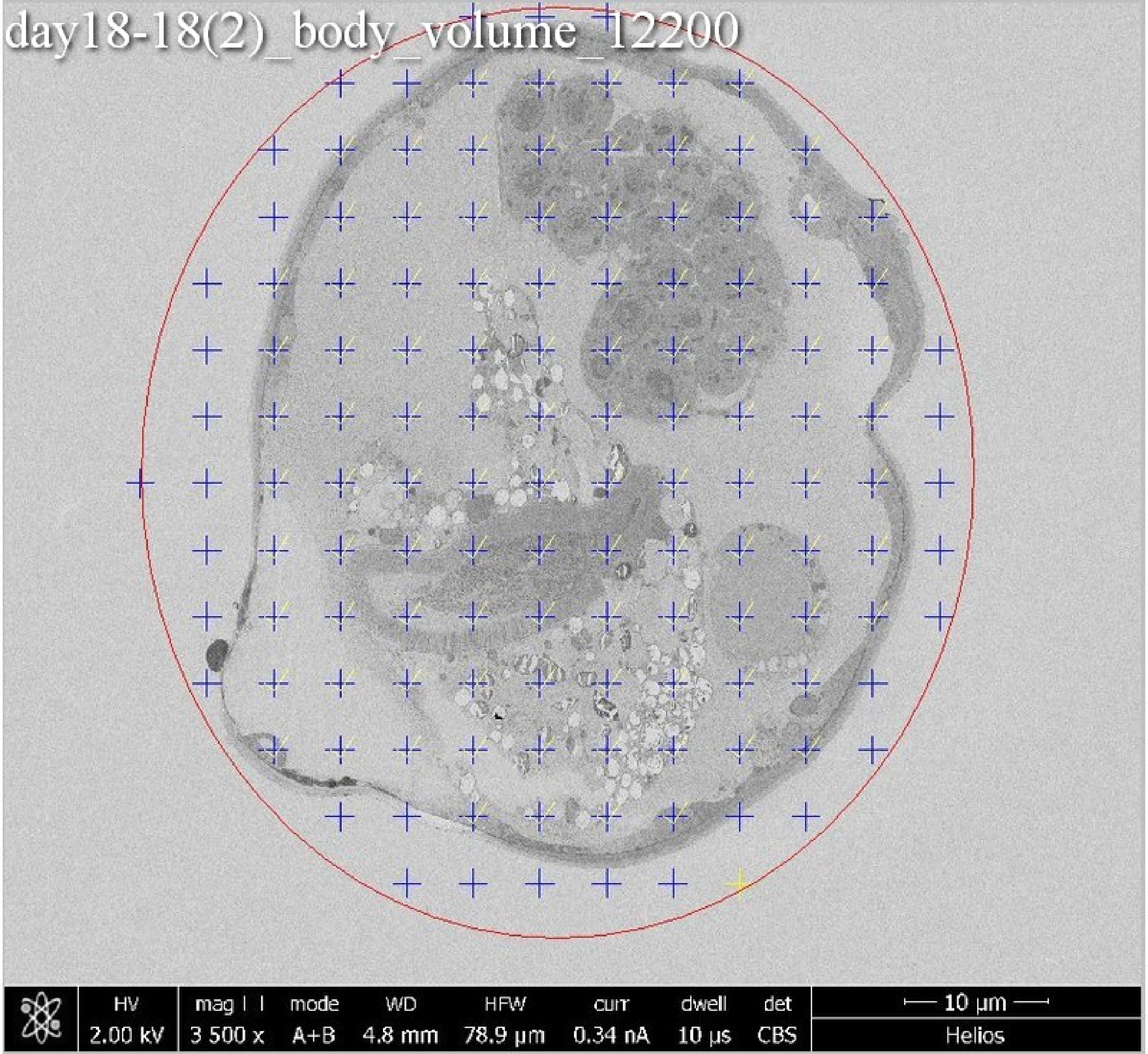
day18-18(2)_body_volume_12200
HV
2.00 kV
mag
3 500 x
mode
A+B
WD
4.8 mm
HFW
78.9 µm
curr
0.34 nA
dwell
10 µs
det
CBS
10 µm
Helios

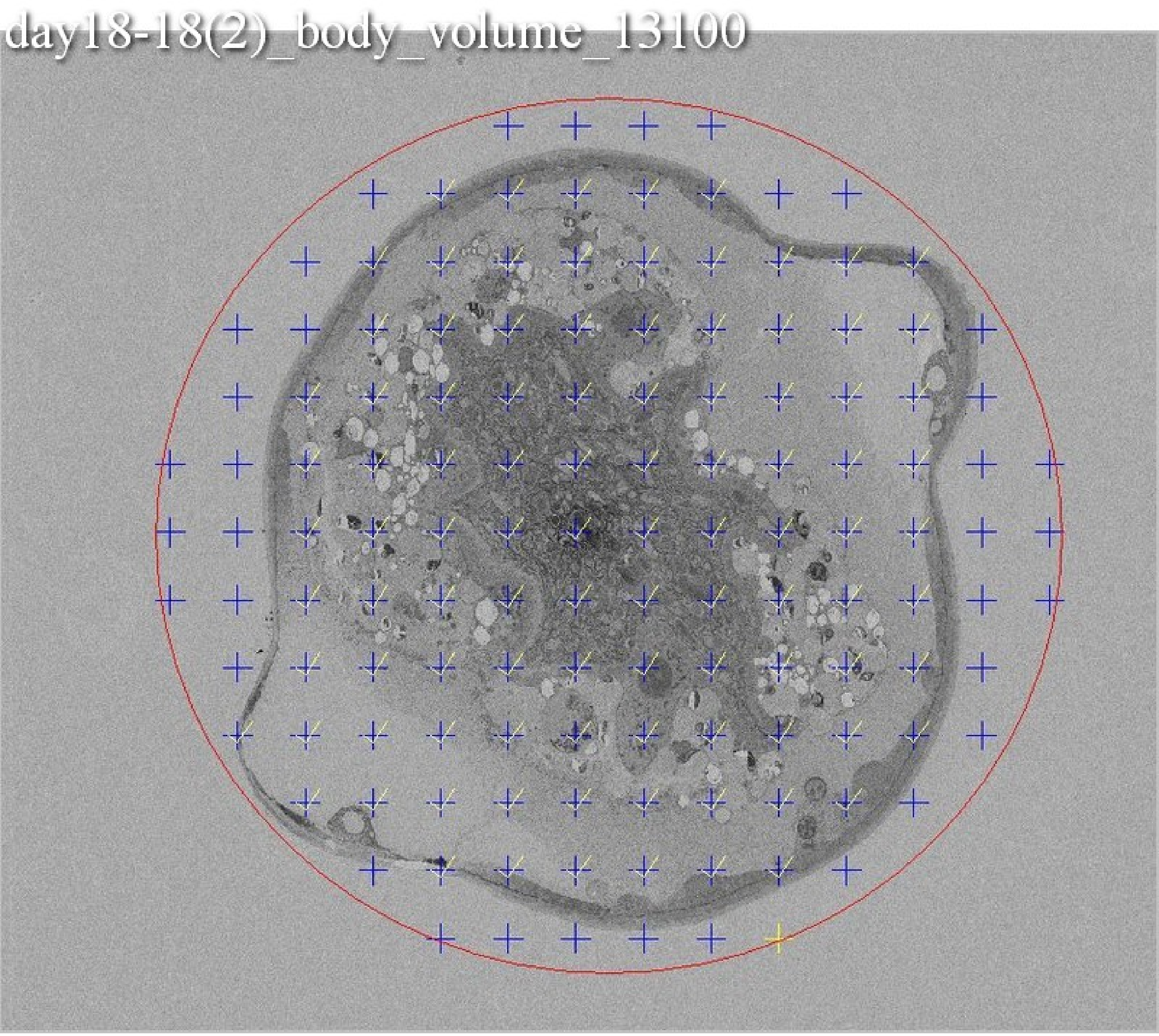
day18-18(2)_body_volume_13100

day18-18(2)_body_volume_13900

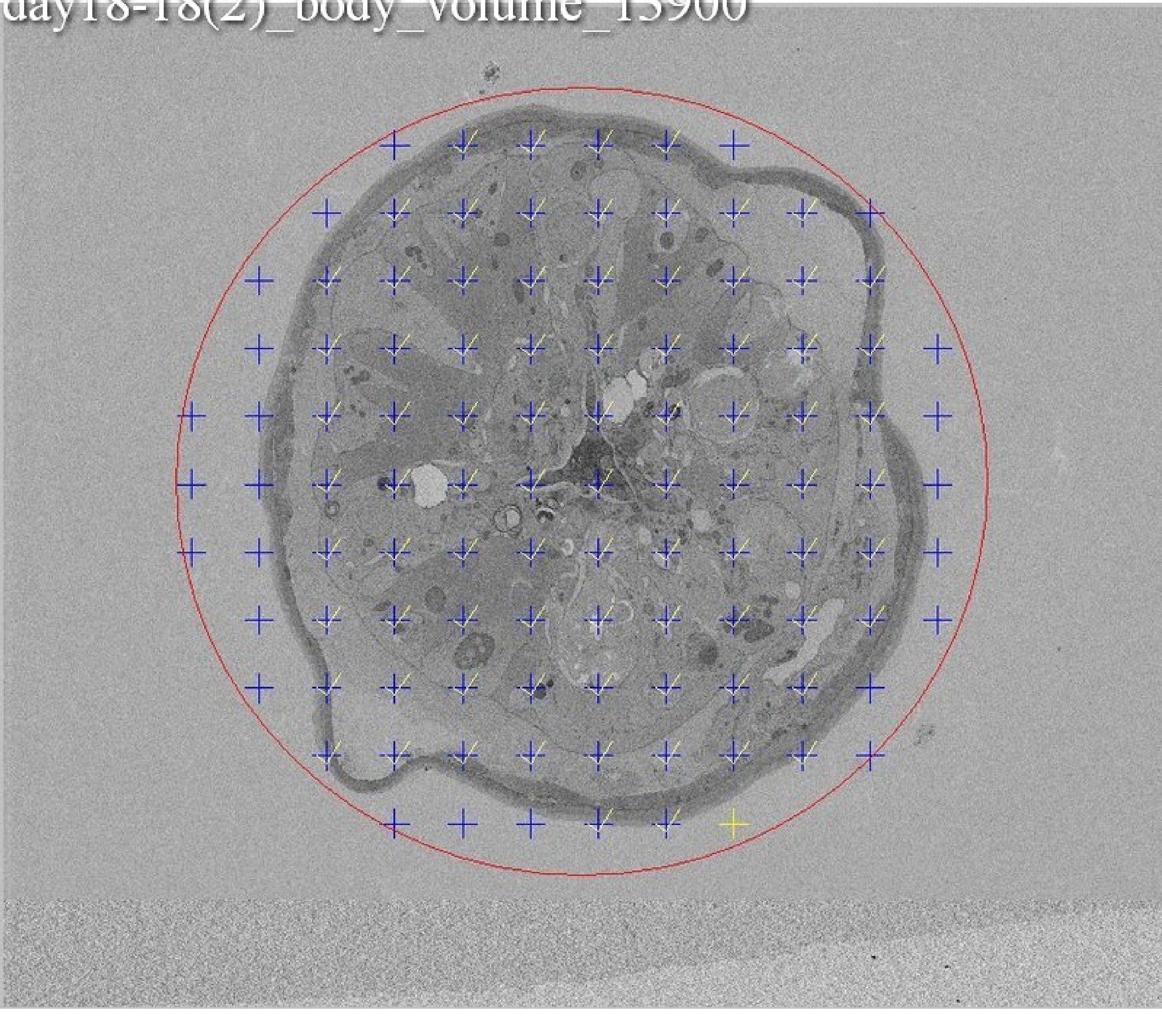

day18-18(2)_body_volume_14700

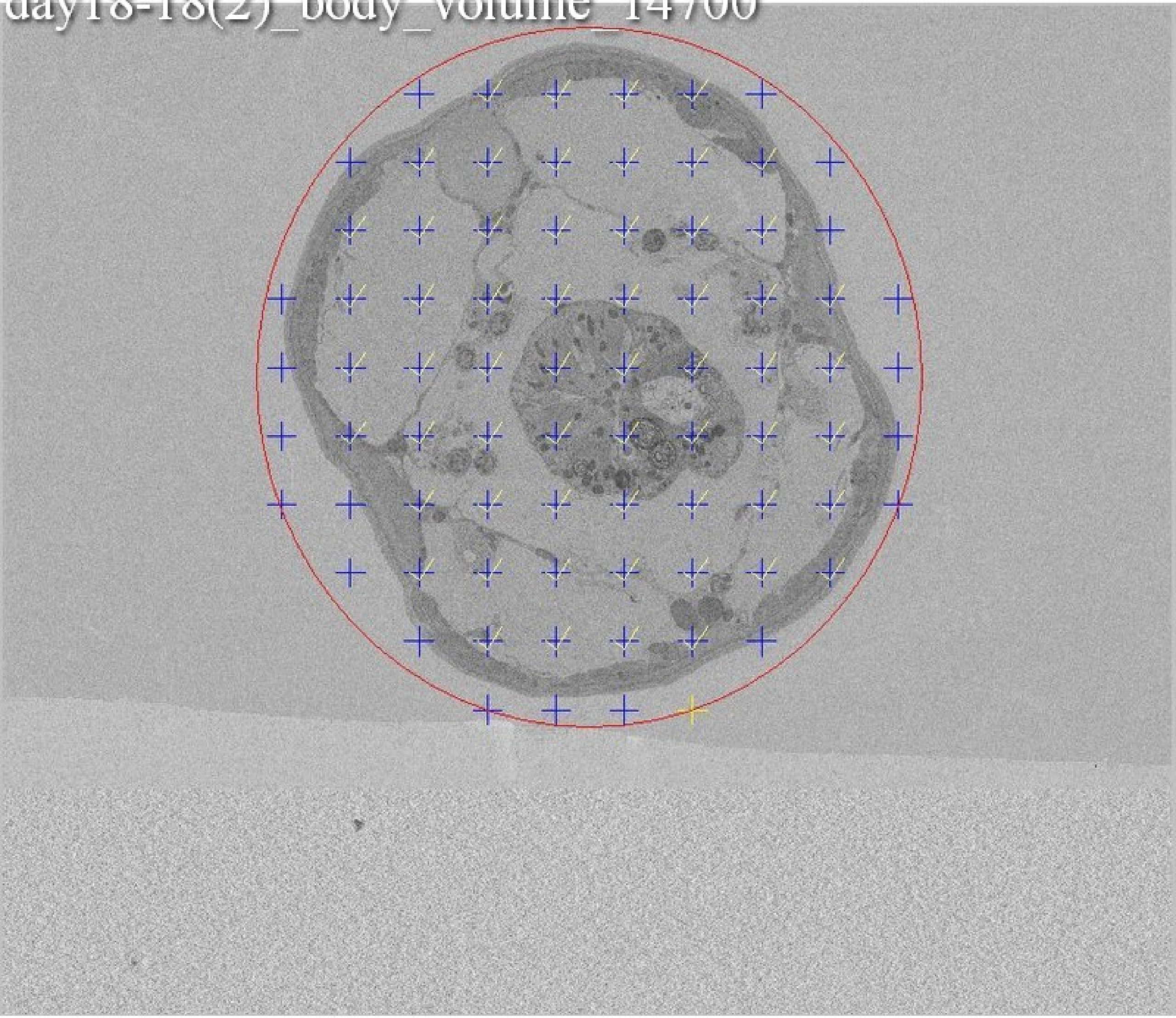

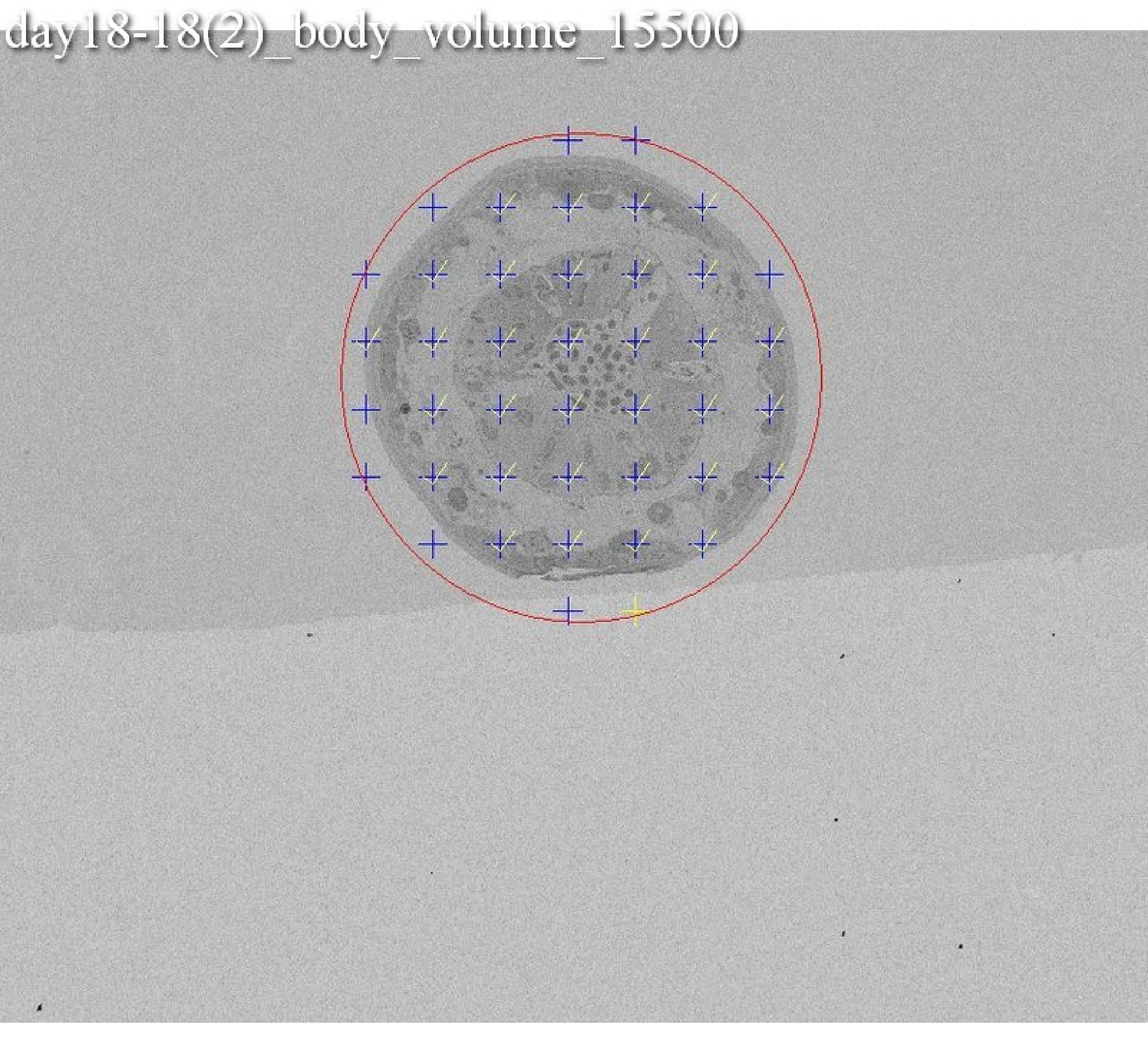
day18-18(2)_body_volume_15500

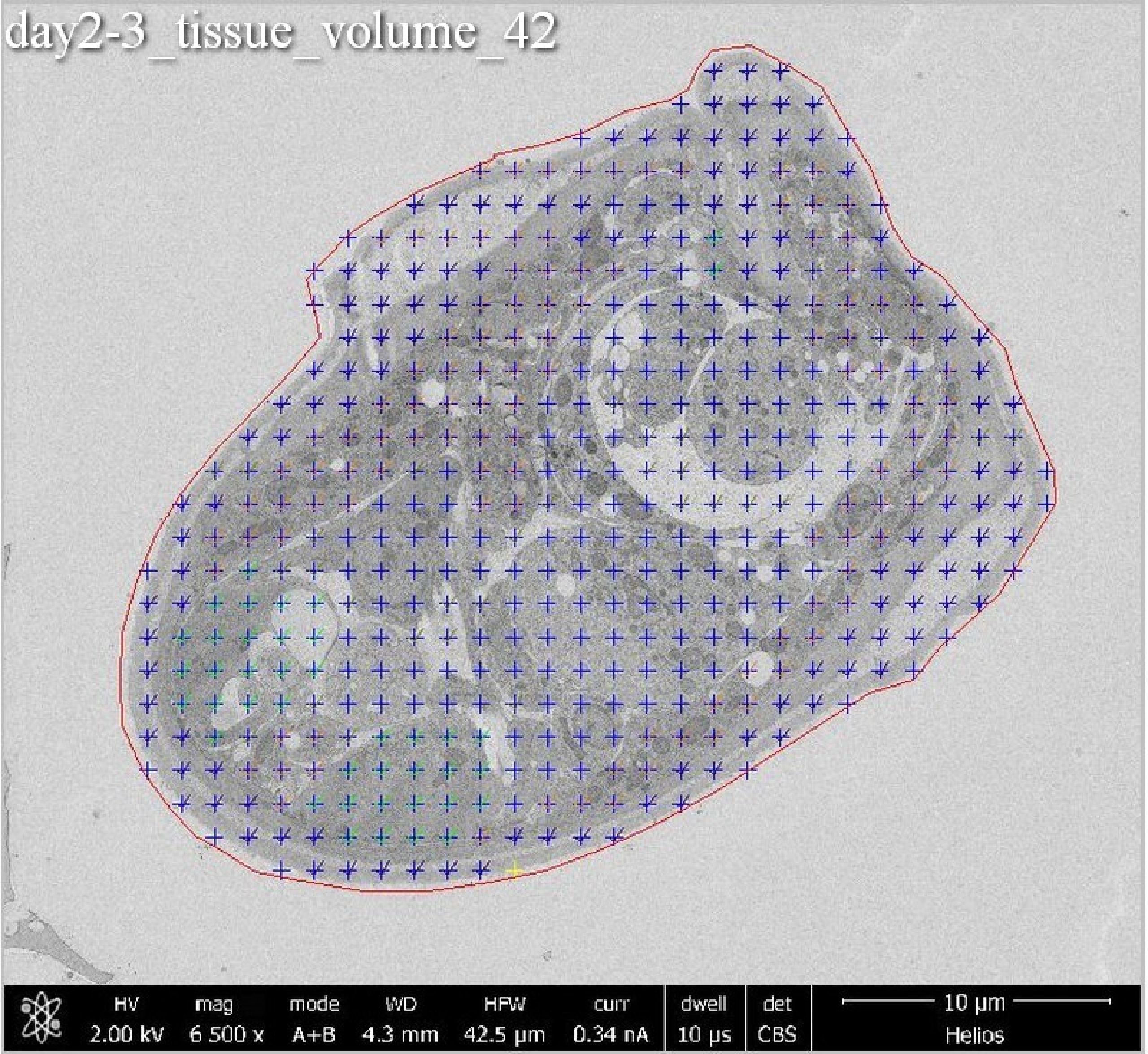
day2-3_tissue_volume_42
HV
2.00 kV
mag
6 500 x
mode
A+B
WD
4.3 mm
HFW
42.5 µm
curr
0.34 nA
dwell
10 µs
det
CBS
10 µm
Helios

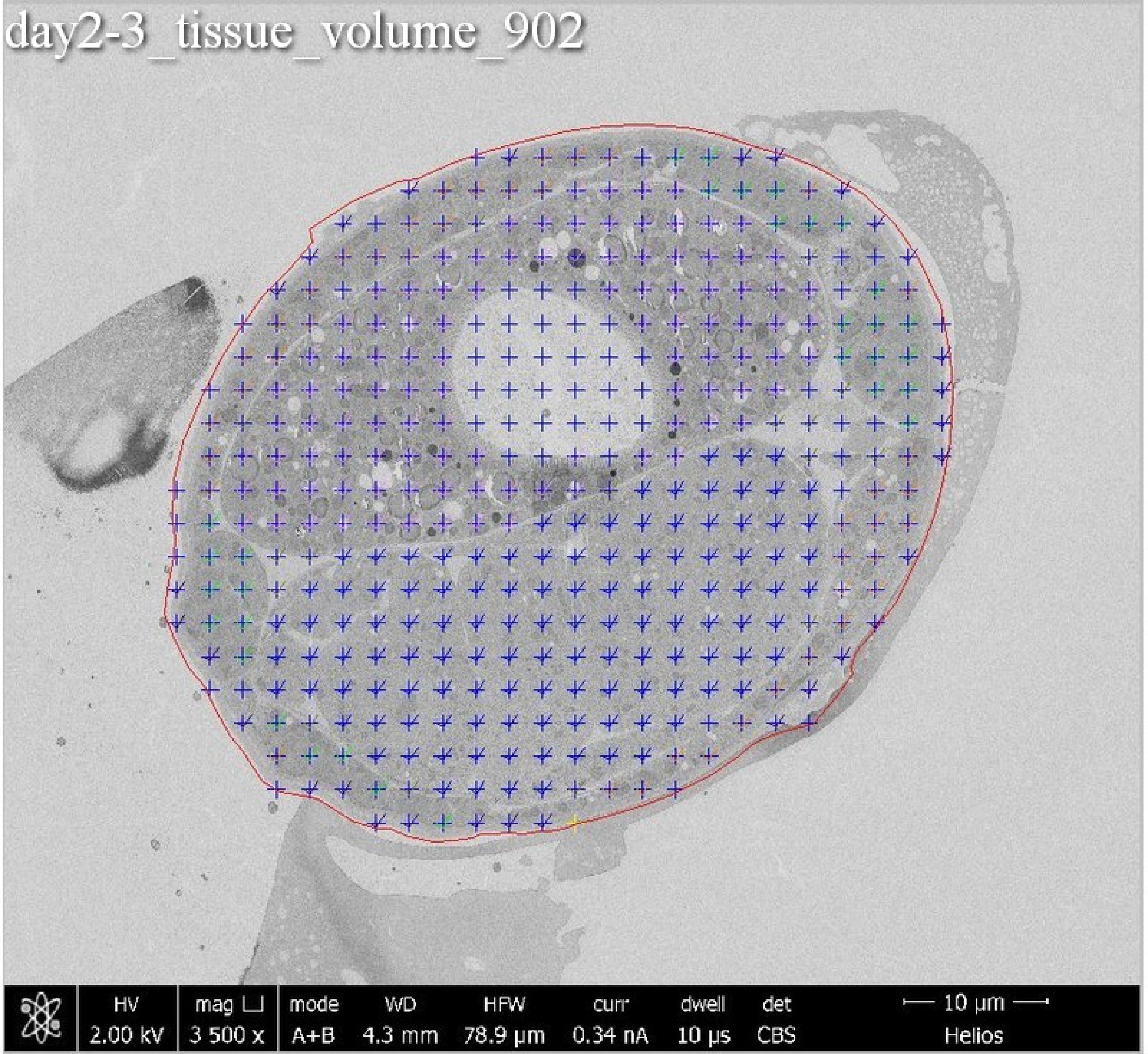
day2-3_tissue_volume_902
HV 2.00 kV
mag ⊔ 3 500 x
mode A+B
WD 4.3 mm
HFW 78.9 μm
curr 0.34 nA
dwell 10 μs
det CBS
10 μm
Helios

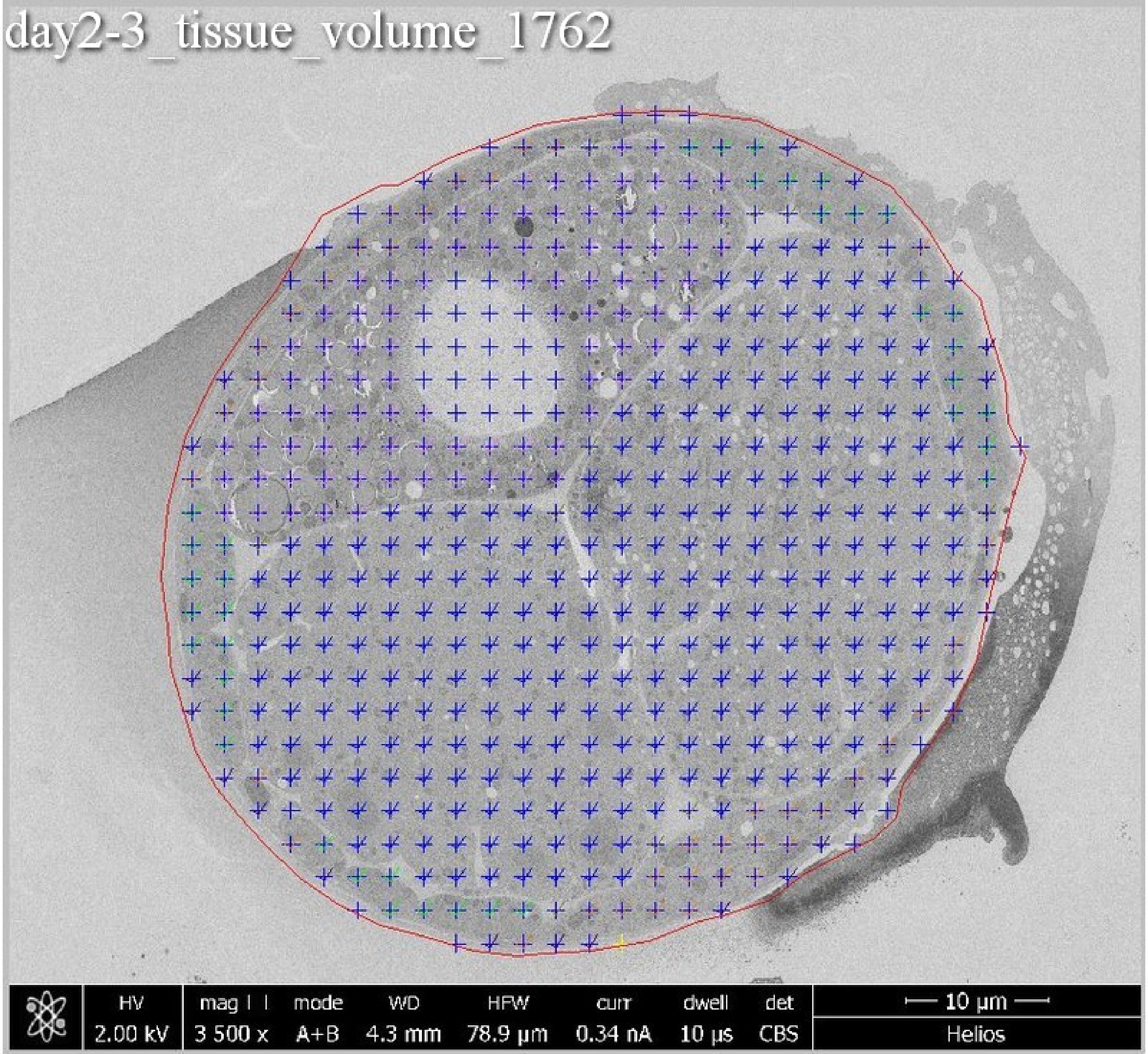
day2-3_tissue_volume_1762
HV
2.00 kV
mag
3 500 x
mode
A+B
WD
4.3 mm
HFW
78.9 µm
curr
0.34 nA
dwell
10 µs
det
CBS
10 µm
Helios

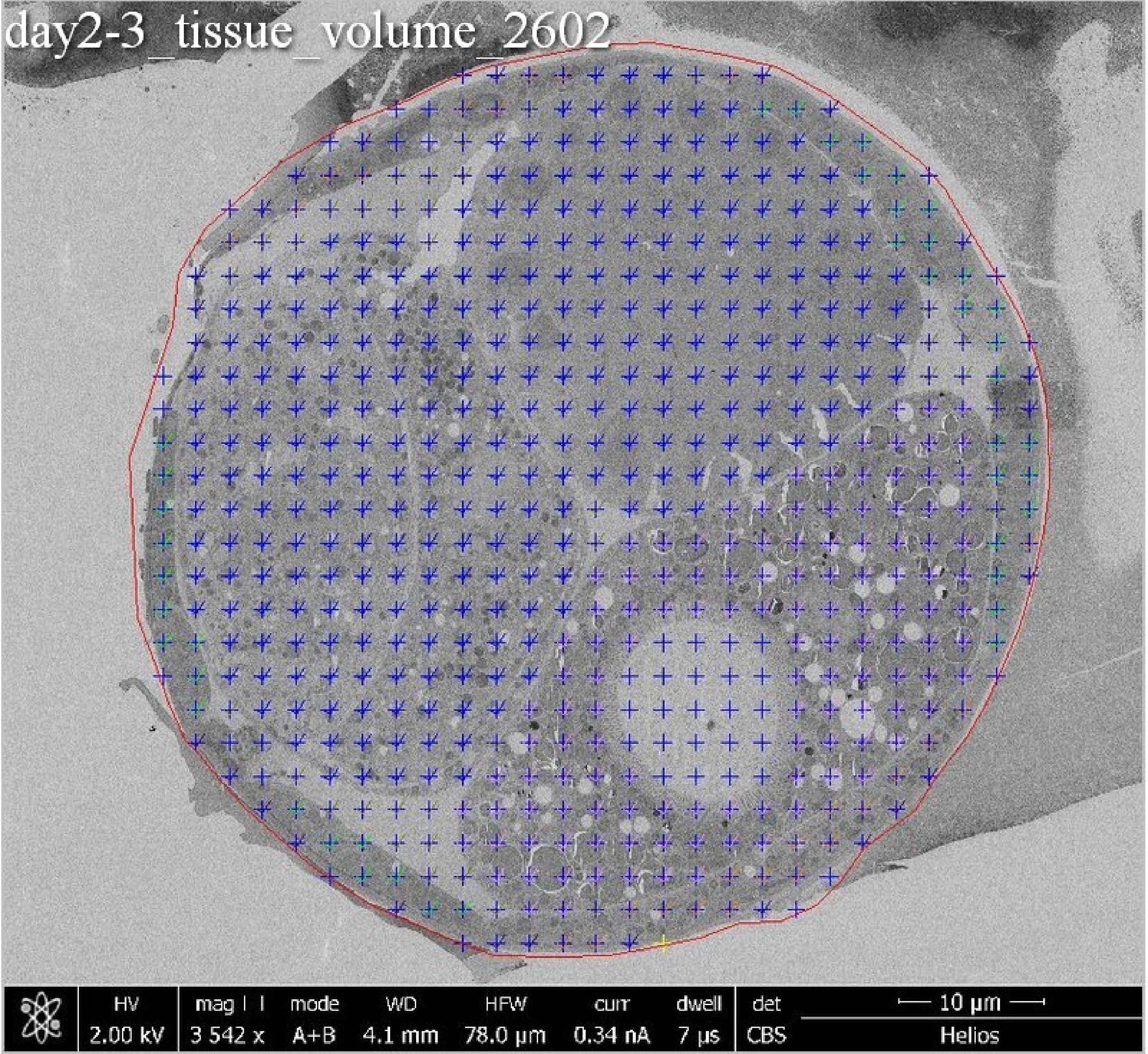
day2-3_tissue_volume_2602
HV
2.00 kV
mag
3 542 x
mode
A+B
WD
4.1 mm
HFW
78.0 μm
curr
0.34 nA
dwell
7 μs
det
CBS
10 μm
Helios

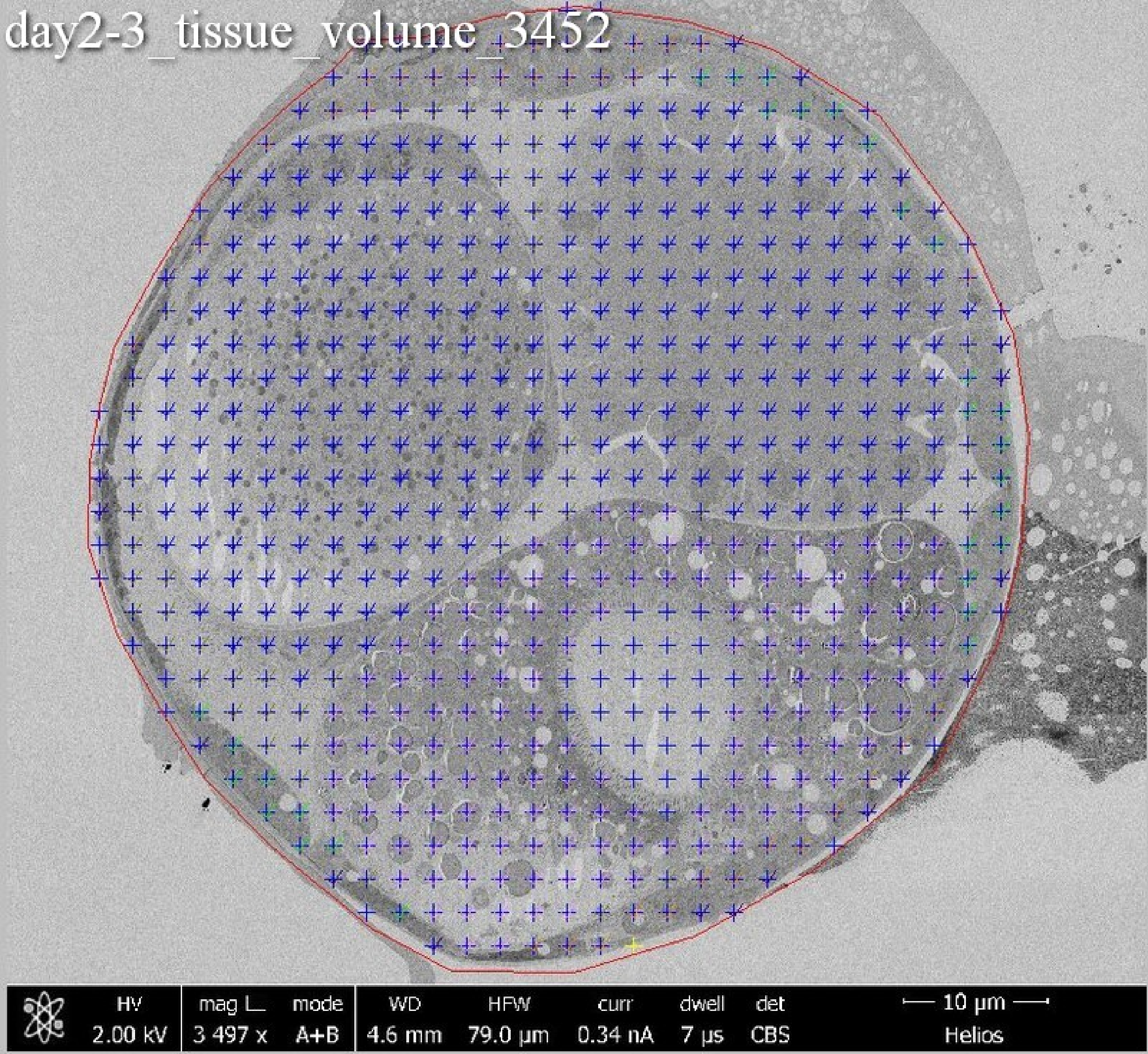
day2-3_tissue_volume_3452
HV
2.00 kV
mag L
3 497 x
mode
A+B
WD
4.6 mm
HFW
79.0 µm
curr
0.34 nA
dwell
7 µs
det
CBS
10 µm
Helios

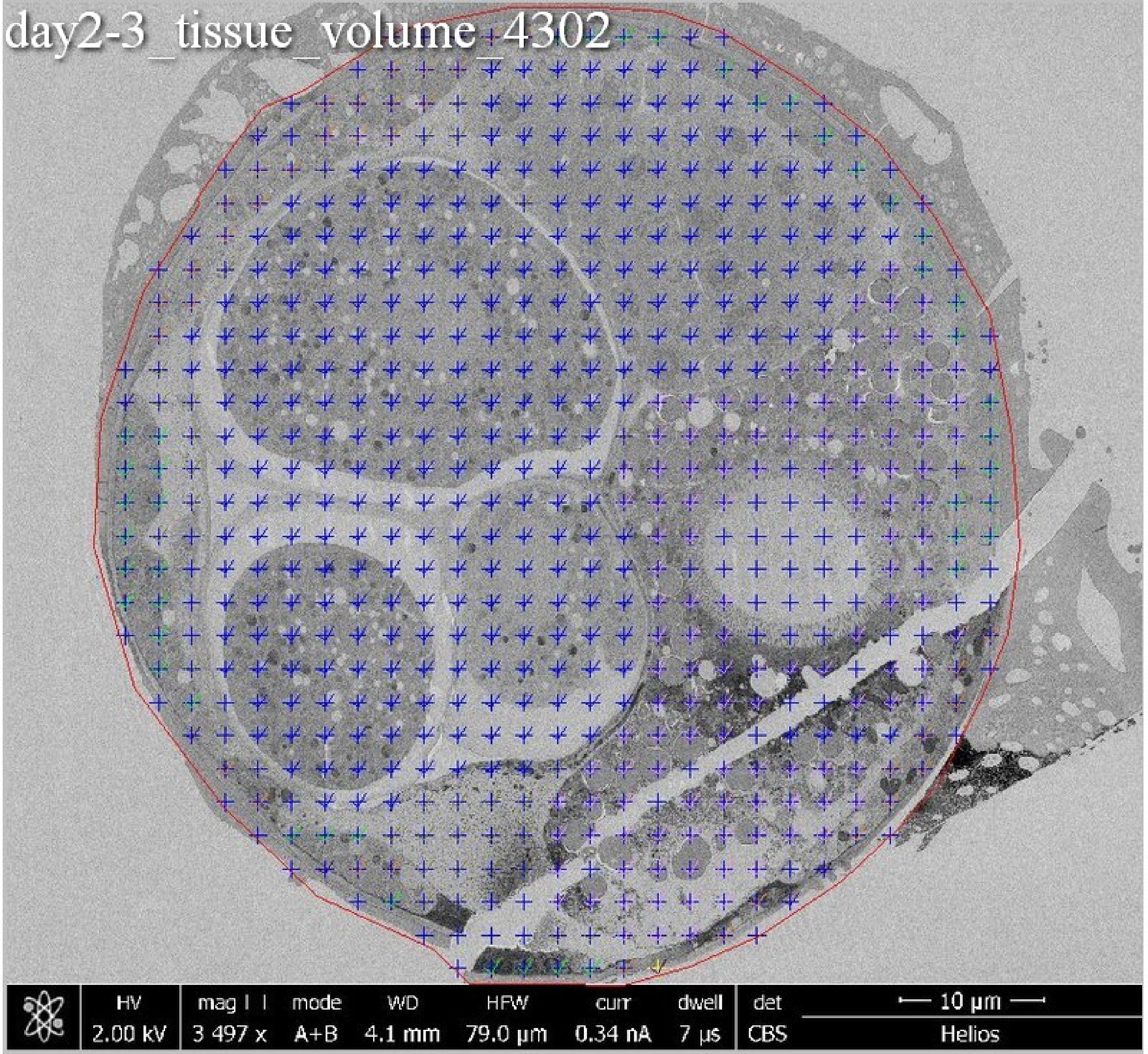
day2-3_tissue_volume_4302
HV
2.00 kV
mag
3 497 x
mode
A+B
WD
4.1 mm
HFW
79.0 μm
curr
0.34 nA
dwell
7 μs
det
CBS
10 μm
Helios

day2-3_tissue_volume_5152

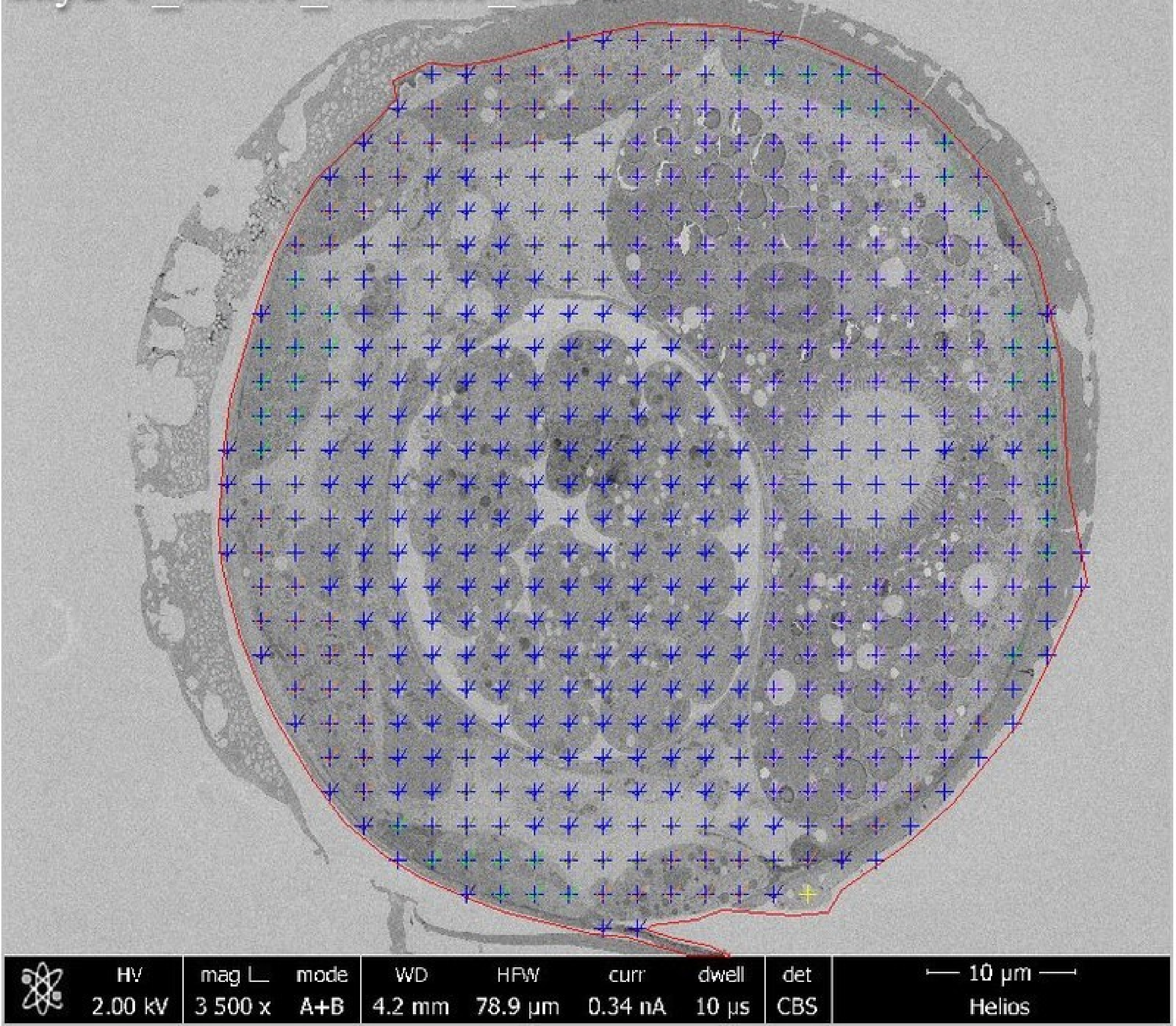

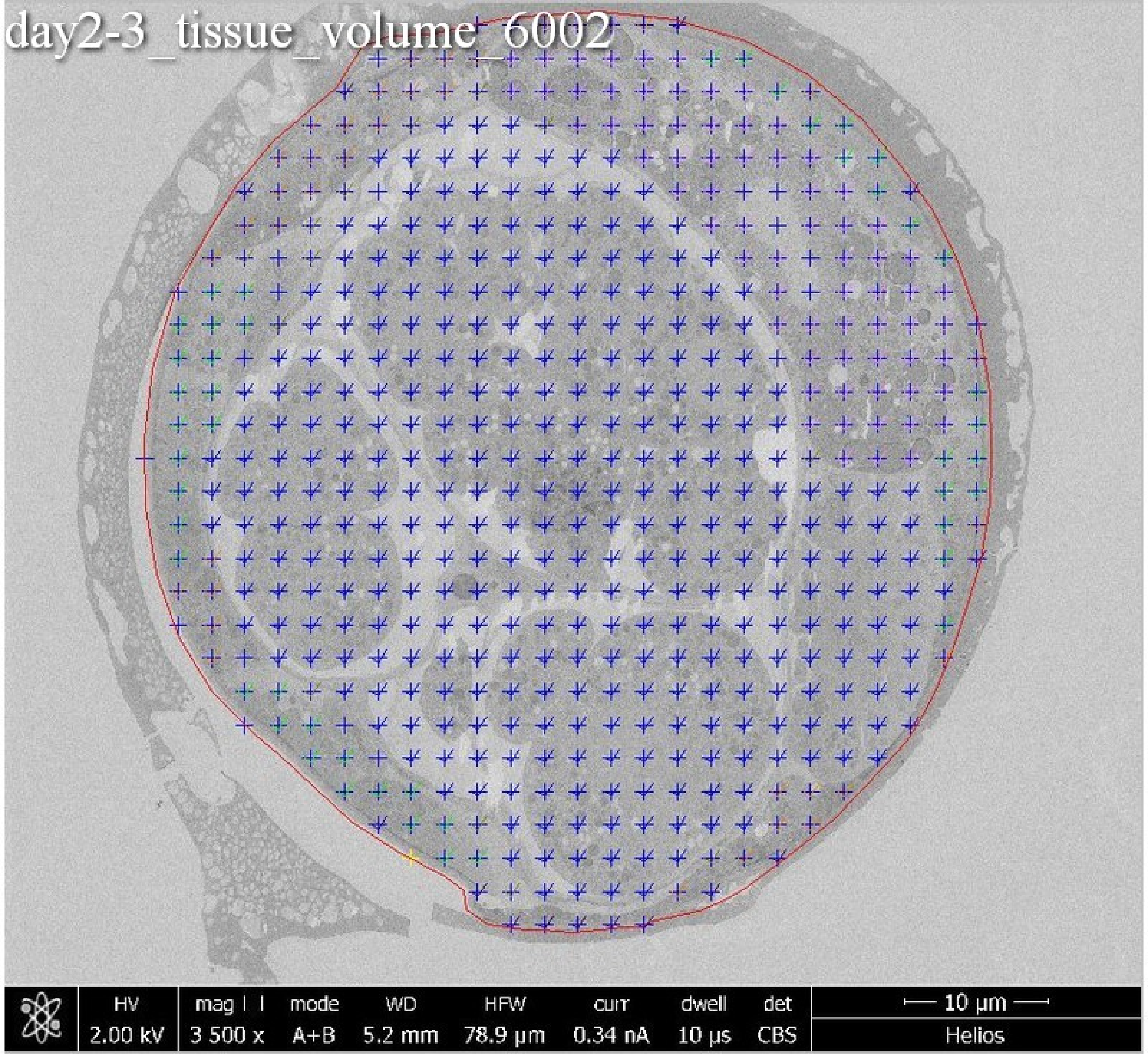
day2-3_tissue_volume_6002
HV
2.00 kV
mag
3 500 x
mode
A+B
WD
5.2 mm
HFW
78.9 µm
curr
0.34 nA
dwell
10 µs
det
CBS
10 µm
Helios

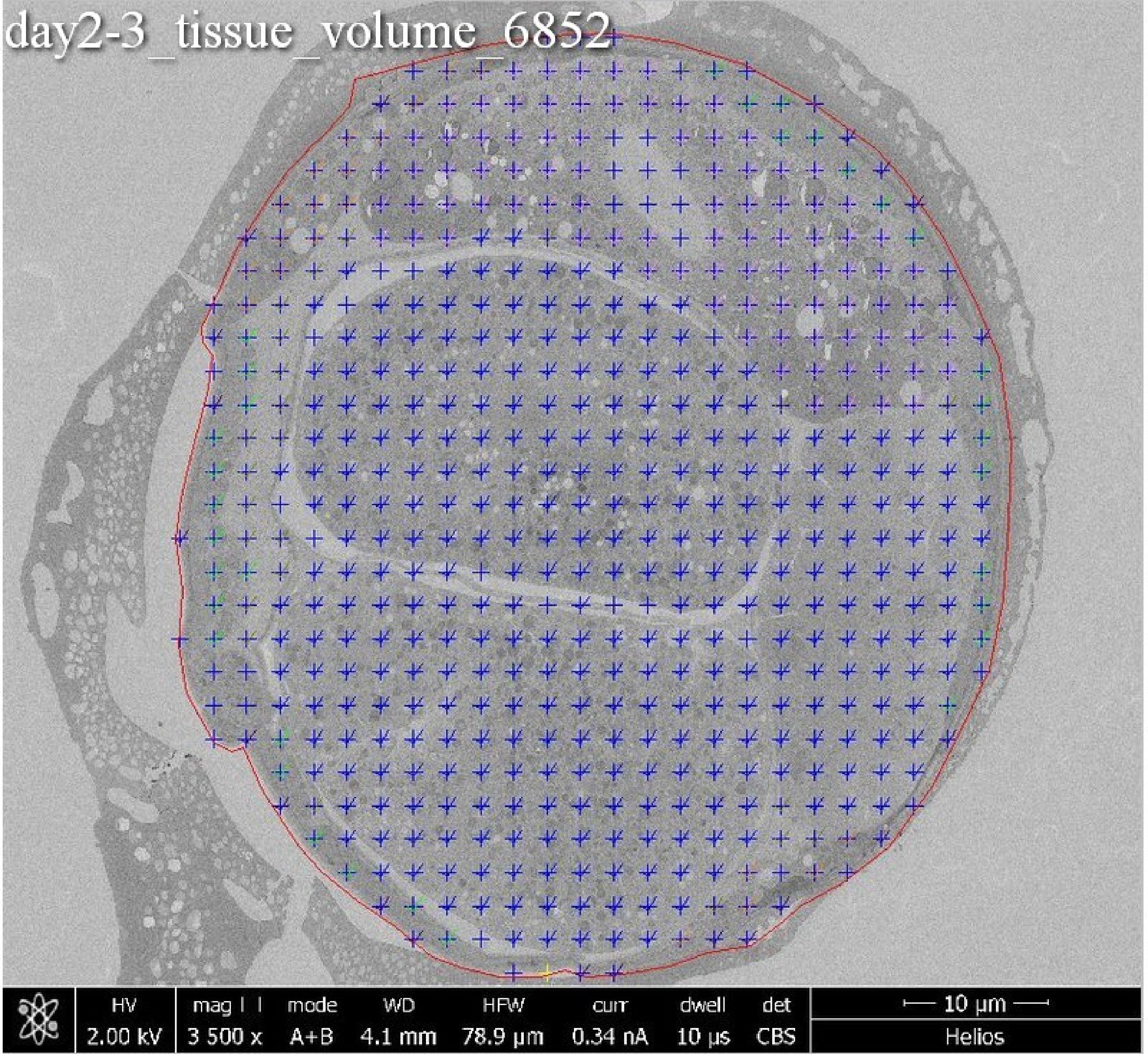
day2-3_tissue_volume_6852
HV
2.00 kV
mag
3 500 x
mode
A+B
WD
4.1 mm
HFW
78.9 µm
curr
0.34 nA
dwell
10 µs
det
CBS
10 µm
Helios

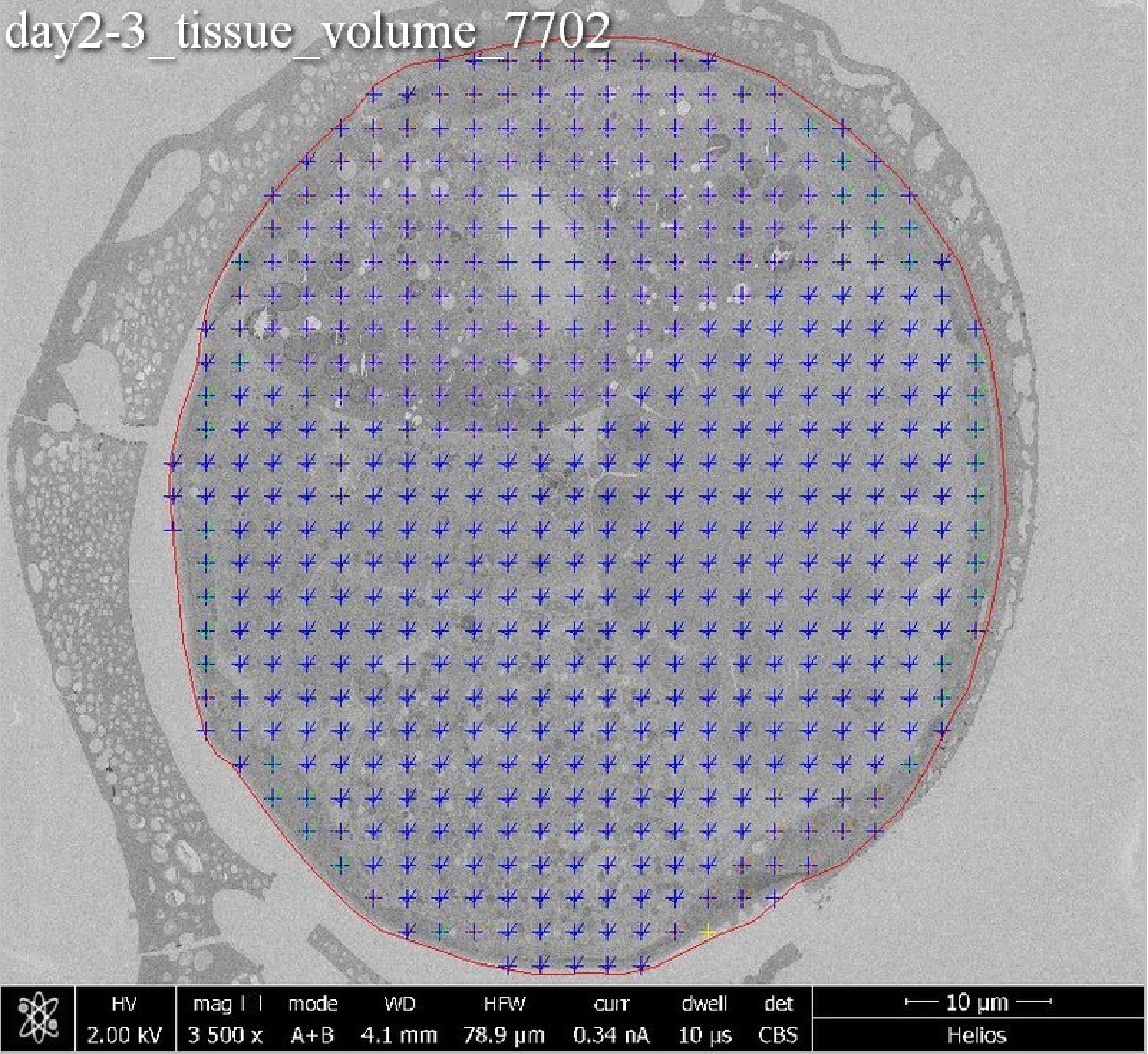
day2-3_tissue_volume_7702
HV
2.00 kV
mag
3 500 x
mode
A+B
WD
4.1 mm
HFW
78.9 µm
curr
0.34 nA
dwell
10 µs
det
CBS
10 µm
Helios

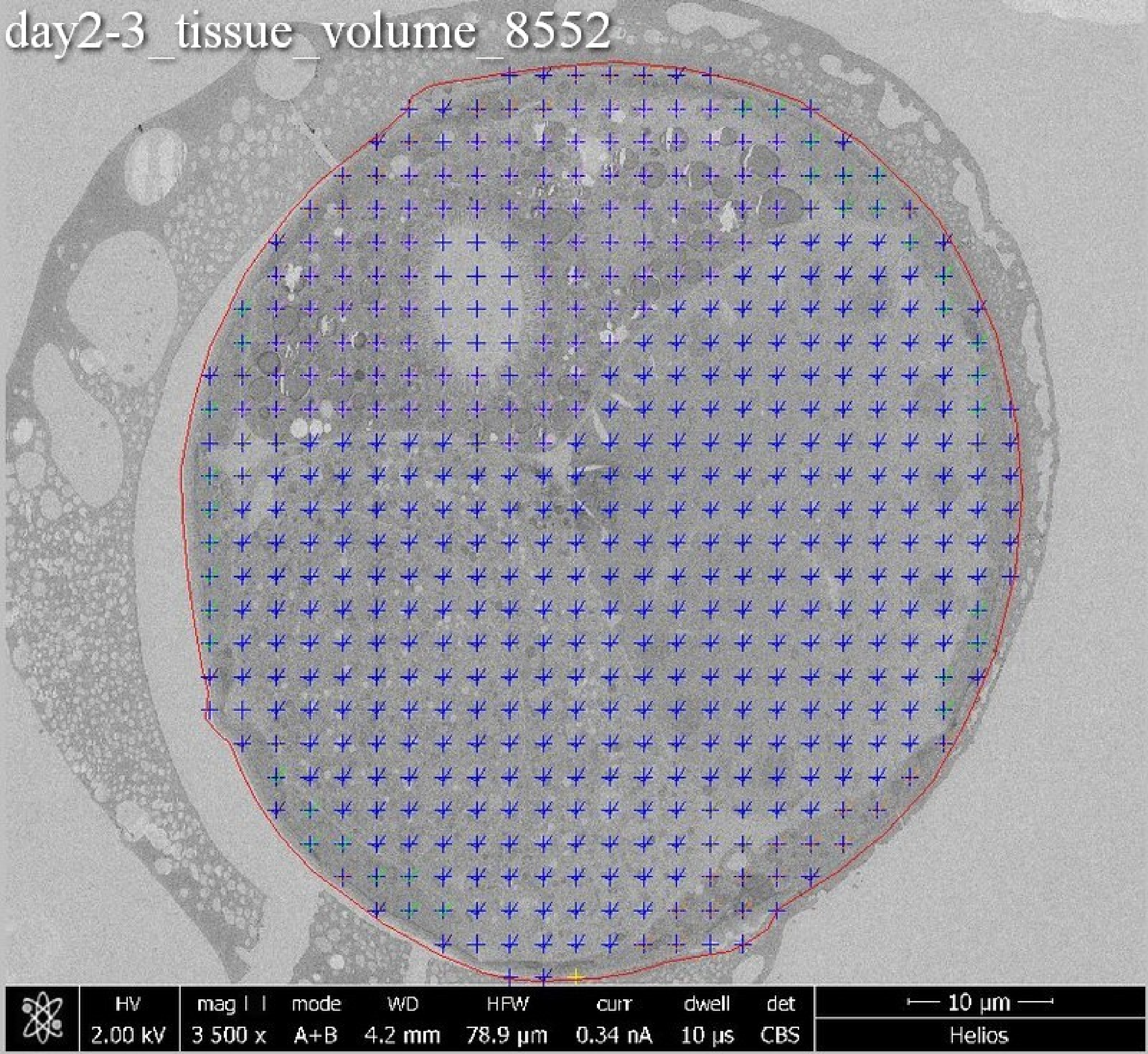
day2-3_tissue_volume_8552
HV
2.00 kV
mag
3 500 x
mode
A+B
WD
4.2 mm
HFW
78.9 µm
curr
0.34 nA
dwell
10 µs
det
CBS
10 µm
Helios

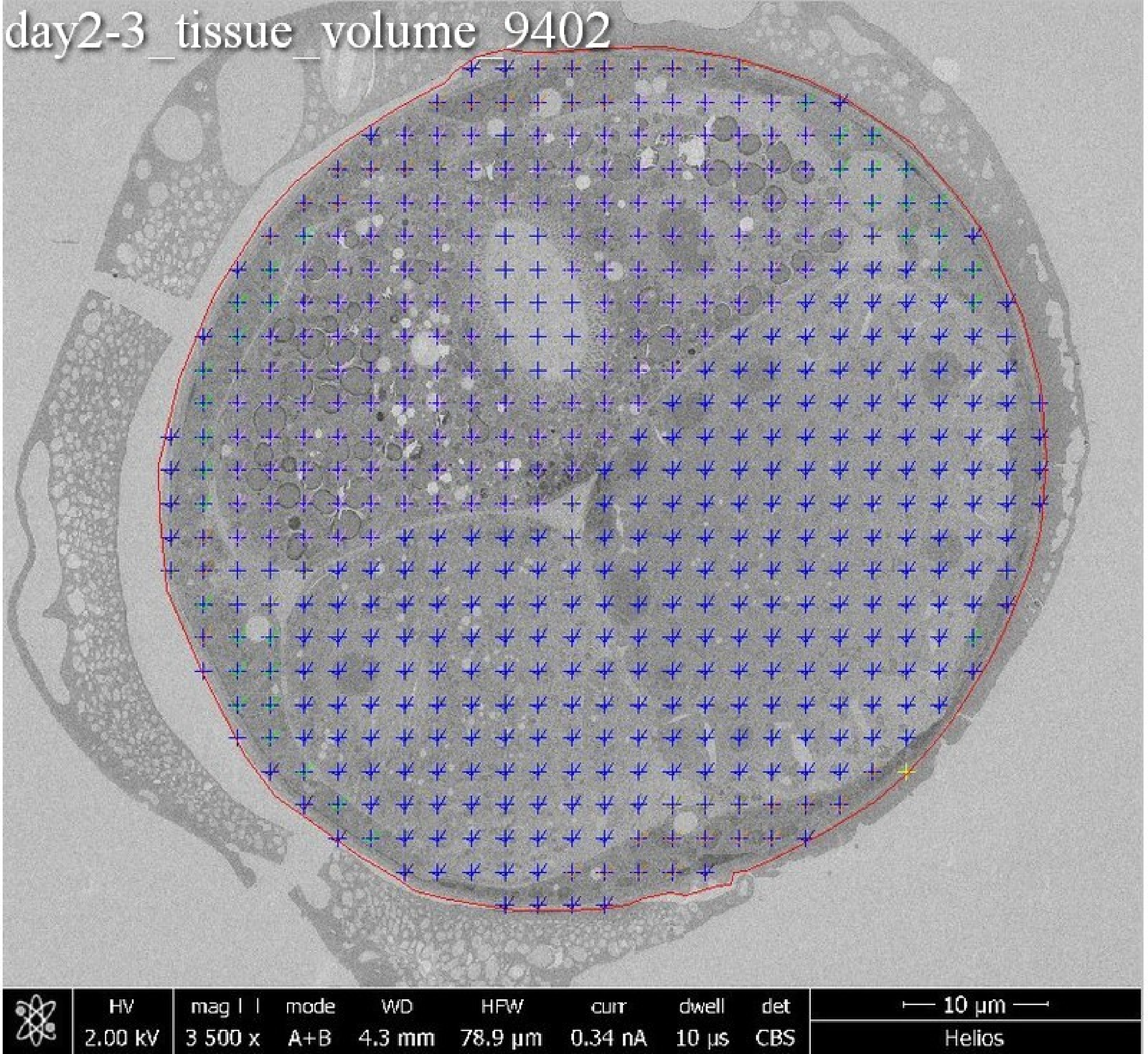
day2-3_tissue_volume_9402
HV
2.00 kV
mag
3 500 x
mode
A+B
WD
4.3 mm
HFW
78.9 µm
curr
0.34 nA
dwell
10 µs
det
CBS
10 µm
Helios

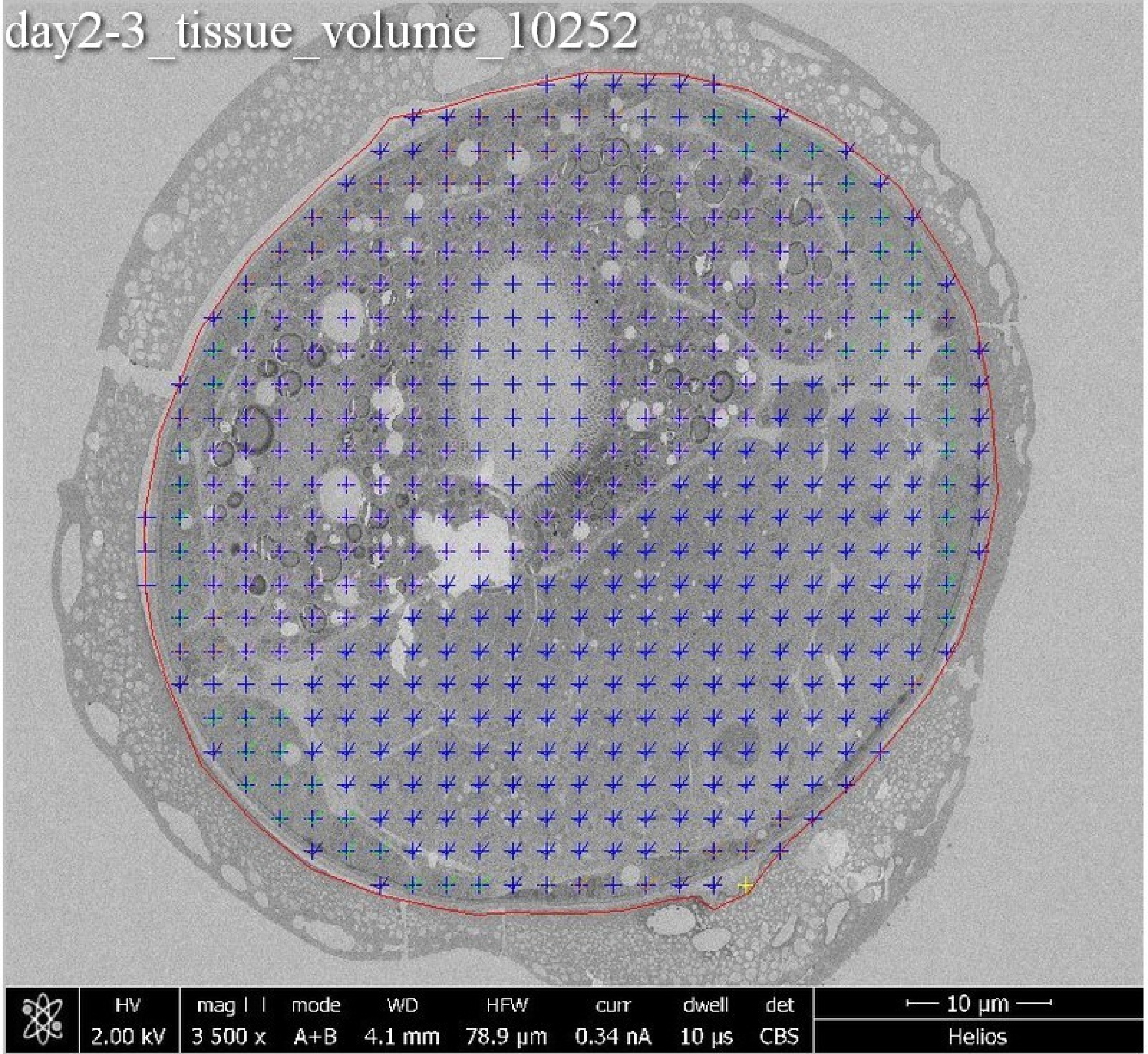
day2-3_tissue_volume_10252
HV
2.00 kV
mag
3 500 x
mode
A+B
WD
4.1 mm
HFW
78.9 μm
curr
0.34 nA
dwell
10 μs
det
CBS
10 μm
Helios

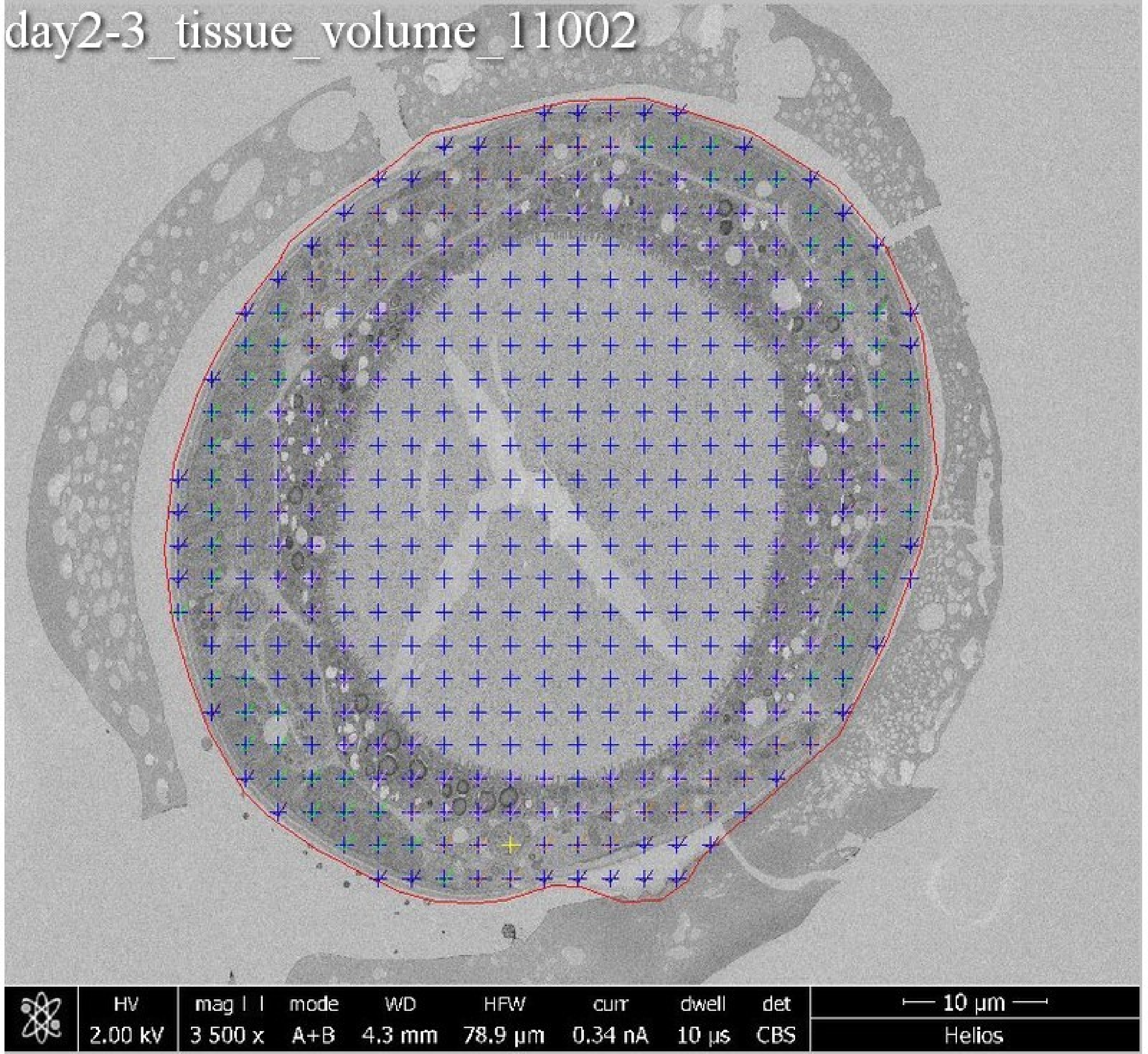

day2-3_tissue_volume_11002
HV
2.00 kV
mag ⊡
3 500 x
mode
A+B
WD
4.3 mm
HFW
78.9 µm
curr
0.34 nA
dwell
10 µs
det
CBS
10 µm
Helios

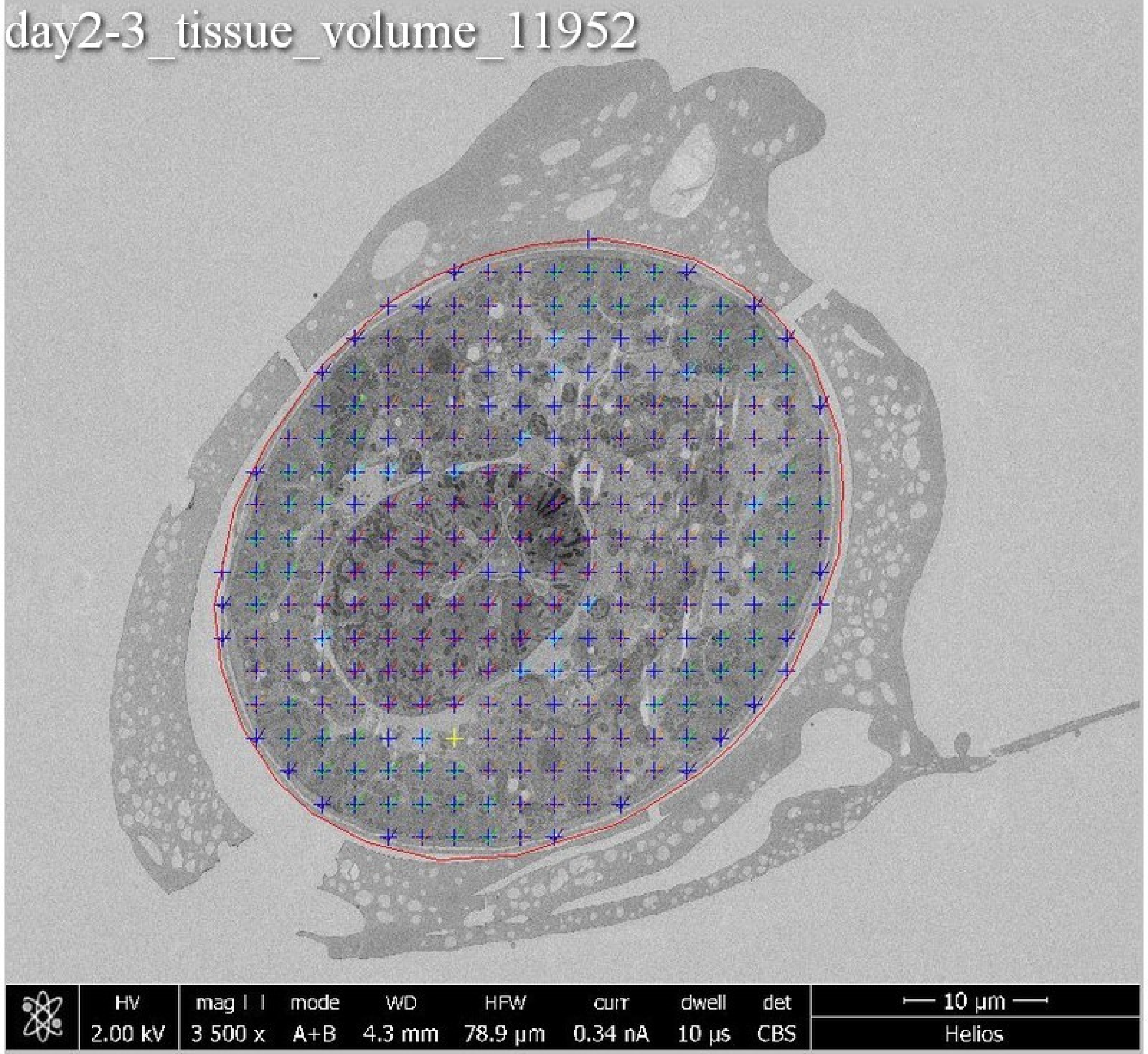
day2-3_tissue_volume_11952
HV
2.00 kV
mag
3 500 x
mode
A+B
WD
4.3 mm
HFW
78.9 µm
curr
0.34 nA
dwell
10 µs
det
CBS
10 µm
Helios

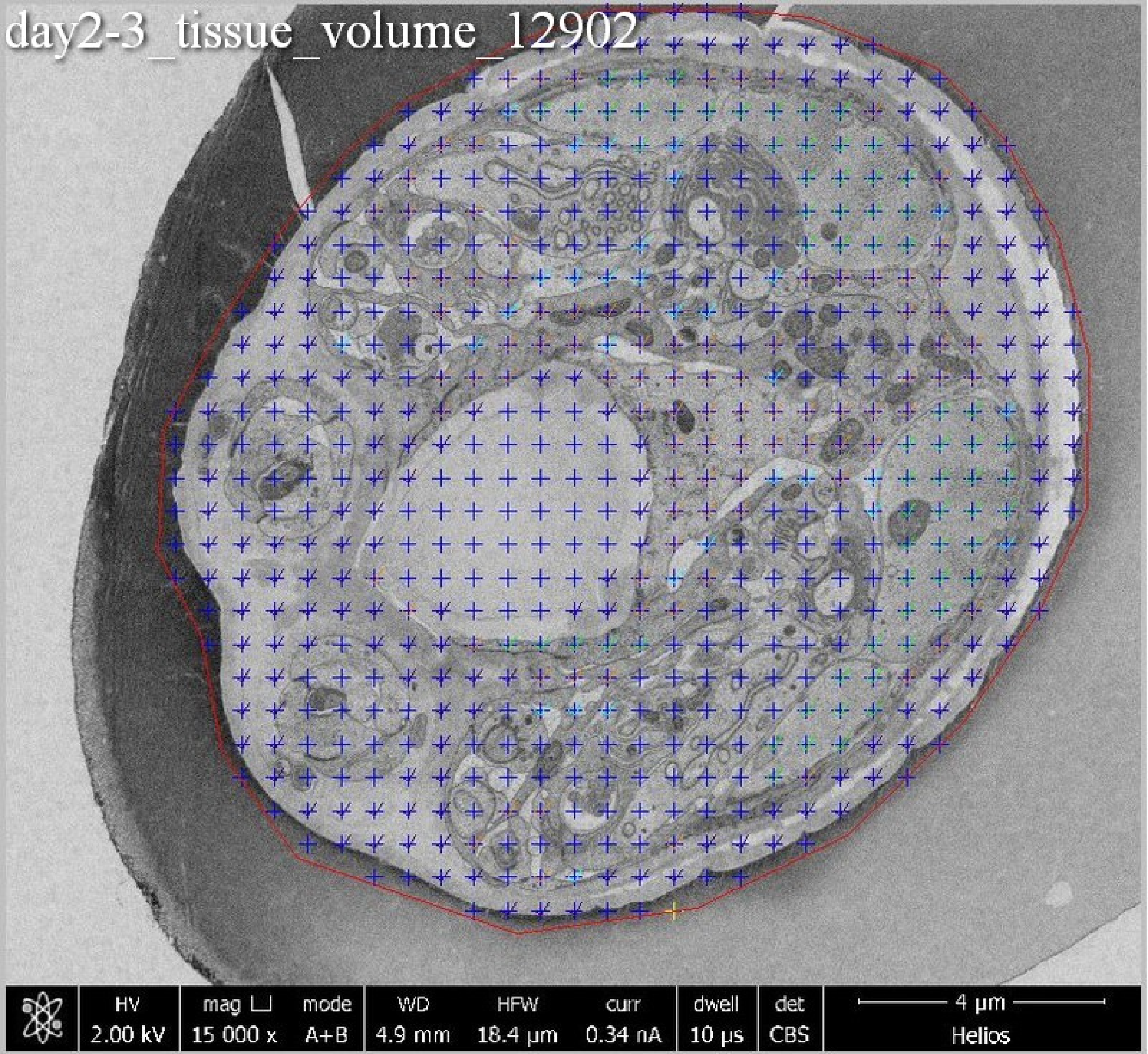
day2-3_tissue_volume_12902
HV
2.00 kV
mag ⊔
15 000 x
mode
A+B
WD
4.9 mm
HFW
18.4 µm
curr
0.34 nA
dwell
10 µs
det
CBS
4 µm
Helios

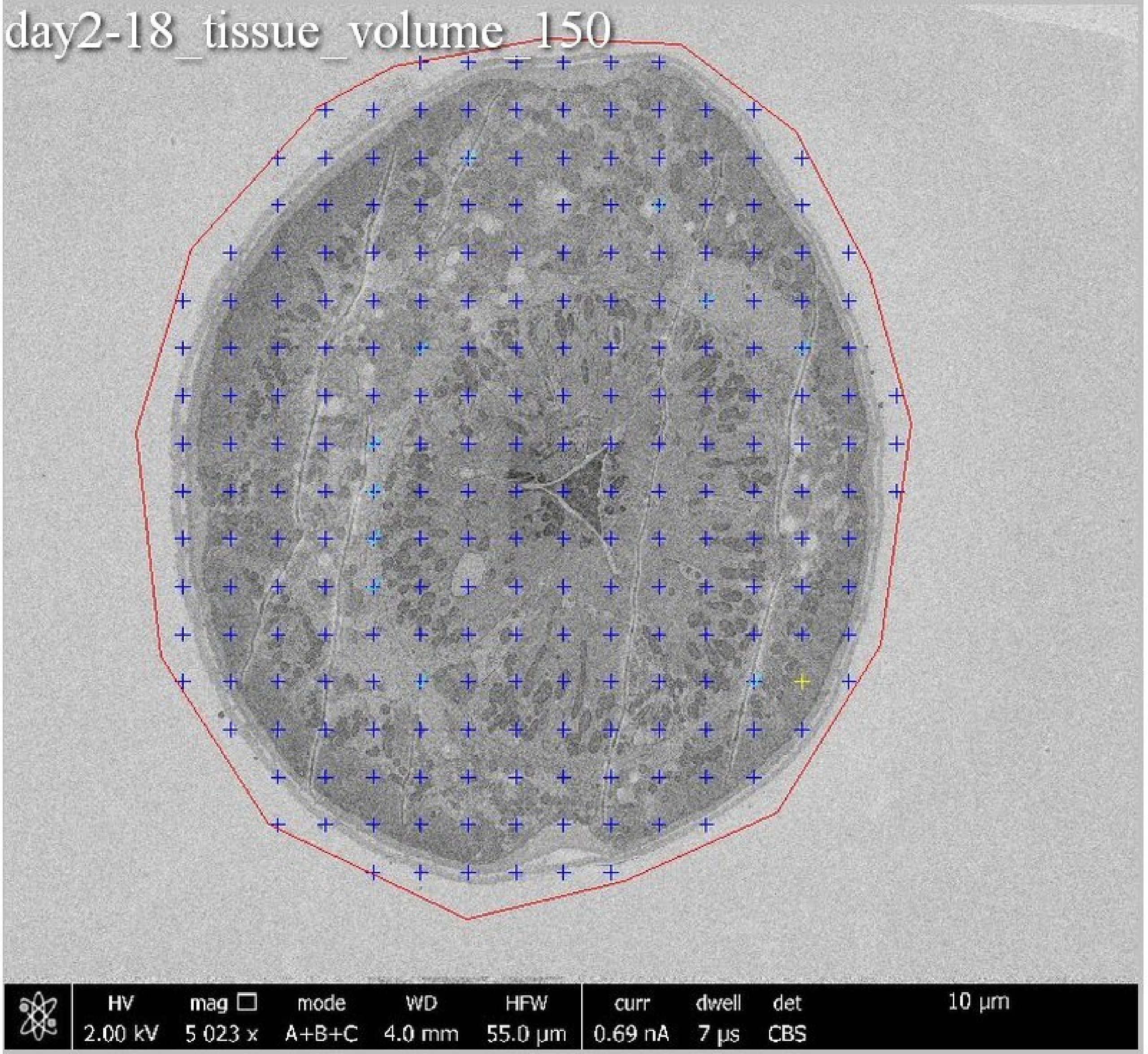
day2-18_tissue_volume_150
HV 2.00 kV
mag 5 023 x
mode A+B+C
WD 4.0 mm
HFW 55.0 µm
curr 0.69 nA
dwell 7 µs
det CBS
10 µm

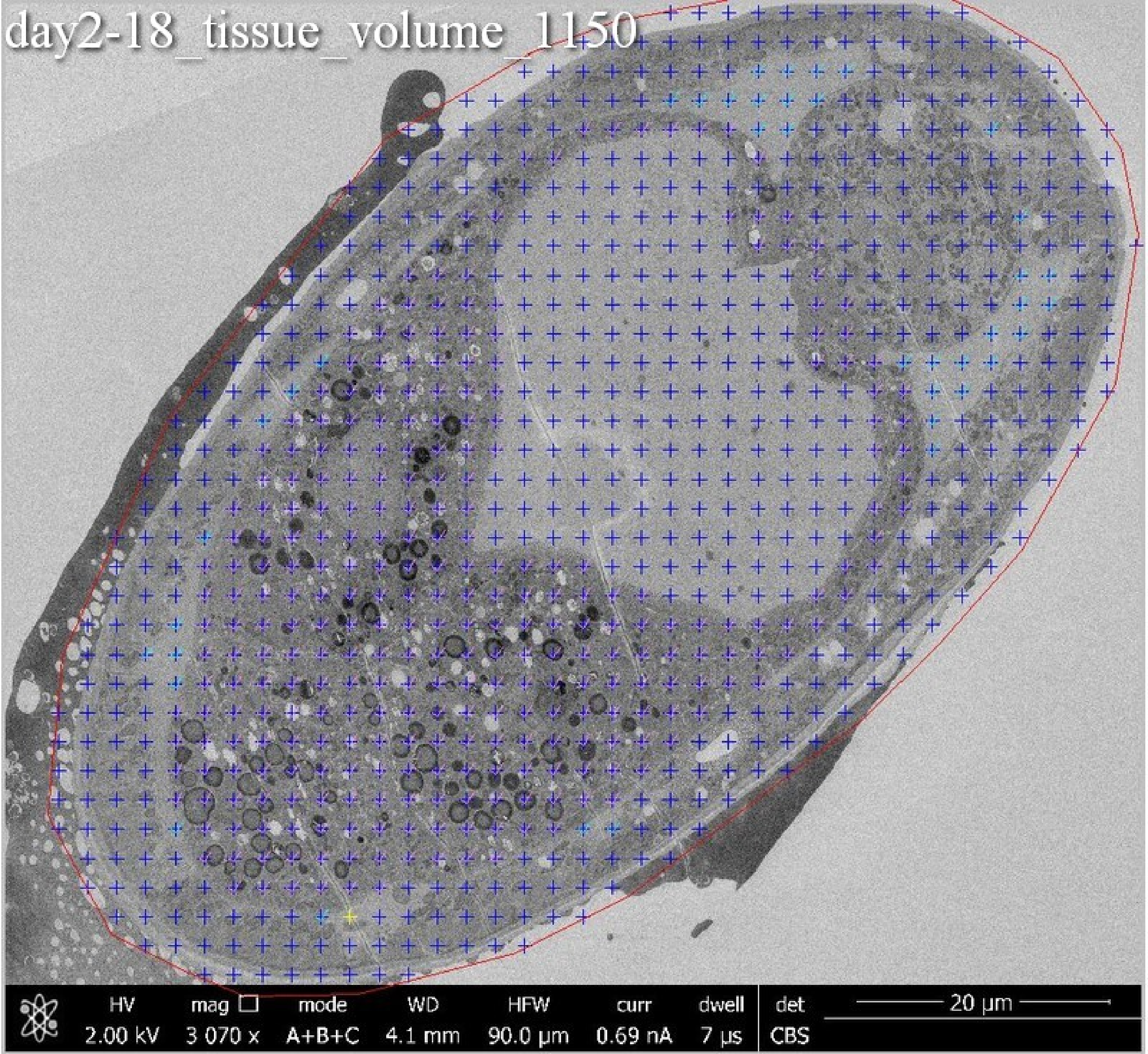
day2-18_tissue_volume_1150
HV
2.00 kV
mag
3 070 x
mode
A+B+C
WD
4.1 mm
HFW
90.0 µm
curr
0.69 nA
dwell
7 µs
det
CBS
20 µm

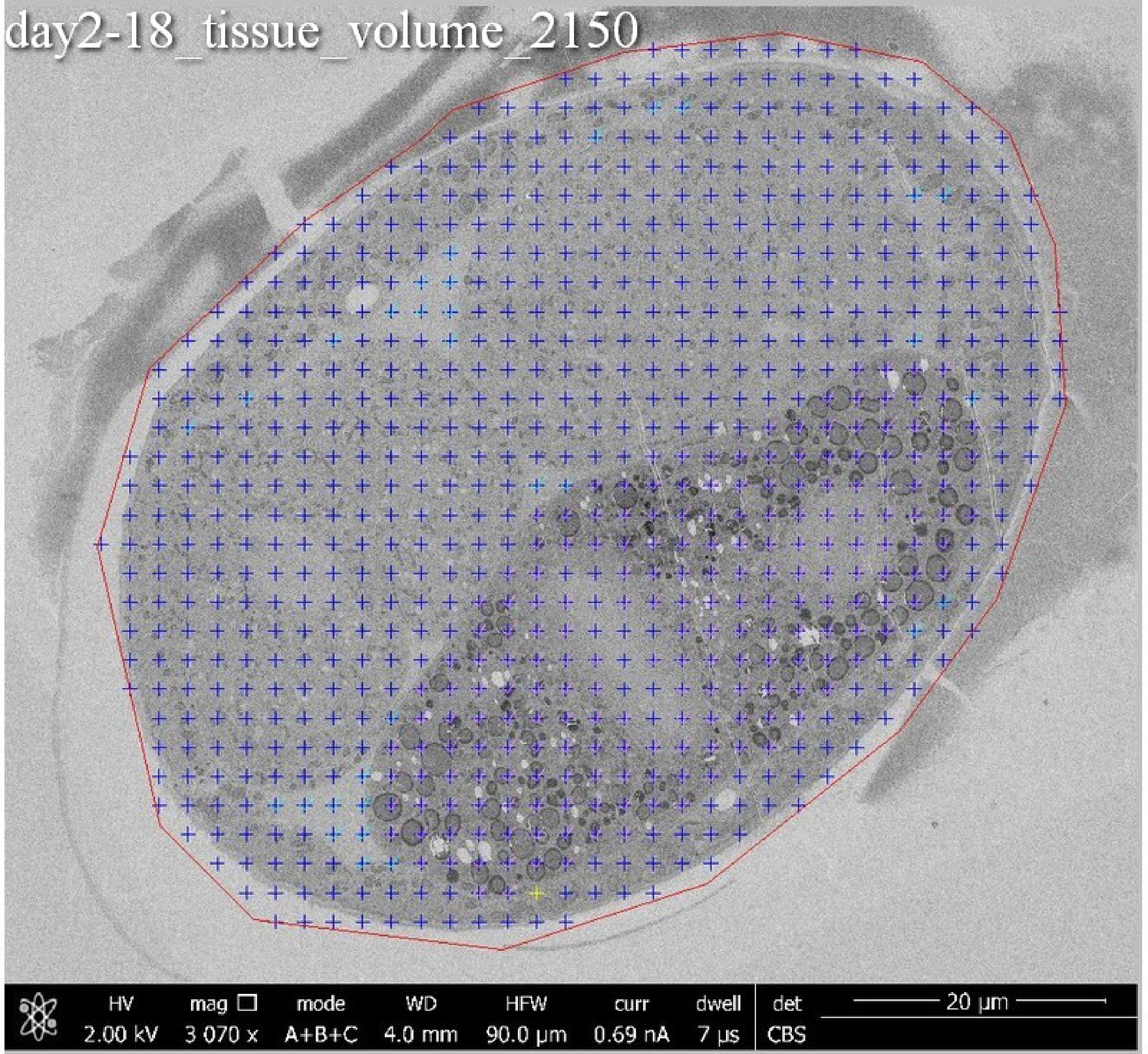
day2-18_tissue_volume_2150
HV 2.00 kV
mag □ 3 070 x
mode A+B+C
WD 4.0 mm
HFW 90.0 μm
curr 0.69 nA
dwell 7 μs
det CBS
20 μm

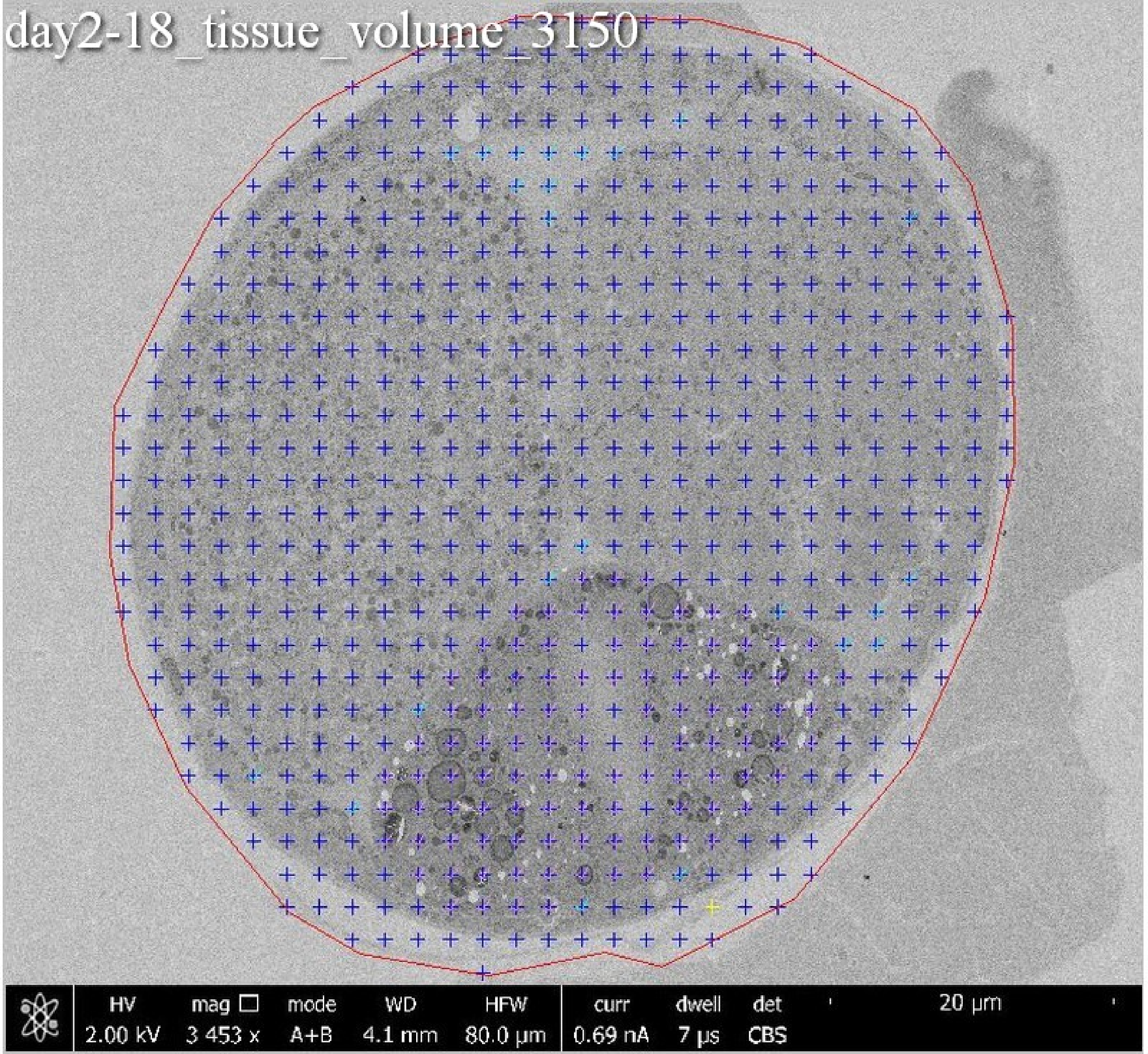
day2-18_tissue_volume_3150
HV
2.00 kV
mag
3 453 x
mode
A+B
WD
4.1 mm
HFW
80.0 µm
curr
0.69 nA
dwell
7 µs
det
CBS
20 µm

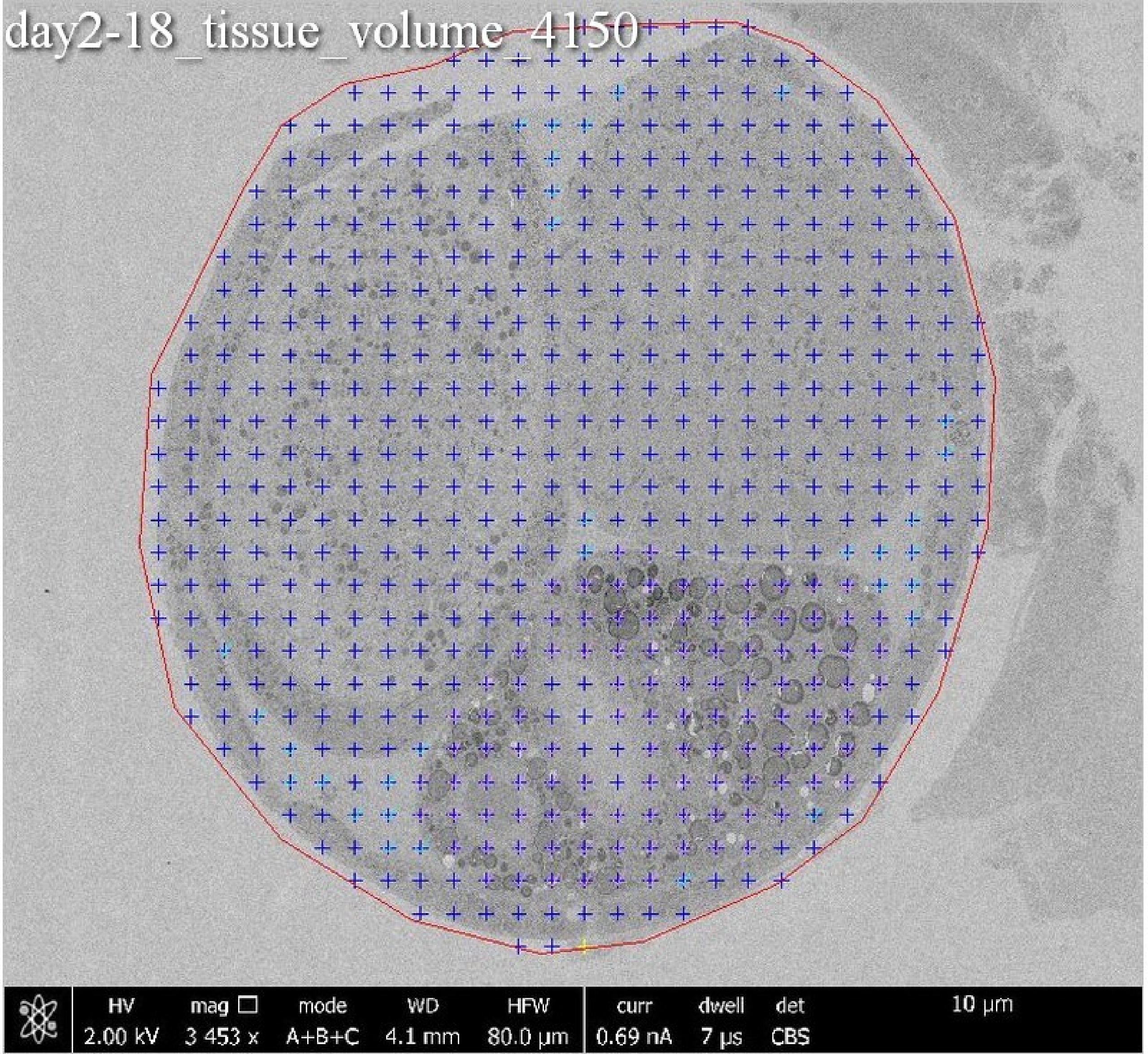
day2-18_tissue_volume_4150
HV 2.00 kV
mag 3 453 x
mode A+B+C
WD 4.1 mm
HFW 80.0 µm
curr 0.69 nA
dwell 7 µs
det CBS
10 µm

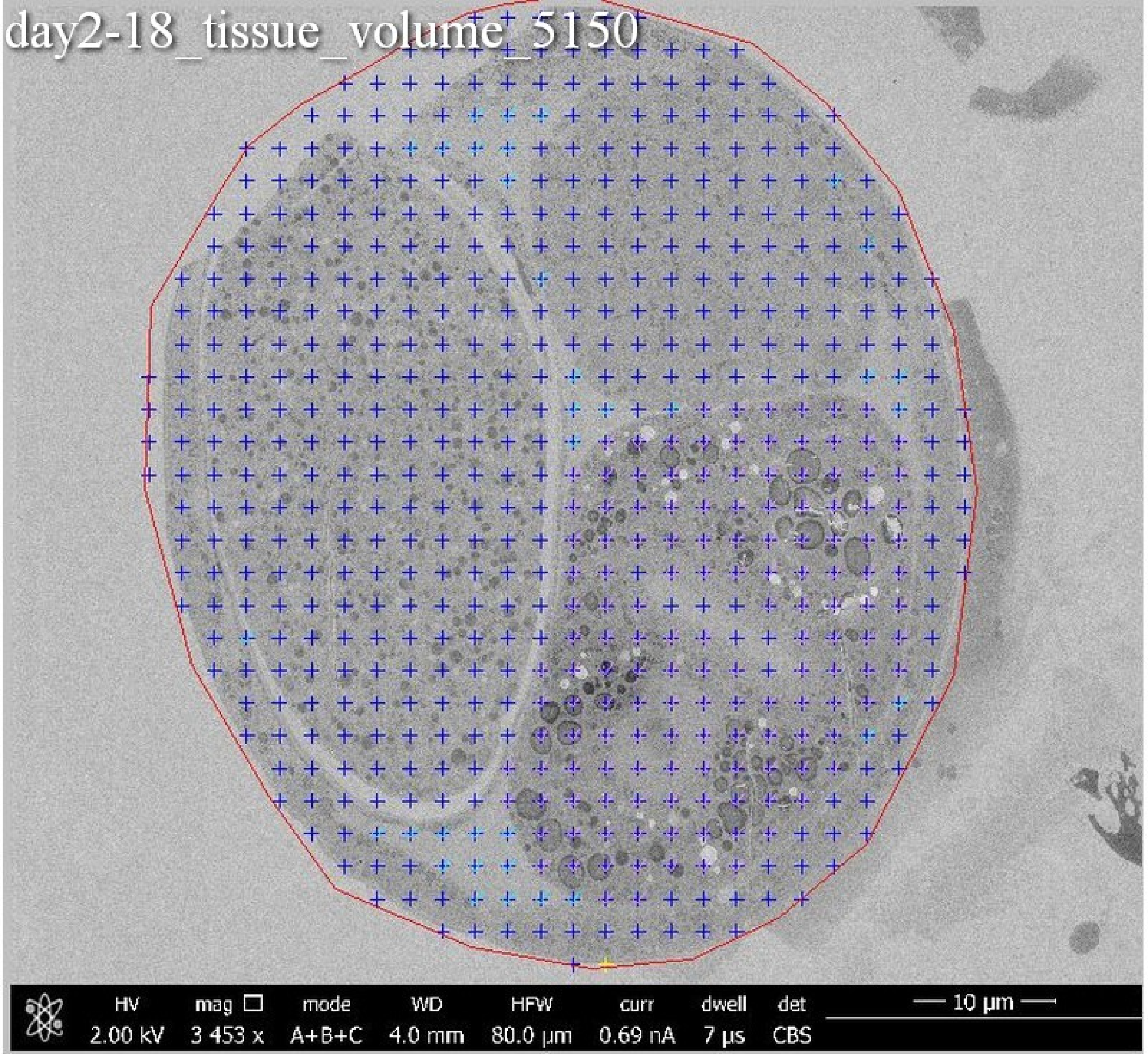
day2-18_tissue_volume_5150
HV 2.00 kV
mag 3 453 x
mode A+B+C
WD 4.0 mm
HFW 80.0 µm
curr 0.69 nA
dwell 7 µs
det CBS
10 µm

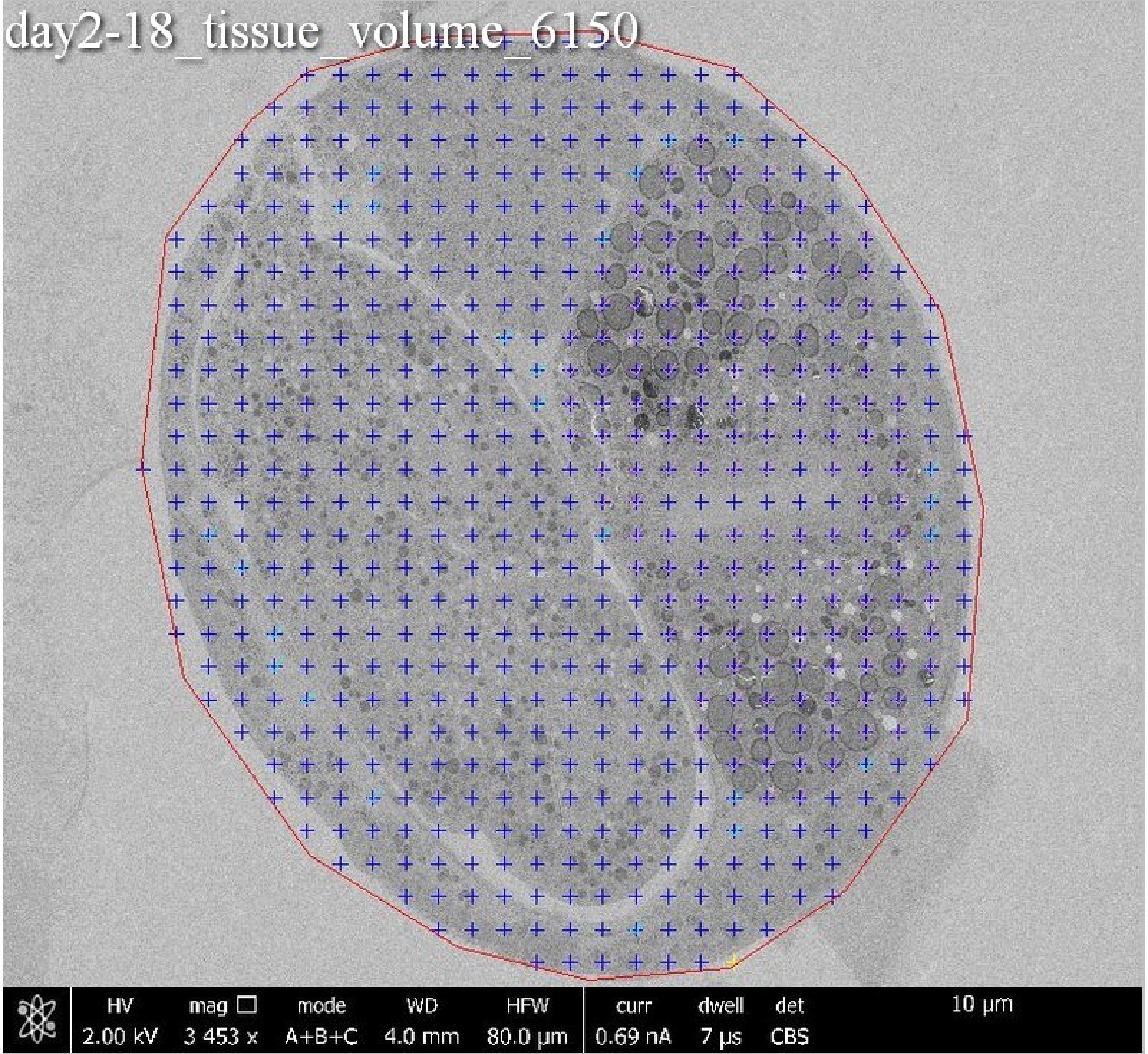

day2-18_tissue_volume_6150
HV
2.00 kV
mag
3 453 x
mode
A+B+C
WD
4.0 mm
HFW
80.0 μm
curr
0.69 nA
dwell
7 μs
det
CBS
10 μm

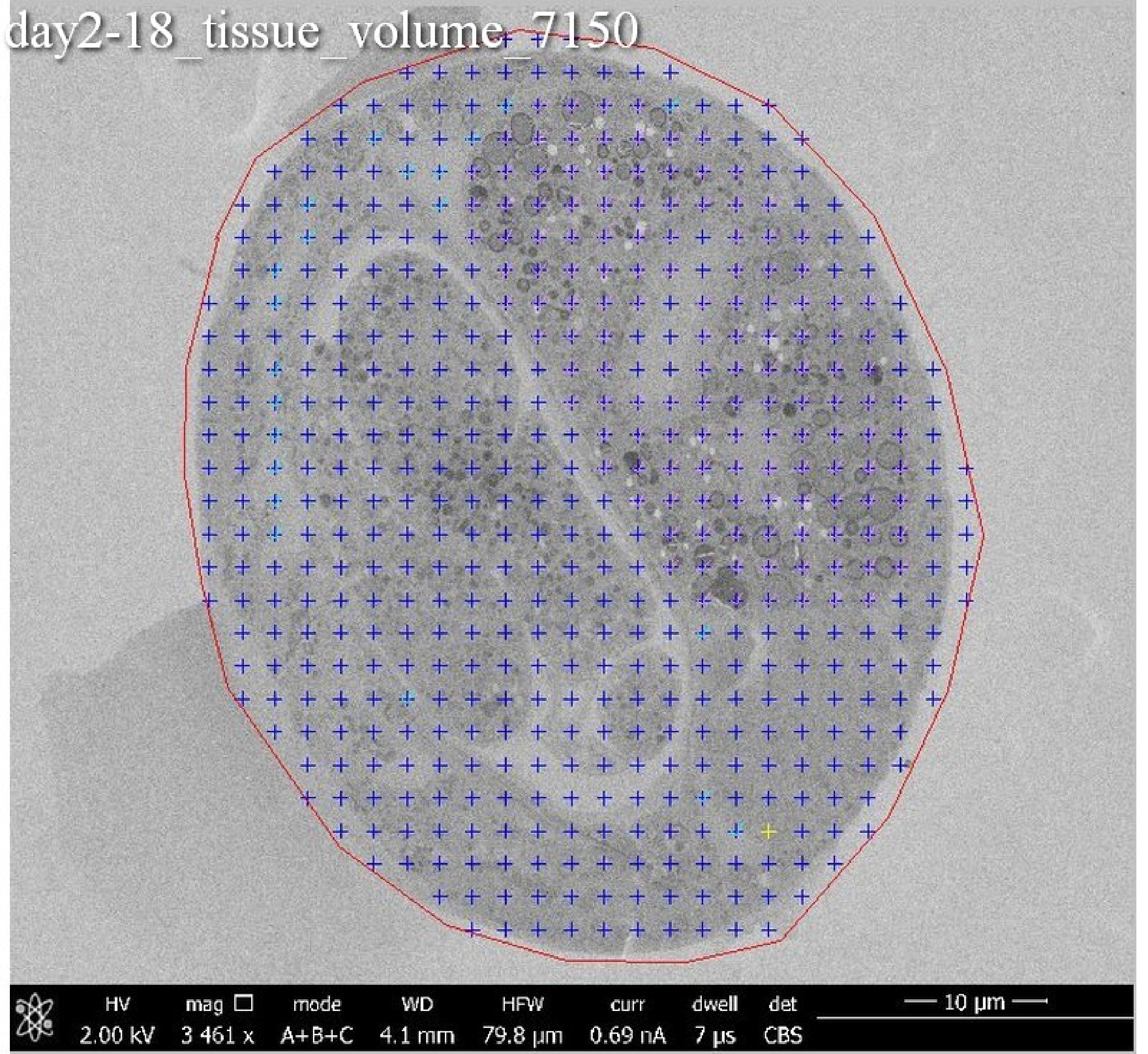
day2-18_tissue_volume_7150
HV 2.00 kV
mag 3 461 x
mode A+B+C
WD 4.1 mm
HFW 79.8 µm
curr 0.69 nA
dwell 7 µs
det CBS
10 µm

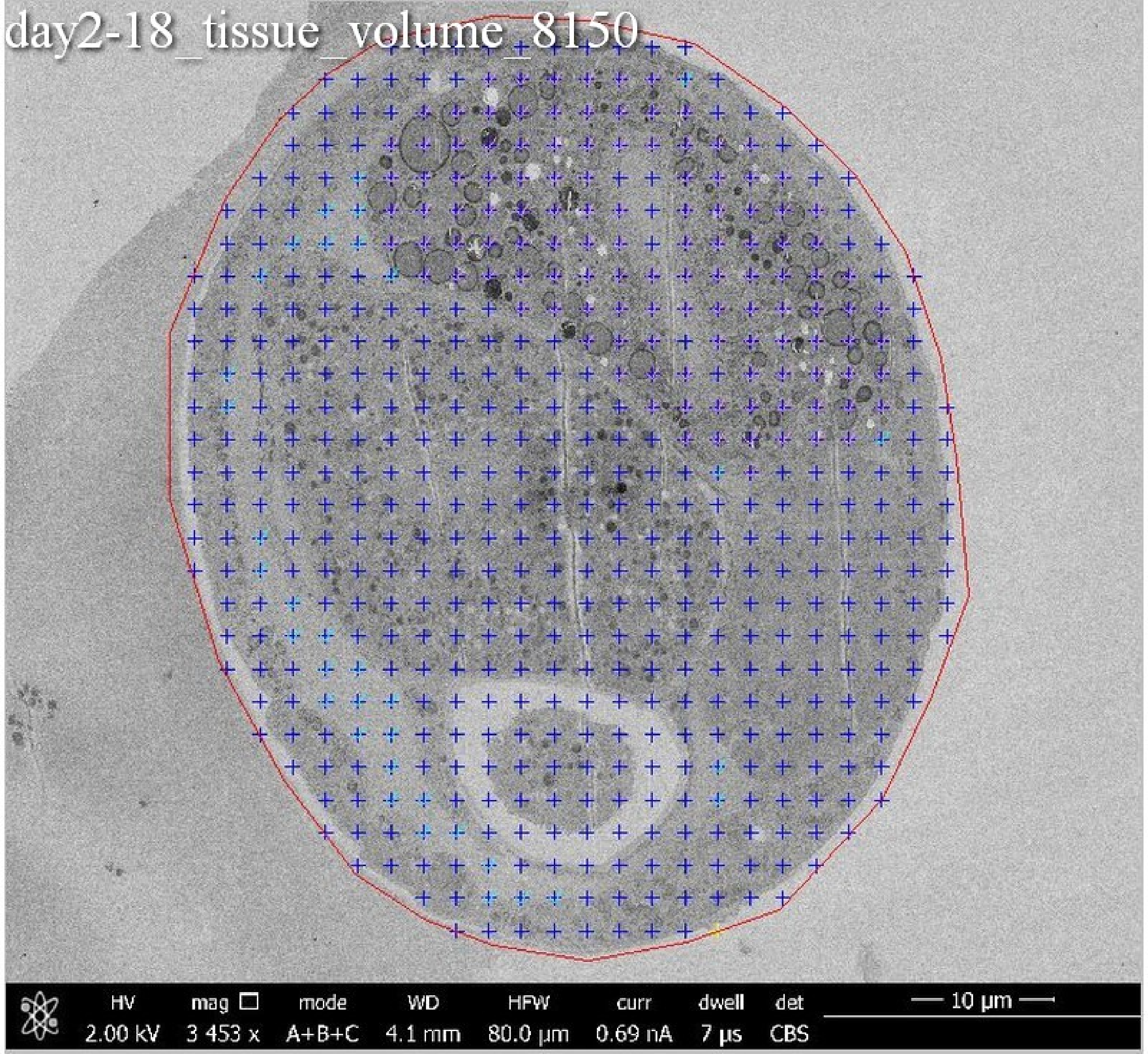
day2-18_tissue_volume_8150
HV 2.00 kV
mag 3 453 x
mode A+B+C
WD 4.1 mm
HFW 80.0 µm
curr 0.69 nA
dwell 7 µs
det CBS
10 µm

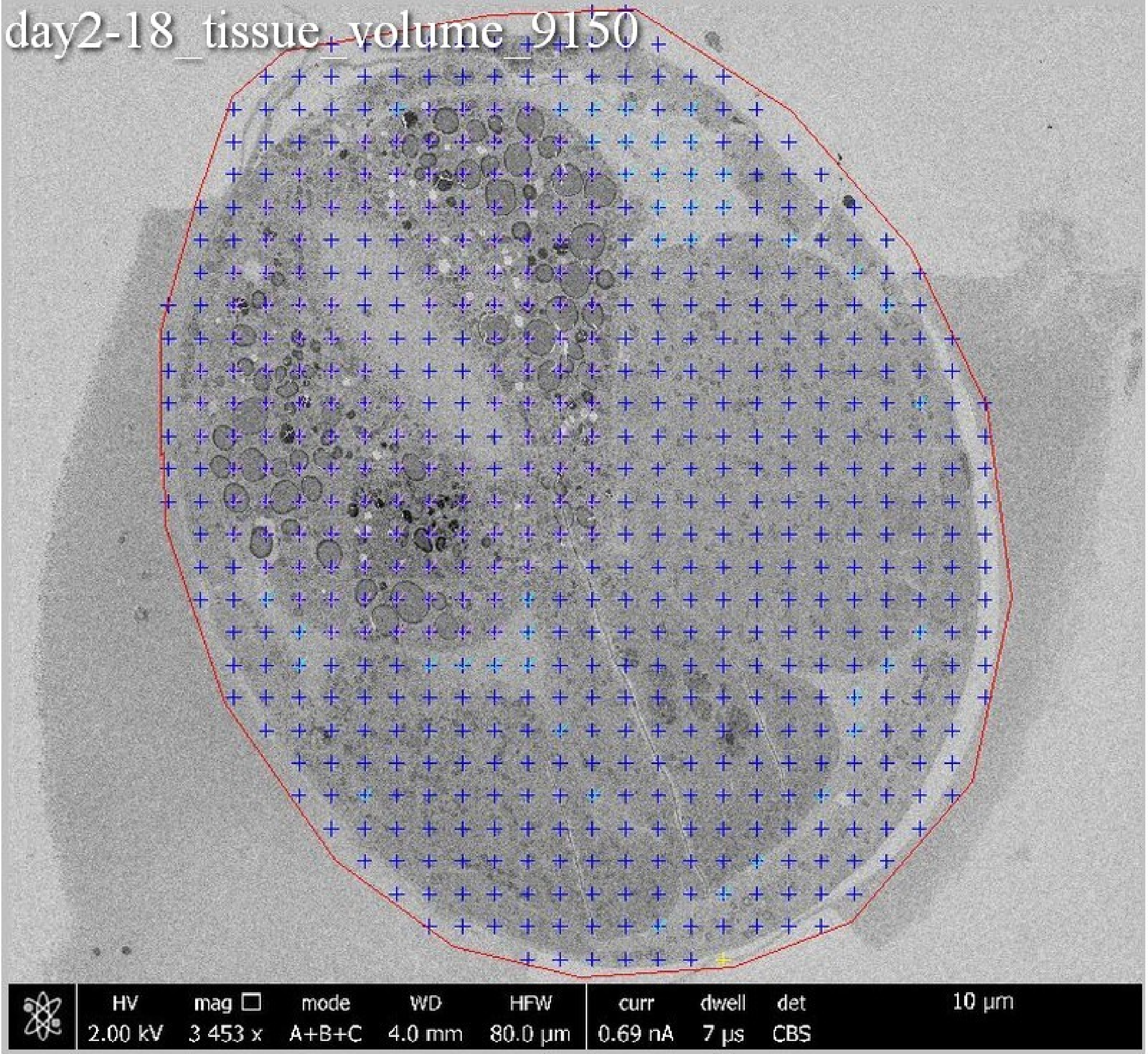
day2-18_tissue_volume_9150
HV
2.00 kV
mag
3 453 x
mode
A+B+C
WD
4.0 mm
HFW
80.0 µm
curr
0.69 nA
dwell
7 µs
det
CBS
10 µm

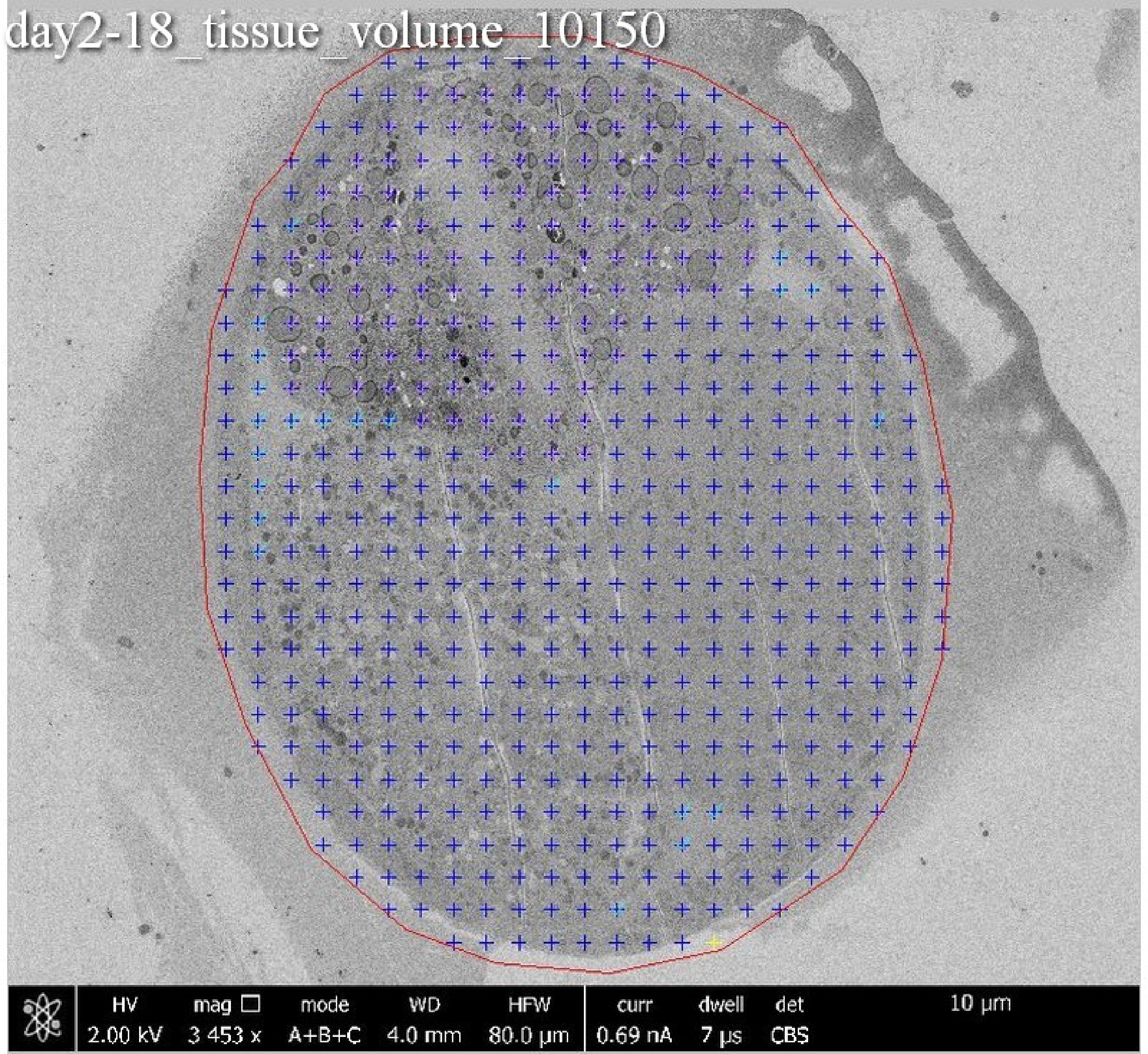
day2-18_tissue_volume_10150
HV 2.00 kV
mag 3 453 x
mode A+B+C
WD 4.0 mm
HFW 80.0 µm
curr 0.69 nA
dwell 7 µs
det CBS
10 µm

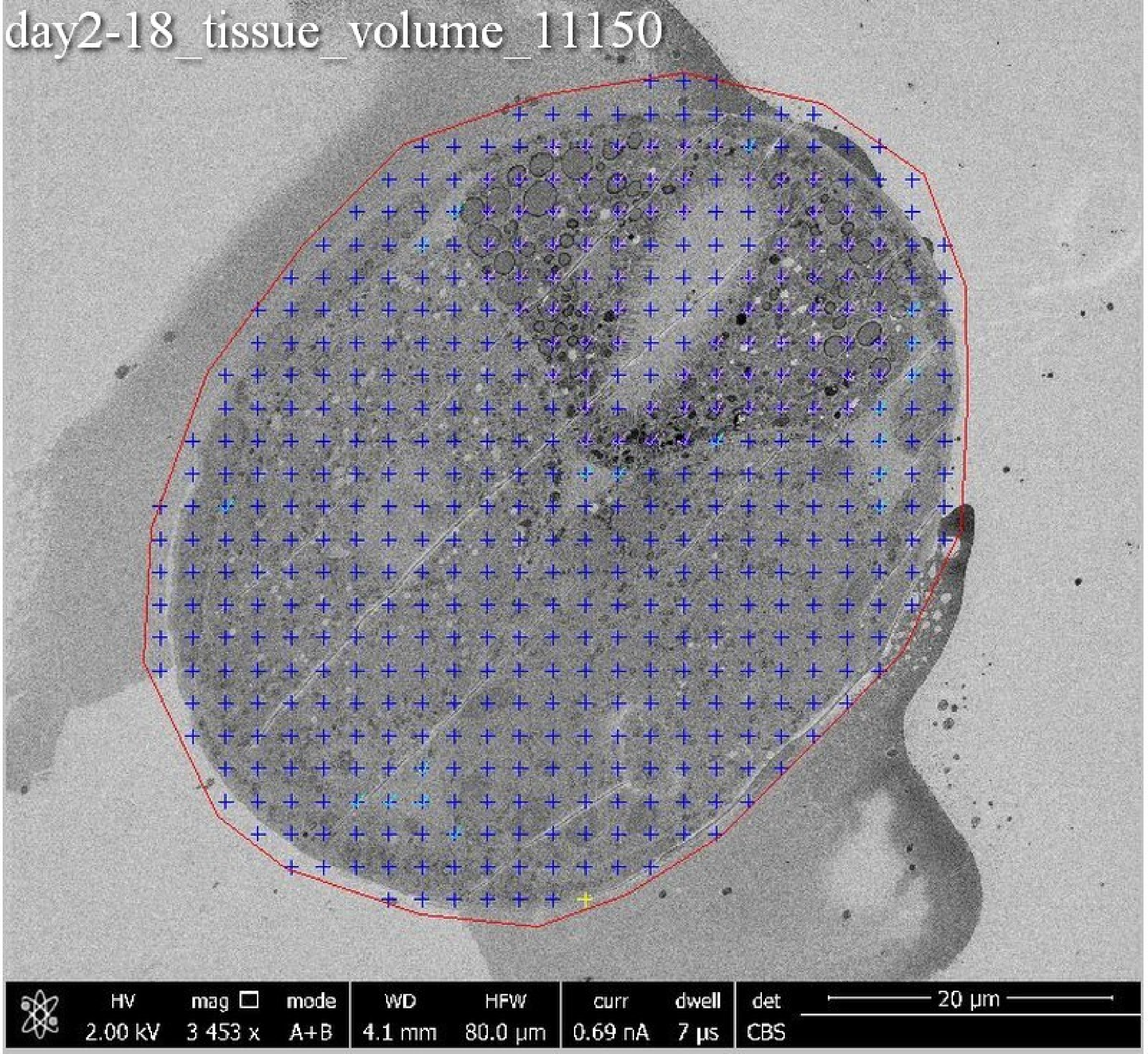

day2-18_tissue_volume_11150
HV
2.00 kV
mag
3 453 x
mode
A+B
WD
4.1 mm
HFW
80.0 µm
curr
0.69 nA
dwell
7 µs
det
CBS
20 µm

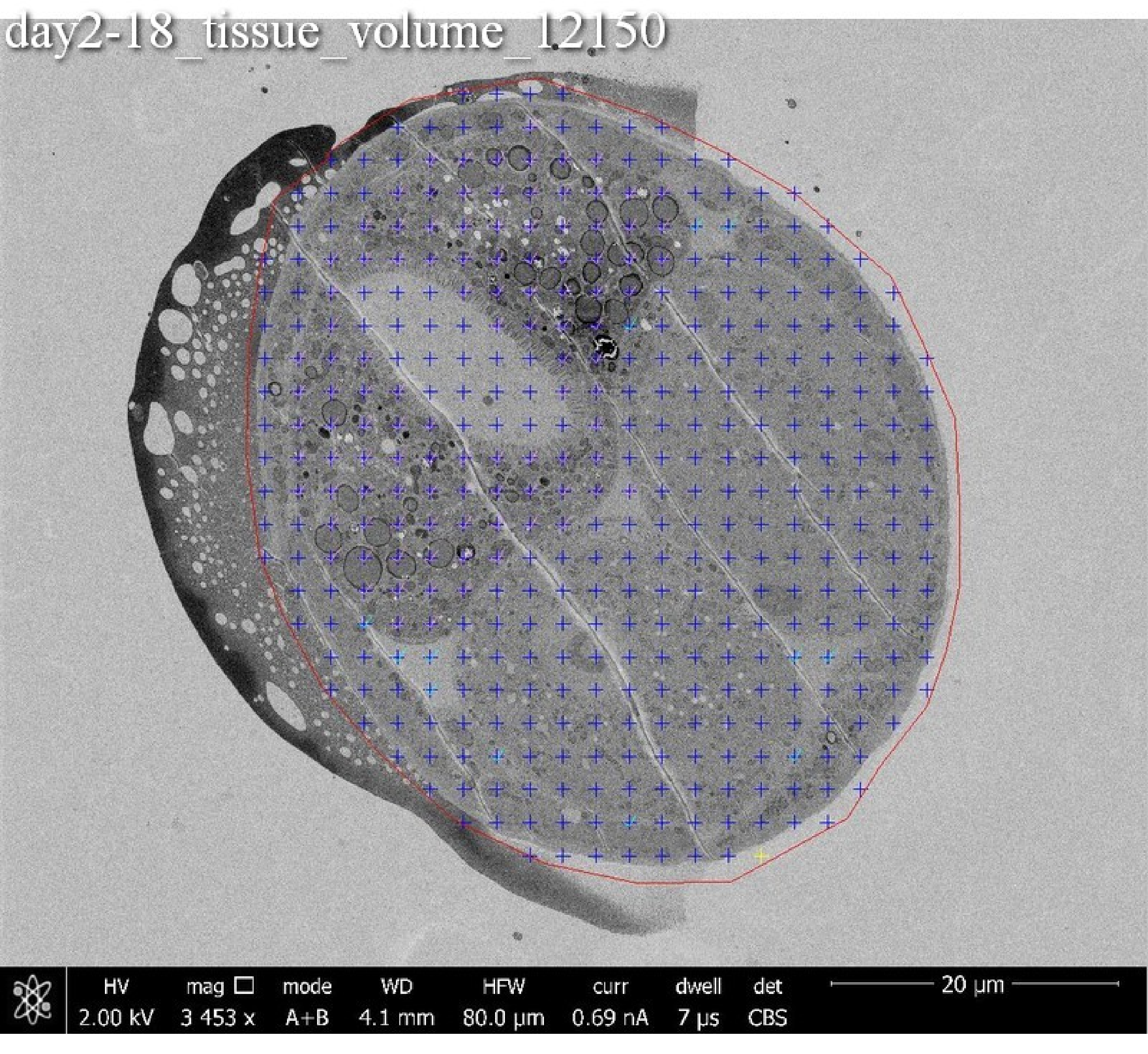

day2-18_tissue_volume_12150
HV 2.00 kV
mag ☐ 3 453 x
mode A+B
WD 4.1 mm
HFW 80.0 μm
curr 0.69 nA
dwell 7 μs
det CBS
20 μm

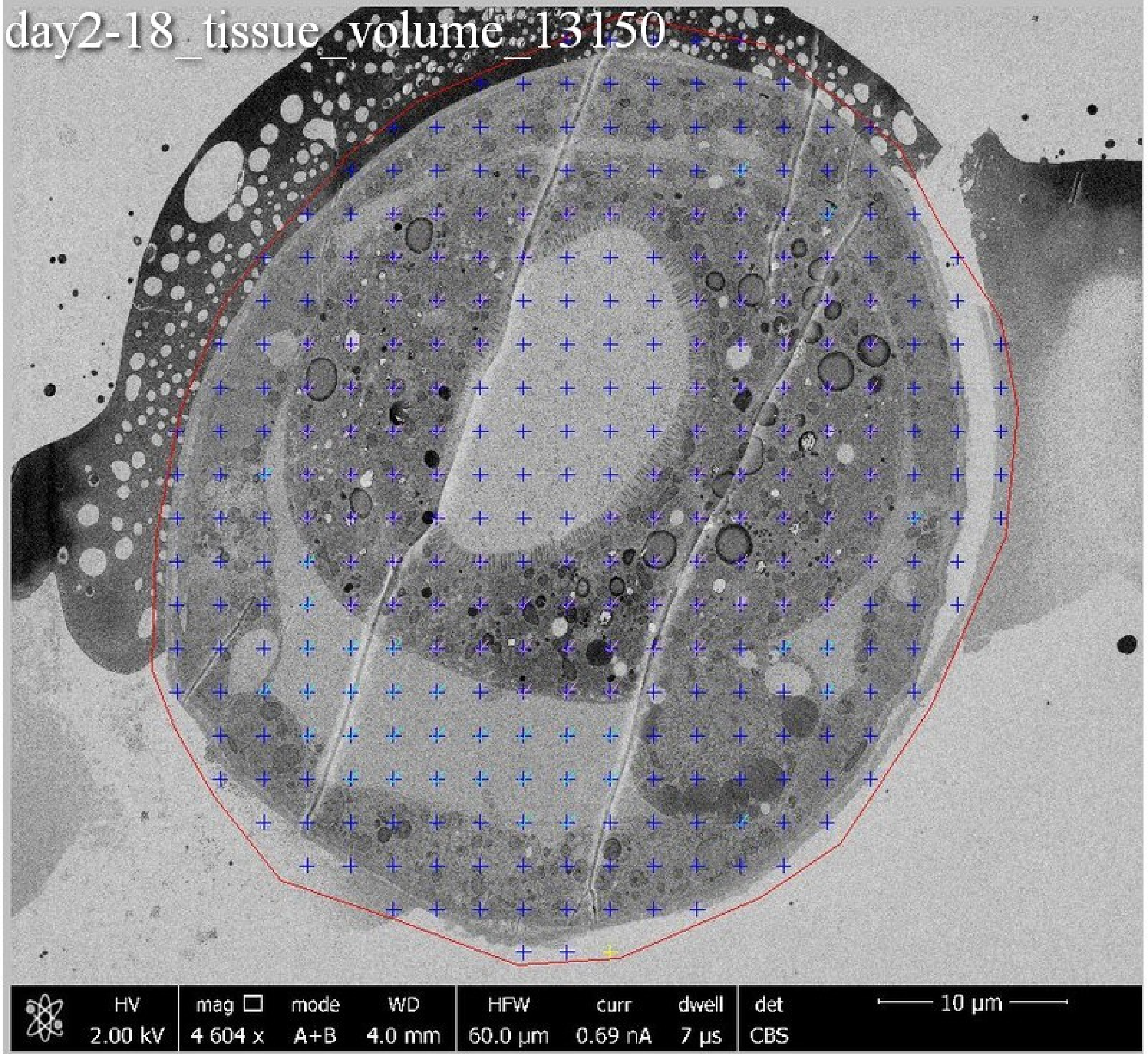
day2-18_tissue_volume_13150
HV
2.00 kV
mag
4 604 x
mode
A+B
WD
4.0 mm
HFW
60.0 μm
curr
0.69 nA
dwell
7 μs
det
CBS
10 μm

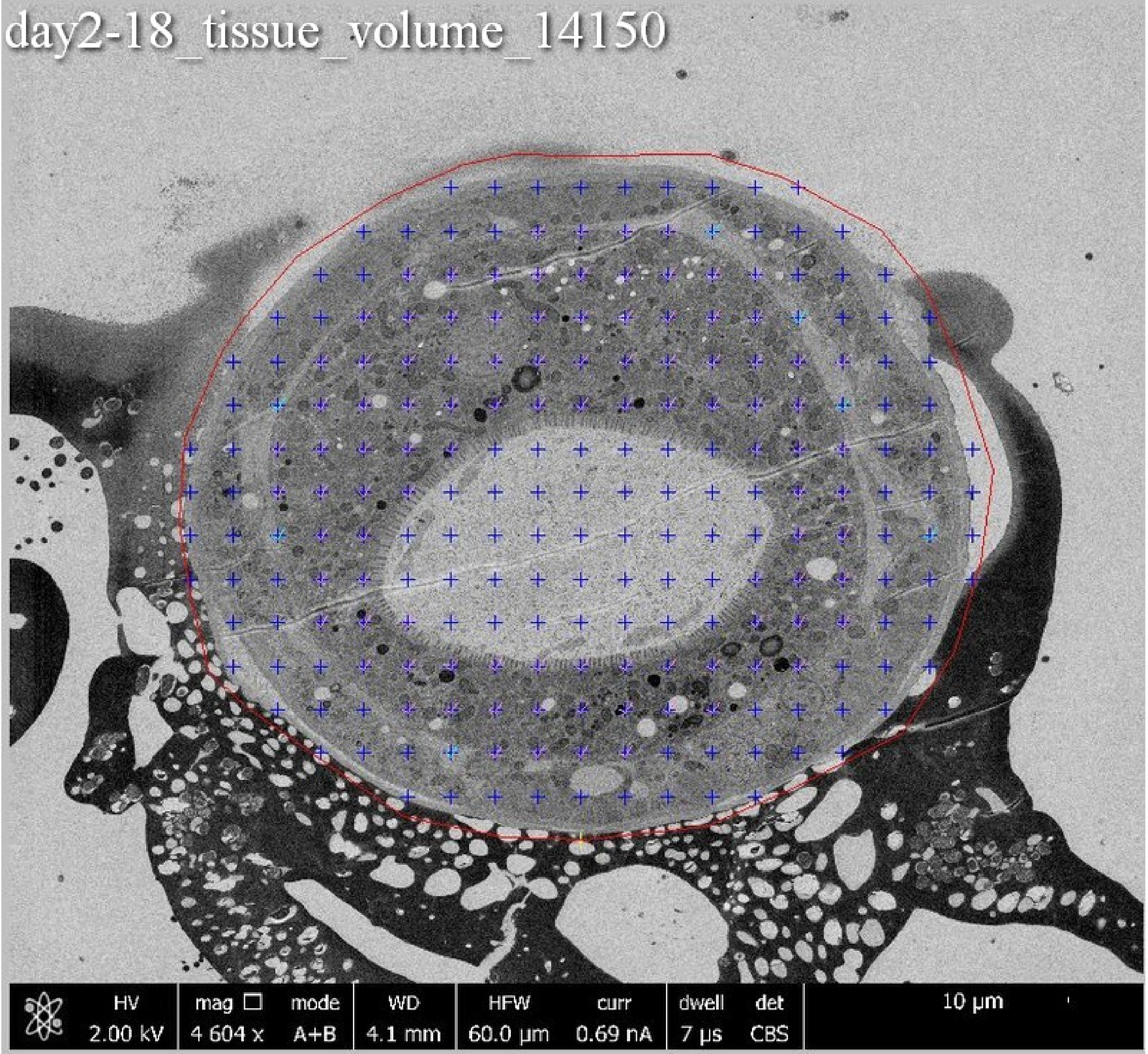
day2-18_tissue_volume_14150
HV 2.00 kV
mag 4 604 x
mode A+B
WD 4.1 mm
HFW 60.0 µm
curr 0.69 nA
dwell 7 µs
det CBS
10 µm

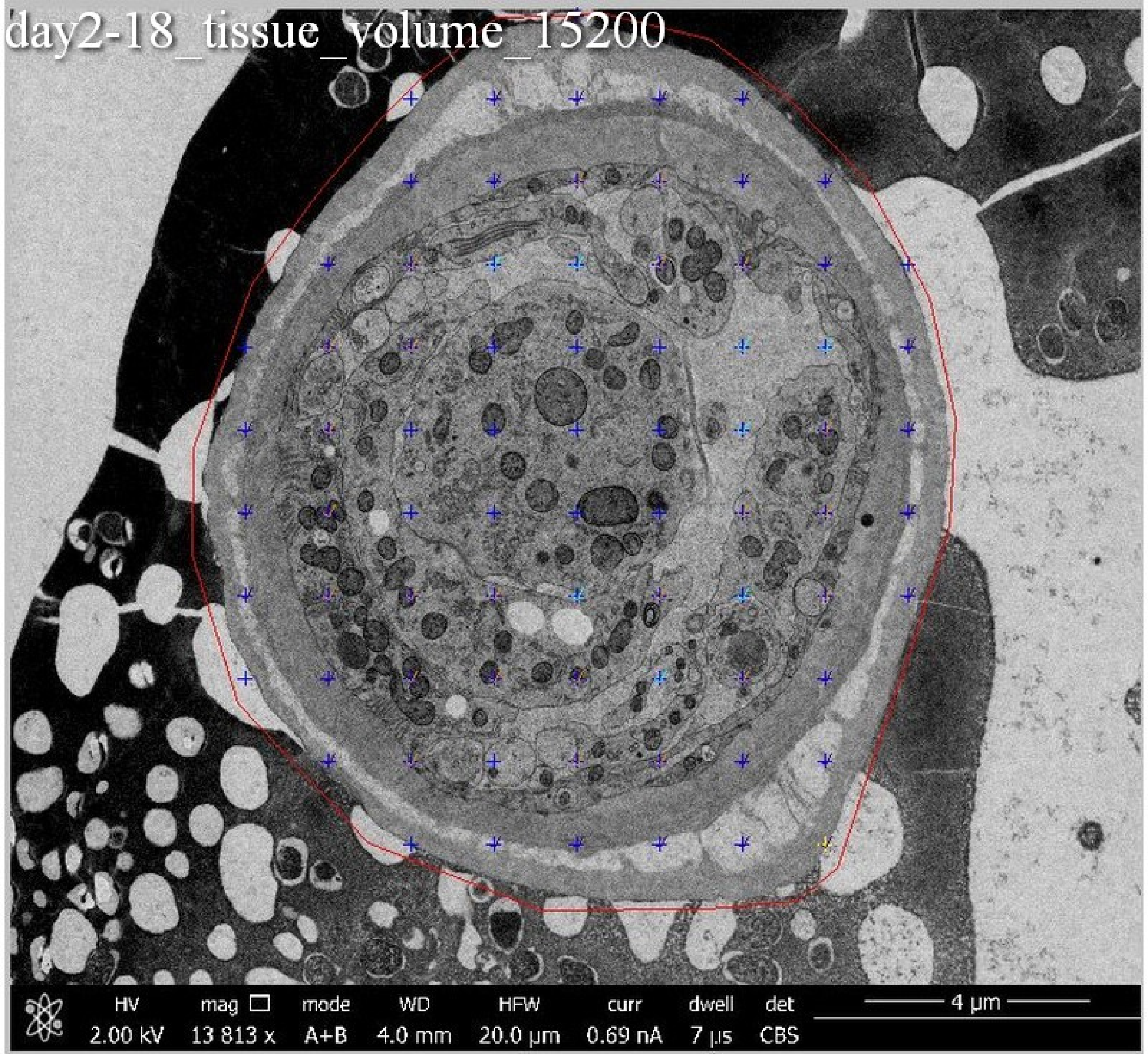
day2-18_tissue_volume_15200
HV 2.00 kV
mag 13 813 x
mode A+B
WD 4.0 mm
HFW 20.0 µm
curr 0.69 nA
dwell 7 µs
det CBS
4 µm

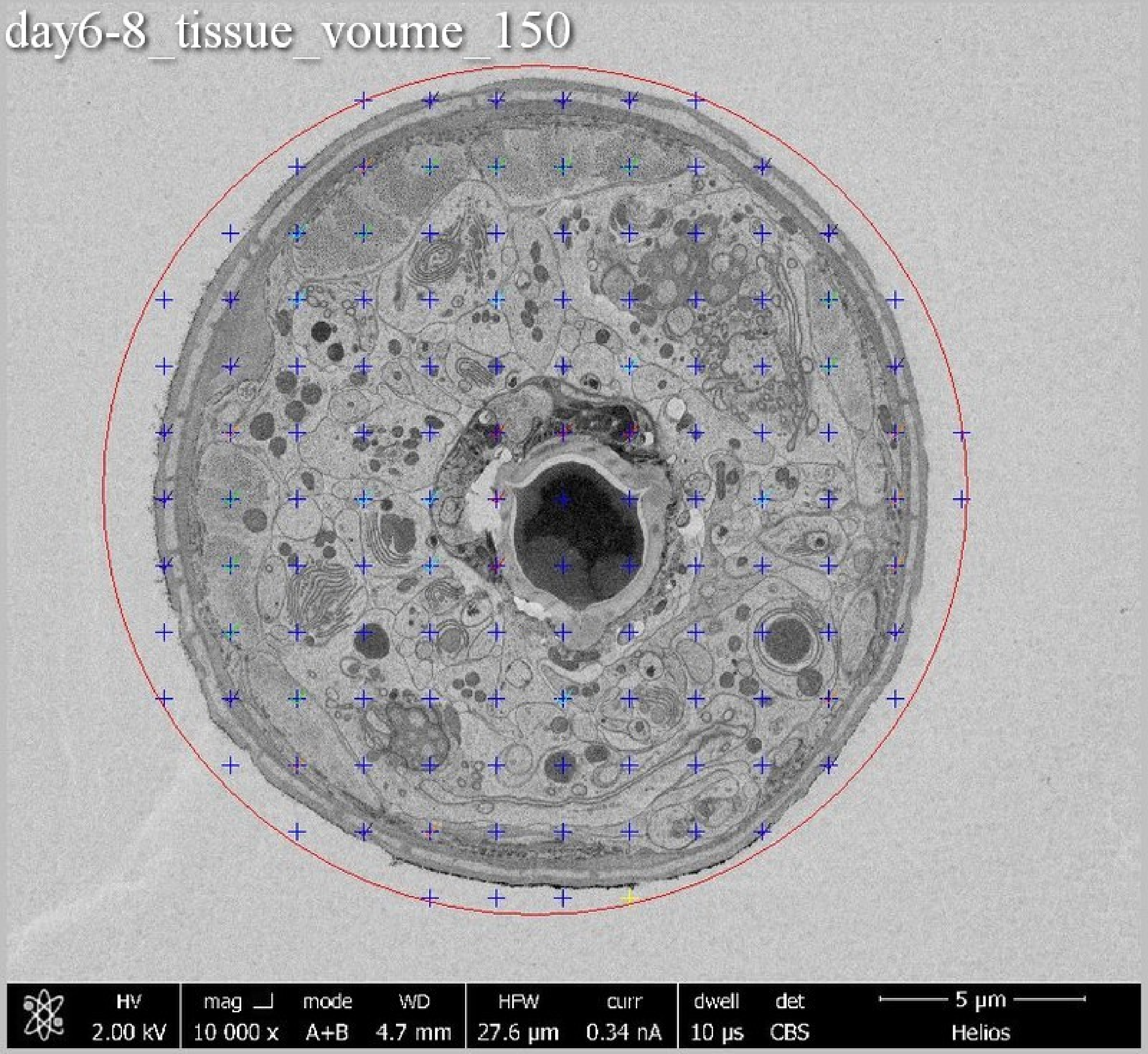
day6-8_tissue_voume_150
HV
2.00 kV
mag
10 000 x
mode
A+B
WD
4.7 mm
HFW
27.6 µm
curr
0.34 nA
dwell
10 µs
det
CBS
5 µm
Helios

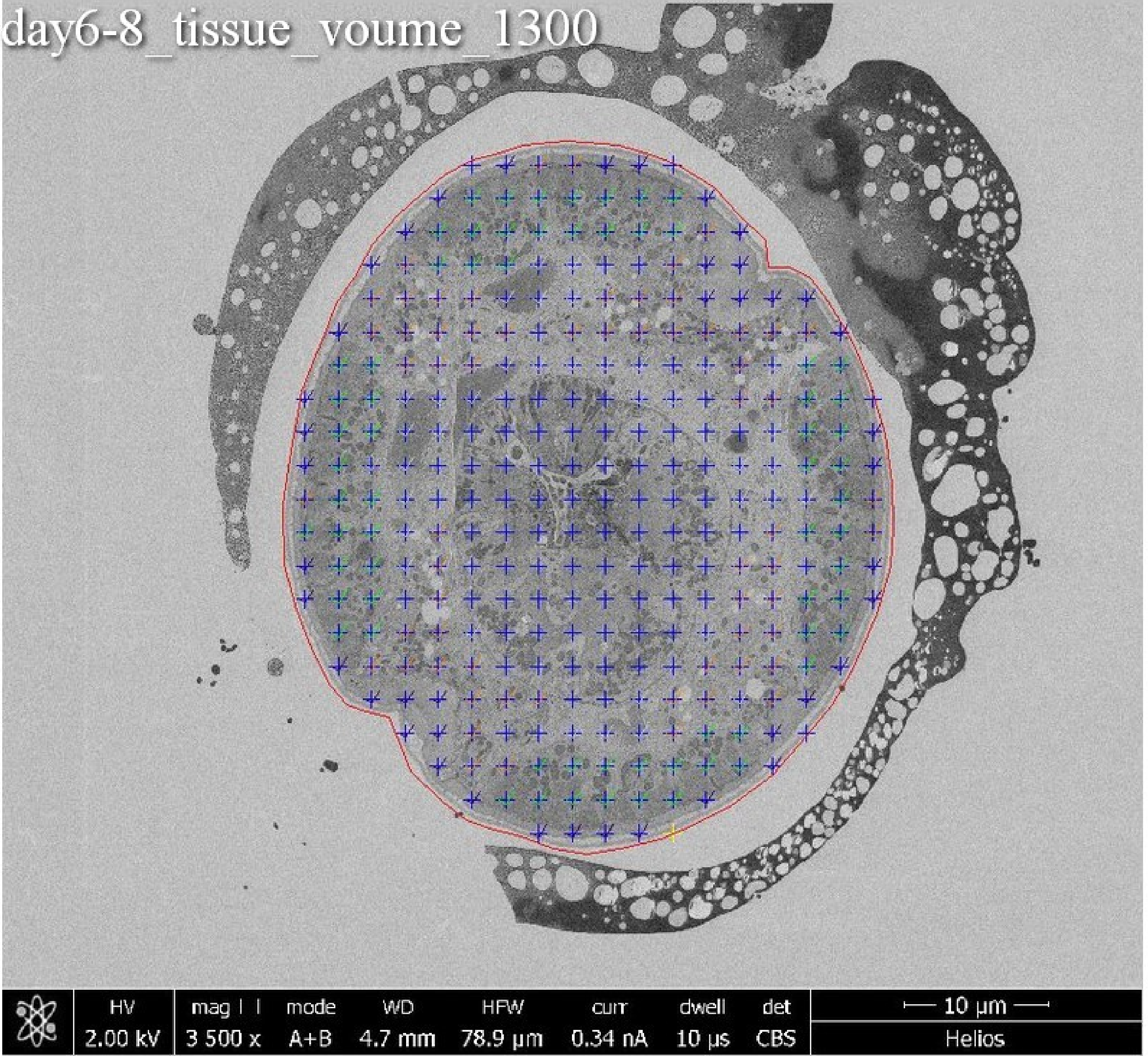
day6-8_tissue_voume_1300
HV
2.00 kV
mag
3 500 x
mode
A+B
WD
4.7 mm
HFW
78.9 µm
curr
0.34 nA
dwell
10 µs
det
CBS
10 µm
Helios

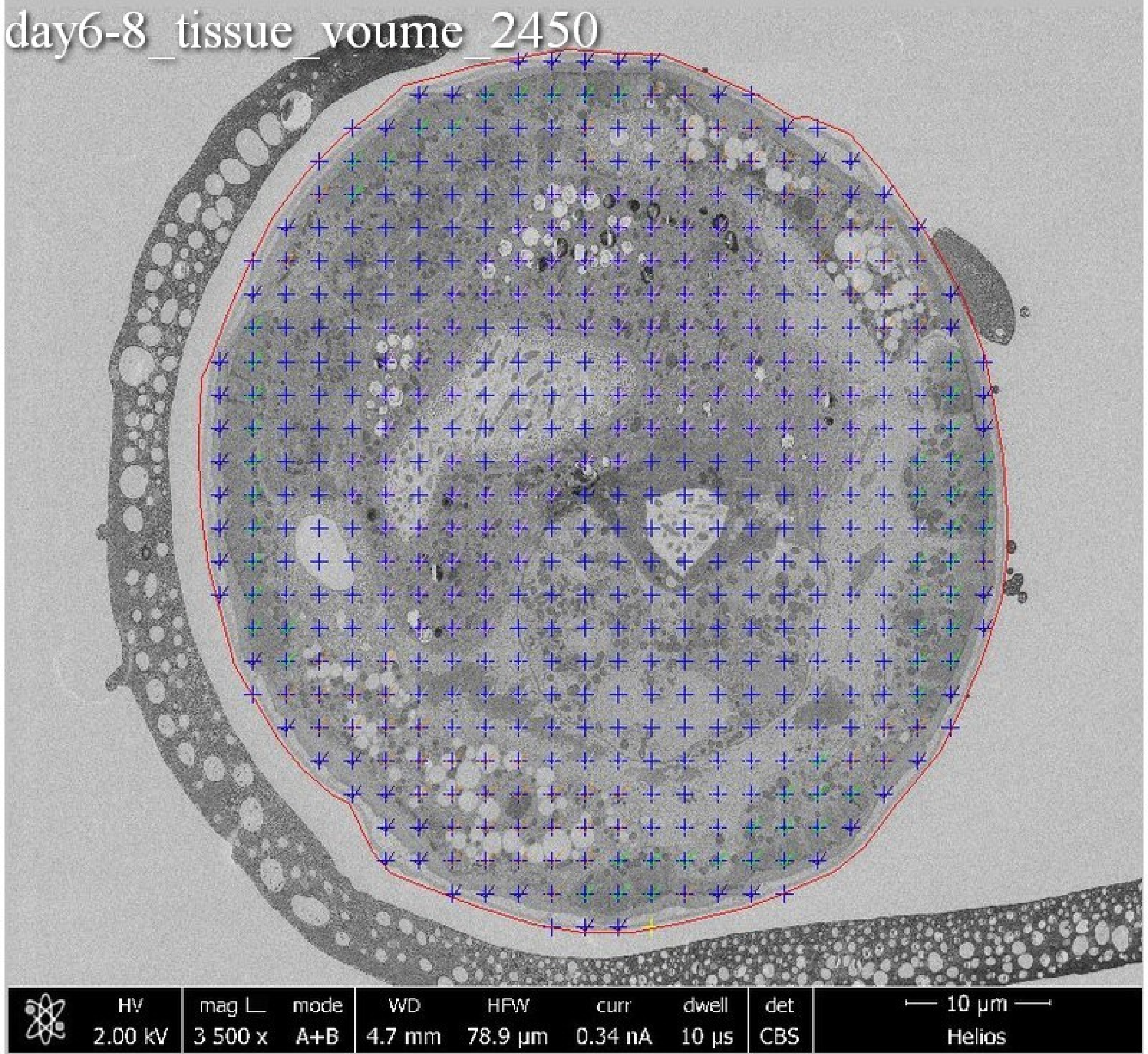
day6-8_tissue_voume_2450
HV
2.00 kV
mag L
3 500 x
mode
A+B
WD
4.7 mm
HFW
78.9 µm
curr
0.34 nA
dwell
10 µs
det
CBS
10 µm
Helios

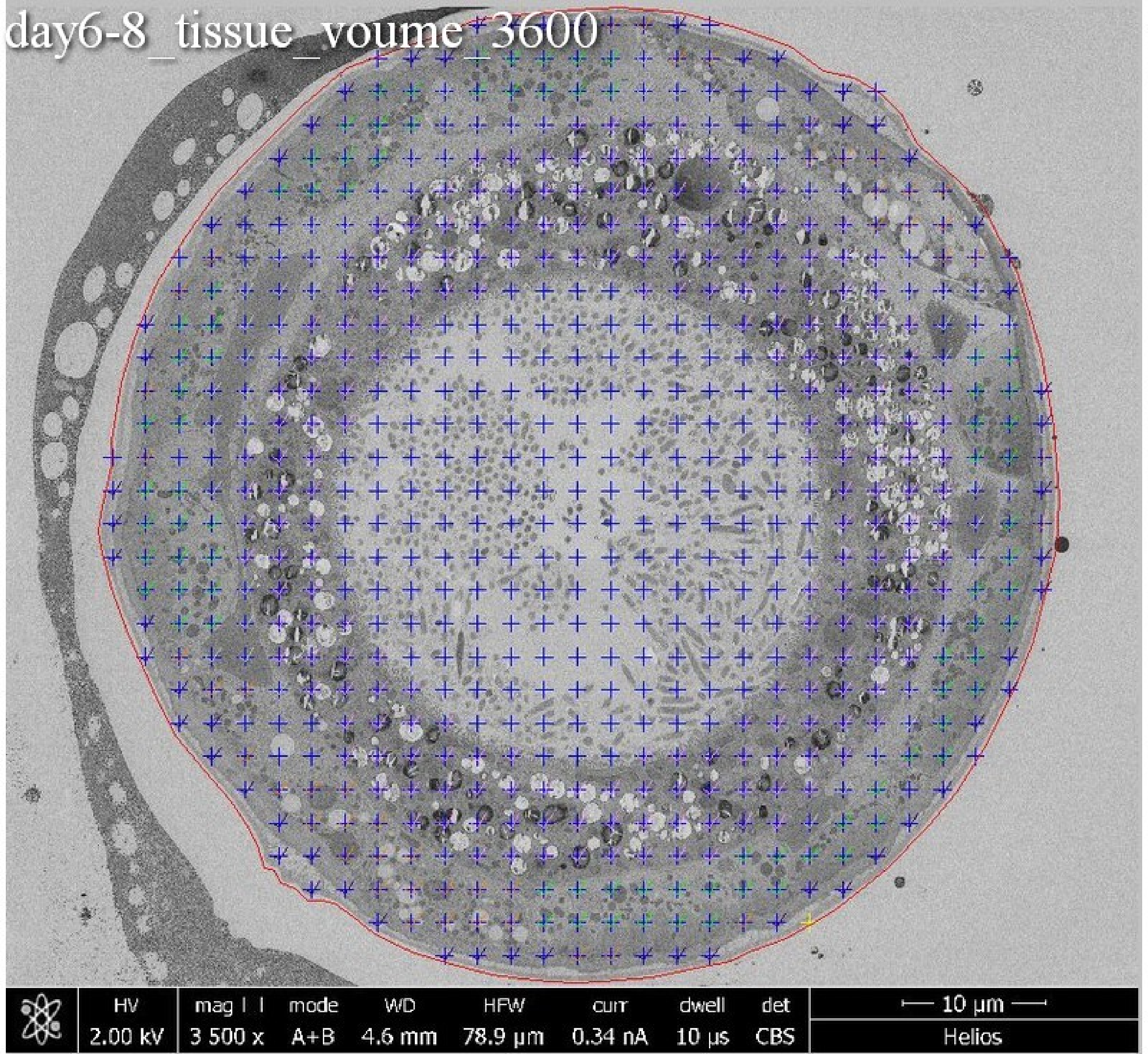
day6-8_tissue_voume_3600
HV
2.00 kV
mag
3 500 x
mode
A+B
WD
4.6 mm
HFW
78.9 µm
curr
0.34 nA
dwell
10 µs
det
CBS
10 µm
Helios

day6-8_tissue_voume_4750

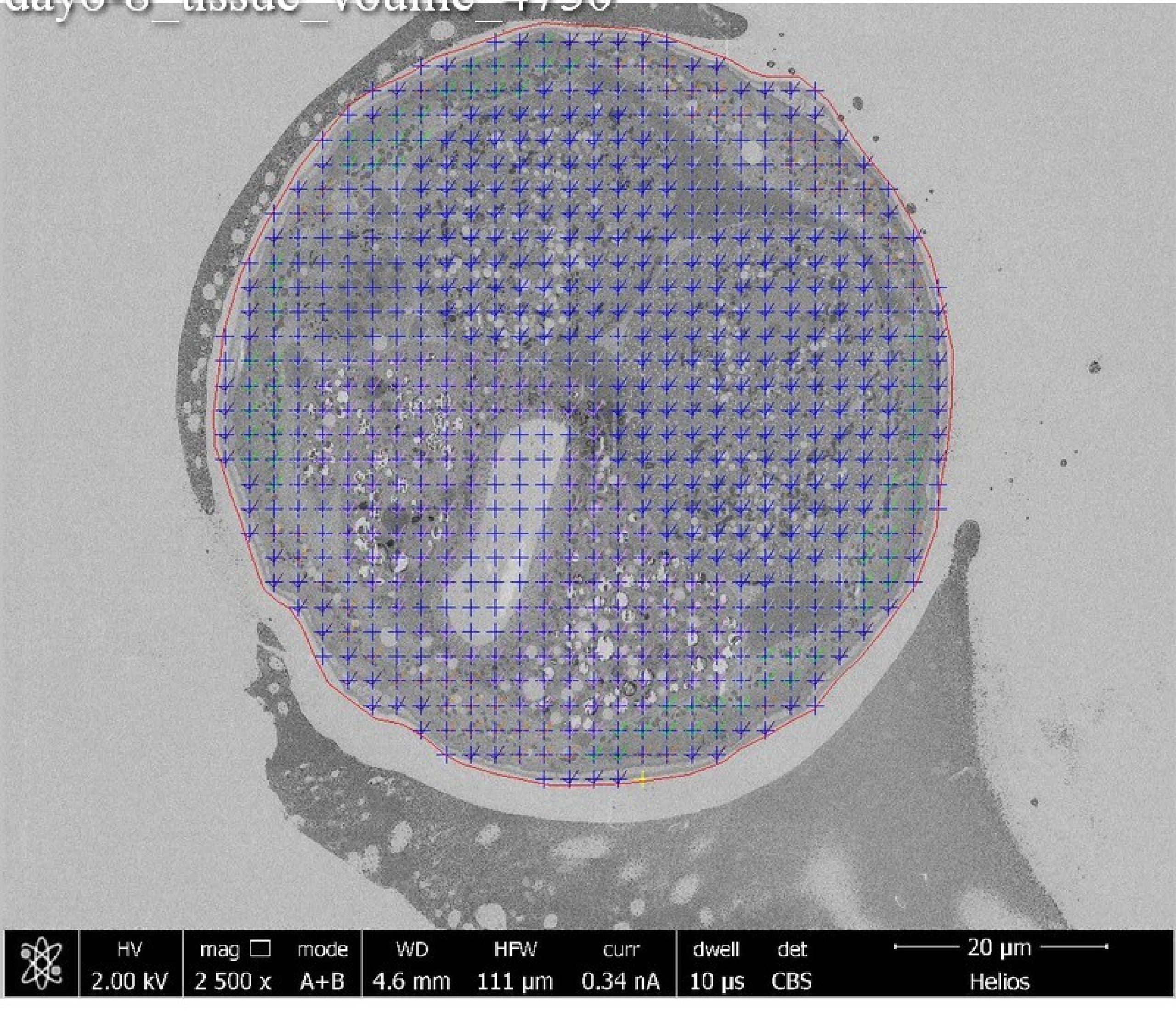

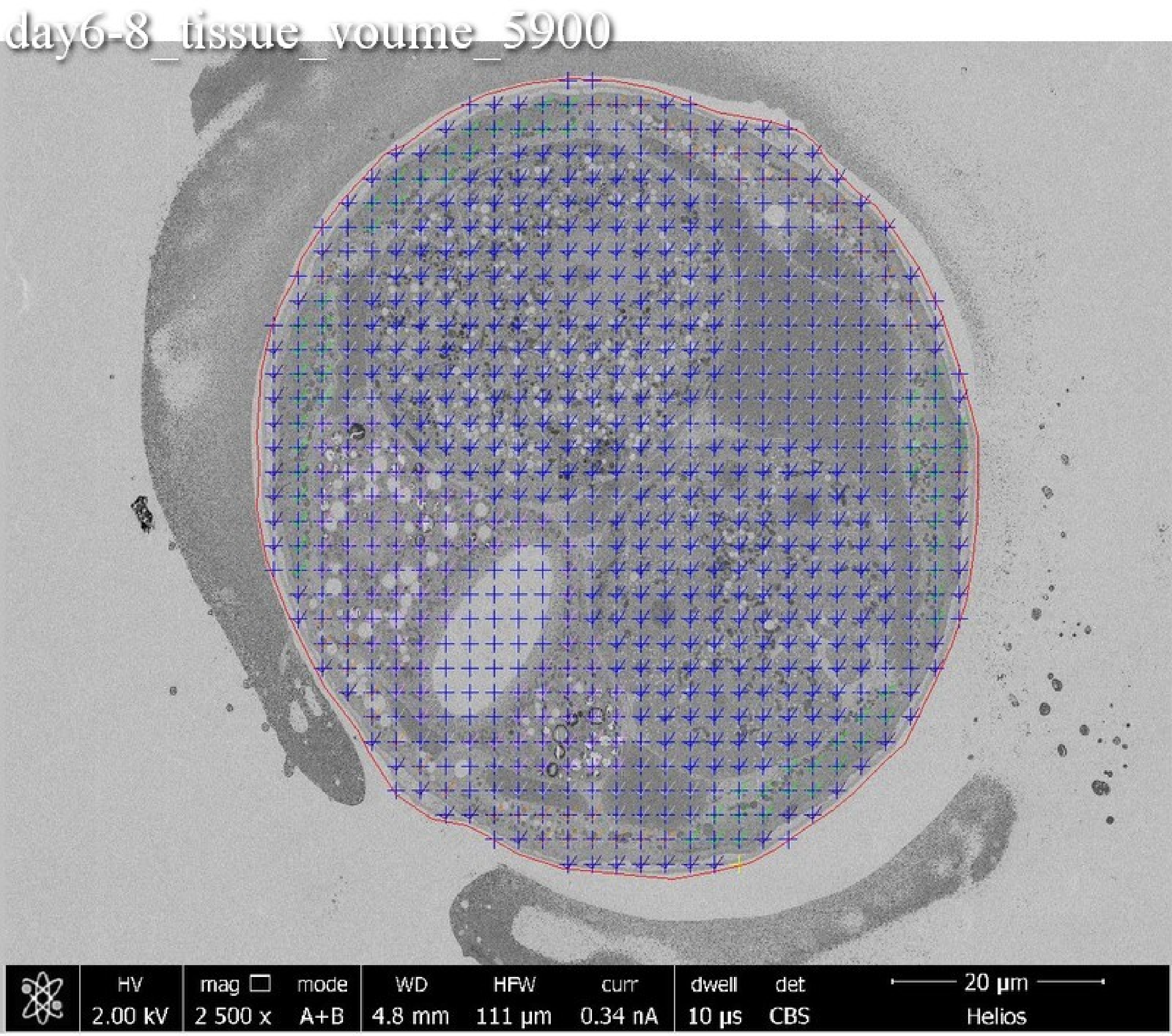
day6-8_tissue_voume_5900
HV 2.00 kV
mag 2 500 x
mode A+B
WD 4.8 mm
HFW 111 μm
curr 0.34 nA
dwell 10 μs
det CBS
20 μm
Helios

day6-8_tissue_voume_7050

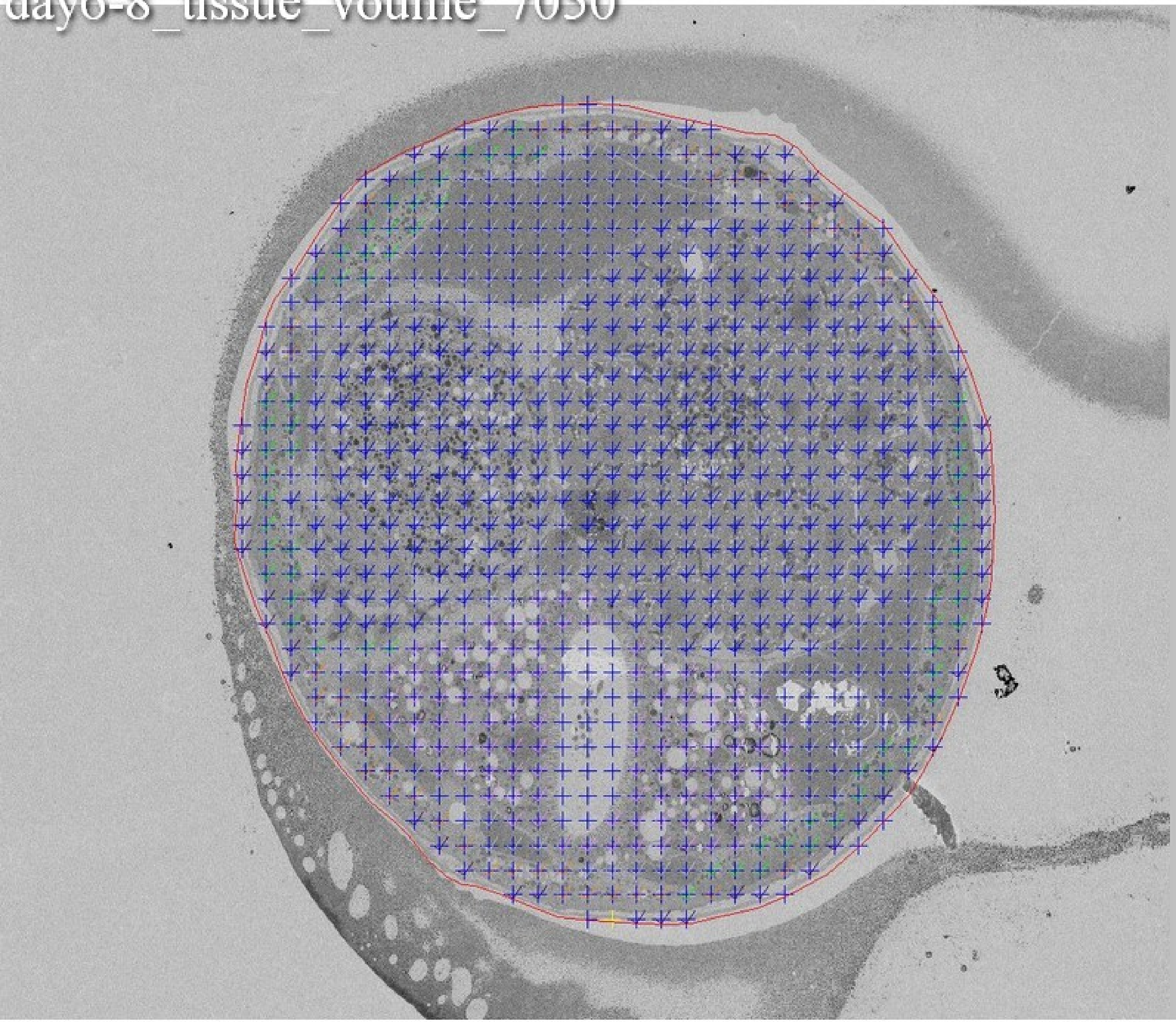

day6-8_tissue_voume_8200

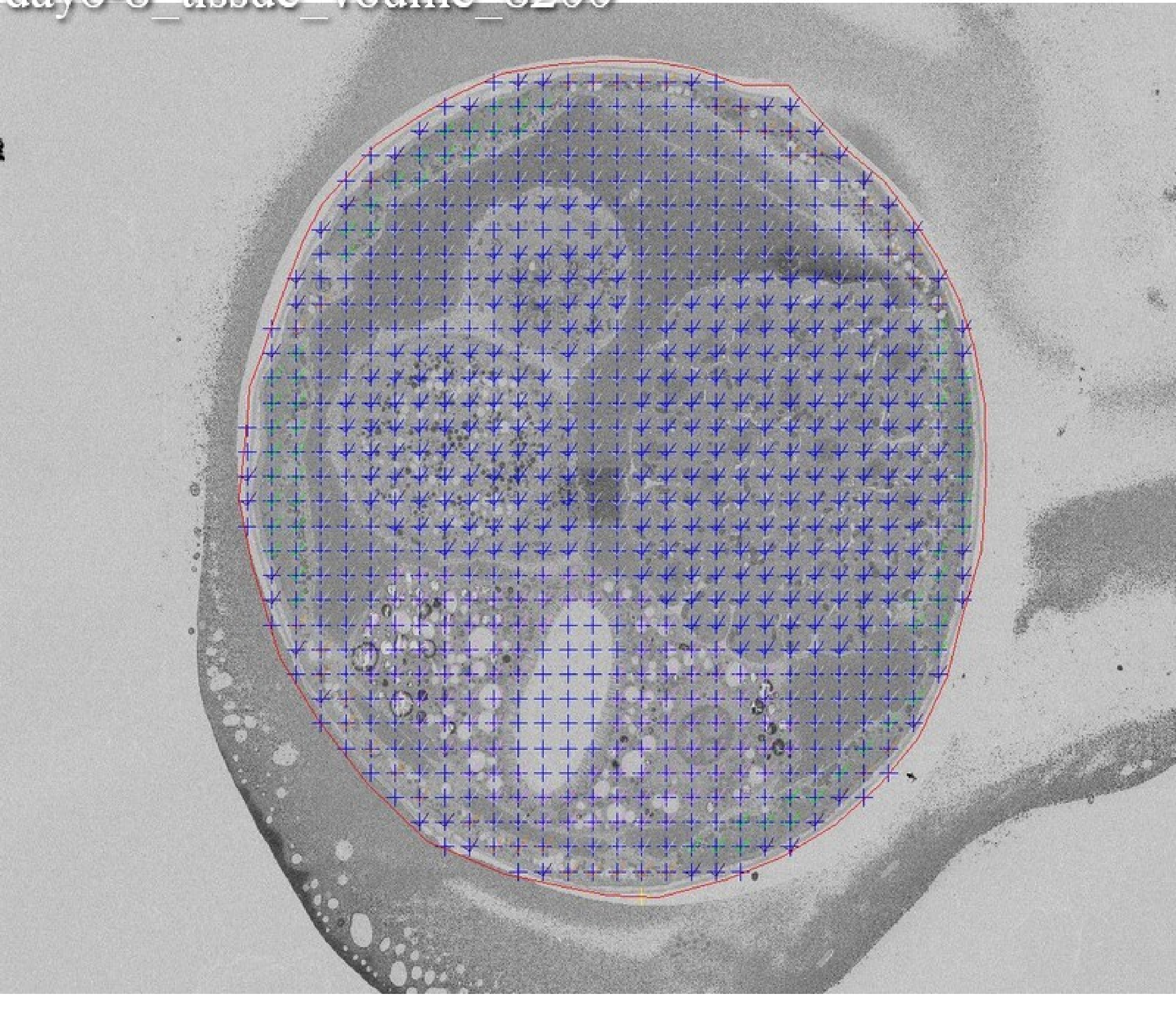

day6-8_tissue_voume_9350

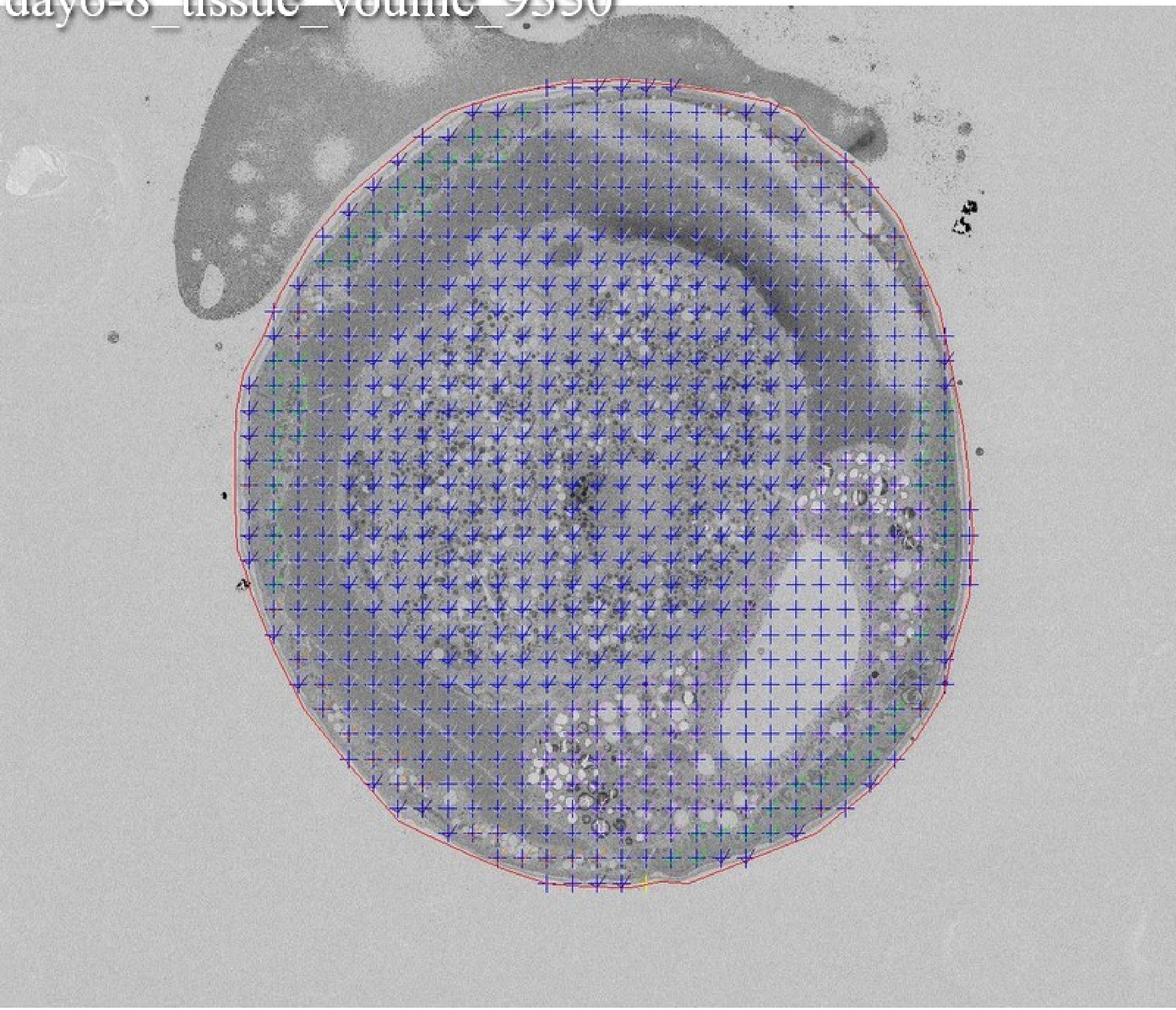

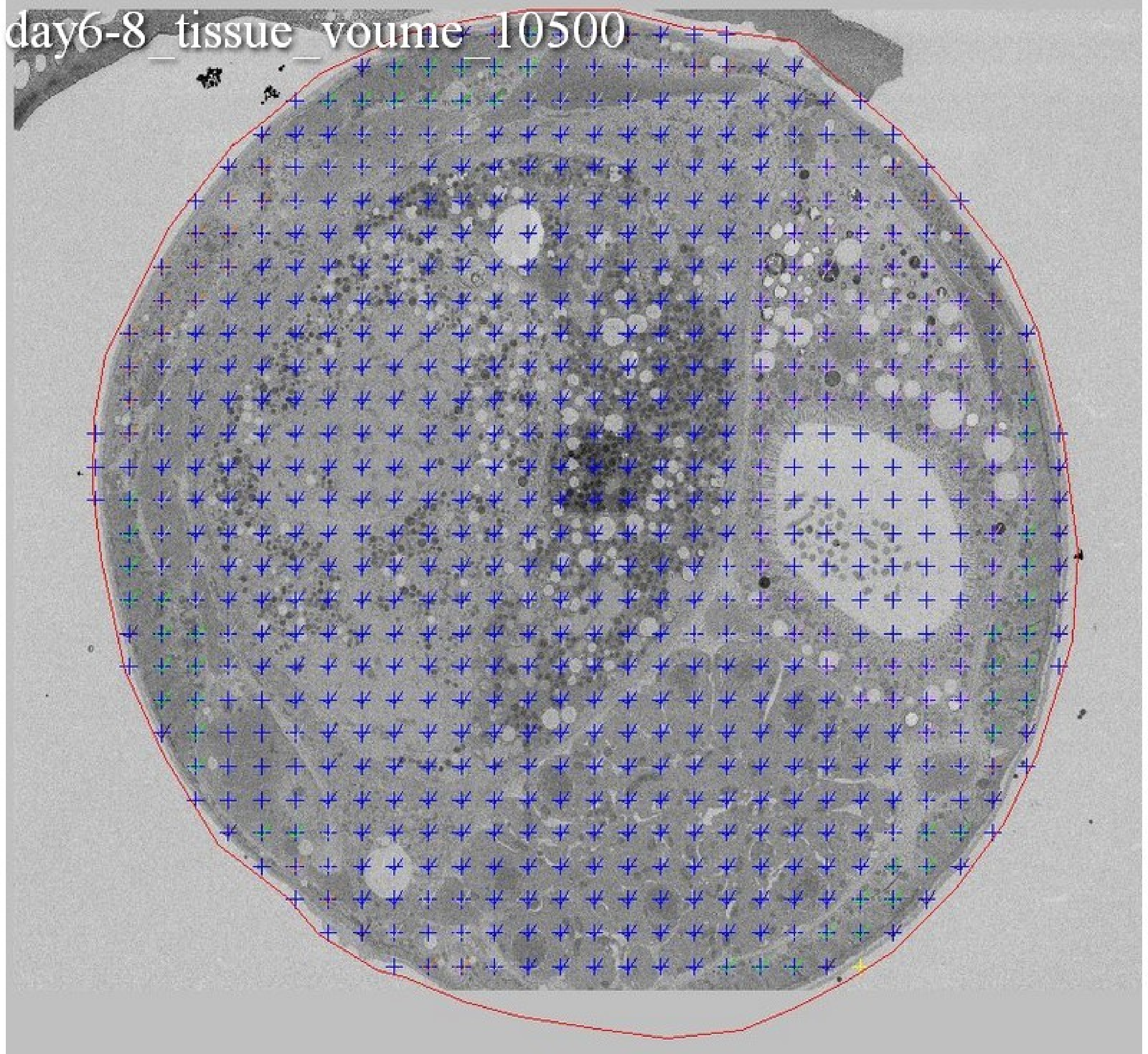
day6-8_tissue_voume_10500

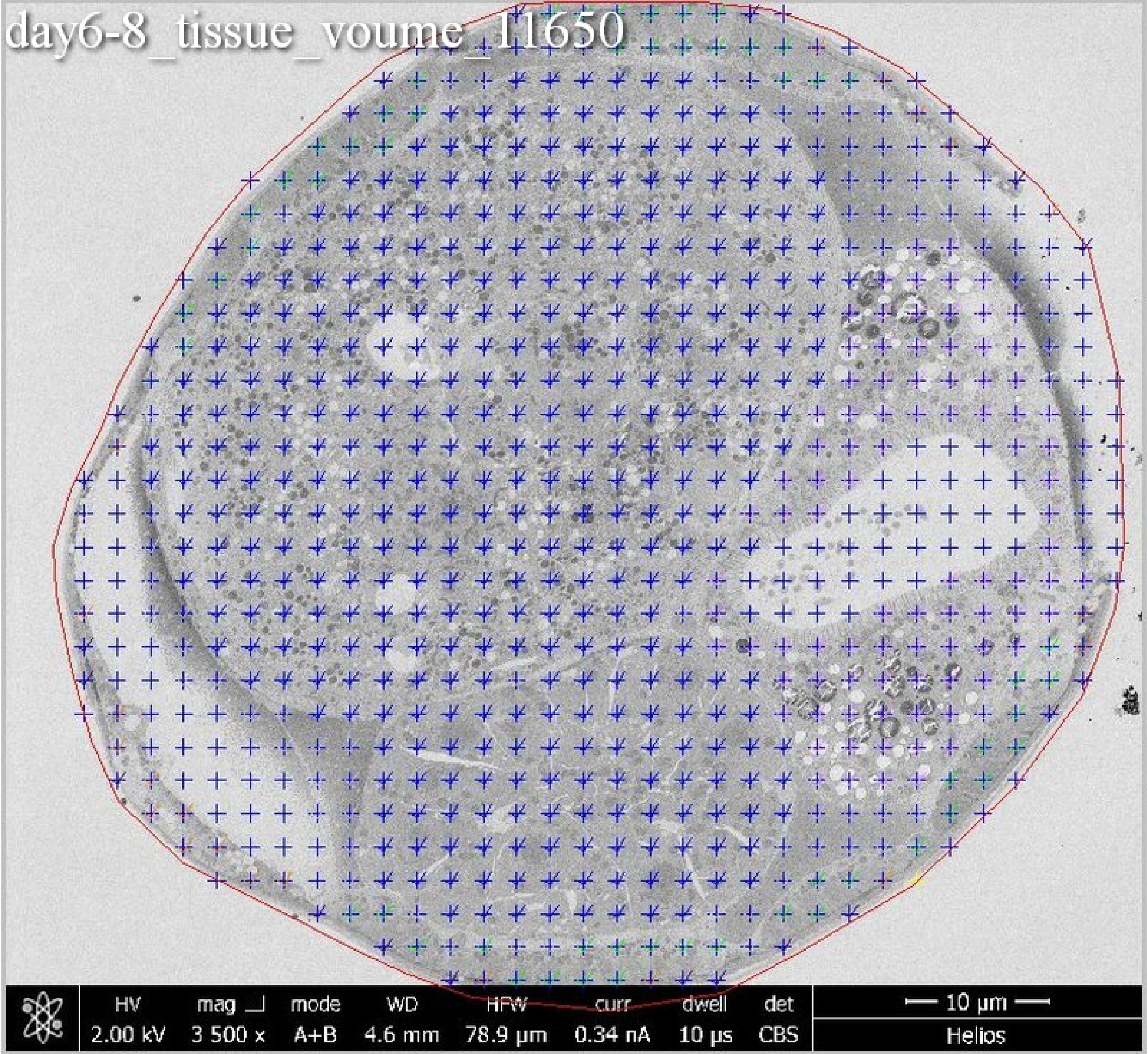
day6-8_tissue_voume_11650
HV 2.00 kV
mag 3 500 x
mode A+B
WD 4.6 mm
HFW 78.9 µm
curr 0.34 nA
dwell 10 µs
det CBS
10 µm
Helios

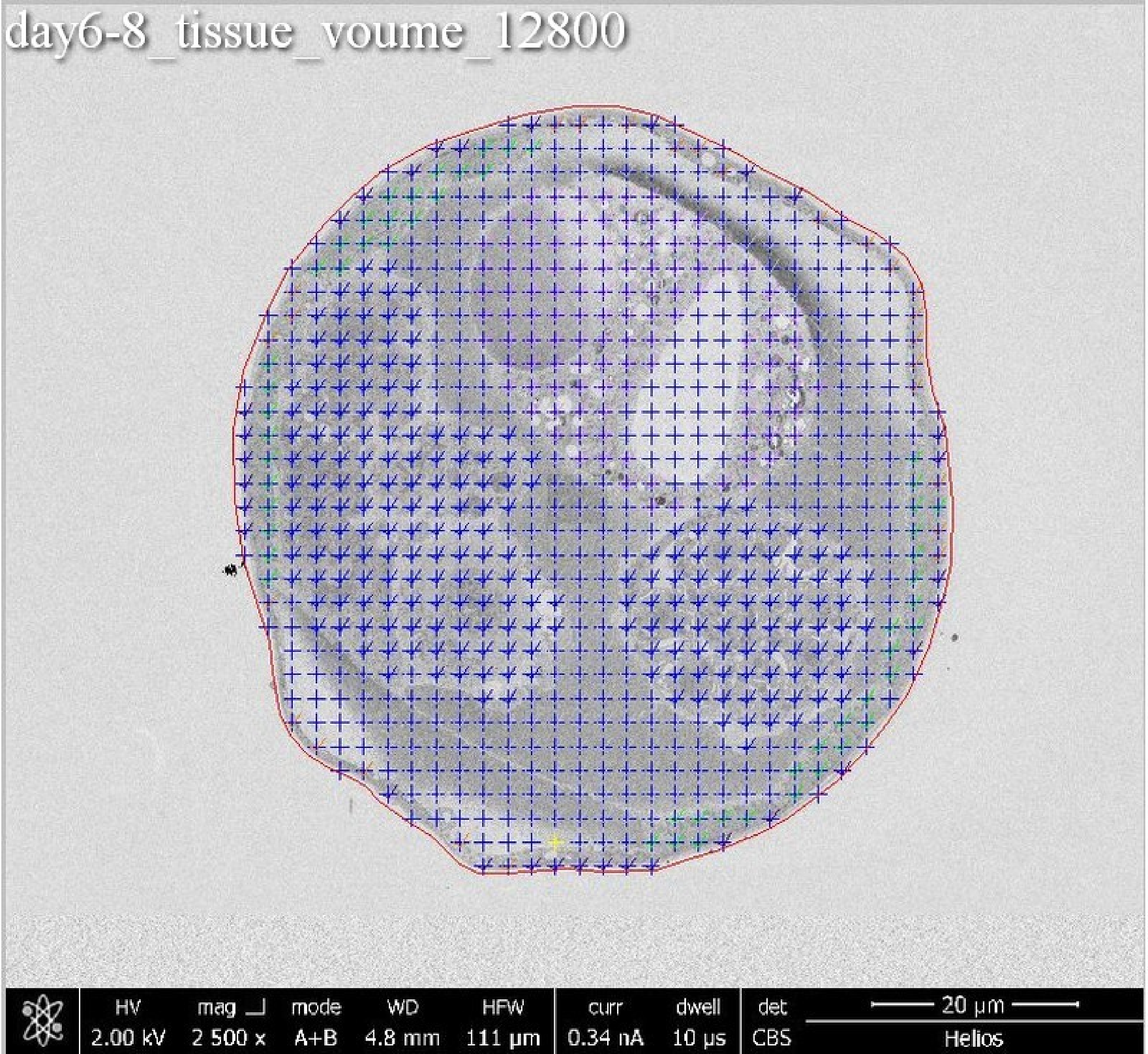
day6-8_tissue_voume_12800
HV 2.00 kV
mag 2 500 x
mode A+B
WD 4.8 mm
HFW 111 μm
curr 0.34 nA
dwell 10 μs
det CBS
20 μm
Helios

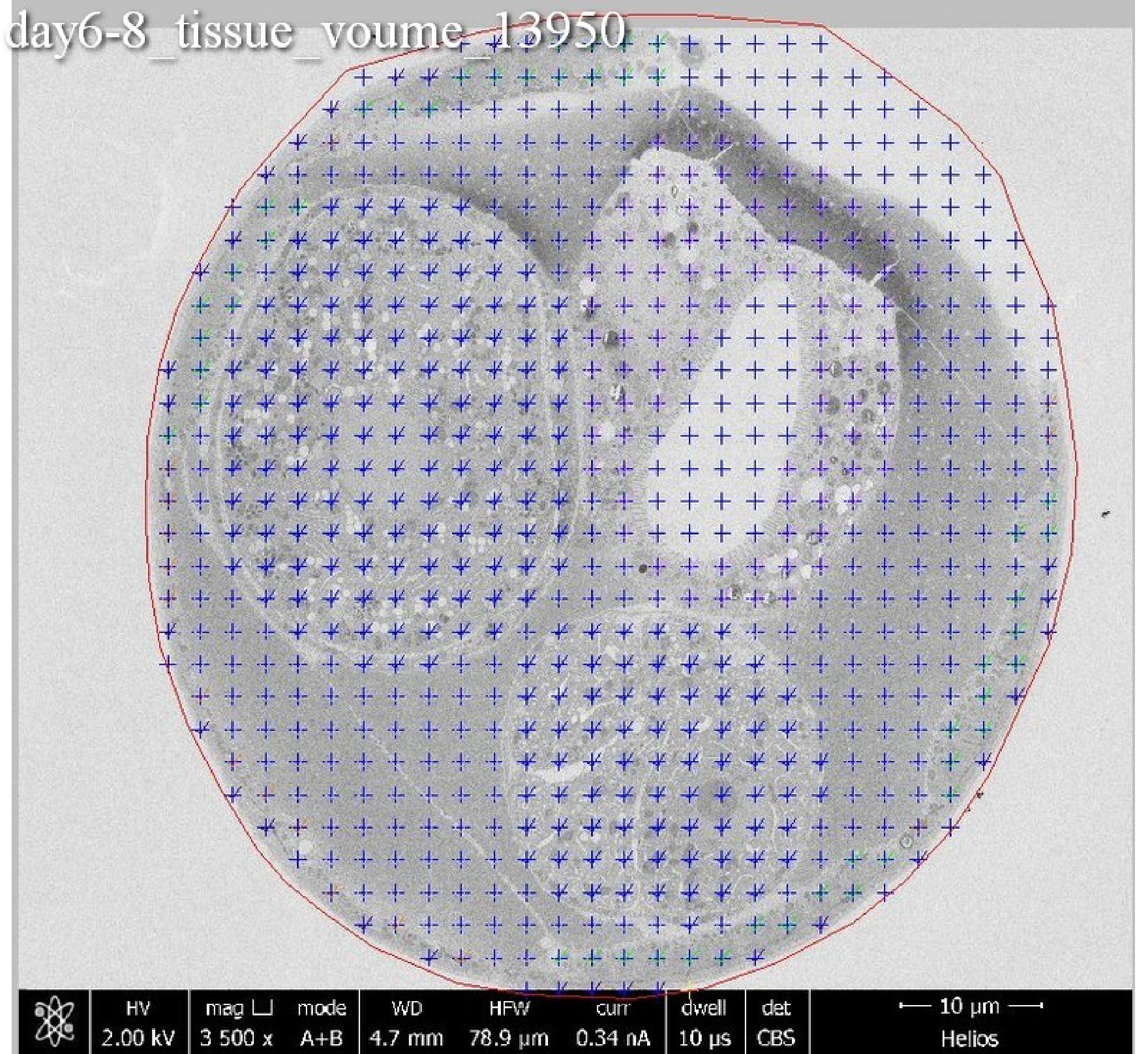
day6-8_tissue_voume_13950
HV
2.00 kV
mag ⊔
3 500 x
mode
A+B
WD
4.7 mm
HFW
78.9 µm
curr
0.34 nA
dwell
10 µs
det
CBS
10 µm
Helios

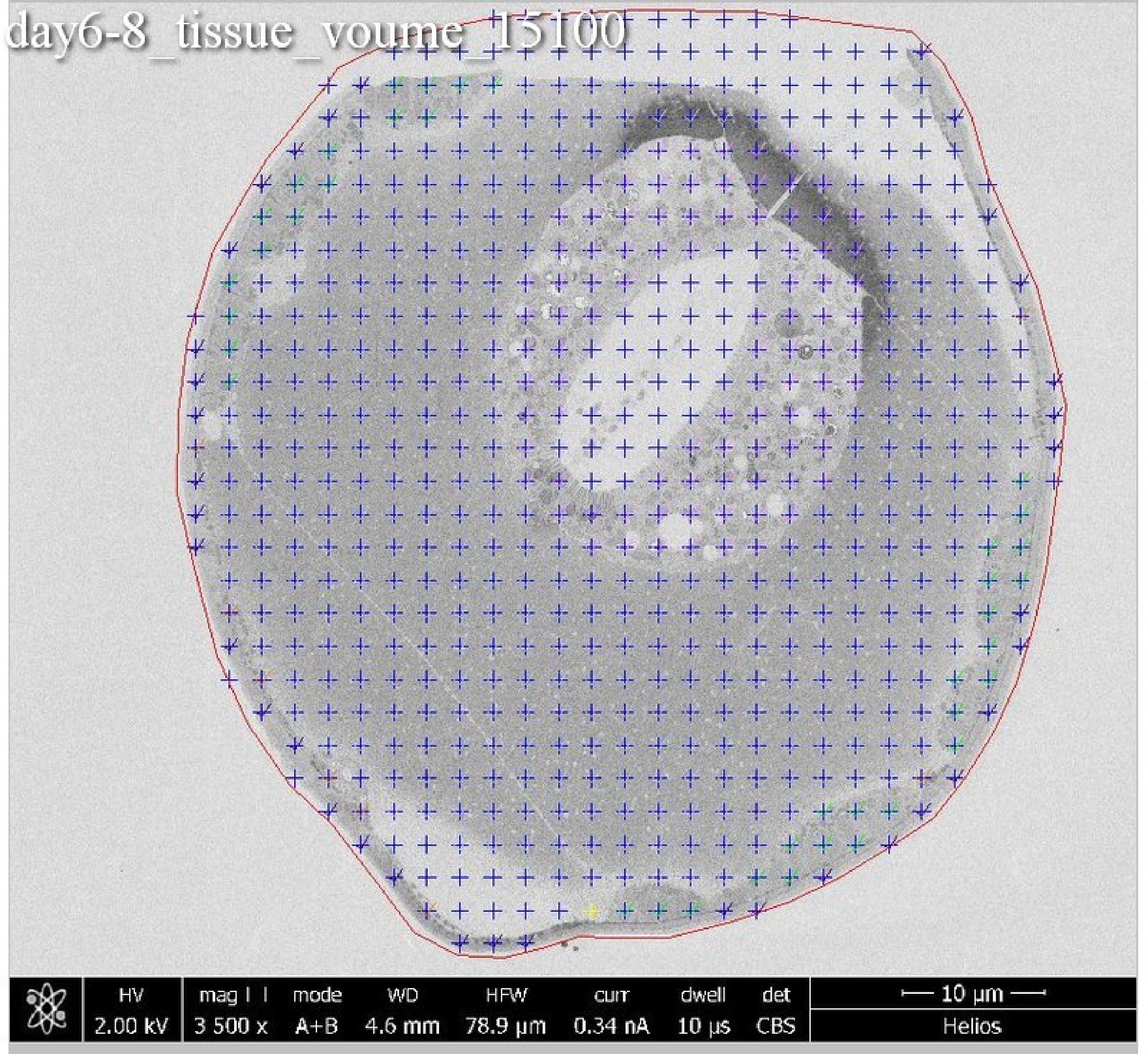
day6-8_tissue_voume_15100
HV 2.00 kV
mag 3 500 x
mode A+B
WD 4.6 mm
HFW 78.9 µm
curr 0.34 nA
dwell 10 µs
det CBS
10 µm
Helios

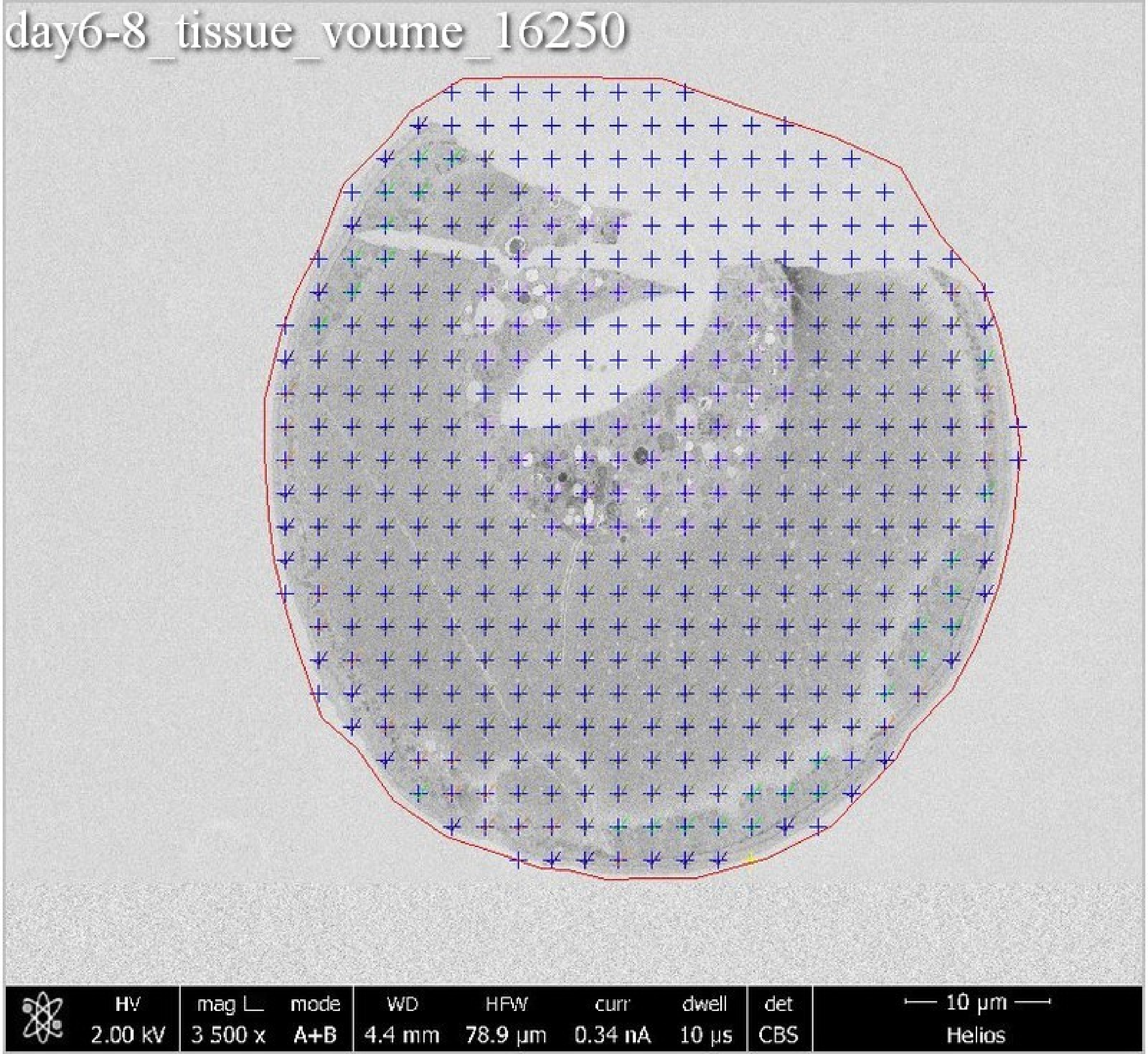
day6-8_tissue_voume_16250
HV
2.00 kV
mag ⊔
3 500 x
mode
A+B
WD
4.4 mm
HFW
78.9 μm
curr
0.34 nA
dwell
10 μs
det
CBS
10 μm
Helios

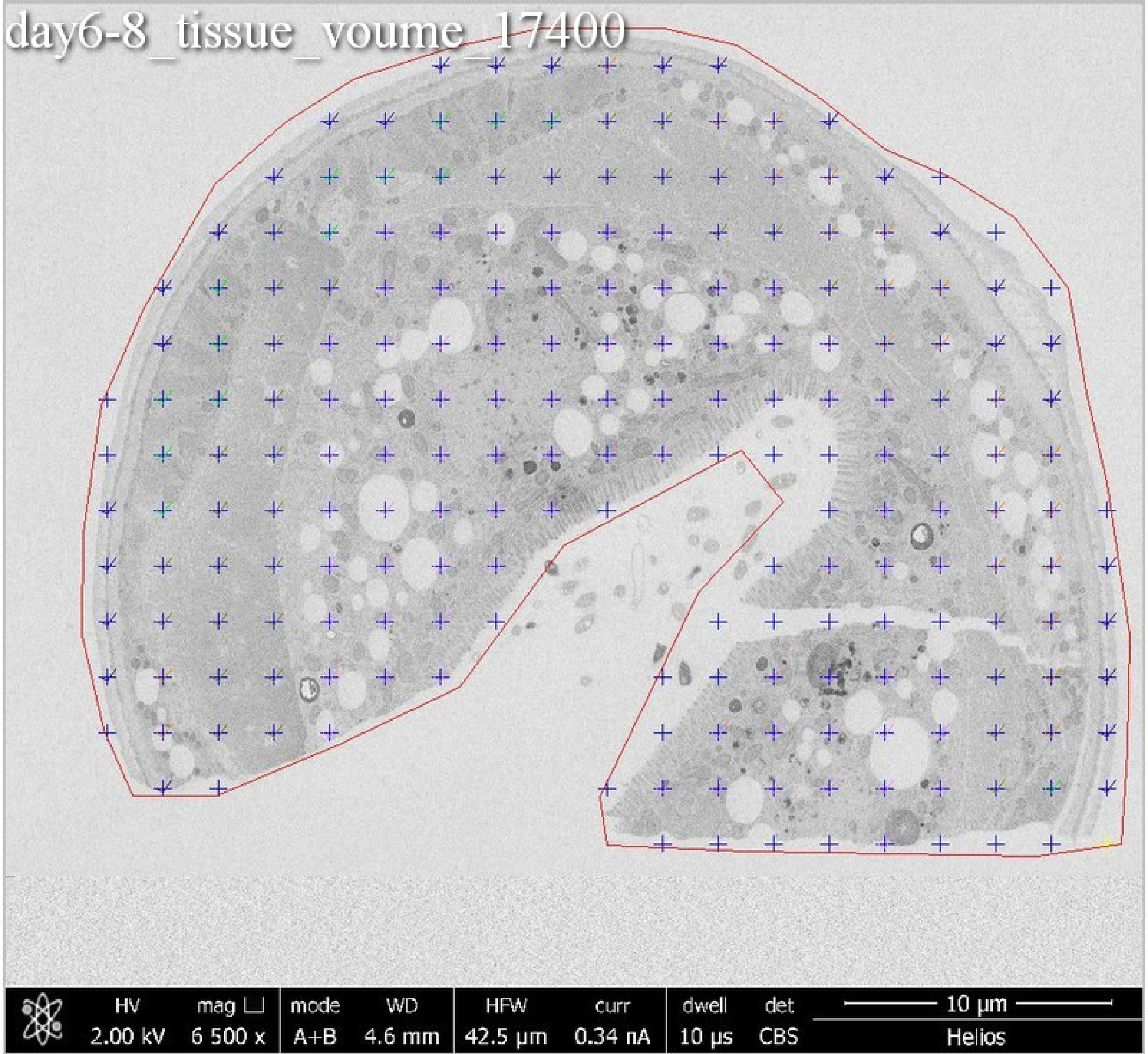
day6-8_tissue_voume_17400
HV
2.00 kV
mag ⊔
6 500 x
mode
A+B
WD
4.6 mm
HFW
42.5 μm
curr
0.34 nA
dwell
10 μs
det
CBS
10 μm
Helios

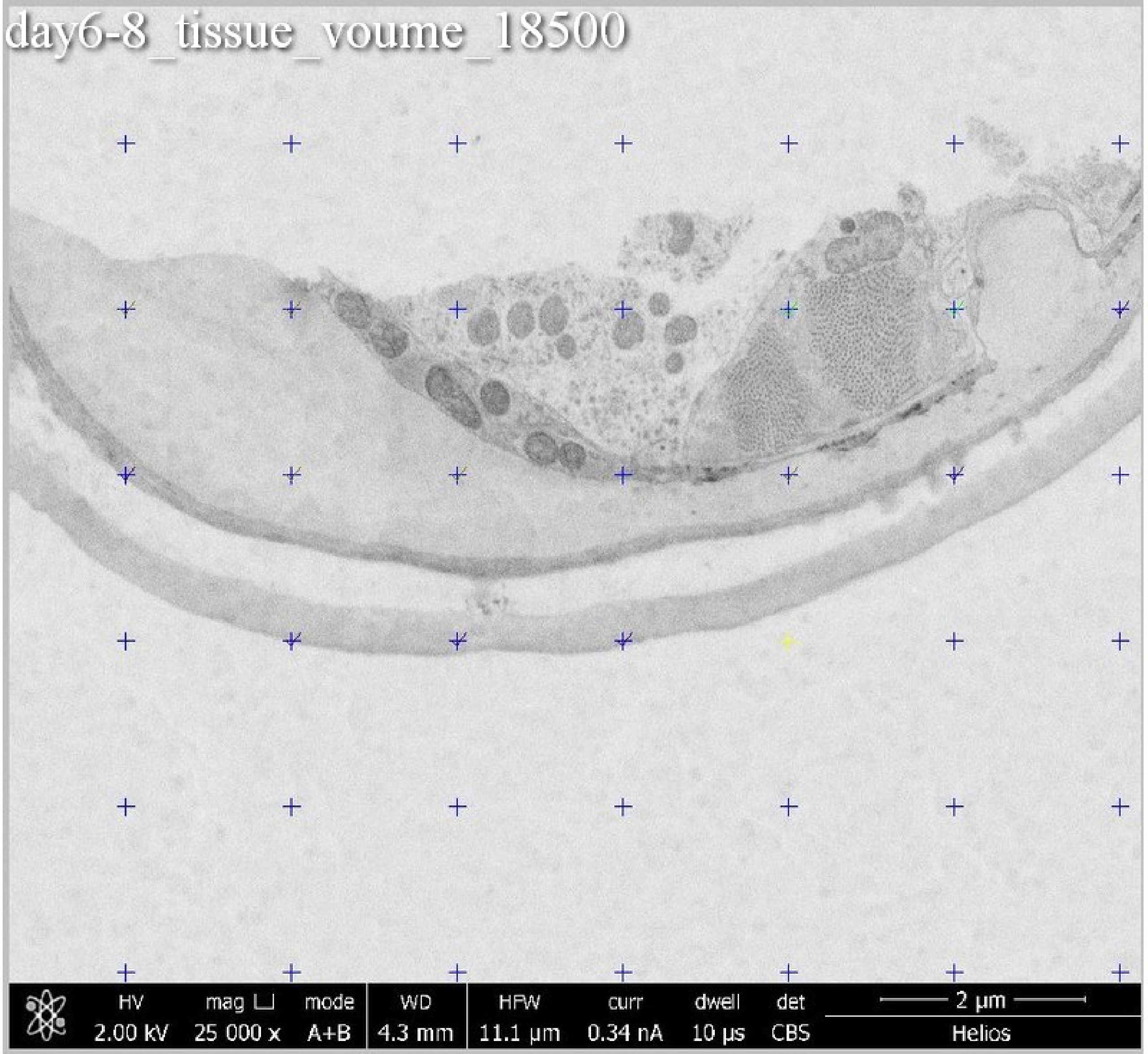
day6-8_tissue_voume_18500
HV 2.00 kV
mag ⊔ 25 000 x
mode A+B
WD 4.3 mm
HFW 11.1 µm
curr 0.34 nA
dwell 10 µs
det CBS
2 µm
Helios

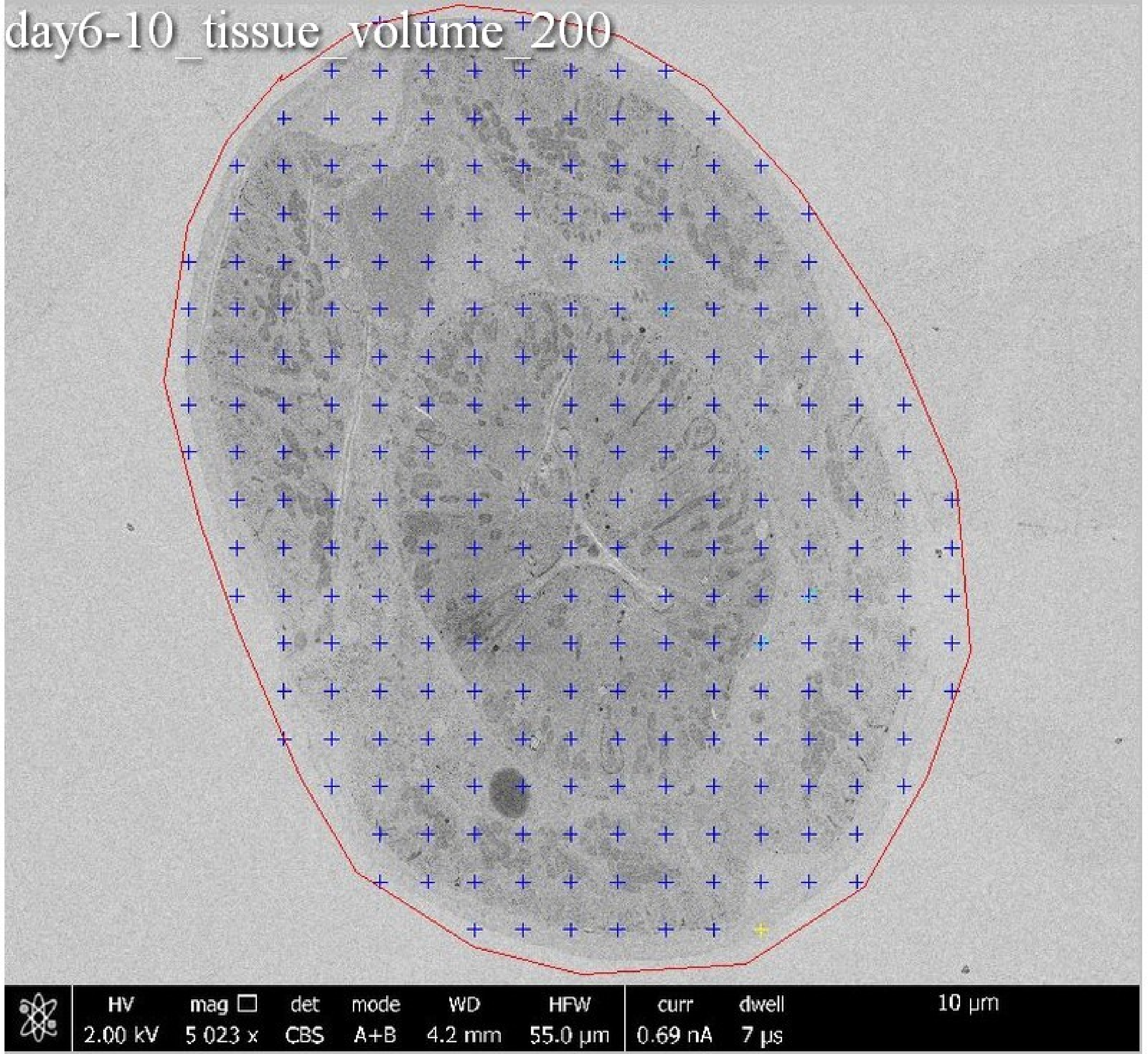
day6-10_tissue_volume_200
HV
2.00 kV
mag □
5 023 x
det
CBS
mode
A+B
WD
4.2 mm
HFW
55.0 μm
curr
0.69 nA
dwell
7 μs
10 μm

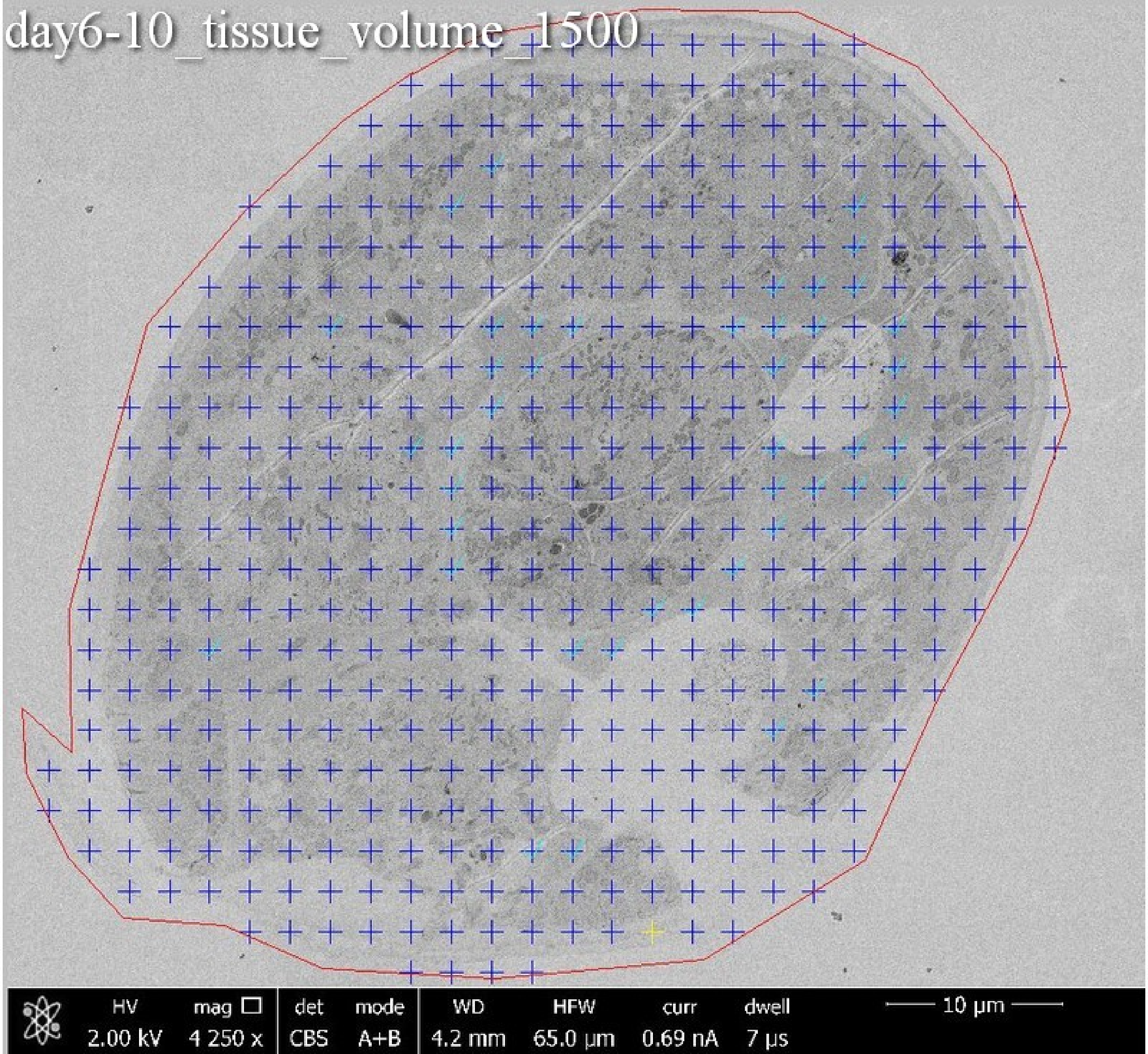
day6-10_tissue_volume_1500
HV
2.00 kV
mag
4 250 x
det
CBS
mode
A+B
WD
4.2 mm
HFW
65.0 µm
curr
0.69 nA
dwell
7 µs
10 µm

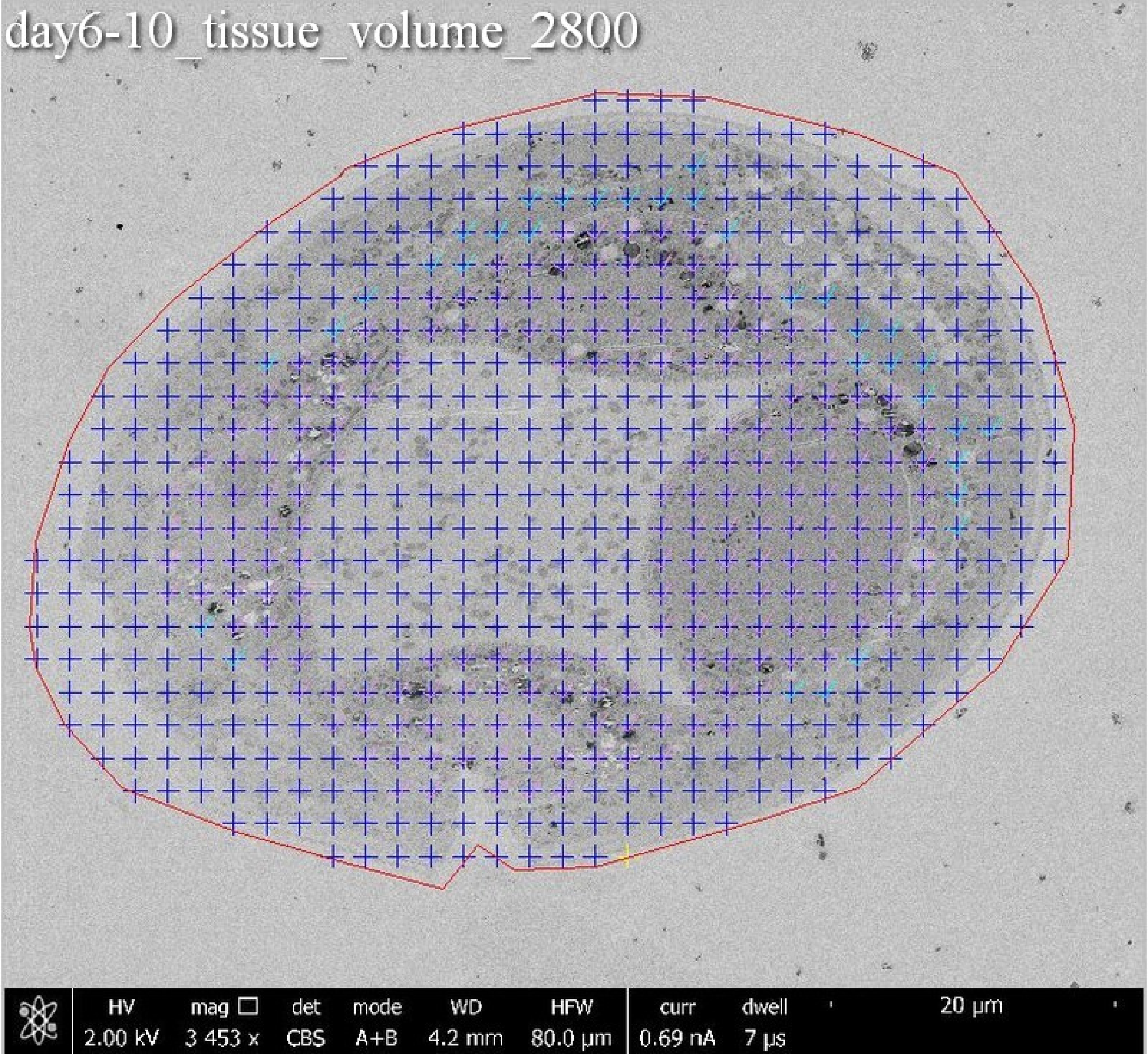
day6-10_tissue_volume_2800
HV
2.00 kV
mag
3 453 x
det
CBS
mode
A+B
WD
4.2 mm
HFW
80.0 µm
curr
0.69 nA
dwell
7 µs
20 µm

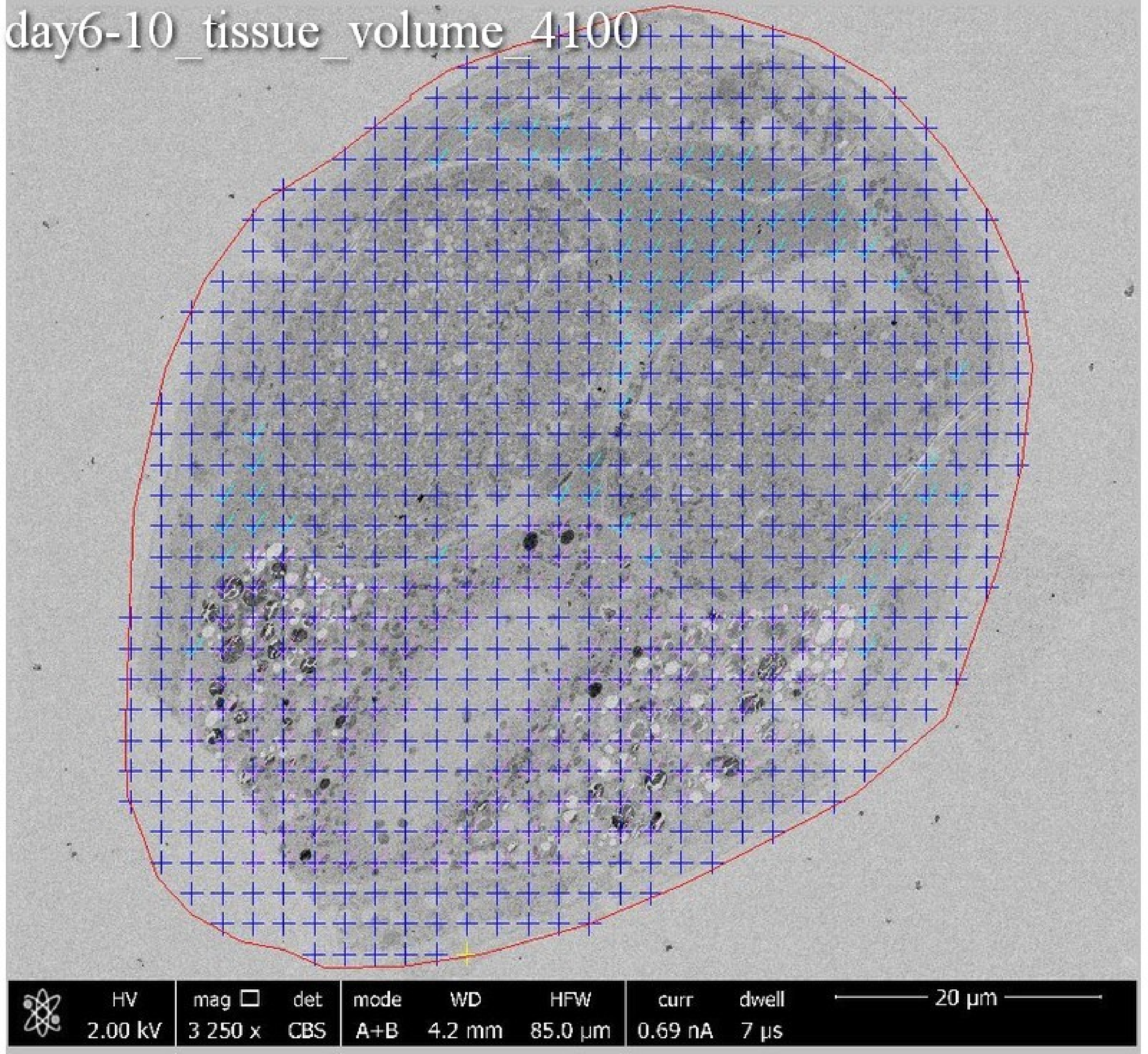
day6-10_tissue_volume_4100
HV
2.00 kV
mag ☐
3 250 x
det
CBS
mode
A+B
WD
4.2 mm
HFW
85.0 µm
curr
0.69 nA
dwell
7 µs
20 µm

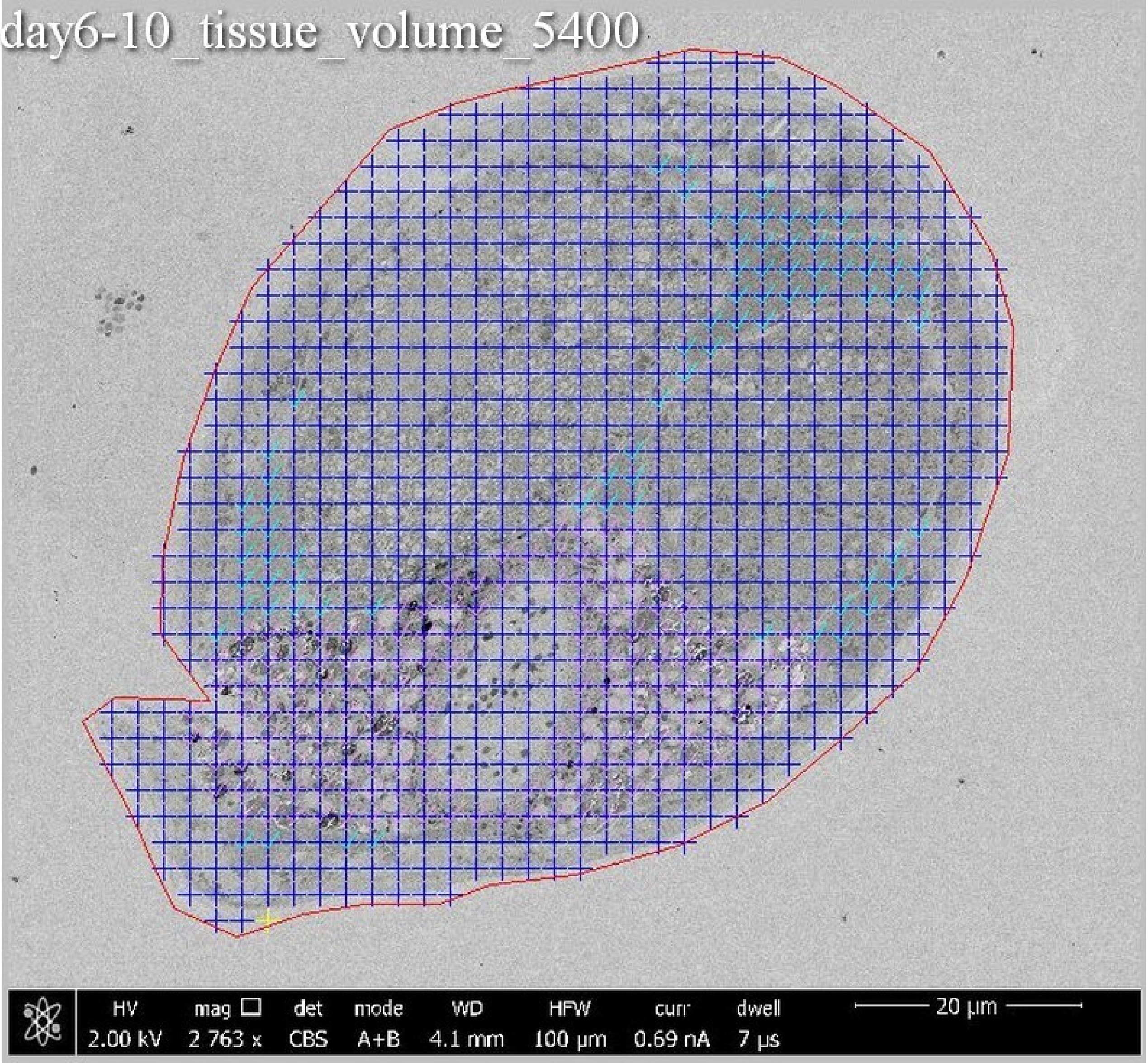
day6-10_tissue_volume_5400
HV 2.00 kV
mag 2 763 x
det CBS
mode A+B
WD 4.1 mm
HFW 100 µm
curr 0.69 nA
dwell 7 µs
20 µm

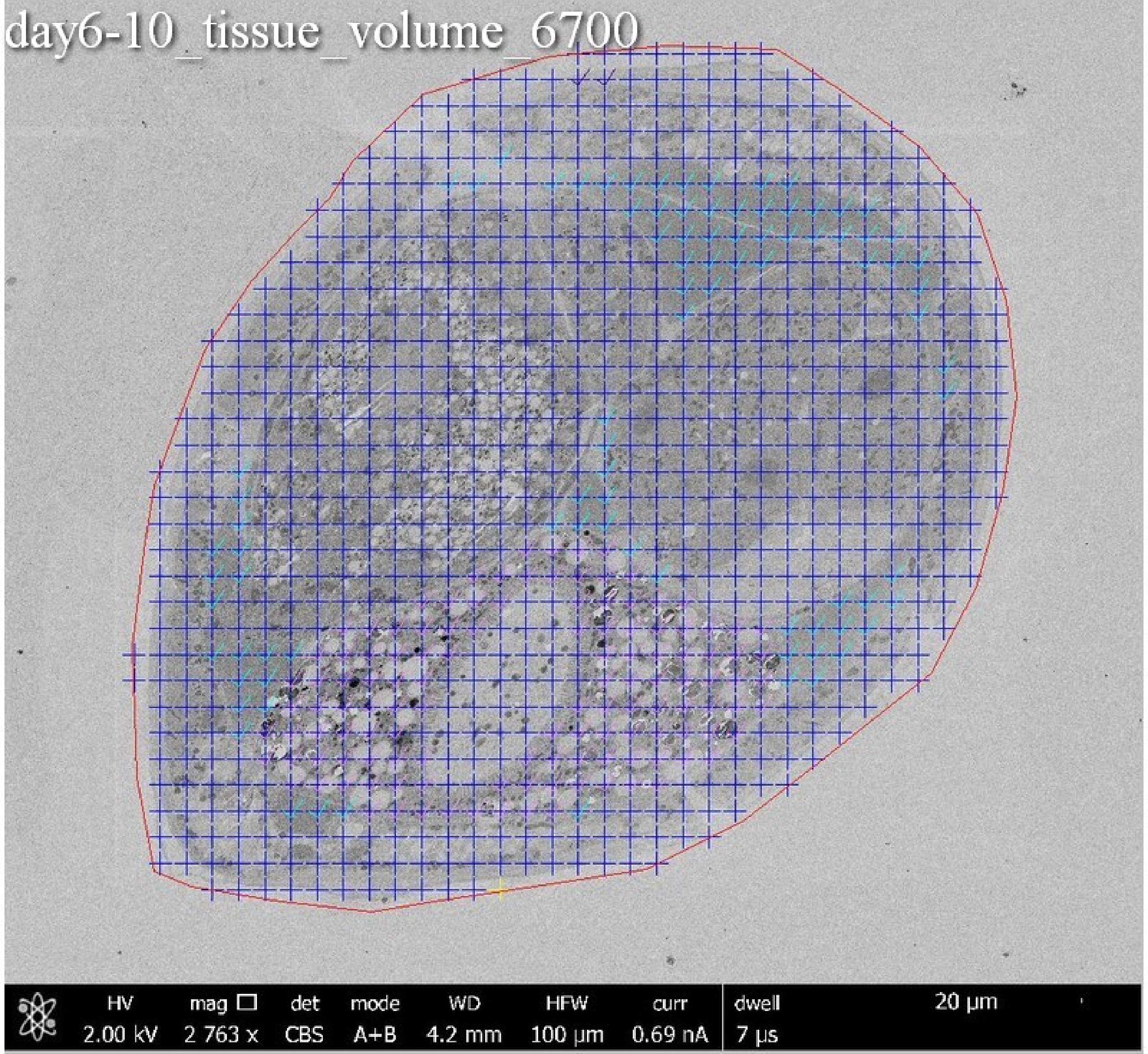
day6-10_tissue_volume_6700
HV 2.00 kV
mag ☐ 2 763 x
det CBS
mode A+B
WD 4.2 mm
HFW 100 µm
curr 0.69 nA
dwell 7 µs
20 µm

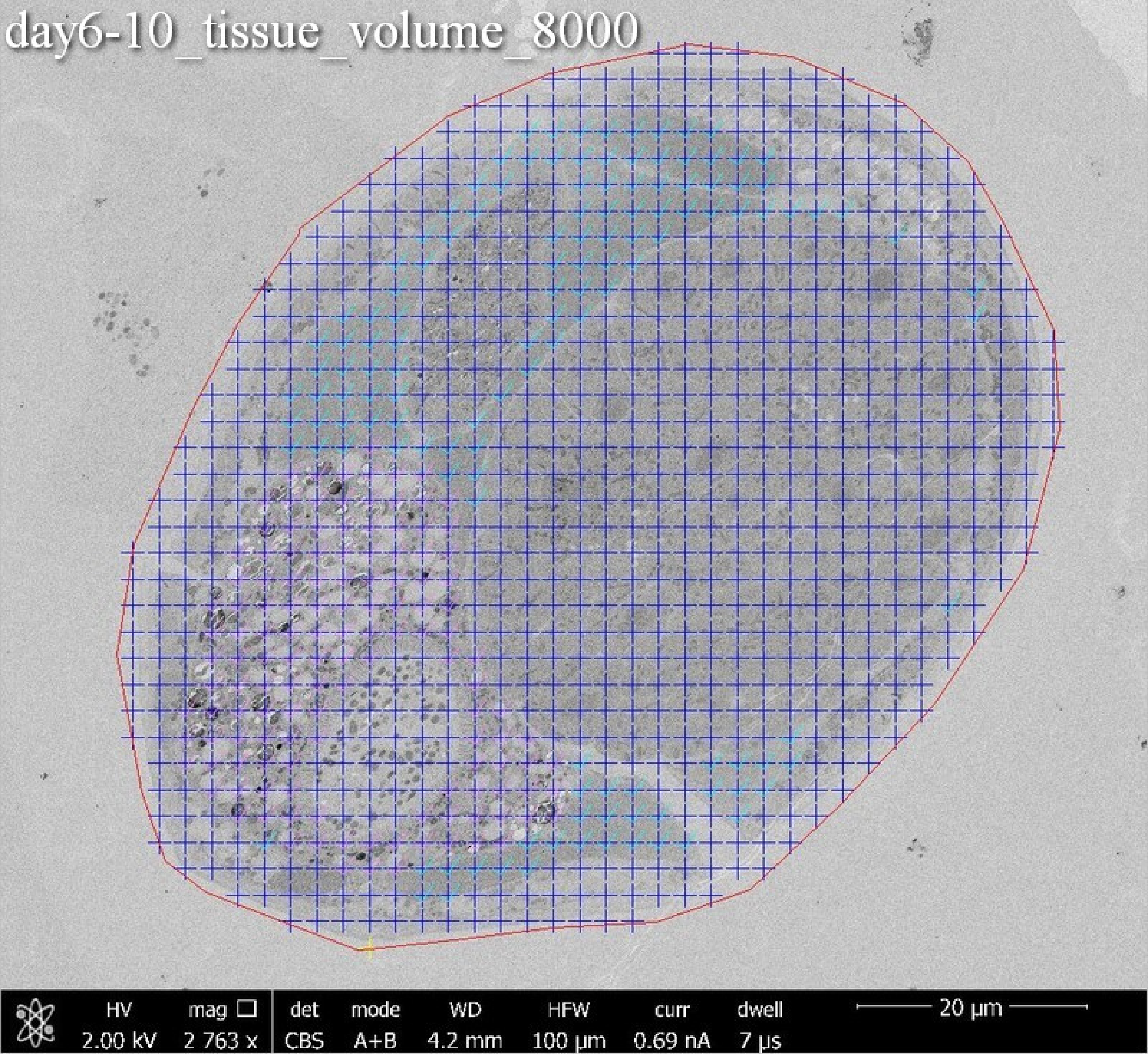
day6-10_tissue_volume_8000
HV
2.00 kV
mag
2 763 x
det
CBS
mode
A+B
WD
4.2 mm
HFW
100 µm
curr
0.69 nA
dwell
7 µs
20 µm

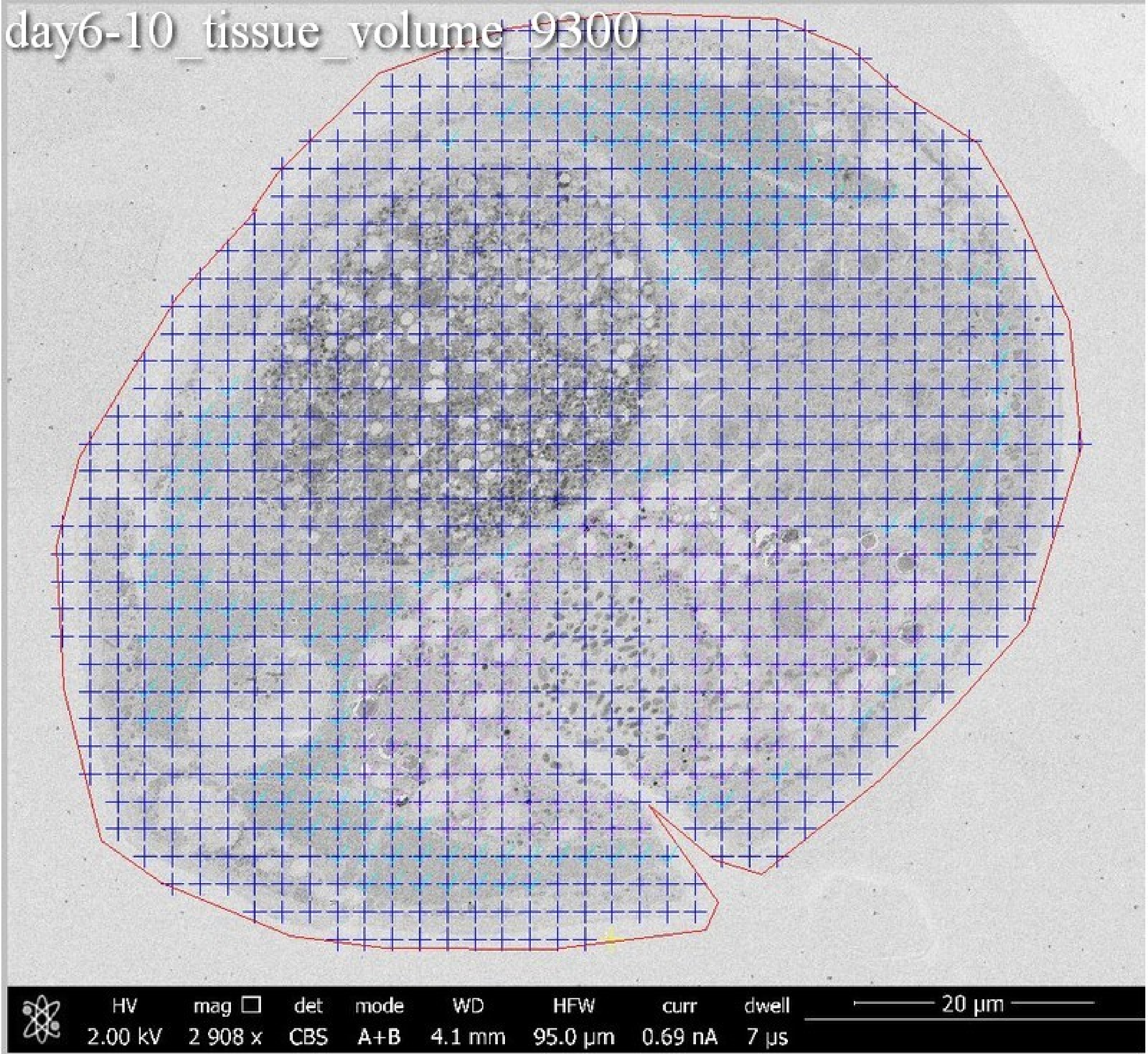
day6-10_tissue_volume_9300
HV 2.00 kV
mag 2 908 x
det CBS
mode A+B
WD 4.1 mm
HFW 95.0 µm
curr 0.69 nA
dwell 7 µs
20 µm

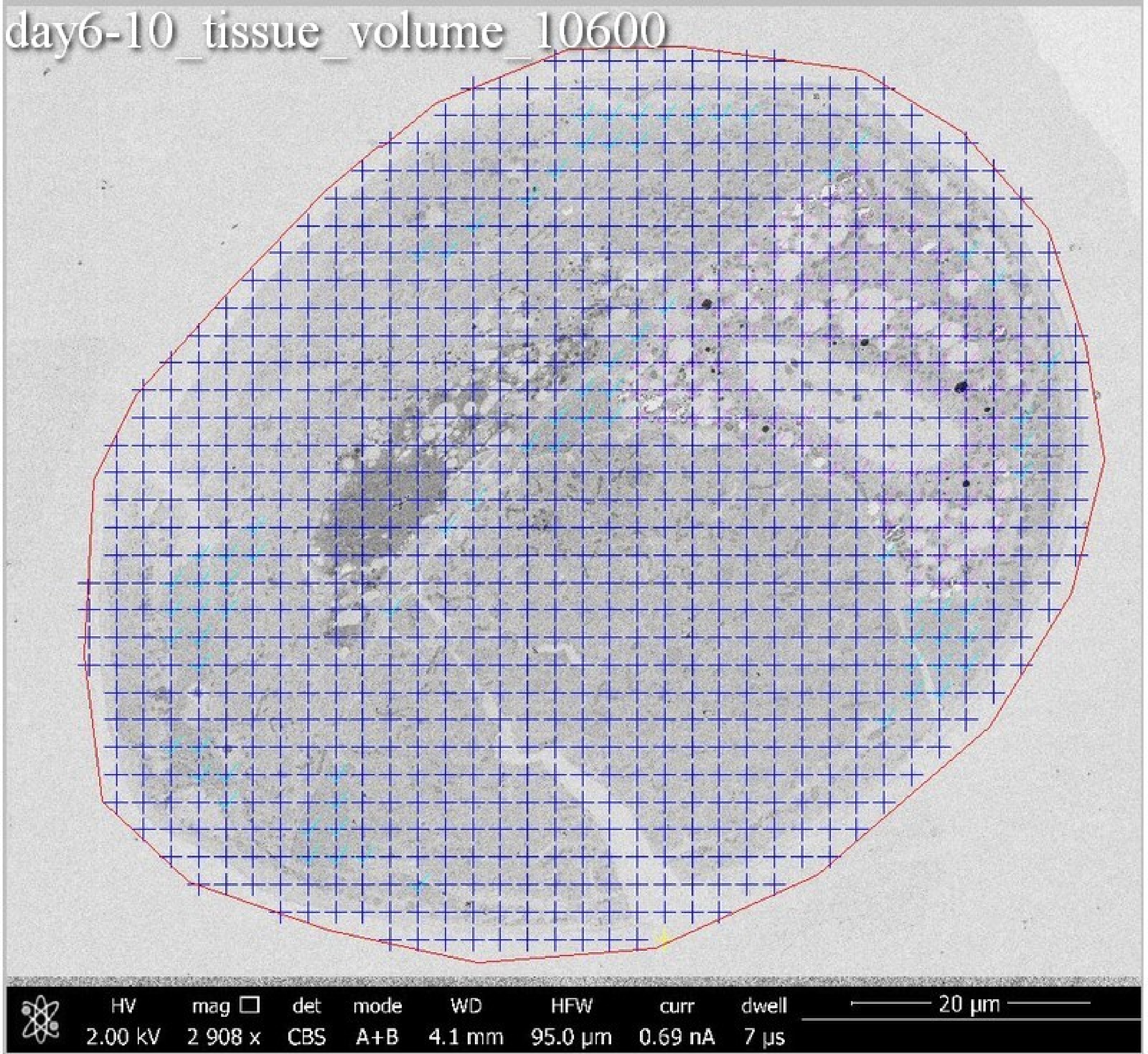
day6-10_tissue_volume_10600
HV 2.00 kV
mag 2 908 x
det CBS
mode A+B
WD 4.1 mm
HFW 95.0 µm
curr 0.69 nA
dwell 7 µs
20 µm

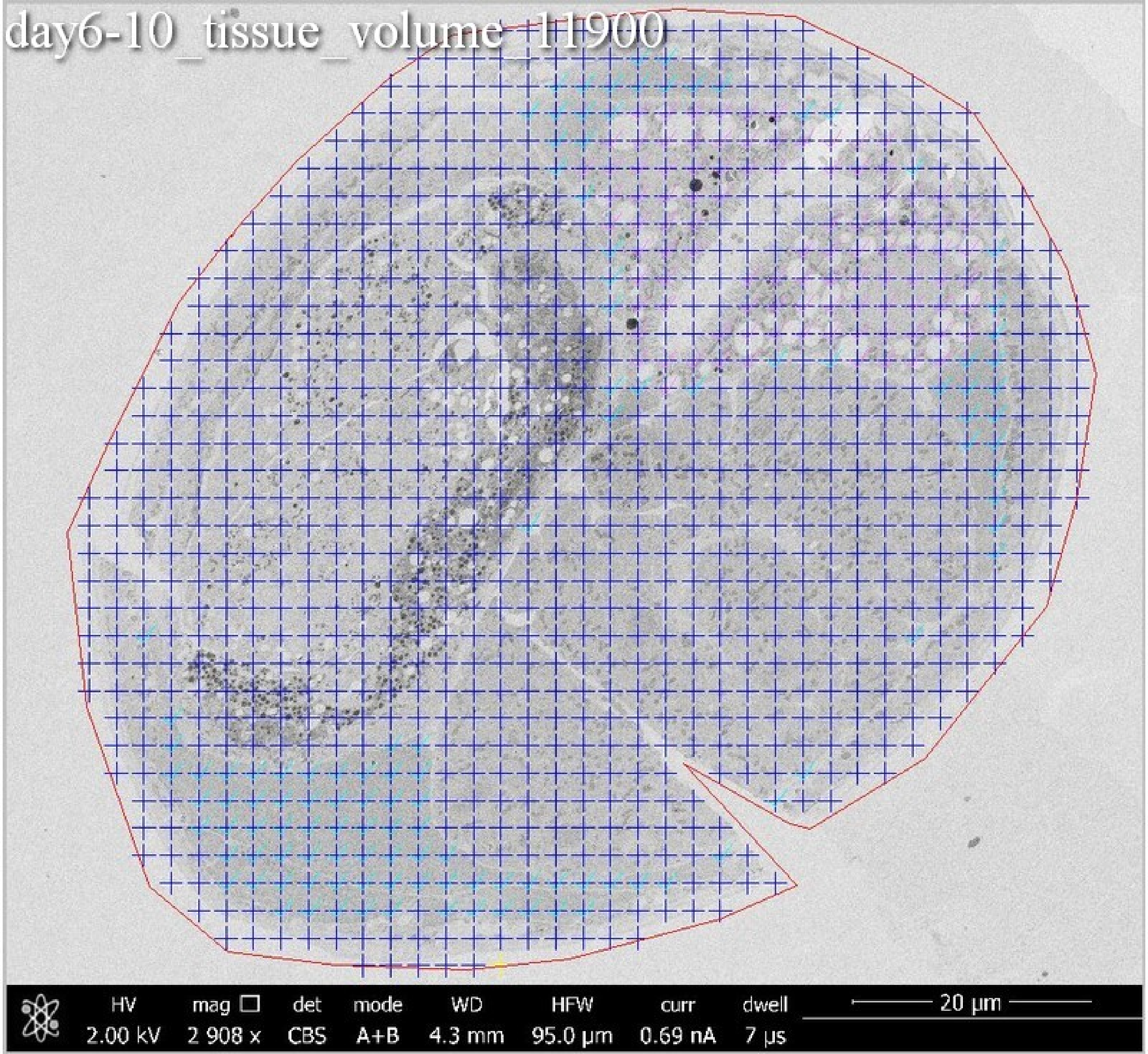
day6-10_tissue_volume_11900
HV 2.00 kV
mag 2 908 x
det CBS
mode A+B
WD 4.3 mm
HFW 95.0 µm
curr 0.69 nA
dwell 7 µs
20 µm

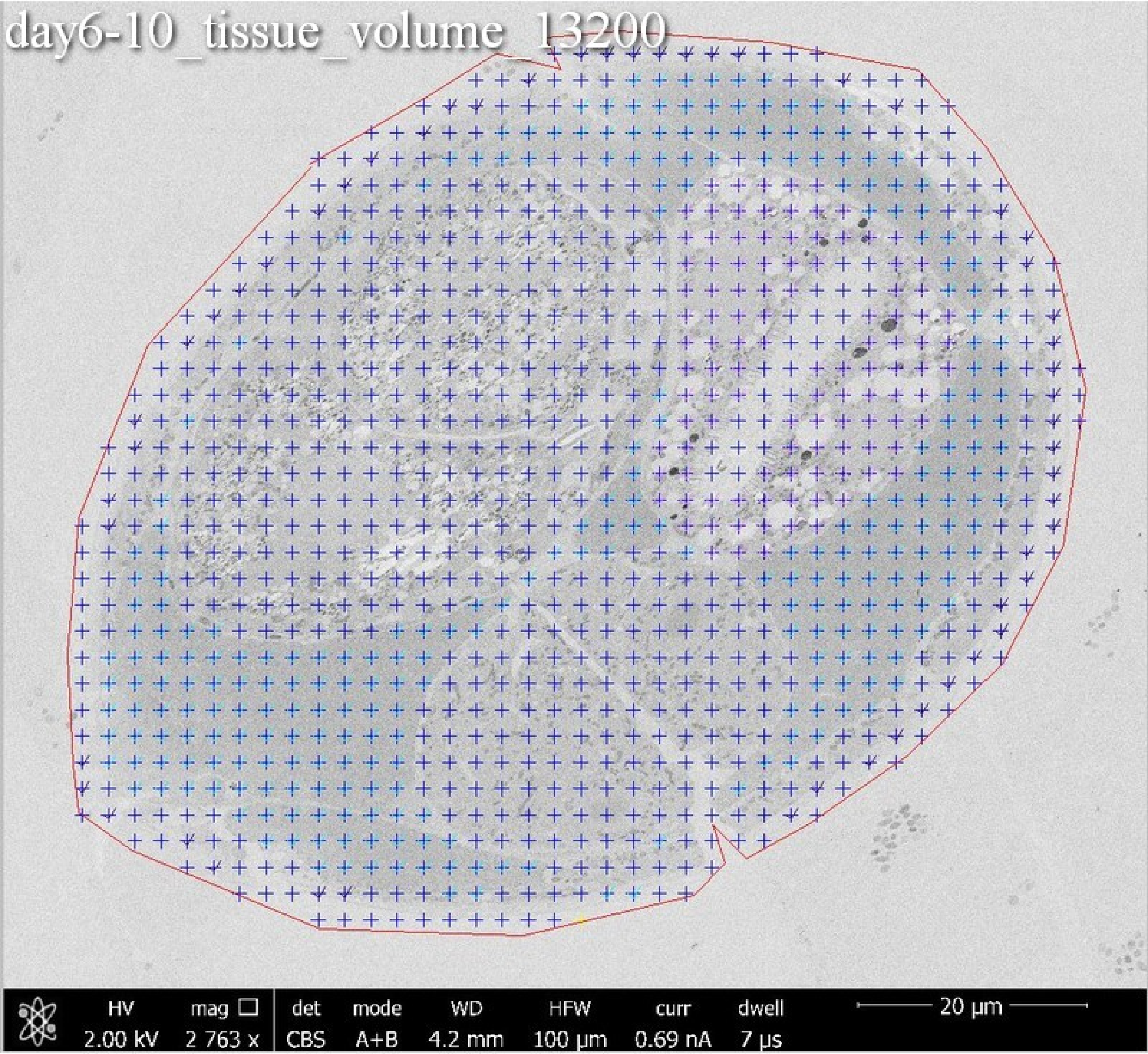
day6-10_tissue_volume_13200
HV 2.00 kV
mag 2 763 x
det CBS
mode A+B
WD 4.2 mm
HFW 100 µm
curr 0.69 nA
dwell 7 µs
20 µm

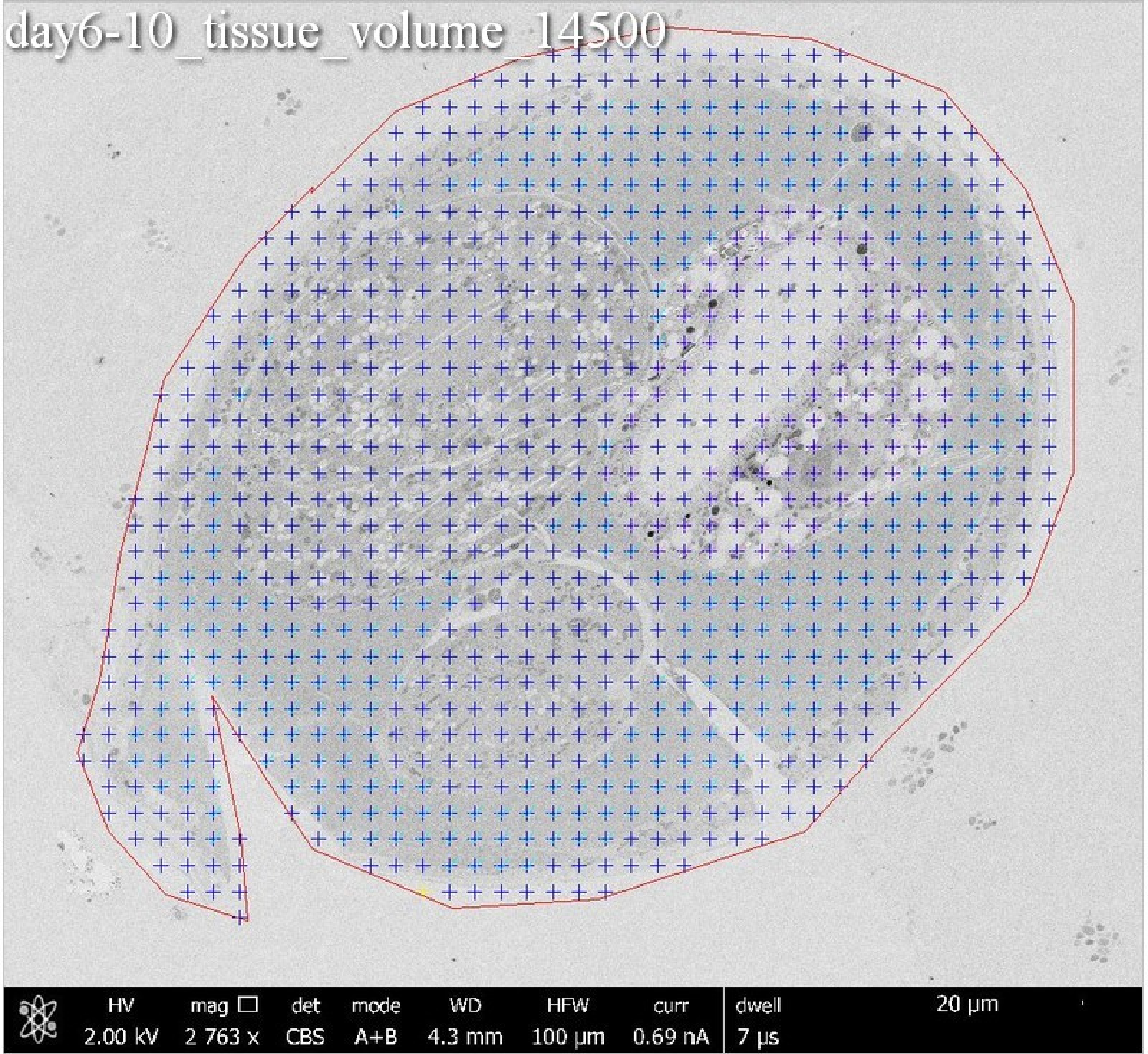
day6-10_tissue_volume_14500
HV 2.00 kV
mag 2 763 x
det CBS
mode A+B
WD 4.3 mm
HFW 100 µm
curr 0.69 nA
dwell 7 µs
20 µm

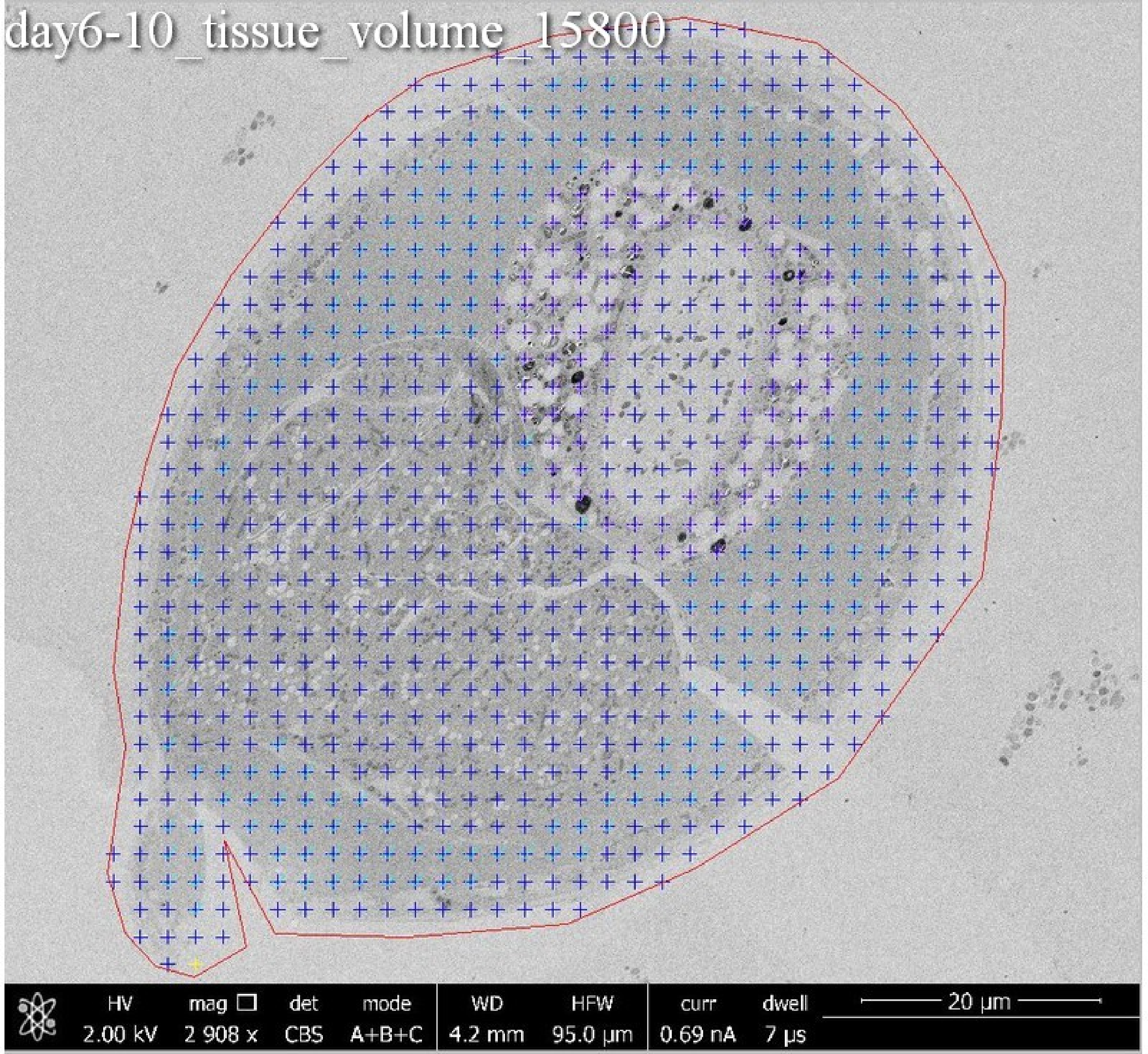
day6-10_tissue_volume_15800
HV 2.00 kV
mag 2 908 x
det CBS
mode A+B+C
WD 4.2 mm
HFW 95.0 µm
curr 0.69 nA
dwell 7 µs
20 µm

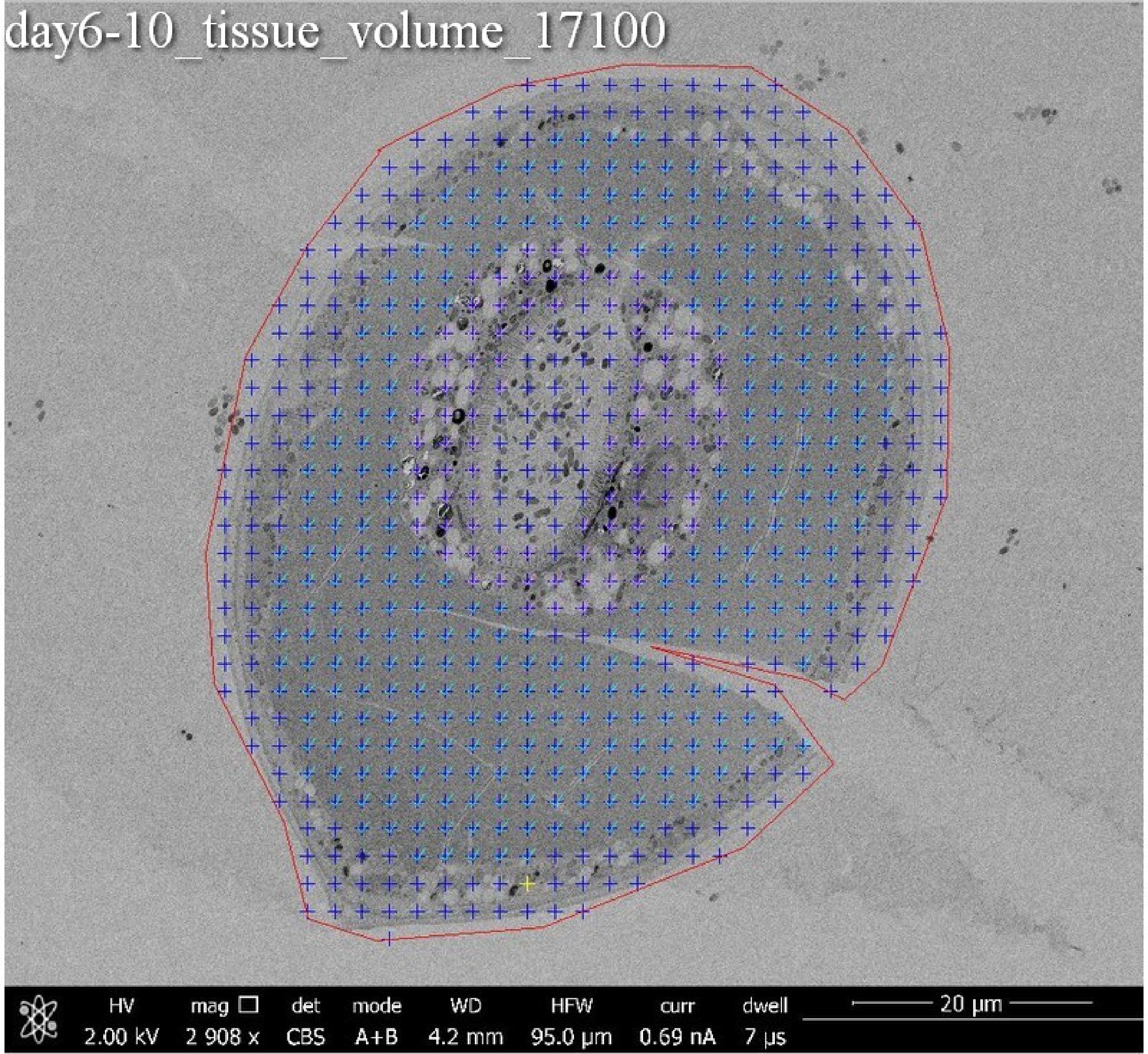
day6-10_tissue_volume_17100
HV 2.00 kV
mag 2 908 x
det CBS
mode A+B
WD 4.2 mm
HFW 95.0 μm
curr 0.69 nA
dwell 7 μs
20 μm

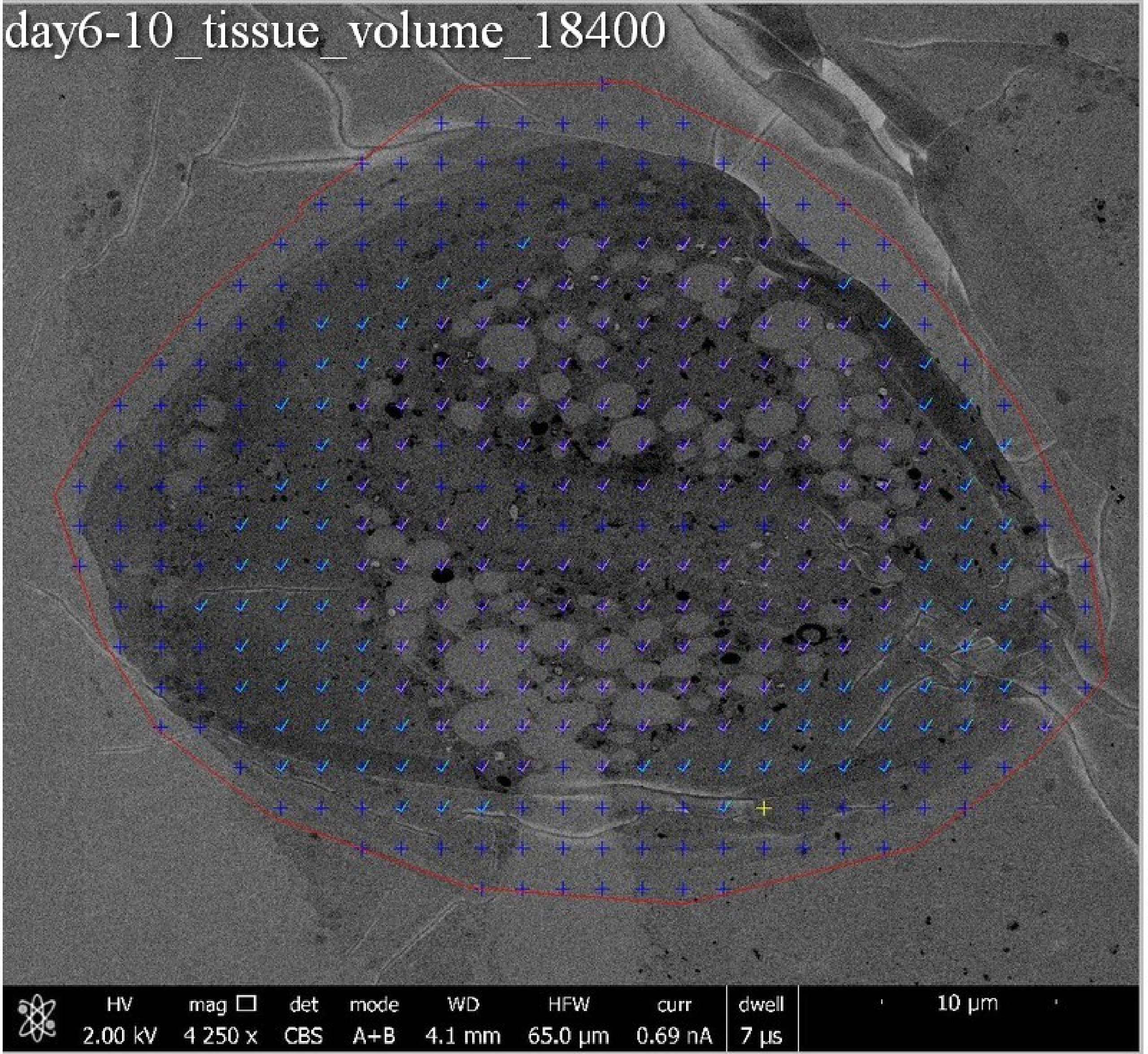
day6-10_tissue_volume_18400
HV
2.00 kV
mag
4 250 x
det
CBS
mode
A+B
WD
4.1 mm
HFW
65.0 µm
curr
0.69 nA
dwell
7 µs
10 µm

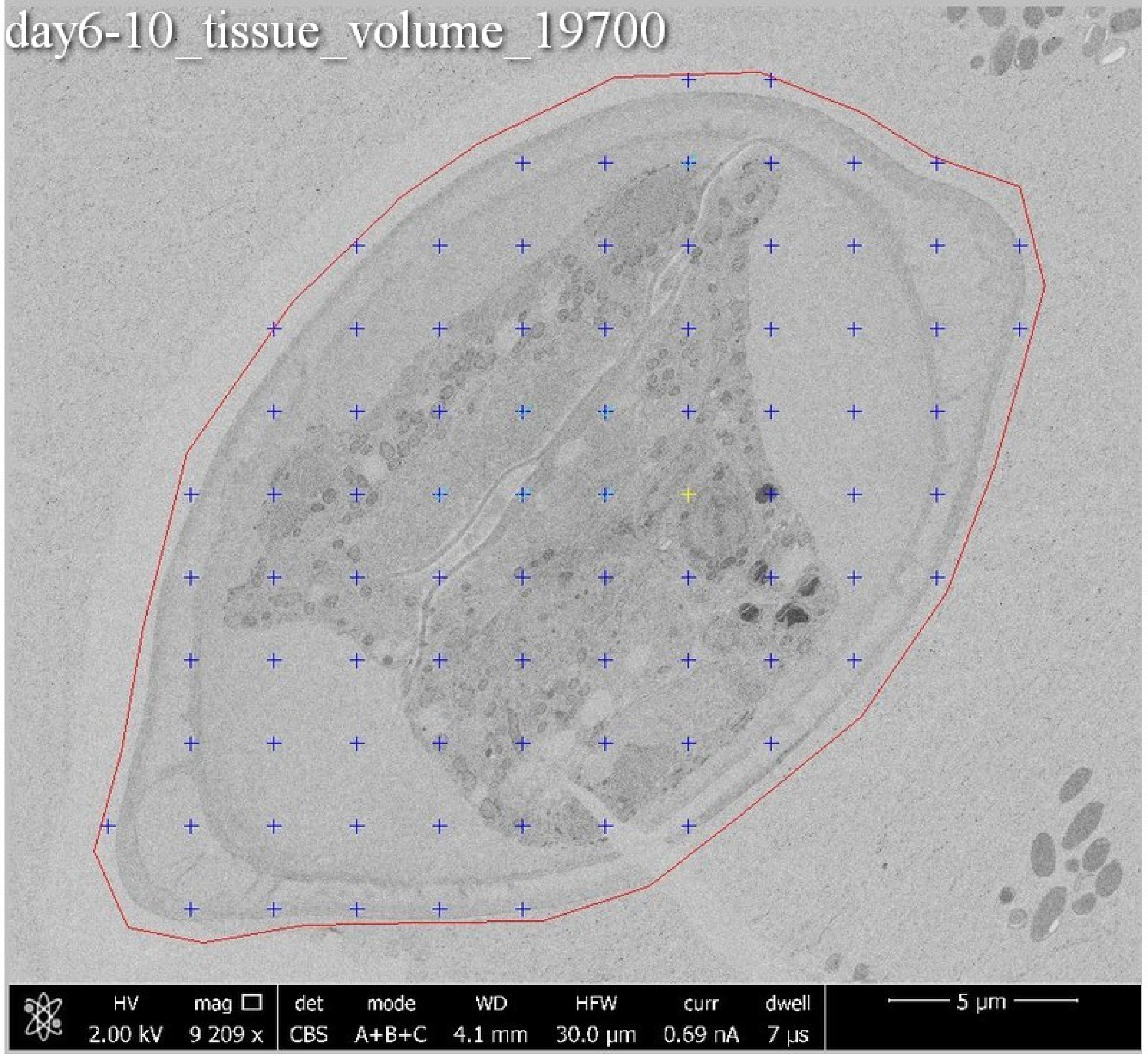
day6-10_tissue_volume_19700
HV
2.00 kV
mag
9 209 x
det
CBS
mode
A+B+C
WD
4.1 mm
HFW
30.0 μm
curr
0.69 nA
dwell
7 μs
5 μm

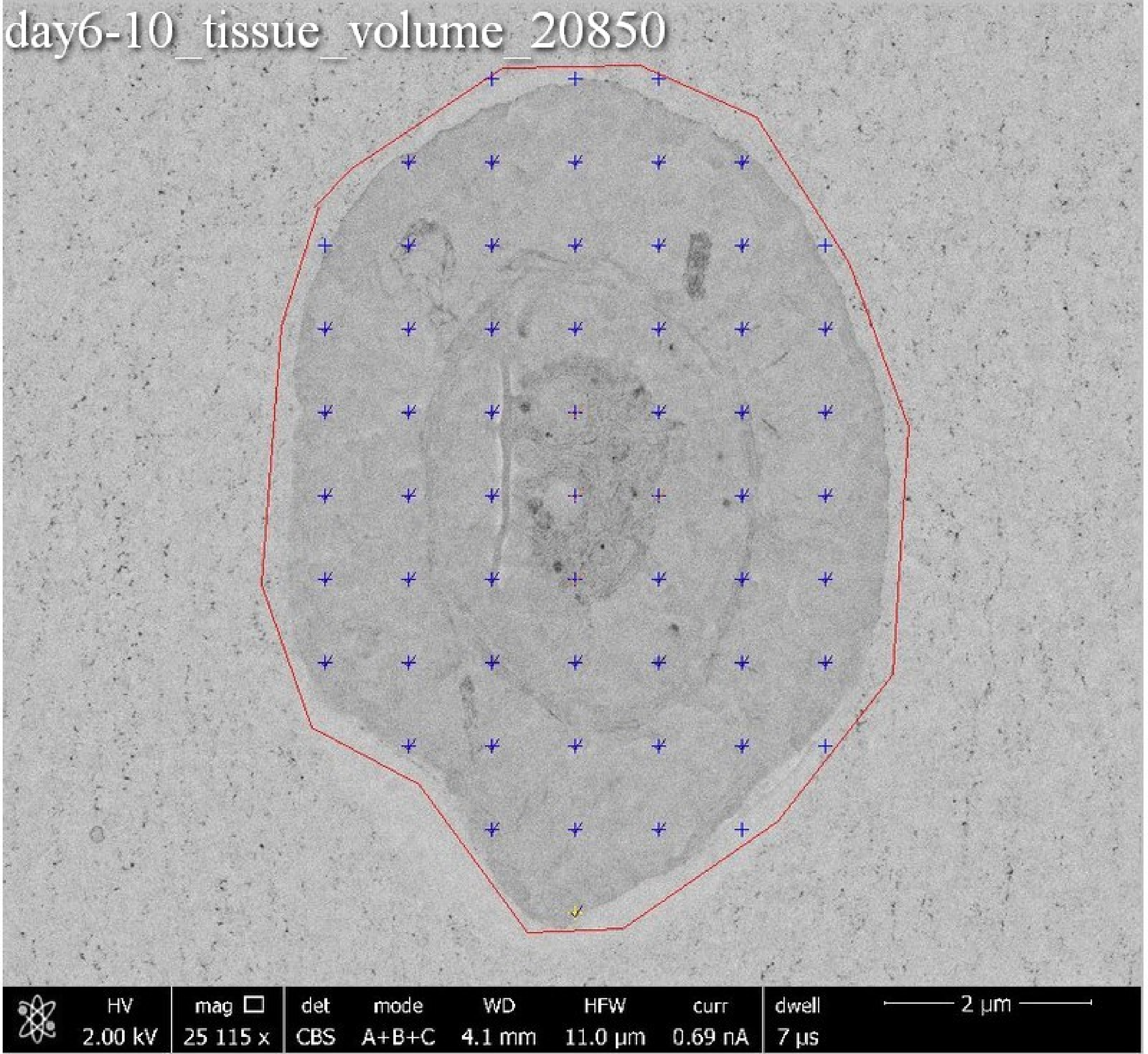
day6-10_tissue_volume_20850
HV 2.00 kV
mag 25 115 x
det CBS
mode A+B+C
WD 4.1 mm
HFW 11.0 µm
curr 0.69 nA
dwell 7 µs
2 µm

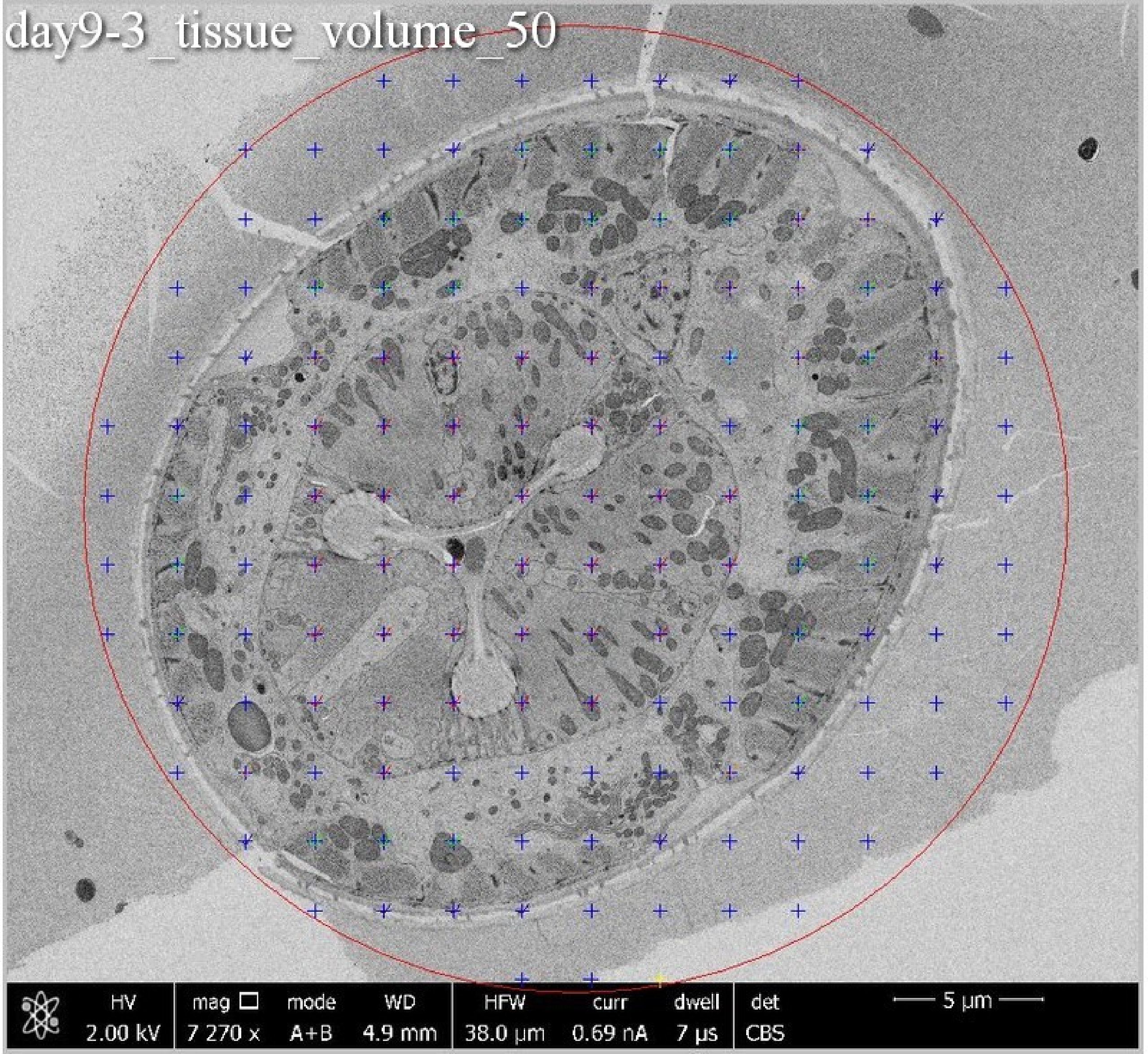
day9-3_tissue_volume_50
HV
2.00 kV
mag □
7 270 x
mode
A+B
WD
4.9 mm
HFW
38.0 µm
curr
0.69 nA
dwell
7 µs
det
CBS
5 µm

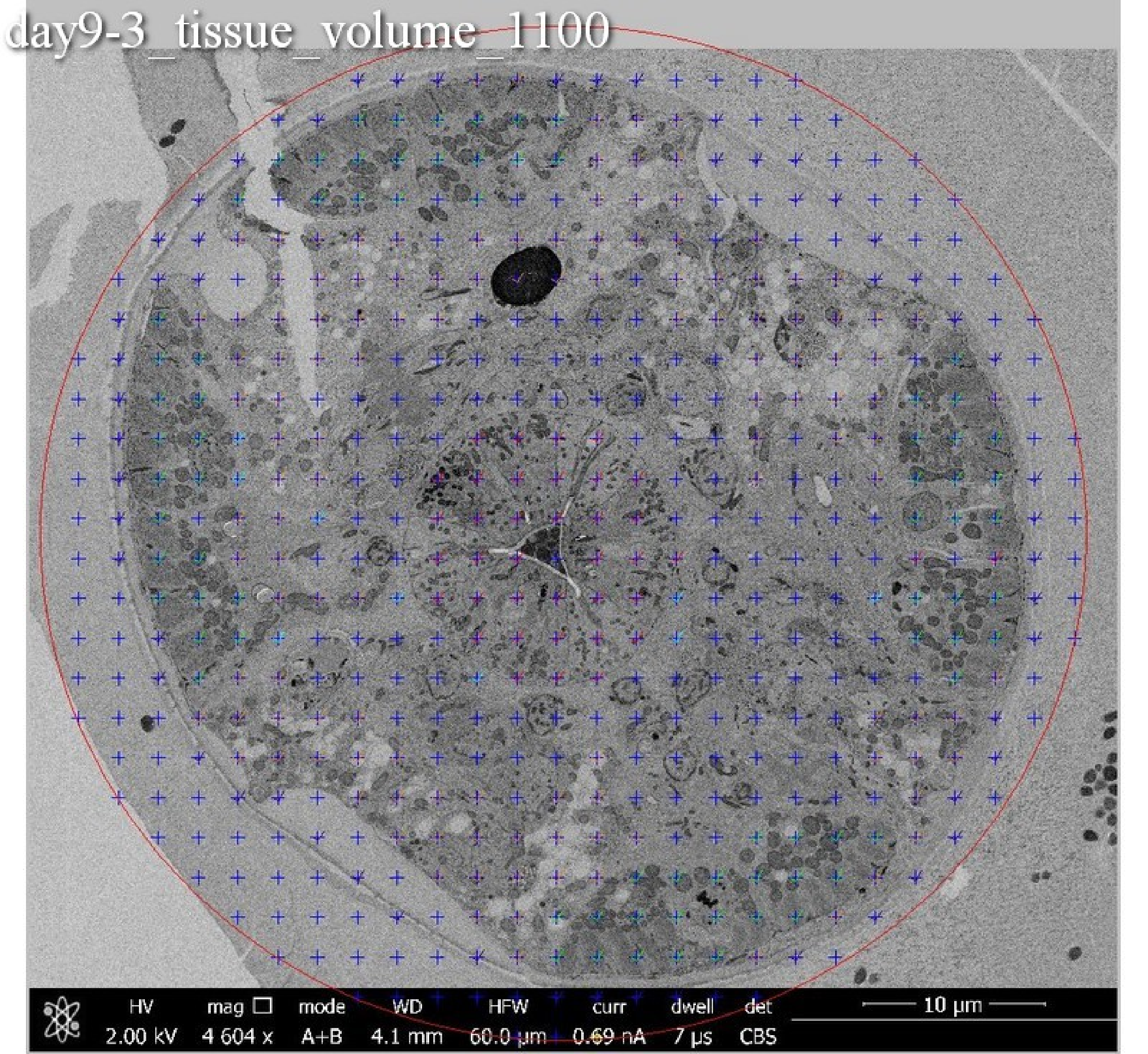
day9-3_tissue_volume_1100
HV
2.00 kV
mag
4 604 x
mode
A+B
WD
4.1 mm
HFW
60.0 µm
curr
0.69 nA
dwell
7 µs
det
CBS
10 µm

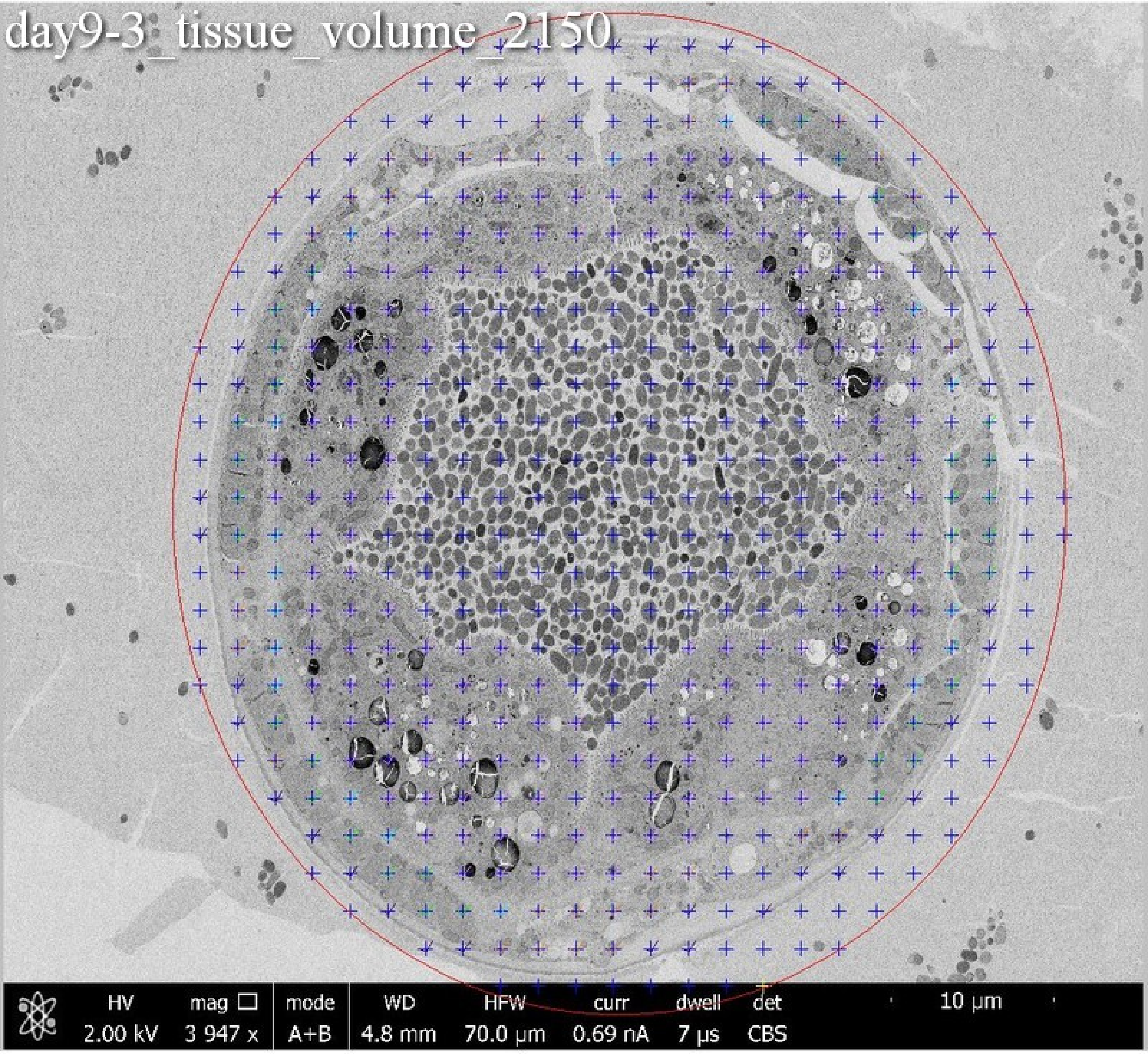
day9-3_tissue_volume_2150
HV 2.00 kV
mag □ 3 947 x
mode A+B
WD 4.8 mm
HFW 70.0 µm
curr 0.69 nA
dwell 7 µs
det CBS
10 µm

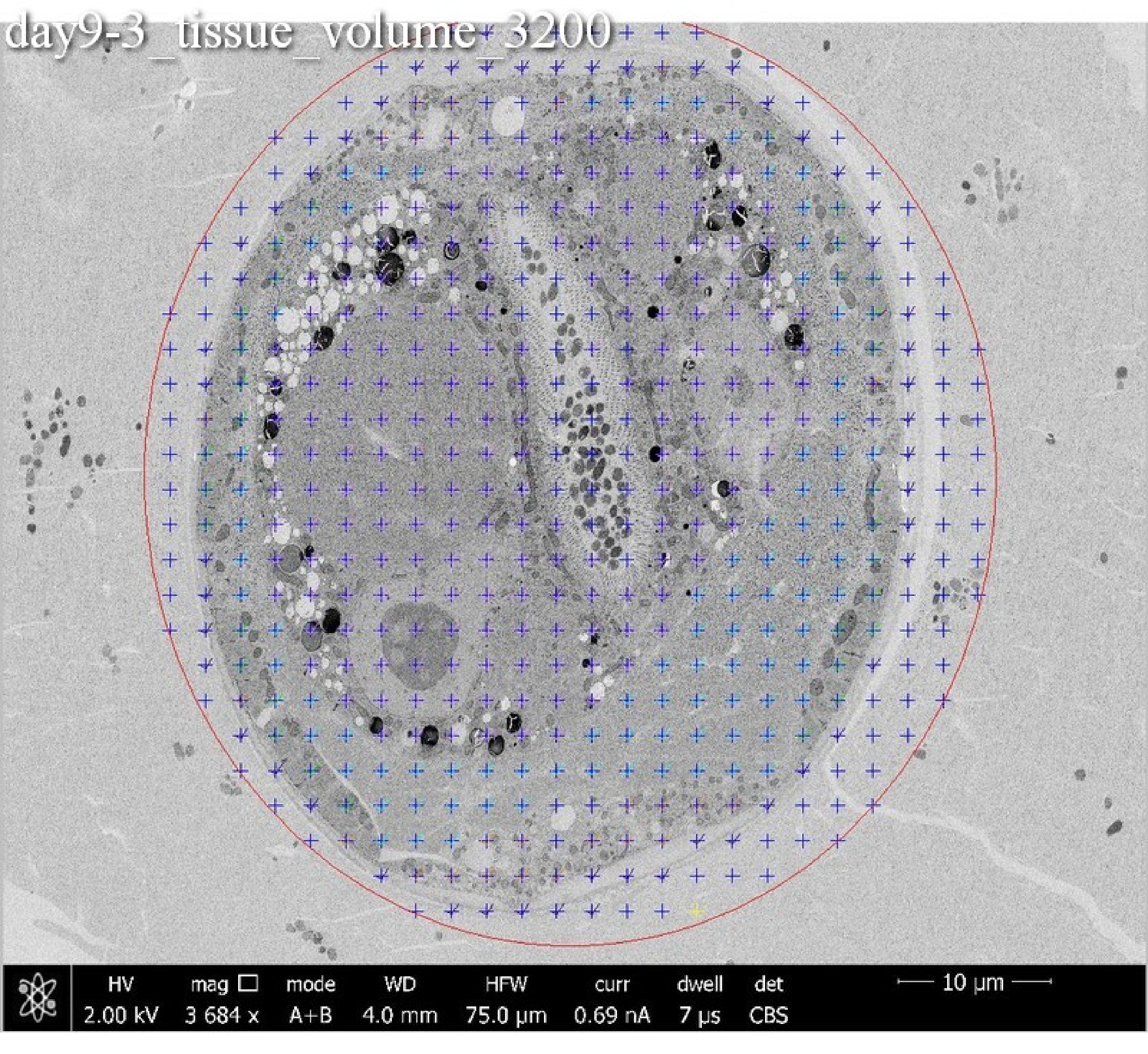
day9-3_tissue_volume_3200
HV 2.00 kV
mag □ 3 684 x
mode A+B
WD 4.0 mm
HFW 75.0 μm
curr 0.69 nA
dwell 7 μs
det CBS
10 μm

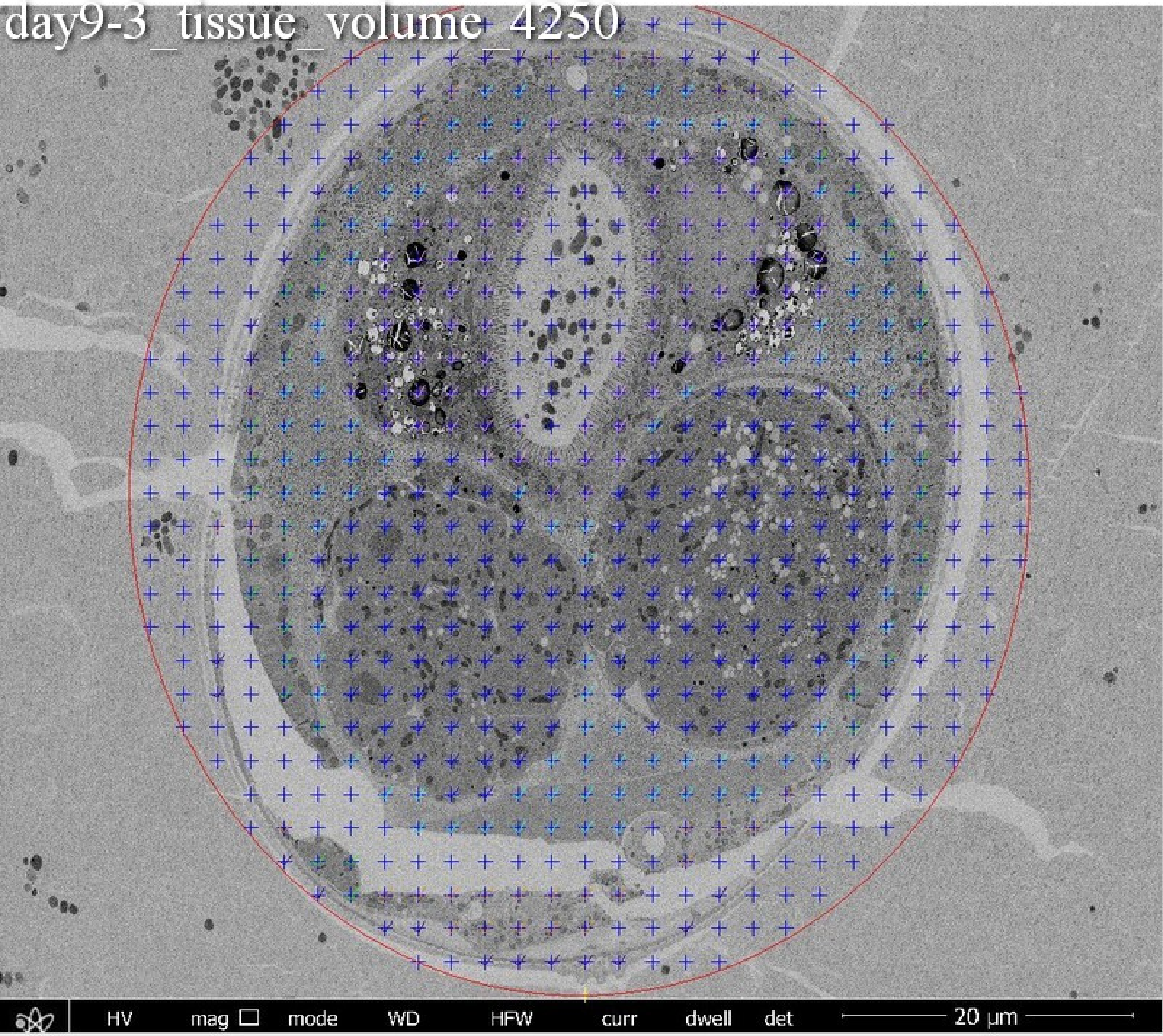
day9-3_tissue_volume_4250
HV
mag
mode
WD
HFW
curr
dwell
det
20 µm

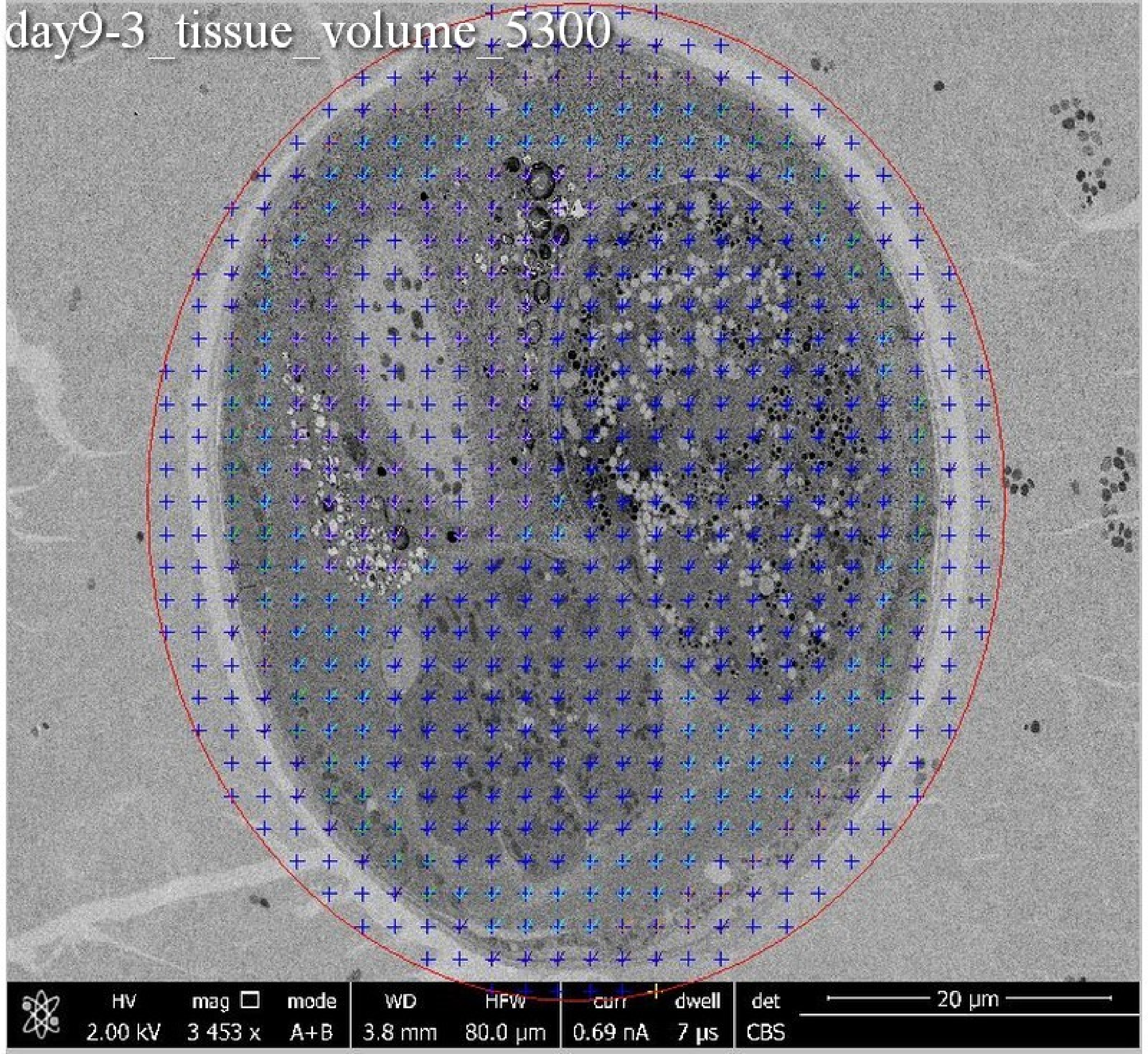
day9-3_tissue_volume_5300
HV
2.00 kV
mag
3 453 x
mode
A+B
WD
3.8 mm
HFW
80.0 µm
curr
0.69 nA
dwell
7 µs
det
CBS
20 µm

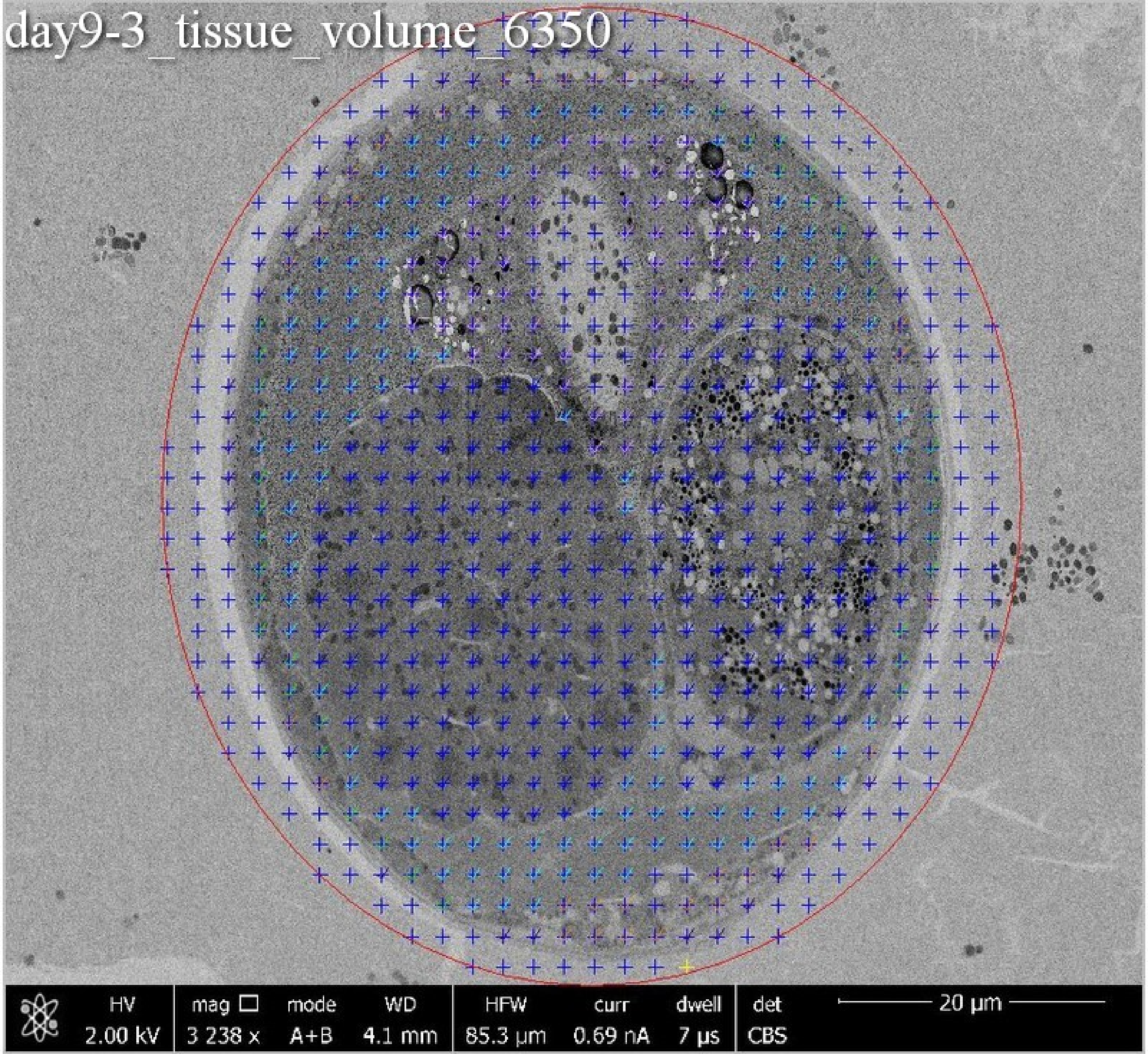
day9-3_tissue_volume_6350
HV
2.00 kV
mag
3 238 x
mode
A+B
WD
4.1 mm
HFW
85.3 μm
curr
0.69 nA
dwell
7 μs
det
CBS
20 μm

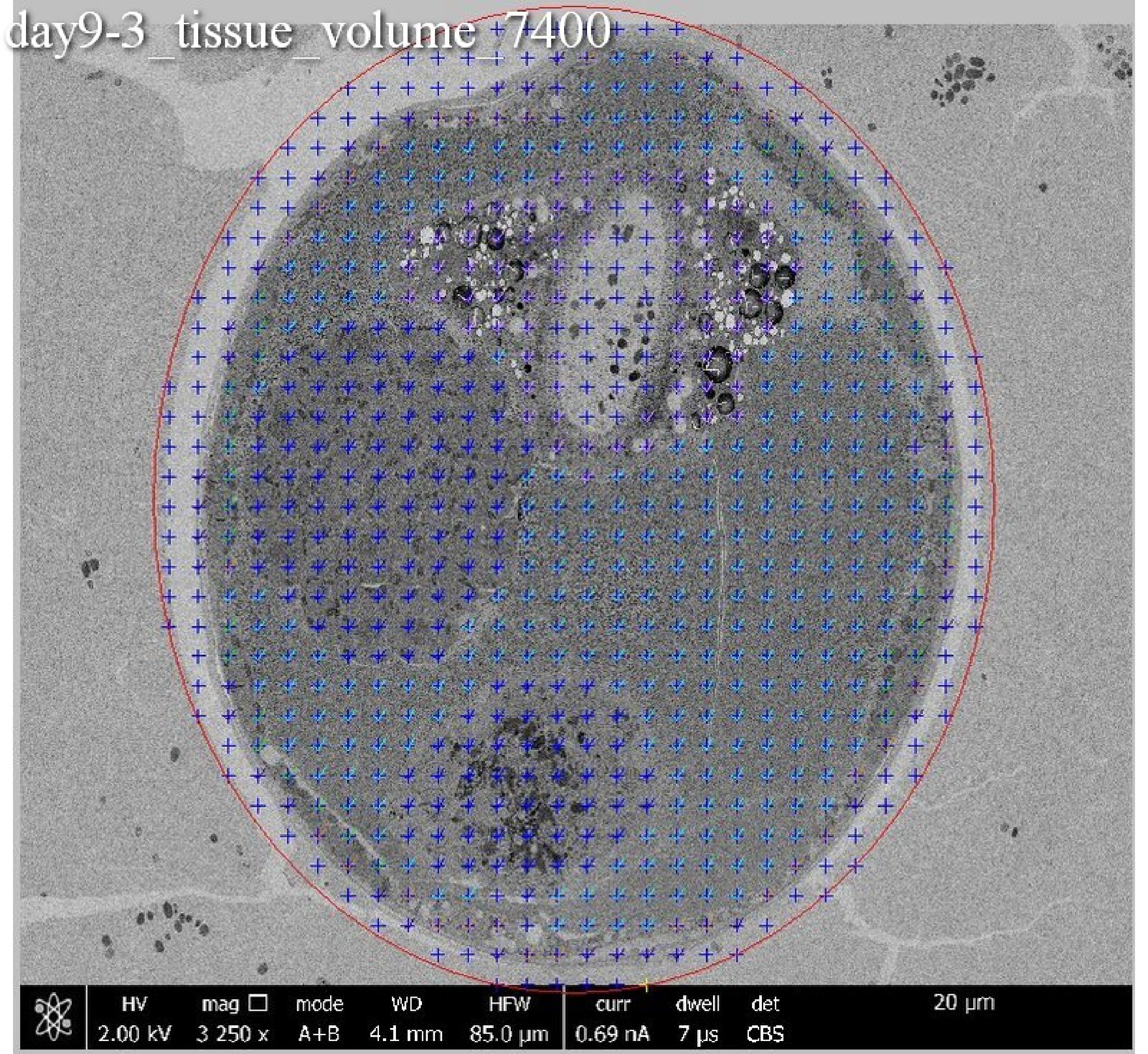
day9-3_tissue_volume 7400
HV
2.00 kV
mag
3 250 x
mode
A+B
WD
4.1 mm
HFW
85.0 µm
curr
0.69 nA
dwell
7 µs
det
CBS
20 µm

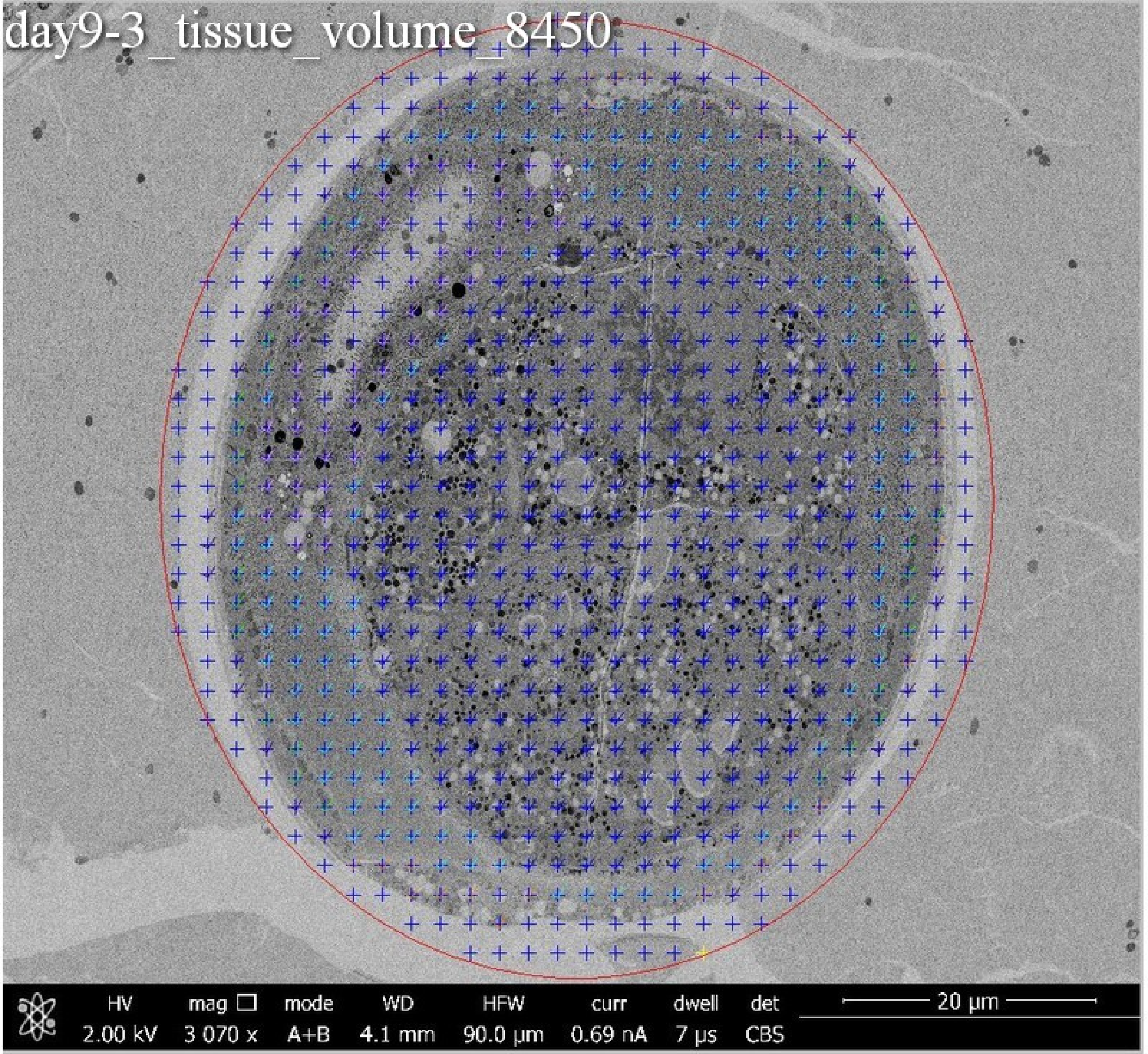

day9-3_tissue_volume_8450
HV 2.00 kV
mag 3 070 x
mode A+B
WD 4.1 mm
HFW 90.0 µm
curr 0.69 nA
dwell 7 µs
det CBS
20 µm

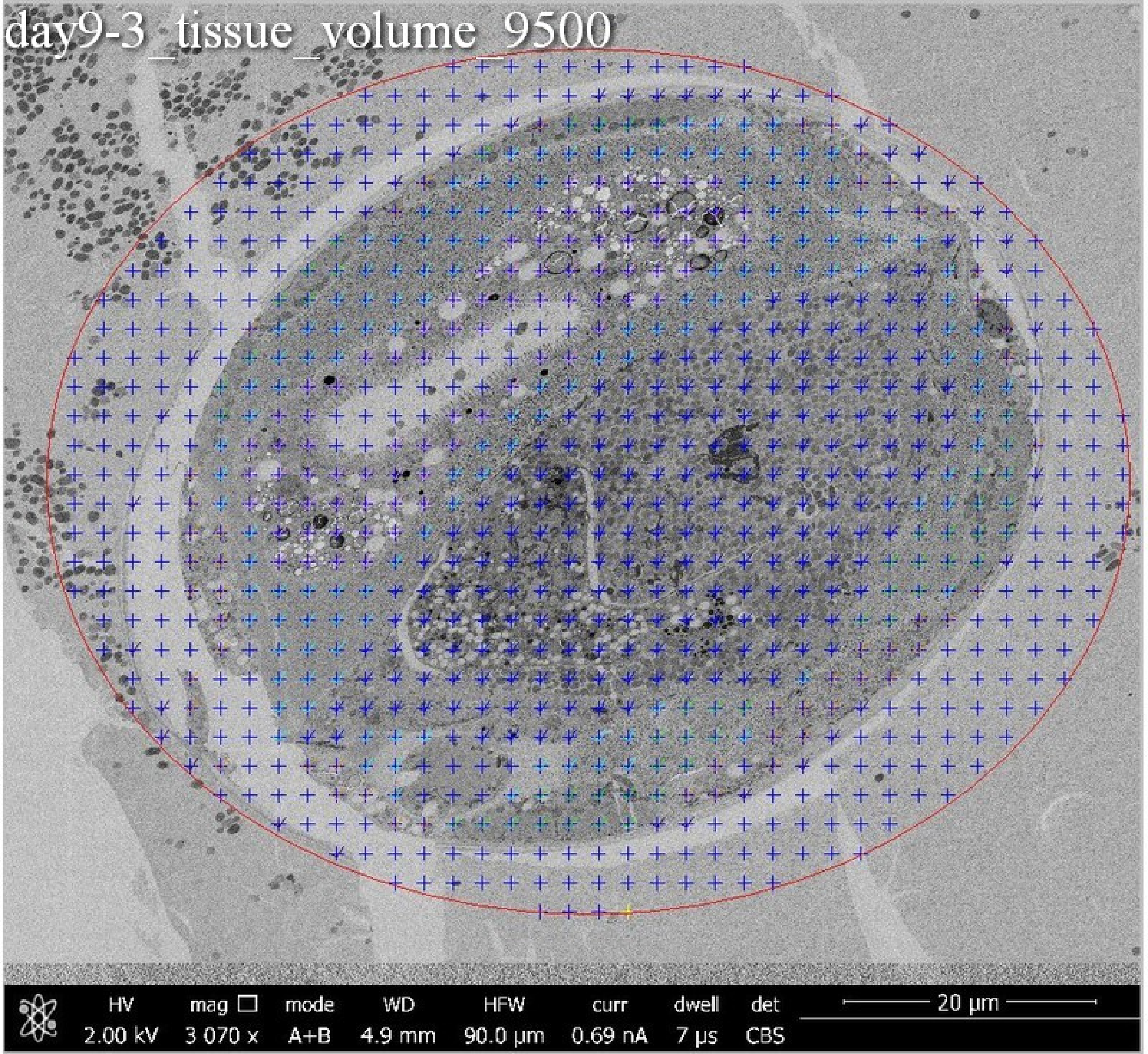
day9-3_tissue_volume_9500
HV 2.00 kV
mag 3 070 x
mode A+B
WD 4.9 mm
HFW 90.0 µm
curr 0.69 nA
dwell 7 µs
det CBS
20 µm

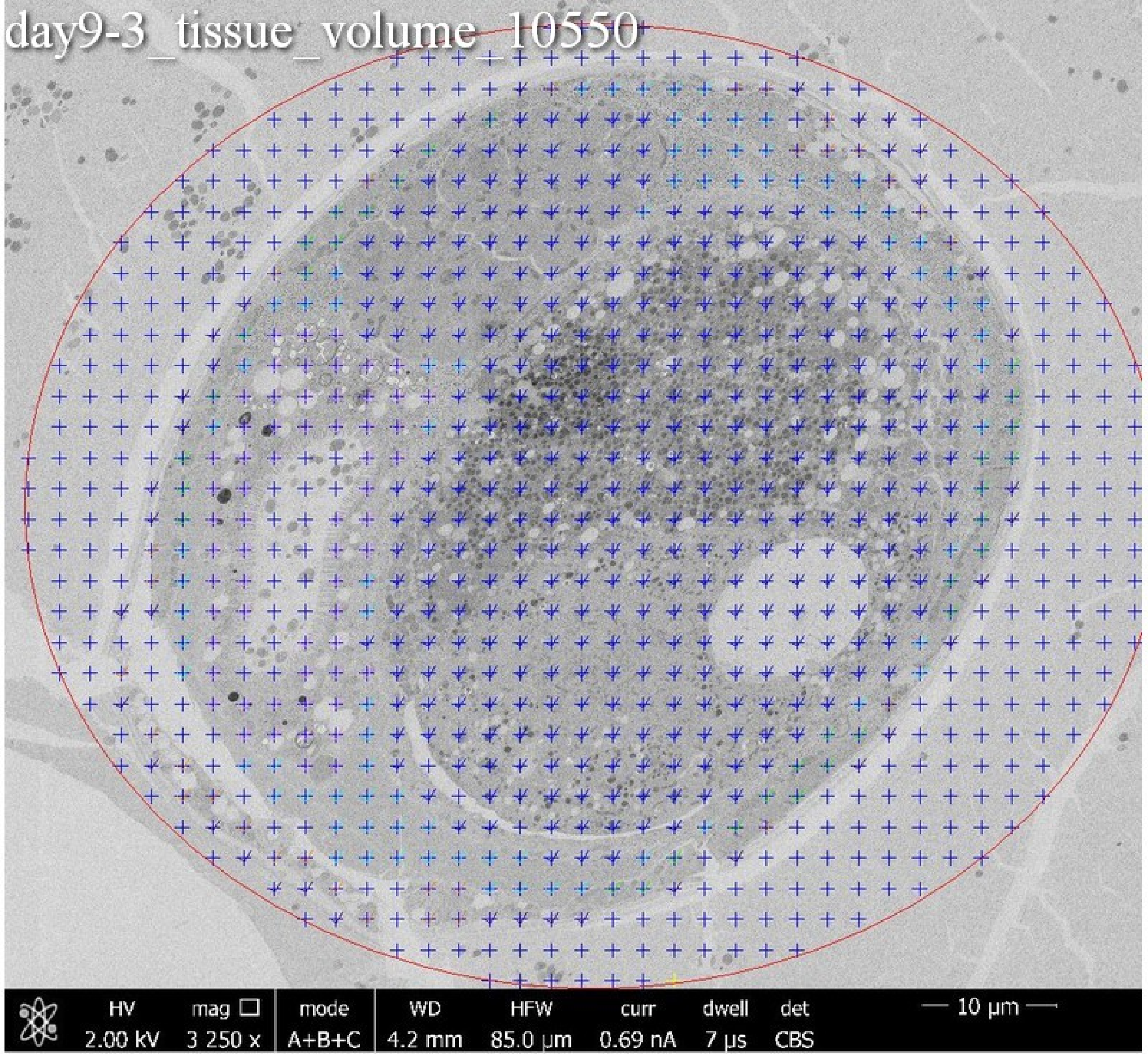
day9-3_tissue_volume_10550
HV 2.00 kV
mag 3 250 x
mode A+B+C
WD 4.2 mm
HFW 85.0 µm
curr 0.69 nA
dwell 7 µs
det CBS
10 µm

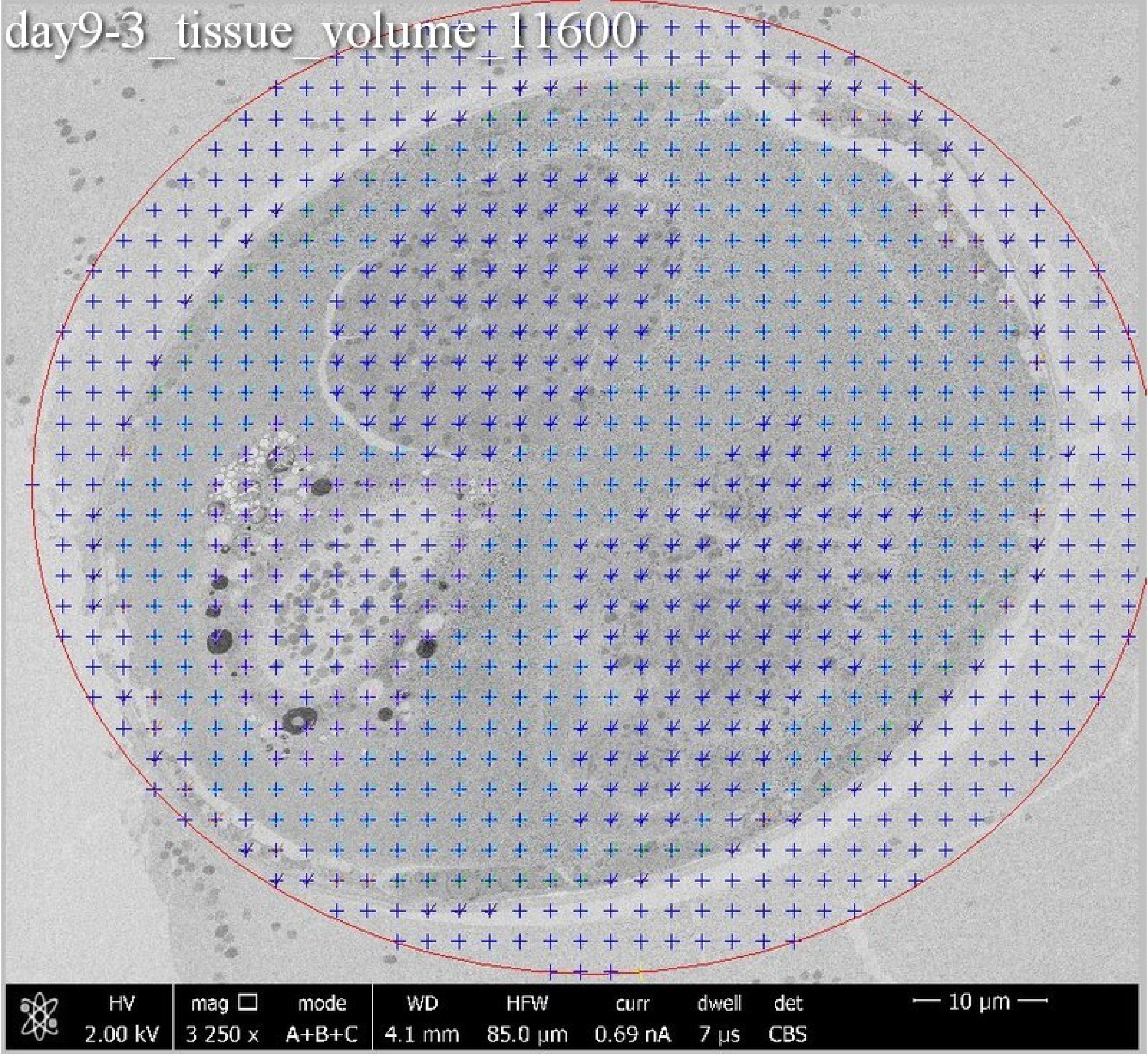
day9-3_tissue_volume_11600
HV
2.00 kV
mag
3 250 x
mode
A+B+C
WD
4.1 mm
HFW
85.0 µm
curr
0.69 nA
dwell
7 µs
det
CBS
10 µm

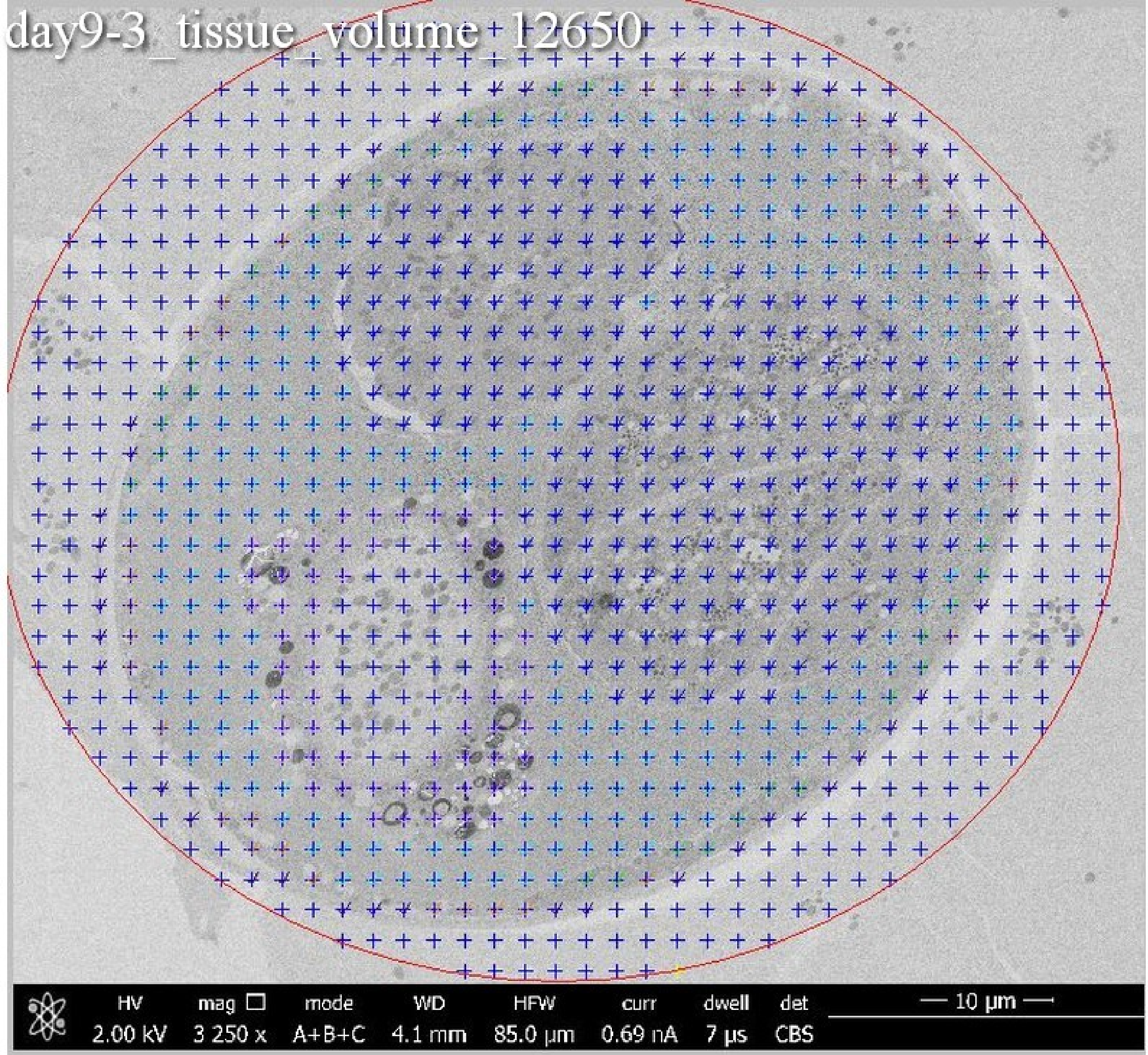
day9-3_tissue_volume 12650
HV 2.00 kV
mag 3 250 x
mode A+B+C
WD 4.1 mm
HFW 85.0 µm
curr 0.69 nA
dwell 7 µs
det CBS
10 µm

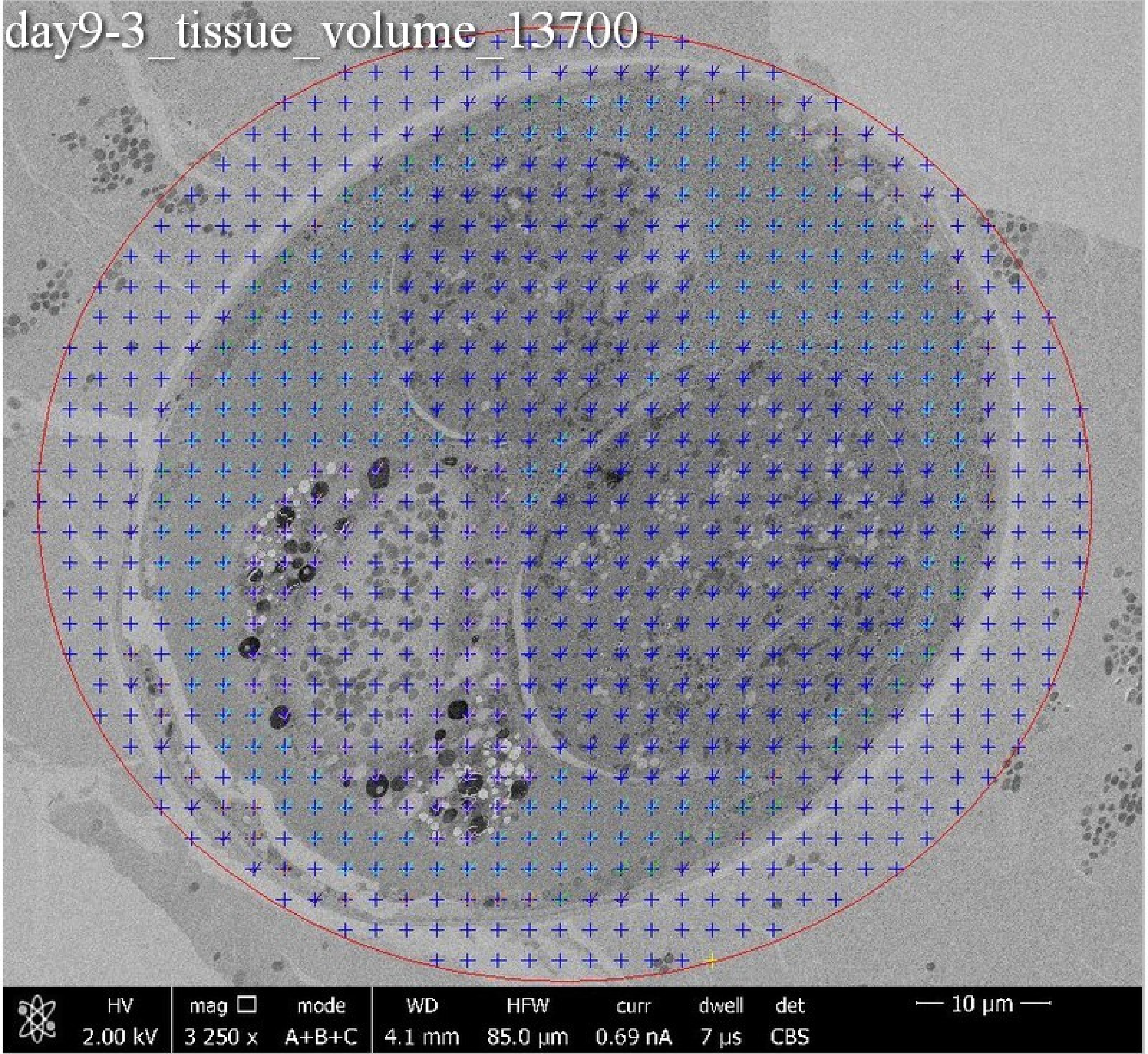
day9-3_tissue_volume_13700
HV
2.00 kV
mag □
3 250 x
mode
A+B+C
WD
4.1 mm
HFW
85.0 µm
curr
0.69 nA
dwell
7 µs
det
CBS
10 µm

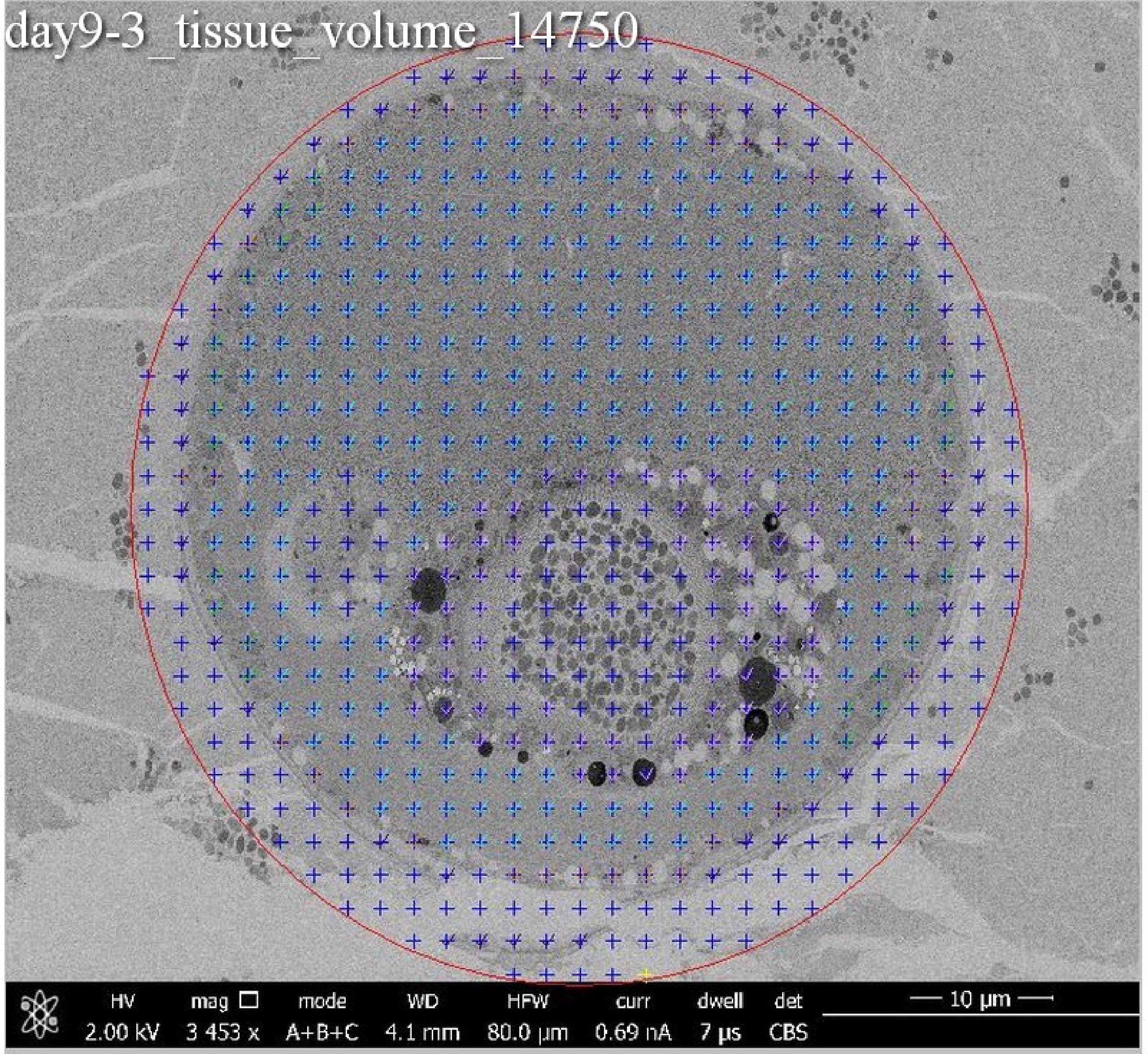
day9-3_tissue_volume_14750
HV
2.00 kV
mag
3 453 x
mode
A+B+C
WD
4.1 mm
HFW
80.0 µm
curr
0.69 nA
dwell
7 µs
det
CBS
10 µm

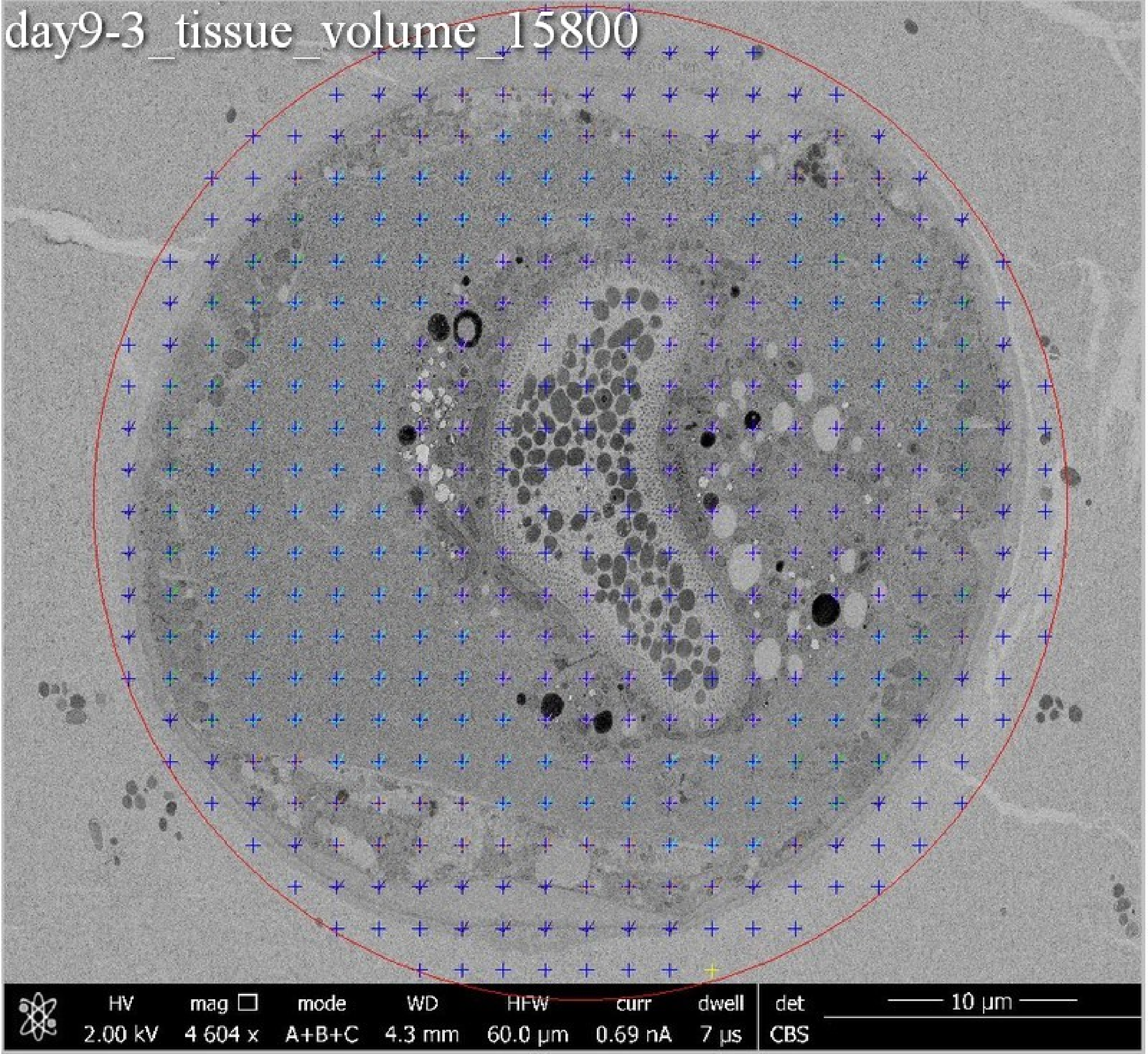
day9-3_tissue_volume_15800
HV 2.00 kV
mag 4 604 x
mode A+B+C
WD 4.3 mm
HFW 60.0 µm
curr 0.69 nA
dwell 7 µs
det CBS
10 µm

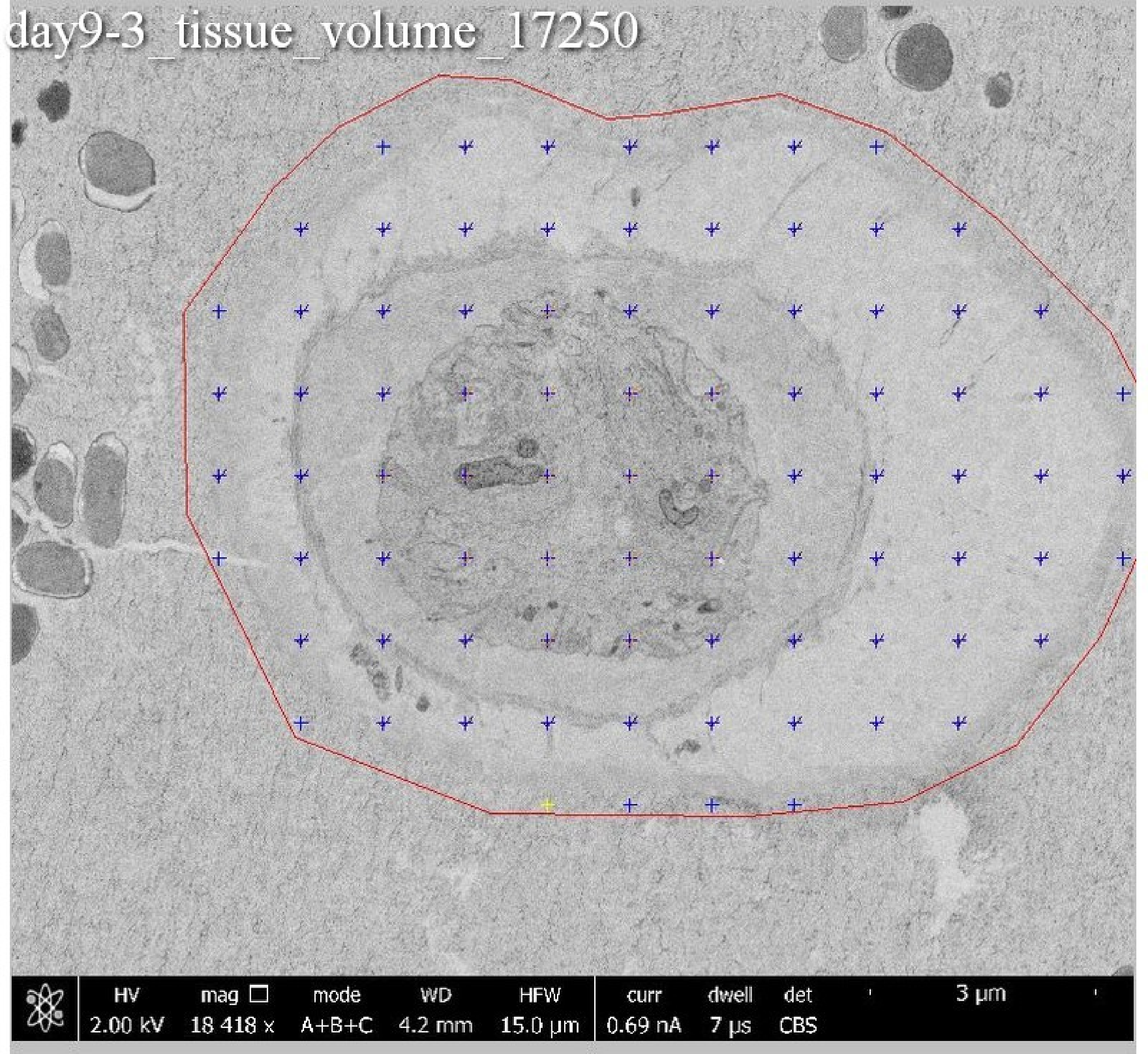
day9-3_tissue_volume_17250
HV 2.00 kV
mag 18 418 x
mode A+B+C
WD 4.2 mm
HFW 15.0 µm
curr 0.69 nA
dwell 7 µs
det CBS
3 µm

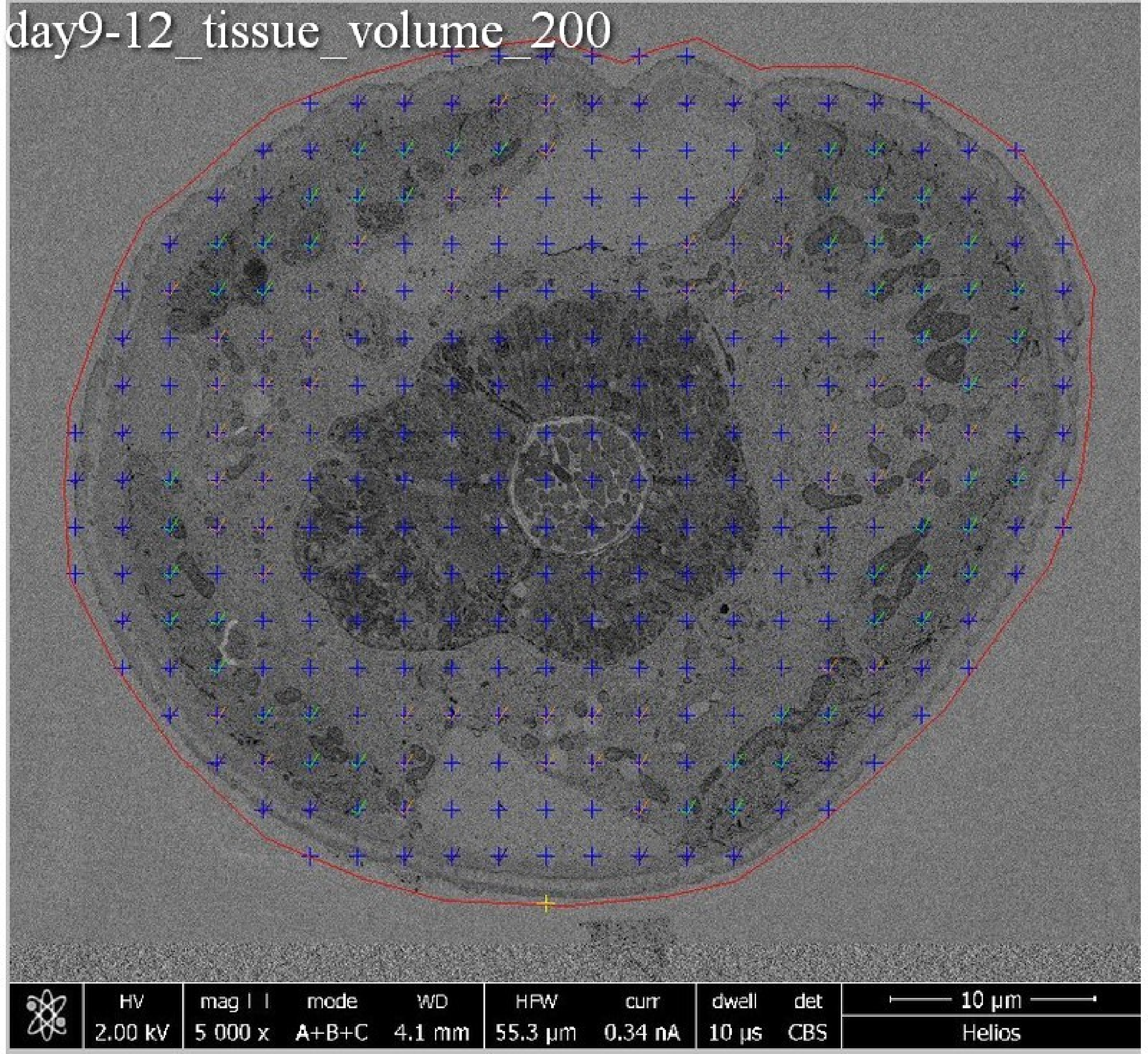
day9-12_tissue_volume_200
HV
2.00 kV
mag
5 000 x
mode
A+B+C
WD
4.1 mm
HFW
55.3 µm
curr
0.34 nA
dwell
10 µs
det
CBS
10 µm
Helios

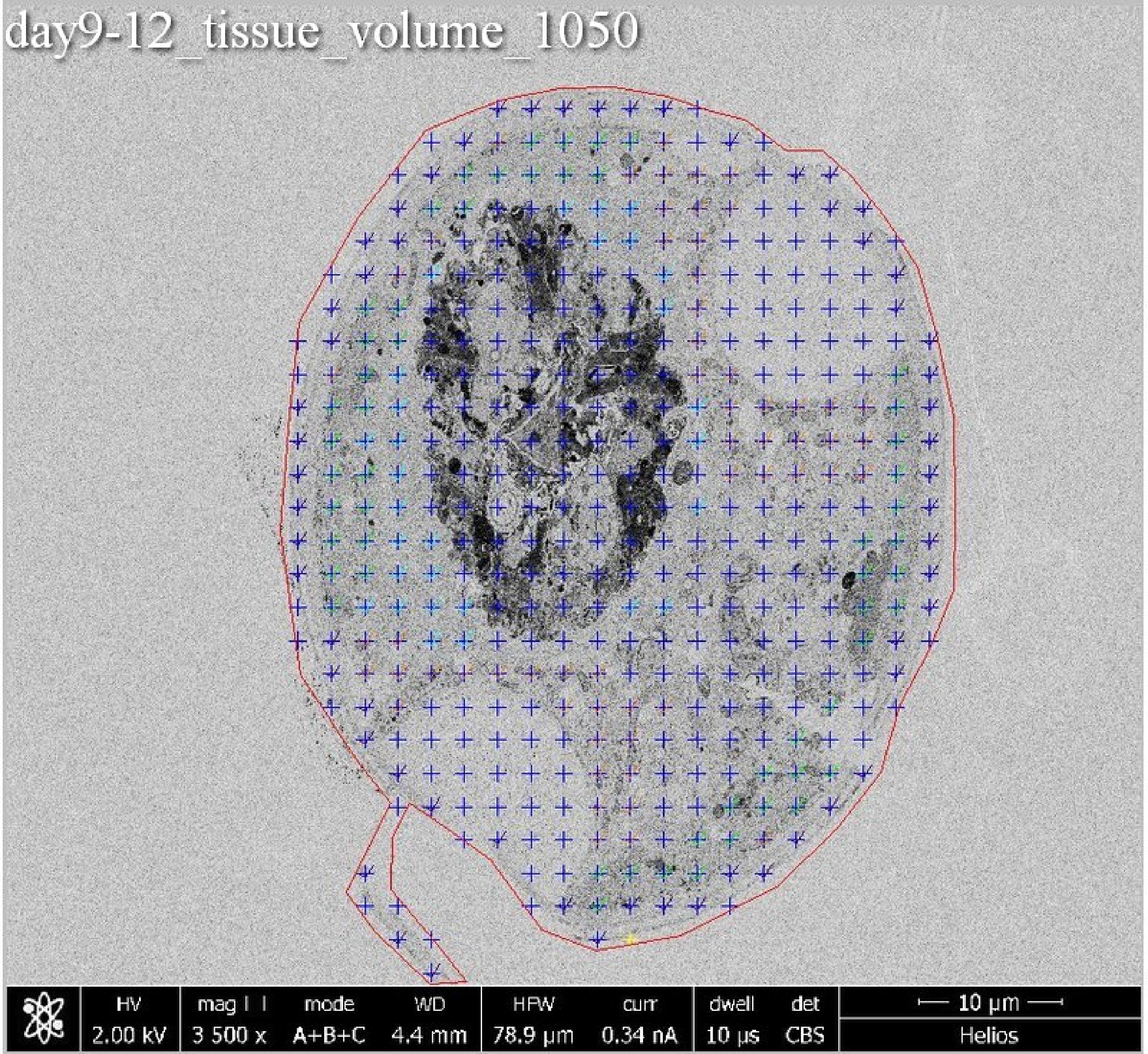

day9-12_tissue_volume_1050
HV
2.00 kV
mag
3 500 x
mode
A+B+C
WD
4.4 mm
HFW
78.9 µm
curr
0.34 nA
dwell
10 µs
det
CBS
10 µm
Helios

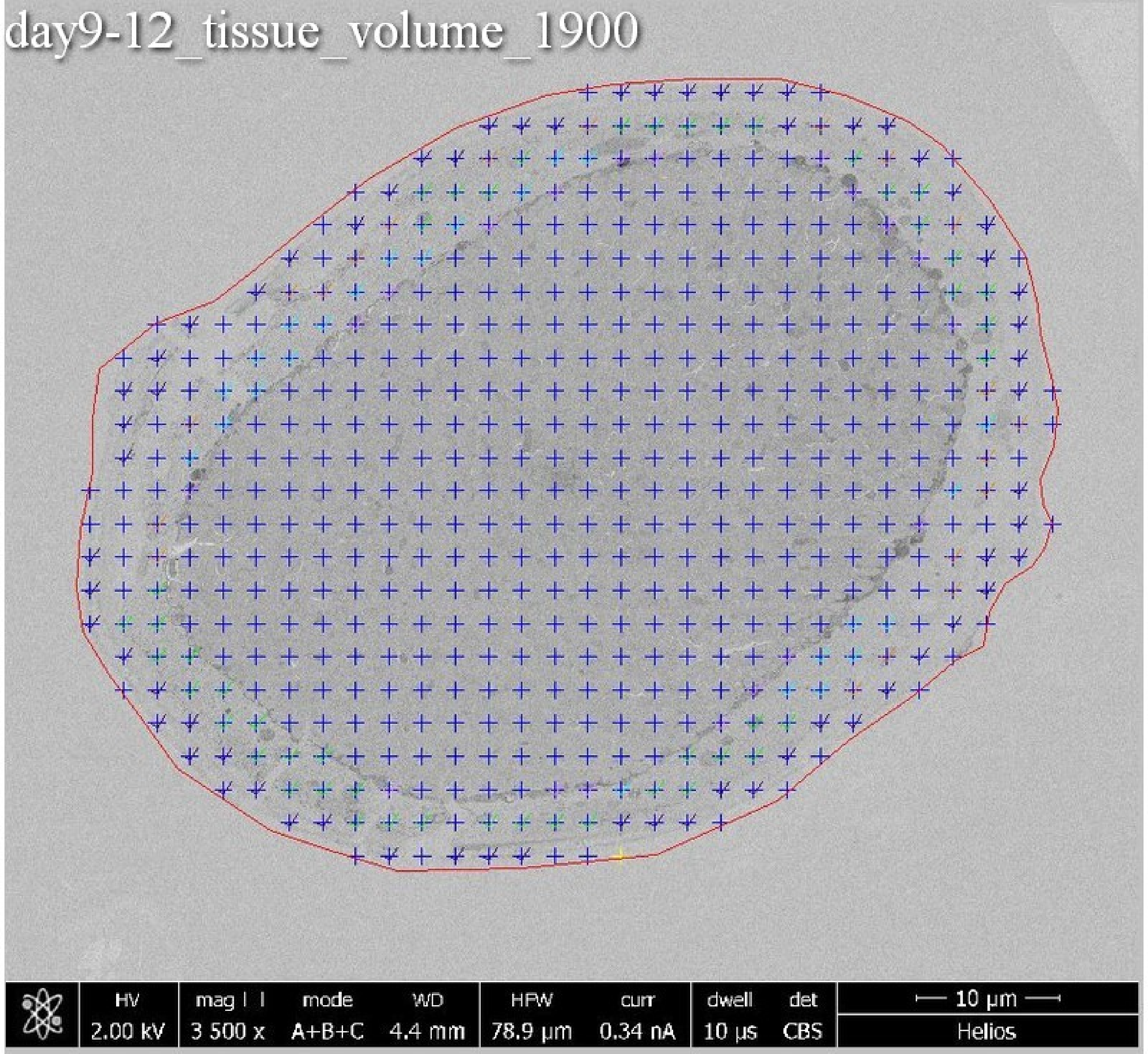
day9-12_tissue_volume_1900
HV
2.00 kV
mag
3 500 x
mode
A+B+C
WD
4.4 mm
HFW
78.9 µm
curr
0.34 nA
dwell
10 µs
det
CBS
10 µm
Helios

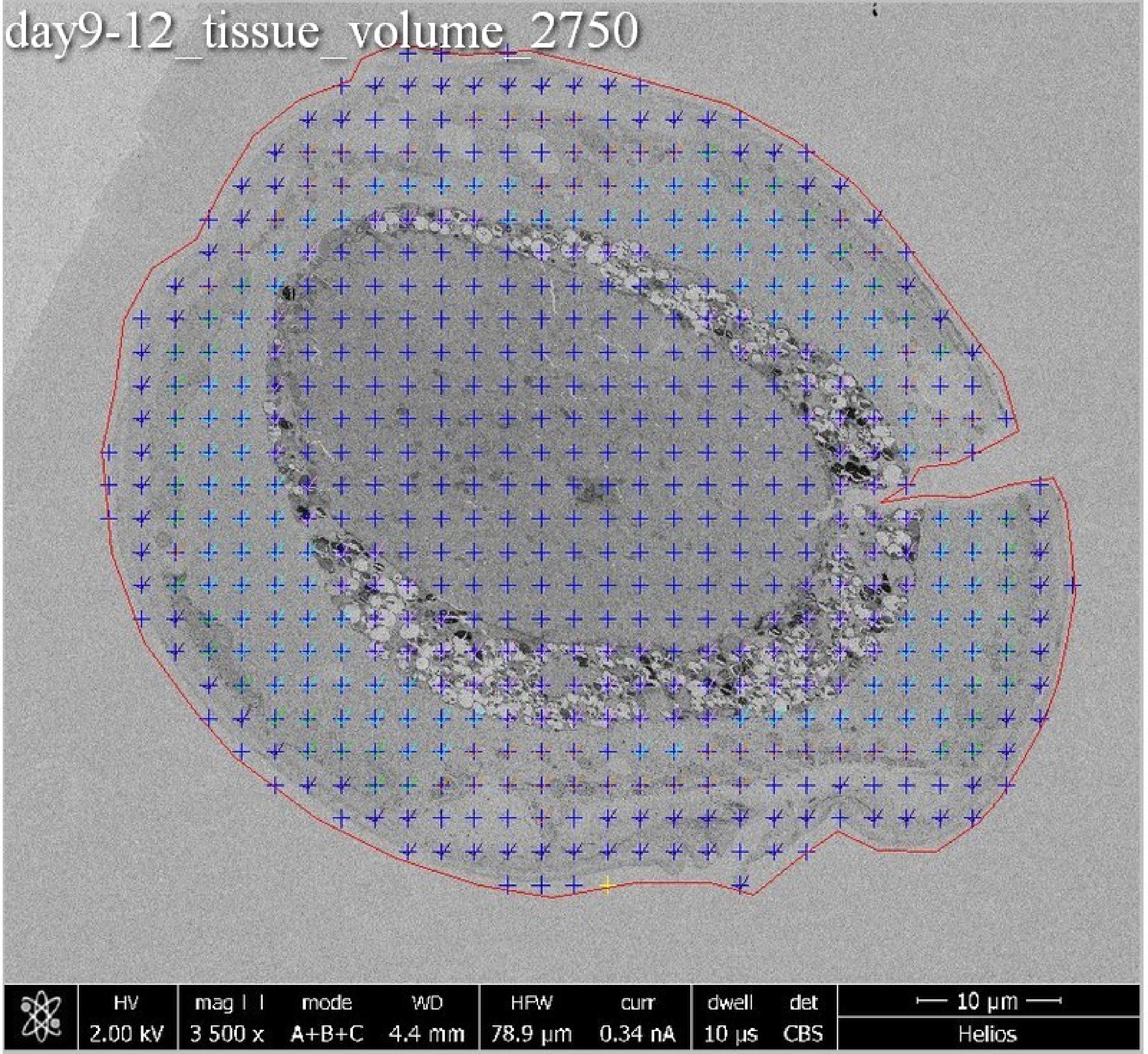
day9-12_tissue_volume_2750
HV
2.00 kV
mag
3 500 x
mode
A+B+C
WD
4.4 mm
HFW
78.9 µm
curr
0.34 nA
dwell
10 µs
det
CBS
10 µm
Helios

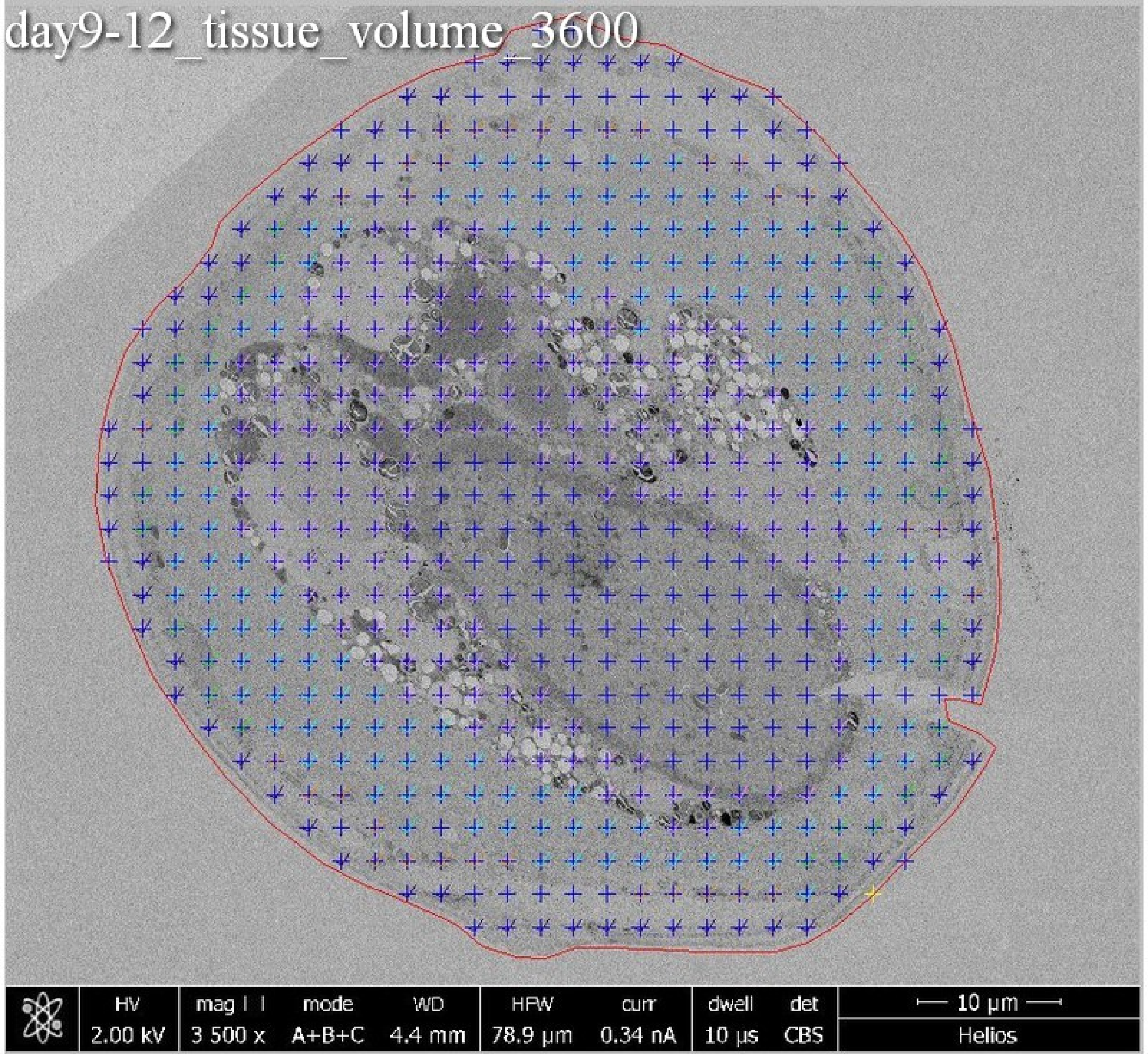
day9-12_tissue_volume_3600
HV 2.00 kV
mag 3 500 x
mode A+B+C
WD 4.4 mm
HFW 78.9 µm
curr 0.34 nA
dwell 10 µs
det CBS
10 µm
Helios

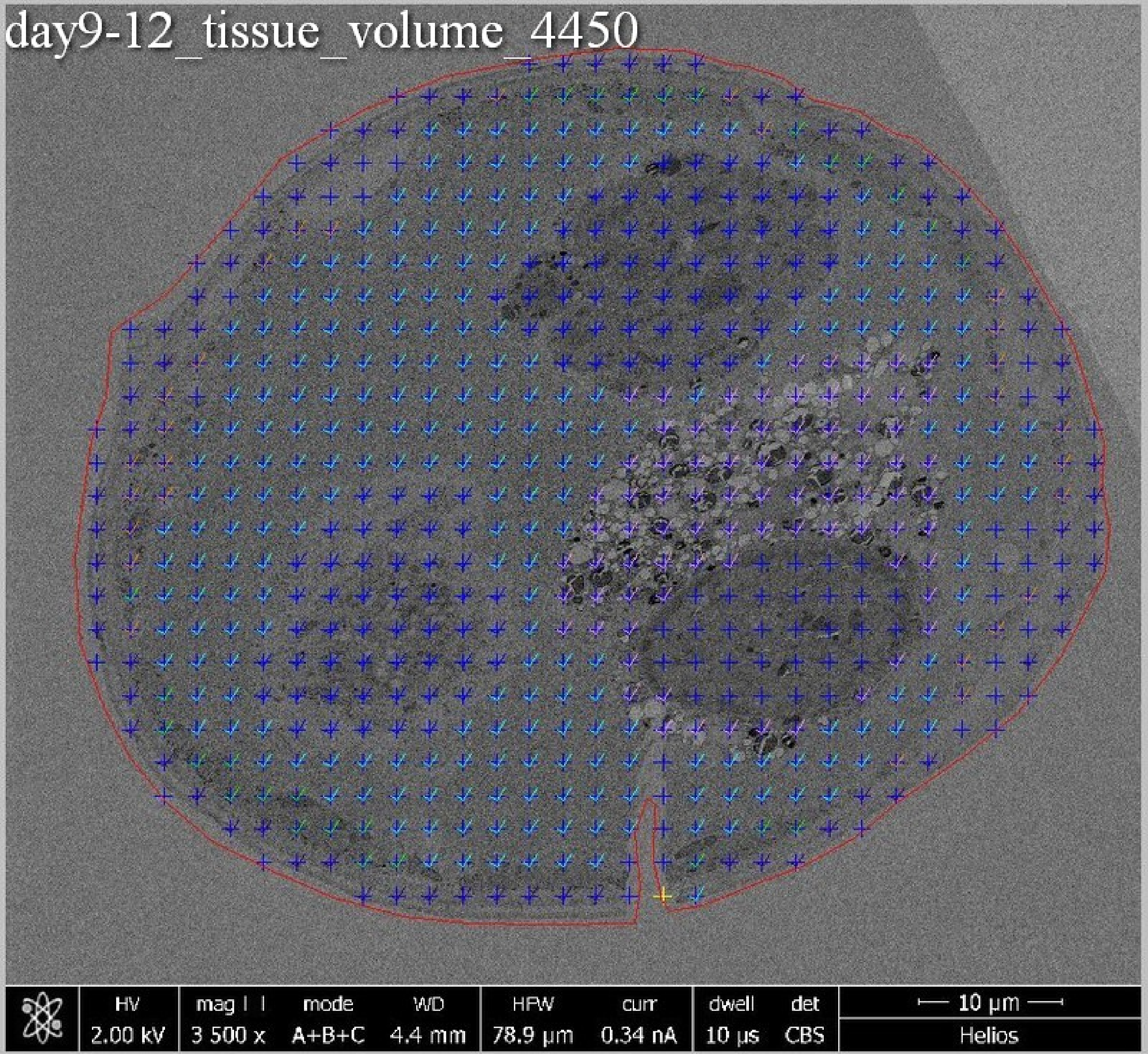
day9-12_tissue_volume_4450
HV
2.00 kV
mag
3 500 x
mode
A+B+C
WD
4.4 mm
HFW
78.9 µm
curr
0.34 nA
dwell
10 µs
det
CBS
10 µm
Helios

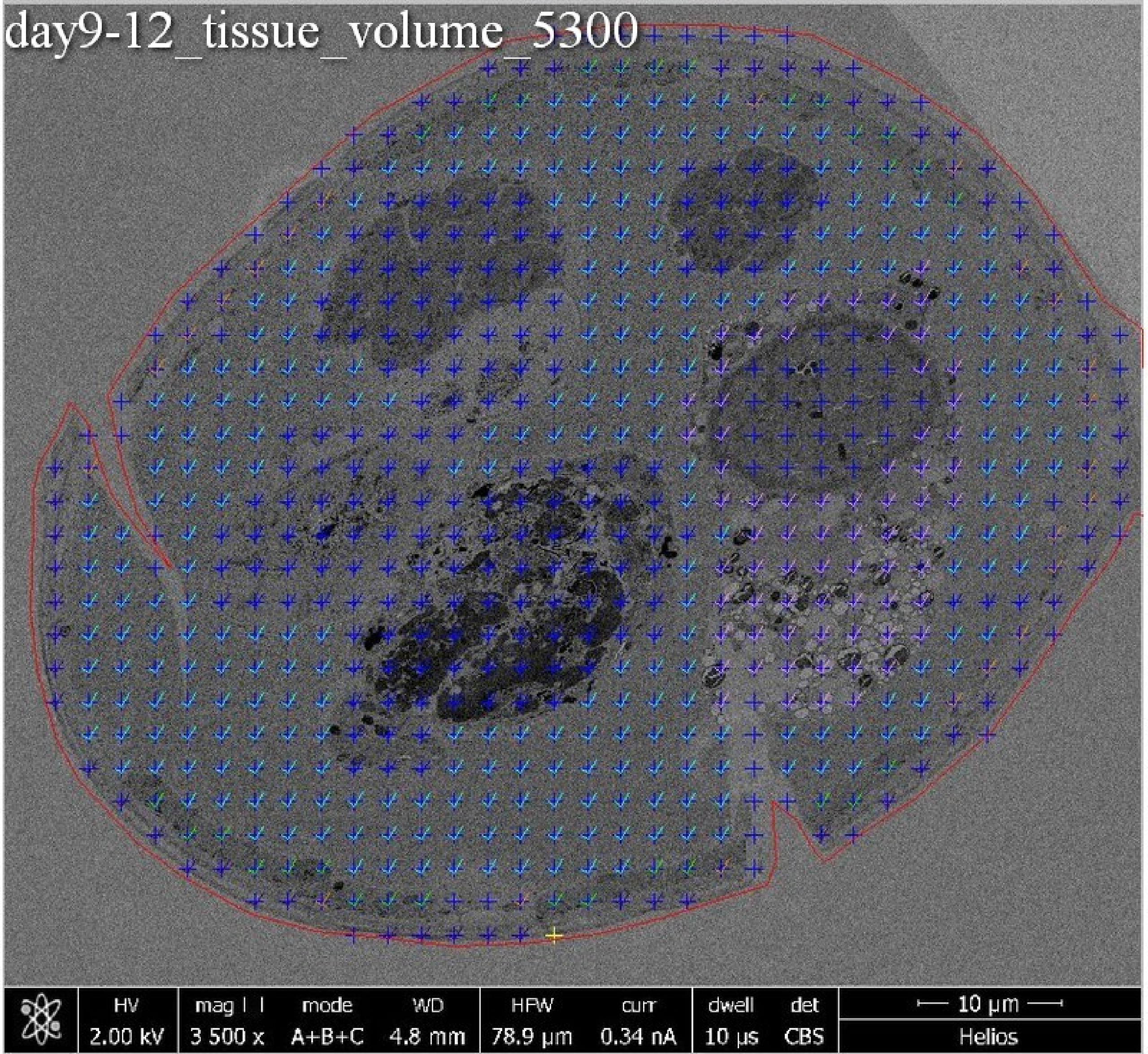
day9-12_tissue_volume_5300
HV
2.00 kV
mag
3 500 x
mode
A+B+C
WD
4.8 mm
HFW
78.9 µm
curr
0.34 nA
dwell
10 µs
det
CBS
10 µm
Helios

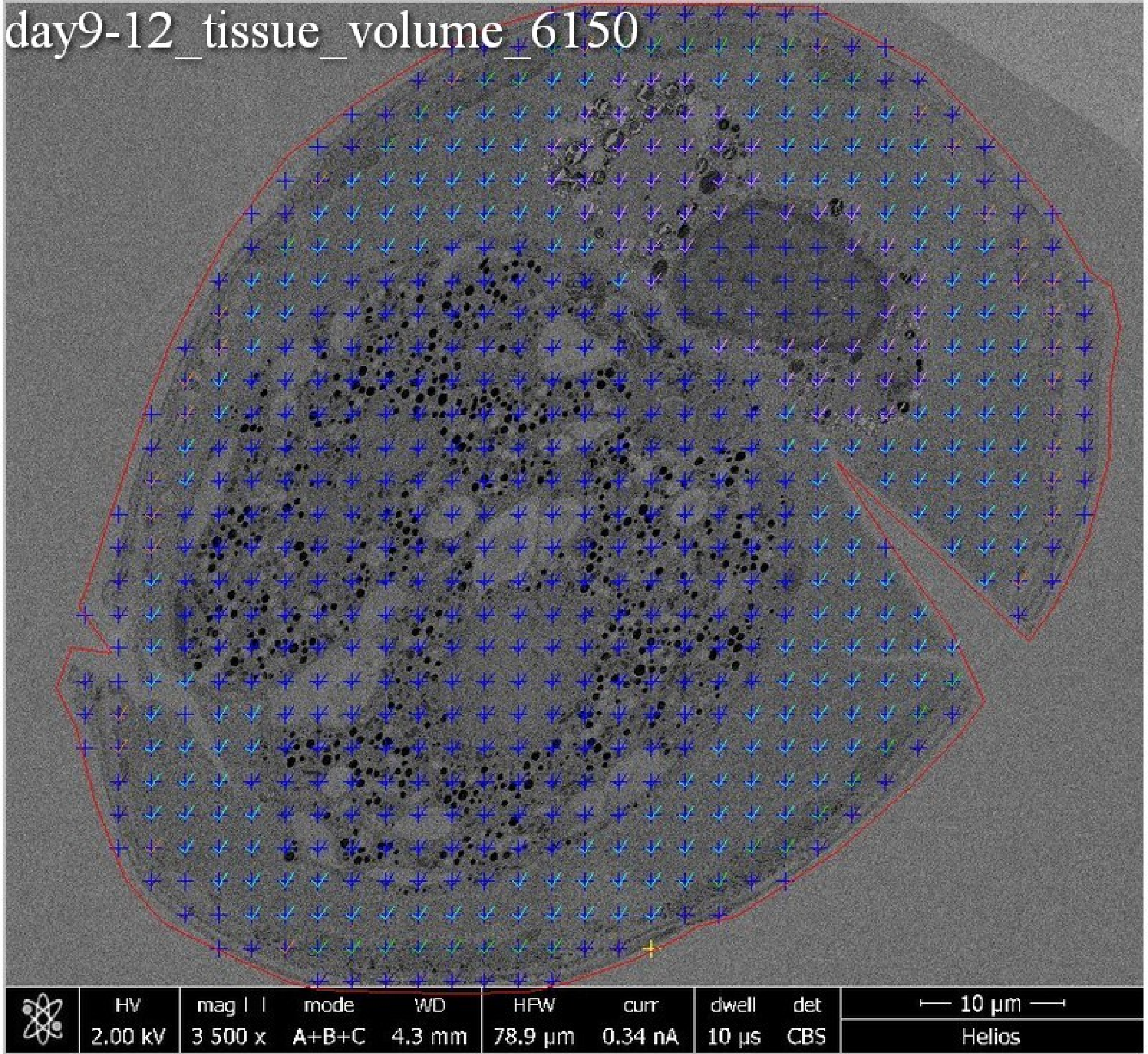
day9-12_tissue_volume_6150
HV
2.00 kV
mag
3 500 x
mode
A+B+C
WD
4.3 mm
HFW
78.9 µm
curr
0.34 nA
dwell
10 µs
det
CBS
10 µm
Helios

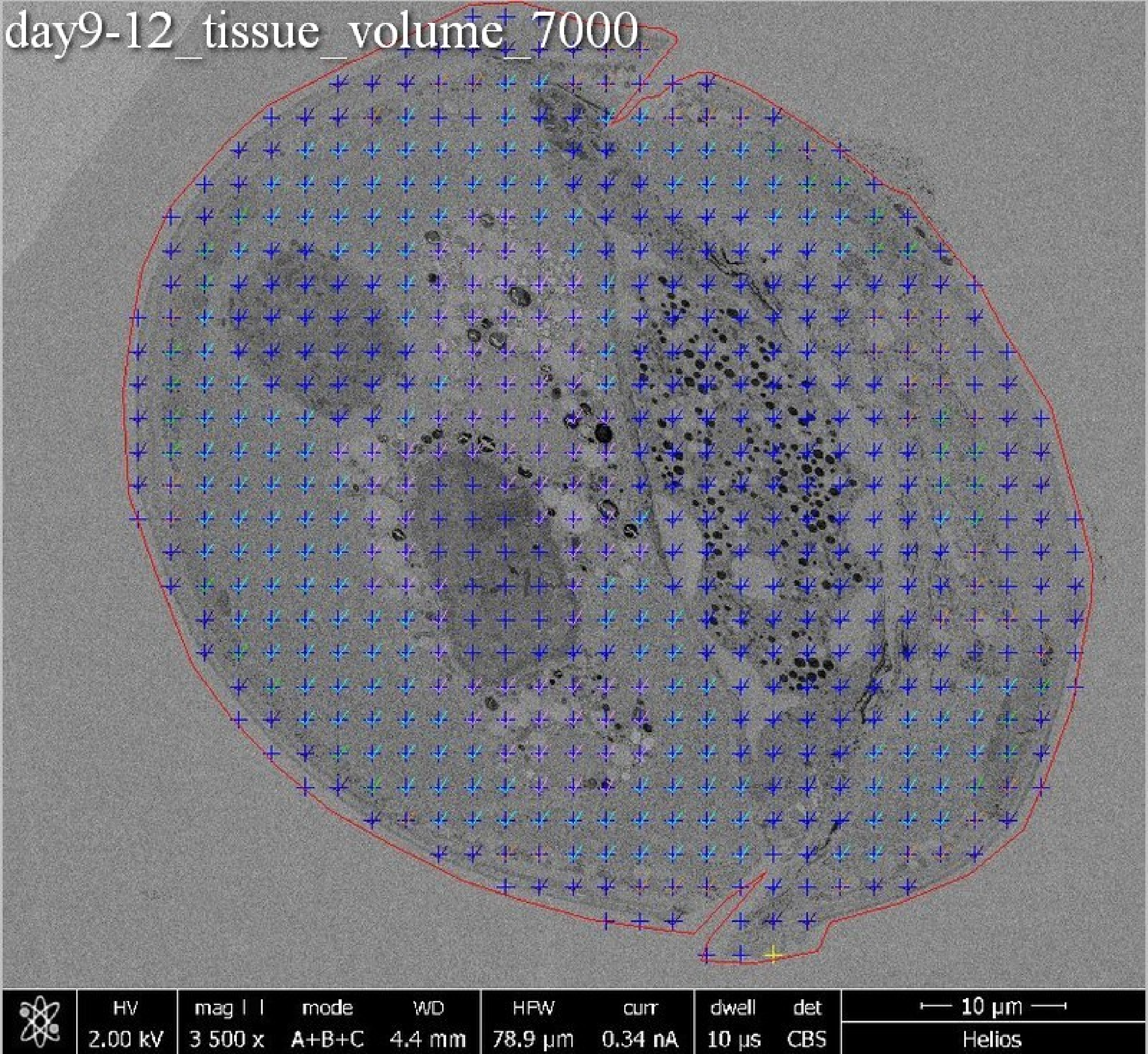
day9-12_tissue_volume_7000
HV 2.00 kV
mag 3 500 x
mode A+B+C
WD 4.4 mm
HFW 78.9 µm
curr 0.34 nA
dwell 10 µs
det CBS
10 µm
Helios

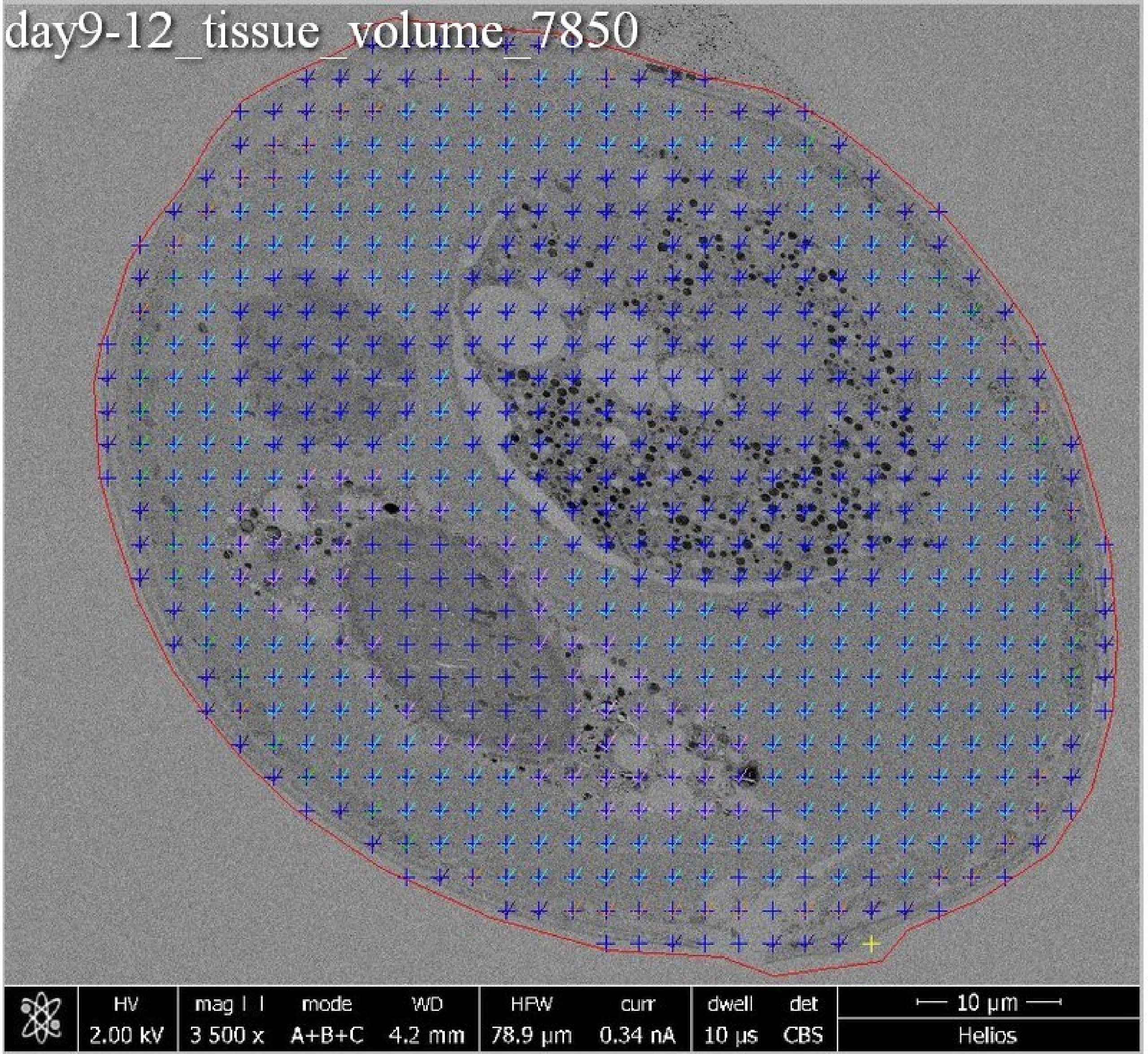
day9-12_tissue_volume_7850
HV
2.00 kV
mag
3 500 x
mode
A+B+C
WD
4.2 mm
HFW
78.9 µm
curr
0.34 nA
dwell
10 µs
det
CBS
10 µm
Helios

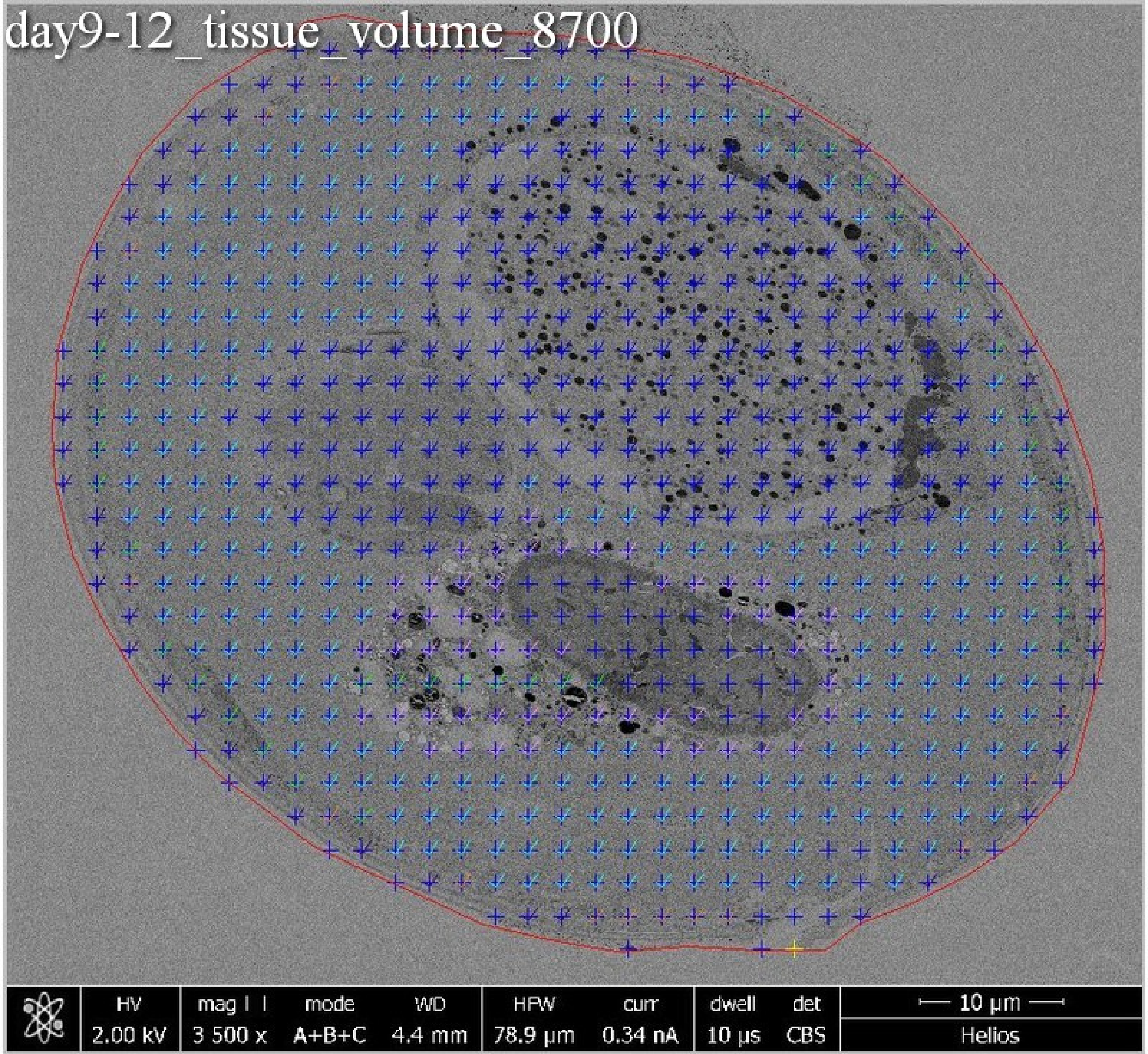
day9-12_tissue_volume_8700
HV
2.00 kV
mag
3 500 x
mode
A+B+C
WD
4.4 mm
HFW
78.9 µm
curr
0.34 nA
dwell
10 µs
det
CBS
10 µm
Helios

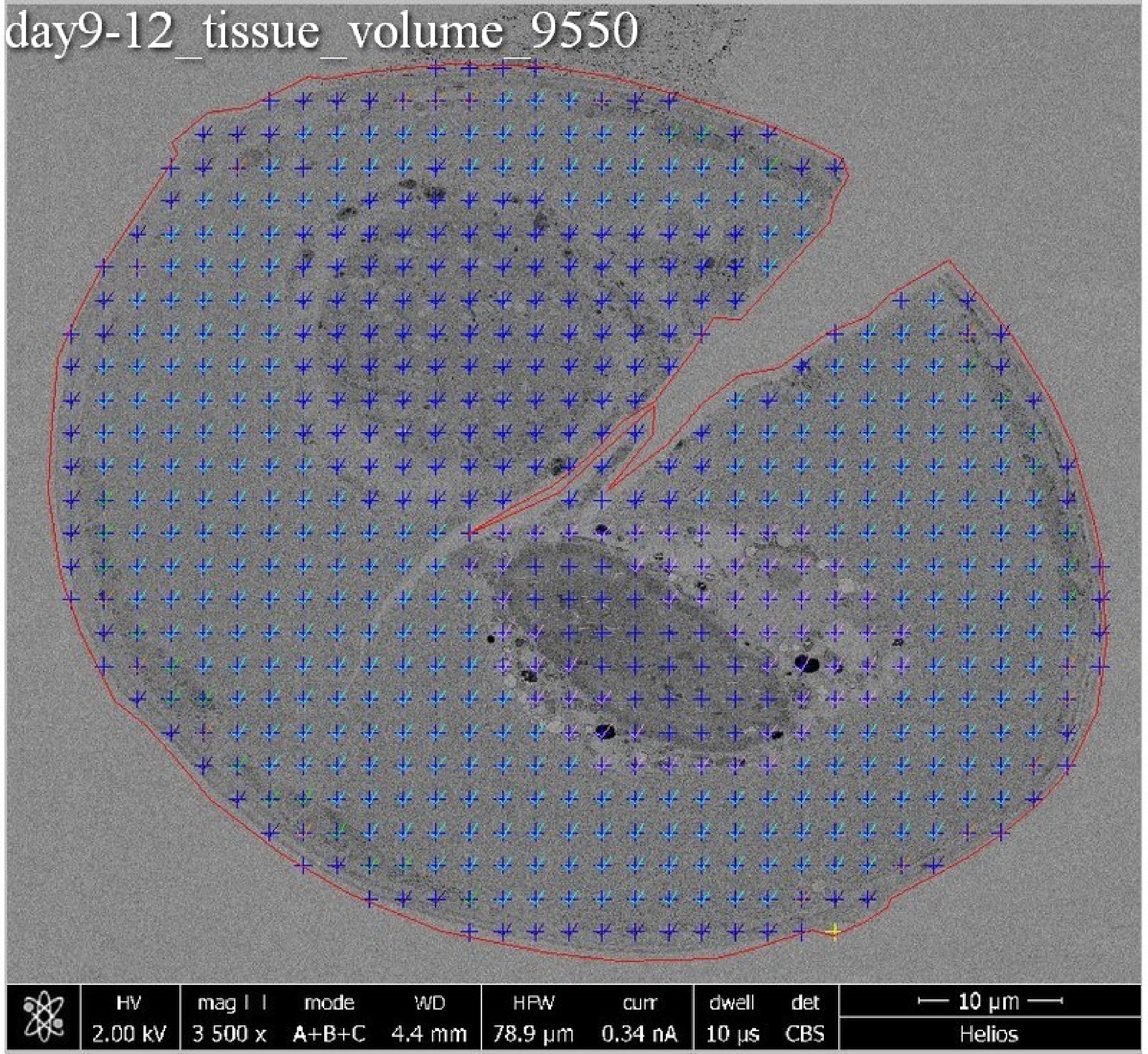
day9-12_tissue_volume_9550
HV 2.00 kV
mag 3 500 x
mode A+B+C
WD 4.4 mm
HFW 78.9 μm
curr 0.34 nA
dwell 10 μs
det CBS
10 μm
Helios

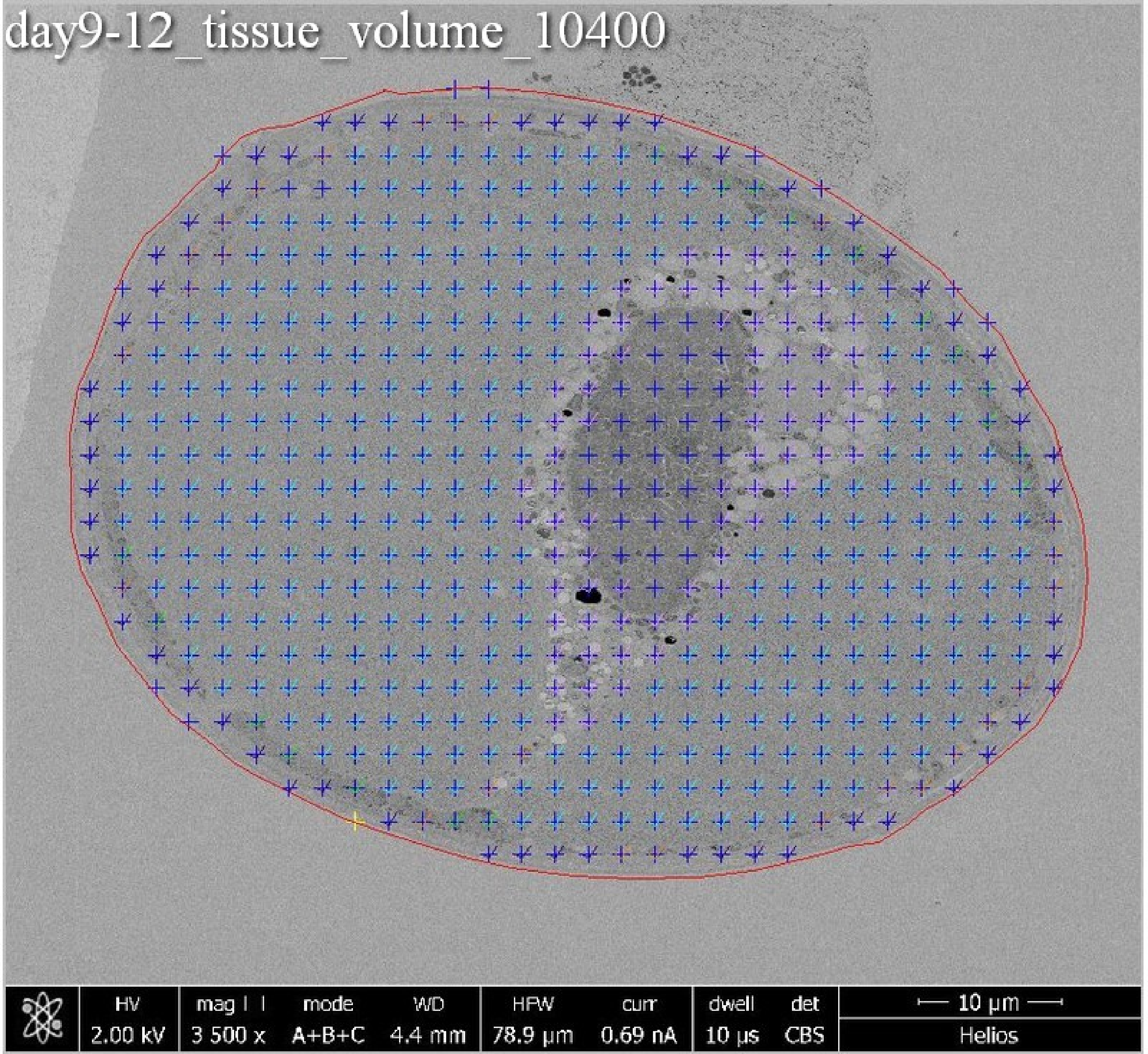
day9-12_tissue_volume_10400
HV
2.00 kV
mag
3 500 x
mode
A+B+C
WD
4.4 mm
HFW
78.9 µm
curr
0.69 nA
dwell
10 µs
det
CBS
10 µm
Helios

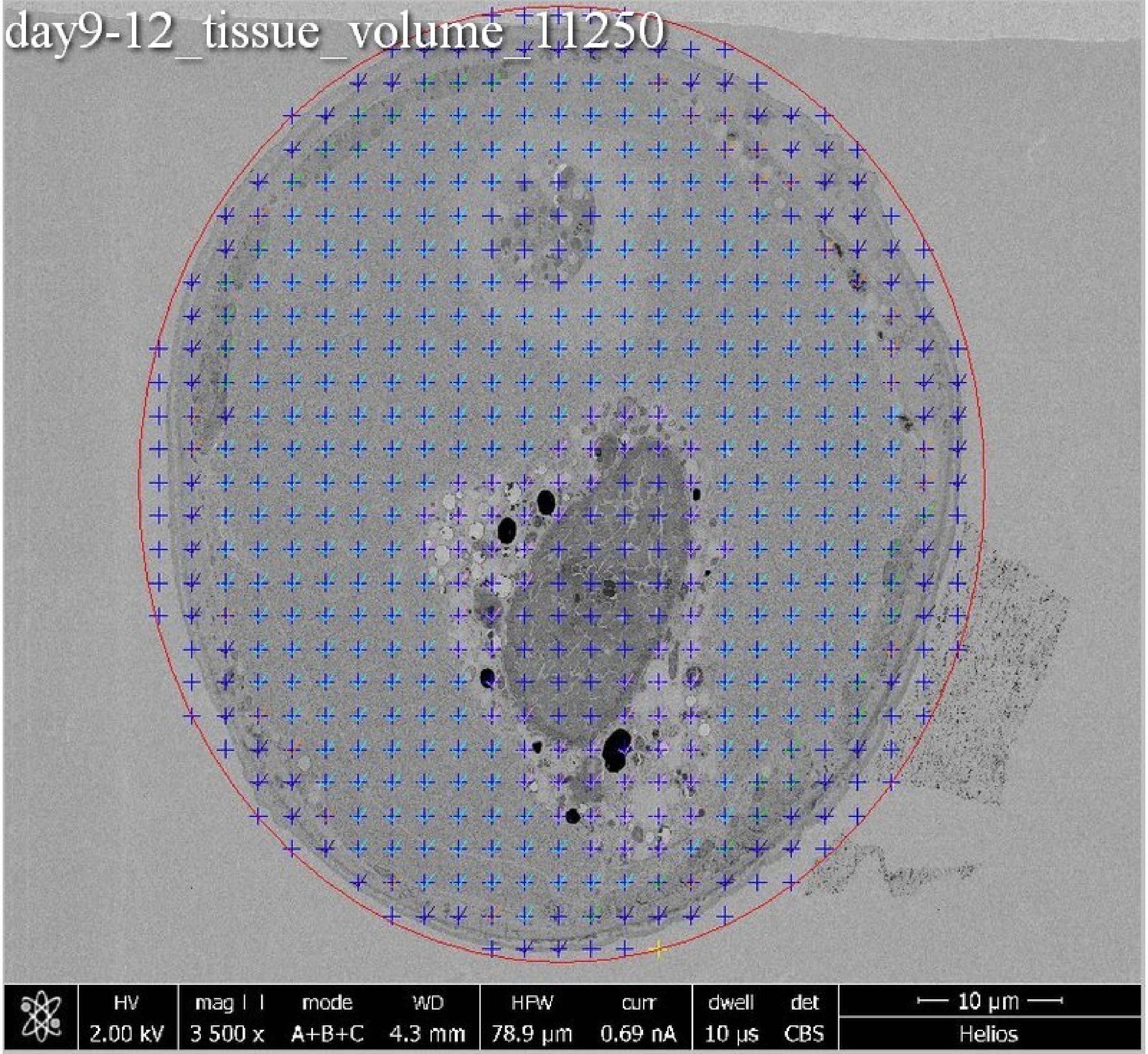
day9-12_tissue_volume_11250
HV
2.00 kV
mag
3 500 x
mode
A+B+C
WD
4.3 mm
HFW
78.9 µm
curr
0.69 nA
dwell
10 µs
det
CBS
10 µm
Helios

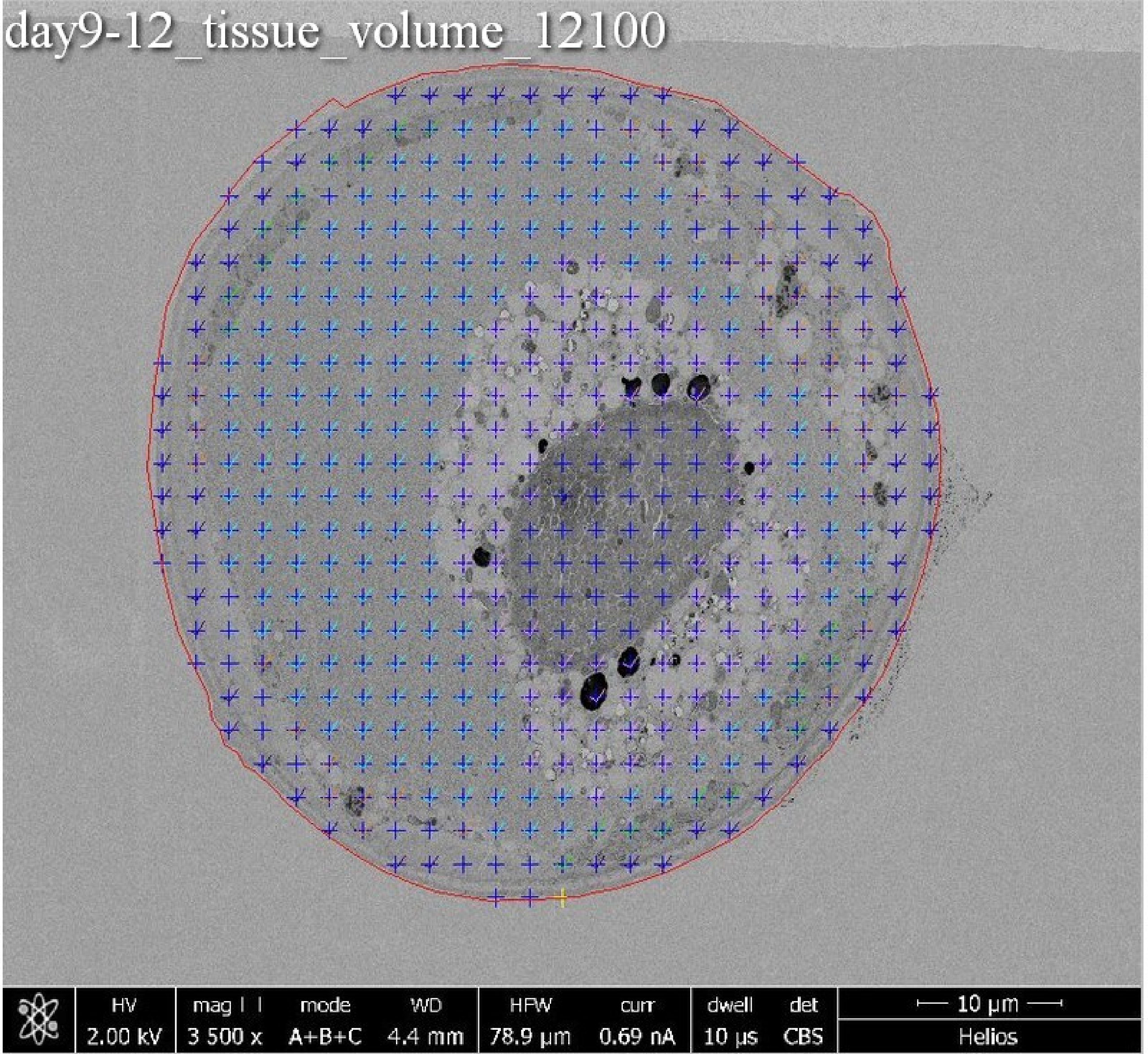
day9-12_tissue_volume_12100
HV 2.00 kV
mag ⊞ 3 500 x
mode A+B+C
WD 4.4 mm
HFW 78.9 µm
curr 0.69 nA
dwell 10 µs
det CBS
10 µm
Helios

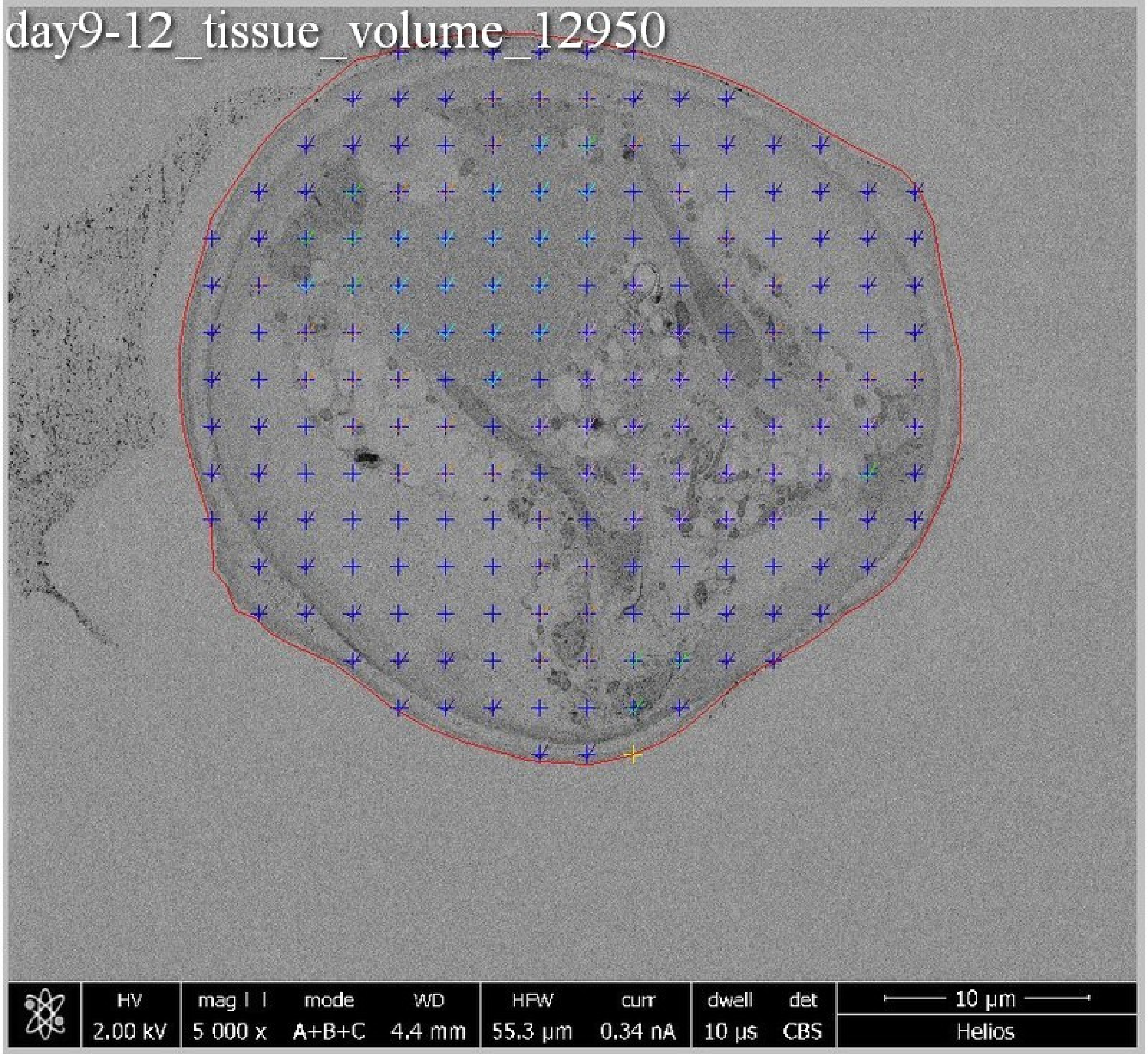

day9-12_tissue_volume_12950
HV
2.00 kV
mag
5 000 x
mode
A+B+C
WD
4.4 mm
HFW
55.3 µm
curr
0.34 nA
dwell
10 µs
det
CBS
10 µm
Helios

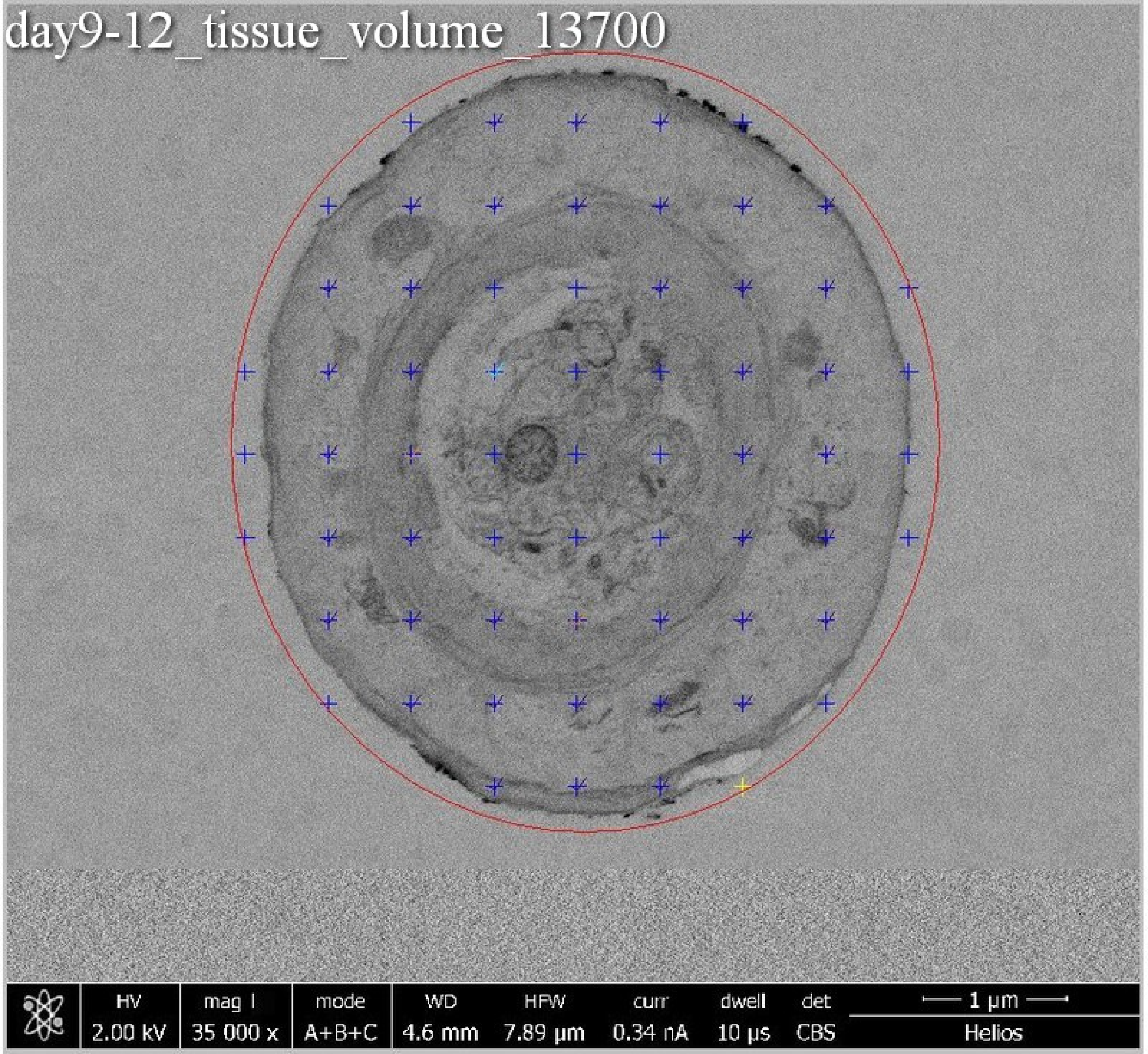
day9-12_tissue_volume_13700
HV 2.00 kV
mag ⊡ 35 000 x
mode A+B+C
WD 4.6 mm
HFW 7.89 µm
curr 0.34 nA
dwell 10 µs
det CBS
1 µm
Helios

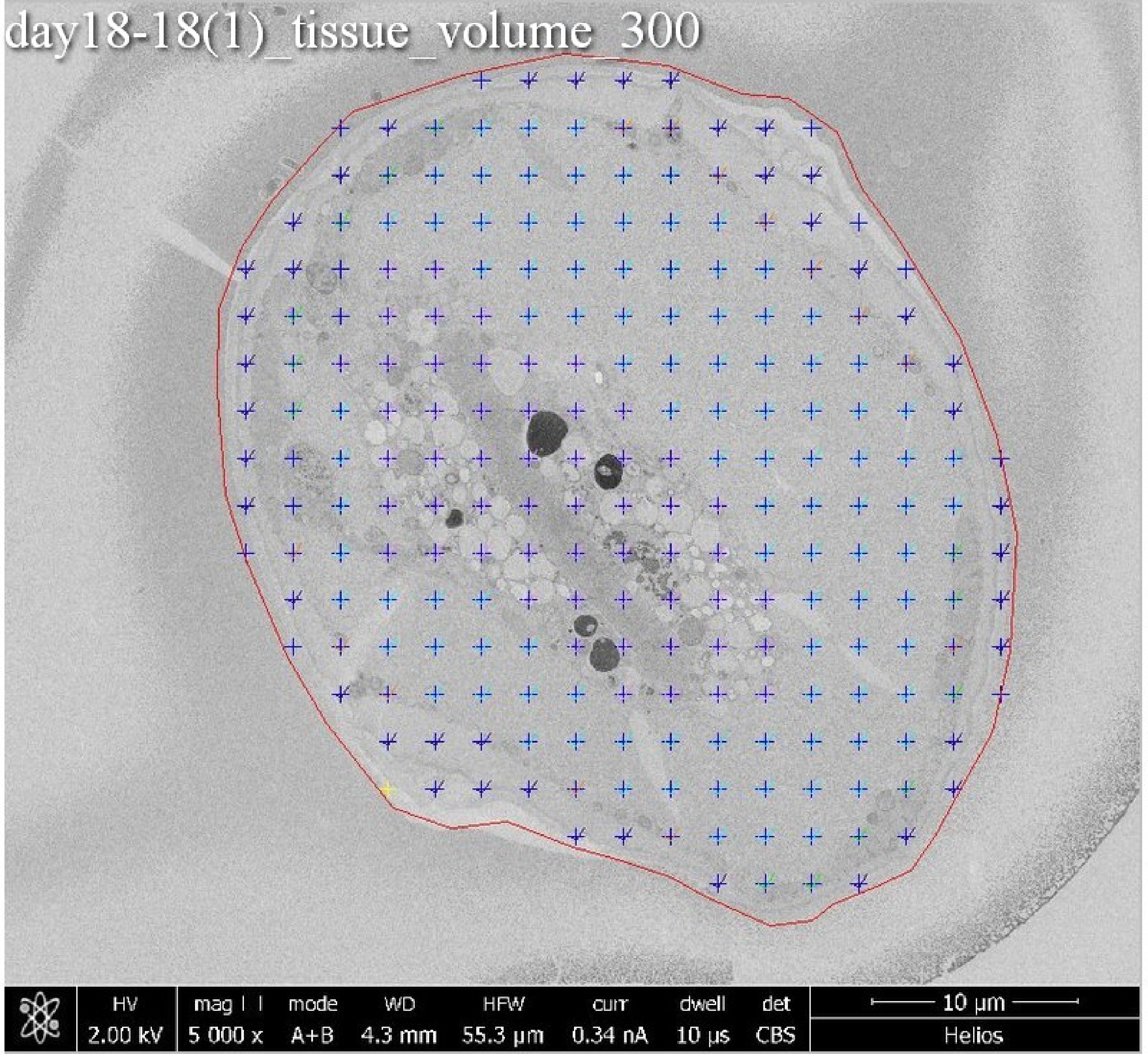
day18-18(1)_tissue_volume_300
HV
2.00 kV
mag
5 000 x
mode
A+B
WD
4.3 mm
HFW
55.3 μm
curr
0.34 nA
dwell
10 μs
det
CBS
10 μm
Helios

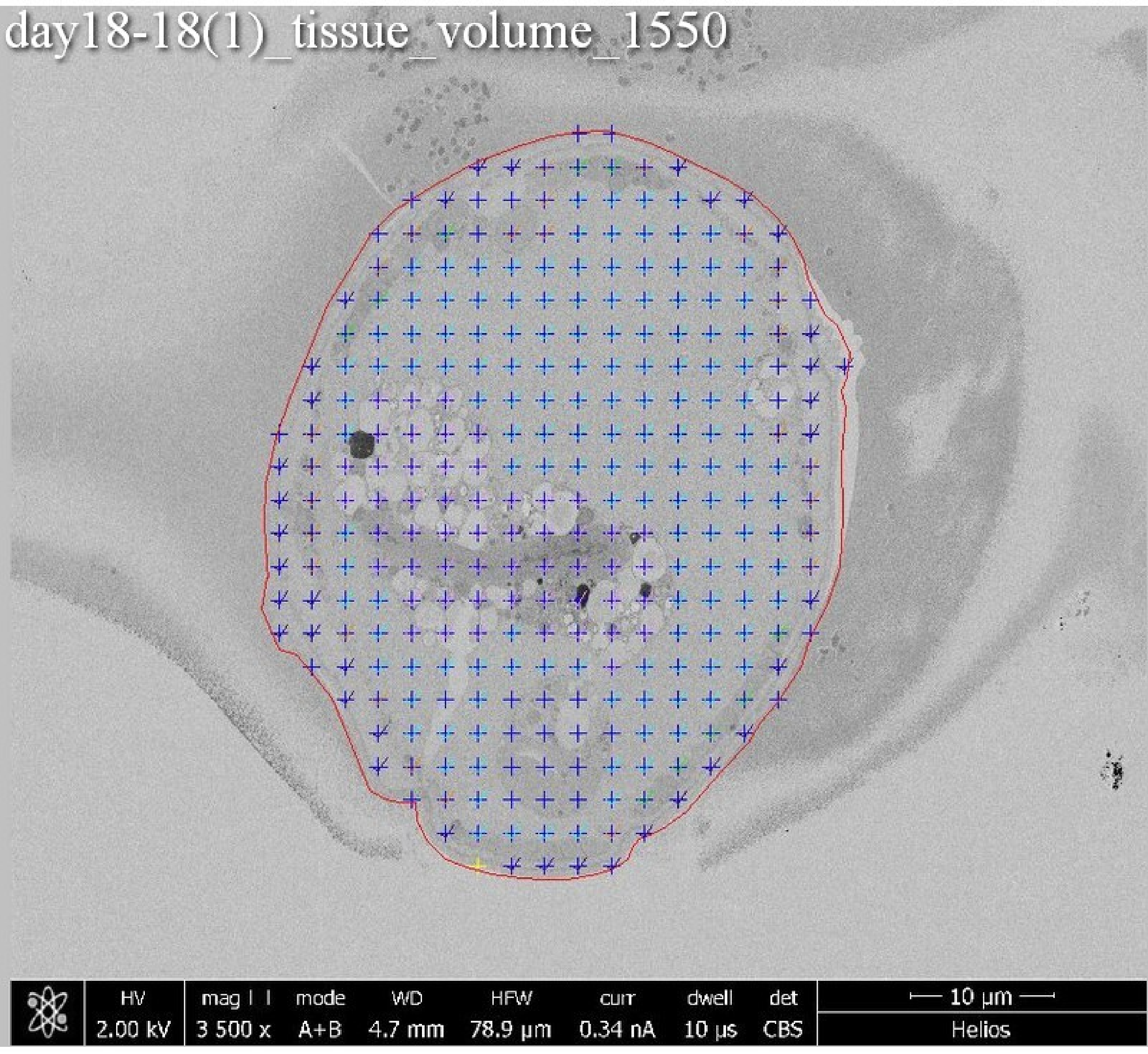
day18-18(1)_tissue_volume_1550
HV
2.00 kV
mag
3 500 x
mode
A+B
WD
4.7 mm
HFW
78.9 µm
curr
0.34 nA
dwell
10 µs
det
CBS
10 µm
Helios

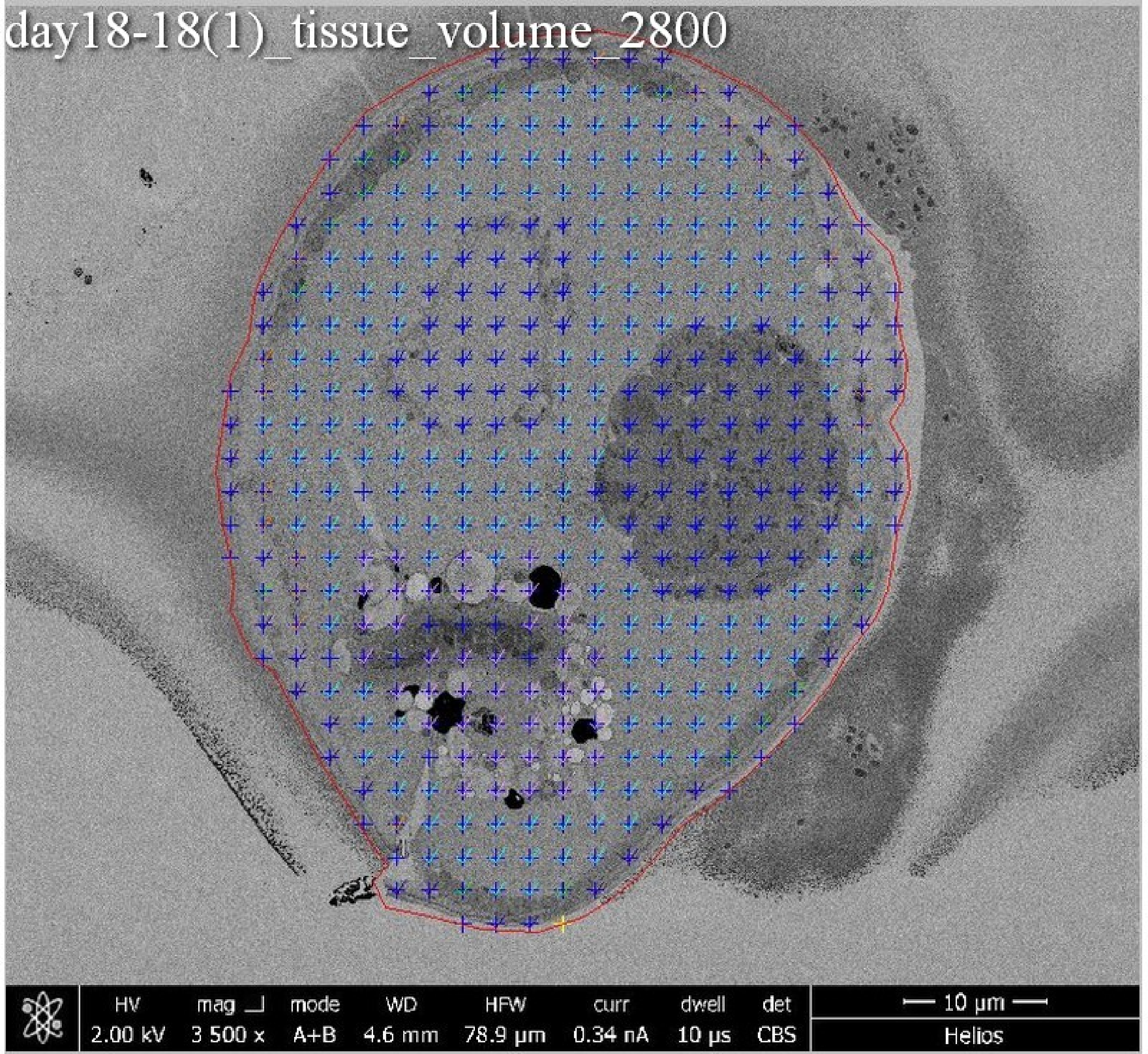
day18-18(1)_tissue_volume_2800
HV
2.00 kV
mag
3 500 x
mode
A+B
WD
4.6 mm
HFW
78.9 μm
curr
0.34 nA
dwell
10 μs
det
CBS
10 μm
Helios

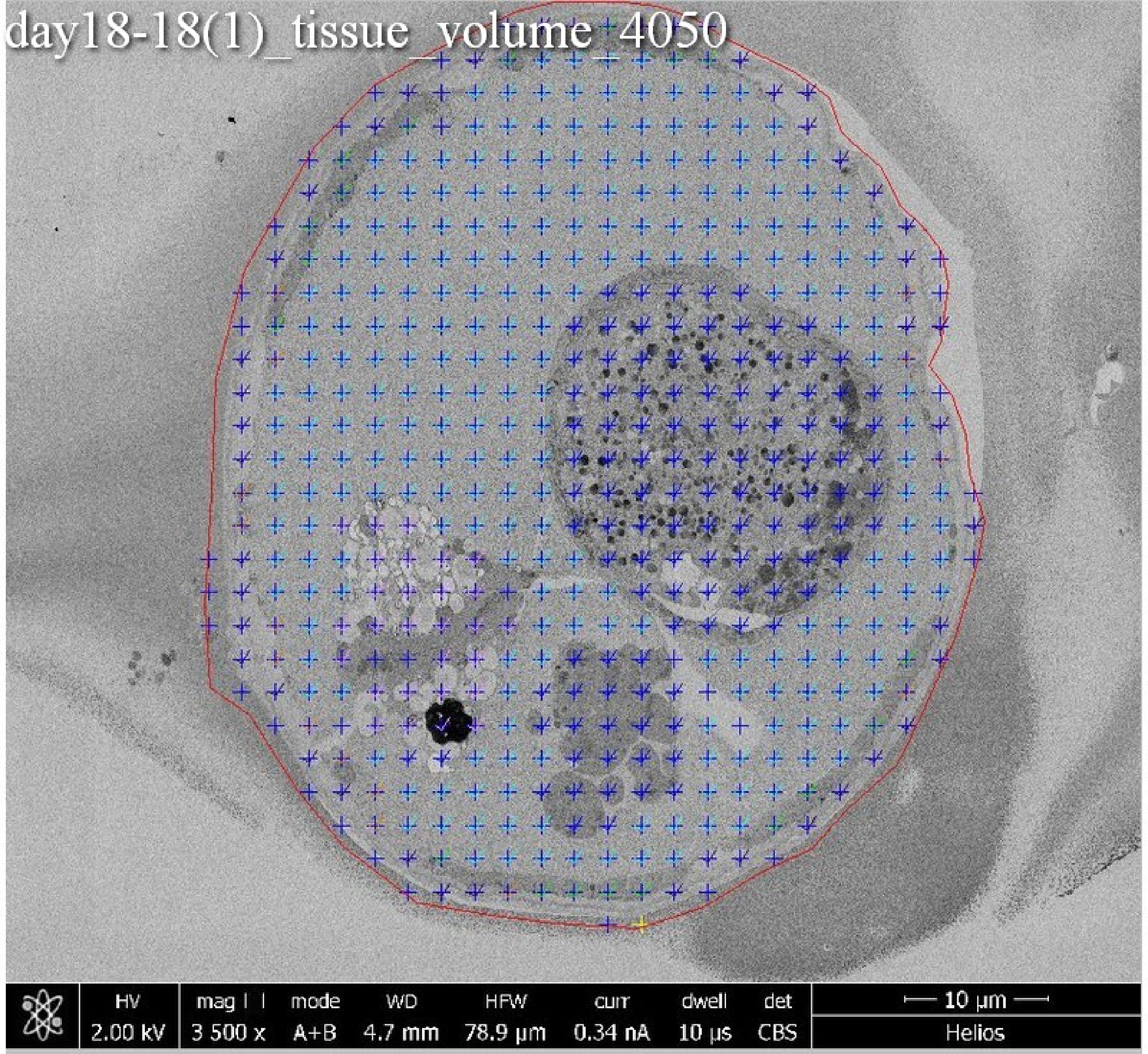
day18-18(1)_tissue_volume_4050
HV
2.00 kV
mag
3 500 x
mode
A+B
WD
4.7 mm
HFW
78.9 µm
curr
0.34 nA
dwell
10 µs
det
CBS
10 µm
Helios

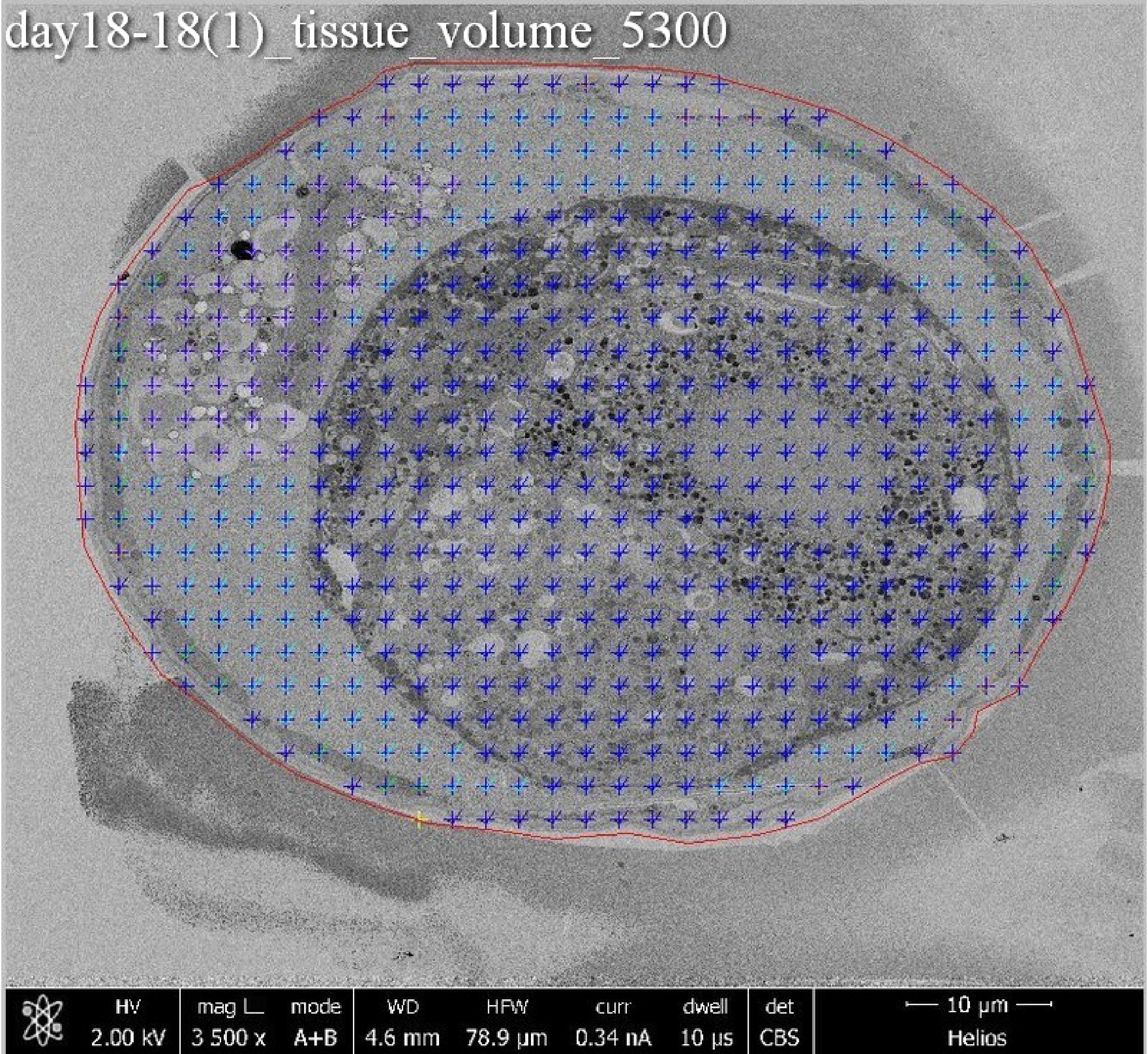
day18-18(1)_tissue_volume_5300
HV
2.00 kV
mag L
3 500 x
mode
A+B
WD
4.6 mm
HFW
78.9 µm
curr
0.34 nA
dwell
10 µs
det
CBS
10 µm
Helios

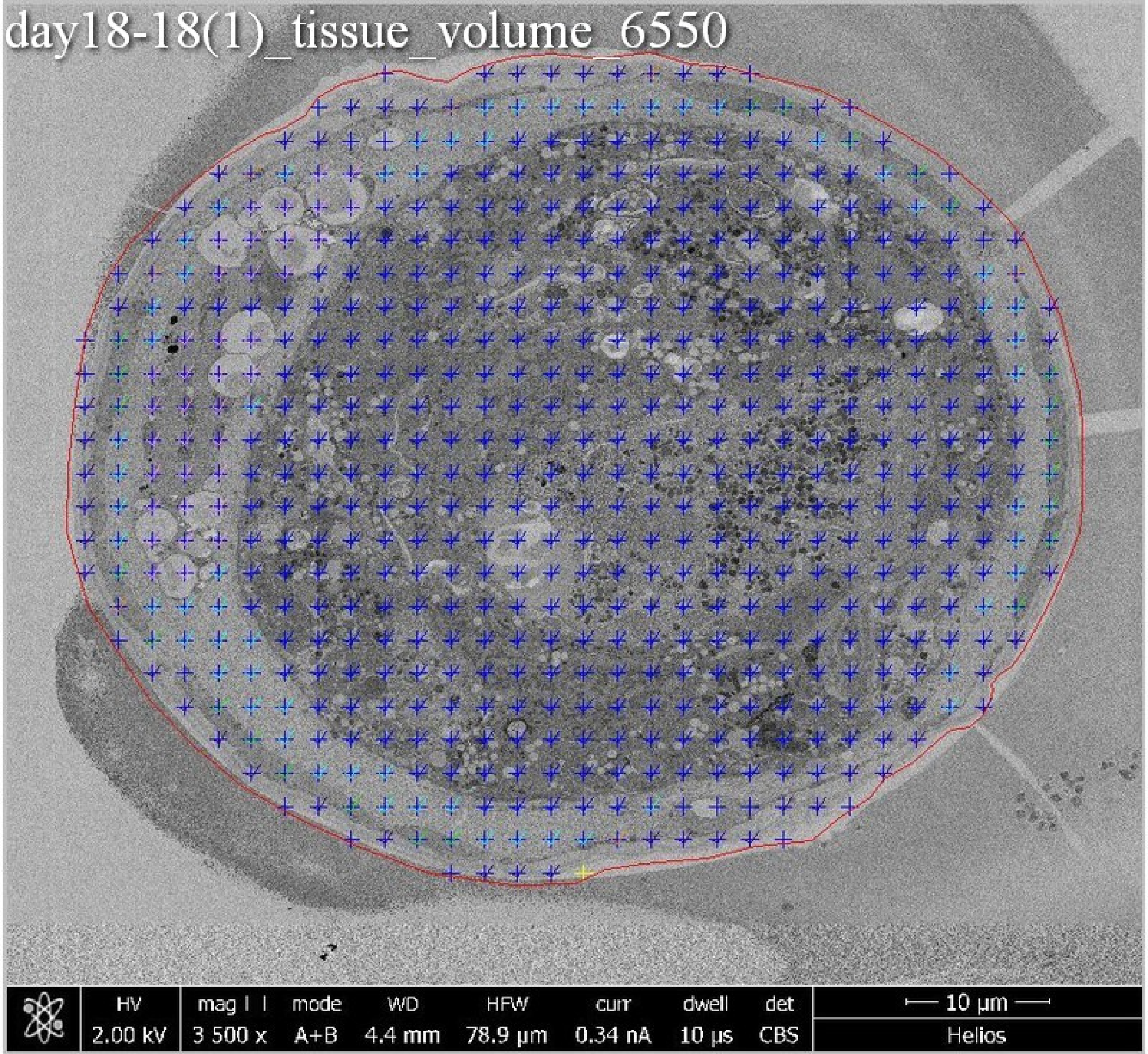
day18-18(1)_tissue_volume_6550
HV
2.00 kV
mag
3 500 x
mode
A+B
WD
4.4 mm
HFW
78.9 µm
curr
0.34 nA
dwell
10 µs
det
CBS
10 µm
Helios

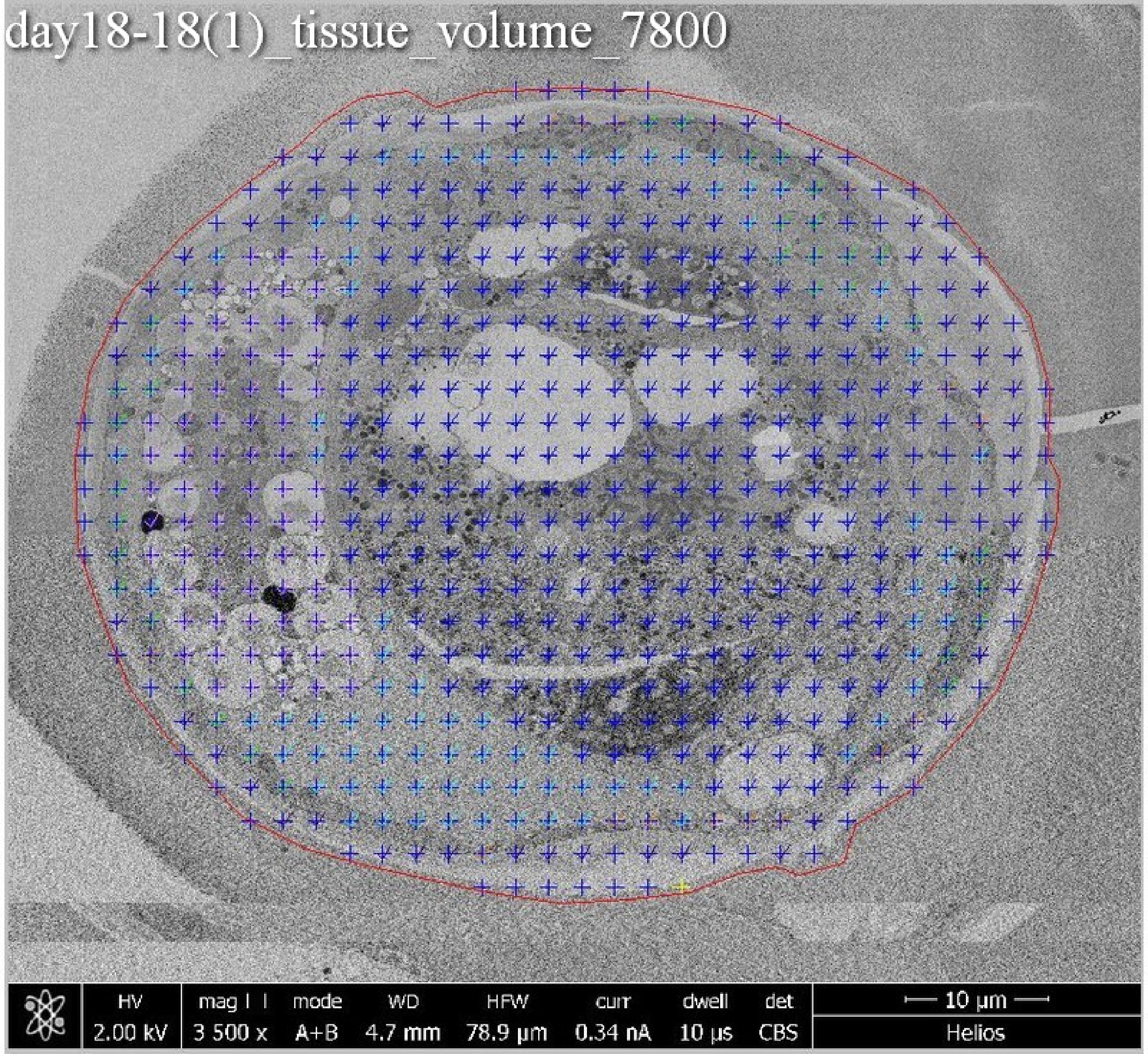

day18-18(1)_tissue_volume_7800
HV
2.00 kV
mag ⊞
3 500 x
mode
A+B
WD
4.7 mm
HFW
78.9 µm
curr
0.34 nA
dwell
10 µs
det
CBS
10 µm
Helios

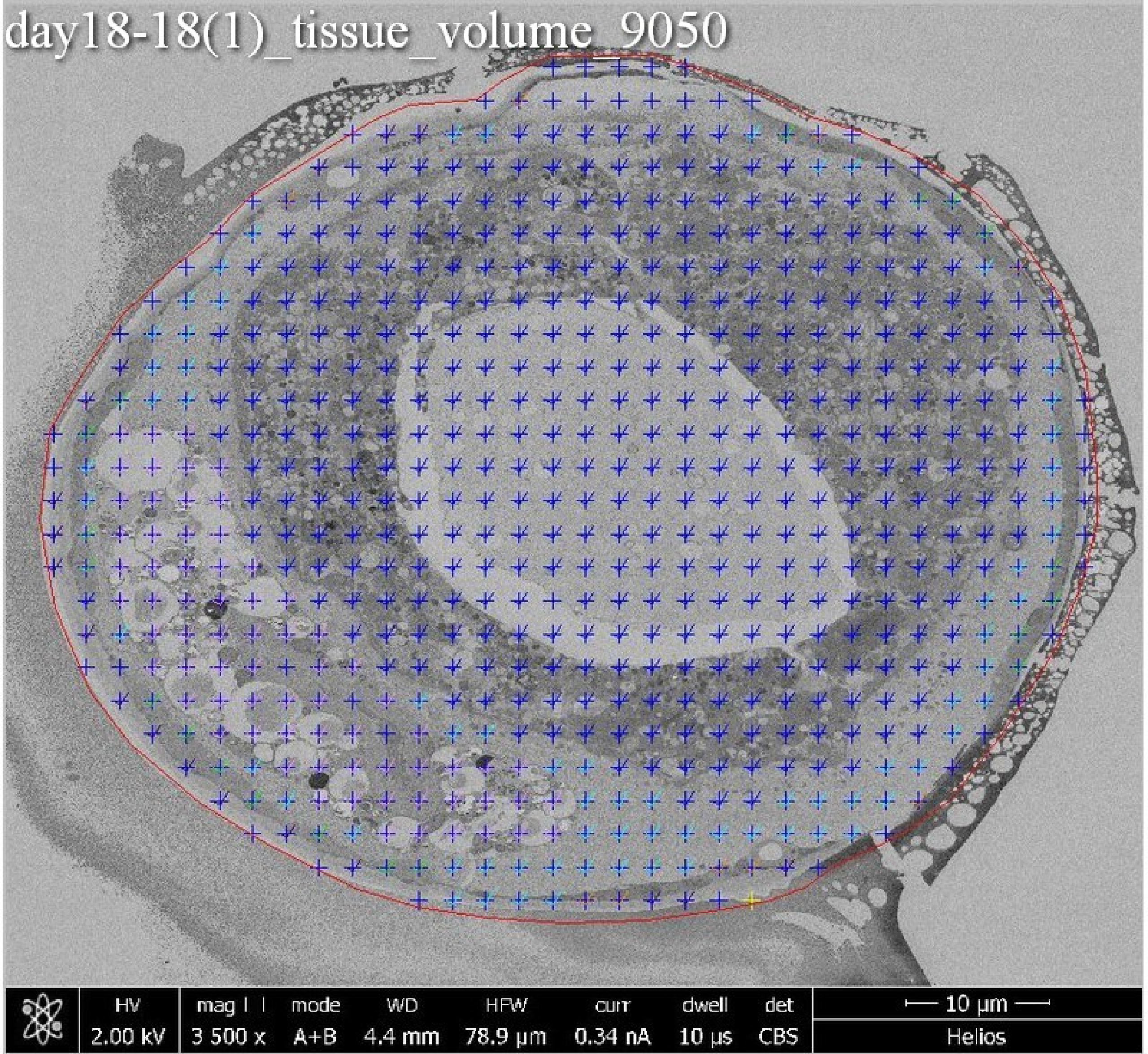
day18-18(1)_tissue_volume_9050
HV
2.00 kV
mag
3 500 x
mode
A+B
WD
4.4 mm
HFW
78.9 µm
curr
0.34 nA
dwell
10 µs
det
CBS
10 µm
Helios

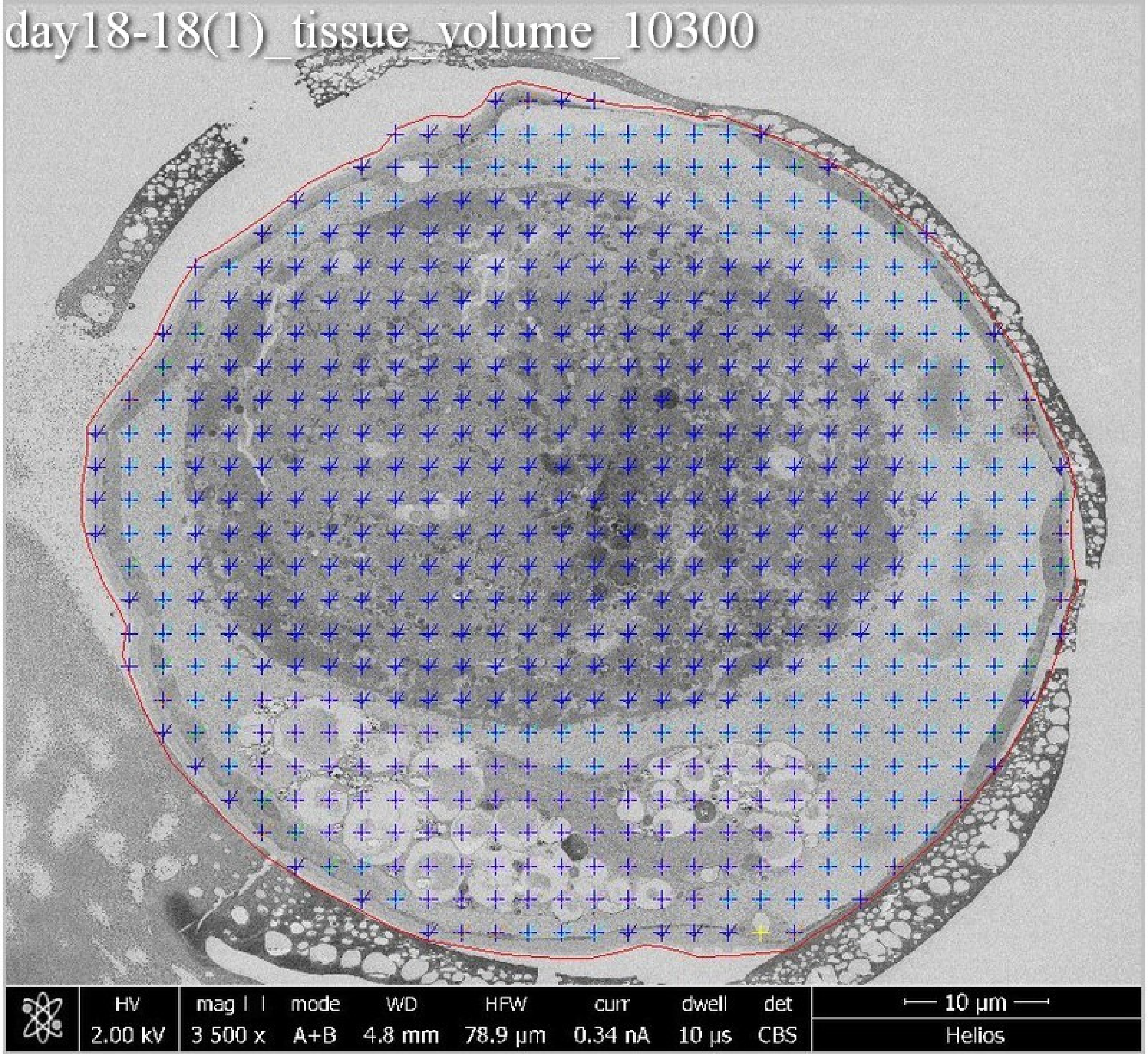
day18-18(1)_tissue_volume_10300
HV
2.00 kV
mag
3 500 x
mode
A+B
WD
4.8 mm
HFW
78.9 µm
curr
0.34 nA
dwell
10 µs
det
CBS
10 µm
Helios

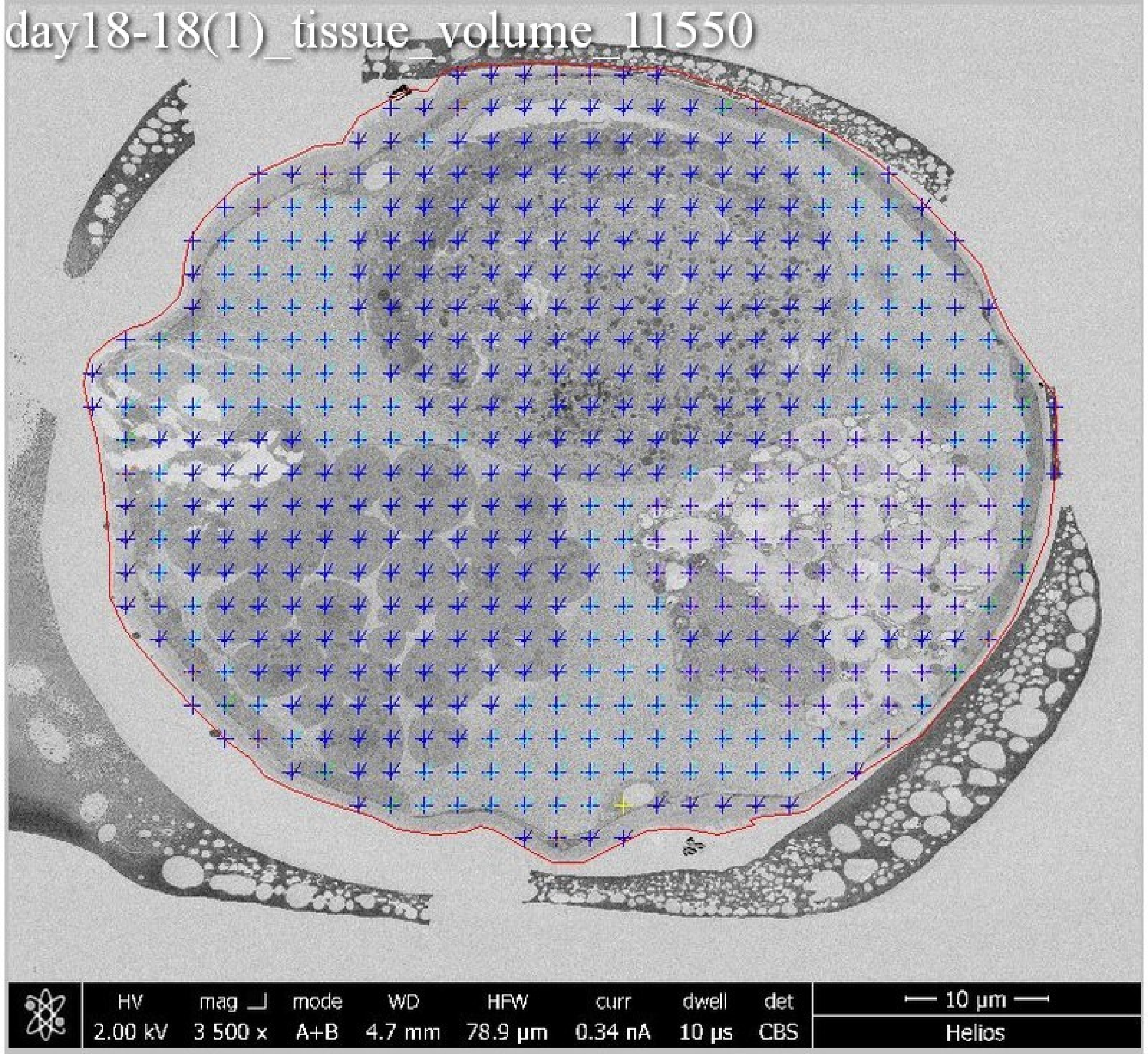
day18-18(1)_tissue_volume_11550
HV
2.00 kV
mag
3 500 x
mode
A+B
WD
4.7 mm
HFW
78.9 µm
curr
0.34 nA
dwell
10 µs
det
CBS
10 µm
Helios

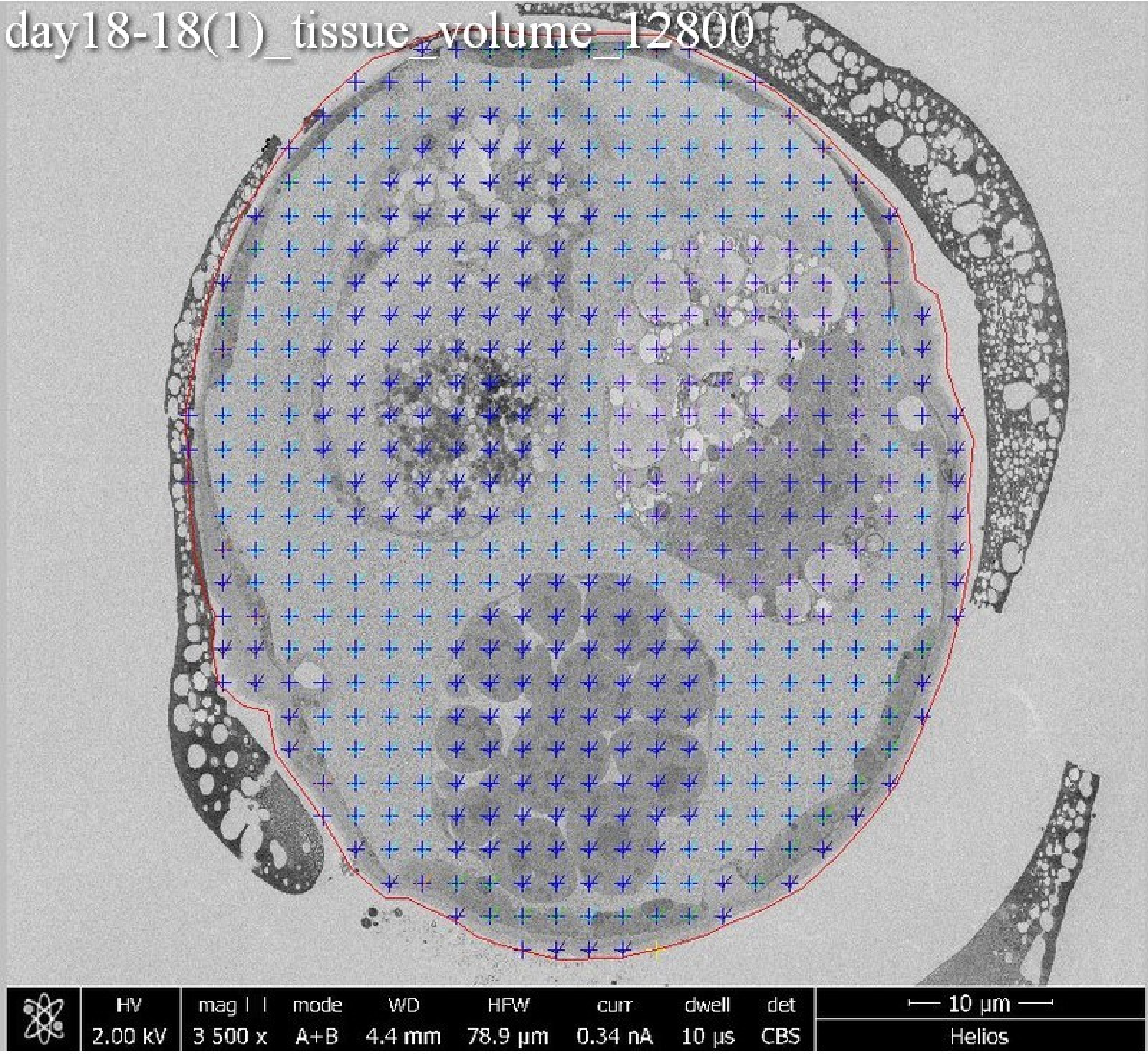
day18-18(1)_tissue_volume_12800
HV 2.00 kV
mag 3 500 x
mode A+B
WD 4.4 mm
HFW 78.9 µm
curr 0.34 nA
dwell 10 µs
det CBS
10 µm
Helios

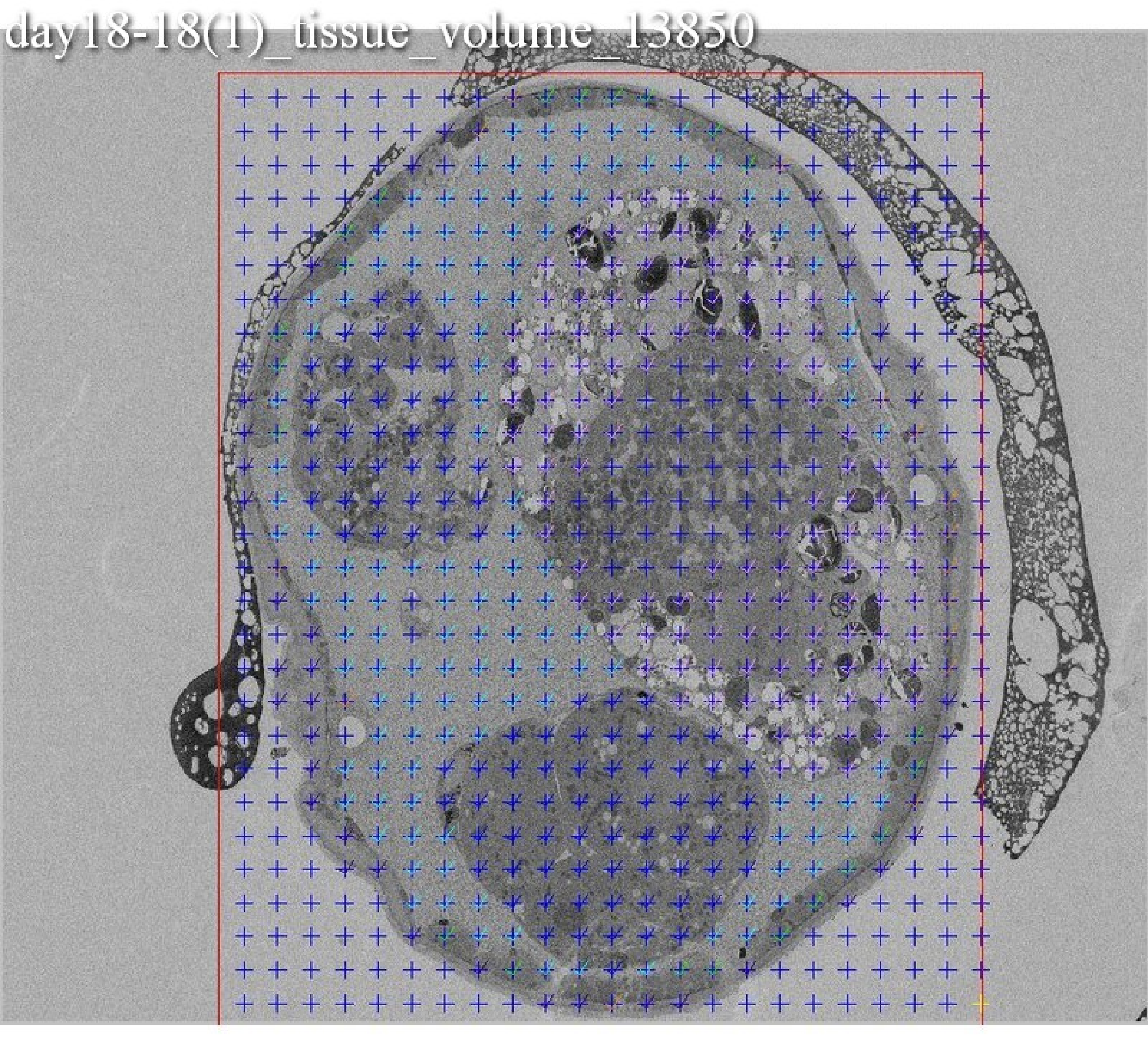
day18-18(1)_tissue_volume_13850

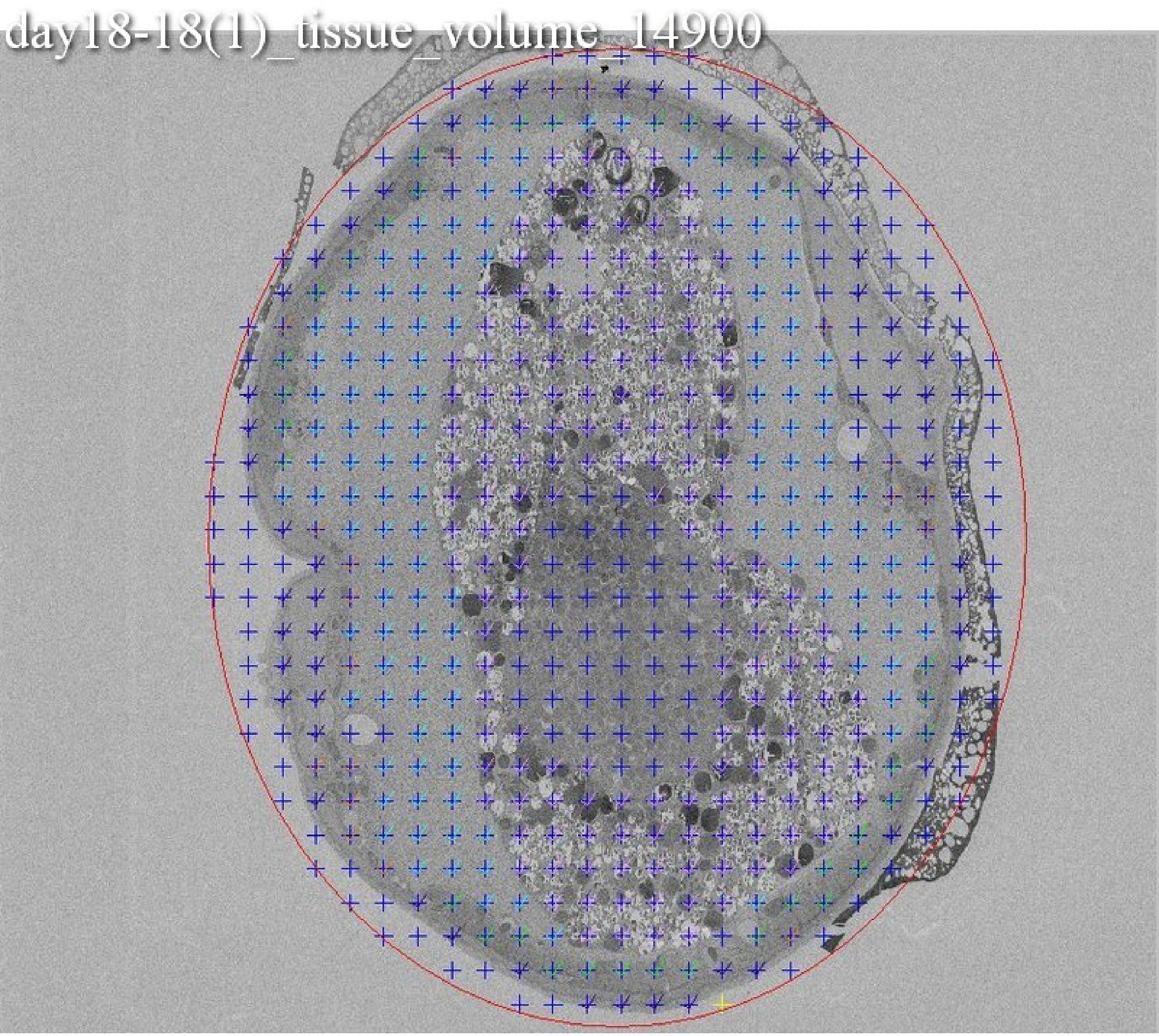
day18-18(1)_tissue_volume_14900

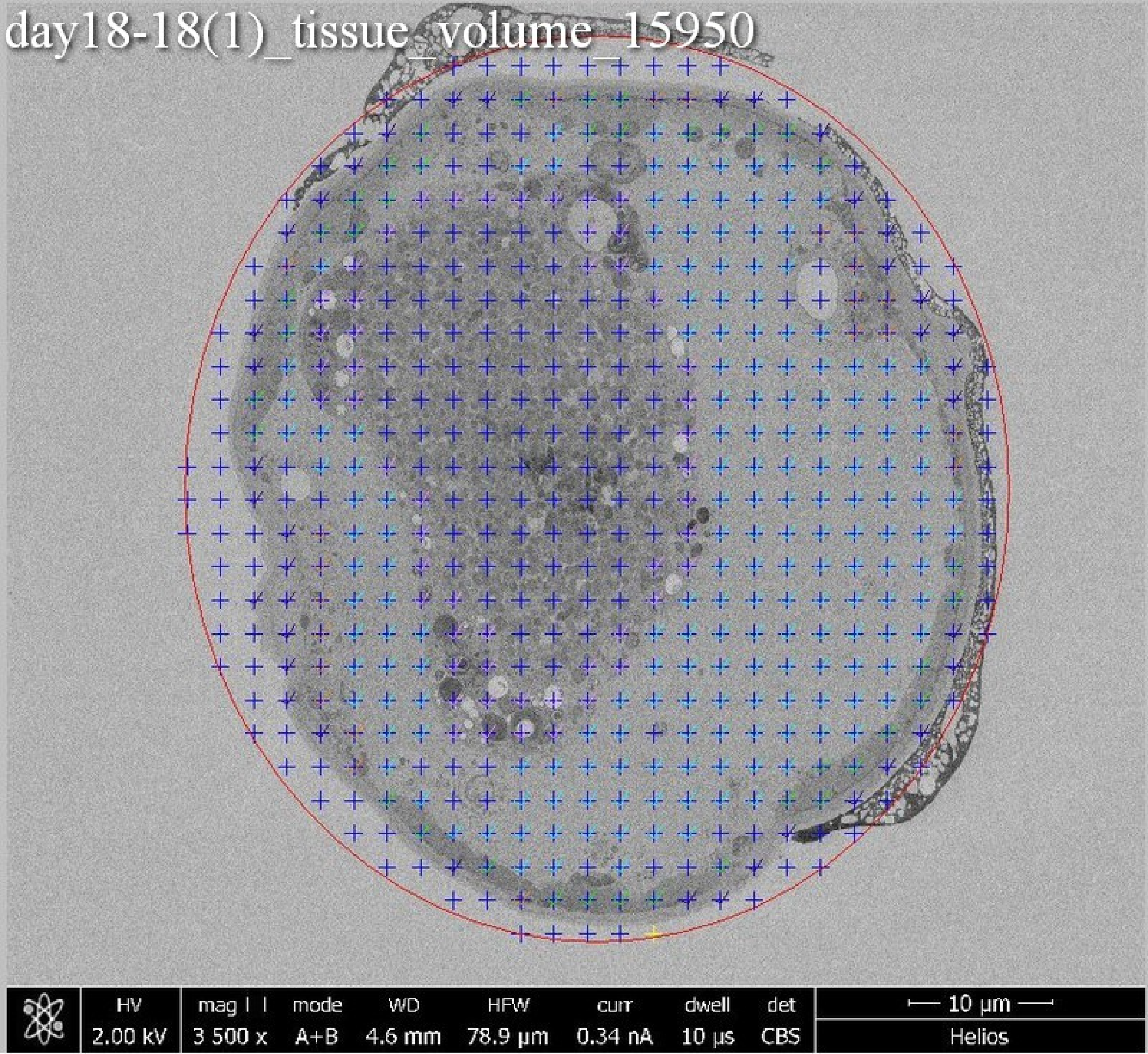
day18-18(1)_tissue_volume_15950
HV 2.00 kV
mag 3 500 x
mode A+B
WD 4.6 mm
HFW 78.9 μm
curr 0.34 nA
dwell 10 μs
det CBS
10 μm
Helios

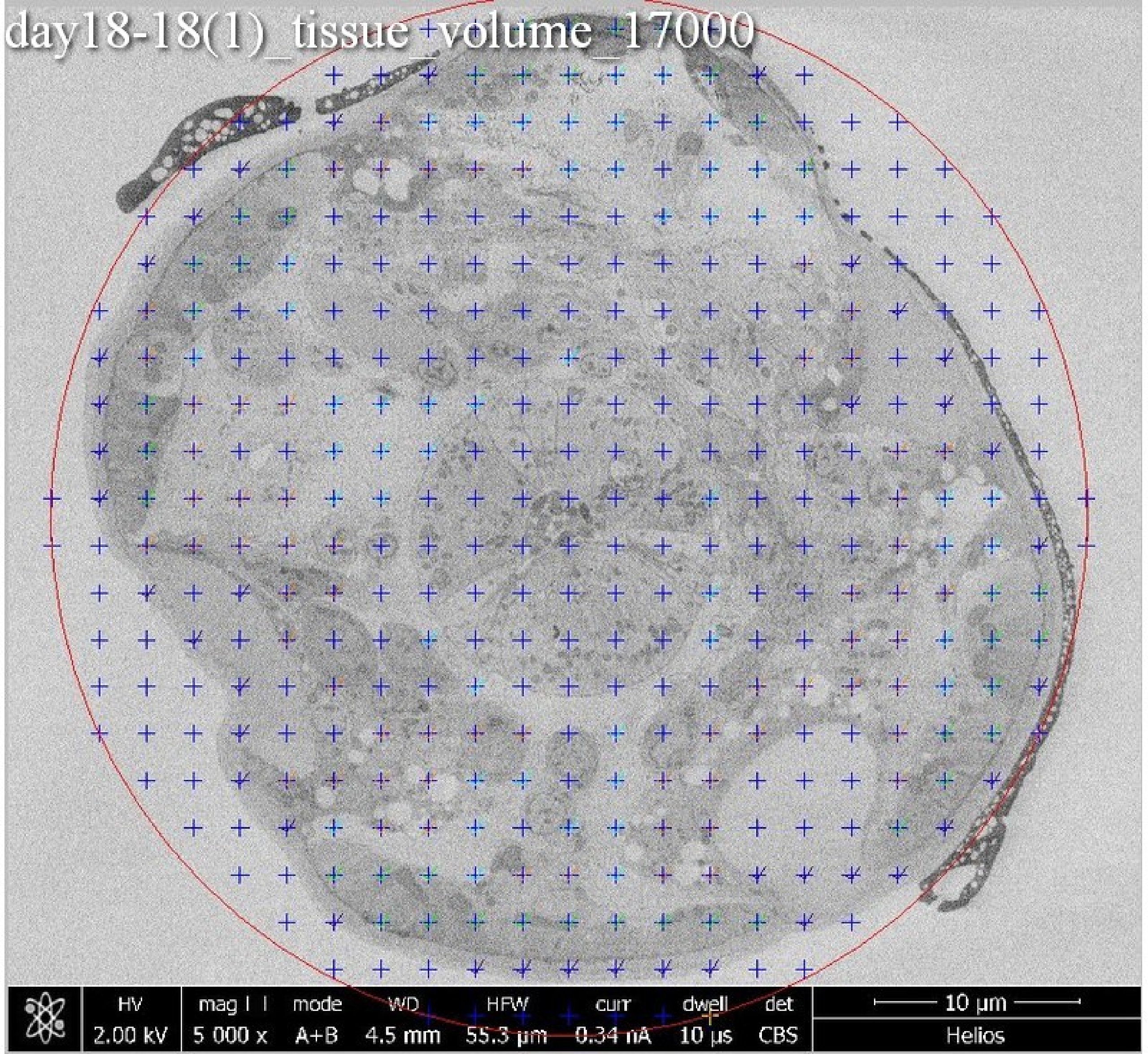
day18-18(1)_tissue_volume_17000
HV
2.00 kV
mag ⊡
5 000 x
mode
A+B
WD
4.5 mm
HFW
55.3 µm
curr
0.34 nA
dwell
10 µs
det
CBS
10 µm
Helios

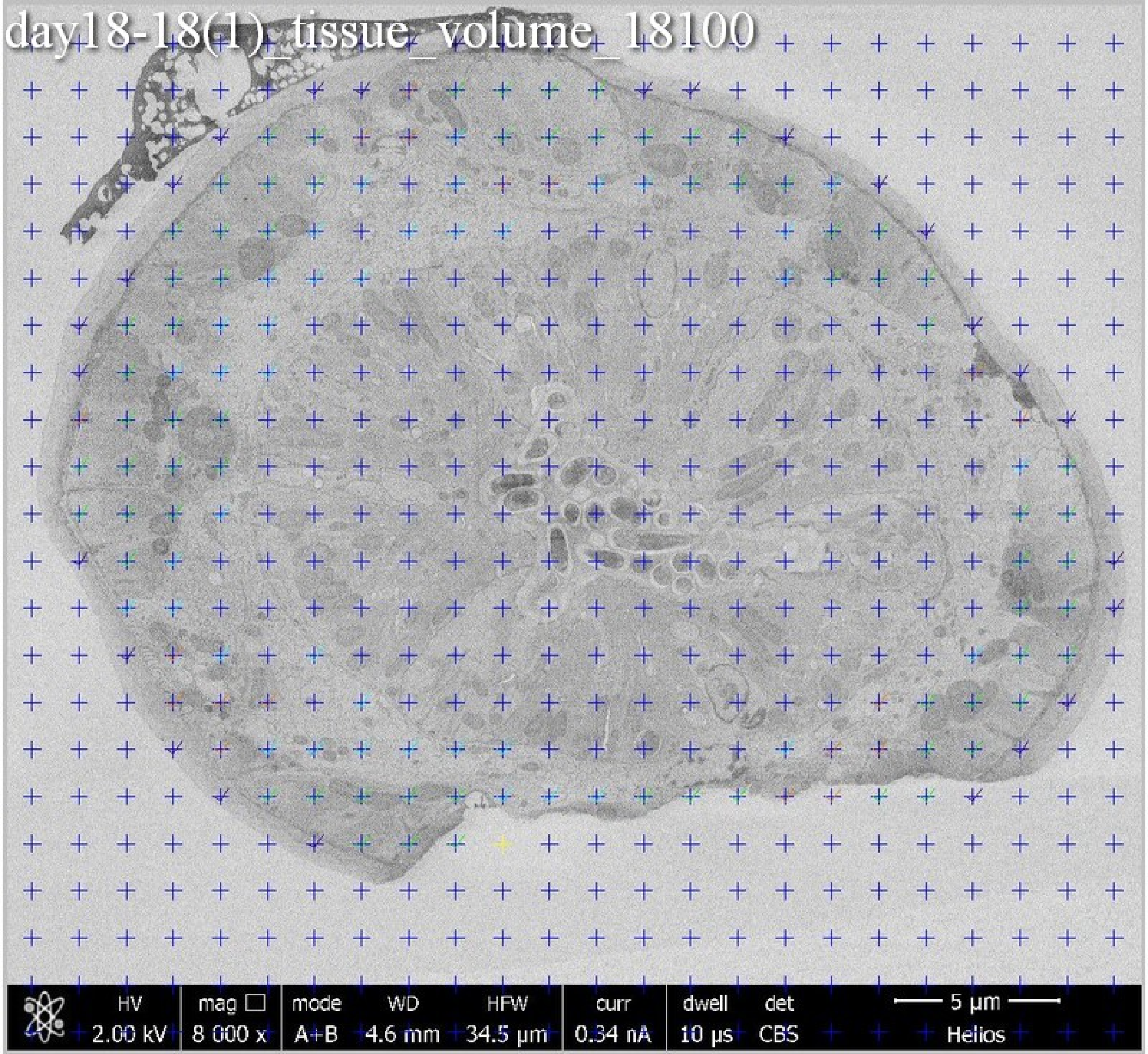
day18-18(1)_tissue_volume_18100
HV
2.00 kV
mag
8 000 x
mode
A+B
WD
4.6 mm
HFW
34.5 µm
curr
0.34 nA
dwell
10 µs
det
CBS
5 µm
Helios

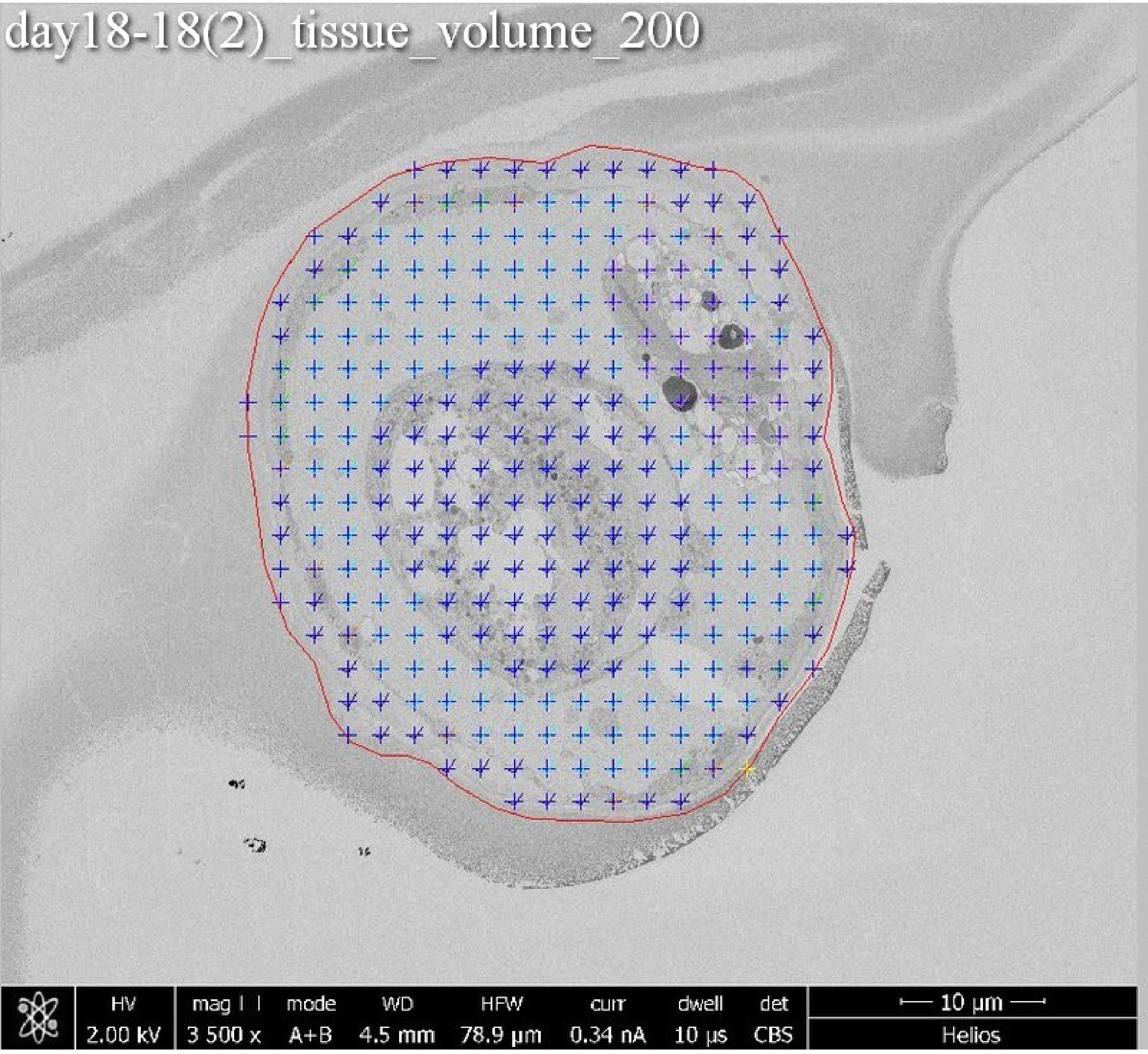
day18-18(2)_tissue_volume_200
HV
2.00 kV
mag
3 500 x
mode
A+B
WD
4.5 mm
HFW
78.9 µm
curr
0.34 nA
dwell
10 µs
det
CBS
10 µm
Helios

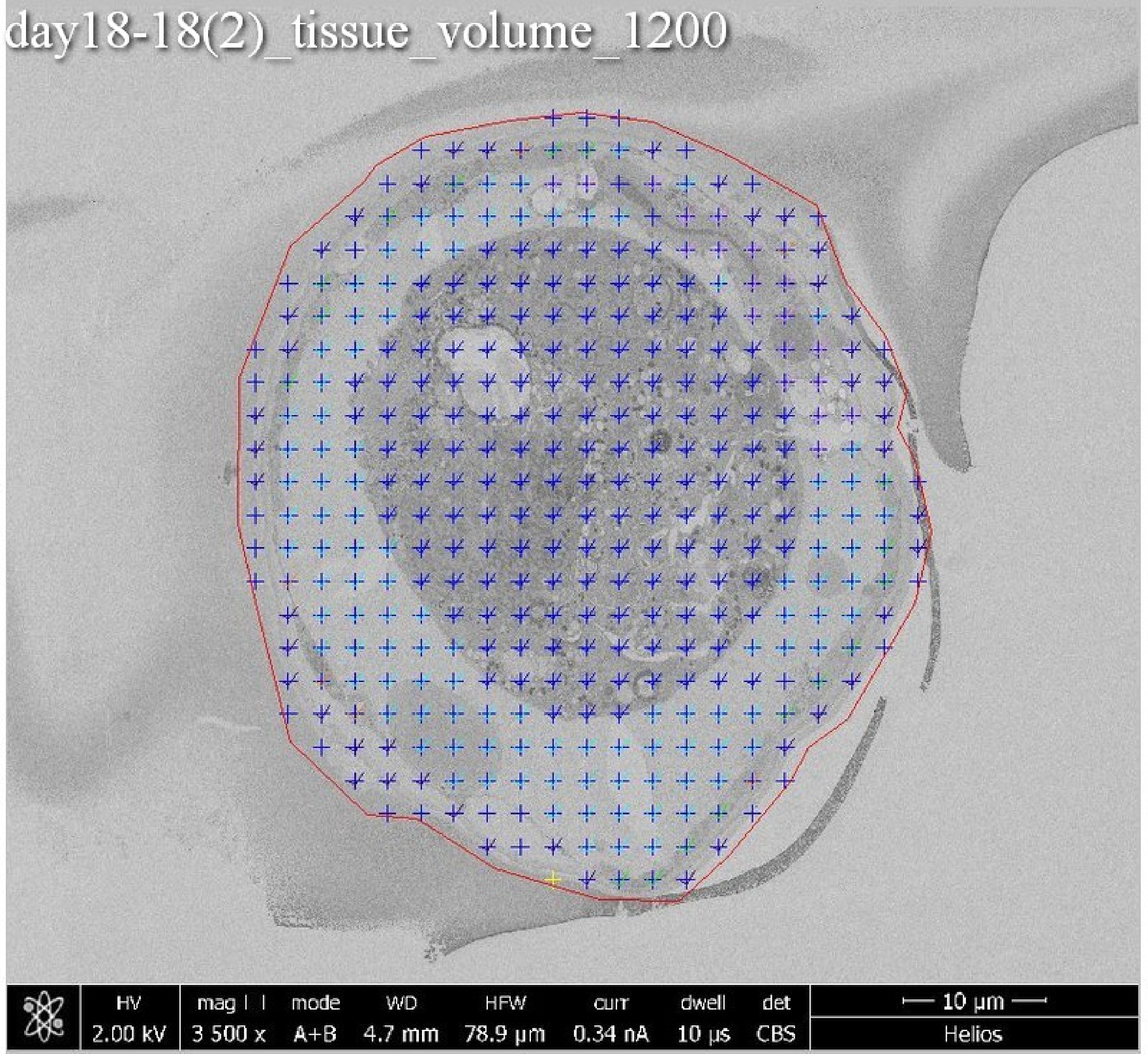
day18-18(2)_tissue_volume_1200
HV 2.00 kV
mag 3 500 x
mode A+B
WD 4.7 mm
HFW 78.9 µm
curr 0.34 nA
dwell 10 µs
det CBS
10 µm
Helios

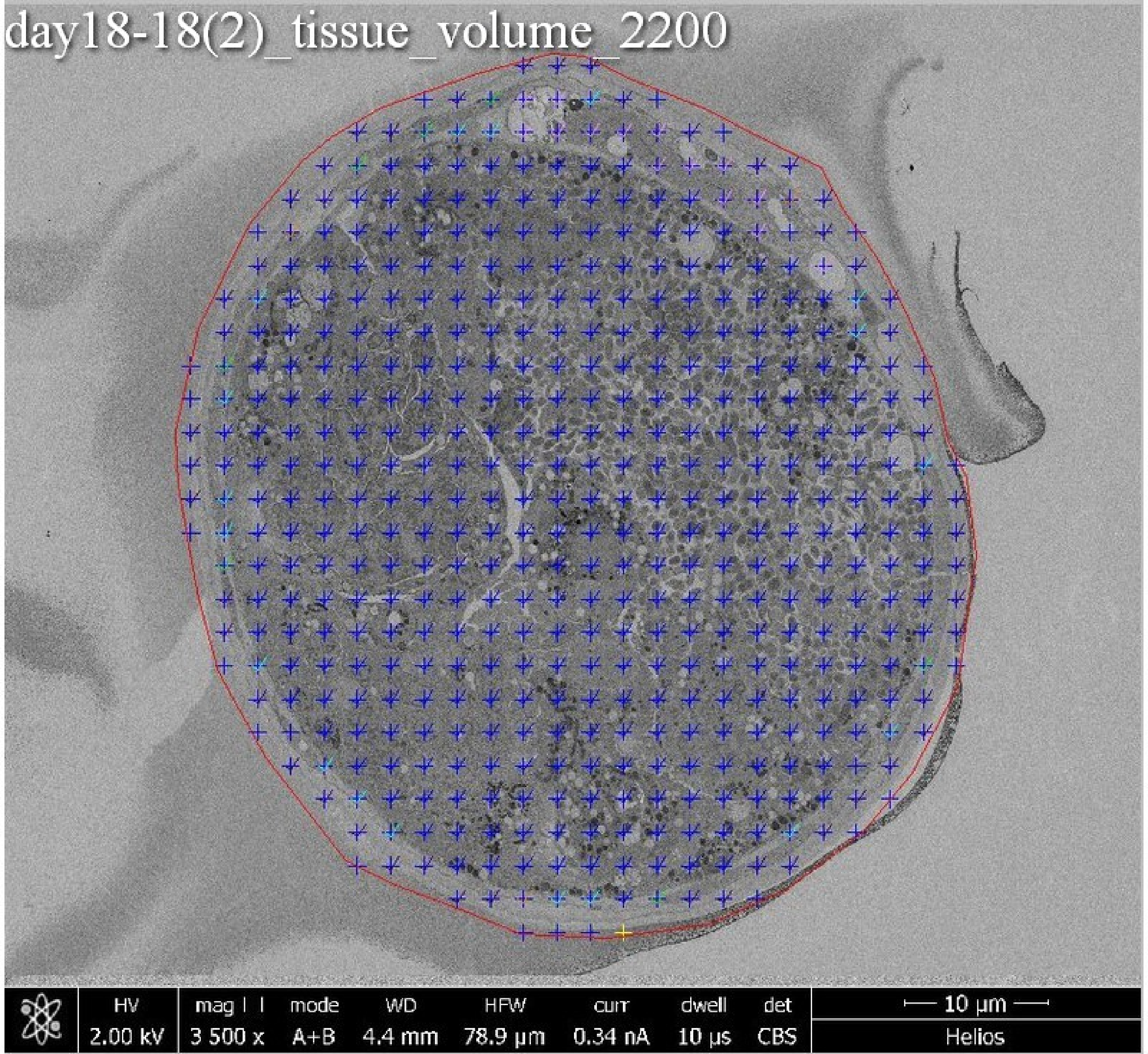
day18-18(2)_tissue_volume_2200
HV 2.00 kV
mag 3 500 x
mode A+B
WD 4.4 mm
HFW 78.9 µm
curr 0.34 nA
dwell 10 µs
det CBS
10 µm
Helios

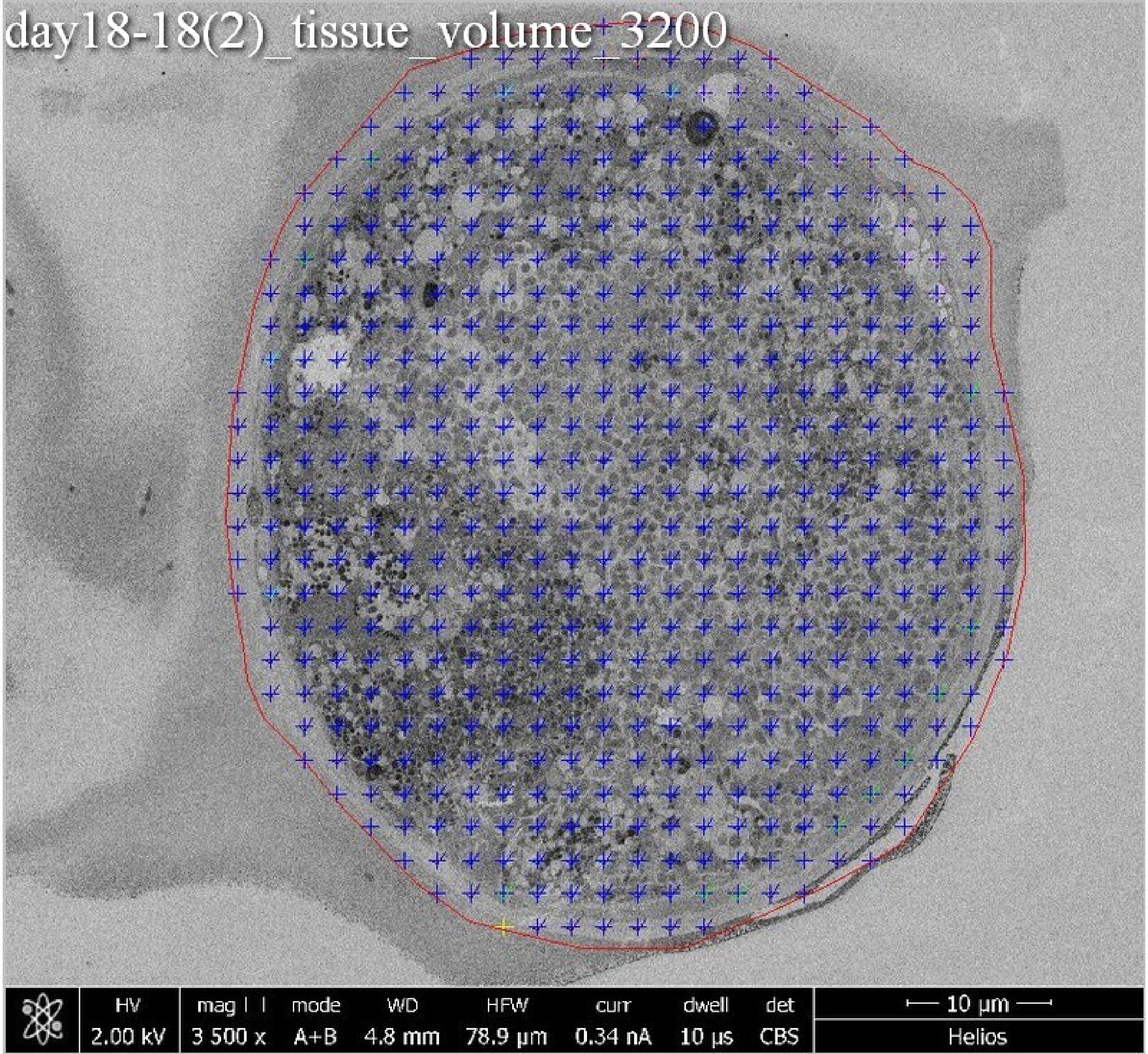
day18-18(2)_tissue_volume_3200
HV
2.00 kV
mag
3 500 x
mode
A+B
WD
4.8 mm
HFW
78.9 μm
curr
0.34 nA
dwell
10 μs
det
CBS
10 μm
Helios

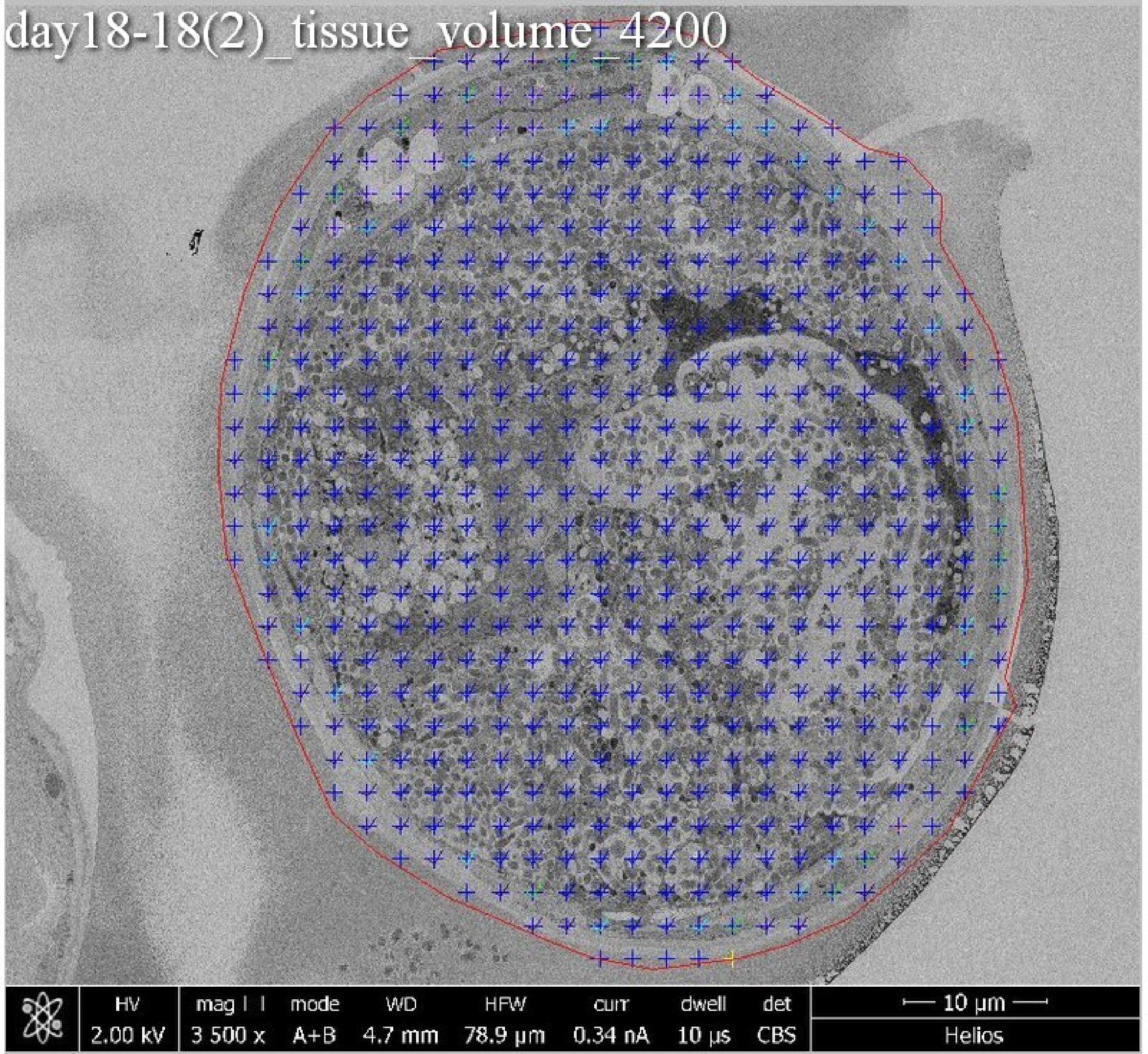
day18-18(2)_tissue_volume_4200
HV 2.00 kV
mag 3 500 x
mode A+B
WD 4.7 mm
HFW 78.9 µm
curr 0.34 nA
dwell 10 µs
det CBS
10 µm
Helios

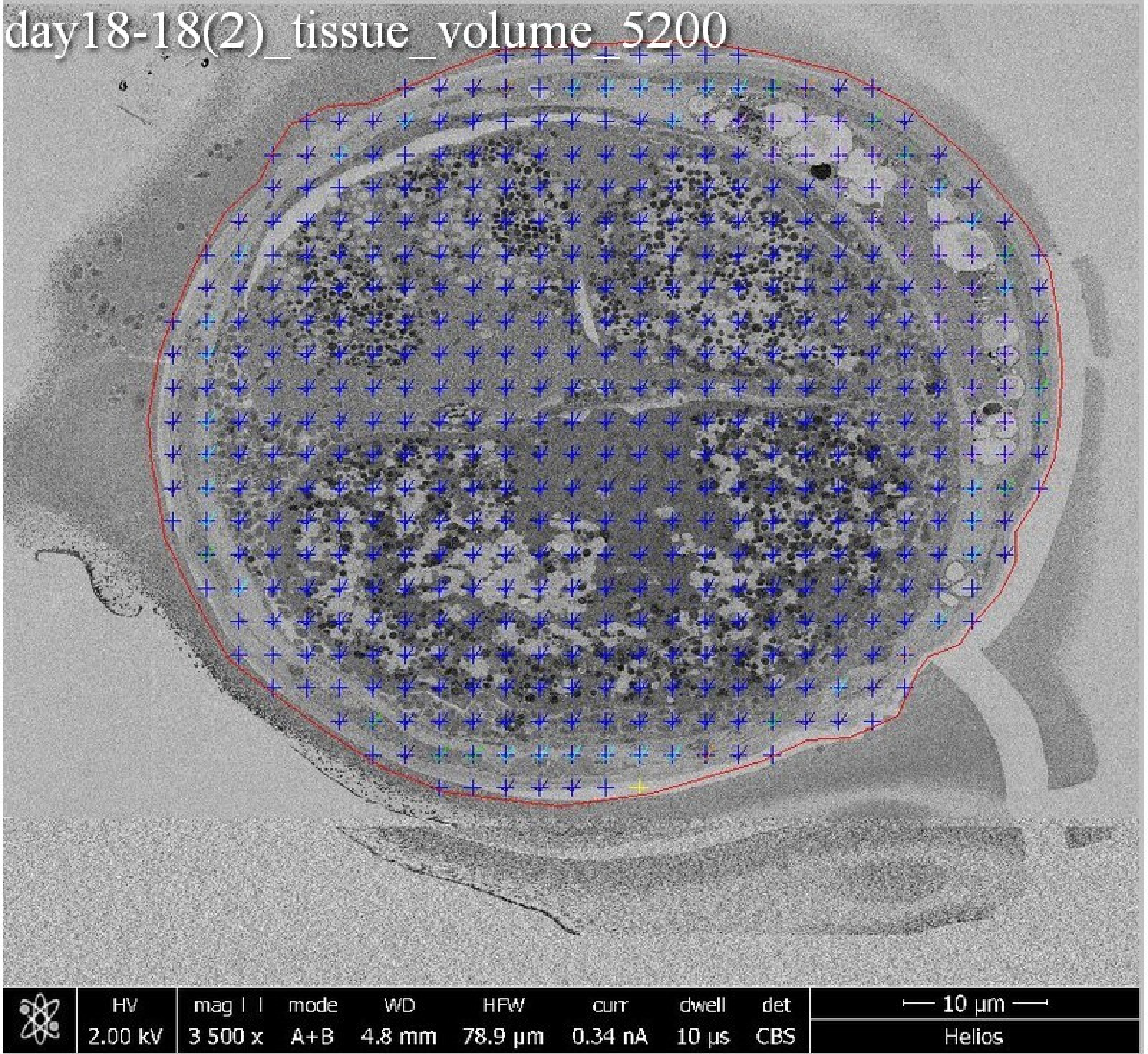
day18-18(2)_tissue_volume_5200
HV
2.00 kV
mag ⊡
3 500 x
mode
A+B
WD
4.8 mm
HFW
78.9 µm
curr
0.34 nA
dwell
10 µs
det
CBS
10 µm
Helios

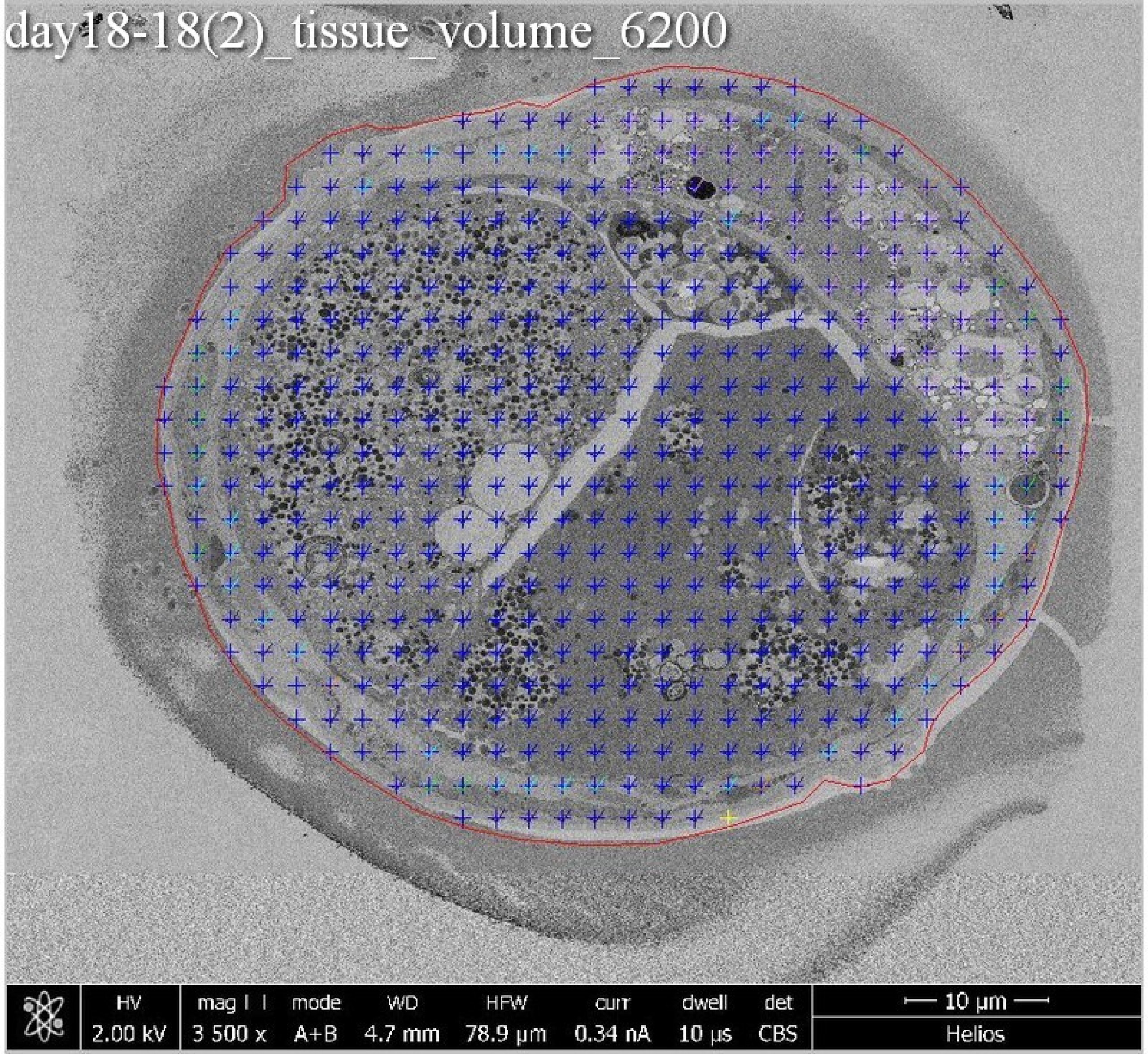

day18-18(2)_tissue_volume_6200
HV
2.00 kV
mag
3 500 x
mode
A+B
WD
4.7 mm
HFW
78.9 µm
curr
0.34 nA
dwell
10 µs
det
CBS
10 µm
Helios

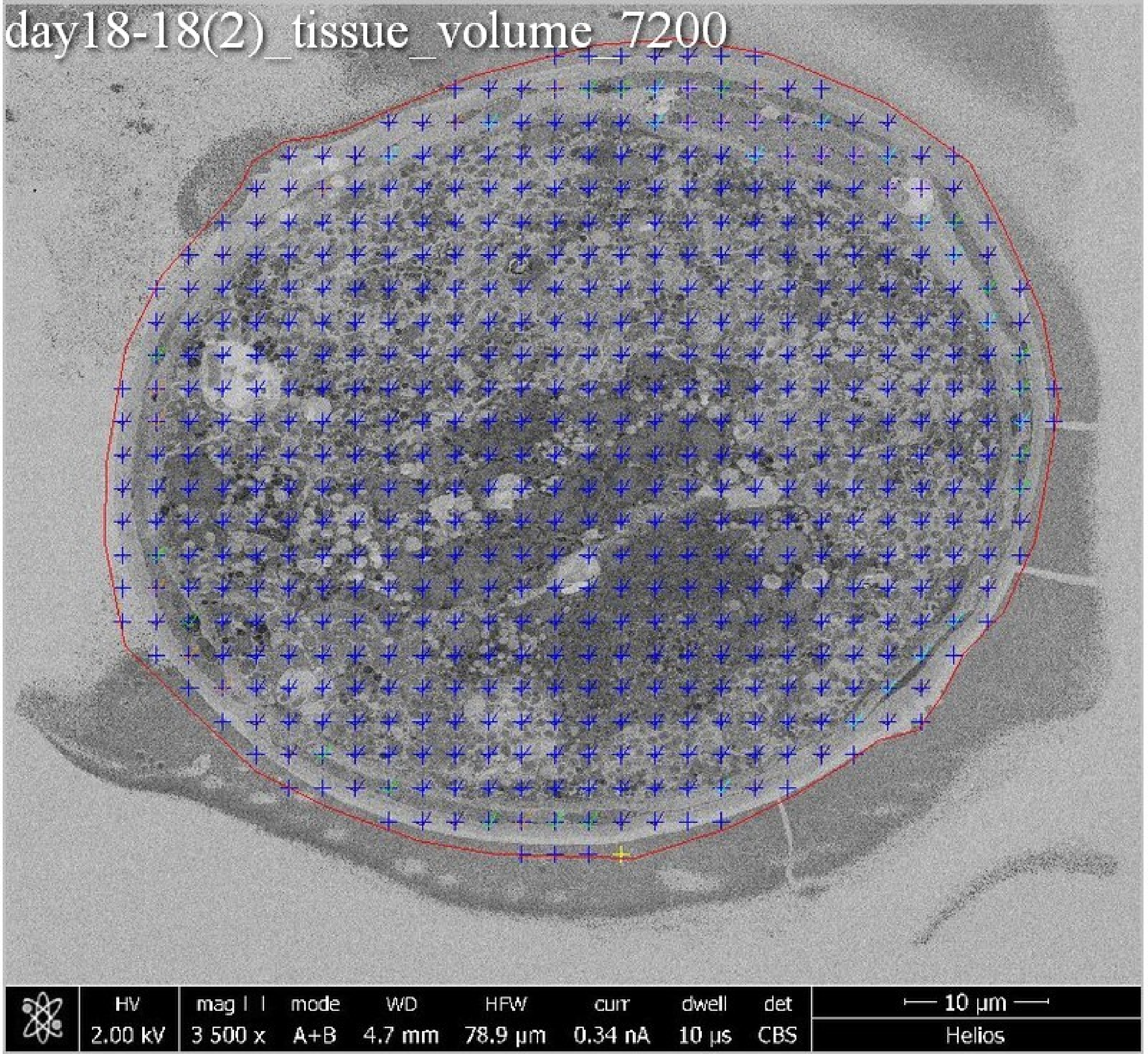
day18-18(2)_tissue_volume_7200
HV
2.00 kV
mag
3 500 x
mode
A+B
WD
4.7 mm
HFW
78.9 µm
curr
0.34 nA
dwell
10 µs
det
CBS
10 µm
Helios

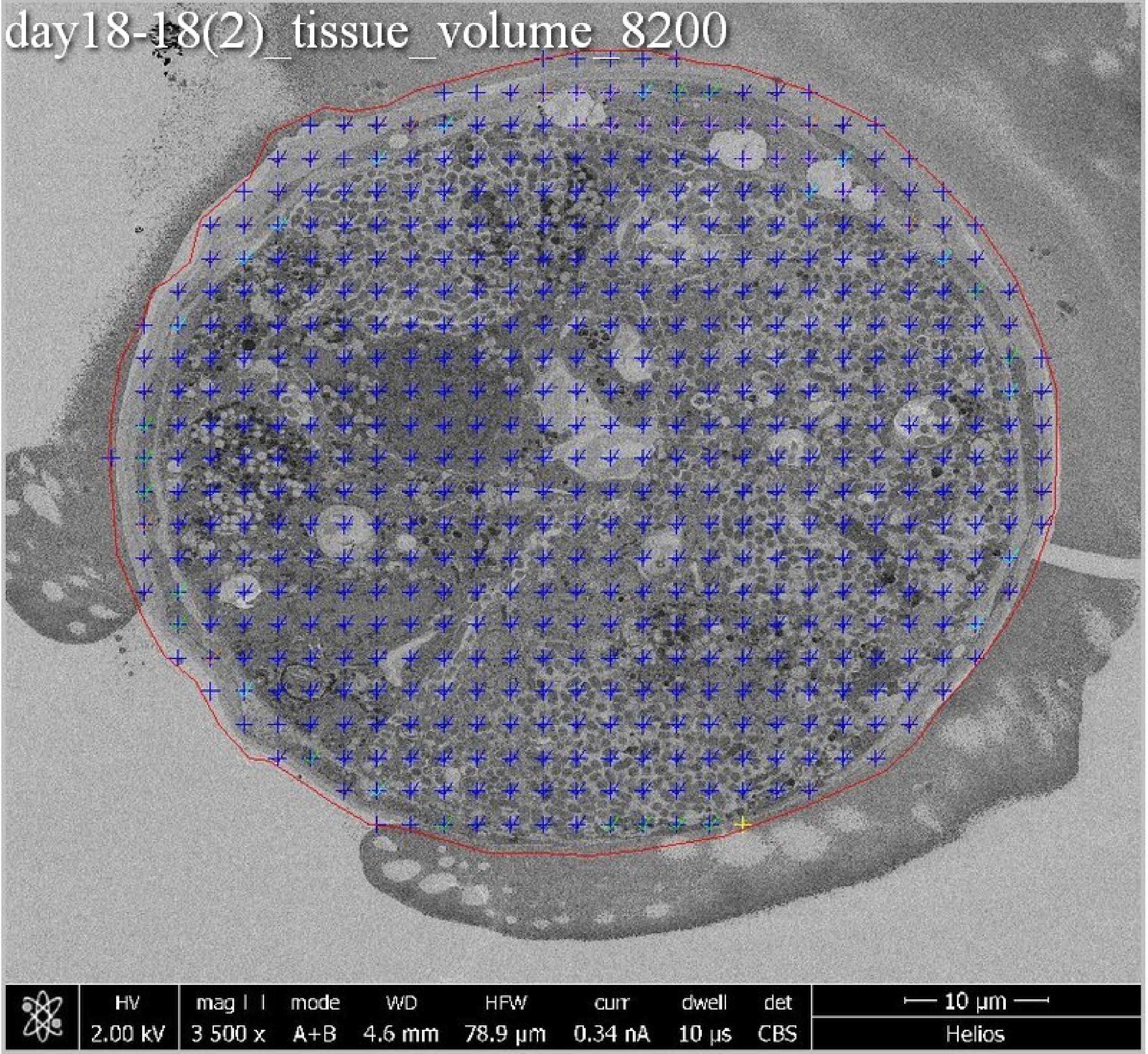
day18-18(2)_tissue_volume_8200
HV
2.00 kV
mag
3 500 x
mode
A+B
WD
4.6 mm
HFW
78.9 µm
curr
0.34 nA
dwell
10 µs
det
CBS
10 µm
Helios

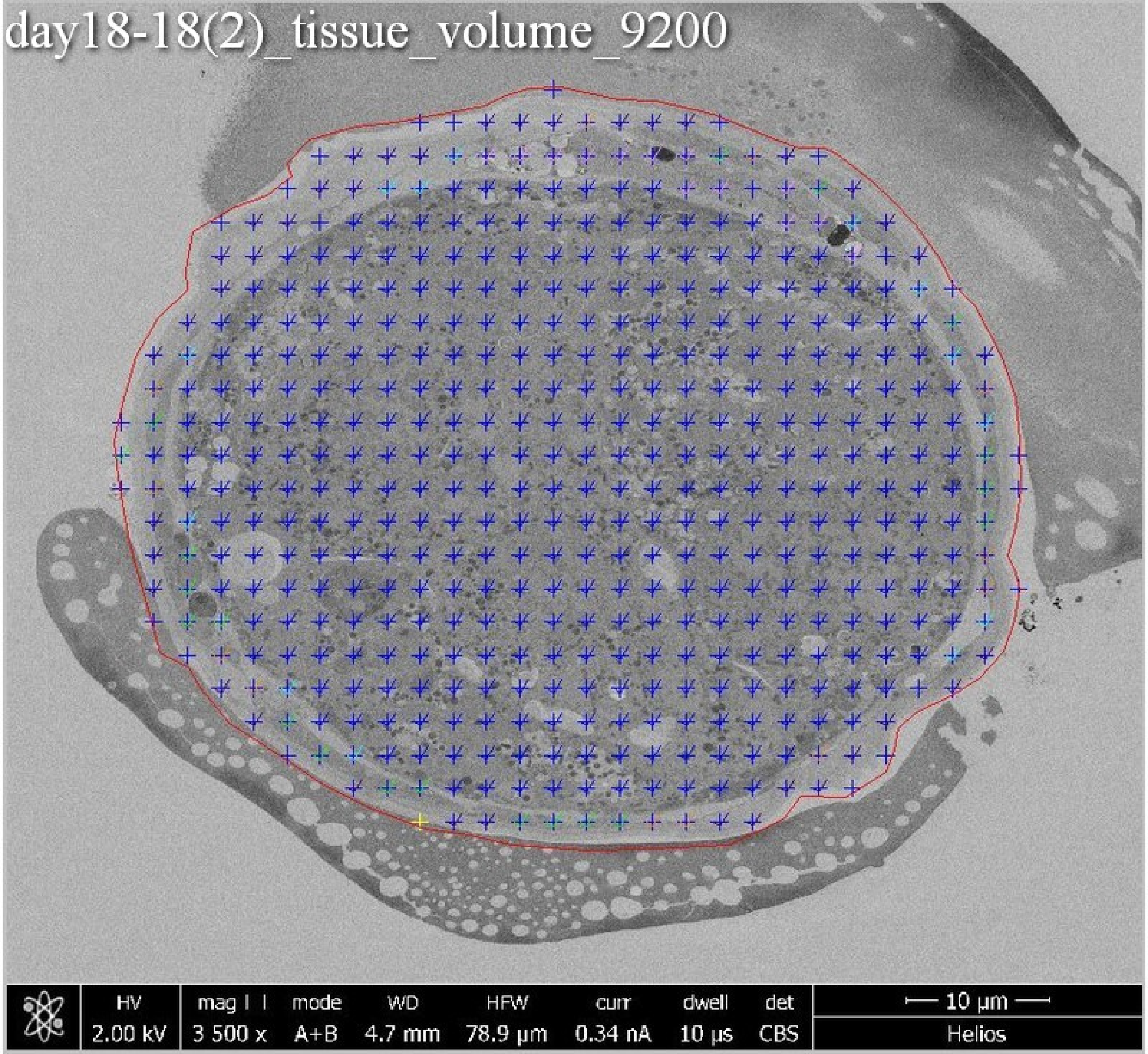
day18-18(2)_tissue_volume_9200
HV
2.00 kV
mag
3 500 x
mode
A+B
WD
4.7 mm
HFW
78.9 µm
curr
0.34 nA
dwell
10 µs
det
CBS
10 µm
Helios

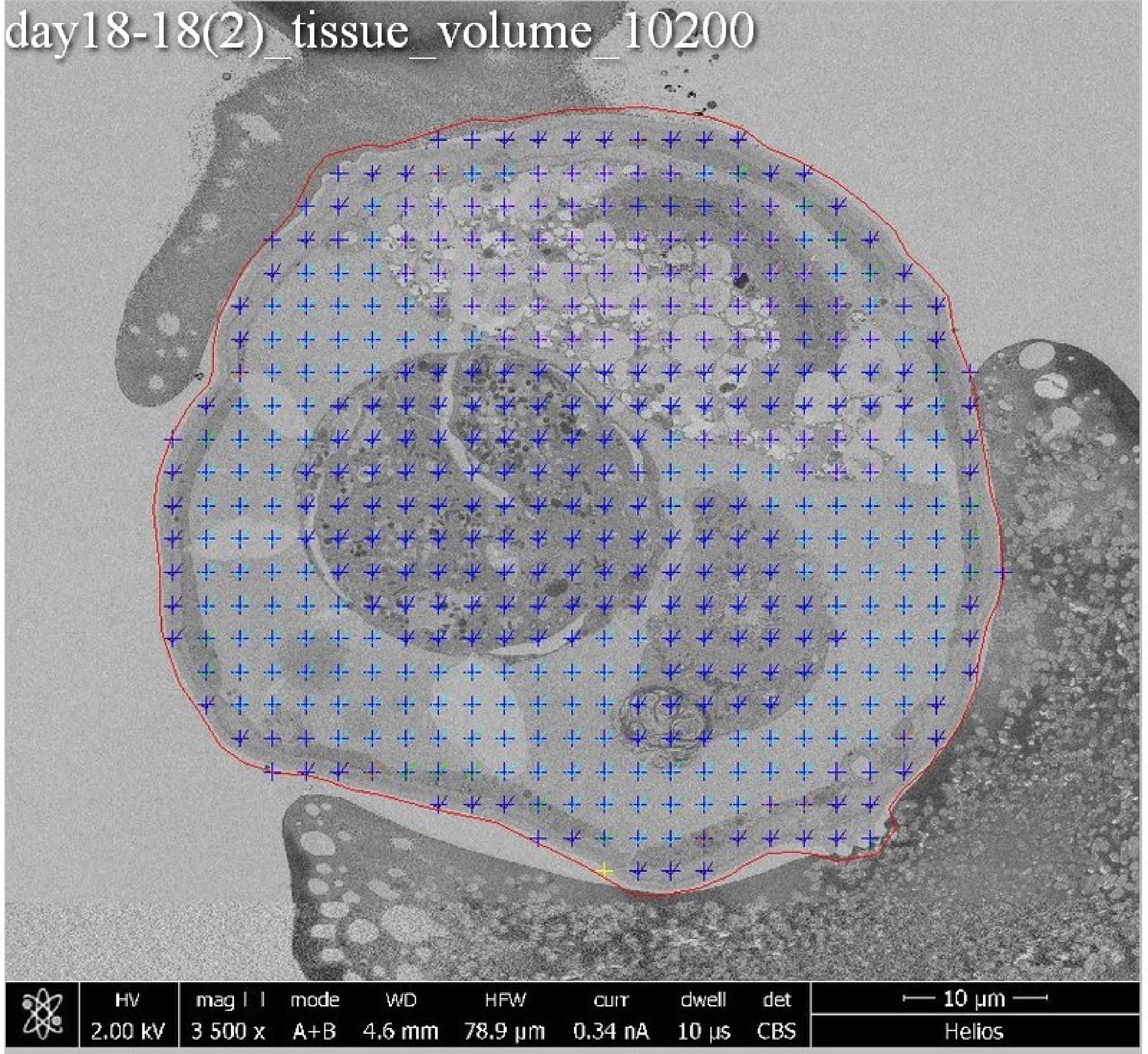
day18-18(2)_tissue_volume_10200
HV
2.00 kV
mag
3 500 x
mode
A+B
WD
4.6 mm
HFW
78.9 µm
curr
0.34 nA
dwell
10 µs
det
CBS
10 µm
Helios

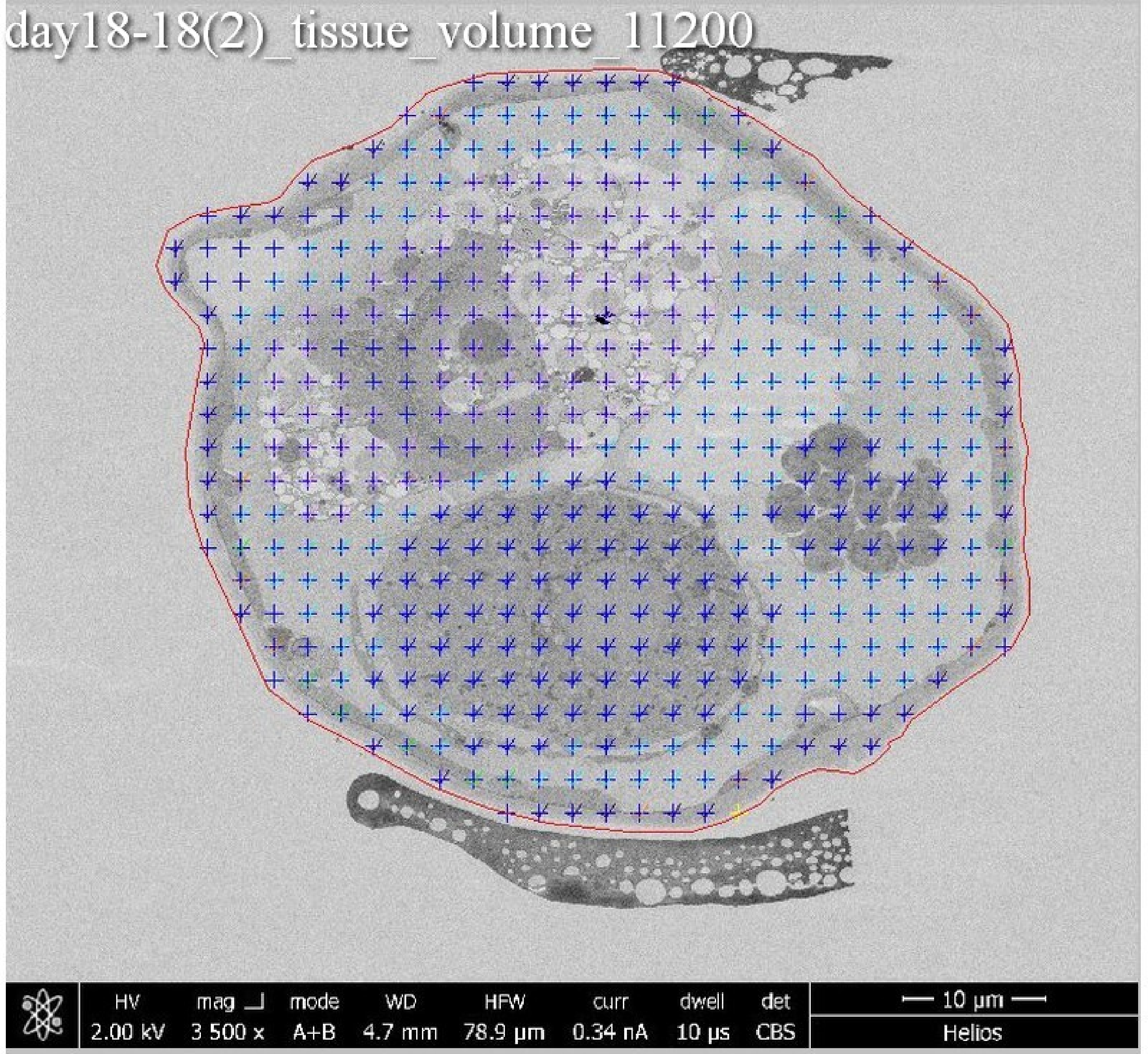
day18-18(2)_tissue_volume_11200
HV 2.00 kV
mag 3 500 x
mode A+B
WD 4.7 mm
HFW 78.9 μm
curr 0.34 nA
dwell 10 μs
det CBS
10 μm
Helios

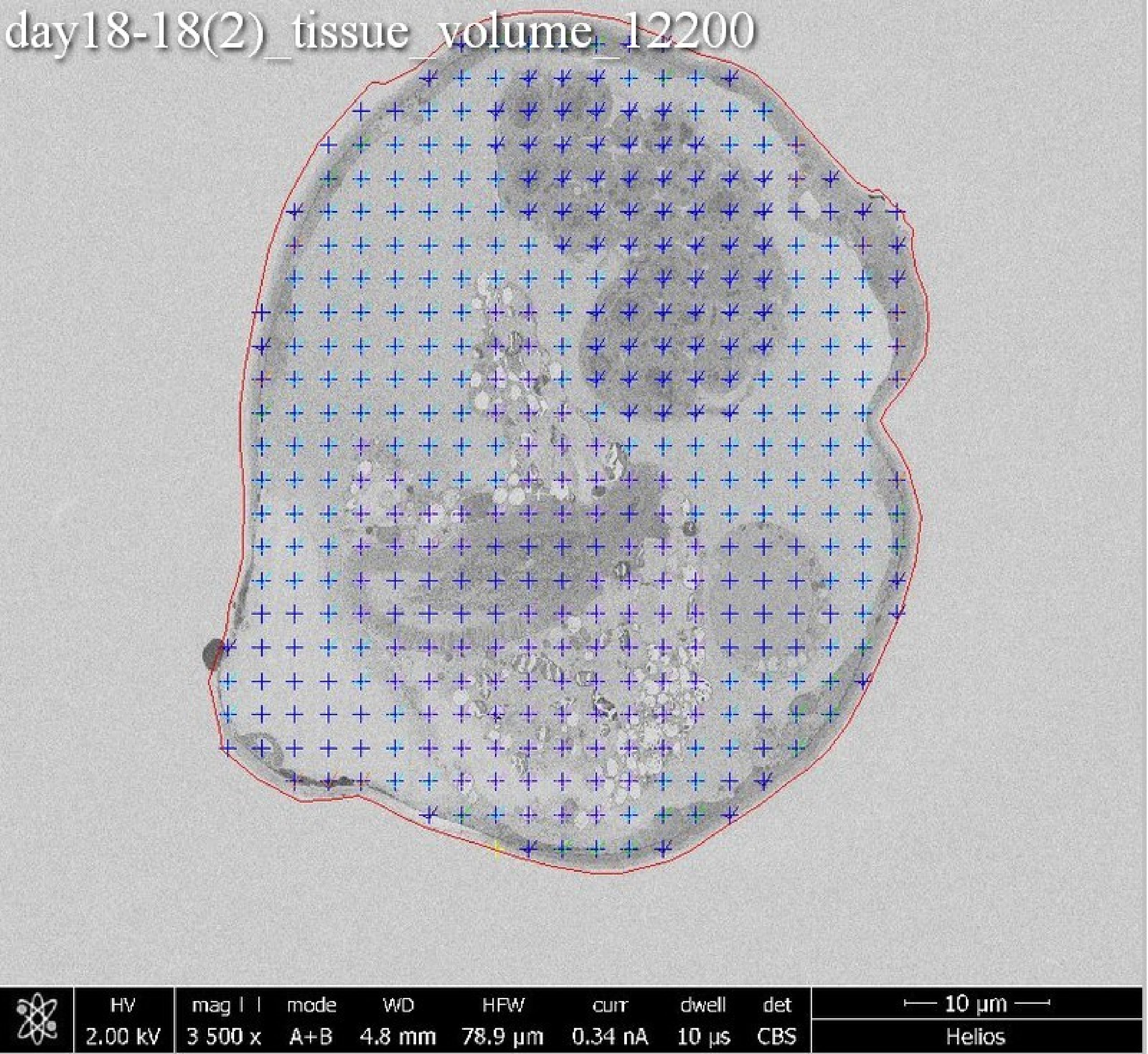
day18-18(2)_tissue_volume_12200
HV
2.00 kV
mag
3 500 x
mode
A+B
WD
4.8 mm
HFW
78.9 µm
curr
0.34 nA
dwell
10 µs
det
CBS
10 µm
Helios

day18-18(2)_tissue_volume_13100

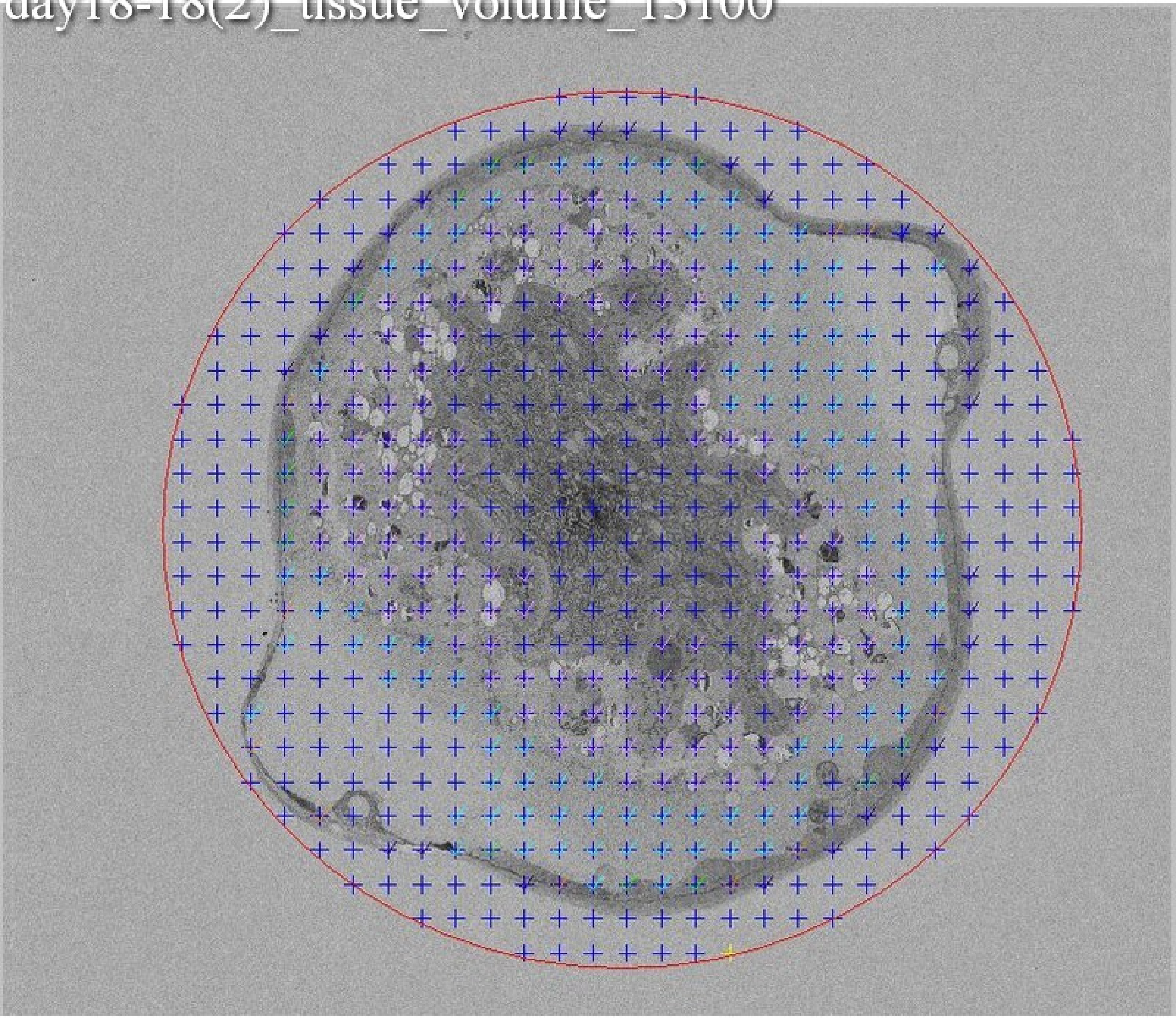

day18-18(2)_tissue_volume_13900

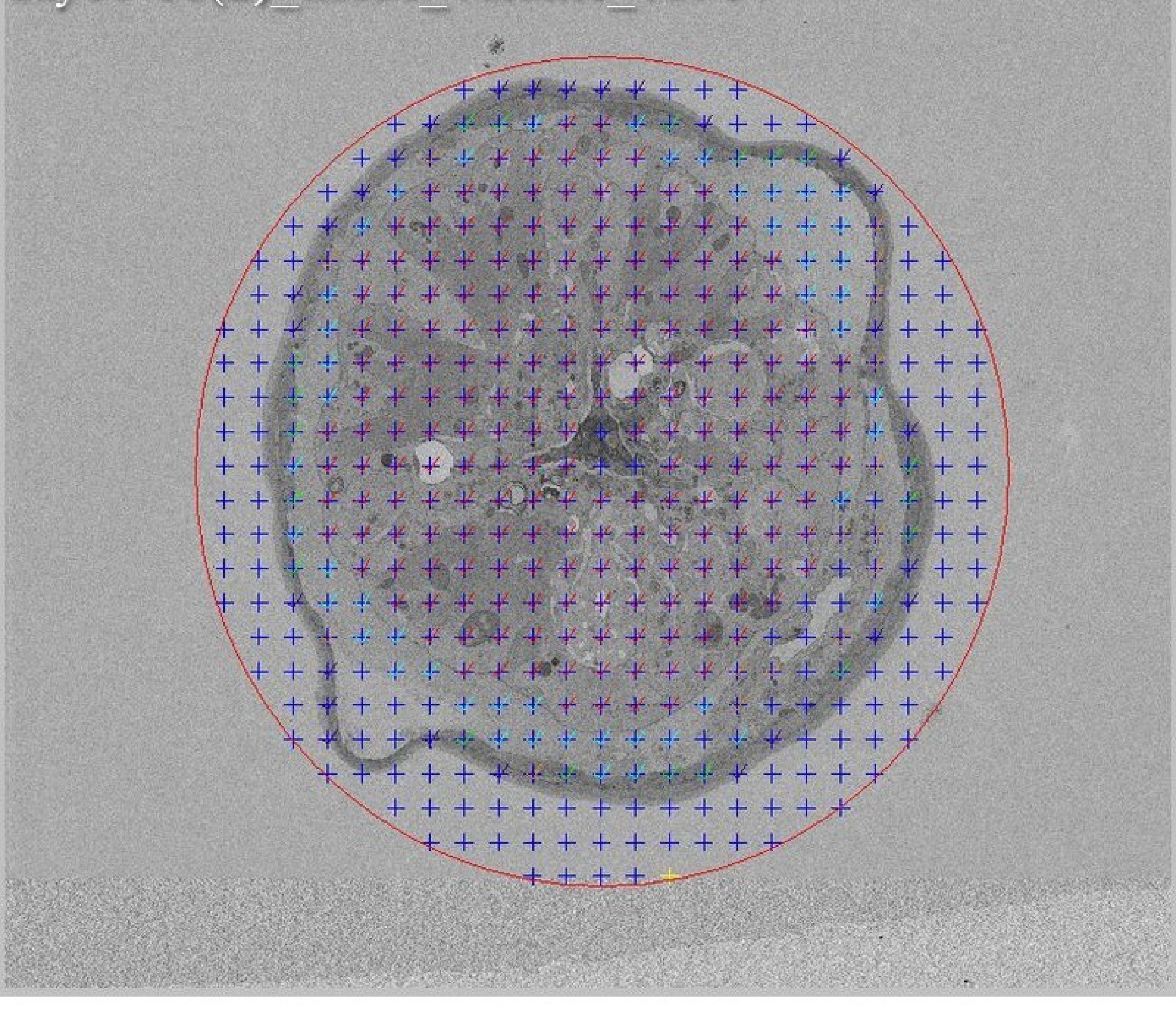

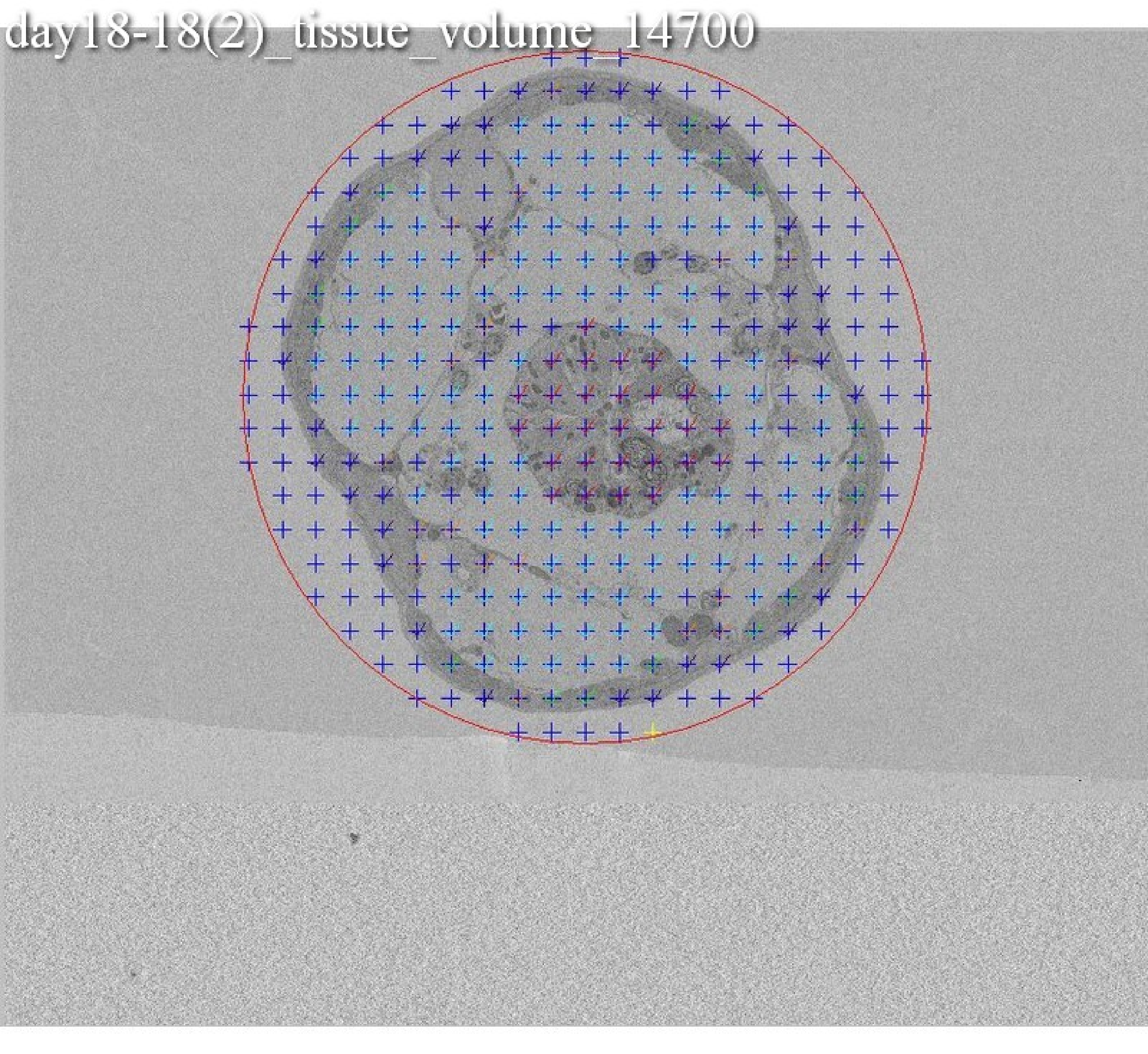
day18-18(2)_tissue_volume_14700

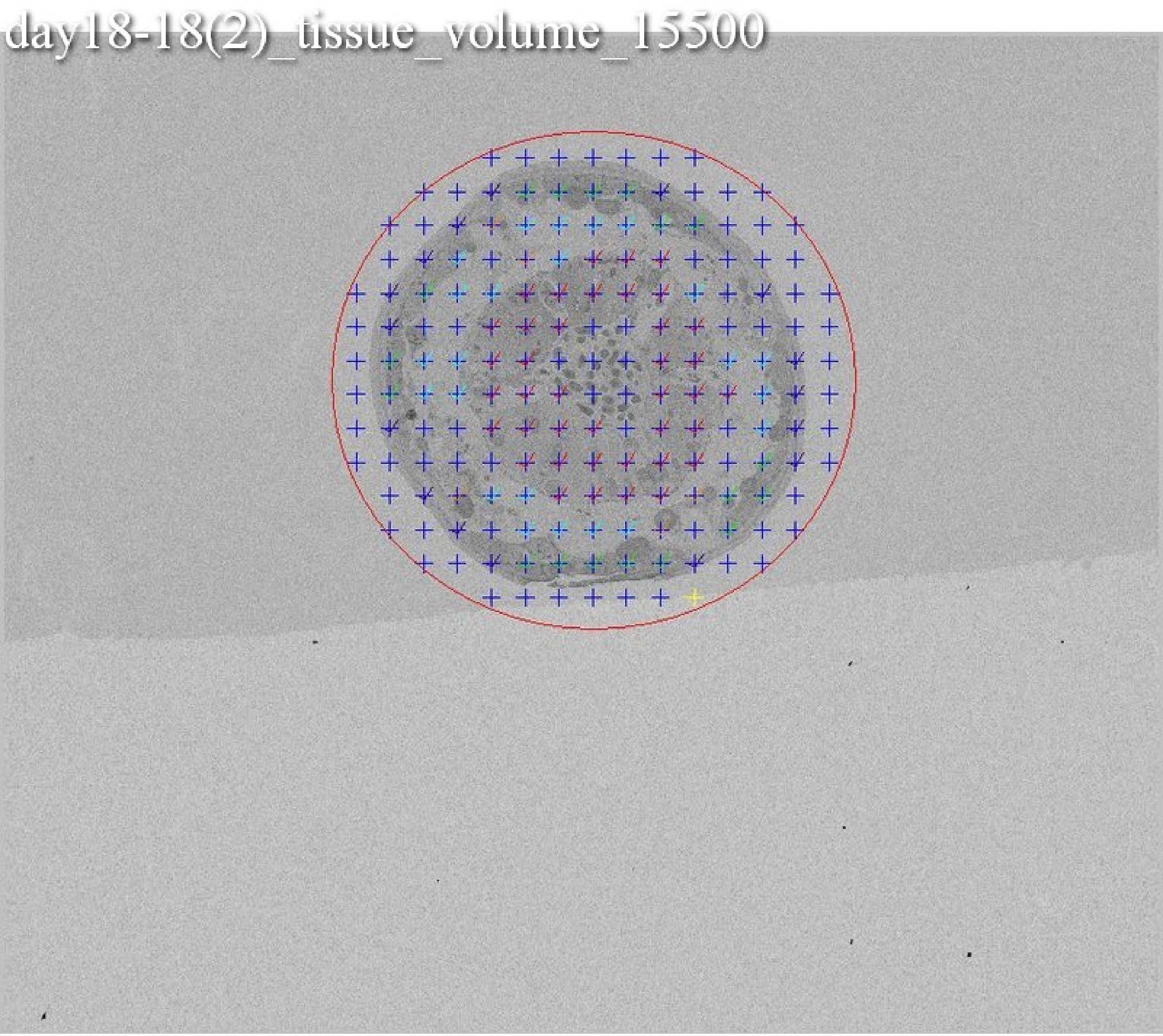
day18-18(2)_tissue_volume_15500
