## Supplementary material for "Old Age Drives Fusion and Expansion of Vitellogenin Vesicles in the Intestine of *Caenorhabditis elegans*": Table S1

day2-3_body volume
section thickness: 70 nm

| Image | Size | pixel size (nm) | Main Grid | Pattern count | object | Hit Points | hfw (µm) | Sampling step (pixels) | interval thickness (nm) | volume of point (nm^3) | volume per portion (nm^3) |
|---|---|---|---|---|---|---|---|---|---|---|---|
| 42.tif | 2048*1887 | 20.75 | Point | 72 | worm body | 33 | 42.5 | 220 | 59500 | 1.24E+12 | 4.09E+13 |
| 902.tif | 2048*1887 | 38.53 | Point | 272 | worm body | 96 | 78.9 | 120 | 59500 | 1.27E+12 | 1.22E+14 |
| 1742.tif | 2048*1887 | 38.53 | Point | 152 | worm body | 122 | 78.9 | 120 | 59500 | 1.27E+12 | 1.55E+14 |
| 2602.tif | 2048*1887 | 38.09 | Point | 178 | worm body | 140 | 78 | 120 | 59500 | 1.24E+12 | 1.74E+14 |
| 3452.tif | 2048*1887 | 38.57 | Point | 167 | worm body | 150 | 79 | 120 | 59500 | 1.27E+12 | 1.91E+14 |
| 4302.tif | 2048*1887 | 38.57 | Point | 178 | worm body | 149 | 79 | 120 | 59500 | 1.27E+12 | 1.90E+14 |
| 5152.tif | 2048*1887 | 38.53 | Point | 152 | worm body | 130 | 78.9 | 120 | 59500 | 1.27E+12 | 1.65E+14 |
| 6002.tif | 2048*1887 | 38.53 | Point | 172 | worm body | 134 | 78.9 | 120 | 59500 | 1.27E+12 | 1.70E+14 |
| 6852.tif | 2048*1887 | 38.53 | Point | 168 | worm body | 138 | 78.9 | 120 | 59500 | 1.27E+12 | 1.75E+14 |
| 7702.tif | 2048*1887 | 38.53 | Point | 159 | worm body | 136 | 78.9 | 120 | 59500 | 1.27E+12 | 1.73E+14 |
| 8552.tif | 2048*1887 | 38.53 | Point | 155 | worm body | 132 | 78.9 | 120 | 59500 | 1.27E+12 | 1.68E+14 |
| 9402.tif | 2048*1887 | 38.53 | Point | 152 | worm body | 133 | 78.9 | 120 | 59500 | 1.27E+12 | 1.69E+14 |
| 10252.tif | 2048*1887 | 38.53 | Point | 150 | worm body | 125 | 78.9 | 120 | 59500 | 1.27E+12 | 1.59E+14 |
| 11102.tif | 2048*1887 | 38.53 | Point | 133 | worm body | 104 | 78.9 | 120 | 59500 | 1.27E+12 | 1.32E+14 |
| 11952.tif | 2048*1887 | 38.53 | Point | 87 | worm body | 67 | 78.9 | 120 | 59500 | 1.27E+12 | 8.52E+13 |
| 12902.tif | 2048*1887 | 8.98 | Point | 153 | worm body | 138 | 18.4 | 120 | 14070 | 1.64E+10 | 2.26E+12 |
| | | | | | | | | | | total volume | 2.27E+15 |

day2-18_body volume
section thickness: 70 nm

| Image | Size | pixel size (nm) | Main Grid | Pattern count | object | Hit Points | hfw (µm) | stampling step (pixel) | interval thickness (nm) | volume per point (nm^3) | volume per portion (nm^3) |
|---|---|---|---|---|---|---|---|---|---|---|---|
| 150.tif | 4096*3775 | 13.43 | Point | 66 | worm body | 51 | 55 | 344 | 70000 | 1.49E+12 | 7.62E+13 |
| 1150.tif | 4096*3775 | 21.97 | Point | 360 | worm body | 199 | 90 | 210 | 70000 | 1.49E+12 | 2.97E+14 |
| 2150.tif | 4096*3775 | 21.97 | Point | 360 | worm body | 183 | 90 | 210 | 70000 | 1.49E+12 | 2.73E+14 |
| 3150.tif | 4096*3775 | 19.53 | Point | 272 | worm body | 148 | 80 | 236 | 70000 | 1.49E+12 | 2.20E+14 |
| 4150.tif | 4096*3775 | 19.53 | Point | 156 | worm body | 142 | 80 | 236 | 70000 | 1.49E+12 | 2.11E+14 |
| 5150.tif | 4096*3775 | 19.53 | Point | 156 | worm body | 146 | 80 | 236 | 70000 | 1.49E+12 | 2.17E+14 |
| 6150.tif | 4096*3775 | 19.53 | Point | 143 | worm body | 140 | 80 | 236 | 70000 | 1.49E+12 | 2.08E+14 |
| 7150.tif | 4096*3775 | 19.26 | Point | 131 | worm body | 123 | 78.9 | 239 | 70000 | 1.48E+12 | 1.82E+14 |
| 8150.tif | 4096*3775 | 19.53 | Point | 140 | worm body | 134 | 80 | 236 | 70000 | 1.49E+12 | 1.99E+14 |
| 9150.tif | 4096*3775 | 19.53 | Point | 153 | worm body | 140 | 80 | 236 | 70000 | 1.49E+12 | 2.08E+14 |
| 10150.tif | 4096*3775 | 19.53 | Point | 272 | worm body | 126 | 80 | 236 | 70000 | 1.49E+12 | 1.87E+14 |
| 11150.tif | 4096*3775 | 19.53 | Point | 121 | worm body | 115 | 80 | 236 | 70000 | 1.49E+12 | 1.71E+14 |
| 12150.tif | 4096*3775 | 19.53 | Point | 98 | worm body | 95 | 80 | 236 | 70000 | 1.49E+12 | 1.41E+14 |
| 13150.tif | 4096*3775 | 14.65 | Point | 81 | worm body | 73 | 60 | 315 | 70000 | 1.49E+12 | 1.09E+14 |
| 14150.tif | 4096*3775 | 14.65 | Point | 57 | worm body | 53 | 60 | 315 | 70000 | 1.49E+12 | 7.90E+13 |
| 15200.tif | 4096*3775 | 4.88 | Point | 69 | worm body | 65 | 20 | 315 | 28000 | 6.62E+10 | 4.31E+12 |
| | | | | | | | | | | total | 2.78E+15 |

day6-8_body_volume
section thickness: 70 nm

| Image | Size | pixel size | Main Grid | Pattern count | object | Hit Points | hfw (µm) | interval thickness (nm) | Sampling step (pixels) | volume per point (nm^3) | volume per portion (nm^3) |
|---|---|---|---|---|---|---|---|---|---|---|---|
| 150.tif | 2048*1887 | 13.48 | Point | 126 | worm body | 111 | 27.6 | 80500 | 120 | 2.11E+11 | 2.34E+13 |
| 1300.tif | 2048*1888 | 38.53 | Point | 85 | worm body | 73 | 78.9 | 80500 | 120 | 1.72E+12 | 1.26E+14 |
| 2450.tif | 2048*1889 | 38.53 | Point | 155 | worm body | 124 | 78.9 | 80500 | 120 | 1.72E+12 | 2.13E+14 |
| 3600.tif | 2048*1890 | 38.53 | Point | 178 | worm body | 164 | 78.9 | 80500 | 120 | 1.72E+12 | 2.82E+14 |
| 4750.tif | 2048*1891 | 54.20 | Point | 225 | worm body | 177 | 111 | 80500 | 86 | 1.75E+12 | 3.10E+14 |
| 5900.tif | 2048*1892 | 54.20 | Point | 235 | worm body | 183 | 111 | 80500 | 86 | 1.75E+12 | 3.20E+14 |
| 7050.tif | 2048*1893 | 54.20 | Point | 221 | worm body | 199 | 111 | 80500 | 86 | 1.75E+12 | 3.48E+14 |
| 8200.tif | 2048*1894 | 54.20 | Point | 222 | worm body | 194 | 111 | 80500 | 86 | 1.75E+12 | 3.39E+14 |
| 9350.tif | 2048*1895 | 54.20 | Point | 224 | worm body | 190 | 111 | 80500 | 86 | 1.75E+12 | 3.32E+14 |
| 10500.tif | 2048*1896 | 38.53 | Point | 189 | worm body | 172 | 78.9 | 80500 | 120 | 1.72E+12 | 2.96E+14 |
| 11650.tif | 2048*1897 | 38.53 | Point | 210 | worm body | 179 | 78.9 | 80500 | 120 | 1.72E+12 | 3.08E+14 |
| 12800.tif | 2048*1898 | 54.20 | Point | 214 | worm body | 187 | 111 | 80500 | 86 | 1.75E+12 | 3.27E+14 |
| 13950.tif | 2048*1899 | 38.53 | Point | 183 | worm body | 158 | 78.9 | 80500 | 120 | 1.72E+12 | 2.72E+14 |
| 15100.tif | 2048*1900 | 38.53 | Point | 183 | worm body | 144 | 78.9 | 80500 | 120 | 1.72E+12 | 2.48E+14 |
| 16250.tif | 2048*1901 | 38.53 | Point | 119 | worm body | 90 | 78.9 | 80500 | 120 | 1.72E+12 | 1.55E+14 |
| 17400.tif | 2048*1902 | 20.75 | Point | 78 | worm body | 55 | 42.5 | 80500 | 200 | 1.39E+12 | 7.63E+13 |
| 18500.tif | 2048*1903 | 5.42 | Point | 60 | worm body | 31 | 11.1 | 10500 | 200 | 1.23E+10 | 3.82E+11 |
| | | | | | | | | | | total | 3.97576E+15 |

day6-10_body_volume
Section thickness: 70 nm

| Image | Size | pixel size | Main Grid | Pattern count | object | Hit Points | hfw (µm) | sampling step (pixels) | interval thickness (nm) | volume per point (nm^3) | volume per part (nm^3) |
|---|---|---|---|---|---|---|---|---|---|---|---|
| 201.tif | 4096*3775 | 13.43 | Point | 64 | worm body | 59 | 55 | 344 | 91000 | 1.94E+12 | 1.15E+14 |
| 1500.tif | 4096*3775 | 15.87 | Point | 117 | worm body | 107 | 65 | 291 | 91000 | 1.94E+12 | 2.08E+14 |
| 2800.tif | 4096*3775 | 19.53 | Point | 145 | worm body | 133 | 80 | 236 | 91000 | 1.93E+12 | 2.57E+14 |
| 4100.tif | 4096*3775 | 20.75 | Point | 180 | worm body | 170 | 85 | 222 | 91000 | 1.93E+12 | 3.28E+14 |
| 5400.tif | 4096*3775 | 24.41 | Point | 215 | worm body | 194 | 100 | 189 | 91000 | 1.94E+12 | 3.76E+14 |
| 6700.tif | 4096*3775 | 24.41 | Point | 221 | worm body | 200 | 100 | 189 | 91000 | 1.94E+12 | 3.88E+14 |
| 8000.tif | 4096*3775 | 24.41 | Point | 228 | worm body | 212 | 100 | 189 | 91000 | 1.94E+12 | 4.11E+14 |
| 9300.tif | 4096*3775 | 23.19 | Point | 234 | worm body | 221 | 95 | 199 | 91000 | 1.94E+12 | 4.28E+14 |
| 10600.tif | 4096*3775 | 23.19 | Point | 233 | worm body | 211 | 95 | 199 | 91000 | 1.94E+12 | 4.09E+14 |
| 11900.tif | 4096*3775 | 23.19 | Point | 242 | worm body | 226 | 95 | 199 | 91000 | 1.94E+12 | 4.38E+14 |
| 13200.tif | 4096*3775 | 24.41 | Point | 258 | worm body | 239 | 100 | 189 | 91000 | 1.94E+12 | 4.63E+14 |
| 14500.tif | 4096*3775 | 24.41 | Point | 242 | worm body | 216 | 100 | 189 | 91000 | 1.94E+12 | 4.19E+14 |
| 15800.tif | 4096*3775 | 23.19 | Point | 204 | worm body | 190 | 95 | 199 | 91000 | 1.94E+12 | 3.68E+14 |
| 17100.tif | 4096*3775 | 23.19 | Point | 161 | worm body | 150 | 95 | 199 | 91000 | 1.94E+12 | 2.91E+14 |
| 18400.tif | 4096*3775 | 15.87 | Point | 92 | worm body | 81 | 65 | 291 | 91000 | 1.94E+12 | 1.57E+14 |
| 19700.tif | 4096*3775 | 7.32 | Point | 86 | worm body | 77 | 30 | 300 | 91000 | 4.39E+11 | 3.38E+13 |
| 20850.tif | 4096*3775 | 2.69 | Point | 59 | worm body | 56 | 11 | 300 | 7000 | 4.54E+09 | 2.54E+11 |
| | | | | | | | | | | total | 5.09E+15 |

day9-3_body_volume
section thickness: 70 nm

| Image | Size | pixel size (nm) | Main Grid | Pattern count | object | Hit Points | HFW (µm) | sampling step (pixels) | interval thickness (nm) | volume per point (nm^3) | volume per portion (nm^3) |
|---|---|---|---|---|---|---|---|---|---|---|---|
| 50.tif | 4096*3775 | 9.28 | Point | 194 | worm body | 183 | 38 | 200 | 73500 | 2.53E+11 | 4.63E+13 |
| 1100.tif | 4096*3775 | 14.65 | Point | 103 | worm body | 98 | 60 | 315 | 73500 | 1.56E+12 | 1.53E+14 |
| 2150.tif | 4096*3775 | 17.09 | Point | 113 | worm body | 107 | 70 | 270 | 73500 | 1.56E+12 | 1.67E+14 |
| 3200.tif | 4096*3775 | 18.31 | Point | 114 | worm body | 106 | 75 | 252 | 73500 | 1.56E+12 | 1.66E+14 |
| 4250.tif | 4096*3775 | 19.53 | Point | 136 | worm body | 121 | 80 | 235 | 73500 | 1.55E+12 | 1.87E+14 |
| 5300.tif | 4096*3775 | 19.53 | Point | 135 | worm body | 128 | 80 | 236 | 73500 | 1.56E+12 | 2.00E+14 |
| 6350.tif | 4096*3775 | 20.83 | Point | 128 | worm body | 123 | 85.3 | 221 | 73500 | 1.56E+12 | 1.91E+14 |
| 7400.tif | 4096*3775 | 20.75 | Point | 168 | worm body | 152 | 85 | 222 | 73500 | 1.56E+12 | 2.37E+14 |
| 8450.tif | 4096*3775 | 21.97 | Point | 178 | worm body | 155 | 90 | 210 | 73500 | 1.56E+12 | 2.43E+14 |
| 9500.tif | 4096*3775 | 21.97 | Point | 182 | worm body | 153 | 90 | 210 | 73500 | 1.56E+12 | 2.39E+14 |
| 10550.tif | 4096*3775 | 20.75 | Point | 173 | worm body | 156 | 85 | 222 | 73500 | 1.56E+12 | 2.43E+14 |
| 11600.tif | 4096*3775 | 20.75 | Point | 183 | worm body | 168 | 85 | 222 | 73500 | 1.56E+12 | 2.62E+14 |
| 12650.tif | 4096*3775 | 20.75 | Point | 183 | worm body | 164 | 85 | 222 | 73500 | 1.56E+12 | 2.56E+14 |
| 13700.tif | 4096*3775 | 20.75 | Point | 169 | worm body | 155 | 85 | 222 | 73500 | 1.56E+12 | 2.42E+14 |
| 14750.tif | 4096*3775 | 19.53 | Point | 143 | worm body | 130 | 80 | 236 | 73500 | 1.56E+12 | 2.03E+14 |
| 15800.tif | 4096*3775 | 14.65 | Point | 88 | worm body | 86 | 60 | 315 | 73500 | 1.56E+12 | 1.35E+14 |
| 17250.tif | 4096*3775 | 1.03 | Point | 156 | worm body | 71 | 4.2 | 315 | 59500 | 6.21E+09 | 4.41E+11 |
| | | | | | | | | | | total | 3.17E+15 |

day9-12_body_volume
section thickness: 70 nm

| Image | Size | pixel size (nm) | Main Grid | Pattern count | object | Hit Points | hfw (µm) | interval thickness (nm) | Sampling step (pixels) | volume of point (nm^3) | volume per portion (nm^3) |
|---|---|---|---|---|---|---|---|---|---|---|---|
| 200.tif | 2048*1887 | 27.00 | Point | 88 | worm body | 73 | 55.3 | 59500 | 170 | 1.25E+12 | 9.15E+13 |
| 1050.tif | 2048*1887 | 38.53 | Point | 140 | worm body | 95 | 78.9 | 59500 | 120 | 1.27E+12 | 1.21E+14 |
| 1900.tif | 2048*1887 | 38.53 | Point | 178 | worm body | 128 | 78.9 | 59500 | 120 | 1.27E+12 | 1.63E+14 |
| 2750.tif | 2048*1887 | 38.53 | Point | 184 | worm body | 135 | 78.9 | 59500 | 120 | 1.27E+12 | 1.72E+14 |
| 3600.tif | 2048*1887 | 38.53 | Point | 173 | worm body | 136 | 78.9 | 59500 | 120 | 1.27E+12 | 1.73E+14 |
| 4450.tif | 2048*1887 | 38.53 | Point | 179 | worm body | 151 | 78.9 | 59500 | 120 | 1.27E+12 | 1.92E+14 |
| 5300.tif | 2048*1887 | 38.53 | Point | 211 | worm body | 164 | 78.9 | 59500 | 120 | 1.27E+12 | 2.09E+14 |
| 6150.tif | 2048*1887 | 38.53 | Point | 233 | worm body | 166 | 78.9 | 59500 | 120 | 1.27E+12 | 2.11E+14 |
| 7000.tif | 2048*1887 | 38.53 | Point | 272 | worm body | 146 | 78.9 | 59500 | 120 | 1.27E+12 | 1.86E+14 |
| 7850.tif | 2048*1887 | 38.53 | Point | 272 | worm body | 158 | 78.9 | 59500 | 120 | 1.27E+12 | 2.01E+14 |
| 8700.tif | 2048*1887 | 38.53 | Point | 272 | worm body | 165 | 78.9 | 59500 | 120 | 1.27E+12 | 2.10E+14 |
| 9550.tif | 2048*1887 | 38.53 | Point | 272 | worm body | 158 | 78.9 | 59500 | 120 | 1.27E+12 | 2.01E+14 |
| 10400.tif | 2048*1887 | 38.53 | Point | 255 | worm body | 140 | 78.9 | 59500 | 120 | 1.27E+12 | 1.78E+14 |
| 11250.tif | 2048*1887 | 38.53 | Point | 146 | worm body | 134 | 78.9 | 59500 | 120 | 1.27E+12 | 1.70E+14 |
| 12100.tif | 2048*1887 | 38.53 | Point | 125 | worm body | 117 | 78.9 | 59500 | 120 | 1.27E+12 | 1.49E+14 |
| 12950_001.tif | 2048*1887 | 27.00 | Point | 64 | worm body | 50 | 55.3 | 59500 | 170 | 1.25E+12 | 6.27E+13 |
| 13700_001.tif | 2048*1887 | 3.85 | Point | 16 | worm body | 13 | 7.89 | 3.852539 | 300 | 1.40E+10 | 1.82E+11 |
| | | | | | | | | | | total | 2.69E+15 |

day18-18(1)_body_volume

section number: 1-12660,  section thickness 50nm; section number: 12661-18500, section thickness 60 nm

| Image | Size | pixel size (nm) | Main Grid | Pattern count | object | Hit Points | hfw (µm) | Sampling step (pixels) | interval thickness (nm) | volume of point (nm^3) | volume per portion (nm^3) |
|---|---|---|---|---|---|---|---|---|---|---|---|
| 1_300_001.tif | X = 2048; Y = 1887 | 27.00 | Point | 71 | worm body | 58 | 55.3 | 170 | 62500 | 1.32E+12 | 7.64E+13 |
| 1_1550.tif | X = 2048; Y = 1887 | 38.53 | Point | 92 | worm body | 73 | 78.9 | 120 | 62500 | 1.34E+12 | 9.75E+13 |
| 1_2800.tif | X = 2048; Y = 1887 | 38.53 | Point | 132 | worm body | 102 | 78.9 | 120 | 62500 | 1.34E+12 | 1.36E+14 |
| 1_3850_001.tif | X = 2048; Y = 1887 | 38.53 | Point | 146 | worm body | 118 | 78.9 | 120 | 62500 | 1.34E+12 | 1.58E+14 |
| 1_4050_001.tif | X = 2048; Y = 1887 | 38.53 | Point | 139 | worm body | 121 | 78.9 | 120 | 62500 | 1.34E+12 | 1.62E+14 |
| 1_5300_001.tif | X = 2048; Y = 1887 | 38.53 | Point | 161 | worm body | 137 | 78.9 | 120 | 62500 | 1.34E+12 | 1.83E+14 |
| 1_6550_001.tif | X = 2048; Y = 1887 | 38.53 | Point | 167 | worm body | 144 | 78.9 | 120 | 62500 | 1.34E+12 | 1.92E+14 |
| 1_7800_001.tif | X = 2048; Y = 1887 | 38.53 | Point | 161 | worm body | 133 | 78.9 | 120 | 62500 | 1.34E+12 | 1.78E+14 |
| 1_9050_001.tif | X = 2048; Y = 1887 | 38.53 | Point | 174 | worm body | 148 | 78.9 | 120 | 62500 | 1.34E+12 | 1.98E+14 |
| 1_10300_001.tif | X = 2048; Y = 1887 | 38.53 | Point | 171 | worm body | 149 | 78.9 | 120 | 62500 | 1.34E+12 | 1.99E+14 |
| 1_11550_001.tif | X = 2048; Y = 1887 | 38.53 | Point | 151 | worm body | 131 | 78.9 | 120 | 61400 | 1.31E+12 | 1.72E+14 |
| 1_12800_001.tif | X = 2048; Y = 1887 | 38.53 | Point | 148 | worm body | 126 | 78.9 | 120 | 63000 | 1.35E+12 | 1.70E+14 |
| 1_14900_001.tif | X = 2048; Y = 1768 | 38.53 | Point | 135 | worm body | 116 | 78.9 | 120 | 63000 | 1.35E+12 | 1.56E+14 |
| 1_15950_001.tif | X = 2048; Y = 1887 | 38.53 | Point | 133 | worm body | 117 | 78.9 | 120 | 63000 | 1.35E+12 | 1.58E+14 |
| 1_17000_002.tif | X = 2048; Y = 1887 | 20.75 | Point | 96 | worm body | 78 | 42.5 | 170 | 63000 | 7.84E+11 | 6.12E+13 |
| 1_18100_002.tif | X = 2048; Y = 1887 | 16.85 | Point | 132 | worm body | 81 | 34.5 | 170 | 63000 | 5.17E+11 | 4.19E+13 |
| | | | | | | | | | | total | 2.34E+15 |

day18-18(2)_body_volume
section number: 1-12660,  section thickness 50nm; section number: 12661-18500, section thickness 60 nm

| Image | Size | pixel size (nm) | Main Grid | Pattern count | object | Hit Points | interval thickness (nm) | hfw (µm) | Sampling step (pixels) | volume per point (nm^3) | volumme per portion (nm^3) |
|---|---|---|---|---|---|---|---|---|---|---|---|
| 2_200.tif | X = 2048; Y = 1887 | 38.52539 | Point | 97 | worm body | 73 | 50000 | 78.9 | 120 | 1.07E+12 | 7.80E+13 |
| 2_1200.tif | X = 2048; Y = 1887 | 38.52539 | Point | 111 | worm body | 91 | 50000 | 78.9 | 120 | 1.07E+12 | 9.72E+13 |
| 2_2200.tif | X = 2048; Y = 1887 | 38.52539 | Point | 136 | worm body | 113 | 50000 | 78.9 | 120 | 1.07E+12 | 1.21E+14 |
| 2_3200_001.tif | X = 2048; Y = 1887 | 38.52539 | Point | 140 | worm body | 127 | 50000 | 78.9 | 120 | 1.07E+12 | 1.36E+14 |
| 2_4200_001.tif | X = 2048; Y = 1887 | 38.52539 | Point | 154 | worm body | 130 | 50000 | 78.9 | 120 | 1.07E+12 | 1.39E+14 |
| 2_5250_001.tif | X = 2048; Y = 1887 | 38.52539 | Point | 148 | worm body | 118 | 50000 | 78.9 | 120 | 1.07E+12 | 1.26E+14 |
| 2_6200_001.tif | X = 2048; Y = 1887 | 38.52539 | Point | 147 | worm body | 122 | 50000 | 78.9 | 120 | 1.07E+12 | 1.30E+14 |
| 2_7200_001.tif | X = 2048; Y = 1887 | 38.52539 | Point | 149 | worm body | 131 | 50000 | 78.9 | 120 | 1.07E+12 | 1.40E+14 |
| 2_8200_001.tif | X = 2048; Y = 1887 | 38.52539 | Point | 151 | worm body | 129 | 50000 | 78.9 | 120 | 1.07E+12 | 1.38E+14 |
| 2_9200_001.tif | X = 2048; Y = 1887 | 38.52539 | Point | 139 | worm body | 117 | 50000 | 78.9 | 120 | 1.07E+12 | 1.25E+14 |
| 2_10200_001.tif | X = 2048; Y = 1887 | 38.52539 | Point | 144 | worm body | 111 | 50000 | 78.9 | 120 | 1.07E+12 | 1.19E+14 |
| 2_11200_001.tif | X = 2048; Y = 1887 | 38.52539 | Point | 142 | worm body | 111 | 50000 | 78.9 | 120 | 1.07E+12 | 1.19E+14 |
| 2_12200_001.tif | X = 2048; Y = 1887 | 38.52539 | Point | 136 | worm body | 104 | 50400 | 78.9 | 120 | 1.08E+12 | 1.12E+14 |
| 2_13100_001.tif | X = 2048; Y = 1768 | 38.52539 | Point | 136 | worm body | 99 | 48000 | 78.9 | 120 | 1.03E+12 | 1.02E+14 |
| 2_13900_001.tif | X = 2048; Y = 1768 | 38.52539 | Point | 108 | worm body | 83 | 48000 | 78.9 | 120 | 1.03E+12 | 8.51E+13 |
| 2_14700_001.tif | X = 2048; Y = 1768 | 38.52539 | Point | 80 | worm body | 59 | 48000 | 78.9 | 120 | 1.03E+12 | 6.05E+13 |
| 2_15500_001.tif | X = 2048; Y = 1768 | 38.52539 | Point | 42 | worm body | 32 | 51000 | 78.9 | 120 | 1.09E+12 | 3.49E+13 |
| | | | | | | | | | | total volume | 1.86E+15 |

day2-3_tissue volume
section thickness: 70 nm

| Image | Size | pixel size (nm) | Main Grid | Pattern count | object | Hit Points | object | Hit Points | hfw (µm) | interval thickness (nm) | Sampling step (pixels) | volume of point (nm^3) | intestinl volume per part (nm^3) | volume of body cavity per part (nm^3) |
|---|---|---|---|---|---|---|---|---|---|---|---|---|---|---|
| 42.tif | X = 2048; Y = 1887 | 20.75 | Point | 467 | intestine | 0 | body cavity | 30 | 42.5 | 59500 | 60 | 9.22E+10 | 0.00E+00 | 2.77E+12 |
| 902.tif | X = 2048; Y = 1887 | 38.53 | Point | 397 | intestine | 104 | body cavity | 16 | 78.9 | 59500 | 60 | 3.18E+11 | 3.31E+13 | 5.09E+12 |
| 1762.tif | X = 2048; Y = 1887 | 38.53 | Point | 513 | intestine | 92 | body cavity | 10 | 78.9 | 59500 | 60 | 3.18E+11 | 2.92E+13 | 3.18E+12 |
| 2602.tif | X = 2048; Y = 1887 | 38.09 | Point | 592 | intestine | 126 | body cavity | 36 | 78 | 59500 | 60 | 3.11E+11 | 3.91E+13 | 1.12E+13 |
| 3452.tif | X = 2048; Y = 1887 | 38.57 | Point | 622 | intestine | 149 | body cavity | 50 | 79 | 59500 | 60 | 3.19E+11 | 4.75E+13 | 1.59E+13 |
| 4302.tif | X = 2048; Y = 1887 | 38.57 | Point | 632 | intestine | 126 | body cavity | 36 | 79 | 59500 | 60 | 3.19E+11 | 4.02E+13 | 1.15E+13 |
| 5152.tif | X = 2048; Y = 1887 | 38.53 | Point | 537 | intestine | 150 | body cavity | 47 | 78.9 | 59500 | 60 | 3.18E+11 | 4.77E+13 | 1.49E+13 |
| 6002.tif | X = 2048; Y = 1887 | 38.53 | Point | 546 | intestine | 71 | body cavity | 13 | 78.9 | 59500 | 60 | 3.18E+11 | 2.26E+13 | 4.13E+12 |
| 6852.tif | X = 2048; Y = 1887 | 38.53 | Point | 562 | intestine | 92 | body cavity | 24 | 78.9 | 59500 | 60 | 3.18E+11 | 2.92E+13 | 7.63E+12 |
| 7702.tif | X = 2048; Y = 1887 | 38.53 | Point | 553 | intestine | 118 | body cavity | 15 | 78.9 | 59500 | 60 | 3.18E+11 | 3.75E+13 | 4.77E+12 |
| 8552.tif | X = 2048; Y = 1887 | 38.53 | Point | 547 | intestine | 101 | body cavity | 11 | 78.9 | 59500 | 60 | 3.18E+11 | 3.21E+13 | 3.50E+12 |
| 9402.tif | X = 2048; Y = 1887 | 38.53 | Point | 536 | intestine | 155 | body cavity | 11 | 78.9 | 59500 | 60 | 3.18E+11 | 4.93E+13 | 3.50E+12 |
| 10252.tif | X = 2048; Y = 1887 | 38.53 | Point | 511 | intestine | 168 | body cavity | 11 | 78.9 | 59500 | 60 | 3.18E+11 | 5.34E+13 | 3.50E+12 |
| 11002.tif | X = 2048; Y = 1887 | 38.53 | Point | 437 | intestine | 102 | body cavity | 11 | 78.9 | 59500 | 60 | 3.18E+11 | 3.24E+13 | 3.50E+12 |
| 11952.tif | X = 2048; Y = 1887 | 38.53 | Point | 270 | intestine | 0 | body cavity | 14 | 78.9 | 59500 | 60 | 3.18E+11 | 0.00E+00 | 4.45E+12 |
| 12902_001.tif | X = 2048; Y = 1887 | 8.98 | Point | 598 | intestine | 0 | body cavity | 31 | 18.4 | 14070 | 60 | 4.09E+09 | 0.00E+00 | 1.27E+11 |

day2-18_tissue volume
section thickness: 70 nm

| Image | Size | pixel size (nm) | Main Grid | Pattern count | object | Hit Points | object | Hit Points | hfw (μm) | Sampling step (pixels) | interval thickness (nm) | volume per point (nm^3) | intestinal volume of per part (nm^3) | volume of the body cavity per part (nm^3) |
|---|---|---|---|---|---|---|---|---|---|---|---|---|---|---|
| 150.tif | 4096*3775 | 13.43 | Point | 232 | intestine | 0 | body cavity | 11 | 55 | 172 | 70000 | 3.73E+11 | 0.00E+00 | 4.11E+12 |
| 1150.tif | 4096*3775 | 21.97 | Point | 868 | intestine | 302 | body cavity | 40 | 90 | 105 | 70000 | 3.73E+11 | 1.13E+14 | 1.49E+13 |
| 2150.tif | 4096*3775 | 21.97 | Point | 779 | intestine | 209 | body cavity | 31 | 90 | 105 | 70000 | 3.73E+11 | 7.79E+13 | 1.16E+13 |
| 3150.tif | 4096*3775 | 19.53 | Point | 645 | intestine | 104 | body cavity | 23 | 80 | 118 | 70000 | 3.72E+11 | 3.87E+13 | 8.55E+12 |
| 4150.tif | 4096*3775 | 19.53 | Point | 589 | intestine | 109 | body cavity | 32 | 80 | 118 | 70000 | 3.72E+11 | 4.05E+13 | 1.19E+13 |
| 5150.tif | 4096*3775 | 19.53 | Point | 593 | intestine | 146 | body cavity | 34 | 80 | 118 | 70000 | 3.72E+11 | 5.43E+13 | 1.26E+13 |
| 6150.tif | 4096*3775 | 19.53 | Point | 588 | intestine | 156 | body cavity | 24 | 80 | 118 | 70000 | 3.72E+11 | 5.80E+13 | 8.92E+12 |
| 7150.tif | 4096*3775 | 19.26 | Point | 528 | intestine | 139 | body cavity | 21 | 78.9 | 119 | 70000 | 3.68E+11 | 5.11E+13 | 7.72E+12 |
| 8150.tif | 4096*3775 | 19.53 | Point | 551 | intestine | 122 | body cavity | 36 | 80 | 118 | 70000 | 3.72E+11 | 4.54E+13 | 1.34E+13 |
| 9150.tif | 4096*3775 | 19.53 | Point | 608 | intestine | 157 | body cavity | 42 | 80 | 118 | 70000 | 3.72E+11 | 5.84E+13 | 1.56E+13 |
| 10150.tif | 4096*3775 | 19.53 | Point | 530 | intestine | 119 | body cavity | 22 | 80 | 118 | 70000 | 3.72E+11 | 4.42E+13 | 8.18E+12 |
| 11150.tif | 4096*3775 | 19.53 | Point | 496 | intestine | 84 | body cavity | 19 | 80 | 118 | 70000 | 3.72E+11 | 3.12E+13 | 7.06E+12 |
| 12150.tif | 4096*3775 | 19.53 | Point | 409 | intestine | 107 | body cavity | 12 | 80 | 118 | 70000 | 3.72E+11 | 3.98E+13 | 4.46E+12 |
| 13150.tif | 4096*3775 | 14.65 | Point | 328 | intestine | 111 | body cavity | 35 | 60 | 157 | 70000 | 3.70E+11 | 4.11E+13 | 1.30E+13 |
| 14150.tif | 4096*3775 | 14.65 | Point | 233 | intestine | 100 | body cavity | 7 | 60 | 157 | 70000 | 3.70E+11 | 3.70E+13 | 2.59E+12 |
| 15200.tif | 4096*3775 | 4.88 | Point | 77 | intestine | 0 | body cavity | 8 | 20 | 300 | 28000 | 6.01E+10 | 0.00E+00 | 4.81E+11 |

day6-8_tissue volume
section thickness: 70 nm

| Image | Size | pixel size (nm) | Main Grid | Pattern count | object | Hit Points | object | Hit Points | hfw (µm) | interval thickness (nm) | Sampling step (pixels) | volume per point (nm^3) | intestinal volume per part (nm^3) | volume of body cavity per part (nm^3) |
|---|---|---|---|---|---|---|---|---|---|---|---|---|---|---|
| 150.tif | 2048*1887 | 13.48 | Point | 132 | intestine | 0 | body cavity | 9 | 27.6 | 80500 | 120 | 2.11E+11 | 0.00E+00 | 1.89E+12 |
| 1300.tif | 2048*1887 | 38.53 | Point | 294 | intestine | 0 | body cavity | 37 | 78.9 | 80500 | 60 | 4.30E+11 | 0.00E+00 | 1.59E+13 |
| 2450.tif | 2048*1887 | 38.53 | Point | 511 | intestine | 108 | body cavity | 58 | 78.9 | 80500 | 60 | 4.30E+11 | 4.65E+13 | 2.49E+13 |
| 3600.tif | 2048*1887 | 38.53 | Point | 658 | intestine | 219 | body cavity | 57 | 78.9 | 80500 | 60 | 4.30E+11 | 9.42E+13 | 2.45E+13 |
| 4750.tif | 2048*1887 | 54.20 | Point | 730 | intestine | 151 | body cavity | 77 | 111 | 80500 | 43 | 4.37E+11 | 6.60E+13 | 3.37E+13 |
| 5900.tif | 2048*1887 | 54.20 | Point | 758 | intestine | 93 | body cavity | 128 | 111 | 80500 | 43 | 4.37E+11 | 4.07E+13 | 5.60E+13 |
| 7050.tif | 2048*1887 | 54.20 | Point | 809 | intestine | 103 | body cavity | 147 | 111 | 80500 | 43 | 4.37E+11 | 4.50E+13 | 6.43E+13 |
| 8200.tif | 2048*1887 | 54.20 | Point | 804 | intestine | 115 | body cavity | 213 | 111 | 80500 | 43 | 4.37E+11 | 5.03E+13 | 9.31E+13 |
| 9350.tif | 2048*1887 | 54.20 | Point | 770 | intestine | 97 | body cavity | 199 | 111 | 80500 | 43 | 4.37E+11 | 4.24E+13 | 8.70E+13 |
| 10500.tif | 2048*1887 | 38.53 | Point | 707 | intestine | 74 | body cavity | 56 | 78.9 | 80500 | 60 | 4.30E+11 | 3.18E+13 | 2.41E+13 |
| 11650.tif | 2048*1887 | 38.53 | Point | 751 | intestine | 91 | body cavity | 82 | 78.9 | 80500 | 60 | 4.30E+11 | 3.91E+13 | 3.53E+13 |
| 12800.tif | 2048*1887 | 54.20 | Point | 771 | intestine | 115 | body cavity | 242 | 111 | 80500 | 43 | 4.37E+11 | 5.03E+13 | 1.06E+14 |
| 13950.tif | 2048*1887 | 38.53 | Point | 690 | intestine | 94 | body cavity | 206 | 78.9 | 80500 | 60 | 4.30E+11 | 4.04E+13 | 8.86E+13 |
| 15100.tif | 2048*1887 | 38.53 | Point | 622 | intestine | 91 | body cavity | 354 | 78.9 | 80500 | 60 | 4.30E+11 | 3.91E+13 | 1.52E+14 |
| 16250.tif | 2048*1887 | 38.53 | Point | 437 | intestine | 57 | body cavity | 212 | 78.9 | 80500 | 60 | 4.30E+11 | 2.45E+13 | 9.12E+13 |
| 17400.tif | 2048*1887 | 20.75 | Point | 212 | intestine | 80 | body cavity | 35 | 42.5 | 80500 | 100 | 3.47E+11 | 2.77E+13 | 1.21E+13 |
| 18500.tif | 2048*1887 | 5.42 | Point | 42 | intestine | 0 | body cavity | 5 | 11.1 | 10500 | 300 | 2.78E+10 | 0.00E+00 | 1.39E+11 |

day6-10_tissue volume
Section thickness: 70 nm

| Image | Size | pixel size (nm) | Main Grid | Pattern count | object | Hit Points | object | Hit Points | hfw (µm) | Sampling step (pixels) | interval thickness (nm) | volume per point (nm^3) | intestinal volume per part (nm^3) | volume of the body avity per part (nm^3) |
|---|---|---|---|---|---|---|---|---|---|---|---|---|---|---|
| 201.tif | 4096*3775 | 13.43 | Point | 257 | intestine | 0 | body cavity | 6 | 55 | 172 | 91000 | 4.85E+11 | 0.00E+00 | 2.91E+12 |
| 1500.tif | 4096*3775 | 15.87 | Point | 464 | intestine | 0 | body cavity | 44 | 65 | 145 | 91000 | 4.82E+11 | 0.00E+00 | 2.12E+13 |
| 2800.tif | 4096*3775 | 19.53 | Point | 588 | intestine | 208 | body cavity | 34 | 80 | 118 | 91000 | 4.83E+11 | 1.01E+14 | 1.64E+13 |
| 4100.tif | 4096*3775 | 20.75 | Point | 720 | intestine | 161 | body cavity | 74 | 85 | 111 | 91000 | 4.83E+11 | 7.77E+13 | 3.57E+13 |
| 5400.tif | 4096*3775 | 24.41 | Point | 857 | intestine | 164 | body cavity | 78 | 100 | 94 | 91000 | 4.79E+11 | 7.86E+13 | 3.74E+13 |
| 6700.tif | 4096*3775 | 24.41 | Point | 869 | intestine | 124 | body cavity | 90 | 100 | 94 | 91000 | 4.79E+11 | 5.94E+13 | 4.31E+13 |
| 8000.tif | 4096*3775 | 24.41 | Point | 925 | intestine | 134 | body cavity | 133 | 100 | 94 | 91000 | 4.79E+11 | 6.42E+13 | 6.37E+13 |
| 9300.tif | 4096*3775 | 23.19 | Point | 971 | intestine | 155 | body cavity | 164 | 95 | 99 | 91000 | 4.80E+11 | 7.44E+13 | 7.87E+13 |
| 10600.tif | 4096*3775 | 23.19 | Point | 948 | intestine | 114 | body cavity | 75 | 95 | 99 | 91000 | 4.80E+11 | 5.47E+13 | 3.60E+13 |
| 11900.tif | 4096*3775 | 23.19 | Point | 990 | intestine | 99 | body cavity | 120 | 95 | 99 | 91000 | 4.80E+11 | 4.75E+13 | 5.76E+13 |
| 13200.tif | 4096*3775 | 24.41 | Point | 1029 | intestine | 109 | body cavity | 280 | 100 | 94 | 91000 | 4.79E+11 | 5.22E+13 | 1.34E+14 |
| 14500.tif | 4096*3775 | 24.41 | Point | 958 | intestine | 100 | body cavity | 325 | 100 | 94 | 91000 | 4.79E+11 | 4.79E+13 | 1.56E+14 |
| 15800.tif | 4096*3775 | 23.19 | Point | 849 | intestine | 103 | body cavity | 242 | 95 | 99 | 91000 | 4.80E+11 | 4.94E+13 | 1.16E+14 |
| 17100.tif | 4096*3775 | 23.19 | Point | 663 | intestine | 96 | body cavity | 320 | 95 | 99 | 91000 | 4.80E+11 | 4.61E+13 | 1.54E+14 |
| 18400.tif | 4096*3775 | 15.87 | Point | 385 | intestine | 149 | body cavity | 84 | 65 | 145 | 91000 | 4.82E+11 | 7.18E+13 | 4.05E+13 |
| 19700.tif | 4096*3775 | 7.32 | Point | 86 | intestine | 0 | body cavity | 6 | 30 | 300 | 91000 | 4.39E+11 | 0.00E+00 | 2.64E+12 |
| 20850.tif | 4096*3775 | 2.69 | Point | 61 | intestine | 0 | body cavity | 0 | 11 | 300 | 7000 | 4.54E+09 | 0.00E+00 | 0.00E+00 |

day9-3_tissue volume
secton thickness: 70 nm

| Image | Size | pixel size (nm) | Main Grid | Pattern count | object | Hit Points | object | Hit Points | hfw (µm) | Sampling step (pixels) | interval thickness (nm) | volume per point (nm^3) | intestinal volume per part (nm^3) | volume of body cavity per part (nm^3) |
|---|---|---|---|---|---|---|---|---|---|---|---|---|---|---|
| 50.tif | 4096*3775 | 9.28 | Point | 156 | intestine | 0 | body cavity | 1 | 38 | 250 | 73500 | 3.95E+11 | 0.00E+00 | 3.95E+11 |
| 1100.tif | 4096*3775 | 14.65 | Point | 521 | intestine | 0 | body cavity | 9 | 60 | 150 | 73500 | 3.55E+11 | 0.00E+00 | 3.19E+12 |
| 2150.tif | 4096*3775 | 17.09 | Point | 498 | intestine | 170 | body cavity | 19 | 70 | 135 | 73500 | 3.91E+11 | 6.65E+13 | 7.43E+12 |
| 3200.tif | 4096*3775 | 18.31 | Point | 508 | intestine | 197 | body cavity | 89 | 75 | 126 | 73500 | 3.91E+11 | 7.71E+13 | 3.48E+13 |
| 4250.tif | 4096*3775 | 19.53 | Point | 633 | intestine | 91 | body cavity | 97 | 80 | 118 | 73500 | 3.90E+11 | 3.55E+13 | 3.79E+13 |
| 5300.tif | 4096*3775 | 19.53 | Point | 634 | intestine | 73 | body cavity | 101 | 80 | 118 | 73500 | 3.90E+11 | 2.85E+13 | 3.94E+13 |
| 6350.tif | 4096*3775 | 20.83 | Point | 707 | intestine | 63 | body cavity | 121 | 85.3 | 110 | 73500 | 3.86E+11 | 2.43E+13 | 4.67E+13 |
| 7400.tif | 4096*3775 | 20.75 | Point | 733 | intestine | 75 | body cavity | 280 | 85 | 110 | 73500 | 3.83E+11 | 2.87E+13 | 1.07E+14 |
| 8450.tif | 4096*3775 | 21.97 | Point | 734 | intestine | 52 | body cavity | 129 | 90 | 105 | 73500 | 3.91E+11 | 2.03E+13 | 5.05E+13 |
| 9500.tif | 4096*3775 | 21.97 | Point | 867 | intestine | 84 | body cavity | 141 | 90 | 105 | 73500 | 3.91E+11 | 3.29E+13 | 5.52E+13 |
| 10550.tif | 4096*3775 | 20.75 | Point | 914 | intestine | 61 | body cavity | 80 | 85 | 110 | 73500 | 3.83E+11 | 2.34E+13 | 3.06E+13 |
| 11600.tif | 4096*3775 | 20.75 | Point | 925 | intestine | 54 | body cavity | 309 | 85 | 110 | 73500 | 3.83E+11 | 2.07E+13 | 1.18E+14 |
| 12650.tif | 4096*3775 | 20.75 | Point | 942 | intestine | 61 | body cavity | 226 | 85 | 110 | 73500 | 3.83E+11 | 2.34E+13 | 8.66E+13 |
| 13700.tif | 4096*3775 | 20.75 | Point | 847 | intestine | 68 | body cavity | 194 | 85 | 110 | 73500 | 3.83E+11 | 2.60E+13 | 7.43E+13 |
| 14750.tif | 4096*3775 | 19.53 | Point | 608 | intestine | 69 | body cavity | 272 | 80 | 120 | 73500 | 4.04E+11 | 2.79E+13 | 1.10E+14 |
| 15800.tif | 4096*3775 | 14.65 | Point | 438 | intestine | 93 | body cavity | 124 | 60 | 150 | 73500 | 3.55E+11 | 3.30E+13 | 4.40E+13 |
| 17250.tif | 4096*3775 | 1.03 | Point | 86 | intestine | 0 | body cavity | 0 | 4.2 | 300 | 59500 | 5.63E+09 | 0.00E+00 | 0.00E+00 |

day9-12 tissue volume
section thickness: 70 nm

| Image | Size | pixel size (nm) | Main Grid | Pattern count | object | Hit Points | object | Hit Points | hfw (µm) | interval thickness (nm) | Sampling step (pixels) | volume of point (nm^3) | intestinal volume per part (nm^3) | volume of body cavity per part (nm^3) |
|---|---|---|---|---|---|---|---|---|---|---|---|---|---|---|
| 200.tif | 2048*1887 | 27.00 | Point | 319 | intestine | 0 | body cavity | 12 | 55.3 | 59500 | 85 | 3.13E+11 | 0.00E+00 | 3.76E+12 |
| 1050.tif | 2048*1887 | 38.53 | Point | 415 | intestine | 0 | body cavity | 29 | 78.9 | 59500 | 60 | 3.18E+11 | 0.00E+00 | 9.22E+12 |
| 1900.tif | 2048*1887 | 38.53 | Point | 544 | intestine | 19 | body cavity | 24 | 78.9 | 59500 | 60 | 3.18E+11 | 6.04E+12 | 7.63E+12 |
| 2750.tif | 2048*1887 | 38.53 | Point | 564 | intestine | 87 | body cavity | 105 | 78.9 | 59500 | 60 | 3.18E+11 | 2.77E+13 | 3.34E+13 |
| 3600.tif | 2048*1887 | 38.53 | Point | 578 | intestine | 177 | body cavity | 153 | 78.9 | 59500 | 60 | 3.18E+11 | 5.63E+13 | 4.86E+13 |
| 4450.tif | 2048*1887 | 38.53 | Point | 634 | intestine | 81 | body cavity | 247 | 78.9 | 59500 | 60 | 3.18E+11 | 2.58E+13 | 7.85E+13 |
| 5300.tif | 2048*1887 | 38.53 | Point | 689 | intestine | 70 | body cavity | 277 | 78.9 | 59500 | 60 | 3.18E+11 | 2.23E+13 | 8.81E+13 |
| 6150.tif | 2048*1887 | 38.53 | Point | 693 | intestine | 55 | body cavity | 196 | 78.9 | 59500 | 60 | 3.18E+11 | 1.75E+13 | 6.23E+13 |
| 7000.tif | 2048*1887 | 38.53 | Point | 611 | intestine | 84 | body cavity | 167 | 78.9 | 59500 | 60 | 3.18E+11 | 2.67E+13 | 5.31E+13 |
| 7850.tif | 2048*1887 | 38.53 | Point | 668 | intestine | 63 | body cavity | 249 | 78.9 | 59500 | 60 | 3.18E+11 | 2.00E+13 | 7.92E+13 |
| 8700.tif | 2048*1887 | 38.53 | Point | 691 | intestine | 55 | body cavity | 267 | 78.9 | 59500 | 60 | 3.18E+11 | 1.75E+13 | 8.49E+13 |
| 9550.tif | 2048*1887 | 38.53 | Point | 647 | intestine | 60 | body cavity | 324 | 78.9 | 59500 | 60 | 3.18E+11 | 1.91E+13 | 1.03E+14 |
| 10400.tif | 2048*1887 | 38.53 | Point | 562 | intestine | 74 | body cavity | 343 | 78.9 | 59500 | 60 | 3.18E+11 | 2.35E+13 | 1.09E+14 |
| 11250.tif | 2048*1887 | 38.53 | Point | 575 | intestine | 66 | body cavity | 291 | 78.9 | 59500 | 60 | 3.18E+11 | 2.10E+13 | 9.25E+13 |
| 12100.tif | 2048*1887 | 38.53 | Point | 471 | intestine | 101 | body cavity | 175 | 78.9 | 59500 | 60 | 3.18E+11 | 3.21E+13 | 5.56E+13 |
| 12950_001.tif | 2048*1887 | 27.00 | Point | 201 | intestine | 27 | body cavity | 19 | 55.3 | 59500 | 85 | 3.13E+11 | 8.46E+12 | 5.96E+12 |
| 13700_001.tif | 2048*1887 | 3.85 | Point | 65 | intestine | 0 | body cavity | 1 | 7.89 | 10500 | 150 | 3.51E+09 | 0.00E+00 | 3.51E+09 |

day18-18(1)_tissue volume
section number: 1-12660,  section thickness 50nm; section number: 12661-18500, section thickness 60 nm

| Image | Size | pixel size (nm) | Main Grid | Pattern count | object | Hit Points | object | Hit Points | hfw (µm) | Sampling step (pixels) | interval thickness (nm) | volume of point (nm^3) | intestinal volume per part (nm^3) | volume of the body cavity per part (nm^3) |
|---|---|---|---|---|---|---|---|---|---|---|---|---|---|---|
| 1_300_001.tif | 2048*1887 | 27.00 | Point | 238 | intestine | 55 | body cavity | 101 | 55.3 | 85 | 62500 | 3.29E+11 | 1.81E+13 | 3.33E+13 |
| 1_1550.tif | 2048*1887 | 38.53 | Point | 301 | intestine | 54 | body cavity | 143 | 78.9 | 60 | 62500 | 3.34E+11 | 1.80E+13 | 4.78E+13 |
| 1_2800.tif | 2048*1887 | 38.53 | Point | 423 | intestine | 43 | body cavity | 192 | 78.9 | 60 | 62500 | 3.34E+11 | 1.44E+13 | 6.41E+13 |
| 1_4050_001.tif | 2048*1887 | 38.53 | Point | 507 | intestine | 28 | body cavity | 249 | 78.9 | 60 | 62500 | 3.34E+11 | 9.35E+12 | 8.32E+13 |
| 1_5300_001.tif | 2048*1887 | 38.53 | Point | 573 | intestine | 49 | body cavity | 124 | 78.9 | 60 | 62500 | 3.34E+11 | 1.64E+13 | 4.14E+13 |
| 1_6550_001.tif | 2048*1887 | 38.53 | Point | 604 | intestine | 37 | body cavity | 61 | 78.9 | 60 | 62500 | 3.34E+11 | 1.24E+13 | 2.04E+13 |
| 1_7800_001.tif | 2048*1887 | 38.53 | Point | 586 | intestine | 73 | body cavity | 82 | 78.9 | 60 | 62500 | 3.34E+11 | 2.44E+13 | 2.74E+13 |
| 1_9050_001.tif | 2048*1887 | 38.53 | Point | 634 | intestine | 72 | body cavity | 71 | 78.9 | 60 | 62500 | 3.34E+11 | 2.40E+13 | 2.37E+13 |
| 1_10300_001.tif | 2048*1887 | 38.53 | Point | 600 | intestine | 78 | body cavity | 150 | 78.9 | 60 | 62500 | 3.34E+11 | 2.60E+13 | 5.01E+13 |
| 1_11550_001.tif | 2048*1887 | 38.53 | Point | 534 | intestine | 67 | body cavity | 144 | 78.9 | 60 | 62500 | 3.34E+11 | 2.24E+13 | 4.81E+13 |
| 1_12800_001.tif | 2048*1887 | 38.53 | Point | 523 | intestine | 67 | body cavity | 196 | 78.9 | 60 | 61400 | 3.28E+11 | 2.20E+13 | 6.43E+13 |
| 1_13850_001.tif | 2048*1887 | 38.53 | Point | 644 | intestine | 124 | body cavity | 132 | 78.9 | 60 | 63000 | 3.37E+11 | 4.17E+13 | 4.44E+13 |
| 1_14900_001.tif | 2048*1887 | 38.53 | Point | 549 | intestine | 170 | body cavity | 131 | 78.9 | 60 | 63000 | 3.37E+11 | 5.72E+13 | 4.41E+13 |
| 1_15950_001.tif | 2048*1887 | 38.53 | Point | 528 | intestine | 53 | body cavity | 183 | 78.9 | 60 | 63000 | 3.37E+11 | 1.78E+13 | 6.16E+13 |
| 1_17000_002.tif | 2048*1887 | 20.75 | Point | 384 | intestine | 0 | body cavity | 50 | 42.5 | 85 | 63000 | 1.96E+11 | 0.00E+00 | 9.80E+12 |
| 1_18100_002.tif | 2048*1887 | 16.85 | Point | 528 | intestine | 0 | body cavity | 39 | 34.5 | 85 | 63000 | 1.29E+11 | 0.00E+00 | 5.04E+12 |

day18-18(2)_tissue volume
section number: 1-12660,  section thickness 50nm; section number: 12661-18500, section thickness 60 nm

| Image | Size | pixel size (nm) | Main Grid | Pattern count | object | Hit Points | object | Hit Points | interval thickness (nm) | hfw (µm) | Sampling step (pixels) | volume per point (nm^3) | intetinal volume per part (nm^3) | volume of body cavity per part (nm^3) |
|---|---|---|---|---|---|---|---|---|---|---|---|---|---|---|
| 2_200.tif | 2048*1887 | 38.53 | Point | 299 | intestine | 26 | body cavity | 116 | 50000 | 78.9 | 60 | 2.67E+11 | 6.95E+12 | 3.10E+13 |
| 2_1200.tif | 2048*1887 | 38.53 | Point | 381 | intestine | 19 | body cavity | 108 | 50000 | 78.9 | 60 | 2.67E+11 | 5.08E+12 | 2.89E+13 |
| 2_2200.tif | 2048*1887 | 38.53 | Point | 491 | intestine | 14 | body cavity | 18 | 50000 | 78.9 | 60 | 2.67E+11 | 3.74E+12 | 4.81E+12 |
| 2_3200_001.tif | 2048*1887 | 38.53 | Point | 535 | intestine | 12 | body cavity | 4 | 50000 | 78.9 | 60 | 2.67E+11 | 3.21E+12 | 1.07E+12 |
| 2_4200_001.tif | 2048*1887 | 38.53 | Point | 547 | intestine | 15 | body cavity | 29 | 50000 | 78.9 | 60 | 2.67E+11 | 4.01E+12 | 7.75E+12 |
| 2_5250_001.tif | 2048*1887 | 38.53 | Point | 499 | intestine | 31 | body cavity | 31 | 50000 | 78.9 | 60 | 2.67E+11 | 8.28E+12 | 8.28E+12 |
| 2_6200_001.tif | 2048*1887 | 38.53 | Point | 507 | intestine | 58 | body cavity | 31 | 50000 | 78.9 | 60 | 2.67E+11 | 1.55E+13 | 8.28E+12 |
| 2_7200_001.tif | 2048*1887 | 38.53 | Point | 552 | intestine | 10 | body cavity | 16 | 50000 | 78.9 | 60 | 2.67E+11 | 2.67E+12 | 4.27E+12 |
| 2_8200_001.tif | 2048*1887 | 38.53 | Point | 541 | intestine | 14 | body cavity | 14 | 50000 | 78.9 | 60 | 2.67E+11 | 3.74E+12 | 3.74E+12 |
| 2_9200_001.tif | 2048*1887 | 38.53 | Point | 489 | intestine | 13 | body cavity | 13 | 50000 | 78.9 | 60 | 2.67E+11 | 3.47E+12 | 3.47E+12 |
| 2_10200_001.tif | 2048*1887 | 38.53 | Point | 466 | intestine | 71 | body cavity | 158 | 50000 | 78.9 | 60 | 2.67E+11 | 1.90E+13 | 4.22E+13 |
| 2_11200_001.tif | 2048*1887 | 38.53 | Point | 460 | intestine | 97 | body cavity | 162 | 50000 | 78.9 | 60 | 2.67E+11 | 2.59E+13 | 4.33E+13 |
| 2_12200_001.tif | 2048*1887 | 38.53 | Point | 424 | intestine | 94 | body cavity | 158 | 50400 | 78.9 | 60 | 2.69E+11 | 2.53E+13 | 4.25E+13 |
| 2_13100_001.tif | 2048*1887 | 38.53 | Point | 547 | intestine | 140 | body cavity | 106 | 48000 | 78.9 | 60 | 2.56E+11 | 3.59E+13 | 2.72E+13 |
| 2_13900_001.tif | 2048*1887 | 38.53 | Point | 449 | intestine | 0 | body cavity | 49 | 48000 | 78.9 | 60 | 2.56E+11 | 0.00E+00 | 1.26E+13 |
| 2_14700_001.tif | 2048*1887 | 38.53 | Point | 331 | intestine | 0 | body cavity | 127 | 48000 | 78.9 | 60 | 2.56E+11 | 0.00E+00 | 3.26E+13 |
| 2_15500_001.tif | 2048*1887 | 38.53 | Point | 178 | intestine | 0 | body cavity | 23 | 51000 | 78.9 | 60 | 2.73E+11 | 0.00E+00 | 6.27E+12 |
